# Biosynthesis of macrocyclic peptides with C-terminal β-amino-α-keto acid groups by three different metalloenzymes

**DOI:** 10.1101/2023.10.30.564719

**Authors:** Dinh T. Nguyen, Lingyang Zhu, Danielle L. Gray, Toby J. Woods, Chandrashekhar Padhi, Kristen M. Flatt, Douglas A. Mitchell, Wilfred A. van der Donk

## Abstract

Advances in genome sequencing and bioinformatics methods have identified a myriad of biosynthetic gene clusters (BGCs) encoding uncharacterized molecules. By mining genomes for BGCs containing a prevalent peptide-binding domain used for the biosynthesis of ribosomally synthesized and post-translationally modified peptides (RiPPs), we uncovered a new class involving modifications installed by a cytochrome P450, a multi-nuclear iron-dependent non-heme oxidative enzyme (MNIO, formerly DUF692), a cobalamin- and radical *S*-adenosyl-L-methionine-dependent enzyme (B12-rSAM), and a methyltransferase. All enzymes encoded by the BGC were functionally expressed in *Burkholderia* sp. FERM BP-3421. Structural characterization with 2D-NMR and Marfey’s method on the resulting RiPP demonstrated that the P450 enzyme catalyzed the formation of a biaryl C-C crosslink between two Tyr residues with the B12-rSAM generating β-methyltyrosine. The MNIO transformed a C-terminal Asp residue into aminopyruvic acid while the methyltransferase acted on the β-carbon of the α-keto acid. Exciton-coupled circular dichroism spectroscopy and microcrystal electron diffraction (MicroED) were used to elucidate the stereochemical configurations of the atropisomer that formed upon biaryl crosslinking. The conserved Cys residue in the precursor peptide was not modified as in all other characterized MNIO-containing BGCs; However, mutational analyses demonstrated that it was essential for the MNIO activity on the C-terminal Asp. To the best of our knowledge, the MNIO featured in this pathway is the first to modify a residue other than Cys. This study underscores the utility of genome mining to discover new macrocyclic RiPPs and that RiPPs remain a significant source of previously undiscovered enzyme chemistry.

## Introduction

Natural products are prolific sources of structurally diverse and biologically active compounds of high societal value.^1^ The rapid expansion of genomic sequence databases, combined with the development of high-throughput, accurate, and open-access bioinformatics tools, has unveiled many new natural product biosynthetic pathways.^2-5^ Just as advances in directed evolution revolutionized the use of enzymes in commodity chemical synthesis, enzymes sourced from natural product biosynthetic pathways have become a rich reservoir for the development of innovative biocatalytic processes.^6-9^

Ribosomally synthesized and post-translationally modified peptides (RiPPs) are a large and growing family of natural products featuring a diverse range of molecular scaffolds.^10^ The ribosomal precursor peptide frequently contains an N-terminal leader region typically responsible for recruiting the modifying enzymes, while a C-terminal core region receives the post-translational modifications.^11^ Exceptional transformations catalyzed by RiPP enzymes result in diverse structural moieties^10^ possessing antimicrobial, antiviral, antifungal, herbicidal, cytotoxic, anticancer, and other activities.^12-14^ With nearly 50 reported structural classes, defined by the post-translational modification(s) installed, RiPP biosynthesis is a highly productive arena for the discovery of new enzyme chemistry,^15-29^ identification of versatile bioengineering catalysts,^30-33^ and isolation of structurally exotic compounds with unprecedented biological modes of action.^34,35^ While tools like AlphaFold have provided high quality structures for millions of enzymes,^36^ a major unsolved challenge is the determination of enzyme function. Typically, one does not know the substrate for a novel enzyme of interest.^37^ However, this challenge is simplified for enzymes involved in prokaryotic RiPP biosynthesis, as the substrate(s) are typically encoded near the modifying enzyme(s) in a biosynthetic gene cluster (BGC).^10^ Advances in identifying and analyzing short open-reading frames in prokaryotic genomes enabled reliable annotation of RiPP precursor peptides of known classes and high-confidence prediction of precursor peptides encoded by yet-uncharacterized RiPP classes.^4,38-40^

Our approach to finding BGCs that encode first-in-class RiPPs starts with bioinformatic searches centered on a prevalent, class-agnostic protein domain termed the RiPP precursor recognition element (RRE).^39,41,42^ These domains are found in the majority of prokaryotic RiPP BGCs but have been difficult to identify owing to their small size (80-90 residues), frequent fusion to much larger proteins, and high sequence variability. All known RRE domains contain three alpha helices and a three-stranded beta sheet. The cognate precursor peptides are engaged as the fourth strand of the beta sheet, often with nanomolar affinities. While the structural fold of RREs is well conserved, sequence homology tools like BLAST-P are insufficient to retrieve RRE domains across RiPP classes. This diversity has necessitated the use of Hidden Markov model-(HMM) based retrieval for cataloging RRE domains. The tool RRE-Finder uses a collection of HMMs and secondary structure prediction to identify these domains,^42^ which serve as an incomplete but useful biomarker of a RiPP BGC. This allows the prioritization of RiPP BGCs that encode novel collections of enzymes or hypothetical enzymes with no known function. The uniqueness of the precursor peptide is also considered paramount in BGC prioritization, as it is the foundation on which the final RiPP is constructed.^4,43^ This procedure was recently followed to discover a new class of RiPPs, now termed the daptides, which modify an invariant C-terminal Thr into (*S*)-*N*_2_,*N*_2_-dimethyl-1,2-propanediamine.^39^ Thus, daptides are ribosomal peptides with two N-termini.

In this work, we pursued the structural characterization of a RiPP BGC from *Burkholderia thailandensis* E264 that is unlike any other reported. The BGC uniquely encodes three metalloenzymes that we predicted would transform the precursor peptide into an unprecedented scaffold. Indeed, we show the cumulative actions of a multinuclear non-heme iron-dependent oxidative enzyme (from protein family PF05114, abbreviated as MNIO), a cobalamin- and radical *S*-adenosyl-L-methionine-dependent enzyme, a cytochrome P450, and a methyltransferase. The cytochrome P450 installed a biaryl macrocyclic peptide formed from two Tyr residues, with one Tyr also undergoing *C*-methylation on an unactivated sp^3^ carbon center catalyzed by the B12-rSAM. Subsequent catalysis by the MNIO on the resulting biaryl atropisomer-containing peptide converted the C-terminal Asp residue to aminopyruvic acid. Given that the MNIO is fully conserved in these RiPP BGCs, we consider it to be class-defining. We therefore termed the products aminopyruvatides and name the BGCs that produce them as *apy*. Some of these PTMs were not observed during heterologous expression in *Escherichia coli* (*E. coli)* but were successfully elucidated in *Burkholderia* sp. FERM BP-3421. Overall, this study suggests that a combination of genome mining focusing on RREs and reconstitution of biosynthetic pathways in heterologous hosts other than the traditional *E. coli* can provide advantageous opportunities for expanding our knowledge of the natural product chemical space.

## Results

### Bioinformatic survey and experimental target selection

We sought out an RRE-dependent RiPP BGC containing multiple uncharacterized metalloenzymes, given their unparalleled ability to catalyze difficult and diverse chemical transformations.^44-46^ A BGC obtained from RRE-Finder that caught our interest was from *Burkholderia thailandensis* E264 and is composed of a sequence-unique substrate peptide (NCBI accession code: ABC34935.1), an MNIO enzyme (ApyH, ABC35712.1),^26^ an RRE-containing protein (ApyI, ABC34269.1), a methyltransferase (ApyS, ABC34725.1), an RRE-containing cobalamin- and radical *S*-adenosyl-L-methionine-dependent enzyme (B12-rSAM, ApyD, ABC34219.1),^46,47^ and a cytochrome P450 enzyme (ApyO, ABC35200.1) (Figure 1, Supplementary Dataset 1).^45,48^ To obtain homologous BGCs, we performed a BLASTP search of each enzyme against the non-redundant NCBI protein database. The genome neighborhood and putative precursor peptides corresponding to the collected homologs were obtained using RODEO,^4^ and BGCs containing putative precursor(s) were analyzed. During this process, we noticed that the precursor peptides were diverse in sequence with only a central CBX_2-3_G motif (X = any amino acid, B = hydrophobic amino acid) and a C-terminal Asp as the conserved features. This knowledge was then utilized to identify probable precursor peptides. This workflow identified highly similar BGCs that were prevalent in the *Burkholderia pseudomallei* group. Select strains from other Pseudomonodota and the phylogenetically distant Actinomycetota also contain putative *apy* BGCs (Supplementary Dataset 1). The homologous BGCs contain an MNIO (100% occurrence), a B12-rSAM (97% occurrence), and a methyltransferase (93% occurrence), whereas only 74% contain a P450. A sequence alignment of the predicted precursor peptides highlighted the previously mentioned conserved C-terminal Asp and a conserved CBX_2-3_G motif in the central portion, which *a priori* could not be assigned as being part of the leader or core region (Figure 1B). Further, we noticed that two aromatic (either Trp or Tyr) residues were only present as the 2^nd^ and 4^th^ residue from the C-terminus when the BGC encoded a P450 enzyme. Identification of such correlations are often informative for determining the site(s) of enzymatic action.

**Figure 1.**
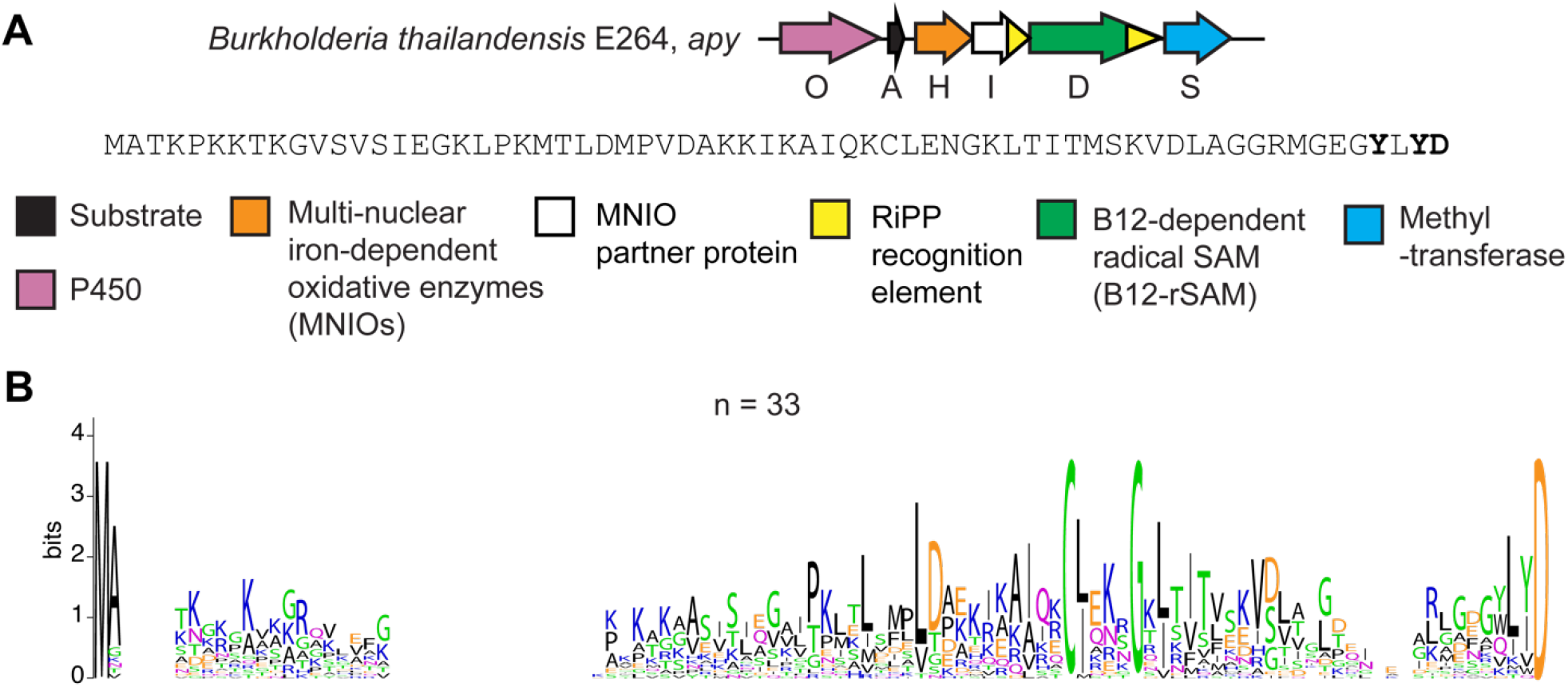
Overview of the *apy* cluster and conserved amino acid motifs in predicted precursor peptides from similar clusters. (A) BGC diagram of the RiPP pathway of interest (*apy*) from *B. thailandensis* E264 and the sequence of the precursor peptide. Amino acid residues that undergo modifications are bolded. (B) Sequence logo of ApyA precursor peptides from homologous BGCs with identical sequences removed (unique sequences = 33, total sequences = 69).

### Assignment of enzymatic activities via heterologous expression in *E. coli*

Initially, we constructed an *E. coli* expression plasmid encoding the precursor peptide (ApyA) containing an N-terminal hexa-His tag and all associated modifying enzymes in a pRSF-based vector (Table S1). All genes were codon-optimized for *E. coli* expression. Given the reliance of the B12-rSAM (ApyD) on cobalamin, and the inability of *E. coli* to synthesize vitamin B12 and derivatives, we co-transformed a pCDF-based vector containing the vitamin B12-uptake pathway (*btu*) and supplemented the medium with cobalamin during expression.^49,50^ The peptide products resulting from co-expression were purified using Ni-NTA affinity chromatography and digested by endoproteinase LysC or GluC, and the samples were subjected to high-resolution electrospray ionization mass spectrometry (HR-ESI-MS).

Upon co-expression of ApyA and the rSAM ApyD and *btu* an increase of +14 Da was observed, suggestive of a single methylation (Figure 2, Figure S1). HR-ESI tandem mass spectrometry (HR-ESI-MS/MS) localized the mass increase to the penultimate residue of ApyA (Tyr), which is found in a Gly-Tyr-Leu-<u>Tyr</u>-Asp motif (Figure 3). Upon co-expression of ApyA with the cytochrome P450 ApyO, a mass loss of 2 Da was observed (Figures 2, S1). HR-ESI-MS/MS analysis localized this change to the Tyr-Leu-Tyr region, and the lack of observable single fragmentation in this part of the peptide was suggestive of a crosslink (Figure S2). Co-expression of ApyA with ApyO and ApyD/*btu* yielded a mass change of +12 Da, consistent with the ApyO-catalyzed 2 Da mass loss and ApyD-catalyzed 14 Da mass gain (Figure S1). Co-expression of ApyA with ApyHI (MNIO and RRE domain) in *E. coli* did not lead to any mass changes. Co-expression of ApyA with all protein components encoded by the BGC (ApyODHIS) yielded the same mass deviations observed upon co-expression with just ApyO and ApyD (Figure S1). These results imply that ApyHI and ApyS were either non-functional in *E. coli* or performed mass-neutral modifications that preserved the molecular formula.

**Figure 2.**
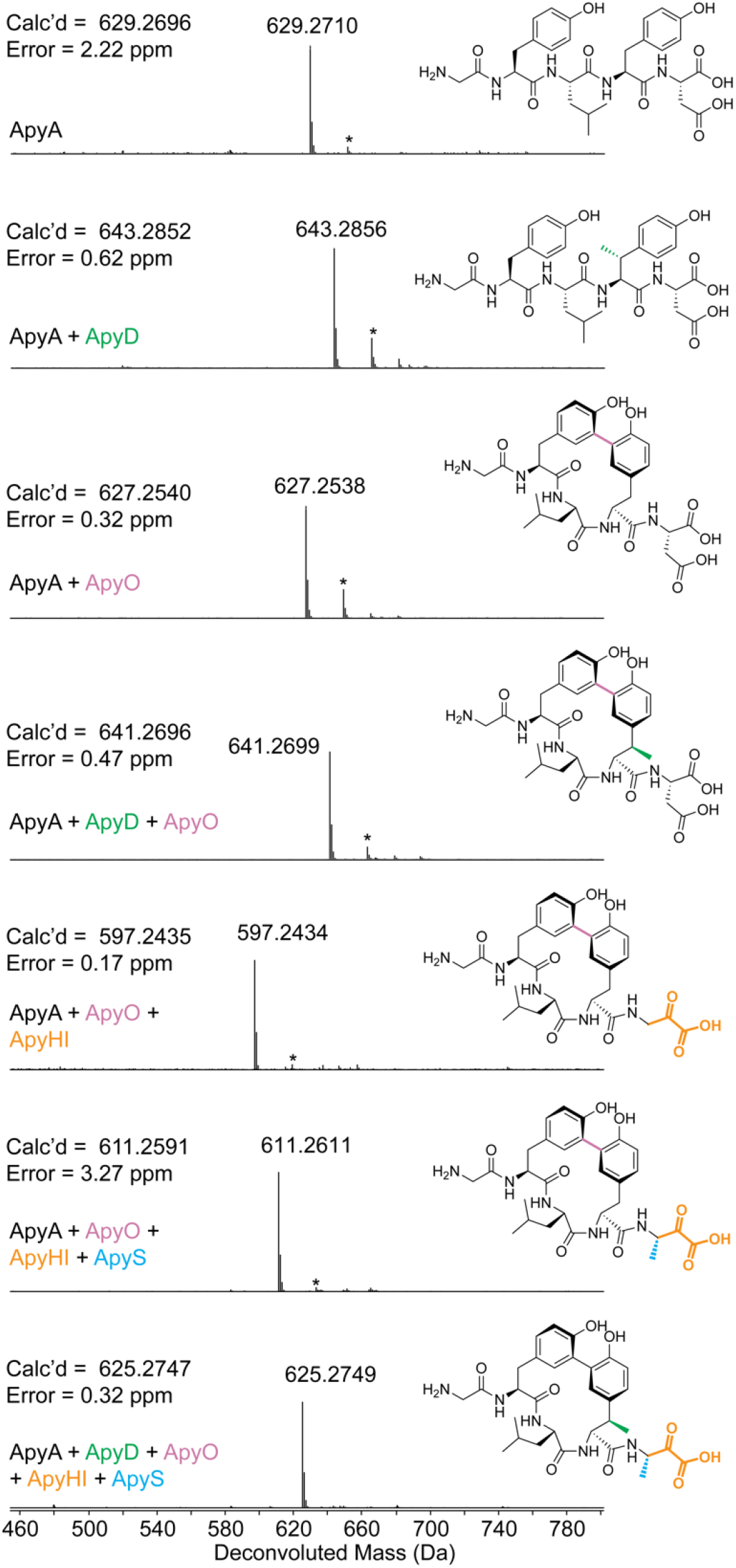
Deconvoluted HR-ESI mass spectra and elucidated structures of ApyA modified by different combinations of enzymes followed by proteolysis with GluC. Structures drawn in 2^nd^, 3^rd^, 4^th^, and 6^th^ spectra were verified by 2D-NMR spectroscopy. The structure drawn in the 3^rd^ spectrum was also verified by MicroED. Other drawn structures were inferred from HR-ESI-MS/MS experiments and determined enzyme functions. * denotes [M+Na]^+^. ApyA, precursor peptide; ApyD (green), B12-rSAM; ApyO (pink), cytochrome P450; ApyHI (orange), MNIO and partner protein; ApyS (blue), methyltransferase. The top four spectra were from expression in *E. coli* and the bottom three spectra were from expression in *Burkholderia* sp. FERM BP-3421. All products were purified by HPLC prior to MS analysis.

**Figure 3.**
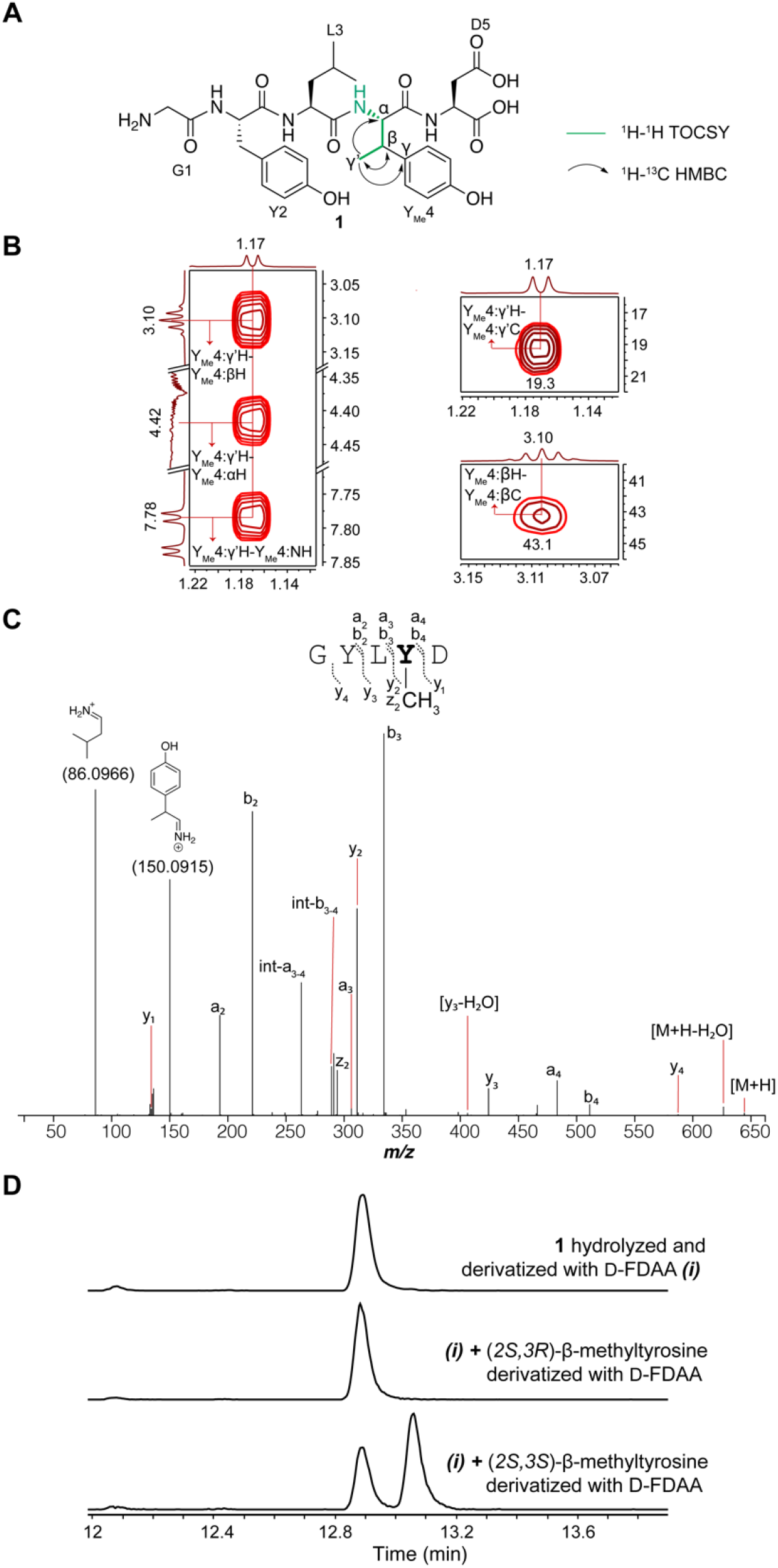
Structural elucidation of the ApyD-catalyzed modification. (A) Key NMR correlations consistent with the presence of β-methyltyrosine in peptide **1** (axes in ppm). (B) NMR spectral data of diagnostic cross peaks indicating the position of the new methyl group. *Left*: ^1^H-^1^H TOCSY. *Right*: ^1^H-^13^C HSQC. The ^1^H-^13^C HMBC key correlations can be found in Figure S17F. (C) HR-ESI-MS/MS data showing the site of methylation. int = internal fragment. The mass table with observed error can be found in the Supplementary Dataset 3. (D) Extracted ion chromatograms (EICs, *m/z* = 700.1937) of **1** or authentic standards after derivatization with Marfey’s reagent (D-FDAA). The EICs of **1** and authentic standards after derivatization with L-FDAA are shown in Figure S25.

### Assignment of enzymatic activities via heterologous expression in *Burkholderia*

To address the first possibility, we performed heterologous expression in *Burkholderia* sp. FERM BP-3421. This emerging chassis strain is closely related to *B. thailandensis* E264, and has been shown to significantly enhance the production of RiPP and non-RiPP natural products.^51-53^ To facilitate expression, we inserted the genomic DNA containing the *apy* BGC downstream of the rhamnose-inducible promoter in a pSCrhaB2 plasmid used previously for protein expression in *Burkholderia* (Table S2).^54^ Additionally, we introduced an N-terminal hexa-His tag on ApyA to facilitate isolation. Upon co-expression of the entire *apy* BGC and Ni-NTA purification, the major species observed was the ApyA precursor peptide with a mass loss of 18 Da (Figures 2, S3). Analysis of tandem MS data suggested that a loss of 2 Da localized to the Tyr-Leu-Tyr region identical to the ApyO-catalyzed modification observed in *E. coli* (Figure S4). The remaining loss of 16 Da was confined to the conserved C-terminal Asp residue (Figure S4). Omission of ApyH or ApyI from the co-expression cultures resulted in no modification of the C-terminal Asp (Figures S5-S8). Omission of ApyS from the expression trials resulted in a 32 Da mass loss from unmodified ApyA (Figure S9). Given the 2 Da mass loss assigned to the P450 reaction, the remaining 30 Da mass loss was attributed to ApyHI. Tandem MS localized this mass deviation (30 Da) to the C-terminal Asp residue (Figure S10). Collectively, these data suggested that ApyHI are active upon expression in *Burkholderia*, but not *E. coli*, and work together to modify the C-terminal Asp. This finding was unexpected since all reported reactions for MNIOs modify Cys residues.^16,26,55^ Furthermore, these data suggest that ApyS catalyzes a methylation reaction on the ApyHI-modified C-terminal Asp residue.

Our initial expression trials in *Burkholderia* sp. FERM BP-3421 appeared to produce active ApyHI, ApyO, and ApyS resulting in an ApyA product decreased in mass by 18 Da. We also observed a minor species with a mass 14 Da larger than the M–18 Da product (i.e. 4 Da decreased from unmodified ApyA; Figure S3), suggesting low efficiency of ApyD-catalyzed methylation. Supplementation of cobalamin to the growth medium and expression optimization resulted in higher efficiency but still incomplete methylation (Figure S11-S13). To improve the ApyD-catalyzed methylation, we redesigned the expression vector (Figure S14, Table S3) to introduce a separate ribosome-binding site for ApyD instead of utilizing the native gene architecture with overlapping *apyH, apyI*, and *apyD* genes. This construct resulted in the ApyA/O/HI/D/S product (Figure 2) being the major product (Figures S11, S15), as supported by HR-ESI-MS/MS data (Figure S16). The mass loss of 4 Da compared to unmodified ApyA was accounted for by a putative Tyr-Tyr crosslink catalyzed by ApyO (–2 Da), methylation of the penultimate Tyr by ApyD (+14 Da), an unknown (–30 Da) modification to the C-terminal Asp catalyzed by ApyHI, and methylation by ApyS (+14 Da).

### Structural elucidation of the ApyD and ApyO reaction products

The MS/MS data described thus far suggested that ApyD methylates the penultimate Tyr of the ApyA peptide; however, the precise site of methylation remained unclear. We therefore prepared a larger quantity of ApyD-modified ApyA in *E. coli*, digested the product with endoproteinase GluC, and purified the resulting methylated Gly-Tyr-Leu-<u>Tyr</u>-Asp pentapeptide by high-performance liquid chromatography (HPLC). Multi-dimensional nuclear magnetic resonance (NMR) spectroscopy was then used to elucidate the structure (Figures 3, S17). A combination of ^1^H-^1^H TOCSY and ^1^H-^1^H ROESY spectra were used to assign the ^1^H chemical shifts (Table S4). Importantly, the ^1^H-^1^H TOCSY spectrum showed a spin system containing the N-H, C_α_-H, C_β_-H, and the new methyl group (γ^’^). ^1^H-^13^C HSQC analysis demonstrated the β-carbon of the Tyr to be CH instead of CH_2_, and ^1^H-^13^C HMBC produced correlations between the new methyl group, C_β_-H and C_α_-H of Tyr, and the C4 carbon of the aromatic side chain (Figure S17F). Overall, these spectroscopic data support ApyD-catalyzed formation of β-methyltyrosine.

To ascertain the stereochemical configuration of the β-methyltyrosine product, Marfey’s method was employed.^56,57^ Specifically, the methylated pentapeptide was hydrolyzed using 6 M DCl/D_2_O, followed by derivatization with both the L- and D-forms of Marfey’s reagent (1-fluoro-2-4-dinitrophenyl-5-alanine amide) (Figure S18). The commercial synthetic standards (2*S*,3*R*)-β-methyltyrosine and (2*S*,3*S*)-β-methyltyrosine were subjected to analogous hydrolysis and derivatization conditions. Coelution in LCMS of the derivatized β-methyltyrosine obtained enzymatically with the similarly derivatized authentic standard of (2*S*,3*R*)-β-methyltyrosine (Figures 3, S18) allowed assignment of the ApyD-catalyzed product as (2*S*,3*R*)-β-methyltyrosine.

We next determined the structure of ApyO-modified ApyA utilizing a similar workflow. Integration of the ^1^H-NMR spectrum of the pentapeptide detected only six aromatic C-H protons, instead of the eight that would be expected for two unmodified Tyr residues. This finding suggested a C-C biaryl crosslink (Figure S19, Table S5). The spin system of the aromatic rings observed by ^1^H and ^1^H-^1^H TOCSY showed two groups of peaks with splitting patterns of doublet (d, 8 Hz), d of doublets (dd, 8 Hz, 2 Hz), and d (2 Hz) (Figure S19G). This change in the aromatic spin system (compared to unmodified Tyr) along with correlations between the meta-hydrogen (d, 2 Hz) of ring 1 and the ortho-carbon of ring 2 (and vice versa, demonstrated by ^1^H-^13^C HMBC), supported a crosslink between the ortho carbons of each Tyr residue (Figure S19J). Analogous 2D-NMR experiments on the peptide resulting from co-expression of ApyA with both ApyO and ApyD were consistent with the product containing the C-C crosslink and β-methyltyrosine (Figure S20, Table S6).

We next assigned the stereochemistry of the atropisomer formed by the ApyO-catalyzed biaryl crosslink. We utilized exciton-coupled circular dichroism (ECD), a non-empirical method for the assignment of the spatial orientation between chromophores absorbing at similar wavelengths.^58^ We first obtained ECD spectra of four chiral synthetic molecules with a similar aromatic C-C crosslink. The ECD spectra of molecules with *R* axial chirality exhibited a negative first and positive second Cotton effect, while the ECD spectra of molecules with *S* axial chirality exhibited the opposite Cotton effect (Figure S21). Analysis of the ApyO-modified peptide yielded a negative first (307 nm) and positive second Cotton effect (285 nm), suggesting a counterclockwise orientation between the two Tyr residues, and hence we assigned *R* axial chirality (Figure S22). Control samples lacking the biaryl crosslink did not exhibit these CD signals (Figure S23).

We next investigated whether the ApyD- and ApyO-modified ApyA product exhibited similar atropisomerism as the ApyO-modified ApyA product, as modifications on the macrocycle can result in changes in the atropisomeric state.^59,60^ ECD spectra of the GluC-cleaved ApyA/D/O product exhibited a significant positive Cotton effect like the ApyA/O product but no negative Cotton effect (Figure S23). This lack of observable exciton coupling has been noted on other chiral molecules.^61,62^ We then turned to the cryo-electron microscopy (cryo-EM) method of microcrystal electron diffraction (MicroED).^63-67^ The diffraction data from two nanocrystals were merged to yield a nearly complete dataset with 1.0 Å resolution (Figure 4). The structure of the ApyA/D/O product was then determined by direct methods, which demonstrated *R* axial chirality for the biaryl ring similar to the ApyA/O product, suggesting that the β-methylation of the Tyr residue did not alter the prevailing atropisomer (Figures 4, S24, Table S7). Furthermore, the assignment by Marfey’s method of the 2*S*,3*R* configuration of β-methyltyrosine was confirmed by MicroED. The structure was deposited under CCDC ID 2324739.

**Figure 4.**
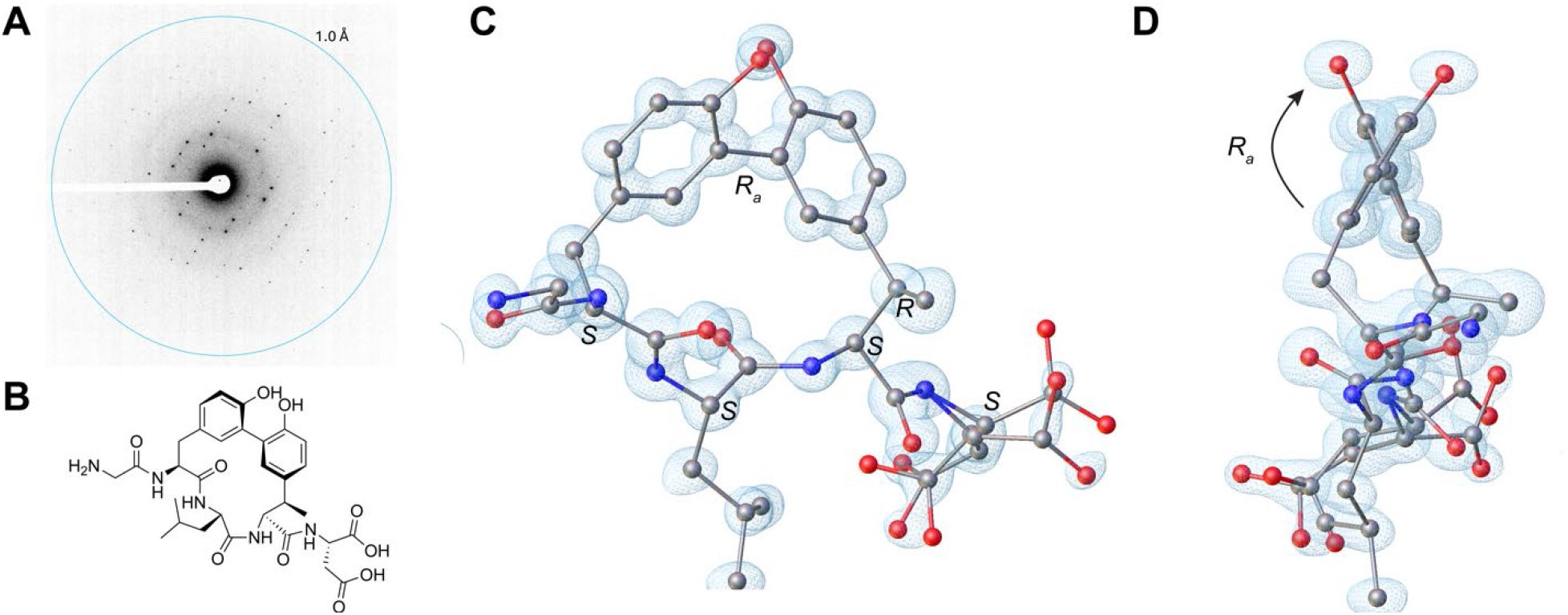
Determination of the ApyD- and ApyO-modified ApyA pentapeptide structure by MicroED. (A) Diffraction pattern of the peptide crystal with resolution rings set at 1.0 Å. (B) Drawing of the chemical structure of ApyD- and ApyO-modified ApyA pentapeptide. (C) MicroED structure of ApyD- and ApyO-modified ApyA pentapeptide at 1.0 Å resolution. CCDC ID 2324739. The blue mesh represents the observed electron density map (F_obs_). The full structure comprises two complete peptide molecules with a Zn atom and water molecules (see Figure S24 and Supplementary Dataset). Here, only one peptide molecule is shown, and hydrogen atoms are not shown for clarity. (D) Side view to highlight the *R* axial chirality.

### Structural elucidation of the ApyO/HI/S reaction product

We next determined the structures of the ApyA modification by ApyHI and ApyS. Omission of ApyO from co-expression resulted in the absence of a modification of the C-terminal Asp, suggesting that the biaryl crosslink is a prerequisite for ApyHI and/or ApyS activity (Figures S25-S26). Consequently, we prepared a larger quantity of ApyA modified by ApyO, ApyHI and ApyS in *Burkholderia* sp. FERM BP-3421 for structure determination (Figure S27, Table S8). Analysis of the pentapeptide product after GluC proteolysis confirmed the C-C crosslink between the ortho carbons of the two Tyr (Figures 5, S27G-H, Table S8). ^1^H-^1^H TOCSY and ^1^H-^13^C HSQC analysis revealed a new spin system for the structure originating from the C-terminal Asp that was comprised of an N-H amide (8.35 ppm), a C-H group with chemical shifts reminiscent of α positions of amino acid residues (4.91 ppm for ^1^H and 51.6 ppm for ^13^C), and a new methyl group (1.31 ppm) (Figure 5, S27B, 27E, S27I). The ^1^H-^13^C HMBC spectrum revealed correlations between the carbon atoms in the C-terminal structure and all the C-H protons and a new carbonyl based on the ^13^C chemical shift (202.5 ppm) (Figure S27I). A secondary species was detected containing an N-H amide (7.81 ppm), a C-H group (4.18 ppm for ^1^H and 50.5 ppm for ^13^C), and a methyl group (1.04 ppm) on the C-terminal residue (Figure S27J, 27K). The ^1^H-^13^C HMBC cross peak between the methyl protons and a carbon exhibiting a chemical shift consistent with a geminal diol (95.1 ppm) suggested this secondary species represented the hydrate of the new ketone, as the data were acquired in 10% D_2_O and 90% H_2_O (Figure S27J, 27K). Owing to this equilibrium, during the mixing period in the pulse sequences of the ^1^H-^1^H NOESY and ^1^H-^1^H ROESY experiments, the signal for the γ-methyl group of the ketone (1.31 ppm) exchanged to the chemical environment of the γ-methyl group of the geminal diol (1.04 ppm), represented by a cross peak connecting the two frequencies observed at the end of the pulse sequences (Figure S27L). This exchange was further supported by this cross peak between the two methyl groups having the same phase as the diagonal peaks in the ^1^H-^1^H ROESY spectrum, whereas all other ROE cross peaks arising from spatial proximity between two nuclei having the opposite phase to the diagonal peaks (Figure S27D, Figure S27L).^68^ These results, combined with the HR-ESI-MS data showing ApyHI catalysis resulted in a loss of 30 Da from unmodified ApyA, suggested that ApyHI and ApyS converted the C-terminal Asp residue into a C-terminal 3-amino-2-oxobutanoic acid (Figure 5).

**Figure 5.**
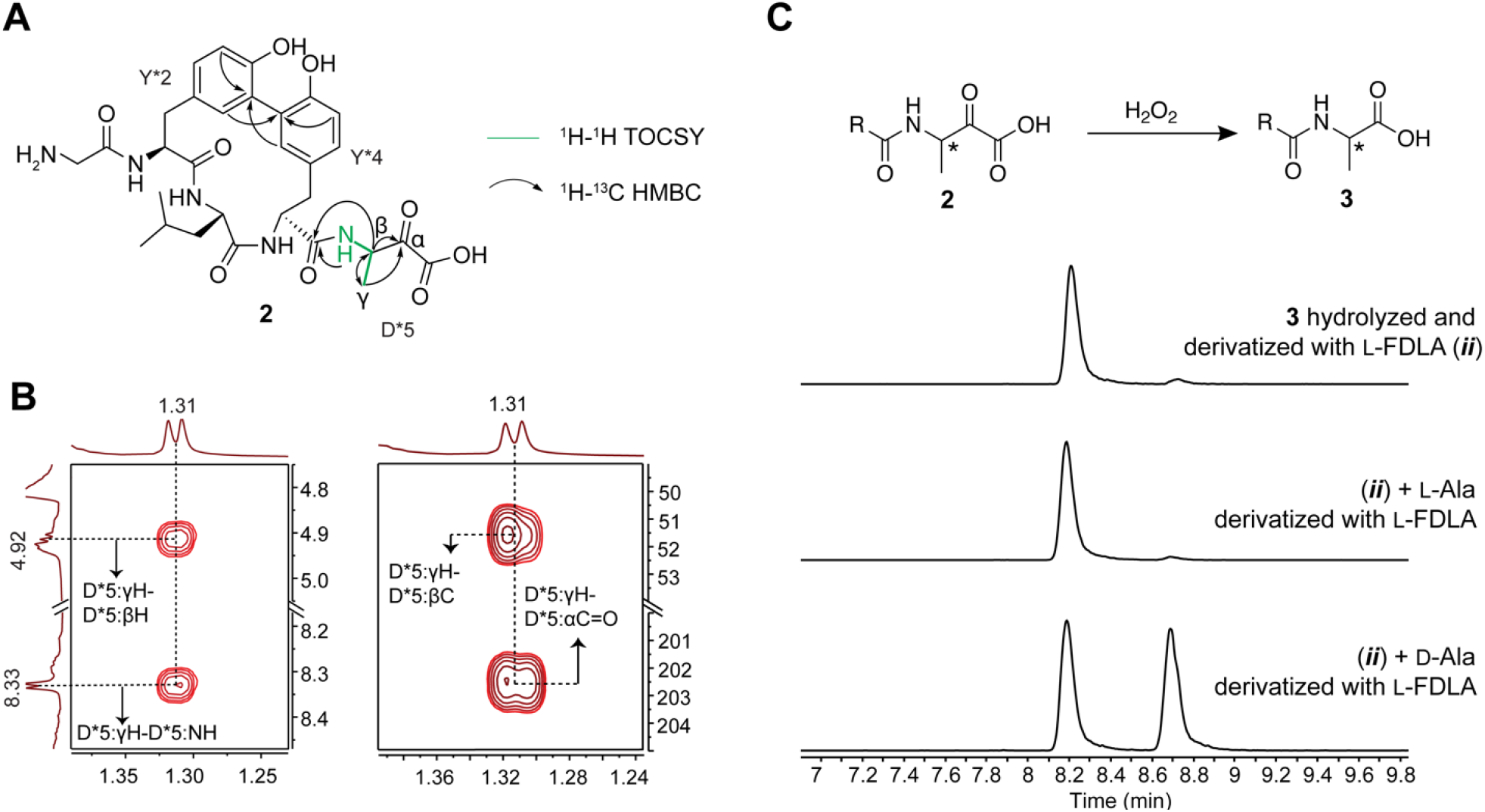
Structural elucidation of the chemistry performed by ApyO, ApyHI and ApyS. (A) Depiction of the 2D-NMR data consistent with a C-C biaryl crosslink and C-terminal 3-amino-2-oxobutanoic acid in peptide **2**. (B) 2D-NMR spectra (*Left*: ^1^H-^13^C HMBC; *Right*: ^1^H-^1^H TOCSY) with key cross peaks elucidating the C-terminal 3-amino-2-oxobutanoic acid. Other key cross peaks can be found in Figure S27I. (C) Schematic depiction of the oxidation of the C-terminal 3-amino-2-oxobutanoic acid by H_2_O_2_ to an Ala residue, and the EIC (*m/z* = 382.1368) of hydrolyzed and derivatized Ala residues after Marfey’s derivatization.

The stereochemistry at the β-carbon of the newly installed α-keto acid was determined by Marfey’s analysis. To circumvent acid-catalyzed epimerization adjacent to the ketone, we first oxidized the α-keto acid to a carboxylic acid using H_2_O_2_ (Figures 5, S28).^69,70^ The successful conversion of the C-terminal 3-amino-2-oxobutanoic acid to Ala further affirmed the structural assignments. We then performed acid hydrolysis and derivatization with the advanced Marfey’s reagent 1-fluoro-2,4-dinitrophenyl-5-L-leucine-amide, and in parallel, derivatized authentic standards of L- and D-Ala.^71^ LC-MS analysis indicated that the derivatized Ala post peptide hydrolysis was L configured, suggesting the new C-terminus installed by ApyHI and ApyS was (*S*)-3-amino-2-oxobutanoic acid.

As previously stated, omission of ApyS from co-expression experiments abolished methyl group transfer to the C-terminus of the modified peptide, implying ApyHI converted the C-terminal Asp residue into aminopyruvic acid (Figures 2, S9-S10). Supporting this hypothesis, we obtained an AlphaFold^36^ model of ApyS and the closest structural homolog identified by DALI^72^ was MppJ, a methyltransferase converting phenylpyruvic acid to 3-methylphenylpyruvic acid (Figure S29).^73,74^ The structural and functional resemblance supports that ApyS *C*-methylates the relatively nucleophilic β-carbon of the α-keto acid generated by ApyHI.

### The conserved Cys of ApyA is essential for MNIO activity

ApyHI is the first MNIO-RRE pair to catalyze the modification of a non-Cys residue, leading us to question whether the centrally located, conserved CBX_2-3_G motif of ApyA was left unmodified (Figure 1B). HR-ESI-MS/MS data indicated that no other segment of ApyA besides the C-terminal Tyr-Leu-Tyr-Asp motif was enzymatically modified in mass (Figures S30-S31). To examine the possibility that ApyHI performed an isobaric modification to the central Cys residue (Figure 1), we reacted ApyHI-treated ApyA with iodoacetamide, which yielded an *S*-alkylated product, indicating the presence of a sulfhydryl group. The peptide was then digested with endoproteinase GluC and trypsin, generating an *S*-alkylated Cys-Leu-Glu tripeptide for examination by NMR spectroscopy (Figure S32, Table S9) and Marfey’s analysis (Figures S33-S34). The ^1^H-^13^C HSQC spectrum of the alkylated peptide revealed two CH_2_ moieties, one from the β-carbon of Cys and one from the carbamidomethyl group, along with a CH moiety from the α-carbon of Cys (Figure S32E). In addition, we observed a ^1^H-^13^C HMBC cross peak between protons on the CH_2_ carbamidomethyl group and the β-carbon of Cys (Figure S32E). Concurrently, the ^1^H-^1^H TOCSY spectrum revealed a spin system consistent with unmodified Cys (Figure S32E). This conclusion was further bolstered by Marfey’s analysis of the *S*-alkylated tripeptide (Figure S33-S34). Authentic standards of L-Cys and D-Cys were reacted with iodoacetamide and Marfey’s reagent. The alkylated Cys residue in ApyHI-modified ApyA after acid hydrolysis and Marfey’s derivatization coeluted with the derivatized L-Cys standard, confirming that the conserved Cys residue was indeed unmodified.

To investigate any possible role of the conserved Cys on ApyA in supporting the activity of ApyHI on the C-terminal Asp, we substituted the Cys with Ser, Ala, Asp, and His. In each case, the activity of ApyHI was abolished, underscoring the importance of the conserved Cys (Figures S35-S37). Replacement of the Gly in the CBX_2-3_G motif with Ala led to a reduced modification efficiency, while Val substitution resulted in no detectable ApyA modification (Figures S35-S36). Thus, while the conserved CBX_2-3_G motif is not chemically modified in the product, it is important for ApyHI activity on ApyA.

## Discussion

Using the class-agnostic RRE domain to identify BGCs encoding an unprecedented arrangement of metalloenzymes, we identified an intriguing case from *B. thailandensis* E264. Similar BGCs were found in Pseudomonodota and Actinomycetota. Among the encoded metalloenzymes, the selected BGC encodes ApyD, a B12-dependent rSAM enzyme. Other members of this family have been reported to perform methylation, C-S bond formation, C-P bond formation, ring contraction, and C-C cyclization.^75^ To the best of our knowledge, ApyD is the first enzyme reported to methylate the Cβ of Tyr. In contrast, ApyO catalyzes biaryl crosslinking between two Tyr residues, a reaction that has been observed with the non-RiPP natural products arylomycin (AryC)^76^ and mycocyclosin (CYP121),^77^ albeit with different substrates and stereochemical outcomes.^78,79^ C-C bond formations involving two aromatic rings have been reported to be catalyzed by cytochrome P450 enzymes in RiPP biosynthesis,^45,48,80,81^ with a recent example (SlyP) catalyzing a similar crosslink as ApyO but without information on atropisomerism.^80^ Some of the precursor peptides in the identified BGCs here have other aromatic residues in the 2^nd^ and 4^th^ positions from the C-terminus (Trp/Tyr; Supplementary Dataset 1), suggesting that other biaryl couplings will be formed. Since P450s are not fully conserved among the BGCs we identified, the RiPPs described here are not classifiable as biarylitides.^21,82^

Perhaps the most interesting modification observed was catalyzed by ApyHI (a member of the multinuclear non-heme iron-dependent oxidative enzyme superfamily, MNIO), which are universally conserved among the BGCs and thus considered class-defining. Previously characterized MNIOs catalyze distinct chemical reactions exclusively on Cys, although a very recent preprint suggests an MNIO catalyzes N–Cα bond cleavage of the penultimate Asn to generate a C-terminally amidated peptide in a methanobactin-like pathway.^83^ In contrast, ApyHI performs an unprecedented enzymatic modification on a C-terminal Asp. We propose a mechanism for the MNIO-catalyzed reaction that first involves a four-electron oxidation of the β-carbon to form a ketone (Figure S37), consistent with the four-electron oxidations catalyzed by all previously characterized MNIOs.^16,26,55^ This mechanism involves hydroxylation at the β-carbon followed by radical formation at the same carbon (Figure S37). This radical intermediate may undergo proton-coupled electron transfer to form a ketone (PCET),^84^ and the resulting β-keto acid could then undergo decarboxylation to yield the observed C-terminal aminopyruvic acid. Alternatively, the radical intermediate could decarboxylate^85-88^ to form an enol that would tautomerize and form the α-keto-acid product (Figure S37). This structure is an oxidized analog of β-alanine, and after *C*-methylation by ApyS an analog of β-homoalanine is formed. Hence, ApyHI provides a route to β-amino acids containing a ketone functionality at the C-terminus of a ribosomally synthesized peptide, akin to the function of spliceases, rSAM enzymes that form such structures within a peptide.^15^ Further work will be necessary to substantiate the proposed enzymatic mechanism for ApyHI.

It is common to utilize to *E. coli* as the heterologous host when characterizing RiPP BGCs. While use of *E. coli* remains highly successful, it is an insufficient vessel to obtain functional enzymes from every desired biosynthetic pathway.^89-91^ We demonstrated here the successful utilization of a *Burkholderia* host to obtain active biosynthetic enzymes from a novel *Burkholderia*-derived BGC. This example illustrates the value of looking beyond *E. coli* for future RiPP characterization.^92-94^ This study also showcases the discovery of new enzyme functions by mining for RiPP BGCs, which contain information on both substrate(s) and biosynthetic enzymes.

In conclusion, we report the bioinformatic discovery and experimental characterization of a novel RiPP class with modifications coming from three distinct families of metalloenzymes. We functionally expressed each enzyme, elucidated the structures of the resulting products by HR-ESI-MS/MS, 2D-NMR, and MicroED, and identified the stereochemistry of the transformations. Our data suggest that the activity of ApyS is dependent on the α-keto acid moiety installed by ApyHI (Figure S5-8). Furthermore, the removal of ApyO resulted in the abolishment of the modifications catalyzed by ApyHI and ApyS (Figure S25-26). We thus suggest the post-translational pathway to start with the installation of the (2*S*,3*R*)-β-methyltyrosine by the B12-dependent rSAM enzyme ApyD and *R*_*a*_ biaryl crosslink formation by the cytochrome P450 ApyO. These steps are apparently independent of each other. Next, the C-terminal Asp residue is converted to aminopyruvic acid by the MNIO ApyHI. Lastly, methylation of the β-carbon of the aminopyruvic acid by the methyltransferase ApyS forms a C-terminal *(S)*-3-amino-2-oxobutanoic acid (Figure 6).

**Figure 6.**
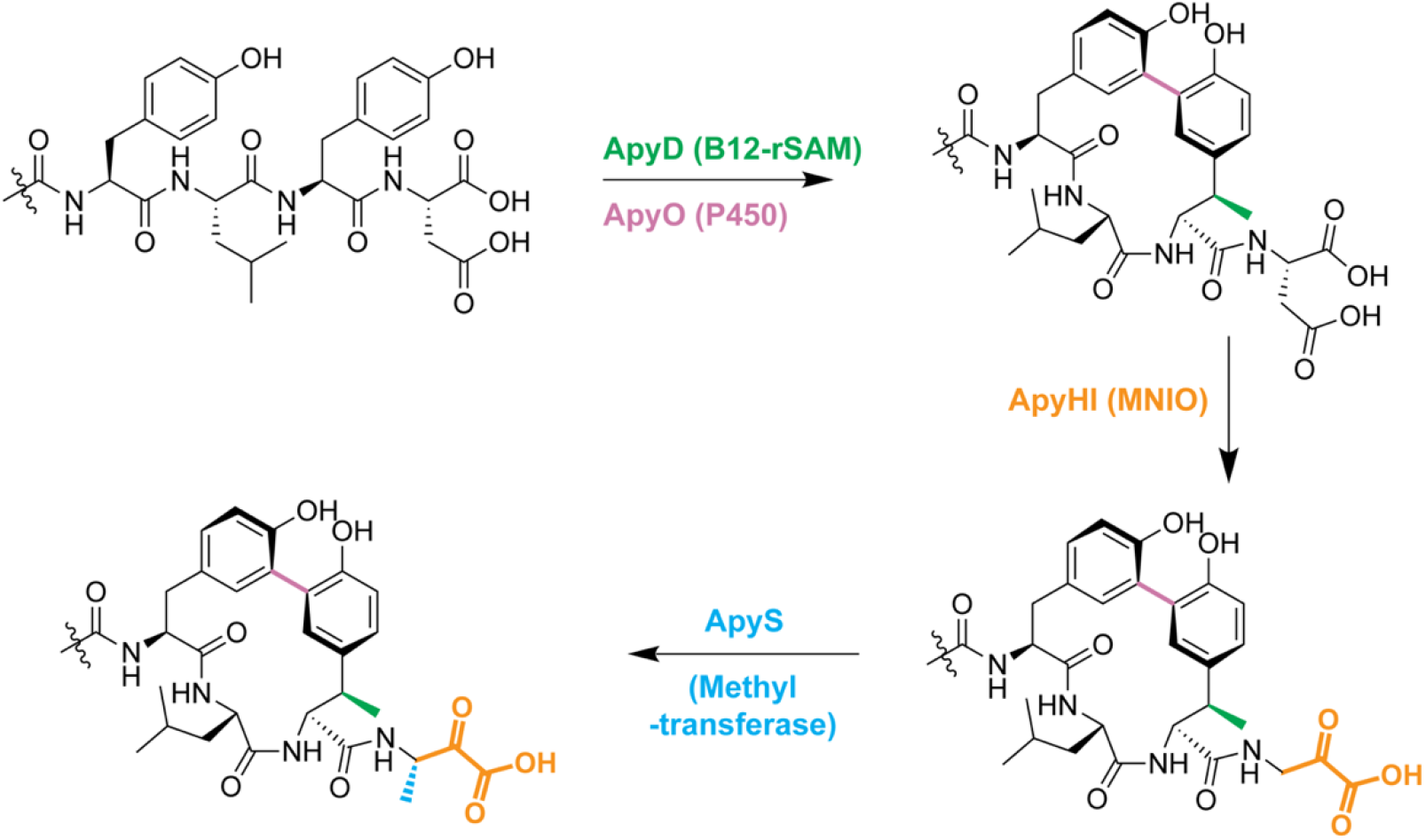
The proposed biosynthetic pathway for the *apy* BGC from *B. thailandensis* E264. B12-rSAM: cobalamin- and radical S-adenosyl-L-methionine-dependent enzyme. P450: cytochrome P450 enzyme. MNIO: multinuclear non-heme iron-dependent oxidative enzyme.

## Supporting information

Annotated BGCs

RODEO Output

## AUTHOR CONTRIBUTION

D.T.N., D.A.M., and W.A.V. designed research. D.T.N. designed and performed experiments and wrote the first draft of the manuscript. D.T.N. and L.Z. performed NMR data acquisition and analysis. D.L.G. and T.J.W. solved the MicroED structure. D.T.N. and C.P. performed crystallization screening. D.T.N. and K.M.F. collected MicroED data. D.A.M. and W.A.V. supervised the study. D.T.N., D.A.M., and W.A.V. wrote the manuscript with input from all authors.

## AUTHOR INFORMATION

### Authors

Dinh T. Nguyen – Carl R. Woese Institute of Genomic Biology, Department of Chemistry and Howard Hughes Medical Institute, University of Illinois at Urbana-Champaign, Urbana, IL, United States; orcid.org/0000-0002-2101-6808.

Lingyang Zhu – School of Chemical Sciences NMR Laboratory, University of Illinois at Urbana-Champaign, Urbana, IL, United States; orcid.org/0000-0002-6657-271X.

Danielle L. Gray – School of Chemical Sciences George L. Clark X-Ray Facility and 3M Materials Laboratory, University of Illinois at Urbana-Champaign, Urbana, IL, United States; orcid.org/0000-0003-0059-2096

Toby J. Woods – School of Chemical Sciences George L. Clark X-Ray Facility and 3M Materials Laboratory, University of Illinois at Urbana-Champaign, Urbana, IL, United States; orcid.org/0000-0002-1737-811X

Chandrashekhar Padhi – Department of Chemistry, University of Illinois at Urbana-Champaign, Urbana, IL, United States; orcid.org/0009-0009-1305-0377

Kristen M. Flatt – Materials Research Laboratory, University of Illinois at Urbana-Champaign, Urbana, IL, United States.

### Funding

This work was supported in part by a grant from the National Institutes of Health (AI144967 to D.A.M. and W.A.vdD.). W.A.vdD is an Investigator of the Howard Hughes Medical Institute.

### Notes

The authors declare there are no competing financial interests.

## Supporting Information

Experimental procedures, Figures S1-S38 and Tables S1-S9 showing spectroscopic data and gene sequences. Supplementary Datasets 1-3 containing nucleotide sequences, mass spectral data, and bioinformatic output. MicroED structural data in CIF format, These materials are associated with this manuscript and available online.

## ACKNOWLEDGMENTS

The authors thank Prof. Alessandra Eustáquio for generously providing *Burkholderia* sp. FERM BP-3421. The MicroED data collection was carried out in part in the Materials Research Laboratory Central Research Facilities, University of Illinois. We thanked the crystallization screening service at National High-Throughput Crystallization Center, which is supported through NIH grant R24GM141256. We also thank Yue Yu, Youran Luo, Mayuresh Gadgil, Shravan Dommaraju, and Dr. Lonnie A. Harris for discussions on the project.

---

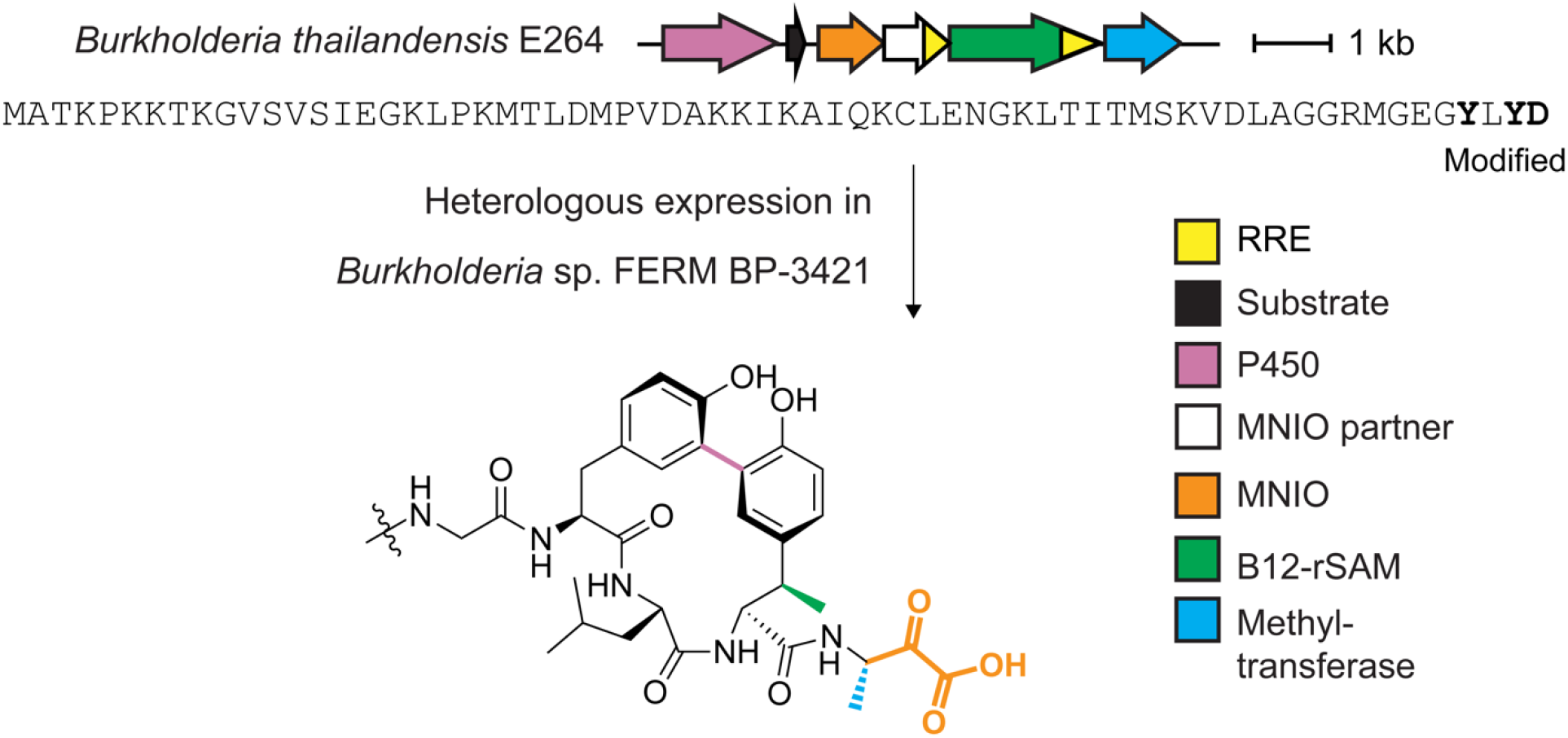

## Notes

### Competing Interest Statement

The authors have declared no competing interest.

### Summary of Updates

microED structure of ApyD/O product was added showing stereochemistry of the biaryl crosslink (new Figure 4); new authors were added that aided in microED structure determination; proposed mechanism of ApyHI was updated; additional supporting in formation figures were added to support the conclusions

