## Supplementary material for "Biosynthesis of macrocyclic peptides with C-terminal β-amino-α-keto acid groups by three different metalloenzymes": RODEO Output

### RODEO2

##### Parameters

|  |  |
| --- | --- |
| Run Time | 06:21PM on October 26, 2023 |
| Version | 2.3.3 |
| Gene Window | +/-8 CDS |
| Peptide Range | 30-300 aa |
| Fetch Distance | 150bp |
| Peptide Type | general |

Annotation Legend

| Appearance | Accession/Name |
| --- | --- |

##### Input Queries (click to navigate)

- ABC35712.1
- WP\_009907054.1
- WP\_009898131.1
- WP\_015603954.1
- WP\_059933631.1
- WP\_063534377.1
- WP\_009914071.1
- WP\_004196528.1
- WP\_038756459.1
- WP\_038754083.1
- WP\_058035158.1
- WP\_122654464.1
- WP\_208813011.1
- WP\_009981063.1
- WP\_264236925.1
- WP\_264253191.1
- WP\_045243502.1
- WP\_270956096.1
- WP\_038725437.1
- WP\_041189437.1
- WP\_038731252.1
- WP\_038763239.1
- WP\_038774393.1
- WP\_208838502.1
- WP\_208205296.1
- WP\_041195603.1
- WP\_316400961.1
- WP\_038760955.1
- TPB63544.1
- MBF3713923.1
- WP\_004557993.1
- WP\_076830717.1
- VBN64128.1
- EIP86375.1
- WP\_004194515.1
- CAG1017483.1
- KGU87554.1
- MBL8238336.1
- MBX3740453.1
- MBN8214163.1
- WP\_263545016.1
- NOT88279.1
- WP\_114844792.1
- MBW8851695.1
- WP\_162002680.1
- WP\_167928014.1
- WP\_209416722.1
- TYC66662.1
- WP\_129271392.1
- MBP1097539.1
- MQY12875.1
- WP\_153452477.1
- WP\_134006280.1
- WP\_224586504.1
- WP\_160330237.1
- MBV8367743.1
- WP\_306383405.1
- WP\_257551926.1
- MDJ0853098.1
- WP\_141343163.1
- WP\_299281768.1
- MTH36435.1
- WP\_170294018.1
- MBP6189622.1
- WP\_074971115.1
- MDI9342369.1
- WP\_083412975.1
- MDH5676845.1

#### Results for ABC35712.1 [Burkholderia thailandensis E264] back to top

#### previous - next

##### Architecture

1000 nucleotides

Link to nucleotide sequence  
  

| Accession | start | end | direction | length (aa) | Pfam/HMM | name | description | E-value |
| --- | --- | --- | --- | --- | --- | --- | --- | --- |
| ABC34322.1 | 19388 | 18341 | - | 348 | PF06314 | ADC | Acetoacetate decarboxylase (ADC) | 1.60E-78 |
| ABC35395.1 | 19492 | 20155 | + | 220 | PF01569 PF14378 | PAP2 PAP2\_3 | PAP2 superfamily PAP2 superfamily | 9.00E-18 2.90E-06 |
| ABC34252.1 | 20936 | 20183 | - | 250 | NO PFAM MATCH | - | - | - |
| ABC35917.1 | 22075 | 21247 | - | 275 | PF04954 PF08021 | SIP FAD\_binding\_9 | Siderophore-interacting protein Siderophore-interacting FAD-binding domain | 2.10E-38 4.80E-37 |
| ABC34311.1 | 22822 | 22114 | - | 235 | PF03551 TIGR03433 | PadR TIGR03433 | Transcriptional regulator PadR-like family padR\_acidobact: transcriptional regulator, Acidobacterial, PadR-family | 1.70E-19 2.60E-12 |
| ABC35679.1 | 22927 | 23764 | + | 278 | NO PFAM MATCH | - | - | - |
| ABC35200.1 | 23760 | 25167 | + | 468 | PF00067 TIGR04538 TIGR04515 TIGR04458 | p450 TIGR04538 TIGR04515 TIGR04458 | Cytochrome P450 P450\_cycloAA\_1: cytochrome P450, cyclodipeptide synthase-associated P450\_rel\_GT\_act: P450-derived glycosyltranferase activator CYP450\_TxtE: 4-nitrotryptophan synthase | 1.70E-36 1.90E-16 1.40E-07 3.70E-07 |
| ABC34935.1 | 25318 | 25519 | + | 66 | NO PFAM MATCH | - | - | - |
| ABC35712.1 | 25688 | 26540 | + | 283 | PF05114 PF01261 | DUF692 AP\_endonuc\_2 | Protein of unknown function (DUF692) Xylose isomerase-like TIM barrel | 1.00E-39 6.00E-05 |
| ABC34269.1 | 26536 | 27370 | + | 277 | NO PFAM MATCH | - | - | - |
| ABC34219.1 | 27366 | 29310 | + | 647 | TIGR02026 TIGR03471 TIGR03975 TIGR04014 TIGR04013 | TIGR02026 TIGR03471 TIGR03975 TIGR04014 TIGR04013 | BchE: magnesium-protoporphyrin IX monomethyl ester anaerobic oxidative cyclase HpnJ: hopanoid biosynthesis associated radical SAM protein HpnJ rSAM\_ocin\_1: ribosomal peptide maturation radical SAM protein 1 B12\_SAM\_MJ\_0865: B12-binding domain/radical SAM domain protein, MJ\_0865 family B12\_SAM\_MJ\_1487: B12-binding domain/radical SAM domain protein, MJ\_1487 family | 1.20E-39 3.50E-34 2.90E-28 3.80E-27 7.60E-26 |
| ABC34725.1 | 29326 | 30313 | + | 328 | TIGR04543 PF13649 PF13847 TIGR03534 PF08241 | TIGR04543 Methyltransf\_25 Methyltransf\_31 TIGR03534 Methyltransf\_11 | ketoArg\_3Met: 2-ketoarginine methyltransferase Methyltransferase domain Methyltransferase domain RF\_mod\_PrmC: protein-(glutamine-N5) methyltransferase, release factor-specific Methyltransferase domain | 1.30E-12 5.70E-08 2.60E-07 7.80E-06 1.30E-05 |
| ABC35534.1 | 30721 | 30547 | - | 57 | NO PFAM MATCH | - | - | - |
| ABC34850.1 | 31378 | 30757 | - | 206 | TIGR00948 PF01810 TIGR00949 | TIGR00948 LysE TIGR00949 | 2a75: L-lysine exporter LysE type translocator 2A76: homoserine/Threonine efflux protein | 5.60E-53 1.20E-36 6.80E-07 |
| ABC35652.1 | 31485 | 32379 | + | 297 | TIGR03298 PF03466 PF00126 TIGR02424 TIGR03418 | TIGR03298 LysR\_substrate HTH\_1 TIGR02424 TIGR03418 | argP: transcriptional regulator, ArgP family LysR substrate binding domain Bacterial regulatory helix-turn-helix protein, lysR family TF\_pcaQ: pca operon transcription factor PcaQ chol\_sulf\_TF: putative choline sulfate-utilization transcription factor | 4.20E-119 7.90E-20 2.90E-14 4.00E-12 2.50E-11 |
| ABC36149.1 | 33185 | 32429 | - | 251 | PF13545 PF00027 PF00325 RREFamAux006 TIGR03697 | HTH\_Crp\_2 cNMP\_binding Crp RRE\_cNMP TIGR03697 | Crp-like helix-turn-helix domain Cyclic nucleotide-binding domain Bacterial regulatory proteins, crp family - NtcA\_cyano: global nitrogen regulator NtcA | 3.50E-20 6.00E-18 2.20E-14 2.60E-08 1.40E-05 |
| ABC34752.1 | 33802 | 33334 | - | 155 | PF00582 | Usp | Universal stress protein family | 3.50E-31 |

#### Results for WP\_009907054.1 [Burkholderia thailandensis] back to top

#### previous - next

##### Architecture

1000 nucleotides

Link to nucleotide sequence  
  

| Accession | start | end | direction | length (aa) | Pfam/HMM | name | description | E-value |
| --- | --- | --- | --- | --- | --- | --- | --- | --- |
| WP\_009894574.1 | 62726 | 62495 | - | 76 | NO PFAM MATCH | - | - | - |
| WP\_009894573.1 | 63146 | 63614 | + | 155 | PF00582 | Usp | Universal stress protein family | 3.50E-31 |
| WP\_009894572.1 | 63763 | 64519 | + | 251 | PF13545 PF00027 PF00325 RREFamAux006 TIGR03697 | HTH\_Crp\_2 cNMP\_binding Crp RRE\_cNMP TIGR03697 | Crp-like helix-turn-helix domain Cyclic nucleotide-binding domain Bacterial regulatory proteins, crp family - NtcA\_cyano: global nitrogen regulator NtcA | 3.50E-20 6.00E-18 2.20E-14 2.60E-08 1.40E-05 |
| WP\_009907056.1 | 65463 | 64569 | - | 297 | TIGR03298 PF03466 PF00126 TIGR02424 TIGR03418 | TIGR03298 LysR\_substrate HTH\_1 TIGR02424 TIGR03418 | argP: transcriptional regulator, ArgP family LysR substrate binding domain Bacterial regulatory helix-turn-helix protein, lysR family TF\_pcaQ: pca operon transcription factor PcaQ chol\_sulf\_TF: putative choline sulfate-utilization transcription factor | 5.10E-120 7.90E-20 5.50E-15 9.00E-13 9.90E-12 |
| WP\_009894570.1 | 65570 | 66191 | + | 206 | TIGR00948 PF01810 TIGR00949 | TIGR00948 LysE TIGR00949 | 2a75: L-lysine exporter LysE type translocator 2A76: homoserine/Threonine efflux protein | 5.60E-53 1.20E-36 6.80E-07 |
| WP\_009894566.1 | 67625 | 66635 | - | 329 | TIGR04543 PF13649 PF13847 TIGR03534 PF08241 | TIGR04543 Methyltransf\_25 Methyltransf\_31 TIGR03534 Methyltransf\_11 | ketoArg\_3Met: 2-ketoarginine methyltransferase Methyltransferase domain Methyltransferase domain RF\_mod\_PrmC: protein-(glutamine-N5) methyltransferase, release factor-specific Methyltransferase domain | 1.30E-12 5.70E-08 2.60E-07 7.80E-06 1.30E-05 |
| WP\_009894565.1 | 69582 | 67638 | - | 647 | TIGR02026 TIGR03471 TIGR03975 TIGR04014 TIGR04013 | TIGR02026 TIGR03471 TIGR03975 TIGR04014 TIGR04013 | BchE: magnesium-protoporphyrin IX monomethyl ester anaerobic oxidative cyclase HpnJ: hopanoid biosynthesis associated radical SAM protein HpnJ rSAM\_ocin\_1: ribosomal peptide maturation radical SAM protein 1 B12\_SAM\_MJ\_0865: B12-binding domain/radical SAM domain protein, MJ\_0865 family B12\_SAM\_MJ\_1487: B12-binding domain/radical SAM domain protein, MJ\_1487 family | 1.20E-39 3.50E-34 2.90E-28 3.80E-27 7.60E-26 |
| WP\_009894564.1 | 70412 | 69578 | - | 277 | NO PFAM MATCH | - | - | - |
| WP\_009907054.1 | 71260 | 70408 | - | 283 | PF05114 PF01261 | DUF692 AP\_endonuc\_2 | Protein of unknown function (DUF692) Xylose isomerase-like TIM barrel | 7.10E-40 6.00E-05 |
| WP\_004201048.1 | 71630 | 71429 | - | 66 | NO PFAM MATCH | - | - | - |
| WP\_009894540.1 | 73188 | 71781 | - | 468 | PF00067 TIGR04538 TIGR04515 TIGR04458 | p450 TIGR04538 TIGR04515 TIGR04458 | Cytochrome P450 P450\_cycloAA\_1: cytochrome P450, cyclodipeptide synthase-associated P450\_rel\_GT\_act: P450-derived glycosyltranferase activator CYP450\_TxtE: 4-nitrotryptophan synthase | 1.70E-36 1.90E-16 1.40E-07 3.70E-07 |
| WP\_230631267.1 | 73565 | 73184 | - | 126 | NO PFAM MATCH | - | - | - |
| WP\_025992048.1 | 74125 | 74833 | + | 235 | PF03551 TIGR03433 | PadR TIGR03433 | Transcriptional regulator PadR-like family padR\_acidobact: transcriptional regulator, Acidobacterial, PadR-family | 1.70E-19 2.70E-12 |
| WP\_009894531.1 | 74872 | 75700 | + | 275 | PF04954 PF08021 | SIP FAD\_binding\_9 | Siderophore-interacting protein Siderophore-interacting FAD-binding domain | 2.10E-38 4.80E-37 |
| WP\_223297137.1 | 76240 | 76006 | - | 77 | NO PFAM MATCH | - | - | - |
| WP\_009894529.1 | 76527 | 76764 | + | 78 | NO PFAM MATCH | - | - | - |
| WP\_009894528.1 | 77389 | 76792 | - | 198 | PF01569 PF14378 | PAP2 PAP2\_3 | PAP2 superfamily PAP2 superfamily | 6.80E-18 1.80E-06 |

#### Results for WP\_009898131.1 [Burkholderia thailandensis] back to top

#### previous - next

##### Architecture

1000 nucleotides

Link to nucleotide sequence  
  

| Accession | start | end | direction | length (aa) | Pfam/HMM | name | description | E-value |
| --- | --- | --- | --- | --- | --- | --- | --- | --- |
| WP\_009894527.1 | 945 | 204 | - | 246 | PF06314 | ADC | Acetoacetate decarboxylase (ADC) | 5.50E-79 |
| WP\_009894528.1 | 1421 | 2018 | + | 198 | PF01569 PF14378 | PAP2 PAP2\_3 | PAP2 superfamily PAP2 superfamily | 6.80E-18 1.80E-06 |
| WP\_009894529.1 | 2283 | 2046 | - | 78 | NO PFAM MATCH | - | - | - |
| WP\_019254851.1 | 3948 | 3120 | - | 275 | PF04954 PF08021 | SIP FAD\_binding\_9 | Siderophore-interacting protein Siderophore-interacting FAD-binding domain | 4.20E-38 1.70E-37 |
| WP\_029227139.1 | 4713 | 3987 | - | 241 | PF03551 TIGR03433 | PadR TIGR03433 | Transcriptional regulator PadR-like family padR\_acidobact: transcriptional regulator, Acidobacterial, PadR-family | 1.80E-19 2.90E-12 |
| WP\_269241812.1 | 5226 | 5655 | + | 142 | NO PFAM MATCH | - | - | - |
| WP\_009898130.1 | 5651 | 7058 | + | 468 | PF00067 TIGR04538 TIGR04515 TIGR04458 | p450 TIGR04538 TIGR04515 TIGR04458 | Cytochrome P450 P450\_cycloAA\_1: cytochrome P450, cyclodipeptide synthase-associated P450\_rel\_GT\_act: P450-derived glycosyltranferase activator CYP450\_TxtE: 4-nitrotryptophan synthase | 1.60E-37 3.80E-16 1.30E-07 5.80E-07 |
| WP\_004201048.1 | 7209 | 7410 | + | 66 | NO PFAM MATCH | - | - | - |
| WP\_009898131.1 | 7579 | 8431 | + | 283 | PF05114 PF01261 | DUF692 AP\_endonuc\_2 | Protein of unknown function (DUF692) Xylose isomerase-like TIM barrel | 1.30E-39 5.70E-05 |
| WP\_019254854.1 | 8427 | 9261 | + | 277 | NO PFAM MATCH | - | - | - |
| WP\_025405954.1 | 9257 | 11201 | + | 647 | TIGR02026 TIGR03471 TIGR03975 TIGR04014 TIGR04013 | TIGR02026 TIGR03471 TIGR03975 TIGR04014 TIGR04013 | BchE: magnesium-protoporphyrin IX monomethyl ester anaerobic oxidative cyclase HpnJ: hopanoid biosynthesis associated radical SAM protein HpnJ rSAM\_ocin\_1: ribosomal peptide maturation radical SAM protein 1 B12\_SAM\_MJ\_0865: B12-binding domain/radical SAM domain protein, MJ\_0865 family B12\_SAM\_MJ\_1487: B12-binding domain/radical SAM domain protein, MJ\_1487 family | 3.20E-39 3.10E-34 1.30E-27 1.20E-26 8.90E-26 |
| WP\_269241806.1 | 11214 | 12204 | + | 329 | TIGR04543 PF13649 PF13847 TIGR03534 PF08241 | TIGR04543 Methyltransf\_25 Methyltransf\_31 TIGR03534 Methyltransf\_11 | ketoArg\_3Met: 2-ketoarginine methyltransferase Methyltransferase domain Methyltransferase domain RF\_mod\_PrmC: protein-(glutamine-N5) methyltransferase, release factor-specific Methyltransferase domain | 1.60E-12 3.20E-08 1.10E-07 2.90E-06 1.40E-05 |
| WP\_009898139.1 | 13269 | 12648 | - | 206 | TIGR00948 PF01810 TIGR00949 | TIGR00948 LysE TIGR00949 | 2a75: L-lysine exporter LysE type translocator 2A76: homoserine/Threonine efflux protein | 8.80E-53 2.20E-36 5.40E-07 |
| WP\_009898141.1 | 13376 | 14270 | + | 297 | TIGR03298 PF03466 PF00126 TIGR02424 TIGR03418 | TIGR03298 LysR\_substrate HTH\_1 TIGR02424 TIGR03418 | argP: transcriptional regulator, ArgP family LysR substrate binding domain Bacterial regulatory helix-turn-helix protein, lysR family TF\_pcaQ: pca operon transcription factor PcaQ chol\_sulf\_TF: putative choline sulfate-utilization transcription factor | 3.60E-120 2.80E-19 5.50E-15 2.10E-12 2.10E-11 |
| WP\_009894572.1 | 15070 | 14314 | - | 251 | PF13545 PF00027 PF00325 RREFamAux006 TIGR03697 | HTH\_Crp\_2 cNMP\_binding Crp RRE\_cNMP TIGR03697 | Crp-like helix-turn-helix domain Cyclic nucleotide-binding domain Bacterial regulatory proteins, crp family - NtcA\_cyano: global nitrogen regulator NtcA | 3.50E-20 6.00E-18 2.20E-14 2.60E-08 1.40E-05 |
| WP\_009894573.1 | 15687 | 15219 | - | 155 | PF00582 | Usp | Universal stress protein family | 3.50E-31 |
| WP\_009898144.1 | 16107 | 16338 | + | 76 | NO PFAM MATCH | - | - | - |

#### Results for WP\_015603954.1 [Burkholderia humptydooensis] back to top

#### previous - next

##### Architecture

1000 nucleotides

Link to nucleotide sequence  
  

| Accession | start | end | direction | length (aa) | Pfam/HMM | name | description | E-value |
| --- | --- | --- | --- | --- | --- | --- | --- | --- |
| WP\_015604046.1 | 26397 | 25680 | - | 238 | PF03551 TIGR03433 | PadR TIGR03433 | Transcriptional regulator PadR-like family padR\_acidobact: transcriptional regulator, Acidobacterial, PadR-family | 2.00E-19 2.20E-12 |
| WP\_012492354.1 | 26510 | 27047 | + | 179 | INFERRED GENE | - | - | - |
| WP\_015602890.1 | 27581 | 28616 | + | 344 | PF08007 PF07883 PF13621 | JmjC\_2 Cupin\_2 Cupin\_8 | JmjC domain Cupin domain Cupin-like domain | 9.30E-27 7.90E-05 2.20E-04 |
| WP\_015603899.1 | 28605 | 29703 | + | 365 | PF07690 TIGR00900 TIGR00880 TIGR00881 TIGR00886 | MFS\_1 TIGR00900 TIGR00880 TIGR00881 TIGR00886 | Major Facilitator Superfamily 2A0121: H+ Antiporter protein 2\_A\_01\_02: multidrug resistance protein 2A0104: phosphoglycerate transporter family protein 2A0108: nitrite transporter | 1.80E-21 1.40E-19 4.30E-14 1.10E-08 1.30E-07 |
| WP\_011235908.1 | 29973 | 31240 | + | 422 | INFERRED GENE | - | - | - |
| WP\_017331931.1 | 31524 | 31836 | + | 104 | INFERRED GENE | - | - | - |
| WP\_041861993.1 | 32126 | 33536 | + | 469 | PF00067 TIGR04538 TIGR04515 TIGR04458 | p450 TIGR04538 TIGR04515 TIGR04458 | Cytochrome P450 P450\_cycloAA\_1: cytochrome P450, cyclodipeptide synthase-associated P450\_rel\_GT\_act: P450-derived glycosyltranferase activator CYP450\_TxtE: 4-nitrotryptophan synthase | 9.60E-35 3.90E-14 1.20E-07 1.20E-06 |
| WP\_006028385.1 | 33686 | 33887 | + | 66 | NO PFAM MATCH | - | - | - |
| WP\_015603954.1 | 33998 | 34850 | + | 283 | PF05114 PF01261 | DUF692 AP\_endonuc\_2 | Protein of unknown function (DUF692) Xylose isomerase-like TIM barrel | 5.50E-41 1.10E-04 |
| WP\_015603030.1 | 34846 | 35680 | + | 277 | NO PFAM MATCH | - | - | - |
| WP\_015604052.1 | 35676 | 37620 | + | 647 | TIGR02026 TIGR03471 TIGR03975 TIGR04014 TIGR04013 | TIGR02026 TIGR03471 TIGR03975 TIGR04014 TIGR04013 | BchE: magnesium-protoporphyrin IX monomethyl ester anaerobic oxidative cyclase HpnJ: hopanoid biosynthesis associated radical SAM protein HpnJ rSAM\_ocin\_1: ribosomal peptide maturation radical SAM protein 1 B12\_SAM\_MJ\_0865: B12-binding domain/radical SAM domain protein, MJ\_0865 family B12\_SAM\_MJ\_1487: B12-binding domain/radical SAM domain protein, MJ\_1487 family | 1.10E-40 2.10E-34 1.60E-28 1.00E-25 3.00E-24 |
| WP\_015603256.1 | 37633 | 38623 | + | 329 | TIGR04543 PF13649 PF13847 PF08242 PF08241 | TIGR04543 Methyltransf\_25 Methyltransf\_31 Methyltransf\_12 Methyltransf\_11 | ketoArg\_3Met: 2-ketoarginine methyltransferase Methyltransferase domain Methyltransferase domain Methyltransferase domain Methyltransferase domain | 1.60E-12 1.10E-08 1.10E-07 1.20E-06 1.40E-05 |
| WP\_015603823.1 | 39479 | 38858 | - | 206 | TIGR00948 PF01810 TIGR00949 | TIGR00948 LysE TIGR00949 | 2a75: L-lysine exporter LysE type translocator 2A76: homoserine/Threonine efflux protein | 1.70E-53 7.80E-37 2.00E-06 |
| WP\_006028391.1 | 39586 | 40480 | + | 297 | TIGR03298 PF03466 PF00126 TIGR02424 TIGR03418 | TIGR03298 LysR\_substrate HTH\_1 TIGR02424 TIGR03418 | argP: transcriptional regulator, ArgP family LysR substrate binding domain Bacterial regulatory helix-turn-helix protein, lysR family TF\_pcaQ: pca operon transcription factor PcaQ chol\_sulf\_TF: putative choline sulfate-utilization transcription factor | 4.70E-121 1.10E-19 5.50E-15 1.20E-12 3.60E-11 |
| WP\_009914074.1 | 41280 | 40524 | - | 251 | PF13545 PF00027 PF00325 RREFamAux006 TIGR03697 | HTH\_Crp\_2 cNMP\_binding Crp RRE\_cNMP TIGR03697 | Crp-like helix-turn-helix domain Cyclic nucleotide-binding domain Bacterial regulatory proteins, crp family - NtcA\_cyano: global nitrogen regulator NtcA | 3.50E-20 4.40E-18 2.20E-14 2.70E-08 1.90E-05 |
| WP\_006028393.1 | 41897 | 41429 | - | 155 | PF00582 | Usp | Universal stress protein family | 1.40E-31 |
| WP\_009914076.1 | 42240 | 42471 | + | 76 | NO PFAM MATCH | - | - | - |

#### Results for WP\_059933631.1 [Burkholderia sp. MSMB1588] back to top

#### previous - next

##### Architecture

1000 nucleotides

Link to nucleotide sequence  
  

| Accession | start | end | direction | length (aa) | Pfam/HMM | name | description | E-value |
| --- | --- | --- | --- | --- | --- | --- | --- | --- |
| WP\_059933641.1 | 786769 | 786061 | - | 235 | PF03551 TIGR03433 | PadR TIGR03433 | Transcriptional regulator PadR-like family padR\_acidobact: transcriptional regulator, Acidobacterial, PadR-family | 2.00E-19 2.10E-12 |
| WP\_012492354.1 | 786882 | 787413 | + | 177 | INFERRED GENE | - | - | - |
| WP\_141667956.1 | 787914 | 788982 | + | 355 | PF08007 PF07883 PF13621 | JmjC\_2 Cupin\_2 Cupin\_8 | JmjC domain Cupin domain Cupin-like domain | 1.60E-24 8.60E-05 9.20E-04 |
| WP\_082743863.1 | 788971 | 790198 | + | 408 | PF07690 TIGR00900 TIGR00880 TIGR00881 TIGR00710 | MFS\_1 TIGR00900 TIGR00880 TIGR00881 TIGR00710 | Major Facilitator Superfamily 2A0121: H+ Antiporter protein 2\_A\_01\_02: multidrug resistance protein 2A0104: phosphoglycerate transporter family protein efflux\_Bcr\_CflA: drug resistance transporter, Bcr/CflA subfamily | 6.40E-25 5.90E-23 5.80E-14 5.30E-09 6.90E-07 |
| WP\_007179151.1 | 790313 | 791604 | + | 430 | INFERRED GENE | - | - | - |
| WP\_004532144.1 | 791847 | 792199 | + | 117 | INFERRED GENE | - | - | - |
| WP\_059933634.1 | 792439 | 793849 | + | 469 | PF00067 TIGR04538 TIGR04515 TIGR04458 | p450 TIGR04538 TIGR04515 TIGR04458 | Cytochrome P450 P450\_cycloAA\_1: cytochrome P450, cyclodipeptide synthase-associated P450\_rel\_GT\_act: P450-derived glycosyltranferase activator CYP450\_TxtE: 4-nitrotryptophan synthase | 3.40E-34 6.20E-15 6.50E-08 1.20E-06 |
| WP\_006028385.1 | 794000 | 794201 | + | 66 | NO PFAM MATCH | - | - | - |
| WP\_059933631.1 | 794312 | 795164 | + | 283 | PF05114 PF01261 | DUF692 AP\_endonuc\_2 | Protein of unknown function (DUF692) Xylose isomerase-like TIM barrel | 4.60E-41 1.30E-04 |
| WP\_059933627.1 | 795160 | 795994 | + | 277 | NO PFAM MATCH | - | - | - |
| WP\_059933624.1 | 795990 | 797934 | + | 647 | TIGR02026 TIGR03471 TIGR03975 TIGR04014 TIGR04013 | TIGR02026 TIGR03471 TIGR03975 TIGR04014 TIGR04013 | BchE: magnesium-protoporphyrin IX monomethyl ester anaerobic oxidative cyclase HpnJ: hopanoid biosynthesis associated radical SAM protein HpnJ rSAM\_ocin\_1: ribosomal peptide maturation radical SAM protein 1 B12\_SAM\_MJ\_0865: B12-binding domain/radical SAM domain protein, MJ\_0865 family B12\_SAM\_MJ\_1487: B12-binding domain/radical SAM domain protein, MJ\_1487 family | 7.80E-41 1.40E-34 1.10E-28 6.80E-26 1.80E-24 |
| WP\_009914073.1 | 797947 | 798937 | + | 329 | TIGR04543 PF13649 PF13847 PF08242 PF08241 | TIGR04543 Methyltransf\_25 Methyltransf\_31 Methyltransf\_12 Methyltransf\_11 | ketoArg\_3Met: 2-ketoarginine methyltransferase Methyltransferase domain Methyltransferase domain Methyltransferase domain Methyltransferase domain | 1.20E-12 1.10E-08 1.10E-07 1.30E-06 2.00E-05 |
| WP\_006028390.1 | 799789 | 799168 | - | 206 | TIGR00948 PF01810 TIGR00949 | TIGR00948 LysE TIGR00949 | 2a75: L-lysine exporter LysE type translocator 2A76: homoserine/Threonine efflux protein | 1.40E-53 2.40E-37 8.20E-07 |
| WP\_006028391.1 | 799896 | 800790 | + | 297 | TIGR03298 PF03466 PF00126 TIGR02424 TIGR03418 | TIGR03298 LysR\_substrate HTH\_1 TIGR02424 TIGR03418 | argP: transcriptional regulator, ArgP family LysR substrate binding domain Bacterial regulatory helix-turn-helix protein, lysR family TF\_pcaQ: pca operon transcription factor PcaQ chol\_sulf\_TF: putative choline sulfate-utilization transcription factor | 4.70E-121 1.10E-19 5.50E-15 1.20E-12 3.60E-11 |
| WP\_009914074.1 | 801590 | 800834 | - | 251 | PF13545 PF00027 PF00325 RREFamAux006 TIGR03697 | HTH\_Crp\_2 cNMP\_binding Crp RRE\_cNMP TIGR03697 | Crp-like helix-turn-helix domain Cyclic nucleotide-binding domain Bacterial regulatory proteins, crp family - NtcA\_cyano: global nitrogen regulator NtcA | 3.50E-20 4.40E-18 2.20E-14 2.70E-08 1.90E-05 |
| WP\_006028393.1 | 802207 | 801739 | - | 155 | PF00582 | Usp | Universal stress protein family | 1.40E-31 |
| WP\_009914076.1 | 802550 | 802781 | + | 76 | NO PFAM MATCH | - | - | - |

#### Results for WP\_063534377.1 [Burkholderia sp. MSMB1589WGS] back to top

#### previous - next

##### Architecture

1000 nucleotides

Link to nucleotide sequence  
  

| Accession | start | end | direction | length (aa) | Pfam/HMM | name | description | E-value |
| --- | --- | --- | --- | --- | --- | --- | --- | --- |
| WP\_015604046.1 | 1387223 | 1386506 | - | 238 | PF03551 TIGR03433 | PadR TIGR03433 | Transcriptional regulator PadR-like family padR\_acidobact: transcriptional regulator, Acidobacterial, PadR-family | 2.00E-19 2.20E-12 |
| WP\_012492354.1 | 1387336 | 1387873 | + | 179 | INFERRED GENE | - | - | - |
| WP\_075645150.1 | 1388586 | 1389444 | + | 285 | PF08007 PF07883 PF13621 | JmjC\_2 Cupin\_2 Cupin\_8 | JmjC domain Cupin domain Cupin-like domain | 1.70E-26 5.60E-05 5.00E-04 |
| WP\_082874982.1 | 1389433 | 1390660 | + | 408 | PF07690 TIGR00900 TIGR00880 TIGR00881 TIGR00886 | MFS\_1 TIGR00900 TIGR00880 TIGR00881 TIGR00886 | Major Facilitator Superfamily 2A0121: H+ Antiporter protein 2\_A\_01\_02: multidrug resistance protein 2A0104: phosphoglycerate transporter family protein 2A0108: nitrite transporter | 2.50E-24 1.50E-22 6.50E-14 4.10E-09 1.70E-07 |
| WP\_007179151.1 | 1390775 | 1392066 | + | 430 | INFERRED GENE | - | - | - |
| WP\_231892106.1 | 1392698 | 1392323 | - | 124 | TIGR02212 TIGR02213 PF12704 | TIGR02212 TIGR02213 MacB\_PCD | lolCE: lipoprotein releasing system, transmembrane protein, LolC/E family lolE\_release: lipoprotein releasing system, transmembrane protein LolE MacB-like periplasmic core domain | 3.40E-23 7.60E-20 2.00E-07 |
| WP\_063534375.1 | 1392952 | 1394362 | + | 469 | PF00067 TIGR04538 TIGR04515 TIGR04458 | p450 TIGR04538 TIGR04515 TIGR04458 | Cytochrome P450 P450\_cycloAA\_1: cytochrome P450, cyclodipeptide synthase-associated P450\_rel\_GT\_act: P450-derived glycosyltranferase activator CYP450\_TxtE: 4-nitrotryptophan synthase | 1.80E-34 2.20E-14 4.20E-07 1.30E-06 |
| WP\_063534376.1 | 1394512 | 1394713 | + | 66 | NO PFAM MATCH | - | - | - |
| WP\_063534377.1 | 1394824 | 1395676 | + | 283 | PF05114 PF01261 | DUF692 AP\_endonuc\_2 | Protein of unknown function (DUF692) Xylose isomerase-like TIM barrel | 7.30E-41 1.10E-04 |
| WP\_009914072.1 | 1395672 | 1396506 | + | 277 | NO PFAM MATCH | - | - | - |
| WP\_063534378.1 | 1396502 | 1398446 | + | 647 | TIGR02026 TIGR03471 TIGR03975 TIGR04014 TIGR04013 | TIGR02026 TIGR03471 TIGR03975 TIGR04014 TIGR04013 | BchE: magnesium-protoporphyrin IX monomethyl ester anaerobic oxidative cyclase HpnJ: hopanoid biosynthesis associated radical SAM protein HpnJ rSAM\_ocin\_1: ribosomal peptide maturation radical SAM protein 1 B12\_SAM\_MJ\_0865: B12-binding domain/radical SAM domain protein, MJ\_0865 family B12\_SAM\_MJ\_1487: B12-binding domain/radical SAM domain protein, MJ\_1487 family | 2.20E-40 8.50E-35 6.80E-29 2.50E-26 5.40E-25 |
| WP\_063534379.1 | 1398459 | 1399449 | + | 329 | TIGR04543 PF13649 PF13847 PF08242 PF08241 | TIGR04543 Methyltransf\_25 Methyltransf\_31 Methyltransf\_12 Methyltransf\_11 | ketoArg\_3Met: 2-ketoarginine methyltransferase Methyltransferase domain Methyltransferase domain Methyltransferase domain Methyltransferase domain | 1.70E-12 1.10E-08 1.10E-07 1.20E-06 8.60E-06 |
| WP\_063534380.1 | 1400309 | 1399688 | - | 206 | TIGR00948 PF01810 TIGR00949 | TIGR00948 LysE TIGR00949 | 2a75: L-lysine exporter LysE type translocator 2A76: homoserine/Threonine efflux protein | 1.70E-53 9.70E-37 9.60E-07 |
| WP\_006028391.1 | 1400416 | 1401310 | + | 297 | TIGR03298 PF03466 PF00126 TIGR02424 TIGR03418 | TIGR03298 LysR\_substrate HTH\_1 TIGR02424 TIGR03418 | argP: transcriptional regulator, ArgP family LysR substrate binding domain Bacterial regulatory helix-turn-helix protein, lysR family TF\_pcaQ: pca operon transcription factor PcaQ chol\_sulf\_TF: putative choline sulfate-utilization transcription factor | 4.70E-121 1.10E-19 5.50E-15 1.20E-12 3.60E-11 |
| WP\_009914074.1 | 1402110 | 1401354 | - | 251 | PF13545 PF00027 PF00325 RREFamAux006 TIGR03697 | HTH\_Crp\_2 cNMP\_binding Crp RRE\_cNMP TIGR03697 | Crp-like helix-turn-helix domain Cyclic nucleotide-binding domain Bacterial regulatory proteins, crp family - NtcA\_cyano: global nitrogen regulator NtcA | 3.50E-20 4.40E-18 2.20E-14 2.70E-08 1.90E-05 |
| WP\_006028393.1 | 1402727 | 1402259 | - | 155 | PF00582 | Usp | Universal stress protein family | 1.40E-31 |
| WP\_009914076.1 | 1403068 | 1403299 | + | 76 | NO PFAM MATCH | - | - | - |

#### Results for WP\_009914071.1 [Burkholderia humptydooensis] back to top

#### previous - next

##### Architecture

1000 nucleotides

Link to nucleotide sequence  
  

| Accession | start | end | direction | length (aa) | Pfam/HMM | name | description | E-value |
| --- | --- | --- | --- | --- | --- | --- | --- | --- |
| WP\_043282959.1 | 2085351 | 2084634 | - | 238 | PF03551 TIGR03433 | PadR TIGR03433 | Transcriptional regulator PadR-like family padR\_acidobact: transcriptional regulator, Acidobacterial, PadR-family | 2.00E-19 2.20E-12 |
| WP\_012492354.1 | 2085464 | 2085995 | + | 177 | INFERRED GENE | - | - | - |
| WP\_157225898.1 | 2086483 | 2087440 | + | 318 | PF08007 PF07883 PF13621 | JmjC\_2 Cupin\_2 Cupin\_8 | JmjC domain Cupin domain Cupin-like domain | 1.20E-22 1.00E-04 6.70E-04 |
| WP\_080595165.1 | 2087520 | 2088747 | + | 408 | PF07690 TIGR00900 TIGR00880 TIGR00881 TIGR00886 | MFS\_1 TIGR00900 TIGR00880 TIGR00881 TIGR00886 | Major Facilitator Superfamily 2A0121: H+ Antiporter protein 2\_A\_01\_02: multidrug resistance protein 2A0104: phosphoglycerate transporter family protein 2A0108: nitrite transporter | 1.50E-23 1.30E-21 1.90E-13 4.50E-08 3.40E-07 |
| WP\_011235908.1 | 2088886 | 2090165 | + | 426 | INFERRED GENE | - | - | - |
| WP\_006028383.1 | 2090761 | 2090422 | - | 112 | TIGR02212 TIGR02213 PF12704 | TIGR02212 TIGR02213 MacB\_PCD | lolCE: lipoprotein releasing system, transmembrane protein, LolC/E family lolE\_release: lipoprotein releasing system, transmembrane protein LolE MacB-like periplasmic core domain | 2.80E-23 1.20E-19 1.20E-06 |
| WP\_006028384.1 | 2091001 | 2092411 | + | 469 | PF00067 TIGR04538 TIGR04515 TIGR04458 | p450 TIGR04538 TIGR04515 TIGR04458 | Cytochrome P450 P450\_cycloAA\_1: cytochrome P450, cyclodipeptide synthase-associated P450\_rel\_GT\_act: P450-derived glycosyltranferase activator CYP450\_TxtE: 4-nitrotryptophan synthase | 9.20E-35 1.80E-14 1.70E-07 1.20E-06 |
| WP\_006028385.1 | 2092561 | 2092762 | + | 66 | NO PFAM MATCH | - | - | - |
| WP\_009914071.1 | 2092873 | 2093725 | + | 283 | PF05114 PF01261 | DUF692 AP\_endonuc\_2 | Protein of unknown function (DUF692) Xylose isomerase-like TIM barrel | 2.20E-40 1.10E-04 |
| WP\_009914072.1 | 2093721 | 2094555 | + | 277 | NO PFAM MATCH | - | - | - |
| WP\_006028388.1 | 2094551 | 2096495 | + | 647 | TIGR02026 TIGR03471 TIGR03975 TIGR04014 TIGR04013 | TIGR02026 TIGR03471 TIGR03975 TIGR04014 TIGR04013 | BchE: magnesium-protoporphyrin IX monomethyl ester anaerobic oxidative cyclase HpnJ: hopanoid biosynthesis associated radical SAM protein HpnJ rSAM\_ocin\_1: ribosomal peptide maturation radical SAM protein 1 B12\_SAM\_MJ\_0865: B12-binding domain/radical SAM domain protein, MJ\_0865 family B12\_SAM\_MJ\_1487: B12-binding domain/radical SAM domain protein, MJ\_1487 family | 2.20E-40 8.50E-35 6.80E-29 2.50E-26 5.40E-25 |
| WP\_009914073.1 | 2096508 | 2097498 | + | 329 | TIGR04543 PF13649 PF13847 PF08242 PF08241 | TIGR04543 Methyltransf\_25 Methyltransf\_31 Methyltransf\_12 Methyltransf\_11 | ketoArg\_3Met: 2-ketoarginine methyltransferase Methyltransferase domain Methyltransferase domain Methyltransferase domain Methyltransferase domain | 1.20E-12 1.10E-08 1.10E-07 1.30E-06 2.00E-05 |
| WP\_006028390.1 | 2098350 | 2097729 | - | 206 | TIGR00948 PF01810 TIGR00949 | TIGR00948 LysE TIGR00949 | 2a75: L-lysine exporter LysE type translocator 2A76: homoserine/Threonine efflux protein | 1.40E-53 2.40E-37 8.20E-07 |
| WP\_006028391.1 | 2098457 | 2099351 | + | 297 | TIGR03298 PF03466 PF00126 TIGR02424 TIGR03418 | TIGR03298 LysR\_substrate HTH\_1 TIGR02424 TIGR03418 | argP: transcriptional regulator, ArgP family LysR substrate binding domain Bacterial regulatory helix-turn-helix protein, lysR family TF\_pcaQ: pca operon transcription factor PcaQ chol\_sulf\_TF: putative choline sulfate-utilization transcription factor | 4.70E-121 1.10E-19 5.50E-15 1.20E-12 3.60E-11 |
| WP\_009914074.1 | 2100151 | 2099395 | - | 251 | PF13545 PF00027 PF00325 RREFamAux006 TIGR03697 | HTH\_Crp\_2 cNMP\_binding Crp RRE\_cNMP TIGR03697 | Crp-like helix-turn-helix domain Cyclic nucleotide-binding domain Bacterial regulatory proteins, crp family - NtcA\_cyano: global nitrogen regulator NtcA | 3.50E-20 4.40E-18 2.20E-14 2.70E-08 1.90E-05 |
| WP\_006028393.1 | 2100768 | 2100300 | - | 155 | PF00582 | Usp | Universal stress protein family | 1.40E-31 |
| WP\_009914076.1 | 2101111 | 2101342 | + | 76 | NO PFAM MATCH | - | - | - |

#### Results for WP\_004196528.1 [Burkholderia pseudomallei] back to top

#### previous - next

##### Architecture

1000 nucleotides

Link to nucleotide sequence  
  

| Accession | start | end | direction | length (aa) | Pfam/HMM | name | description | E-value |
| --- | --- | --- | --- | --- | --- | --- | --- | --- |
| WP\_004194465.1 | 55471 | 55240 | - | 76 | NO PFAM MATCH | - | - | - |
| WP\_004537261.1 | 55894 | 56362 | + | 155 | PF00582 | Usp | Universal stress protein family | 2.40E-31 |
| WP\_004529578.1 | 56511 | 57267 | + | 251 | PF13545 PF00027 PF00325 RREFamAux006 TIGR03896 | HTH\_Crp\_2 cNMP\_binding Crp RRE\_cNMP TIGR03896 | Crp-like helix-turn-helix domain Cyclic nucleotide-binding domain Bacterial regulatory proteins, crp family - cyc\_nuc\_ocin: bacteriocin-type transport-associated protein | 2.90E-20 6.00E-18 2.20E-14 2.60E-08 1.70E-05 |
| WP\_004525603.1 | 58227 | 57333 | - | 297 | TIGR03298 PF03466 PF00126 TIGR02424 TIGR03418 | TIGR03298 LysR\_substrate HTH\_1 TIGR02424 TIGR03418 | argP: transcriptional regulator, ArgP family LysR substrate binding domain Bacterial regulatory helix-turn-helix protein, lysR family TF\_pcaQ: pca operon transcription factor PcaQ chol\_sulf\_TF: putative choline sulfate-utilization transcription factor | 3.90E-120 6.00E-20 5.60E-15 4.10E-12 3.60E-11 |
| WP\_004194492.1 | 58334 | 58955 | + | 206 | TIGR00948 PF01810 TIGR00949 | TIGR00948 LysE TIGR00949 | 2a75: L-lysine exporter LysE type translocator 2A76: homoserine/Threonine efflux protein | 6.90E-53 1.40E-36 5.50E-06 |
| WP\_004529575.1 | 60116 | 59126 | - | 329 | TIGR04543 PF13649 PF13847 PF08242 TIGR03534 | TIGR04543 Methyltransf\_25 Methyltransf\_31 Methyltransf\_12 TIGR03534 | ketoArg\_3Met: 2-ketoarginine methyltransferase Methyltransferase domain Methyltransferase domain Methyltransferase domain RF\_mod\_PrmC: protein-(glutamine-N5) methyltransferase, release factor-specific | 1.10E-11 8.40E-09 2.80E-08 7.50E-07 1.50E-06 |
| WP\_004536689.1 | 62073 | 60129 | - | 647 | TIGR02026 TIGR03471 TIGR03975 TIGR04014 TIGR04013 | TIGR02026 TIGR03471 TIGR03975 TIGR04014 TIGR04013 | BchE: magnesium-protoporphyrin IX monomethyl ester anaerobic oxidative cyclase HpnJ: hopanoid biosynthesis associated radical SAM protein HpnJ rSAM\_ocin\_1: ribosomal peptide maturation radical SAM protein 1 B12\_SAM\_MJ\_0865: B12-binding domain/radical SAM domain protein, MJ\_0865 family B12\_SAM\_MJ\_1487: B12-binding domain/radical SAM domain protein, MJ\_1487 family | 4.60E-39 8.80E-32 3.30E-27 1.00E-25 3.00E-24 |
| WP\_004532432.1 | 62903 | 62069 | - | 277 | NO PFAM MATCH | - | - | - |
| WP\_004196528.1 | 63751 | 62899 | - | 283 | PF05114 PF01261 | DUF692 AP\_endonuc\_2 | Protein of unknown function (DUF692) Xylose isomerase-like TIM barrel | 1.90E-39 1.50E-06 |
| WP\_004201048.1 | 64125 | 63924 | - | 66 | NO PFAM MATCH | - | - | - |
| WP\_004194516.1 | 65681 | 64274 | - | 468 | PF00067 TIGR04538 TIGR04458 TIGR04515 | p450 TIGR04538 TIGR04458 TIGR04515 | Cytochrome P450 P450\_cycloAA\_1: cytochrome P450, cyclodipeptide synthase-associated CYP450\_TxtE: 4-nitrotryptophan synthase P450\_rel\_GT\_act: P450-derived glycosyltranferase activator | 5.60E-34 3.30E-16 4.80E-06 2.50E-05 |
| WP\_004529569.1 | 66083 | 66788 | + | 234 | PF03551 TIGR03433 | PadR TIGR03433 | Transcriptional regulator PadR-like family padR\_acidobact: transcriptional regulator, Acidobacterial, PadR-family | 2.00E-19 2.10E-12 |
| WP\_004194445.1 | 66827 | 67655 | + | 275 | PF04954 PF08021 | SIP FAD\_binding\_9 | Siderophore-interacting protein Siderophore-interacting FAD-binding domain | 2.90E-39 1.90E-37 |
| WP\_004194583.1 | 68221 | 67972 | - | 82 | NO PFAM MATCH | - | - | - |
| WP\_004196519.1 | 68414 | 68651 | + | 78 | NO PFAM MATCH | - | - | - |
| WP\_004194570.1 | 69270 | 68673 | - | 198 | PF01569 PF14378 | PAP2 PAP2\_3 | PAP2 superfamily PAP2 superfamily | 1.30E-17 4.00E-07 |
| WP\_004194452.1 | 69728 | 70469 | + | 246 | PF06314 | ADC | Acetoacetate decarboxylase (ADC) | 4.50E-79 |

#### Results for WP\_038756459.1 [Burkholderia pseudomallei] back to top

#### previous - next

##### Architecture

1000 nucleotides

Link to nucleotide sequence  
  

| Accession | start | end | direction | length (aa) | Pfam/HMM | name | description | E-value |
| --- | --- | --- | --- | --- | --- | --- | --- | --- |
| WP\_004194465.1 | 2279644 | 2279413 | - | 76 | NO PFAM MATCH | - | - | - |
| WP\_004194548.1 | 2280067 | 2280535 | + | 155 | PF00582 | Usp | Universal stress protein family | 2.40E-31 |
| WP\_004529578.1 | 2280684 | 2281440 | + | 251 | PF13545 PF00027 PF00325 RREFamAux006 TIGR03896 | HTH\_Crp\_2 cNMP\_binding Crp RRE\_cNMP TIGR03896 | Crp-like helix-turn-helix domain Cyclic nucleotide-binding domain Bacterial regulatory proteins, crp family - cyc\_nuc\_ocin: bacteriocin-type transport-associated protein | 2.90E-20 6.00E-18 2.20E-14 2.60E-08 1.70E-05 |
| WP\_004525603.1 | 2282391 | 2281497 | - | 297 | TIGR03298 PF03466 PF00126 TIGR02424 TIGR03418 | TIGR03298 LysR\_substrate HTH\_1 TIGR02424 TIGR03418 | argP: transcriptional regulator, ArgP family LysR substrate binding domain Bacterial regulatory helix-turn-helix protein, lysR family TF\_pcaQ: pca operon transcription factor PcaQ chol\_sulf\_TF: putative choline sulfate-utilization transcription factor | 3.90E-120 6.00E-20 5.60E-15 4.10E-12 3.60E-11 |
| WP\_004194492.1 | 2282498 | 2283119 | + | 206 | TIGR00948 PF01810 TIGR00949 | TIGR00948 LysE TIGR00949 | 2a75: L-lysine exporter LysE type translocator 2A76: homoserine/Threonine efflux protein | 6.90E-53 1.40E-36 5.50E-06 |
| WP\_004545057.1 | 2284280 | 2283290 | - | 329 | TIGR04543 PF13649 PF13847 PF08242 TIGR03534 | TIGR04543 Methyltransf\_25 Methyltransf\_31 Methyltransf\_12 TIGR03534 | ketoArg\_3Met: 2-ketoarginine methyltransferase Methyltransferase domain Methyltransferase domain Methyltransferase domain RF\_mod\_PrmC: protein-(glutamine-N5) methyltransferase, release factor-specific | 1.10E-11 8.40E-09 2.80E-08 7.50E-07 1.40E-06 |
| WP\_004544988.1 | 2286237 | 2284293 | - | 647 | TIGR02026 TIGR03471 TIGR03975 TIGR04014 TIGR04013 | TIGR02026 TIGR03471 TIGR03975 TIGR04014 TIGR04013 | BchE: magnesium-protoporphyrin IX monomethyl ester anaerobic oxidative cyclase HpnJ: hopanoid biosynthesis associated radical SAM protein HpnJ rSAM\_ocin\_1: ribosomal peptide maturation radical SAM protein 1 B12\_SAM\_MJ\_0865: B12-binding domain/radical SAM domain protein, MJ\_0865 family B12\_SAM\_MJ\_1487: B12-binding domain/radical SAM domain protein, MJ\_1487 family | 3.70E-39 8.80E-32 3.30E-27 1.00E-25 3.00E-24 |
| WP\_009932366.1 | 2287055 | 2286233 | - | 273 | NO PFAM MATCH | - | - | - |
| WP\_038756459.1 | 2287903 | 2287051 | - | 283 | PF05114 PF01261 | DUF692 AP\_endonuc\_2 | Protein of unknown function (DUF692) Xylose isomerase-like TIM barrel | 3.60E-39 7.40E-06 |
| WP\_004201048.1 | 2288277 | 2288076 | - | 66 | NO PFAM MATCH | - | - | - |
| WP\_038756461.1 | 2289833 | 2288426 | - | 468 | PF00067 TIGR04538 TIGR04458 TIGR04515 | p450 TIGR04538 TIGR04458 TIGR04515 | Cytochrome P450 P450\_cycloAA\_1: cytochrome P450, cyclodipeptide synthase-associated CYP450\_TxtE: 4-nitrotryptophan synthase P450\_rel\_GT\_act: P450-derived glycosyltranferase activator | 5.60E-34 3.30E-16 4.80E-06 2.50E-05 |
| WP\_004196523.1 | 2290240 | 2290948 | + | 235 | PF03551 TIGR03433 | PadR TIGR03433 | Transcriptional regulator PadR-like family padR\_acidobact: transcriptional regulator, Acidobacterial, PadR-family | 2.00E-19 2.20E-12 |
| WP\_004194445.1 | 2290987 | 2291815 | + | 275 | PF04954 PF08021 | SIP FAD\_binding\_9 | Siderophore-interacting protein Siderophore-interacting FAD-binding domain | 2.90E-39 1.90E-37 |
| WP\_004196519.1 | 2292580 | 2292817 | + | 78 | NO PFAM MATCH | - | - | - |
| WP\_038731007.1 | 2293436 | 2292839 | - | 198 | PF01569 PF14378 | PAP2 PAP2\_3 | PAP2 superfamily PAP2 superfamily | 1.30E-17 4.00E-07 |
| WP\_004194452.1 | 2293894 | 2294635 | + | 246 | PF06314 | ADC | Acetoacetate decarboxylase (ADC) | 4.50E-79 |
| WP\_004194455.1 | 2294681 | 2295467 | + | 261 | TIGR01963 PF00106 PF13561 TIGR03971 TIGR02415 | TIGR01963 adh\_short adh\_short\_C2 TIGR03971 TIGR02415 | PHB\_DH: 3-hydroxybutyrate dehydrogenase short chain dehydrogenase Enoyl-(Acyl carrier protein) reductase SDR\_subfam\_1: SDR family mycofactocin-dependent oxidoreductase 23BDH: acetoin reductases | 4.30E-104 1.80E-52 2.40E-52 6.40E-52 8.90E-52 |

#### Results for WP\_058035158.1 [Burkholderia pseudomallei] back to top

#### previous - next

##### Architecture

1000 nucleotides

Link to nucleotide sequence  
  

| Accession | start | end | direction | length (aa) | Pfam/HMM | name | description | E-value |
| --- | --- | --- | --- | --- | --- | --- | --- | --- |
| WP\_004194452.1 | 811 | 70 | - | 246 | PF06314 | ADC | Acetoacetate decarboxylase (ADC) | 4.50E-79 |
| WP\_004525615.1 | 1269 | 1866 | + | 198 | PF01569 PF14378 | PAP2 PAP2\_3 | PAP2 superfamily PAP2 superfamily | 7.40E-18 7.90E-07 |
| WP\_004196519.1 | 2125 | 1888 | - | 78 | NO PFAM MATCH | - | - | - |
| WP\_004194445.1 | 3718 | 2890 | - | 275 | PF04954 PF08021 | SIP FAD\_binding\_9 | Siderophore-interacting protein Siderophore-interacting FAD-binding domain | 2.90E-39 1.90E-37 |
| WP\_038749976.1 | 4474 | 3757 | - | 238 | PF03551 TIGR03433 | PadR TIGR03433 | Transcriptional regulator PadR-like family padR\_acidobact: transcriptional regulator, Acidobacterial, PadR-family | 2.00E-19 2.20E-12 |
| WP\_004549191.1 | 4876 | 6283 | + | 468 | PF00067 TIGR04538 TIGR04515 TIGR04458 | p450 TIGR04538 TIGR04515 TIGR04458 | Cytochrome P450 P450\_cycloAA\_1: cytochrome P450, cyclodipeptide synthase-associated P450\_rel\_GT\_act: P450-derived glycosyltranferase activator CYP450\_TxtE: 4-nitrotryptophan synthase | 1.70E-34 1.90E-16 5.30E-06 6.70E-06 |
| WP\_004201048.1 | 6432 | 6633 | + | 66 | NO PFAM MATCH | - | - | - |
| WP\_058035158.1 | 6806 | 7658 | + | 283 | PF05114 PF01261 | DUF692 AP\_endonuc\_2 | Protein of unknown function (DUF692) Xylose isomerase-like TIM barrel | 1.70E-39 1.30E-06 |
| WP\_038726508.1 | 7654 | 8488 | + | 277 | NO PFAM MATCH | - | - | - |
| WP\_208808628.1 | 8484 | 10428 | + | 647 | TIGR02026 TIGR03471 TIGR03975 TIGR04014 TIGR04013 | TIGR02026 TIGR03471 TIGR03975 TIGR04014 TIGR04013 | BchE: magnesium-protoporphyrin IX monomethyl ester anaerobic oxidative cyclase HpnJ: hopanoid biosynthesis associated radical SAM protein HpnJ rSAM\_ocin\_1: ribosomal peptide maturation radical SAM protein 1 B12\_SAM\_MJ\_0865: B12-binding domain/radical SAM domain protein, MJ\_0865 family B12\_SAM\_MJ\_1487: B12-binding domain/radical SAM domain protein, MJ\_1487 family | 6.60E-39 1.50E-31 1.40E-26 3.00E-25 1.00E-23 |
| WP\_038734007.1 | 10441 | 11431 | + | 329 | TIGR04543 PF13649 PF13847 PF08242 TIGR03534 | TIGR04543 Methyltransf\_25 Methyltransf\_31 Methyltransf\_12 TIGR03534 | ketoArg\_3Met: 2-ketoarginine methyltransferase Methyltransferase domain Methyltransferase domain Methyltransferase domain RF\_mod\_PrmC: protein-(glutamine-N5) methyltransferase, release factor-specific | 7.00E-12 8.00E-09 2.80E-08 7.00E-07 1.30E-06 |
| WP\_004546141.1 | 12223 | 11602 | - | 206 | TIGR00948 PF01810 TIGR00949 | TIGR00948 LysE TIGR00949 | 2a75: L-lysine exporter LysE type translocator 2A76: homoserine/Threonine efflux protein | 4.90E-53 1.00E-36 5.30E-06 |
| WP\_004525603.1 | 12330 | 13224 | + | 297 | TIGR03298 PF03466 PF00126 TIGR02424 TIGR03418 | TIGR03298 LysR\_substrate HTH\_1 TIGR02424 TIGR03418 | argP: transcriptional regulator, ArgP family LysR substrate binding domain Bacterial regulatory helix-turn-helix protein, lysR family TF\_pcaQ: pca operon transcription factor PcaQ chol\_sulf\_TF: putative choline sulfate-utilization transcription factor | 3.90E-120 6.00E-20 5.60E-15 4.10E-12 3.60E-11 |
| WP\_004529578.1 | 14036 | 13280 | - | 251 | PF13545 PF00027 PF00325 RREFamAux006 TIGR03896 | HTH\_Crp\_2 cNMP\_binding Crp RRE\_cNMP TIGR03896 | Crp-like helix-turn-helix domain Cyclic nucleotide-binding domain Bacterial regulatory proteins, crp family - cyc\_nuc\_ocin: bacteriocin-type transport-associated protein | 2.90E-20 6.00E-18 2.20E-14 2.60E-08 1.70E-05 |
| WP\_004194548.1 | 14653 | 14185 | - | 155 | PF00582 | Usp | Universal stress protein family | 2.40E-31 |
| WP\_004194465.1 | 15076 | 15307 | + | 76 | NO PFAM MATCH | - | - | - |

#### Results for WP\_038754083.1 [Burkholderia pseudomallei] back to top

#### previous - next

##### Architecture

1000 nucleotides

Link to nucleotide sequence  
  

| Accession | start | end | direction | length (aa) | Pfam/HMM | name | description | E-value |
| --- | --- | --- | --- | --- | --- | --- | --- | --- |
| WP\_004194455.1 | 1341246 | 1340460 | - | 261 | TIGR01963 PF00106 PF13561 TIGR03971 TIGR02415 | TIGR01963 adh\_short adh\_short\_C2 TIGR03971 TIGR02415 | PHB\_DH: 3-hydroxybutyrate dehydrogenase short chain dehydrogenase Enoyl-(Acyl carrier protein) reductase SDR\_subfam\_1: SDR family mycofactocin-dependent oxidoreductase 23BDH: acetoin reductases | 4.30E-104 1.80E-52 2.40E-52 6.40E-52 8.90E-52 |
| WP\_004194452.1 | 1342033 | 1341292 | - | 246 | PF06314 | ADC | Acetoacetate decarboxylase (ADC) | 4.50E-79 |
| WP\_004194570.1 | 1342490 | 1343087 | + | 198 | PF01569 PF14378 | PAP2 PAP2\_3 | PAP2 superfamily PAP2 superfamily | 1.30E-17 4.00E-07 |
| WP\_004196519.1 | 1343346 | 1343109 | - | 78 | NO PFAM MATCH | - | - | - |
| WP\_038726504.1 | 1344934 | 1344106 | - | 275 | PF04954 PF08021 | SIP FAD\_binding\_9 | Siderophore-interacting protein Siderophore-interacting FAD-binding domain | 3.70E-39 1.90E-37 |
| WP\_004529569.1 | 1345678 | 1344973 | - | 234 | PF03551 TIGR03433 | PadR TIGR03433 | Transcriptional regulator PadR-like family padR\_acidobact: transcriptional regulator, Acidobacterial, PadR-family | 2.00E-19 2.10E-12 |
| WP\_038754080.1 | 1346079 | 1347486 | + | 468 | PF00067 TIGR04538 TIGR04515 TIGR04458 | p450 TIGR04538 TIGR04515 TIGR04458 | Cytochrome P450 P450\_cycloAA\_1: cytochrome P450, cyclodipeptide synthase-associated P450\_rel\_GT\_act: P450-derived glycosyltranferase activator CYP450\_TxtE: 4-nitrotryptophan synthase | 1.70E-34 1.90E-16 4.70E-06 6.80E-06 |
| WP\_004201048.1 | 1347635 | 1347836 | + | 66 | NO PFAM MATCH | - | - | - |
| WP\_038754083.1 | 1348009 | 1348861 | + | 283 | PF05114 PF01261 | DUF692 AP\_endonuc\_2 | Protein of unknown function (DUF692) Xylose isomerase-like TIM barrel | 1.60E-39 1.50E-06 |
| WP\_038754086.1 | 1348857 | 1349691 | + | 277 | NO PFAM MATCH | - | - | - |
| WP\_004544988.1 | 1349687 | 1351631 | + | 647 | TIGR02026 TIGR03471 TIGR03975 TIGR04014 TIGR04013 | TIGR02026 TIGR03471 TIGR03975 TIGR04014 TIGR04013 | BchE: magnesium-protoporphyrin IX monomethyl ester anaerobic oxidative cyclase HpnJ: hopanoid biosynthesis associated radical SAM protein HpnJ rSAM\_ocin\_1: ribosomal peptide maturation radical SAM protein 1 B12\_SAM\_MJ\_0865: B12-binding domain/radical SAM domain protein, MJ\_0865 family B12\_SAM\_MJ\_1487: B12-binding domain/radical SAM domain protein, MJ\_1487 family | 3.70E-39 8.80E-32 3.30E-27 1.00E-25 3.00E-24 |
| WP\_004532676.1 | 1351644 | 1352634 | + | 329 | TIGR04543 PF13649 PF13847 PF08242 TIGR03534 | TIGR04543 Methyltransf\_25 Methyltransf\_31 Methyltransf\_12 TIGR03534 | ketoArg\_3Met: 2-ketoarginine methyltransferase Methyltransferase domain Methyltransferase domain Methyltransferase domain RF\_mod\_PrmC: protein-(glutamine-N5) methyltransferase, release factor-specific | 7.10E-12 8.20E-09 2.80E-08 7.60E-07 1.40E-06 |
| WP\_004546141.1 | 1353426 | 1352805 | - | 206 | TIGR00948 PF01810 TIGR00949 | TIGR00948 LysE TIGR00949 | 2a75: L-lysine exporter LysE type translocator 2A76: homoserine/Threonine efflux protein | 4.90E-53 1.00E-36 5.30E-06 |
| WP\_004525603.1 | 1353533 | 1354427 | + | 297 | TIGR03298 PF03466 PF00126 TIGR02424 TIGR03418 | TIGR03298 LysR\_substrate HTH\_1 TIGR02424 TIGR03418 | argP: transcriptional regulator, ArgP family LysR substrate binding domain Bacterial regulatory helix-turn-helix protein, lysR family TF\_pcaQ: pca operon transcription factor PcaQ chol\_sulf\_TF: putative choline sulfate-utilization transcription factor | 3.90E-120 6.00E-20 5.60E-15 4.10E-12 3.60E-11 |
| WP\_004529578.1 | 1355239 | 1354483 | - | 251 | PF13545 PF00027 PF00325 RREFamAux006 TIGR03896 | HTH\_Crp\_2 cNMP\_binding Crp RRE\_cNMP TIGR03896 | Crp-like helix-turn-helix domain Cyclic nucleotide-binding domain Bacterial regulatory proteins, crp family - cyc\_nuc\_ocin: bacteriocin-type transport-associated protein | 2.90E-20 6.00E-18 2.20E-14 2.60E-08 1.70E-05 |
| WP\_004194548.1 | 1355856 | 1355388 | - | 155 | PF00582 | Usp | Universal stress protein family | 2.40E-31 |
| WP\_004194465.1 | 1356279 | 1356510 | + | 76 | NO PFAM MATCH | - | - | - |

#### Results for WP\_122654464.1 [Burkholderia pseudomallei] back to top

#### previous - next

##### Architecture

1000 nucleotides

Link to nucleotide sequence  
  

| Accession | start | end | direction | length (aa) | Pfam/HMM | name | description | E-value |
| --- | --- | --- | --- | --- | --- | --- | --- | --- |
| WP\_004194452.1 | 1088 | 347 | - | 246 | PF06314 | ADC | Acetoacetate decarboxylase (ADC) | 4.50E-79 |
| WP\_004525615.1 | 1546 | 2143 | + | 198 | PF01569 PF14378 | PAP2 PAP2\_3 | PAP2 superfamily PAP2 superfamily | 7.40E-18 7.90E-07 |
| WP\_004529566.1 | 2402 | 2165 | - | 78 | NO PFAM MATCH | - | - | - |
| WP\_004194583.1 | 2595 | 2844 | + | 82 | NO PFAM MATCH | - | - | - |
| WP\_004532447.1 | 3995 | 3167 | - | 275 | PF04954 PF08021 | SIP FAD\_binding\_9 | Siderophore-interacting protein Siderophore-interacting FAD-binding domain | 2.50E-39 1.80E-37 |
| WP\_004529569.1 | 4739 | 4034 | - | 234 | PF03551 TIGR03433 | PadR TIGR03433 | Transcriptional regulator PadR-like family padR\_acidobact: transcriptional regulator, Acidobacterial, PadR-family | 2.00E-19 2.10E-12 |
| WP\_004529570.1 | 5141 | 6548 | + | 468 | PF00067 TIGR04538 TIGR04458 TIGR04515 | p450 TIGR04538 TIGR04458 TIGR04515 | Cytochrome P450 P450\_cycloAA\_1: cytochrome P450, cyclodipeptide synthase-associated CYP450\_TxtE: 4-nitrotryptophan synthase P450\_rel\_GT\_act: P450-derived glycosyltranferase activator | 5.20E-33 1.40E-15 3.50E-05 6.30E-05 |
| WP\_004201048.1 | 6697 | 6898 | + | 66 | NO PFAM MATCH | - | - | - |
| WP\_122654464.1 | 7071 | 7923 | + | 283 | PF05114 PF01261 | DUF692 AP\_endonuc\_2 | Protein of unknown function (DUF692) Xylose isomerase-like TIM barrel | 3.70E-39 1.30E-06 |
| WP\_004532432.1 | 7919 | 8753 | + | 277 | NO PFAM MATCH | - | - | - |
| WP\_004532412.1 | 8749 | 10693 | + | 647 | TIGR02026 TIGR03471 TIGR03975 TIGR04014 TIGR04013 | TIGR02026 TIGR03471 TIGR03975 TIGR04014 TIGR04013 | BchE: magnesium-protoporphyrin IX monomethyl ester anaerobic oxidative cyclase HpnJ: hopanoid biosynthesis associated radical SAM protein HpnJ rSAM\_ocin\_1: ribosomal peptide maturation radical SAM protein 1 B12\_SAM\_MJ\_0865: B12-binding domain/radical SAM domain protein, MJ\_0865 family B12\_SAM\_MJ\_1487: B12-binding domain/radical SAM domain protein, MJ\_1487 family | 6.30E-40 5.20E-32 7.30E-28 1.60E-26 1.70E-24 |
| WP\_004532434.1 | 10706 | 11696 | + | 329 | TIGR04543 PF13649 PF13847 PF08242 TIGR03534 | TIGR04543 Methyltransf\_25 Methyltransf\_31 Methyltransf\_12 TIGR03534 | ketoArg\_3Met: 2-ketoarginine methyltransferase Methyltransferase domain Methyltransferase domain Methyltransferase domain RF\_mod\_PrmC: protein-(glutamine-N5) methyltransferase, release factor-specific | 7.20E-12 8.20E-09 2.80E-08 7.60E-07 1.50E-06 |
| WP\_004194492.1 | 12488 | 11867 | - | 206 | TIGR00948 PF01810 TIGR00949 | TIGR00948 LysE TIGR00949 | 2a75: L-lysine exporter LysE type translocator 2A76: homoserine/Threonine efflux protein | 6.90E-53 1.40E-36 5.50E-06 |
| WP\_004525603.1 | 12595 | 13489 | + | 297 | TIGR03298 PF03466 PF00126 TIGR02424 TIGR03418 | TIGR03298 LysR\_substrate HTH\_1 TIGR02424 TIGR03418 | argP: transcriptional regulator, ArgP family LysR substrate binding domain Bacterial regulatory helix-turn-helix protein, lysR family TF\_pcaQ: pca operon transcription factor PcaQ chol\_sulf\_TF: putative choline sulfate-utilization transcription factor | 3.90E-120 6.00E-20 5.60E-15 4.10E-12 3.60E-11 |
| WP\_004529578.1 | 14329 | 13573 | - | 251 | PF13545 PF00027 PF00325 RREFamAux006 TIGR03896 | HTH\_Crp\_2 cNMP\_binding Crp RRE\_cNMP TIGR03896 | Crp-like helix-turn-helix domain Cyclic nucleotide-binding domain Bacterial regulatory proteins, crp family - cyc\_nuc\_ocin: bacteriocin-type transport-associated protein | 2.90E-20 6.00E-18 2.20E-14 2.60E-08 1.70E-05 |
| WP\_004194548.1 | 14946 | 14478 | - | 155 | PF00582 | Usp | Universal stress protein family | 2.40E-31 |
| WP\_004194465.1 | 15365 | 15596 | + | 76 | NO PFAM MATCH | - | - | - |

#### Results for WP\_208813011.1 [Burkholderia pseudomallei] back to top

#### previous - next

##### Architecture

1000 nucleotides

Link to nucleotide sequence  
  

| Accession | start | end | direction | length (aa) | Pfam/HMM | name | description | E-value |
| --- | --- | --- | --- | --- | --- | --- | --- | --- |
| WP\_004194465.1 | 58299 | 58068 | - | 76 | NO PFAM MATCH | - | - | - |
| WP\_004194548.1 | 58722 | 59190 | + | 155 | PF00582 | Usp | Universal stress protein family | 2.40E-31 |
| WP\_004529578.1 | 59339 | 60095 | + | 251 | PF13545 PF00027 PF00325 RREFamAux006 TIGR03896 | HTH\_Crp\_2 cNMP\_binding Crp RRE\_cNMP TIGR03896 | Crp-like helix-turn-helix domain Cyclic nucleotide-binding domain Bacterial regulatory proteins, crp family - cyc\_nuc\_ocin: bacteriocin-type transport-associated protein | 2.90E-20 6.00E-18 2.20E-14 2.60E-08 1.70E-05 |
| WP\_004525603.1 | 61035 | 60141 | - | 297 | TIGR03298 PF03466 PF00126 TIGR02424 TIGR03418 | TIGR03298 LysR\_substrate HTH\_1 TIGR02424 TIGR03418 | argP: transcriptional regulator, ArgP family LysR substrate binding domain Bacterial regulatory helix-turn-helix protein, lysR family TF\_pcaQ: pca operon transcription factor PcaQ chol\_sulf\_TF: putative choline sulfate-utilization transcription factor | 3.90E-120 6.00E-20 5.60E-15 4.10E-12 3.60E-11 |
| WP\_004194492.1 | 61142 | 61763 | + | 206 | TIGR00948 PF01810 TIGR00949 | TIGR00948 LysE TIGR00949 | 2a75: L-lysine exporter LysE type translocator 2A76: homoserine/Threonine efflux protein | 6.90E-53 1.40E-36 5.50E-06 |
| WP\_004532676.1 | 62924 | 61934 | - | 329 | TIGR04543 PF13649 PF13847 PF08242 TIGR03534 | TIGR04543 Methyltransf\_25 Methyltransf\_31 Methyltransf\_12 TIGR03534 | ketoArg\_3Met: 2-ketoarginine methyltransferase Methyltransferase domain Methyltransferase domain Methyltransferase domain RF\_mod\_PrmC: protein-(glutamine-N5) methyltransferase, release factor-specific | 7.10E-12 8.20E-09 2.80E-08 7.60E-07 1.40E-06 |
| WP\_004544988.1 | 64881 | 62937 | - | 647 | TIGR02026 TIGR03471 TIGR03975 TIGR04014 TIGR04013 | TIGR02026 TIGR03471 TIGR03975 TIGR04014 TIGR04013 | BchE: magnesium-protoporphyrin IX monomethyl ester anaerobic oxidative cyclase HpnJ: hopanoid biosynthesis associated radical SAM protein HpnJ rSAM\_ocin\_1: ribosomal peptide maturation radical SAM protein 1 B12\_SAM\_MJ\_0865: B12-binding domain/radical SAM domain protein, MJ\_0865 family B12\_SAM\_MJ\_1487: B12-binding domain/radical SAM domain protein, MJ\_1487 family | 3.70E-39 8.80E-32 3.30E-27 1.00E-25 3.00E-24 |
| WP\_004194512.1 | 65711 | 64877 | - | 277 | NO PFAM MATCH | - | - | - |
| WP\_208813011.1 | 66559 | 65707 | - | 283 | PF05114 PF01261 | DUF692 AP\_endonuc\_2 | Protein of unknown function (DUF692) Xylose isomerase-like TIM barrel | 1.90E-39 1.50E-06 |
| WP\_004201048.1 | 66933 | 66732 | - | 66 | NO PFAM MATCH | - | - | - |
| WP\_004194516.1 | 68489 | 67082 | - | 468 | PF00067 TIGR04538 TIGR04458 TIGR04515 | p450 TIGR04538 TIGR04458 TIGR04515 | Cytochrome P450 P450\_cycloAA\_1: cytochrome P450, cyclodipeptide synthase-associated CYP450\_TxtE: 4-nitrotryptophan synthase P450\_rel\_GT\_act: P450-derived glycosyltranferase activator | 5.60E-34 3.30E-16 4.80E-06 2.50E-05 |
| WP\_004196523.1 | 68891 | 69599 | + | 235 | PF03551 TIGR03433 | PadR TIGR03433 | Transcriptional regulator PadR-like family padR\_acidobact: transcriptional regulator, Acidobacterial, PadR-family | 2.00E-19 2.20E-12 |
| WP\_004194445.1 | 69638 | 70466 | + | 275 | PF04954 PF08021 | SIP FAD\_binding\_9 | Siderophore-interacting protein Siderophore-interacting FAD-binding domain | 2.90E-39 1.90E-37 |
| WP\_004196519.1 | 71231 | 71468 | + | 78 | NO PFAM MATCH | - | - | - |
| WP\_004194570.1 | 72087 | 71490 | - | 198 | PF01569 PF14378 | PAP2 PAP2\_3 | PAP2 superfamily PAP2 superfamily | 1.30E-17 4.00E-07 |
| WP\_004194452.1 | 72545 | 73286 | + | 246 | PF06314 | ADC | Acetoacetate decarboxylase (ADC) | 4.50E-79 |

#### Results for WP\_009981063.1 [Burkholderia pseudomallei BCC215] back to top

#### previous - next

##### Architecture

1000 nucleotides

Link to nucleotide sequence  
  

| Accession | start | end | direction | length (aa) | Pfam/HMM | name | description | E-value |
| --- | --- | --- | --- | --- | --- | --- | --- | --- |
| WP\_004194452.1 | 812 | 71 | - | 246 | PF06314 | ADC | Acetoacetate decarboxylase (ADC) | 4.50E-79 |
| WP\_004525615.1 | 1270 | 1867 | + | 198 | PF01569 PF14378 | PAP2 PAP2\_3 | PAP2 superfamily PAP2 superfamily | 7.40E-18 7.90E-07 |
| WP\_004196519.1 | 2126 | 1889 | - | 78 | NO PFAM MATCH | - | - | - |
| WP\_004194445.1 | 3719 | 2891 | - | 275 | PF04954 PF08021 | SIP FAD\_binding\_9 | Siderophore-interacting protein Siderophore-interacting FAD-binding domain | 2.90E-39 1.90E-37 |
| WP\_004529569.1 | 4463 | 3758 | - | 234 | PF03551 TIGR03433 | PadR TIGR03433 | Transcriptional regulator PadR-like family padR\_acidobact: transcriptional regulator, Acidobacterial, PadR-family | 2.00E-19 2.10E-12 |
| WP\_009981062.1 | 4864 | 6271 | + | 468 | PF00067 TIGR04538 TIGR04458 TIGR04515 | p450 TIGR04538 TIGR04458 TIGR04515 | Cytochrome P450 P450\_cycloAA\_1: cytochrome P450, cyclodipeptide synthase-associated CYP450\_TxtE: 4-nitrotryptophan synthase P450\_rel\_GT\_act: P450-derived glycosyltranferase activator | 5.20E-33 2.90E-15 3.50E-05 5.80E-05 |
| WP\_004201048.1 | 6420 | 6621 | + | 66 | NO PFAM MATCH | - | - | - |
| WP\_009981063.1 | 6794 | 7646 | + | 283 | PF05114 PF01261 | DUF692 AP\_endonuc\_2 | Protein of unknown function (DUF692) Xylose isomerase-like TIM barrel | 3.00E-39 9.70E-07 |
| WP\_009967374.1 | 7642 | 8476 | + | 277 | NO PFAM MATCH | - | - | - |
| WP\_004546215.1 | 8472 | 10416 | + | 647 | TIGR02026 TIGR03471 TIGR03975 TIGR04014 TIGR04013 | TIGR02026 TIGR03471 TIGR03975 TIGR04014 TIGR04013 | BchE: magnesium-protoporphyrin IX monomethyl ester anaerobic oxidative cyclase HpnJ: hopanoid biosynthesis associated radical SAM protein HpnJ rSAM\_ocin\_1: ribosomal peptide maturation radical SAM protein 1 B12\_SAM\_MJ\_0865: B12-binding domain/radical SAM domain protein, MJ\_0865 family B12\_SAM\_MJ\_1487: B12-binding domain/radical SAM domain protein, MJ\_1487 family | 3.70E-39 8.80E-32 3.30E-27 1.00E-25 3.00E-24 |
| WP\_009967375.1 | 10429 | 11419 | + | 329 | TIGR04543 PF13649 PF13847 PF08242 TIGR03534 | TIGR04543 Methyltransf\_25 Methyltransf\_31 Methyltransf\_12 TIGR03534 | ketoArg\_3Met: 2-ketoarginine methyltransferase Methyltransferase domain Methyltransferase domain Methyltransferase domain RF\_mod\_PrmC: protein-(glutamine-N5) methyltransferase, release factor-specific | 4.80E-12 7.90E-09 2.80E-08 7.00E-07 1.40E-06 |
| WP\_004194492.1 | 12211 | 11590 | - | 206 | TIGR00948 PF01810 TIGR00949 | TIGR00948 LysE TIGR00949 | 2a75: L-lysine exporter LysE type translocator 2A76: homoserine/Threonine efflux protein | 6.90E-53 1.40E-36 5.50E-06 |
| WP\_004525603.1 | 12318 | 13212 | + | 297 | TIGR03298 PF03466 PF00126 TIGR02424 TIGR03418 | TIGR03298 LysR\_substrate HTH\_1 TIGR02424 TIGR03418 | argP: transcriptional regulator, ArgP family LysR substrate binding domain Bacterial regulatory helix-turn-helix protein, lysR family TF\_pcaQ: pca operon transcription factor PcaQ chol\_sulf\_TF: putative choline sulfate-utilization transcription factor | 3.90E-120 6.00E-20 5.60E-15 4.10E-12 3.60E-11 |
| WP\_004529578.1 | 14034 | 13278 | - | 251 | PF13545 PF00027 PF00325 RREFamAux006 TIGR03896 | HTH\_Crp\_2 cNMP\_binding Crp RRE\_cNMP TIGR03896 | Crp-like helix-turn-helix domain Cyclic nucleotide-binding domain Bacterial regulatory proteins, crp family - cyc\_nuc\_ocin: bacteriocin-type transport-associated protein | 2.90E-20 6.00E-18 2.20E-14 2.60E-08 1.70E-05 |
| WP\_004194548.1 | 14651 | 14183 | - | 155 | PF00582 | Usp | Universal stress protein family | 2.40E-31 |
| WP\_004194465.1 | 15074 | 15305 | + | 76 | NO PFAM MATCH | - | - | - |

#### Results for WP\_264236925.1 [Burkholderia pseudomallei] back to top

#### previous - next

##### Architecture

1000 nucleotides

Link to nucleotide sequence  
  

| Accession | start | end | direction | length (aa) | Pfam/HMM | name | description | E-value |
| --- | --- | --- | --- | --- | --- | --- | --- | --- |
| WP\_038797612.1 | 151 | 0 | - | 50 | TIGR01963 TIGR02632 TIGR02415 PF00106 TIGR03206 | TIGR01963 TIGR02632 TIGR02415 adh\_short TIGR03206 | PHB\_DH: 3-hydroxybutyrate dehydrogenase RhaD\_aldol-ADH: rhamnulose-1-phosphate aldolase/alcohol dehydrogenase 23BDH: acetoin reductases short chain dehydrogenase benzo\_BadH: 2-hydroxycyclohexanecarboxyl-CoA dehydrogenase | 5.00E-12 3.60E-11 1.30E-10 6.90E-10 9.50E-10 |
| WP\_004194452.1 | 938 | 197 | - | 246 | PF06314 | ADC | Acetoacetate decarboxylase (ADC) | 4.50E-79 |
| WP\_004525615.1 | 1396 | 1993 | + | 198 | PF01569 PF14378 | PAP2 PAP2\_3 | PAP2 superfamily PAP2 superfamily | 7.40E-18 7.90E-07 |
| WP\_004196519.1 | 2252 | 2015 | - | 78 | NO PFAM MATCH | - | - | - |
| WP\_004194583.1 | 2445 | 2694 | + | 82 | NO PFAM MATCH | - | - | - |
| WP\_004194445.1 | 3845 | 3017 | - | 275 | PF04954 PF08021 | SIP FAD\_binding\_9 | Siderophore-interacting protein Siderophore-interacting FAD-binding domain | 2.90E-39 1.90E-37 |
| WP\_004553039.1 | 4592 | 3884 | - | 235 | PF03551 TIGR03433 | PadR TIGR03433 | Transcriptional regulator PadR-like family padR\_acidobact: transcriptional regulator, Acidobacterial, PadR-family | 1.40E-19 2.00E-12 |
| WP\_004529570.1 | 4999 | 6406 | + | 468 | PF00067 TIGR04538 TIGR04458 TIGR04515 | p450 TIGR04538 TIGR04458 TIGR04515 | Cytochrome P450 P450\_cycloAA\_1: cytochrome P450, cyclodipeptide synthase-associated CYP450\_TxtE: 4-nitrotryptophan synthase P450\_rel\_GT\_act: P450-derived glycosyltranferase activator | 5.20E-33 1.40E-15 3.50E-05 6.30E-05 |
| WP\_004201048.1 | 6555 | 6756 | + | 66 | NO PFAM MATCH | - | - | - |
| WP\_264236925.1 | 6929 | 7781 | + | 283 | PF05114 PF01261 | DUF692 AP\_endonuc\_2 | Protein of unknown function (DUF692) Xylose isomerase-like TIM barrel | 2.60E-39 2.00E-06 |
| WP\_004532432.1 | 7777 | 8611 | + | 277 | NO PFAM MATCH | - | - | - |
| WP\_179104781.1 | 8607 | 10551 | + | 647 | TIGR02026 TIGR03471 TIGR03975 TIGR04014 TIGR04013 | TIGR02026 TIGR03471 TIGR03975 TIGR04014 TIGR04013 | BchE: magnesium-protoporphyrin IX monomethyl ester anaerobic oxidative cyclase HpnJ: hopanoid biosynthesis associated radical SAM protein HpnJ rSAM\_ocin\_1: ribosomal peptide maturation radical SAM protein 1 B12\_SAM\_MJ\_0865: B12-binding domain/radical SAM domain protein, MJ\_0865 family B12\_SAM\_MJ\_1487: B12-binding domain/radical SAM domain protein, MJ\_1487 family | 5.60E-40 5.10E-32 1.80E-28 2.90E-26 9.60E-25 |
| WP\_004532676.1 | 10564 | 11554 | + | 329 | TIGR04543 PF13649 PF13847 PF08242 TIGR03534 | TIGR04543 Methyltransf\_25 Methyltransf\_31 Methyltransf\_12 TIGR03534 | ketoArg\_3Met: 2-ketoarginine methyltransferase Methyltransferase domain Methyltransferase domain Methyltransferase domain RF\_mod\_PrmC: protein-(glutamine-N5) methyltransferase, release factor-specific | 7.10E-12 8.20E-09 2.80E-08 7.60E-07 1.40E-06 |
| WP\_264236923.1 | 12329 | 11708 | - | 206 | TIGR00948 PF01810 TIGR00949 | TIGR00948 LysE TIGR00949 | 2a75: L-lysine exporter LysE type translocator 2A76: homoserine/Threonine efflux protein | 7.80E-53 2.10E-36 4.40E-06 |
| WP\_004532664.1 | 12436 | 13330 | + | 297 | TIGR03298 PF03466 PF00126 TIGR02424 TIGR03418 | TIGR03298 LysR\_substrate HTH\_1 TIGR02424 TIGR03418 | argP: transcriptional regulator, ArgP family LysR substrate binding domain Bacterial regulatory helix-turn-helix protein, lysR family TF\_pcaQ: pca operon transcription factor PcaQ chol\_sulf\_TF: putative choline sulfate-utilization transcription factor | 1.30E-119 7.50E-20 5.50E-15 4.00E-12 2.70E-11 |
| WP\_004529578.1 | 14150 | 13394 | - | 251 | PF13545 PF00027 PF00325 RREFamAux006 TIGR03896 | HTH\_Crp\_2 cNMP\_binding Crp RRE\_cNMP TIGR03896 | Crp-like helix-turn-helix domain Cyclic nucleotide-binding domain Bacterial regulatory proteins, crp family - cyc\_nuc\_ocin: bacteriocin-type transport-associated protein | 2.90E-20 6.00E-18 2.20E-14 2.60E-08 1.70E-05 |
| WP\_004194548.1 | 14767 | 14299 | - | 155 | PF00582 | Usp | Universal stress protein family | 2.40E-31 |
| WP\_004194465.1 | 15190 | 15421 | + | 76 | NO PFAM MATCH | - | - | - |

#### Results for WP\_264253191.1 [Burkholderia pseudomallei] back to top

#### previous - next

##### Architecture

1000 nucleotides

Link to nucleotide sequence  
  

| Accession | start | end | direction | length (aa) | Pfam/HMM | name | description | E-value |
| --- | --- | --- | --- | --- | --- | --- | --- | --- |
| WP\_004194465.1 | 60217 | 59986 | - | 76 | NO PFAM MATCH | - | - | - |
| WP\_004194548.1 | 60640 | 61108 | + | 155 | PF00582 | Usp | Universal stress protein family | 2.40E-31 |
| WP\_004529578.1 | 61257 | 62013 | + | 251 | PF13545 PF00027 PF00325 RREFamAux006 TIGR03896 | HTH\_Crp\_2 cNMP\_binding Crp RRE\_cNMP TIGR03896 | Crp-like helix-turn-helix domain Cyclic nucleotide-binding domain Bacterial regulatory proteins, crp family - cyc\_nuc\_ocin: bacteriocin-type transport-associated protein | 2.90E-20 6.00E-18 2.20E-14 2.60E-08 1.70E-05 |
| WP\_004525603.1 | 62964 | 62070 | - | 297 | TIGR03298 PF03466 PF00126 TIGR02424 TIGR03418 | TIGR03298 LysR\_substrate HTH\_1 TIGR02424 TIGR03418 | argP: transcriptional regulator, ArgP family LysR substrate binding domain Bacterial regulatory helix-turn-helix protein, lysR family TF\_pcaQ: pca operon transcription factor PcaQ chol\_sulf\_TF: putative choline sulfate-utilization transcription factor | 3.90E-120 6.00E-20 5.60E-15 4.10E-12 3.60E-11 |
| WP\_004194492.1 | 63071 | 63692 | + | 206 | TIGR00948 PF01810 TIGR00949 | TIGR00948 LysE TIGR00949 | 2a75: L-lysine exporter LysE type translocator 2A76: homoserine/Threonine efflux protein | 6.90E-53 1.40E-36 5.50E-06 |
| WP\_004532434.1 | 64853 | 63863 | - | 329 | TIGR04543 PF13649 PF13847 PF08242 TIGR03534 | TIGR04543 Methyltransf\_25 Methyltransf\_31 Methyltransf\_12 TIGR03534 | ketoArg\_3Met: 2-ketoarginine methyltransferase Methyltransferase domain Methyltransferase domain Methyltransferase domain RF\_mod\_PrmC: protein-(glutamine-N5) methyltransferase, release factor-specific | 7.20E-12 8.20E-09 2.80E-08 7.60E-07 1.50E-06 |
| WP\_004539589.1 | 66810 | 64866 | - | 647 | TIGR02026 TIGR03471 TIGR03975 TIGR04014 TIGR04013 | TIGR02026 TIGR03471 TIGR03975 TIGR04014 TIGR04013 | BchE: magnesium-protoporphyrin IX monomethyl ester anaerobic oxidative cyclase HpnJ: hopanoid biosynthesis associated radical SAM protein HpnJ rSAM\_ocin\_1: ribosomal peptide maturation radical SAM protein 1 B12\_SAM\_MJ\_0865: B12-binding domain/radical SAM domain protein, MJ\_0865 family B12\_SAM\_MJ\_1487: B12-binding domain/radical SAM domain protein, MJ\_1487 family | 5.60E-40 5.10E-32 1.70E-28 2.80E-26 9.60E-25 |
| WP\_004532432.1 | 67640 | 66806 | - | 277 | NO PFAM MATCH | - | - | - |
| WP\_264253191.1 | 68488 | 67636 | - | 283 | PF05114 PF01261 | DUF692 AP\_endonuc\_2 | Protein of unknown function (DUF692) Xylose isomerase-like TIM barrel | 6.10E-39 1.70E-06 |
| WP\_004201048.1 | 68862 | 68661 | - | 66 | NO PFAM MATCH | - | - | - |
| WP\_009971234.1 | 70418 | 69011 | - | 468 | PF00067 TIGR04538 TIGR04458 TIGR04515 | p450 TIGR04538 TIGR04458 TIGR04515 | Cytochrome P450 P450\_cycloAA\_1: cytochrome P450, cyclodipeptide synthase-associated CYP450\_TxtE: 4-nitrotryptophan synthase P450\_rel\_GT\_act: P450-derived glycosyltranferase activator | 2.10E-34 2.20E-16 2.10E-06 1.20E-05 |
| WP\_050866163.1 | 70820 | 71555 | + | 244 | PF03551 TIGR03433 | PadR TIGR03433 | Transcriptional regulator PadR-like family padR\_acidobact: transcriptional regulator, Acidobacterial, PadR-family | 2.10E-19 2.30E-12 |
| WP\_264259594.1 | 71594 | 72422 | + | 275 | PF04954 PF08021 | SIP FAD\_binding\_9 | Siderophore-interacting protein Siderophore-interacting FAD-binding domain | 2.90E-39 1.90E-37 |
| WP\_004194583.1 | 72988 | 72739 | - | 82 | NO PFAM MATCH | - | - | - |
| WP\_004196519.1 | 73181 | 73418 | + | 78 | NO PFAM MATCH | - | - | - |
| WP\_004194570.1 | 74037 | 73440 | - | 198 | PF01569 PF14378 | PAP2 PAP2\_3 | PAP2 superfamily PAP2 superfamily | 1.30E-17 4.00E-07 |
| WP\_004194452.1 | 74495 | 75236 | + | 246 | PF06314 | ADC | Acetoacetate decarboxylase (ADC) | 4.50E-79 |
| WP\_038797612.1 | 75282 | 75433 | + | 50 | TIGR01963 TIGR02632 TIGR02415 PF00106 TIGR03206 | TIGR01963 TIGR02632 TIGR02415 adh\_short TIGR03206 | PHB\_DH: 3-hydroxybutyrate dehydrogenase RhaD\_aldol-ADH: rhamnulose-1-phosphate aldolase/alcohol dehydrogenase 23BDH: acetoin reductases short chain dehydrogenase benzo\_BadH: 2-hydroxycyclohexanecarboxyl-CoA dehydrogenase | 5.00E-12 3.60E-11 1.30E-10 6.90E-10 9.50E-10 |

#### Results for WP\_270956096.1 [Burkholderia pseudomallei] back to top

#### previous - next

##### Architecture

1000 nucleotides

Link to nucleotide sequence  
  

| Accession | start | end | direction | length (aa) | Pfam/HMM | name | description | E-value |
| --- | --- | --- | --- | --- | --- | --- | --- | --- |
| WP\_004194465.1 | 457890 | 457659 | - | 76 | NO PFAM MATCH | - | - | - |
| WP\_004194548.1 | 458313 | 458781 | + | 155 | PF00582 | Usp | Universal stress protein family | 2.40E-31 |
| WP\_004529578.1 | 458930 | 459686 | + | 251 | PF13545 PF00027 PF00325 RREFamAux006 TIGR03896 | HTH\_Crp\_2 cNMP\_binding Crp RRE\_cNMP TIGR03896 | Crp-like helix-turn-helix domain Cyclic nucleotide-binding domain Bacterial regulatory proteins, crp family - cyc\_nuc\_ocin: bacteriocin-type transport-associated protein | 2.90E-20 6.00E-18 2.20E-14 2.60E-08 1.70E-05 |
| WP\_004525603.1 | 460691 | 459797 | - | 297 | TIGR03298 PF03466 PF00126 TIGR02424 TIGR03418 | TIGR03298 LysR\_substrate HTH\_1 TIGR02424 TIGR03418 | argP: transcriptional regulator, ArgP family LysR substrate binding domain Bacterial regulatory helix-turn-helix protein, lysR family TF\_pcaQ: pca operon transcription factor PcaQ chol\_sulf\_TF: putative choline sulfate-utilization transcription factor | 3.90E-120 6.00E-20 5.60E-15 4.10E-12 3.60E-11 |
| WP\_004194492.1 | 460798 | 461419 | + | 206 | TIGR00948 PF01810 TIGR00949 | TIGR00948 LysE TIGR00949 | 2a75: L-lysine exporter LysE type translocator 2A76: homoserine/Threonine efflux protein | 6.90E-53 1.40E-36 5.50E-06 |
| WP\_004532676.1 | 462580 | 461590 | - | 329 | TIGR04543 PF13649 PF13847 PF08242 TIGR03534 | TIGR04543 Methyltransf\_25 Methyltransf\_31 Methyltransf\_12 TIGR03534 | ketoArg\_3Met: 2-ketoarginine methyltransferase Methyltransferase domain Methyltransferase domain Methyltransferase domain RF\_mod\_PrmC: protein-(glutamine-N5) methyltransferase, release factor-specific | 7.10E-12 8.20E-09 2.80E-08 7.60E-07 1.40E-06 |
| WP\_160447439.1 | 464537 | 462593 | - | 647 | TIGR02026 TIGR03471 TIGR03975 TIGR04014 TIGR04013 | TIGR02026 TIGR03471 TIGR03975 TIGR04014 TIGR04013 | BchE: magnesium-protoporphyrin IX monomethyl ester anaerobic oxidative cyclase HpnJ: hopanoid biosynthesis associated radical SAM protein HpnJ rSAM\_ocin\_1: ribosomal peptide maturation radical SAM protein 1 B12\_SAM\_MJ\_0865: B12-binding domain/radical SAM domain protein, MJ\_0865 family B12\_SAM\_MJ\_1487: B12-binding domain/radical SAM domain protein, MJ\_1487 family | 5.60E-40 5.10E-32 1.80E-28 2.80E-26 9.60E-25 |
| WP\_004529573.1 | 465355 | 464533 | - | 273 | NO PFAM MATCH | - | - | - |
| WP\_270956096.1 | 466203 | 465351 | - | 283 | PF05114 PF01261 | DUF692 AP\_endonuc\_2 | Protein of unknown function (DUF692) Xylose isomerase-like TIM barrel | 1.10E-39 1.50E-06 |
| WP\_004201048.1 | 466577 | 466376 | - | 66 | NO PFAM MATCH | - | - | - |
| WP\_024428486.1 | 468133 | 466726 | - | 468 | PF00067 TIGR04538 TIGR04458 TIGR04515 | p450 TIGR04538 TIGR04458 TIGR04515 | Cytochrome P450 P450\_cycloAA\_1: cytochrome P450, cyclodipeptide synthase-associated CYP450\_TxtE: 4-nitrotryptophan synthase P450\_rel\_GT\_act: P450-derived glycosyltranferase activator | 7.00E-34 4.60E-16 2.40E-06 9.80E-06 |
| WP\_004529569.1 | 468535 | 469240 | + | 234 | PF03551 TIGR03433 | PadR TIGR03433 | Transcriptional regulator PadR-like family padR\_acidobact: transcriptional regulator, Acidobacterial, PadR-family | 2.00E-19 2.10E-12 |
| WP\_004194445.1 | 469279 | 470107 | + | 275 | PF04954 PF08021 | SIP FAD\_binding\_9 | Siderophore-interacting protein Siderophore-interacting FAD-binding domain | 2.90E-39 1.90E-37 |
| WP\_004529568.1 | 470679 | 470430 | - | 82 | NO PFAM MATCH | - | - | - |
| WP\_004196519.1 | 470872 | 471109 | + | 78 | NO PFAM MATCH | - | - | - |
| WP\_004525615.1 | 471728 | 471131 | - | 198 | PF01569 PF14378 | PAP2 PAP2\_3 | PAP2 superfamily PAP2 superfamily | 7.40E-18 7.90E-07 |
| WP\_004194452.1 | 472186 | 472927 | + | 246 | PF06314 | ADC | Acetoacetate decarboxylase (ADC) | 4.50E-79 |

#### Results for WP\_045243502.1 [Burkholderia pseudomallei] back to top

#### previous - next

##### Architecture

1000 nucleotides

Link to nucleotide sequence  
  

| Accession | start | end | direction | length (aa) | Pfam/HMM | name | description | E-value |
| --- | --- | --- | --- | --- | --- | --- | --- | --- |
| WP\_038797612.1 | 151 | 0 | - | 50 | TIGR01963 TIGR02632 TIGR02415 PF00106 TIGR03206 | TIGR01963 TIGR02632 TIGR02415 adh\_short TIGR03206 | PHB\_DH: 3-hydroxybutyrate dehydrogenase RhaD\_aldol-ADH: rhamnulose-1-phosphate aldolase/alcohol dehydrogenase 23BDH: acetoin reductases short chain dehydrogenase benzo\_BadH: 2-hydroxycyclohexanecarboxyl-CoA dehydrogenase | 5.00E-12 3.60E-11 1.30E-10 6.90E-10 9.50E-10 |
| WP\_004194452.1 | 938 | 197 | - | 246 | PF06314 | ADC | Acetoacetate decarboxylase (ADC) | 4.50E-79 |
| WP\_004525615.1 | 1396 | 1993 | + | 198 | PF01569 PF14378 | PAP2 PAP2\_3 | PAP2 superfamily PAP2 superfamily | 7.40E-18 7.90E-07 |
| WP\_004196519.1 | 2252 | 2015 | - | 78 | NO PFAM MATCH | - | - | - |
| WP\_004194583.1 | 2445 | 2694 | + | 82 | NO PFAM MATCH | - | - | - |
| WP\_004532447.1 | 3017 | 3852 | + | 278 | INFERRED GENE | - | - | - |
| WP\_004196523.1 | 4599 | 3891 | - | 235 | PF03551 TIGR03433 | PadR TIGR03433 | Transcriptional regulator PadR-like family padR\_acidobact: transcriptional regulator, Acidobacterial, PadR-family | 2.00E-19 2.20E-12 |
| WP\_004194516.1 | 5006 | 6413 | + | 468 | PF00067 TIGR04538 TIGR04458 TIGR04515 | p450 TIGR04538 TIGR04458 TIGR04515 | Cytochrome P450 P450\_cycloAA\_1: cytochrome P450, cyclodipeptide synthase-associated CYP450\_TxtE: 4-nitrotryptophan synthase P450\_rel\_GT\_act: P450-derived glycosyltranferase activator | 5.60E-34 3.30E-16 4.80E-06 2.50E-05 |
| WP\_004201048.1 | 6562 | 6763 | + | 66 | NO PFAM MATCH | - | - | - |
| WP\_045243502.1 | 6936 | 7788 | + | 283 | PF05114 PF01261 | DUF692 AP\_endonuc\_2 | Protein of unknown function (DUF692) Xylose isomerase-like TIM barrel | 2.80E-39 1.50E-06 |
| WP\_004529573.1 | 7784 | 8606 | + | 273 | NO PFAM MATCH | - | - | - |
| WP\_045243618.1 | 8602 | 10546 | + | 647 | TIGR02026 TIGR03471 TIGR03975 TIGR04014 TIGR04013 | TIGR02026 TIGR03471 TIGR03975 TIGR04014 TIGR04013 | BchE: magnesium-protoporphyrin IX monomethyl ester anaerobic oxidative cyclase HpnJ: hopanoid biosynthesis associated radical SAM protein HpnJ rSAM\_ocin\_1: ribosomal peptide maturation radical SAM protein 1 B12\_SAM\_MJ\_0865: B12-binding domain/radical SAM domain protein, MJ\_0865 family B12\_SAM\_MJ\_1487: B12-binding domain/radical SAM domain protein, MJ\_1487 family | 5.90E-40 5.20E-32 1.80E-28 2.80E-26 9.60E-25 |
| WP\_004545057.1 | 10559 | 11549 | + | 329 | TIGR04543 PF13649 PF13847 PF08242 TIGR03534 | TIGR04543 Methyltransf\_25 Methyltransf\_31 Methyltransf\_12 TIGR03534 | ketoArg\_3Met: 2-ketoarginine methyltransferase Methyltransferase domain Methyltransferase domain Methyltransferase domain RF\_mod\_PrmC: protein-(glutamine-N5) methyltransferase, release factor-specific | 1.10E-11 8.40E-09 2.80E-08 7.50E-07 1.40E-06 |
| WP\_004194492.1 | 12341 | 11720 | - | 206 | TIGR00948 PF01810 TIGR00949 | TIGR00948 LysE TIGR00949 | 2a75: L-lysine exporter LysE type translocator 2A76: homoserine/Threonine efflux protein | 6.90E-53 1.40E-36 5.50E-06 |
| WP\_004525603.1 | 12448 | 13342 | + | 297 | TIGR03298 PF03466 PF00126 TIGR02424 TIGR03418 | TIGR03298 LysR\_substrate HTH\_1 TIGR02424 TIGR03418 | argP: transcriptional regulator, ArgP family LysR substrate binding domain Bacterial regulatory helix-turn-helix protein, lysR family TF\_pcaQ: pca operon transcription factor PcaQ chol\_sulf\_TF: putative choline sulfate-utilization transcription factor | 3.90E-120 6.00E-20 5.60E-15 4.10E-12 3.60E-11 |
| WP\_004529578.1 | 14164 | 13408 | - | 251 | PF13545 PF00027 PF00325 RREFamAux006 TIGR03896 | HTH\_Crp\_2 cNMP\_binding Crp RRE\_cNMP TIGR03896 | Crp-like helix-turn-helix domain Cyclic nucleotide-binding domain Bacterial regulatory proteins, crp family - cyc\_nuc\_ocin: bacteriocin-type transport-associated protein | 2.90E-20 6.00E-18 2.20E-14 2.60E-08 1.70E-05 |
| WP\_004194548.1 | 14781 | 14313 | - | 155 | PF00582 | Usp | Universal stress protein family | 2.40E-31 |
| WP\_004194465.1 | 15204 | 15435 | + | 76 | NO PFAM MATCH | - | - | - |

#### Results for WP\_038725437.1 [Burkholderia pseudomallei] back to top

#### previous - next

##### Architecture

1000 nucleotides

Link to nucleotide sequence  
  

| Accession | start | end | direction | length (aa) | Pfam/HMM | name | description | E-value |
| --- | --- | --- | --- | --- | --- | --- | --- | --- |
| WP\_004194455.1 | 84989 | 84203 | - | 261 | TIGR01963 PF00106 PF13561 TIGR03971 TIGR02415 | TIGR01963 adh\_short adh\_short\_C2 TIGR03971 TIGR02415 | PHB\_DH: 3-hydroxybutyrate dehydrogenase short chain dehydrogenase Enoyl-(Acyl carrier protein) reductase SDR\_subfam\_1: SDR family mycofactocin-dependent oxidoreductase 23BDH: acetoin reductases | 4.30E-104 1.80E-52 2.40E-52 6.40E-52 8.90E-52 |
| WP\_004194452.1 | 85776 | 85035 | - | 246 | PF06314 | ADC | Acetoacetate decarboxylase (ADC) | 4.50E-79 |
| WP\_041194472.1 | 86234 | 86831 | + | 198 | PF01569 PF14378 | PAP2 PAP2\_3 | PAP2 superfamily PAP2 superfamily | 7.70E-18 8.00E-07 |
| WP\_004196519.1 | 87090 | 86853 | - | 78 | NO PFAM MATCH | - | - | - |
| WP\_004194445.1 | 88683 | 87855 | - | 275 | PF04954 PF08021 | SIP FAD\_binding\_9 | Siderophore-interacting protein Siderophore-interacting FAD-binding domain | 2.90E-39 1.90E-37 |
| WP\_004196523.1 | 89430 | 88722 | - | 235 | PF03551 TIGR03433 | PadR TIGR03433 | Transcriptional regulator PadR-like family padR\_acidobact: transcriptional regulator, Acidobacterial, PadR-family | 2.00E-19 2.20E-12 |
| WP\_024428486.1 | 89837 | 91244 | + | 468 | PF00067 TIGR04538 TIGR04458 TIGR04515 | p450 TIGR04538 TIGR04458 TIGR04515 | Cytochrome P450 P450\_cycloAA\_1: cytochrome P450, cyclodipeptide synthase-associated CYP450\_TxtE: 4-nitrotryptophan synthase P450\_rel\_GT\_act: P450-derived glycosyltranferase activator | 7.00E-34 4.60E-16 2.40E-06 9.80E-06 |
| WP\_004201048.1 | 91393 | 91594 | + | 66 | NO PFAM MATCH | - | - | - |
| WP\_038725437.1 | 91767 | 92619 | + | 283 | PF05114 PF01261 | DUF692 AP\_endonuc\_2 | Protein of unknown function (DUF692) Xylose isomerase-like TIM barrel | 1.90E-39 1.50E-06 |
| WP\_038798248.1 | 92615 | 93449 | + | 277 | NO PFAM MATCH | - | - | - |
| WP\_043289401.1 | 93445 | 95389 | + | 647 | TIGR02026 TIGR03471 TIGR03975 TIGR04014 TIGR04013 | TIGR02026 TIGR03471 TIGR03975 TIGR04014 TIGR04013 | BchE: magnesium-protoporphyrin IX monomethyl ester anaerobic oxidative cyclase HpnJ: hopanoid biosynthesis associated radical SAM protein HpnJ rSAM\_ocin\_1: ribosomal peptide maturation radical SAM protein 1 B12\_SAM\_MJ\_0865: B12-binding domain/radical SAM domain protein, MJ\_0865 family B12\_SAM\_MJ\_1487: B12-binding domain/radical SAM domain protein, MJ\_1487 family | 4.50E-39 8.80E-32 3.30E-27 1.00E-25 3.00E-24 |
| WP\_004545057.1 | 95402 | 96392 | + | 329 | TIGR04543 PF13649 PF13847 PF08242 TIGR03534 | TIGR04543 Methyltransf\_25 Methyltransf\_31 Methyltransf\_12 TIGR03534 | ketoArg\_3Met: 2-ketoarginine methyltransferase Methyltransferase domain Methyltransferase domain Methyltransferase domain RF\_mod\_PrmC: protein-(glutamine-N5) methyltransferase, release factor-specific | 1.10E-11 8.40E-09 2.80E-08 7.50E-07 1.40E-06 |
| WP\_004194492.1 | 97184 | 96563 | - | 206 | TIGR00948 PF01810 TIGR00949 | TIGR00948 LysE TIGR00949 | 2a75: L-lysine exporter LysE type translocator 2A76: homoserine/Threonine efflux protein | 6.90E-53 1.40E-36 5.50E-06 |
| WP\_076859412.1 | 97291 | 98185 | + | 297 | TIGR03298 PF03466 PF00126 TIGR02424 TIGR03418 | TIGR03298 LysR\_substrate HTH\_1 TIGR02424 TIGR03418 | argP: transcriptional regulator, ArgP family LysR substrate binding domain Bacterial regulatory helix-turn-helix protein, lysR family TF\_pcaQ: pca operon transcription factor PcaQ chol\_sulf\_TF: putative choline sulfate-utilization transcription factor | 2.30E-119 9.70E-20 5.80E-15 5.80E-12 3.70E-11 |
| WP\_004529578.1 | 99005 | 98249 | - | 251 | PF13545 PF00027 PF00325 RREFamAux006 TIGR03896 | HTH\_Crp\_2 cNMP\_binding Crp RRE\_cNMP TIGR03896 | Crp-like helix-turn-helix domain Cyclic nucleotide-binding domain Bacterial regulatory proteins, crp family - cyc\_nuc\_ocin: bacteriocin-type transport-associated protein | 2.90E-20 6.00E-18 2.20E-14 2.60E-08 1.70E-05 |
| WP\_004194548.1 | 99622 | 99154 | - | 155 | PF00582 | Usp | Universal stress protein family | 2.40E-31 |
| WP\_004194465.1 | 100045 | 100276 | + | 76 | NO PFAM MATCH | - | - | - |

#### Results for WP\_041189437.1 [Burkholderia pseudomallei MSHR4000] back to top

#### previous - next

##### Architecture

1000 nucleotides

Link to nucleotide sequence  
  

| Accession | start | end | direction | length (aa) | Pfam/HMM | name | description | E-value |
| --- | --- | --- | --- | --- | --- | --- | --- | --- |
| WP\_004194455.1 | 2486250 | 2485464 | - | 261 | TIGR01963 PF00106 PF13561 TIGR03971 TIGR02415 | TIGR01963 adh\_short adh\_short\_C2 TIGR03971 TIGR02415 | PHB\_DH: 3-hydroxybutyrate dehydrogenase short chain dehydrogenase Enoyl-(Acyl carrier protein) reductase SDR\_subfam\_1: SDR family mycofactocin-dependent oxidoreductase 23BDH: acetoin reductases | 4.30E-104 1.80E-52 2.40E-52 6.40E-52 8.90E-52 |
| WP\_004194452.1 | 2487037 | 2486296 | - | 246 | PF06314 | ADC | Acetoacetate decarboxylase (ADC) | 4.50E-79 |
| WP\_004525615.1 | 2487495 | 2488092 | + | 198 | PF01569 PF14378 | PAP2 PAP2\_3 | PAP2 superfamily PAP2 superfamily | 7.40E-18 7.90E-07 |
| WP\_009922809.1 | 2488351 | 2488114 | - | 78 | NO PFAM MATCH | - | - | - |
| WP\_004194445.1 | 2489944 | 2489116 | - | 275 | PF04954 PF08021 | SIP FAD\_binding\_9 | Siderophore-interacting protein Siderophore-interacting FAD-binding domain | 2.90E-39 1.90E-37 |
| WP\_004548999.1 | 2490688 | 2489983 | - | 234 | PF03551 TIGR03433 | PadR TIGR03433 | Transcriptional regulator PadR-like family padR\_acidobact: transcriptional regulator, Acidobacterial, PadR-family | 2.00E-19 2.10E-12 |
| WP\_004194516.1 | 2491101 | 2492508 | + | 468 | PF00067 TIGR04538 TIGR04458 TIGR04515 | p450 TIGR04538 TIGR04458 TIGR04515 | Cytochrome P450 P450\_cycloAA\_1: cytochrome P450, cyclodipeptide synthase-associated CYP450\_TxtE: 4-nitrotryptophan synthase P450\_rel\_GT\_act: P450-derived glycosyltranferase activator | 5.60E-34 3.30E-16 4.80E-06 2.50E-05 |
| WP\_004201048.1 | 2492657 | 2492858 | + | 66 | NO PFAM MATCH | - | - | - |
| WP\_041189437.1 | 2493031 | 2493883 | + | 283 | PF05114 PF01261 | DUF692 AP\_endonuc\_2 | Protein of unknown function (DUF692) Xylose isomerase-like TIM barrel | 1.90E-39 1.50E-06 |
| WP\_038733402.1 | 2493879 | 2494701 | + | 273 | NO PFAM MATCH | - | - | - |
| WP\_041189555.1 | 2494697 | 2496641 | + | 647 | TIGR02026 TIGR03471 TIGR03975 TIGR04014 TIGR04013 | TIGR02026 TIGR03471 TIGR03975 TIGR04014 TIGR04013 | BchE: magnesium-protoporphyrin IX monomethyl ester anaerobic oxidative cyclase HpnJ: hopanoid biosynthesis associated radical SAM protein HpnJ rSAM\_ocin\_1: ribosomal peptide maturation radical SAM protein 1 B12\_SAM\_MJ\_0865: B12-binding domain/radical SAM domain protein, MJ\_0865 family B12\_SAM\_MJ\_1487: B12-binding domain/radical SAM domain protein, MJ\_1487 family | 5.70E-39 1.20E-31 1.30E-26 4.50E-25 1.00E-23 |
| WP\_004545057.1 | 2496654 | 2497644 | + | 329 | TIGR04543 PF13649 PF13847 PF08242 TIGR03534 | TIGR04543 Methyltransf\_25 Methyltransf\_31 Methyltransf\_12 TIGR03534 | ketoArg\_3Met: 2-ketoarginine methyltransferase Methyltransferase domain Methyltransferase domain Methyltransferase domain RF\_mod\_PrmC: protein-(glutamine-N5) methyltransferase, release factor-specific | 1.10E-11 8.40E-09 2.80E-08 7.50E-07 1.40E-06 |
| WP\_004194492.1 | 2498436 | 2497815 | - | 206 | TIGR00948 PF01810 TIGR00949 | TIGR00948 LysE TIGR00949 | 2a75: L-lysine exporter LysE type translocator 2A76: homoserine/Threonine efflux protein | 6.90E-53 1.40E-36 5.50E-06 |
| WP\_004525603.1 | 2498543 | 2499437 | + | 297 | TIGR03298 PF03466 PF00126 TIGR02424 TIGR03418 | TIGR03298 LysR\_substrate HTH\_1 TIGR02424 TIGR03418 | argP: transcriptional regulator, ArgP family LysR substrate binding domain Bacterial regulatory helix-turn-helix protein, lysR family TF\_pcaQ: pca operon transcription factor PcaQ chol\_sulf\_TF: putative choline sulfate-utilization transcription factor | 3.90E-120 6.00E-20 5.60E-15 4.10E-12 3.60E-11 |
| WP\_004529578.1 | 2500331 | 2499575 | - | 251 | PF13545 PF00027 PF00325 RREFamAux006 TIGR03896 | HTH\_Crp\_2 cNMP\_binding Crp RRE\_cNMP TIGR03896 | Crp-like helix-turn-helix domain Cyclic nucleotide-binding domain Bacterial regulatory proteins, crp family - cyc\_nuc\_ocin: bacteriocin-type transport-associated protein | 2.90E-20 6.00E-18 2.20E-14 2.60E-08 1.70E-05 |
| WP\_004194548.1 | 2500948 | 2500480 | - | 155 | PF00582 | Usp | Universal stress protein family | 2.40E-31 |
| WP\_004194465.1 | 2501371 | 2501602 | + | 76 | NO PFAM MATCH | - | - | - |

#### Results for WP\_038731252.1 [Burkholderia pseudomallei 576] back to top

#### previous - next

##### Architecture

1000 nucleotides

Link to nucleotide sequence  
  

| Accession | start | end | direction | length (aa) | Pfam/HMM | name | description | E-value |
| --- | --- | --- | --- | --- | --- | --- | --- | --- |
| WP\_004194465.1 | 448210 | 447979 | - | 76 | NO PFAM MATCH | - | - | - |
| WP\_004194548.1 | 448633 | 449101 | + | 155 | PF00582 | Usp | Universal stress protein family | 2.40E-31 |
| WP\_004529578.1 | 449250 | 450006 | + | 251 | PF13545 PF00027 PF00325 RREFamAux006 TIGR03896 | HTH\_Crp\_2 cNMP\_binding Crp RRE\_cNMP TIGR03896 | Crp-like helix-turn-helix domain Cyclic nucleotide-binding domain Bacterial regulatory proteins, crp family - cyc\_nuc\_ocin: bacteriocin-type transport-associated protein | 2.90E-20 6.00E-18 2.20E-14 2.60E-08 1.70E-05 |
| WP\_004194503.1 | 450964 | 450070 | - | 297 | TIGR03298 PF03466 PF00126 TIGR02424 TIGR03418 | TIGR03298 LysR\_substrate HTH\_1 TIGR02424 TIGR03418 | argP: transcriptional regulator, ArgP family LysR substrate binding domain Bacterial regulatory helix-turn-helix protein, lysR family TF\_pcaQ: pca operon transcription factor PcaQ chol\_sulf\_TF: putative choline sulfate-utilization transcription factor | 3.90E-120 4.80E-20 5.30E-15 5.90E-12 2.30E-11 |
| WP\_004194492.1 | 451071 | 451692 | + | 206 | TIGR00948 PF01810 TIGR00949 | TIGR00948 LysE TIGR00949 | 2a75: L-lysine exporter LysE type translocator 2A76: homoserine/Threonine efflux protein | 6.90E-53 1.40E-36 5.50E-06 |
| WP\_004545057.1 | 452853 | 451863 | - | 329 | TIGR04543 PF13649 PF13847 PF08242 TIGR03534 | TIGR04543 Methyltransf\_25 Methyltransf\_31 Methyltransf\_12 TIGR03534 | ketoArg\_3Met: 2-ketoarginine methyltransferase Methyltransferase domain Methyltransferase domain Methyltransferase domain RF\_mod\_PrmC: protein-(glutamine-N5) methyltransferase, release factor-specific | 1.10E-11 8.40E-09 2.80E-08 7.50E-07 1.40E-06 |
| WP\_004544988.1 | 454810 | 452866 | - | 647 | TIGR02026 TIGR03471 TIGR03975 TIGR04014 TIGR04013 | TIGR02026 TIGR03471 TIGR03975 TIGR04014 TIGR04013 | BchE: magnesium-protoporphyrin IX monomethyl ester anaerobic oxidative cyclase HpnJ: hopanoid biosynthesis associated radical SAM protein HpnJ rSAM\_ocin\_1: ribosomal peptide maturation radical SAM protein 1 B12\_SAM\_MJ\_0865: B12-binding domain/radical SAM domain protein, MJ\_0865 family B12\_SAM\_MJ\_1487: B12-binding domain/radical SAM domain protein, MJ\_1487 family | 3.70E-39 8.80E-32 3.30E-27 1.00E-25 3.00E-24 |
| WP\_004544974.1 | 455640 | 454806 | - | 277 | NO PFAM MATCH | - | - | - |
| WP\_038731252.1 | 456488 | 455636 | - | 283 | PF05114 PF01261 | DUF692 AP\_endonuc\_2 | Protein of unknown function (DUF692) Xylose isomerase-like TIM barrel | 8.90E-40 1.50E-06 |
| WP\_004201048.1 | 456862 | 456661 | - | 66 | NO PFAM MATCH | - | - | - |
| WP\_004194516.1 | 458418 | 457011 | - | 468 | PF00067 TIGR04538 TIGR04458 TIGR04515 | p450 TIGR04538 TIGR04458 TIGR04515 | Cytochrome P450 P450\_cycloAA\_1: cytochrome P450, cyclodipeptide synthase-associated CYP450\_TxtE: 4-nitrotryptophan synthase P450\_rel\_GT\_act: P450-derived glycosyltranferase activator | 5.60E-34 3.30E-16 4.80E-06 2.50E-05 |
| WP\_004196523.1 | 458825 | 459533 | + | 235 | PF03551 TIGR03433 | PadR TIGR03433 | Transcriptional regulator PadR-like family padR\_acidobact: transcriptional regulator, Acidobacterial, PadR-family | 2.00E-19 2.20E-12 |
| WP\_004194445.1 | 459572 | 460400 | + | 275 | PF04954 PF08021 | SIP FAD\_binding\_9 | Siderophore-interacting protein Siderophore-interacting FAD-binding domain | 2.90E-39 1.90E-37 |
| WP\_004194583.1 | 460972 | 460723 | - | 82 | NO PFAM MATCH | - | - | - |
| WP\_004529566.1 | 461165 | 461402 | + | 78 | NO PFAM MATCH | - | - | - |
| WP\_004525615.1 | 462021 | 461424 | - | 198 | PF01569 PF14378 | PAP2 PAP2\_3 | PAP2 superfamily PAP2 superfamily | 7.40E-18 7.90E-07 |
| WP\_004194452.1 | 462479 | 463220 | + | 246 | PF06314 | ADC | Acetoacetate decarboxylase (ADC) | 4.50E-79 |

#### Results for WP\_038763239.1 [Burkholderia pseudomallei] back to top

#### previous - next

##### Architecture

1000 nucleotides

Link to nucleotide sequence  
  

| Accession | start | end | direction | length (aa) | Pfam/HMM | name | description | E-value |
| --- | --- | --- | --- | --- | --- | --- | --- | --- |
| WP\_122650933.1 | 101 | 0 | - | 34 | TIGR01963 TIGR01832 TIGR03971 PF00106 TIGR02632 | TIGR01963 TIGR01832 TIGR03971 adh\_short TIGR02632 | PHB\_DH: 3-hydroxybutyrate dehydrogenase kduD: 2-deoxy-D-gluconate 3-dehydrogenase SDR\_subfam\_1: SDR family mycofactocin-dependent oxidoreductase short chain dehydrogenase RhaD\_aldol-ADH: rhamnulose-1-phosphate aldolase/alcohol dehydrogenase | 1.90E-08 5.50E-08 1.80E-07 4.40E-07 1.20E-06 |
| WP\_004194452.1 | 888 | 147 | - | 246 | PF06314 | ADC | Acetoacetate decarboxylase (ADC) | 4.50E-79 |
| WP\_041194472.1 | 1346 | 1943 | + | 198 | PF01569 PF14378 | PAP2 PAP2\_3 | PAP2 superfamily PAP2 superfamily | 7.70E-18 8.00E-07 |
| WP\_004196519.1 | 2202 | 1965 | - | 78 | NO PFAM MATCH | - | - | - |
| WP\_076850552.1 | 2395 | 2644 | + | 82 | NO PFAM MATCH | - | - | - |
| WP\_004194445.1 | 3795 | 2967 | - | 275 | PF04954 PF08021 | SIP FAD\_binding\_9 | Siderophore-interacting protein Siderophore-interacting FAD-binding domain | 2.90E-39 1.90E-37 |
| WP\_004529569.1 | 4539 | 3834 | - | 234 | PF03551 TIGR03433 | PadR TIGR03433 | Transcriptional regulator PadR-like family padR\_acidobact: transcriptional regulator, Acidobacterial, PadR-family | 2.00E-19 2.10E-12 |
| WP\_004529570.1 | 4941 | 6348 | + | 468 | PF00067 TIGR04538 TIGR04458 TIGR04515 | p450 TIGR04538 TIGR04458 TIGR04515 | Cytochrome P450 P450\_cycloAA\_1: cytochrome P450, cyclodipeptide synthase-associated CYP450\_TxtE: 4-nitrotryptophan synthase P450\_rel\_GT\_act: P450-derived glycosyltranferase activator | 5.20E-33 1.40E-15 3.50E-05 6.30E-05 |
| WP\_004201048.1 | 6497 | 6698 | + | 66 | NO PFAM MATCH | - | - | - |
| WP\_038763239.1 | 6871 | 7723 | + | 283 | PF05114 PF01261 | DUF692 AP\_endonuc\_2 | Protein of unknown function (DUF692) Xylose isomerase-like TIM barrel | 2.50E-39 1.90E-06 |
| WP\_038731003.1 | 7719 | 8541 | + | 273 | NO PFAM MATCH | - | - | - |
| WP\_038736602.1 | 8537 | 10481 | + | 647 | TIGR02026 TIGR03471 TIGR03975 TIGR04014 TIGR04013 | TIGR02026 TIGR03471 TIGR03975 TIGR04014 TIGR04013 | BchE: magnesium-protoporphyrin IX monomethyl ester anaerobic oxidative cyclase HpnJ: hopanoid biosynthesis associated radical SAM protein HpnJ rSAM\_ocin\_1: ribosomal peptide maturation radical SAM protein 1 B12\_SAM\_MJ\_0865: B12-binding domain/radical SAM domain protein, MJ\_0865 family B12\_SAM\_MJ\_1487: B12-binding domain/radical SAM domain protein, MJ\_1487 family | 1.30E-38 7.30E-32 3.30E-27 8.50E-26 1.70E-24 |
| WP\_038739802.1 | 10494 | 11484 | + | 329 | TIGR04543 PF13649 PF13847 PF08242 TIGR03534 | TIGR04543 Methyltransf\_25 Methyltransf\_31 Methyltransf\_12 TIGR03534 | ketoArg\_3Met: 2-ketoarginine methyltransferase Methyltransferase domain Methyltransferase domain Methyltransferase domain RF\_mod\_PrmC: protein-(glutamine-N5) methyltransferase, release factor-specific | 6.60E-12 8.40E-09 2.70E-08 7.00E-07 1.30E-06 |
| WP\_004194492.1 | 12276 | 11655 | - | 206 | TIGR00948 PF01810 TIGR00949 | TIGR00948 LysE TIGR00949 | 2a75: L-lysine exporter LysE type translocator 2A76: homoserine/Threonine efflux protein | 6.90E-53 1.40E-36 5.50E-06 |
| WP\_004525603.1 | 12383 | 13277 | + | 297 | TIGR03298 PF03466 PF00126 TIGR02424 TIGR03418 | TIGR03298 LysR\_substrate HTH\_1 TIGR02424 TIGR03418 | argP: transcriptional regulator, ArgP family LysR substrate binding domain Bacterial regulatory helix-turn-helix protein, lysR family TF\_pcaQ: pca operon transcription factor PcaQ chol\_sulf\_TF: putative choline sulfate-utilization transcription factor | 3.90E-120 6.00E-20 5.60E-15 4.10E-12 3.60E-11 |
| WP\_004529578.1 | 14091 | 13335 | - | 251 | PF13545 PF00027 PF00325 RREFamAux006 TIGR03896 | HTH\_Crp\_2 cNMP\_binding Crp RRE\_cNMP TIGR03896 | Crp-like helix-turn-helix domain Cyclic nucleotide-binding domain Bacterial regulatory proteins, crp family - cyc\_nuc\_ocin: bacteriocin-type transport-associated protein | 2.90E-20 6.00E-18 2.20E-14 2.60E-08 1.70E-05 |
| WP\_004194548.1 | 14708 | 14240 | - | 155 | PF00582 | Usp | Universal stress protein family | 2.40E-31 |
| WP\_004194465.1 | 15131 | 15362 | + | 76 | NO PFAM MATCH | - | - | - |

#### Results for WP\_038774393.1 [Burkholderia pseudomallei] back to top

#### previous - next

##### Architecture

1000 nucleotides

Link to nucleotide sequence  
  

| Accession | start | end | direction | length (aa) | Pfam/HMM | name | description | E-value |
| --- | --- | --- | --- | --- | --- | --- | --- | --- |
| WP\_203561953.1 | 402 | 0 | - | 134 | TIGR01963 PF00106 TIGR02632 TIGR02415 TIGR03206 | TIGR01963 adh\_short TIGR02632 TIGR02415 TIGR03206 | PHB\_DH: 3-hydroxybutyrate dehydrogenase short chain dehydrogenase RhaD\_aldol-ADH: rhamnulose-1-phosphate aldolase/alcohol dehydrogenase 23BDH: acetoin reductases benzo\_BadH: 2-hydroxycyclohexanecarboxyl-CoA dehydrogenase | 4.00E-45 1.10E-32 3.80E-32 2.40E-29 1.20E-28 |
| WP\_004194452.1 | 1189 | 448 | - | 246 | PF06314 | ADC | Acetoacetate decarboxylase (ADC) | 4.50E-79 |
| WP\_041194472.1 | 1647 | 2244 | + | 198 | PF01569 PF14378 | PAP2 PAP2\_3 | PAP2 superfamily PAP2 superfamily | 7.70E-18 8.00E-07 |
| WP\_004196519.1 | 2503 | 2266 | - | 78 | NO PFAM MATCH | - | - | - |
| WP\_004194445.1 | 4096 | 3268 | - | 275 | PF04954 PF08021 | SIP FAD\_binding\_9 | Siderophore-interacting protein Siderophore-interacting FAD-binding domain | 2.90E-39 1.90E-37 |
| WP\_004529569.1 | 4840 | 4135 | - | 234 | PF03551 TIGR03433 | PadR TIGR03433 | Transcriptional regulator PadR-like family padR\_acidobact: transcriptional regulator, Acidobacterial, PadR-family | 2.00E-19 2.10E-12 |
| WP\_004529570.1 | 5242 | 6649 | + | 468 | PF00067 TIGR04538 TIGR04458 TIGR04515 | p450 TIGR04538 TIGR04458 TIGR04515 | Cytochrome P450 P450\_cycloAA\_1: cytochrome P450, cyclodipeptide synthase-associated CYP450\_TxtE: 4-nitrotryptophan synthase P450\_rel\_GT\_act: P450-derived glycosyltranferase activator | 5.20E-33 1.40E-15 3.50E-05 6.30E-05 |
| WP\_004201048.1 | 6798 | 6999 | + | 66 | NO PFAM MATCH | - | - | - |
| WP\_038774393.1 | 7172 | 8024 | + | 283 | PF05114 PF01261 | DUF692 AP\_endonuc\_2 | Protein of unknown function (DUF692) Xylose isomerase-like TIM barrel | 1.80E-39 1.50E-06 |
| WP\_038733402.1 | 8020 | 8842 | + | 273 | NO PFAM MATCH | - | - | - |
| WP\_038800877.1 | 8838 | 10782 | + | 647 | TIGR02026 TIGR03471 TIGR03975 TIGR04014 TIGR04013 | TIGR02026 TIGR03471 TIGR03975 TIGR04014 TIGR04013 | BchE: magnesium-protoporphyrin IX monomethyl ester anaerobic oxidative cyclase HpnJ: hopanoid biosynthesis associated radical SAM protein HpnJ rSAM\_ocin\_1: ribosomal peptide maturation radical SAM protein 1 B12\_SAM\_MJ\_0865: B12-binding domain/radical SAM domain protein, MJ\_0865 family B12\_SAM\_MJ\_1487: B12-binding domain/radical SAM domain protein, MJ\_1487 family | 5.10E-39 7.20E-32 3.20E-27 1.50E-25 3.00E-24 |
| WP\_004532676.1 | 10795 | 11785 | + | 329 | TIGR04543 PF13649 PF13847 PF08242 TIGR03534 | TIGR04543 Methyltransf\_25 Methyltransf\_31 Methyltransf\_12 TIGR03534 | ketoArg\_3Met: 2-ketoarginine methyltransferase Methyltransferase domain Methyltransferase domain Methyltransferase domain RF\_mod\_PrmC: protein-(glutamine-N5) methyltransferase, release factor-specific | 7.10E-12 8.20E-09 2.80E-08 7.60E-07 1.40E-06 |
| WP\_004194492.1 | 12577 | 11956 | - | 206 | TIGR00948 PF01810 TIGR00949 | TIGR00948 LysE TIGR00949 | 2a75: L-lysine exporter LysE type translocator 2A76: homoserine/Threonine efflux protein | 6.90E-53 1.40E-36 5.50E-06 |
| WP\_038734008.1 | 12684 | 13578 | + | 297 | TIGR03298 PF03466 PF00126 TIGR02424 TIGR03418 | TIGR03298 LysR\_substrate HTH\_1 TIGR02424 TIGR03418 | argP: transcriptional regulator, ArgP family LysR substrate binding domain Bacterial regulatory helix-turn-helix protein, lysR family TF\_pcaQ: pca operon transcription factor PcaQ chol\_sulf\_TF: putative choline sulfate-utilization transcription factor | 4.60E-120 8.40E-20 5.60E-15 4.40E-12 3.60E-11 |
| WP\_004529578.1 | 14398 | 13642 | - | 251 | PF13545 PF00027 PF00325 RREFamAux006 TIGR03896 | HTH\_Crp\_2 cNMP\_binding Crp RRE\_cNMP TIGR03896 | Crp-like helix-turn-helix domain Cyclic nucleotide-binding domain Bacterial regulatory proteins, crp family - cyc\_nuc\_ocin: bacteriocin-type transport-associated protein | 2.90E-20 6.00E-18 2.20E-14 2.60E-08 1.70E-05 |
| WP\_004194548.1 | 15015 | 14547 | - | 155 | PF00582 | Usp | Universal stress protein family | 2.40E-31 |
| WP\_004194465.1 | 15438 | 15669 | + | 76 | NO PFAM MATCH | - | - | - |

#### Results for WP\_208838502.1 [Burkholderia pseudomallei] back to top

#### previous - next

##### Architecture

1000 nucleotides

Link to nucleotide sequence  
  

| Accession | start | end | direction | length (aa) | Pfam/HMM | name | description | E-value |
| --- | --- | --- | --- | --- | --- | --- | --- | --- |
| WP\_004194465.1 | 59989 | 59758 | - | 76 | NO PFAM MATCH | - | - | - |
| WP\_004194548.1 | 60412 | 60880 | + | 155 | PF00582 | Usp | Universal stress protein family | 2.40E-31 |
| WP\_004529578.1 | 61029 | 61785 | + | 251 | PF13545 PF00027 PF00325 RREFamAux006 TIGR03896 | HTH\_Crp\_2 cNMP\_binding Crp RRE\_cNMP TIGR03896 | Crp-like helix-turn-helix domain Cyclic nucleotide-binding domain Bacterial regulatory proteins, crp family - cyc\_nuc\_ocin: bacteriocin-type transport-associated protein | 2.90E-20 6.00E-18 2.20E-14 2.60E-08 1.70E-05 |
| WP\_004525603.1 | 62763 | 61869 | - | 297 | TIGR03298 PF03466 PF00126 TIGR02424 TIGR03418 | TIGR03298 LysR\_substrate HTH\_1 TIGR02424 TIGR03418 | argP: transcriptional regulator, ArgP family LysR substrate binding domain Bacterial regulatory helix-turn-helix protein, lysR family TF\_pcaQ: pca operon transcription factor PcaQ chol\_sulf\_TF: putative choline sulfate-utilization transcription factor | 3.90E-120 6.00E-20 5.60E-15 4.10E-12 3.60E-11 |
| WP\_004546141.1 | 62870 | 63491 | + | 206 | TIGR00948 PF01810 TIGR00949 | TIGR00948 LysE TIGR00949 | 2a75: L-lysine exporter LysE type translocator 2A76: homoserine/Threonine efflux protein | 4.90E-53 1.00E-36 5.30E-06 |
| WP\_004545057.1 | 64652 | 63662 | - | 329 | TIGR04543 PF13649 PF13847 PF08242 TIGR03534 | TIGR04543 Methyltransf\_25 Methyltransf\_31 Methyltransf\_12 TIGR03534 | ketoArg\_3Met: 2-ketoarginine methyltransferase Methyltransferase domain Methyltransferase domain Methyltransferase domain RF\_mod\_PrmC: protein-(glutamine-N5) methyltransferase, release factor-specific | 1.10E-11 8.40E-09 2.80E-08 7.50E-07 1.40E-06 |
| WP\_004551133.1 | 66609 | 64665 | - | 647 | TIGR02026 TIGR03471 TIGR03975 TIGR04014 TIGR04013 | TIGR02026 TIGR03471 TIGR03975 TIGR04014 TIGR04013 | BchE: magnesium-protoporphyrin IX monomethyl ester anaerobic oxidative cyclase HpnJ: hopanoid biosynthesis associated radical SAM protein HpnJ rSAM\_ocin\_1: ribosomal peptide maturation radical SAM protein 1 B12\_SAM\_MJ\_0865: B12-binding domain/radical SAM domain protein, MJ\_0865 family B12\_SAM\_MJ\_1487: B12-binding domain/radical SAM domain protein, MJ\_1487 family | 7.40E-39 1.20E-31 1.30E-26 4.50E-25 1.00E-23 |
| WP\_208838501.1 | 67439 | 66605 | - | 277 | NO PFAM MATCH | - | - | - |
| WP\_208838502.1 | 68287 | 67435 | - | 283 | PF05114 PF01261 | DUF692 AP\_endonuc\_2 | Protein of unknown function (DUF692) Xylose isomerase-like TIM barrel | 1.10E-39 1.20E-06 |
| WP\_004201048.1 | 68661 | 68460 | - | 66 | NO PFAM MATCH | - | - | - |
| WP\_004539595.1 | 70217 | 68810 | - | 468 | PF00067 TIGR04538 TIGR04458 TIGR04515 | p450 TIGR04538 TIGR04458 TIGR04515 | Cytochrome P450 P450\_cycloAA\_1: cytochrome P450, cyclodipeptide synthase-associated CYP450\_TxtE: 4-nitrotryptophan synthase P450\_rel\_GT\_act: P450-derived glycosyltranferase activator | 1.90E-33 6.70E-16 5.70E-06 2.00E-05 |
| WP\_004196523.1 | 70624 | 71332 | + | 235 | PF03551 TIGR03433 | PadR TIGR03433 | Transcriptional regulator PadR-like family padR\_acidobact: transcriptional regulator, Acidobacterial, PadR-family | 2.00E-19 2.20E-12 |
| WP\_004194445.1 | 71371 | 72199 | + | 275 | PF04954 PF08021 | SIP FAD\_binding\_9 | Siderophore-interacting protein Siderophore-interacting FAD-binding domain | 2.90E-39 1.90E-37 |
| WP\_004529568.1 | 72771 | 72522 | - | 82 | NO PFAM MATCH | - | - | - |
| WP\_004196519.1 | 72964 | 73201 | + | 78 | NO PFAM MATCH | - | - | - |
| WP\_004525615.1 | 73820 | 73223 | - | 198 | PF01569 PF14378 | PAP2 PAP2\_3 | PAP2 superfamily PAP2 superfamily | 7.40E-18 7.90E-07 |
| WP\_004194452.1 | 74278 | 75019 | + | 246 | PF06314 | ADC | Acetoacetate decarboxylase (ADC) | 4.50E-79 |

#### Results for WP\_208205296.1 [Burkholderia pseudomallei] back to top

#### previous - next

##### Architecture

1000 nucleotides

Link to nucleotide sequence  
  

| Accession | start | end | direction | length (aa) | Pfam/HMM | name | description | E-value |
| --- | --- | --- | --- | --- | --- | --- | --- | --- |
| WP\_038760963.1 | 811 | 70 | - | 246 | PF06314 | ADC | Acetoacetate decarboxylase (ADC) | 4.50E-79 |
| WP\_208794800.1 | 1269 | 1866 | + | 198 | PF01569 PF14378 | PAP2 PAP2\_3 | PAP2 superfamily PAP2 superfamily | 3.30E-17 2.80E-06 |
| WP\_038733905.1 | 2125 | 1888 | - | 78 | NO PFAM MATCH | - | - | - |
| WP\_004194583.1 | 2318 | 2567 | + | 82 | NO PFAM MATCH | - | - | - |
| WP\_004532447.1 | 3718 | 2890 | - | 275 | PF04954 PF08021 | SIP FAD\_binding\_9 | Siderophore-interacting protein Siderophore-interacting FAD-binding domain | 2.50E-39 1.80E-37 |
| WP\_004196523.1 | 4465 | 3757 | - | 235 | PF03551 TIGR03433 | PadR TIGR03433 | Transcriptional regulator PadR-like family padR\_acidobact: transcriptional regulator, Acidobacterial, PadR-family | 2.00E-19 2.20E-12 |
| WP\_004539595.1 | 4872 | 6279 | + | 468 | PF00067 TIGR04538 TIGR04458 TIGR04515 | p450 TIGR04538 TIGR04458 TIGR04515 | Cytochrome P450 P450\_cycloAA\_1: cytochrome P450, cyclodipeptide synthase-associated CYP450\_TxtE: 4-nitrotryptophan synthase P450\_rel\_GT\_act: P450-derived glycosyltranferase activator | 1.90E-33 6.70E-16 5.70E-06 2.00E-05 |
| WP\_004201048.1 | 6428 | 6629 | + | 66 | NO PFAM MATCH | - | - | - |
| WP\_208205296.1 | 6802 | 7654 | + | 283 | PF05114 PF01261 | DUF692 AP\_endonuc\_2 | Protein of unknown function (DUF692) Xylose isomerase-like TIM barrel | 3.70E-39 3.90E-06 |
| WP\_208794798.1 | 7650 | 8484 | + | 277 | NO PFAM MATCH | - | - | - |
| WP\_041221553.1 | 8480 | 10424 | + | 647 | TIGR02026 TIGR03471 TIGR03975 TIGR04014 TIGR04013 | TIGR02026 TIGR03471 TIGR03975 TIGR04014 TIGR04013 | BchE: magnesium-protoporphyrin IX monomethyl ester anaerobic oxidative cyclase HpnJ: hopanoid biosynthesis associated radical SAM protein HpnJ rSAM\_ocin\_1: ribosomal peptide maturation radical SAM protein 1 B12\_SAM\_MJ\_0865: B12-binding domain/radical SAM domain protein, MJ\_0865 family B12\_SAM\_MJ\_1487: B12-binding domain/radical SAM domain protein, MJ\_1487 family | 4.20E-39 7.20E-32 3.20E-27 1.50E-25 3.00E-24 |
| WP\_004532676.1 | 10437 | 11427 | + | 329 | TIGR04543 PF13649 PF13847 PF08242 TIGR03534 | TIGR04543 Methyltransf\_25 Methyltransf\_31 Methyltransf\_12 TIGR03534 | ketoArg\_3Met: 2-ketoarginine methyltransferase Methyltransferase domain Methyltransferase domain Methyltransferase domain RF\_mod\_PrmC: protein-(glutamine-N5) methyltransferase, release factor-specific | 7.10E-12 8.20E-09 2.80E-08 7.60E-07 1.40E-06 |
| WP\_004194492.1 | 12219 | 11598 | - | 206 | TIGR00948 PF01810 TIGR00949 | TIGR00948 LysE TIGR00949 | 2a75: L-lysine exporter LysE type translocator 2A76: homoserine/Threonine efflux protein | 6.90E-53 1.40E-36 5.50E-06 |
| WP\_004194503.1 | 12326 | 13220 | + | 297 | TIGR03298 PF03466 PF00126 TIGR02424 TIGR03418 | TIGR03298 LysR\_substrate HTH\_1 TIGR02424 TIGR03418 | argP: transcriptional regulator, ArgP family LysR substrate binding domain Bacterial regulatory helix-turn-helix protein, lysR family TF\_pcaQ: pca operon transcription factor PcaQ chol\_sulf\_TF: putative choline sulfate-utilization transcription factor | 3.90E-120 4.80E-20 5.30E-15 5.90E-12 2.30E-11 |
| WP\_004529578.1 | 14069 | 13313 | - | 251 | PF13545 PF00027 PF00325 RREFamAux006 TIGR03896 | HTH\_Crp\_2 cNMP\_binding Crp RRE\_cNMP TIGR03896 | Crp-like helix-turn-helix domain Cyclic nucleotide-binding domain Bacterial regulatory proteins, crp family - cyc\_nuc\_ocin: bacteriocin-type transport-associated protein | 2.90E-20 6.00E-18 2.20E-14 2.60E-08 1.70E-05 |
| WP\_004194548.1 | 14686 | 14218 | - | 155 | PF00582 | Usp | Universal stress protein family | 2.40E-31 |
| WP\_004194465.1 | 15109 | 15340 | + | 76 | NO PFAM MATCH | - | - | - |

#### Results for WP\_041195603.1 [Burkholderia pseudomallei MSHR3335] back to top

#### previous - next

##### Architecture

1000 nucleotides

Link to nucleotide sequence  
  

| Accession | start | end | direction | length (aa) | Pfam/HMM | name | description | E-value |
| --- | --- | --- | --- | --- | --- | --- | --- | --- |
| WP\_004194465.1 | 1922125 | 1921894 | - | 76 | NO PFAM MATCH | - | - | - |
| WP\_004194548.1 | 1922548 | 1923016 | + | 155 | PF00582 | Usp | Universal stress protein family | 2.40E-31 |
| WP\_004529578.1 | 1923165 | 1923921 | + | 251 | PF13545 PF00027 PF00325 RREFamAux006 TIGR03896 | HTH\_Crp\_2 cNMP\_binding Crp RRE\_cNMP TIGR03896 | Crp-like helix-turn-helix domain Cyclic nucleotide-binding domain Bacterial regulatory proteins, crp family - cyc\_nuc\_ocin: bacteriocin-type transport-associated protein | 2.90E-20 6.00E-18 2.20E-14 2.60E-08 1.70E-05 |
| WP\_004525603.1 | 1924881 | 1923987 | - | 297 | TIGR03298 PF03466 PF00126 TIGR02424 TIGR03418 | TIGR03298 LysR\_substrate HTH\_1 TIGR02424 TIGR03418 | argP: transcriptional regulator, ArgP family LysR substrate binding domain Bacterial regulatory helix-turn-helix protein, lysR family TF\_pcaQ: pca operon transcription factor PcaQ chol\_sulf\_TF: putative choline sulfate-utilization transcription factor | 3.90E-120 6.00E-20 5.60E-15 4.10E-12 3.60E-11 |
| WP\_004194492.1 | 1924988 | 1925609 | + | 206 | TIGR00948 PF01810 TIGR00949 | TIGR00948 LysE TIGR00949 | 2a75: L-lysine exporter LysE type translocator 2A76: homoserine/Threonine efflux protein | 6.90E-53 1.40E-36 5.50E-06 |
| WP\_004545057.1 | 1926770 | 1925780 | - | 329 | TIGR04543 PF13649 PF13847 PF08242 TIGR03534 | TIGR04543 Methyltransf\_25 Methyltransf\_31 Methyltransf\_12 TIGR03534 | ketoArg\_3Met: 2-ketoarginine methyltransferase Methyltransferase domain Methyltransferase domain Methyltransferase domain RF\_mod\_PrmC: protein-(glutamine-N5) methyltransferase, release factor-specific | 1.10E-11 8.40E-09 2.80E-08 7.50E-07 1.40E-06 |
| WP\_041195775.1 | 1928727 | 1926783 | - | 647 | TIGR02026 TIGR03471 TIGR03975 TIGR04014 TIGR04013 | TIGR02026 TIGR03471 TIGR03975 TIGR04014 TIGR04013 | BchE: magnesium-protoporphyrin IX monomethyl ester anaerobic oxidative cyclase HpnJ: hopanoid biosynthesis associated radical SAM protein HpnJ rSAM\_ocin\_1: ribosomal peptide maturation radical SAM protein 1 B12\_SAM\_MJ\_0865: B12-binding domain/radical SAM domain protein, MJ\_0865 family B12\_SAM\_MJ\_1487: B12-binding domain/radical SAM domain protein, MJ\_1487 family | 3.00E-39 7.10E-32 2.00E-27 5.30E-26 1.70E-24 |
| WP\_041195602.1 | 1929557 | 1928723 | - | 277 | NO PFAM MATCH | - | - | - |
| WP\_041195603.1 | 1930405 | 1929553 | - | 283 | PF05114 PF01261 | DUF692 AP\_endonuc\_2 | Protein of unknown function (DUF692) Xylose isomerase-like TIM barrel | 1.10E-38 1.90E-06 |
| WP\_004201048.1 | 1930779 | 1930578 | - | 66 | NO PFAM MATCH | - | - | - |
| WP\_004539595.1 | 1932335 | 1930928 | - | 468 | PF00067 TIGR04538 TIGR04458 TIGR04515 | p450 TIGR04538 TIGR04458 TIGR04515 | Cytochrome P450 P450\_cycloAA\_1: cytochrome P450, cyclodipeptide synthase-associated CYP450\_TxtE: 4-nitrotryptophan synthase P450\_rel\_GT\_act: P450-derived glycosyltranferase activator | 1.90E-33 6.70E-16 5.70E-06 2.00E-05 |
| WP\_004196523.1 | 1932742 | 1933450 | + | 235 | PF03551 TIGR03433 | PadR TIGR03433 | Transcriptional regulator PadR-like family padR\_acidobact: transcriptional regulator, Acidobacterial, PadR-family | 2.00E-19 2.20E-12 |
| WP\_004194445.1 | 1933489 | 1934317 | + | 275 | PF04954 PF08021 | SIP FAD\_binding\_9 | Siderophore-interacting protein Siderophore-interacting FAD-binding domain | 2.90E-39 1.90E-37 |
| WP\_004196519.1 | 1935082 | 1935319 | + | 78 | NO PFAM MATCH | - | - | - |
| WP\_038731007.1 | 1935938 | 1935341 | - | 198 | PF01569 PF14378 | PAP2 PAP2\_3 | PAP2 superfamily PAP2 superfamily | 1.30E-17 4.00E-07 |
| WP\_004194452.1 | 1936396 | 1937137 | + | 246 | PF06314 | ADC | Acetoacetate decarboxylase (ADC) | 4.50E-79 |
| WP\_004194455.1 | 1937183 | 1937969 | + | 261 | TIGR01963 PF00106 PF13561 TIGR03971 TIGR02415 | TIGR01963 adh\_short adh\_short\_C2 TIGR03971 TIGR02415 | PHB\_DH: 3-hydroxybutyrate dehydrogenase short chain dehydrogenase Enoyl-(Acyl carrier protein) reductase SDR\_subfam\_1: SDR family mycofactocin-dependent oxidoreductase 23BDH: acetoin reductases | 4.30E-104 1.80E-52 2.40E-52 6.40E-52 8.90E-52 |

#### Results for WP\_316400961.1 [Burkholderia pseudomallei] back to top

#### previous - next

##### Architecture

1000 nucleotides

Link to nucleotide sequence  
  

| Accession | start | end | direction | length (aa) | Pfam/HMM | name | description | E-value |
| --- | --- | --- | --- | --- | --- | --- | --- | --- |
| WP\_122650933.1 | 101 | 0 | - | 34 | TIGR01963 TIGR01832 TIGR03971 PF00106 TIGR02632 | TIGR01963 TIGR01832 TIGR03971 adh\_short TIGR02632 | PHB\_DH: 3-hydroxybutyrate dehydrogenase kduD: 2-deoxy-D-gluconate 3-dehydrogenase SDR\_subfam\_1: SDR family mycofactocin-dependent oxidoreductase short chain dehydrogenase RhaD\_aldol-ADH: rhamnulose-1-phosphate aldolase/alcohol dehydrogenase | 1.90E-08 5.50E-08 1.80E-07 4.40E-07 1.20E-06 |
| WP\_004194452.1 | 888 | 147 | - | 246 | PF06314 | ADC | Acetoacetate decarboxylase (ADC) | 4.50E-79 |
| WP\_004525615.1 | 1346 | 1943 | + | 198 | PF01569 PF14378 | PAP2 PAP2\_3 | PAP2 superfamily PAP2 superfamily | 7.40E-18 7.90E-07 |
| WP\_004529566.1 | 2202 | 1965 | - | 78 | NO PFAM MATCH | - | - | - |
| WP\_004194583.1 | 2395 | 2644 | + | 82 | NO PFAM MATCH | - | - | - |
| WP\_004532447.1 | 3795 | 2967 | - | 275 | PF04954 PF08021 | SIP FAD\_binding\_9 | Siderophore-interacting protein Siderophore-interacting FAD-binding domain | 2.50E-39 1.80E-37 |
| WP\_004529569.1 | 4539 | 3834 | - | 234 | PF03551 TIGR03433 | PadR TIGR03433 | Transcriptional regulator PadR-like family padR\_acidobact: transcriptional regulator, Acidobacterial, PadR-family | 2.00E-19 2.10E-12 |
| WP\_004194516.1 | 4941 | 6348 | + | 468 | PF00067 TIGR04538 TIGR04458 TIGR04515 | p450 TIGR04538 TIGR04458 TIGR04515 | Cytochrome P450 P450\_cycloAA\_1: cytochrome P450, cyclodipeptide synthase-associated CYP450\_TxtE: 4-nitrotryptophan synthase P450\_rel\_GT\_act: P450-derived glycosyltranferase activator | 5.60E-34 3.30E-16 4.80E-06 2.50E-05 |
| WP\_004201048.1 | 6497 | 6698 | + | 66 | NO PFAM MATCH | - | - | - |
| WP\_316400961.1 | 6871 | 7723 | + | 283 | PF05114 PF01261 | DUF692 AP\_endonuc\_2 | Protein of unknown function (DUF692) Xylose isomerase-like TIM barrel | 1.20E-39 1.60E-06 |
| WP\_004551132.1 | 7719 | 8553 | + | 277 | NO PFAM MATCH | - | - | - |
| WP\_004551133.1 | 8549 | 10493 | + | 647 | TIGR02026 TIGR03471 TIGR03975 TIGR04014 TIGR04013 | TIGR02026 TIGR03471 TIGR03975 TIGR04014 TIGR04013 | BchE: magnesium-protoporphyrin IX monomethyl ester anaerobic oxidative cyclase HpnJ: hopanoid biosynthesis associated radical SAM protein HpnJ rSAM\_ocin\_1: ribosomal peptide maturation radical SAM protein 1 B12\_SAM\_MJ\_0865: B12-binding domain/radical SAM domain protein, MJ\_0865 family B12\_SAM\_MJ\_1487: B12-binding domain/radical SAM domain protein, MJ\_1487 family | 7.40E-39 1.20E-31 1.30E-26 4.50E-25 1.00E-23 |
| WP\_004551134.1 | 10506 | 11496 | + | 329 | TIGR04543 PF13649 PF13847 PF08242 TIGR03534 | TIGR04543 Methyltransf\_25 Methyltransf\_31 Methyltransf\_12 TIGR03534 | ketoArg\_3Met: 2-ketoarginine methyltransferase Methyltransferase domain Methyltransferase domain Methyltransferase domain RF\_mod\_PrmC: protein-(glutamine-N5) methyltransferase, release factor-specific | 7.10E-12 8.10E-09 2.80E-08 7.00E-07 1.40E-06 |
| WP\_004556447.1 | 12288 | 11667 | - | 206 | TIGR00948 PF01810 TIGR00949 | TIGR00948 LysE TIGR00949 | 2a75: L-lysine exporter LysE type translocator 2A76: homoserine/Threonine efflux protein | 2.40E-53 6.80E-37 1.10E-06 |
| WP\_004525603.1 | 12395 | 13289 | + | 297 | TIGR03298 PF03466 PF00126 TIGR02424 TIGR03418 | TIGR03298 LysR\_substrate HTH\_1 TIGR02424 TIGR03418 | argP: transcriptional regulator, ArgP family LysR substrate binding domain Bacterial regulatory helix-turn-helix protein, lysR family TF\_pcaQ: pca operon transcription factor PcaQ chol\_sulf\_TF: putative choline sulfate-utilization transcription factor | 3.90E-120 6.00E-20 5.60E-15 4.10E-12 3.60E-11 |
| WP\_004529578.1 | 14100 | 13344 | - | 251 | PF13545 PF00027 PF00325 RREFamAux006 TIGR03896 | HTH\_Crp\_2 cNMP\_binding Crp RRE\_cNMP TIGR03896 | Crp-like helix-turn-helix domain Cyclic nucleotide-binding domain Bacterial regulatory proteins, crp family - cyc\_nuc\_ocin: bacteriocin-type transport-associated protein | 2.90E-20 6.00E-18 2.20E-14 2.60E-08 1.70E-05 |
| WP\_004194548.1 | 14717 | 14249 | - | 155 | PF00582 | Usp | Universal stress protein family | 2.40E-31 |
| WP\_004194465.1 | 15140 | 15371 | + | 76 | NO PFAM MATCH | - | - | - |

#### Results for TPB63544.1 [Burkholderia pseudomallei] back to top

#### previous - next

##### Architecture

1000 nucleotides

Link to nucleotide sequence  
  

| Accession | start | end | direction | length (aa) | Pfam/HMM | name | description | E-value |
| --- | --- | --- | --- | --- | --- | --- | --- | --- |
| TPB63560.1 | 6507 | 6240 | - | 88 | NO PFAM MATCH | - | - | - |
| TPB63537.1 | 6894 | 7362 | + | 155 | PF00582 | Usp | Universal stress protein family | 2.40E-31 |
| TPB63538.1 | 7511 | 8267 | + | 251 | PF13545 PF00027 PF00325 RREFamAux006 TIGR03896 | HTH\_Crp\_2 cNMP\_binding Crp RRE\_cNMP TIGR03896 | Crp-like helix-turn-helix domain Cyclic nucleotide-binding domain Bacterial regulatory proteins, crp family - cyc\_nuc\_ocin: bacteriocin-type transport-associated protein | 3.00E-20 3.20E-17 2.30E-14 1.60E-07 3.70E-05 |
| TPB63539.1 | 9218 | 8324 | - | 297 | TIGR03298 PF03466 PF00126 TIGR02424 TIGR03418 | TIGR03298 LysR\_substrate HTH\_1 TIGR02424 TIGR03418 | argP: transcriptional regulator, ArgP family LysR substrate binding domain Bacterial regulatory helix-turn-helix protein, lysR family TF\_pcaQ: pca operon transcription factor PcaQ chol\_sulf\_TF: putative choline sulfate-utilization transcription factor | 1.20E-119 1.40E-19 5.60E-15 1.70E-11 3.10E-11 |
| TPB63540.1 | 9325 | 9946 | + | 206 | TIGR00948 PF01810 TIGR00949 | TIGR00948 LysE TIGR00949 | 2a75: L-lysine exporter LysE type translocator 2A76: homoserine/Threonine efflux protein | 6.90E-53 1.40E-36 5.50E-06 |
| TPB63541.1 | 11107 | 10117 | - | 329 | TIGR04543 PF13649 PF13847 PF08242 TIGR03534 | TIGR04543 Methyltransf\_25 Methyltransf\_31 Methyltransf\_12 TIGR03534 | ketoArg\_3Met: 2-ketoarginine methyltransferase Methyltransferase domain Methyltransferase domain Methyltransferase domain RF\_mod\_PrmC: protein-(glutamine-N5) methyltransferase, release factor-specific | 7.20E-12 8.20E-09 2.80E-08 7.60E-07 1.50E-06 |
| TPB63542.1 | 13076 | 11120 | - | 651 | TIGR02026 TIGR03471 TIGR03975 TIGR04014 TIGR04013 | TIGR02026 TIGR03471 TIGR03975 TIGR04014 TIGR04013 | BchE: magnesium-protoporphyrin IX monomethyl ester anaerobic oxidative cyclase HpnJ: hopanoid biosynthesis associated radical SAM protein HpnJ rSAM\_ocin\_1: ribosomal peptide maturation radical SAM protein 1 B12\_SAM\_MJ\_0865: B12-binding domain/radical SAM domain protein, MJ\_0865 family B12\_SAM\_MJ\_1487: B12-binding domain/radical SAM domain protein, MJ\_1487 family | 2.90E-39 2.00E-31 1.70E-27 5.20E-25 1.20E-23 |
| TPB63543.1 | 13894 | 13060 | - | 277 | NO PFAM MATCH | - | - | - |
| TPB63544.1 | 14742 | 13890 | - | 283 | PF05114 PF01261 | DUF692 AP\_endonuc\_2 | Protein of unknown function (DUF692) Xylose isomerase-like TIM barrel | 1.90E-39 1.50E-06 |
| TPB63561.1 | 14658 | 14925 | + | 88 | NO PFAM MATCH | - | - | - |
| TPB63545.1 | 15116 | 14915 | - | 66 | NO PFAM MATCH | - | - | - |
| TPB63546.1 | 16672 | 15265 | - | 468 | PF00067 TIGR04538 TIGR04458 TIGR04515 | p450 TIGR04538 TIGR04458 TIGR04515 | Cytochrome P450 P450\_cycloAA\_1: cytochrome P450, cyclodipeptide synthase-associated CYP450\_TxtE: 4-nitrotryptophan synthase P450\_rel\_GT\_act: P450-derived glycosyltranferase activator | 2.10E-34 2.20E-16 2.10E-06 1.20E-05 |
| TPB63562.1 | 16927 | 16744 | - | 60 | NO PFAM MATCH | - | - | - |
| TPB63547.1 | 17074 | 17800 | + | 241 | PF03551 TIGR03433 | PadR TIGR03433 | Transcriptional regulator PadR-like family padR\_acidobact: transcriptional regulator, Acidobacterial, PadR-family | 2.10E-19 2.30E-12 |
| TPB63548.1 | 17839 | 18667 | + | 275 | PF04954 PF08021 | SIP FAD\_binding\_9 | Siderophore-interacting protein Siderophore-interacting FAD-binding domain | 2.90E-39 1.90E-37 |
| TPB63549.1 | 18632 | 18893 | + | 86 | NO PFAM MATCH | - | - | - |
| TPB63550.1 | 19233 | 18984 | - | 82 | NO PFAM MATCH | - | - | - |

#### Results for WP\_038760955.1 [Burkholderia pseudomallei TSV 31] back to top

#### previous - next

##### Architecture

1000 nucleotides

Link to nucleotide sequence  
  

| Accession | start | end | direction | length (aa) | Pfam/HMM | name | description | E-value |
| --- | --- | --- | --- | --- | --- | --- | --- | --- |
| WP\_038760963.1 | 905774 | 905033 | - | 246 | PF06314 | ADC | Acetoacetate decarboxylase (ADC) | 4.50E-79 |
| WP\_004525615.1 | 906232 | 906829 | + | 198 | PF01569 PF14378 | PAP2 PAP2\_3 | PAP2 superfamily PAP2 superfamily | 7.40E-18 7.90E-07 |
| WP\_038733905.1 | 907088 | 906851 | - | 78 | NO PFAM MATCH | - | - | - |
| WP\_004194583.1 | 907281 | 907530 | + | 82 | NO PFAM MATCH | - | - | - |
| WP\_038760960.1 | 908681 | 907853 | - | 275 | PF04954 PF08021 | SIP FAD\_binding\_9 | Siderophore-interacting protein Siderophore-interacting FAD-binding domain | 3.10E-39 1.80E-37 |
| WP\_025988206.1 | 909446 | 908720 | - | 241 | PF03551 TIGR03433 | PadR TIGR03433 | Transcriptional regulator PadR-like family padR\_acidobact: transcriptional regulator, Acidobacterial, PadR-family | 2.10E-19 2.30E-12 |
| WP\_038760957.1 | 909853 | 911260 | + | 468 | PF00067 TIGR04538 TIGR04515 TIGR04458 | p450 TIGR04538 TIGR04515 TIGR04458 | Cytochrome P450 P450\_cycloAA\_1: cytochrome P450, cyclodipeptide synthase-associated P450\_rel\_GT\_act: P450-derived glycosyltranferase activator CYP450\_TxtE: 4-nitrotryptophan synthase | 2.10E-34 2.20E-16 6.20E-06 1.60E-05 |
| WP\_004201048.1 | 911409 | 911610 | + | 66 | NO PFAM MATCH | - | - | - |
| WP\_038760955.1 | 911783 | 912635 | + | 283 | PF05114 PF01261 | DUF692 AP\_endonuc\_2 | Protein of unknown function (DUF692) Xylose isomerase-like TIM barrel | 1.50E-39 1.40E-06 |
| WP\_038760953.1 | 912631 | 913453 | + | 273 | NO PFAM MATCH | - | - | - |
| WP\_038761491.1 | 913449 | 915393 | + | 647 | TIGR02026 TIGR03471 TIGR03975 TIGR04014 TIGR04013 | TIGR02026 TIGR03471 TIGR03975 TIGR04014 TIGR04013 | BchE: magnesium-protoporphyrin IX monomethyl ester anaerobic oxidative cyclase HpnJ: hopanoid biosynthesis associated radical SAM protein HpnJ rSAM\_ocin\_1: ribosomal peptide maturation radical SAM protein 1 B12\_SAM\_MJ\_0865: B12-binding domain/radical SAM domain protein, MJ\_0865 family B12\_SAM\_MJ\_1487: B12-binding domain/radical SAM domain protein, MJ\_1487 family | 8.20E-40 5.30E-32 2.90E-28 5.30E-26 1.60E-24 |
| WP\_004532676.1 | 915406 | 916396 | + | 329 | TIGR04543 PF13649 PF13847 PF08242 TIGR03534 | TIGR04543 Methyltransf\_25 Methyltransf\_31 Methyltransf\_12 TIGR03534 | ketoArg\_3Met: 2-ketoarginine methyltransferase Methyltransferase domain Methyltransferase domain Methyltransferase domain RF\_mod\_PrmC: protein-(glutamine-N5) methyltransferase, release factor-specific | 7.10E-12 8.20E-09 2.80E-08 7.60E-07 1.40E-06 |
| WP\_004194492.1 | 917188 | 916567 | - | 206 | TIGR00948 PF01810 TIGR00949 | TIGR00948 LysE TIGR00949 | 2a75: L-lysine exporter LysE type translocator 2A76: homoserine/Threonine efflux protein | 6.90E-53 1.40E-36 5.50E-06 |
| WP\_011852412.1 | 917295 | 918189 | + | 297 | TIGR03298 PF03466 PF00126 TIGR02424 TIGR03418 | TIGR03298 LysR\_substrate HTH\_1 TIGR02424 TIGR03418 | argP: transcriptional regulator, ArgP family LysR substrate binding domain Bacterial regulatory helix-turn-helix protein, lysR family TF\_pcaQ: pca operon transcription factor PcaQ chol\_sulf\_TF: putative choline sulfate-utilization transcription factor | 4.20E-120 4.10E-20 5.50E-15 6.10E-12 3.20E-11 |
| WP\_004529578.1 | 919018 | 918262 | - | 251 | PF13545 PF00027 PF00325 RREFamAux006 TIGR03896 | HTH\_Crp\_2 cNMP\_binding Crp RRE\_cNMP TIGR03896 | Crp-like helix-turn-helix domain Cyclic nucleotide-binding domain Bacterial regulatory proteins, crp family - cyc\_nuc\_ocin: bacteriocin-type transport-associated protein | 2.90E-20 6.00E-18 2.20E-14 2.60E-08 1.70E-05 |
| WP\_011852413.1 | 919635 | 919167 | - | 155 | PF00582 | Usp | Universal stress protein family | 2.50E-31 |
| WP\_004194465.1 | 920058 | 920289 | + | 76 | NO PFAM MATCH | - | - | - |

#### Results for MBF3713923.1 [Burkholderia pseudomallei] back to top

#### previous - next

##### Architecture

1000 nucleotides

Link to nucleotide sequence  
  

| Accession | start | end | direction | length (aa) | Pfam/HMM | name | description | E-value |
| --- | --- | --- | --- | --- | --- | --- | --- | --- |
| MBF3713915.1 | 58338 | 58107 | - | 76 | NO PFAM MATCH | - | - | - |
| MBF3713916.1 | 58761 | 59229 | + | 155 | PF00582 | Usp | Universal stress protein family | 2.40E-31 |
| MBF3713917.1 | 59378 | 60134 | + | 251 | PF13545 PF00027 PF00325 RREFamAux006 TIGR03896 | HTH\_Crp\_2 cNMP\_binding Crp RRE\_cNMP TIGR03896 | Crp-like helix-turn-helix domain Cyclic nucleotide-binding domain Bacterial regulatory proteins, crp family - cyc\_nuc\_ocin: bacteriocin-type transport-associated protein | 2.90E-20 6.00E-18 2.20E-14 2.60E-08 1.70E-05 |
| MBF3713918.1 | 61148 | 60254 | - | 297 | TIGR03298 PF03466 PF00126 TIGR02424 TIGR03418 | TIGR03298 LysR\_substrate HTH\_1 TIGR02424 TIGR03418 | argP: transcriptional regulator, ArgP family LysR substrate binding domain Bacterial regulatory helix-turn-helix protein, lysR family TF\_pcaQ: pca operon transcription factor PcaQ chol\_sulf\_TF: putative choline sulfate-utilization transcription factor | 3.90E-120 6.00E-20 5.60E-15 4.10E-12 3.60E-11 |
| MBF3713919.1 | 61255 | 61876 | + | 206 | TIGR00948 PF01810 TIGR00949 | TIGR00948 LysE TIGR00949 | 2a75: L-lysine exporter LysE type translocator 2A76: homoserine/Threonine efflux protein | 6.90E-53 1.40E-36 5.50E-06 |
| MBF3713920.1 | 63037 | 62047 | - | 329 | TIGR04543 PF13649 PF13847 PF08242 TIGR03534 | TIGR04543 Methyltransf\_25 Methyltransf\_31 Methyltransf\_12 TIGR03534 | ketoArg\_3Met: 2-ketoarginine methyltransferase Methyltransferase domain Methyltransferase domain Methyltransferase domain RF\_mod\_PrmC: protein-(glutamine-N5) methyltransferase, release factor-specific | 7.20E-12 8.20E-09 2.80E-08 7.60E-07 1.50E-06 |
| MBF3713921.1 | 64994 | 63050 | - | 647 | TIGR02026 TIGR03471 TIGR03975 TIGR04014 TIGR04013 | TIGR02026 TIGR03471 TIGR03975 TIGR04014 TIGR04013 | BchE: magnesium-protoporphyrin IX monomethyl ester anaerobic oxidative cyclase HpnJ: hopanoid biosynthesis associated radical SAM protein HpnJ rSAM\_ocin\_1: ribosomal peptide maturation radical SAM protein 1 B12\_SAM\_MJ\_0865: B12-binding domain/radical SAM domain protein, MJ\_0865 family B12\_SAM\_MJ\_1487: B12-binding domain/radical SAM domain protein, MJ\_1487 family | 4.80E-39 1.50E-31 1.40E-26 2.80E-25 9.70E-24 |
| MBF3713922.1 | 65812 | 64990 | - | 273 | NO PFAM MATCH | - | - | - |
| MBF3713923.1 | 66660 | 65808 | - | 283 | PF05114 PF01261 | DUF692 AP\_endonuc\_2 | Protein of unknown function (DUF692) Xylose isomerase-like TIM barrel | 8.20E-39 3.20E-06 |
| MBF3713924.1 | 67034 | 66833 | - | 66 | NO PFAM MATCH | - | - | - |
| MBF3713925.1 | 68590 | 67183 | - | 468 | PF00067 TIGR04538 TIGR04458 TIGR04515 | p450 TIGR04538 TIGR04458 TIGR04515 | Cytochrome P450 P450\_cycloAA\_1: cytochrome P450, cyclodipeptide synthase-associated CYP450\_TxtE: 4-nitrotryptophan synthase P450\_rel\_GT\_act: P450-derived glycosyltranferase activator | 5.60E-34 3.30E-16 4.80E-06 2.50E-05 |
| MBF3713926.1 | 68997 | 69705 | + | 235 | PF03551 TIGR03433 | PadR TIGR03433 | Transcriptional regulator PadR-like family padR\_acidobact: transcriptional regulator, Acidobacterial, PadR-family | 2.00E-19 2.20E-12 |
| MBF3713927.1 | 69744 | 70572 | + | 275 | PF04954 PF08021 | SIP FAD\_binding\_9 | Siderophore-interacting protein Siderophore-interacting FAD-binding domain | 2.90E-39 1.90E-37 |
| MBF3713928.1 | 71337 | 71574 | + | 78 | NO PFAM MATCH | - | - | - |
| MBF3713929.1 | 72193 | 71596 | - | 198 | PF01569 PF14378 | PAP2 PAP2\_3 | PAP2 superfamily PAP2 superfamily | 1.30E-17 4.00E-07 |
| MBF3713930.1 | 72651 | 73392 | + | 246 | PF06314 | ADC | Acetoacetate decarboxylase (ADC) | 4.50E-79 |
| MBF3713931.1 | 73438 | 73573 | + | 45 | TIGR01963 TIGR02632 TIGR02415 TIGR03971 PF00106 | TIGR01963 TIGR02632 TIGR02415 TIGR03971 adh\_short | PHB\_DH: 3-hydroxybutyrate dehydrogenase RhaD\_aldol-ADH: rhamnulose-1-phosphate aldolase/alcohol dehydrogenase 23BDH: acetoin reductases SDR\_subfam\_1: SDR family mycofactocin-dependent oxidoreductase short chain dehydrogenase | 2.20E-11 4.80E-10 8.30E-10 3.10E-09 4.00E-09 |

#### Results for WP\_004557993.1 [Burkholderia pseudomallei] back to top

#### previous - next

##### Architecture

1000 nucleotides

Link to nucleotide sequence  
  

| Accession | start | end | direction | length (aa) | Pfam/HMM | name | description | E-value |
| --- | --- | --- | --- | --- | --- | --- | --- | --- |
| WP\_004194465.1 | 2268266 | 2268035 | - | 76 | NO PFAM MATCH | - | - | - |
| WP\_004194548.1 | 2268689 | 2269157 | + | 155 | PF00582 | Usp | Universal stress protein family | 2.40E-31 |
| WP\_004529578.1 | 2269306 | 2270062 | + | 251 | PF13545 PF00027 PF00325 RREFamAux006 TIGR03896 | HTH\_Crp\_2 cNMP\_binding Crp RRE\_cNMP TIGR03896 | Crp-like helix-turn-helix domain Cyclic nucleotide-binding domain Bacterial regulatory proteins, crp family - cyc\_nuc\_ocin: bacteriocin-type transport-associated protein | 2.90E-20 6.00E-18 2.20E-14 2.60E-08 1.70E-05 |
| WP\_004525603.1 | 2271011 | 2270117 | - | 297 | TIGR03298 PF03466 PF00126 TIGR02424 TIGR03418 | TIGR03298 LysR\_substrate HTH\_1 TIGR02424 TIGR03418 | argP: transcriptional regulator, ArgP family LysR substrate binding domain Bacterial regulatory helix-turn-helix protein, lysR family TF\_pcaQ: pca operon transcription factor PcaQ chol\_sulf\_TF: putative choline sulfate-utilization transcription factor | 3.90E-120 6.00E-20 5.60E-15 4.10E-12 3.60E-11 |
| WP\_004194492.1 | 2271118 | 2271739 | + | 206 | TIGR00948 PF01810 TIGR00949 | TIGR00948 LysE TIGR00949 | 2a75: L-lysine exporter LysE type translocator 2A76: homoserine/Threonine efflux protein | 6.90E-53 1.40E-36 5.50E-06 |
| WP\_004525604.1 | 2272900 | 2271910 | - | 329 | TIGR04543 PF13649 PF13847 PF08242 TIGR03534 | TIGR04543 Methyltransf\_25 Methyltransf\_31 Methyltransf\_12 TIGR03534 | ketoArg\_3Met: 2-ketoarginine methyltransferase Methyltransferase domain Methyltransferase domain Methyltransferase domain RF\_mod\_PrmC: protein-(glutamine-N5) methyltransferase, release factor-specific | 2.00E-11 1.00E-08 2.90E-08 8.60E-07 1.40E-06 |
| WP\_004557994.1 | 2274857 | 2272913 | - | 647 | TIGR02026 TIGR03471 TIGR03975 TIGR04014 TIGR04013 | TIGR02026 TIGR03471 TIGR03975 TIGR04014 TIGR04013 | BchE: magnesium-protoporphyrin IX monomethyl ester anaerobic oxidative cyclase HpnJ: hopanoid biosynthesis associated radical SAM protein HpnJ rSAM\_ocin\_1: ribosomal peptide maturation radical SAM protein 1 B12\_SAM\_MJ\_0865: B12-binding domain/radical SAM domain protein, MJ\_0865 family B12\_SAM\_MJ\_1487: B12-binding domain/radical SAM domain protein, MJ\_1487 family | 6.80E-40 4.80E-32 2.90E-28 7.80E-26 1.60E-24 |
| WP\_004525606.1 | 2275687 | 2274853 | - | 277 | NO PFAM MATCH | - | - | - |
| WP\_004557993.1 | 2276535 | 2275683 | - | 283 | PF05114 PF01261 | DUF692 AP\_endonuc\_2 | Protein of unknown function (DUF692) Xylose isomerase-like TIM barrel | 6.80E-39 9.90E-07 |
| WP\_004525608.1 | 2276909 | 2276708 | - | 66 | NO PFAM MATCH | - | - | - |
| WP\_004194516.1 | 2278465 | 2277058 | - | 468 | PF00067 TIGR04538 TIGR04458 TIGR04515 | p450 TIGR04538 TIGR04458 TIGR04515 | Cytochrome P450 P450\_cycloAA\_1: cytochrome P450, cyclodipeptide synthase-associated CYP450\_TxtE: 4-nitrotryptophan synthase P450\_rel\_GT\_act: P450-derived glycosyltranferase activator | 5.60E-34 3.30E-16 4.80E-06 2.50E-05 |
| WP\_004196523.1 | 2278872 | 2279580 | + | 235 | PF03551 TIGR03433 | PadR TIGR03433 | Transcriptional regulator PadR-like family padR\_acidobact: transcriptional regulator, Acidobacterial, PadR-family | 2.00E-19 2.20E-12 |
| WP\_004557992.1 | 2279619 | 2280447 | + | 275 | PF04954 PF08021 | SIP FAD\_binding\_9 | Siderophore-interacting protein Siderophore-interacting FAD-binding domain | 2.90E-39 1.70E-36 |
| WP\_004194583.1 | 2281019 | 2280770 | - | 82 | NO PFAM MATCH | - | - | - |
| WP\_004529566.1 | 2281212 | 2281449 | + | 78 | NO PFAM MATCH | - | - | - |
| WP\_004525615.1 | 2282068 | 2281471 | - | 198 | PF01569 PF14378 | PAP2 PAP2\_3 | PAP2 superfamily PAP2 superfamily | 7.40E-18 7.90E-07 |
| WP\_004194452.1 | 2282526 | 2283267 | + | 246 | PF06314 | ADC | Acetoacetate decarboxylase (ADC) | 4.50E-79 |

#### Results for WP\_076830717.1 [Burkholderia pseudomallei] back to top

#### previous - next

##### Architecture

1000 nucleotides

Link to nucleotide sequence  
  

| Accession | start | end | direction | length (aa) | Pfam/HMM | name | description | E-value |
| --- | --- | --- | --- | --- | --- | --- | --- | --- |
| WP\_004194452.1 | 866 | 125 | - | 246 | PF06314 | ADC | Acetoacetate decarboxylase (ADC) | 4.50E-79 |
| WP\_004525615.1 | 1324 | 1921 | + | 198 | PF01569 PF14378 | PAP2 PAP2\_3 | PAP2 superfamily PAP2 superfamily | 7.40E-18 7.90E-07 |
| WP\_038780596.1 | 2180 | 1943 | - | 78 | NO PFAM MATCH | - | - | - |
| WP\_004194583.1 | 2373 | 2586 | + | 71 | INFERRED GENE | - | - | - |
| WP\_076830708.1 | 3710 | 2882 | - | 275 | PF04954 PF08021 | SIP FAD\_binding\_9 | Siderophore-interacting protein Siderophore-interacting FAD-binding domain | 6.10E-39 1.90E-37 |
| WP\_076830710.1 | 4454 | 3749 | - | 234 | PF03551 TIGR03433 | PadR TIGR03433 | Transcriptional regulator PadR-like family padR\_acidobact: transcriptional regulator, Acidobacterial, PadR-family | 2.00E-19 2.10E-12 |
| WP\_038774396.1 | 4856 | 6263 | + | 468 | PF00067 TIGR04538 TIGR04458 TIGR04515 | p450 TIGR04538 TIGR04458 TIGR04515 | Cytochrome P450 P450\_cycloAA\_1: cytochrome P450, cyclodipeptide synthase-associated CYP450\_TxtE: 4-nitrotryptophan synthase P450\_rel\_GT\_act: P450-derived glycosyltranferase activator | 6.70E-34 6.30E-16 4.80E-06 3.40E-05 |
| WP\_004201048.1 | 6412 | 6613 | + | 66 | NO PFAM MATCH | - | - | - |
| WP\_076830717.1 | 6786 | 7638 | + | 283 | PF05114 PF01261 | DUF692 AP\_endonuc\_2 | Protein of unknown function (DUF692) Xylose isomerase-like TIM barrel | 4.90E-39 5.70E-06 |
| WP\_038787684.1 | 7634 | 8468 | + | 277 | NO PFAM MATCH | - | - | - |
| WP\_045243618.1 | 8464 | 10408 | + | 647 | TIGR02026 TIGR03471 TIGR03975 TIGR04014 TIGR04013 | TIGR02026 TIGR03471 TIGR03975 TIGR04014 TIGR04013 | BchE: magnesium-protoporphyrin IX monomethyl ester anaerobic oxidative cyclase HpnJ: hopanoid biosynthesis associated radical SAM protein HpnJ rSAM\_ocin\_1: ribosomal peptide maturation radical SAM protein 1 B12\_SAM\_MJ\_0865: B12-binding domain/radical SAM domain protein, MJ\_0865 family B12\_SAM\_MJ\_1487: B12-binding domain/radical SAM domain protein, MJ\_1487 family | 5.90E-40 5.20E-32 1.80E-28 2.80E-26 9.60E-25 |
| WP\_076830723.1 | 10421 | 11411 | + | 329 | TIGR04543 PF13649 PF13847 PF08242 TIGR03534 | TIGR04543 Methyltransf\_25 Methyltransf\_31 Methyltransf\_12 TIGR03534 | ketoArg\_3Met: 2-ketoarginine methyltransferase Methyltransferase domain Methyltransferase domain Methyltransferase domain RF\_mod\_PrmC: protein-(glutamine-N5) methyltransferase, release factor-specific | 2.30E-11 7.80E-09 2.80E-08 7.80E-07 1.50E-06 |
| WP\_004194492.1 | 12203 | 11582 | - | 206 | TIGR00948 PF01810 TIGR00949 | TIGR00948 LysE TIGR00949 | 2a75: L-lysine exporter LysE type translocator 2A76: homoserine/Threonine efflux protein | 6.90E-53 1.40E-36 5.50E-06 |
| WP\_009922820.1 | 12310 | 13204 | + | 297 | TIGR03298 PF03466 PF00126 TIGR02424 TIGR03418 | TIGR03298 LysR\_substrate HTH\_1 TIGR02424 TIGR03418 | argP: transcriptional regulator, ArgP family LysR substrate binding domain Bacterial regulatory helix-turn-helix protein, lysR family TF\_pcaQ: pca operon transcription factor PcaQ chol\_sulf\_TF: putative choline sulfate-utilization transcription factor | 3.90E-120 6.00E-20 5.60E-15 4.10E-12 3.60E-11 |
| WP\_004529578.1 | 14025 | 13269 | - | 251 | PF13545 PF00027 PF00325 RREFamAux006 TIGR03896 | HTH\_Crp\_2 cNMP\_binding Crp RRE\_cNMP TIGR03896 | Crp-like helix-turn-helix domain Cyclic nucleotide-binding domain Bacterial regulatory proteins, crp family - cyc\_nuc\_ocin: bacteriocin-type transport-associated protein | 2.90E-20 6.00E-18 2.20E-14 2.60E-08 1.70E-05 |
| WP\_004194548.1 | 14642 | 14174 | - | 155 | PF00582 | Usp | Universal stress protein family | 2.40E-31 |
| WP\_004194465.1 | 15065 | 15296 | + | 76 | NO PFAM MATCH | - | - | - |

#### Results for CAG1017483.1 [Burkholderiaceae bacterium] back to top

#### previous - next

##### Architecture

1000 nucleotides

Link to nucleotide sequence  
  

| Accession | start | end | direction | length (aa) | Pfam/HMM | name | description | E-value |
| --- | --- | --- | --- | --- | --- | --- | --- | --- |
| CAG1017478.1 | 228 | 651 | + | 140 | PF00582 | Usp | Universal stress protein family | 3.70E-19 |
| CAG1017479.1 | 803 | 2681 | + | 625 | TIGR01241 TIGR01242 TIGR01243 PF01434 PF00004 | TIGR01241 TIGR01242 TIGR01243 Peptidase\_M41 AAA | FtsH\_fam: ATP-dependent metallopeptidase HflB 26Sp45: 26S proteasome subunit P45 family CDC48: AAA family ATPase, CDC48 subfamily Peptidase family M41 ATPase family associated with various cellular activities (AAA) | 4.50E-218 1.40E-79 1.20E-69 6.60E-64 8.10E-43 |
| CAG1017480.1 | 2792 | 3461 | + | 222 | PF13444 PF00583 | Acetyltransf\_5 Acetyltransf\_1 | Acetyltransferase (GNAT) domain Acetyltransferase (GNAT) family | 1.40E-04 2.90E-04 |
| CAG1017481.1 | 3548 | 4871 | + | 440 | PF00067 TIGR04538 TIGR04515 TIGR04458 | p450 TIGR04538 TIGR04515 TIGR04458 | Cytochrome P450 P450\_cycloAA\_1: cytochrome P450, cyclodipeptide synthase-associated P450\_rel\_GT\_act: P450-derived glycosyltranferase activator CYP450\_TxtE: 4-nitrotryptophan synthase | 6.30E-27 2.90E-15 4.50E-13 1.60E-09 |
| CAG1017482.1 | 4967 | 5174 | + | 68 | NO PFAM MATCH | - | - | - |
| CAG1017483.1 | 5238 | 6084 | + | 281 | PF05114 PF01261 | DUF692 AP\_endonuc\_2 | Protein of unknown function (DUF692) Xylose isomerase-like TIM barrel | 1.70E-44 5.80E-06 |
| CAG1017484.1 | 6080 | 6905 | + | 274 | NO PFAM MATCH | - | - | - |
| CAG1017485.1 | 6901 | 8863 | + | 653 | TIGR02026 TIGR03471 TIGR04014 TIGR03975 TIGR04479 | TIGR02026 TIGR03471 TIGR04014 TIGR03975 TIGR04479 | BchE: magnesium-protoporphyrin IX monomethyl ester anaerobic oxidative cyclase HpnJ: hopanoid biosynthesis associated radical SAM protein HpnJ B12\_SAM\_MJ\_0865: B12-binding domain/radical SAM domain protein, MJ\_0865 family rSAM\_ocin\_1: ribosomal peptide maturation radical SAM protein 1 bcpD\_PhpK\_rSAM: radical SAM P-methyltransferase, PhpK family | 3.70E-41 1.10E-36 9.40E-30 5.00E-29 3.70E-26 |
| CAG1017486.1 | 8859 | 9729 | + | 290 | TIGR04543 PF13649 TIGR00091 PF02390 TIGR02469 | TIGR04543 Methyltransf\_25 TIGR00091 Methyltransf\_4 TIGR02469 | ketoArg\_3Met: 2-ketoarginine methyltransferase Methyltransferase domain TIGR00091: tRNA (guanine-N(7)-)-methyltransferase Putative methyltransferase CbiT: precorrin-6Y C5,15-methyltransferase (decarboxylating), CbiT subunit | 1.30E-15 1.80E-06 1.20E-05 1.30E-04 3.20E-04 |

#### Results for VBN64128.1 [Burkholderia pseudomallei] back to top

#### previous - next

##### Architecture

1000 nucleotides

Link to nucleotide sequence  
  

| Accession | start | end | direction | length (aa) | Pfam/HMM | name | description | E-value |
| --- | --- | --- | --- | --- | --- | --- | --- | --- |
| VBN64078.1 | 1094 | 353 | - | 246 | PF06314 | ADC | Acetoacetate decarboxylase (ADC) | 4.50E-79 |
| VBN64087.1 | 1552 | 2149 | + | 198 | PF01569 PF14378 | PAP2 PAP2\_3 | PAP2 superfamily PAP2 superfamily | 1.30E-17 4.00E-07 |
| VBN64090.1 | 2408 | 2171 | - | 78 | NO PFAM MATCH | - | - | - |
| VBN64099.1 | 4001 | 3173 | - | 275 | PF04954 PF08021 | SIP FAD\_binding\_9 | Siderophore-interacting protein Siderophore-interacting FAD-binding domain | 2.90E-39 1.90E-37 |
| VBN64107.1 | 4748 | 4040 | - | 235 | PF03551 TIGR03433 | PadR TIGR03433 | Transcriptional regulator PadR-like family padR\_acidobact: transcriptional regulator, Acidobacterial, PadR-family | 1.40E-19 2.00E-12 |
| VBN64110.1 | 4894 | 5083 | + | 62 | NO PFAM MATCH | - | - | - |
| VBN64117.1 | 5155 | 6562 | + | 468 | PF00067 TIGR04538 TIGR04458 TIGR04515 | p450 TIGR04538 TIGR04458 TIGR04515 | Cytochrome P450 P450\_cycloAA\_1: cytochrome P450, cyclodipeptide synthase-associated CYP450\_TxtE: 4-nitrotryptophan synthase P450\_rel\_GT\_act: P450-derived glycosyltranferase activator | 7.00E-34 4.60E-16 2.40E-06 9.80E-06 |
| VBN64121.1 | 6711 | 6912 | + | 66 | NO PFAM MATCH | - | - | - |
| VBN64128.1 | 7085 | 8765 | + | 559 | PF05114 PF01261 | DUF692 AP\_endonuc\_2 | Protein of unknown function (DUF692) Xylose isomerase-like TIM barrel | 3.60E-38 2.90E-06 |
| VBN64134.1 | 8761 | 10705 | + | 647 | TIGR02026 TIGR03471 TIGR03975 TIGR04014 TIGR04013 | TIGR02026 TIGR03471 TIGR03975 TIGR04014 TIGR04013 | BchE: magnesium-protoporphyrin IX monomethyl ester anaerobic oxidative cyclase HpnJ: hopanoid biosynthesis associated radical SAM protein HpnJ rSAM\_ocin\_1: ribosomal peptide maturation radical SAM protein 1 B12\_SAM\_MJ\_0865: B12-binding domain/radical SAM domain protein, MJ\_0865 family B12\_SAM\_MJ\_1487: B12-binding domain/radical SAM domain protein, MJ\_1487 family | 5.60E-40 5.10E-32 1.70E-28 2.80E-26 9.60E-25 |
| VBN64142.1 | 10718 | 11708 | + | 329 | TIGR04543 PF13649 PF13847 PF08242 TIGR03534 | TIGR04543 Methyltransf\_25 Methyltransf\_31 Methyltransf\_12 TIGR03534 | ketoArg\_3Met: 2-ketoarginine methyltransferase Methyltransferase domain Methyltransferase domain Methyltransferase domain RF\_mod\_PrmC: protein-(glutamine-N5) methyltransferase, release factor-specific | 7.20E-12 8.20E-09 2.80E-08 7.60E-07 1.50E-06 |
| VBN64148.1 | 12500 | 11879 | - | 206 | TIGR00948 PF01810 TIGR00949 | TIGR00948 LysE TIGR00949 | 2a75: L-lysine exporter LysE type translocator 2A76: homoserine/Threonine efflux protein | 6.90E-53 1.40E-36 5.50E-06 |
| VBN64153.1 | 12607 | 13501 | + | 297 | TIGR03298 PF03466 PF00126 TIGR02424 TIGR03418 | TIGR03298 LysR\_substrate HTH\_1 TIGR02424 TIGR03418 | argP: transcriptional regulator, ArgP family LysR substrate binding domain Bacterial regulatory helix-turn-helix protein, lysR family TF\_pcaQ: pca operon transcription factor PcaQ chol\_sulf\_TF: putative choline sulfate-utilization transcription factor | 3.90E-120 6.00E-20 5.60E-15 4.10E-12 3.60E-11 |
| VBN64158.1 | 14323 | 13567 | - | 251 | PF13545 PF00027 PF00325 RREFamAux006 TIGR03896 | HTH\_Crp\_2 cNMP\_binding Crp RRE\_cNMP TIGR03896 | Crp-like helix-turn-helix domain Cyclic nucleotide-binding domain Bacterial regulatory proteins, crp family - cyc\_nuc\_ocin: bacteriocin-type transport-associated protein | 2.90E-20 6.00E-18 2.20E-14 2.60E-08 1.70E-05 |
| VBN64163.1 | 14940 | 14472 | - | 155 | PF00582 | Usp | Universal stress protein family | 2.40E-31 |
| VBN64166.1 | 15363 | 15594 | + | 76 | NO PFAM MATCH | - | - | - |
| VBN64172.1 | 16377 | 17958 | + | 526 | PF00171 TIGR02299 TIGR01780 TIGR03216 TIGR03240 | Aldedh TIGR02299 TIGR01780 TIGR03216 TIGR03240 | Aldehyde dehydrogenase family HpaE: 5-carboxymethyl-2-hydroxymuconate semialdehyde dehydrogenase SSADH: succinate-semialdehyde dehydrogenase OH\_muco\_semi\_DH: 2-hydroxymuconic semialdehyde dehydrogenase arg\_catab\_astD: succinylglutamate-semialdehyde dehydrogenase | 6.50E-46 6.80E-28 5.90E-24 2.60E-21 2.80E-20 |

#### Results for WP\_004194515.1 [Burkholderia mallei] back to top

#### previous - next

##### Architecture

1000 nucleotides

Link to nucleotide sequence  
  

| Accession | start | end | direction | length (aa) | Pfam/HMM | name | description | E-value |
| --- | --- | --- | --- | --- | --- | --- | --- | --- |
| WP\_004194465.1 | 2090508 | 2090277 | - | 76 | NO PFAM MATCH | - | - | - |
| WP\_004194548.1 | 2090931 | 2091399 | + | 155 | PF00582 | Usp | Universal stress protein family | 2.40E-31 |
| WP\_004194557.1 | 2091548 | 2092304 | + | 251 | PF13545 PF00027 PF00325 RREFamAux006 TIGR03896 | HTH\_Crp\_2 cNMP\_binding Crp RRE\_cNMP TIGR03896 | Crp-like helix-turn-helix domain Cyclic nucleotide-binding domain Bacterial regulatory proteins, crp family - cyc\_nuc\_ocin: bacteriocin-type transport-associated protein | 2.70E-20 6.00E-18 2.20E-14 1.10E-08 1.30E-05 |
| WP\_004194503.1 | 2093244 | 2092350 | - | 297 | TIGR03298 PF03466 PF00126 TIGR02424 TIGR03418 | TIGR03298 LysR\_substrate HTH\_1 TIGR02424 TIGR03418 | argP: transcriptional regulator, ArgP family LysR substrate binding domain Bacterial regulatory helix-turn-helix protein, lysR family TF\_pcaQ: pca operon transcription factor PcaQ chol\_sulf\_TF: putative choline sulfate-utilization transcription factor | 3.90E-120 4.80E-20 5.30E-15 5.90E-12 2.30E-11 |
| WP\_004194492.1 | 2093351 | 2093972 | + | 206 | TIGR00948 PF01810 TIGR00949 | TIGR00948 LysE TIGR00949 | 2a75: L-lysine exporter LysE type translocator 2A76: homoserine/Threonine efflux protein | 6.90E-53 1.40E-36 5.50E-06 |
| WP\_004194471.1 | 2095133 | 2094143 | - | 329 | TIGR04543 PF13649 PF13847 PF08242 TIGR03534 | TIGR04543 Methyltransf\_25 Methyltransf\_31 Methyltransf\_12 TIGR03534 | ketoArg\_3Met: 2-ketoarginine methyltransferase Methyltransferase domain Methyltransferase domain Methyltransferase domain RF\_mod\_PrmC: protein-(glutamine-N5) methyltransferase, release factor-specific | 3.70E-12 4.00E-08 1.70E-07 1.60E-06 4.70E-06 |
| WP\_004196531.1 | 2097090 | 2095146 | - | 647 | TIGR02026 TIGR03471 TIGR03975 TIGR04014 TIGR04013 | TIGR02026 TIGR03471 TIGR03975 TIGR04014 TIGR04013 | BchE: magnesium-protoporphyrin IX monomethyl ester anaerobic oxidative cyclase HpnJ: hopanoid biosynthesis associated radical SAM protein HpnJ rSAM\_ocin\_1: ribosomal peptide maturation radical SAM protein 1 B12\_SAM\_MJ\_0865: B12-binding domain/radical SAM domain protein, MJ\_0865 family B12\_SAM\_MJ\_1487: B12-binding domain/radical SAM domain protein, MJ\_1487 family | 4.50E-39 6.80E-32 3.10E-27 9.50E-26 3.10E-24 |
| WP\_004194512.1 | 2097920 | 2097086 | - | 277 | NO PFAM MATCH | - | - | - |
| WP\_004194515.1 | 2098708 | 2097916 | - | 263 | PF05114 PF01261 | DUF692 AP\_endonuc\_2 | Protein of unknown function (DUF692) Xylose isomerase-like TIM barrel | 1.80E-38 1.30E-06 |
| WP\_232515405.1 | 2098588 | 2098951 | + | 120 | NO PFAM MATCH | - | - | - |
| WP\_004201048.1 | 2099142 | 2098941 | - | 66 | NO PFAM MATCH | - | - | - |
| WP\_004194516.1 | 2100698 | 2099291 | - | 468 | PF00067 TIGR04538 TIGR04458 TIGR04515 | p450 TIGR04538 TIGR04458 TIGR04515 | Cytochrome P450 P450\_cycloAA\_1: cytochrome P450, cyclodipeptide synthase-associated CYP450\_TxtE: 4-nitrotryptophan synthase P450\_rel\_GT\_act: P450-derived glycosyltranferase activator | 5.60E-34 3.30E-16 4.80E-06 2.50E-05 |
| WP\_004196523.1 | 2101105 | 2101813 | + | 235 | PF03551 TIGR03433 | PadR TIGR03433 | Transcriptional regulator PadR-like family padR\_acidobact: transcriptional regulator, Acidobacterial, PadR-family | 2.00E-19 2.20E-12 |
| WP\_004194445.1 | 2101852 | 2102680 | + | 275 | PF04954 PF08021 | SIP FAD\_binding\_9 | Siderophore-interacting protein Siderophore-interacting FAD-binding domain | 2.90E-39 1.90E-37 |
| WP\_004194583.1 | 2103252 | 2103003 | - | 82 | NO PFAM MATCH | - | - | - |
| WP\_004196519.1 | 2103445 | 2103682 | + | 78 | NO PFAM MATCH | - | - | - |
| WP\_004194570.1 | 2104301 | 2103704 | - | 198 | PF01569 PF14378 | PAP2 PAP2\_3 | PAP2 superfamily PAP2 superfamily | 1.30E-17 4.00E-07 |

#### Results for EIP86375.1 [Burkholderia humptydooensis MSMB43] back to top

#### previous - next

##### Architecture

1000 nucleotides

Link to nucleotide sequence  
  

| Accession | start | end | direction | length (aa) | Pfam/HMM | name | description | E-value |
| --- | --- | --- | --- | --- | --- | --- | --- | --- |
| EIP86367.1 | 673336 | 673972 | + | 211 | PF01569 PF14378 | PAP2 PAP2\_3 | PAP2 superfamily PAP2 superfamily | 1.10E-17 5.10E-07 |
| EIP86368.1 | 674241 | 674004 | - | 78 | NO PFAM MATCH | - | - | - |
| EIP86369.1 | 674739 | 674556 | - | 60 | NO PFAM MATCH | - | - | - |
| EIP86370.1 | 675530 | 674738 | - | 263 | PF04954 PF08021 | SIP FAD\_binding\_9 | Siderophore-interacting protein Siderophore-interacting FAD-binding domain | 1.40E-39 7.30E-36 |
| EIP86371.1 | 676241 | 675608 | - | 210 | PF03551 TIGR03433 | PadR TIGR03433 | Transcriptional regulator PadR-like family padR\_acidobact: transcriptional regulator, Acidobacterial, PadR-family | 1.60E-19 1.70E-12 |
| EIP86372.1 | 681735 | 681396 | - | 112 | TIGR02212 TIGR02213 PF12704 | TIGR02212 TIGR02213 MacB\_PCD | lolCE: lipoprotein releasing system, transmembrane protein, LolC/E family lolE\_release: lipoprotein releasing system, transmembrane protein LolE MacB-like periplasmic core domain | 2.80E-23 1.20E-19 1.20E-06 |
| EIP86373.1 | 681975 | 683385 | + | 469 | PF00067 TIGR04538 TIGR04515 TIGR04458 | p450 TIGR04538 TIGR04515 TIGR04458 | Cytochrome P450 P450\_cycloAA\_1: cytochrome P450, cyclodipeptide synthase-associated P450\_rel\_GT\_act: P450-derived glycosyltranferase activator CYP450\_TxtE: 4-nitrotryptophan synthase | 9.20E-35 1.80E-14 1.70E-07 1.20E-06 |
| EIP86374.1 | 683535 | 683736 | + | 66 | NO PFAM MATCH | - | - | - |
| EIP86375.1 | 683907 | 684699 | + | 263 | PF05114 PF01261 | DUF692 AP\_endonuc\_2 | Protein of unknown function (DUF692) Xylose isomerase-like TIM barrel | 1.80E-39 9.20E-05 |
| EIP86376.1 | 684782 | 685529 | + | 248 | NO PFAM MATCH | - | - | - |
| EIP86377.1 | 685525 | 687469 | + | 647 | TIGR02026 TIGR03471 TIGR03975 TIGR04014 TIGR04013 | TIGR02026 TIGR03471 TIGR03975 TIGR04014 TIGR04013 | BchE: magnesium-protoporphyrin IX monomethyl ester anaerobic oxidative cyclase HpnJ: hopanoid biosynthesis associated radical SAM protein HpnJ rSAM\_ocin\_1: ribosomal peptide maturation radical SAM protein 1 B12\_SAM\_MJ\_0865: B12-binding domain/radical SAM domain protein, MJ\_0865 family B12\_SAM\_MJ\_1487: B12-binding domain/radical SAM domain protein, MJ\_1487 family | 2.20E-40 8.50E-35 6.80E-29 2.50E-26 5.40E-25 |
| EIP86378.1 | 687551 | 688472 | + | 306 | TIGR04543 PF13649 PF13847 PF08242 PF08241 | TIGR04543 Methyltransf\_25 Methyltransf\_31 Methyltransf\_12 Methyltransf\_11 | ketoArg\_3Met: 2-ketoarginine methyltransferase Methyltransferase domain Methyltransferase domain Methyltransferase domain Methyltransferase domain | 3.30E-12 1.00E-08 9.70E-08 1.40E-06 1.80E-05 |
| EIP86379.1 | 689324 | 688703 | - | 206 | TIGR00948 PF01810 TIGR00949 | TIGR00948 LysE TIGR00949 | 2a75: L-lysine exporter LysE type translocator 2A76: homoserine/Threonine efflux protein | 1.40E-53 2.40E-37 8.20E-07 |
| EIP86380.1 | 689431 | 690325 | + | 297 | TIGR03298 PF03466 PF00126 TIGR02424 TIGR03418 | TIGR03298 LysR\_substrate HTH\_1 TIGR02424 TIGR03418 | argP: transcriptional regulator, ArgP family LysR substrate binding domain Bacterial regulatory helix-turn-helix protein, lysR family TF\_pcaQ: pca operon transcription factor PcaQ chol\_sulf\_TF: putative choline sulfate-utilization transcription factor | 4.70E-121 1.10E-19 5.50E-15 1.20E-12 3.60E-11 |
| EIP86381.1 | 691026 | 690369 | - | 218 | PF13545 PF00027 PF00325 RREFamAux006 TIGR03697 | HTH\_Crp\_2 cNMP\_binding Crp RRE\_cNMP TIGR03697 | Crp-like helix-turn-helix domain Cyclic nucleotide-binding domain Bacterial regulatory proteins, crp family - NtcA\_cyano: global nitrogen regulator NtcA | 2.60E-20 3.30E-18 1.70E-14 2.40E-08 1.20E-05 |
| EIP86382.1 | 691742 | 691274 | - | 155 | PF00582 | Usp | Universal stress protein family | 1.40E-31 |
| EIP86383.1 | 692718 | 691983 | - | 244 | NO PFAM MATCH | - | - | - |

#### Results for MBL8238336.1 [Bryobacterales bacterium] back to top

#### previous - next

##### Architecture

1000 nucleotides

Link to nucleotide sequence  
  

| Accession | start | end | direction | length (aa) | Pfam/HMM | name | description | E-value |
| --- | --- | --- | --- | --- | --- | --- | --- | --- |
| MBL8238328.1 | 41435 | 42686 | + | 416 | PF07632 | DUF1593 | Protein of unknown function (DUF1593) | 9.00E-56 |
| MBL8238329.1 | 42751 | 46417 | + | 1221 | PF13620 PF13715 PF02369 PF17210 PF09134 | CarboxypepD\_reg CarbopepD\_reg\_2 Big\_1 SdrD\_B Invasin\_D3 | Carboxypeptidase regulatory-like domain CarboxypepD\_reg-like domain Bacterial Ig-like domain (group 1) SdrD B-like domain Invasin, domain 3 | 6.10E-20 1.90E-08 1.00E-04 4.60E-04 6.00E-04 |
| MBL8238330.1 | 46427 | 47645 | + | 405 | PF13378 TIGR02534 PF02746 TIGR01928 TIGR01927 | MR\_MLE\_C TIGR02534 MR\_MLE\_N TIGR01928 TIGR01927 | Enolase C-terminal domain-like mucon\_cyclo: muconate and chloromuconate cycloisomerases Mandelate racemase / muconate lactonizing enzyme, N-terminal domain menC\_lowGC/arch: o-succinylbenzoate synthase menC\_gamma/gm+: o-succinylbenzoate synthase | 1.20E-51 2.30E-22 6.80E-19 3.00E-07 2.80E-04 |
| MBL8238331.1 | 48092 | 47693 | - | 132 | PF04143 PF20398 | Sulf\_transp DUF6691 | Sulphur transport Family of unknown function (DUF6691) | 3.30E-08 1.40E-06 |
| MBL8238332.1 | 48675 | 48105 | - | 189 | PF04143 | Sulf\_transp | Sulphur transport | 8.60E-12 |
| MBL8238333.1 | 49711 | 48724 | - | 328 | TIGR04543 PF13649 PF08241 TIGR03534 PF13847 | TIGR04543 Methyltransf\_25 Methyltransf\_11 TIGR03534 Methyltransf\_31 | ketoArg\_3Met: 2-ketoarginine methyltransferase Methyltransferase domain Methyltransferase domain RF\_mod\_PrmC: protein-(glutamine-N5) methyltransferase, release factor-specific Methyltransferase domain | 7.80E-10 1.20E-08 1.90E-06 2.20E-05 9.60E-05 |
| MBL8238334.1 | 51657 | 49707 | - | 649 | TIGR02026 TIGR03471 TIGR04014 TIGR03975 TIGR04013 | TIGR02026 TIGR03471 TIGR04014 TIGR03975 TIGR04013 | BchE: magnesium-protoporphyrin IX monomethyl ester anaerobic oxidative cyclase HpnJ: hopanoid biosynthesis associated radical SAM protein HpnJ B12\_SAM\_MJ\_0865: B12-binding domain/radical SAM domain protein, MJ\_0865 family rSAM\_ocin\_1: ribosomal peptide maturation radical SAM protein 1 B12\_SAM\_MJ\_1487: B12-binding domain/radical SAM domain protein, MJ\_1487 family | 8.00E-37 1.70E-31 5.80E-29 7.70E-28 1.90E-27 |
| MBL8238335.1 | 52448 | 51653 | - | 264 | NO PFAM MATCH | - | - | - |
| MBL8238336.1 | 53293 | 52444 | - | 282 | PF05114 PF01261 | DUF692 AP\_endonuc\_2 | Protein of unknown function (DUF692) Xylose isomerase-like TIM barrel | 1.90E-45 3.90E-09 |
| MBL8238337.1 | 53512 | 53311 | - | 66 | NO PFAM MATCH | - | - | - |
| MBL8238338.1 | 54993 | 53670 | - | 440 | PF00067 TIGR04538 TIGR04515 TIGR04458 | p450 TIGR04538 TIGR04515 TIGR04458 | Cytochrome P450 P450\_cycloAA\_1: cytochrome P450, cyclodipeptide synthase-associated P450\_rel\_GT\_act: P450-derived glycosyltranferase activator CYP450\_TxtE: 4-nitrotryptophan synthase | 2.60E-32 1.20E-10 7.40E-08 4.40E-04 |

#### Results for KGU87554.1 [Burkholderia pseudomallei MSHR4372] back to top

#### previous - next

##### Architecture

1000 nucleotides

Link to nucleotide sequence  
  

| Accession | start | end | direction | length (aa) | Pfam/HMM | name | description | E-value |
| --- | --- | --- | --- | --- | --- | --- | --- | --- |
| KGU87679.1 | 2211873 | 2211642 | - | 76 | NO PFAM MATCH | - | - | - |
| KGU89001.1 | 2212296 | 2212764 | + | 155 | PF00582 | Usp | Universal stress protein family | 2.40E-31 |
| KGU88117.1 | 2212913 | 2213669 | + | 251 | PF13545 PF00027 PF00325 RREFamAux006 TIGR03896 | HTH\_Crp\_2 cNMP\_binding Crp RRE\_cNMP TIGR03896 | Crp-like helix-turn-helix domain Cyclic nucleotide-binding domain Bacterial regulatory proteins, crp family - cyc\_nuc\_ocin: bacteriocin-type transport-associated protein | 2.90E-20 6.00E-18 2.20E-14 2.60E-08 1.70E-05 |
| KGU87941.1 | 2214647 | 2213753 | - | 297 | TIGR03298 PF03466 PF00126 TIGR02424 TIGR03418 | TIGR03298 LysR\_substrate HTH\_1 TIGR02424 TIGR03418 | argP: transcriptional regulator, ArgP family LysR substrate binding domain Bacterial regulatory helix-turn-helix protein, lysR family TF\_pcaQ: pca operon transcription factor PcaQ chol\_sulf\_TF: putative choline sulfate-utilization transcription factor | 3.90E-120 6.00E-20 5.60E-15 4.10E-12 3.60E-11 |
| KGU88261.1 | 2214754 | 2215375 | + | 206 | TIGR00948 PF01810 TIGR00949 | TIGR00948 LysE TIGR00949 | 2a75: L-lysine exporter LysE type translocator 2A76: homoserine/Threonine efflux protein | 6.90E-53 1.40E-36 5.50E-06 |
| KGU88428.1 | 2216536 | 2215546 | - | 329 | TIGR04543 PF13649 PF13847 PF08242 TIGR03534 | TIGR04543 Methyltransf\_25 Methyltransf\_31 Methyltransf\_12 TIGR03534 | ketoArg\_3Met: 2-ketoarginine methyltransferase Methyltransferase domain Methyltransferase domain Methyltransferase domain RF\_mod\_PrmC: protein-(glutamine-N5) methyltransferase, release factor-specific | 1.10E-11 8.40E-09 2.80E-08 7.50E-07 1.40E-06 |
| KGU88305.1 | 2218493 | 2216549 | - | 647 | TIGR02026 TIGR03471 TIGR03975 TIGR04014 TIGR04013 | TIGR02026 TIGR03471 TIGR03975 TIGR04014 TIGR04013 | BchE: magnesium-protoporphyrin IX monomethyl ester anaerobic oxidative cyclase HpnJ: hopanoid biosynthesis associated radical SAM protein HpnJ rSAM\_ocin\_1: ribosomal peptide maturation radical SAM protein 1 B12\_SAM\_MJ\_0865: B12-binding domain/radical SAM domain protein, MJ\_0865 family B12\_SAM\_MJ\_1487: B12-binding domain/radical SAM domain protein, MJ\_1487 family | 3.70E-39 8.80E-32 3.30E-27 1.00E-25 3.00E-24 |
| KGU89134.1 | 2219311 | 2218489 | - | 273 | NO PFAM MATCH | - | - | - |
| KGU87554.1 | 2220099 | 2219307 | - | 263 | PF05114 PF01261 | DUF692 AP\_endonuc\_2 | Protein of unknown function (DUF692) Xylose isomerase-like TIM barrel | 2.40E-38 1.60E-06 |
| KGU89305.1 | 2220533 | 2220332 | - | 66 | NO PFAM MATCH | - | - | - |
| KGU87856.1 | 2222089 | 2220682 | - | 468 | PF00067 TIGR04538 TIGR04515 TIGR04458 | p450 TIGR04538 TIGR04515 TIGR04458 | Cytochrome P450 P450\_cycloAA\_1: cytochrome P450, cyclodipeptide synthase-associated P450\_rel\_GT\_act: P450-derived glycosyltranferase activator CYP450\_TxtE: 4-nitrotryptophan synthase | 5.60E-34 3.80E-16 4.50E-06 8.00E-06 |
| KGU89267.1 | 2222491 | 2223196 | + | 234 | PF03551 TIGR03433 | PadR TIGR03433 | Transcriptional regulator PadR-like family padR\_acidobact: transcriptional regulator, Acidobacterial, PadR-family | 2.00E-19 2.10E-12 |
| KGU89096.1 | 2223235 | 2224063 | + | 275 | PF04954 PF08021 | SIP FAD\_binding\_9 | Siderophore-interacting protein Siderophore-interacting FAD-binding domain | 4.50E-39 1.90E-37 |
| KGU89227.1 | 2224832 | 2225069 | + | 78 | NO PFAM MATCH | - | - | - |
| KGU88141.1 | 2225688 | 2225091 | - | 198 | PF01569 PF14378 | PAP2 PAP2\_3 | PAP2 superfamily PAP2 superfamily | 1.30E-17 4.00E-07 |
| KGU88801.1 | 2226146 | 2226887 | + | 246 | PF06314 | ADC | Acetoacetate decarboxylase (ADC) | 4.50E-79 |
| KGU88459.1 | 2226933 | 2227719 | + | 261 | TIGR01963 PF00106 PF13561 TIGR03971 TIGR02415 | TIGR01963 adh\_short adh\_short\_C2 TIGR03971 TIGR02415 | PHB\_DH: 3-hydroxybutyrate dehydrogenase short chain dehydrogenase Enoyl-(Acyl carrier protein) reductase SDR\_subfam\_1: SDR family mycofactocin-dependent oxidoreductase 23BDH: acetoin reductases | 4.30E-104 1.80E-52 2.40E-52 6.40E-52 8.90E-52 |

#### Results for MBN8214163.1 [Xanthomonadales bacterium] back to top

#### previous - next

##### Architecture

1000 nucleotides

Link to nucleotide sequence  
  

| Accession | start | end | direction | length (aa) | Pfam/HMM | name | description | E-value |
| --- | --- | --- | --- | --- | --- | --- | --- | --- |
| MBN8214160.1 | 621 | 0 | - | 207 | TIGR04543 PF13649 PF13489 TIGR04074 | TIGR04543 Methyltransf\_25 Methyltransf\_23 TIGR04074 | ketoArg\_3Met: 2-ketoarginine methyltransferase Methyltransferase domain Methyltransferase domain bacter\_Hen1: 3' terminal RNA ribose 2'-O-methyltransferase Hen1 | 7.80E-07 3.00E-05 9.60E-05 4.70E-04 |
| MBN8214161.1 | 2651 | 623 | - | 675 | TIGR03471 TIGR02026 TIGR04479 TIGR04014 TIGR03975 | TIGR03471 TIGR02026 TIGR04479 TIGR04014 TIGR03975 | HpnJ: hopanoid biosynthesis associated radical SAM protein HpnJ BchE: magnesium-protoporphyrin IX monomethyl ester anaerobic oxidative cyclase bcpD\_PhpK\_rSAM: radical SAM P-methyltransferase, PhpK family B12\_SAM\_MJ\_0865: B12-binding domain/radical SAM domain protein, MJ\_0865 family rSAM\_ocin\_1: ribosomal peptide maturation radical SAM protein 1 | 7.40E-38 5.80E-36 9.60E-27 3.30E-26 7.10E-25 |
| MBN8214162.1 | 3461 | 2651 | - | 269 | NO PFAM MATCH | - | - | - |
| MBN8214163.1 | 4342 | 3457 | - | 294 | PF05114 PF01261 | DUF692 AP\_endonuc\_2 | Protein of unknown function (DUF692) Xylose isomerase-like TIM barrel | 7.40E-42 5.00E-04 |
| MBN8214164.1 | 4753 | 4507 | - | 81 | TIGR03824 | TIGR03824 | FlgM\_jcvi: flagellar biosynthesis anti-sigma factor FlgM | 3.60E-05 |
| MBN8214165.1 | 6357 | 4956 | - | 466 | PF00067 TIGR04538 TIGR04515 TIGR04458 | p450 TIGR04538 TIGR04515 TIGR04458 | Cytochrome P450 P450\_cycloAA\_1: cytochrome P450, cyclodipeptide synthase-associated P450\_rel\_GT\_act: P450-derived glycosyltranferase activator CYP450\_TxtE: 4-nitrotryptophan synthase | 6.70E-28 9.70E-13 3.20E-09 3.20E-04 |
| MBN8214166.1 | 6657 | 7116 | + | 152 | NO PFAM MATCH | - | - | - |
| MBN8214167.1 | 8694 | 7206 | - | 495 | TIGR00836 PF00909 TIGR03644 PF19779 | TIGR00836 Ammonium\_transp TIGR03644 DUF6264 | amt: ammonium transporter Ammonium Transporter Family marine\_trans\_1: probable ammonium transporter, marine subtype Family of unknown function (DUF6264) | 8.70E-134 6.70E-122 1.40E-54 8.20E-04 |
| MBN8214168.1 | 9135 | 8796 | - | 112 | PF00543 | P-II | Nitrogen regulatory protein P-II | 2.20E-44 |
| MBN8214169.1 | 9770 | 9314 | - | 151 | PF00583 PF13508 PF13673 TIGR01575 PF13527 | Acetyltransf\_1 Acetyltransf\_7 Acetyltransf\_10 TIGR01575 Acetyltransf\_9 | Acetyltransferase (GNAT) family Acetyltransferase (GNAT) domain Acetyltransferase (GNAT) domain rimI: ribosomal-protein-alanine acetyltransferase Acetyltransferase (GNAT) domain | 3.80E-12 2.90E-08 4.40E-08 6.10E-07 6.90E-04 |
| MBN8214170.1 | 11253 | 9843 | - | 469 | TIGR00653 PF00120 TIGR03105 PF03951 | TIGR00653 Gln-synt\_C TIGR03105 Gln-synt\_N | GlnA: glutamine synthetase, type I Glutamine synthetase, catalytic domain gln\_synth\_III: glutamine synthetase, type III Glutamine synthetase, beta-Grasp domain | 1.50E-199 1.50E-137 3.30E-69 9.40E-31 |
| MBN8214171.1 | 11510 | 12425 | + | 304 | PF04116 | FA\_hydroxylase | Fatty acid hydroxylase | 3.20E-24 |

#### Results for MBX3740453.1 [Akkermansiaceae bacterium] back to top

#### previous - next

##### Architecture

1000 nucleotides

Link to nucleotide sequence  
  

| Accession | start | end | direction | length (aa) | Pfam/HMM | name | description | E-value |
| --- | --- | --- | --- | --- | --- | --- | --- | --- |
| MBX3740445.1 | 1316266 | 1317538 | + | 423 | PF04413 PF13579 | Glycos\_transf\_N Glyco\_trans\_4\_4 | 3-Deoxy-D-manno-octulosonic-acid transferase (kdotransferase) Glycosyl transferase 4-like domain | 2.10E-50 2.50E-04 |
| MBX3740446.1 | 1317687 | 1319376 | + | 562 | NO PFAM MATCH | - | - | - |
| MBX3740447.1 | 1320753 | 1319754 | - | 332 | NO PFAM MATCH | - | - | - |
| MBX3740448.1 | 1324622 | 1321376 | - | 1081 | NO PFAM MATCH | - | - | - |
| MBX3740449.1 | 1325706 | 1325100 | - | 201 | PF13340 PF01609 PF13612 | DUF4096 DDE\_Tnp\_1 DDE\_Tnp\_1\_3 | Putative transposase of IS4/5 family (DUF4096) Transposase DDE domain Transposase DDE domain | 1.70E-13 3.40E-12 6.80E-07 |
| MBX3740450.1 | 1327099 | 1326103 | - | 331 | TIGR04543 TIGR00091 PF13649 PF02390 PF13489 | TIGR04543 TIGR00091 Methyltransf\_25 Methyltransf\_4 Methyltransf\_23 | ketoArg\_3Met: 2-ketoarginine methyltransferase TIGR00091: tRNA (guanine-N(7)-)-methyltransferase Methyltransferase domain Putative methyltransferase Methyltransferase domain | 2.70E-14 3.10E-06 6.80E-06 2.10E-04 2.10E-04 |
| MBX3740451.1 | 1329083 | 1327091 | - | 663 | TIGR02026 TIGR03471 TIGR03975 TIGR04014 TIGR04479 | TIGR02026 TIGR03471 TIGR03975 TIGR04014 TIGR04479 | BchE: magnesium-protoporphyrin IX monomethyl ester anaerobic oxidative cyclase HpnJ: hopanoid biosynthesis associated radical SAM protein HpnJ rSAM\_ocin\_1: ribosomal peptide maturation radical SAM protein 1 B12\_SAM\_MJ\_0865: B12-binding domain/radical SAM domain protein, MJ\_0865 family bcpD\_PhpK\_rSAM: radical SAM P-methyltransferase, PhpK family | 1.60E-41 1.60E-37 2.30E-27 2.90E-26 5.70E-25 |
| MBX3740452.1 | 1329901 | 1329082 | - | 272 | NO PFAM MATCH | - | - | - |
| MBX3740453.1 | 1330746 | 1329897 | - | 282 | PF05114 PF01261 | DUF692 AP\_endonuc\_2 | Protein of unknown function (DUF692) Xylose isomerase-like TIM barrel | 1.70E-44 1.30E-07 |
| MBX3740454.1 | 1331100 | 1330878 | - | 73 | NO PFAM MATCH | - | - | - |
| MBX3740455.1 | 1332622 | 1331212 | - | 469 | PF00067 TIGR04538 TIGR04458 TIGR04515 | p450 TIGR04538 TIGR04458 TIGR04515 | Cytochrome P450 P450\_cycloAA\_1: cytochrome P450, cyclodipeptide synthase-associated CYP450\_TxtE: 4-nitrotryptophan synthase P450\_rel\_GT\_act: P450-derived glycosyltranferase activator | 2.40E-39 5.60E-13 2.40E-07 2.90E-07 |
| MBX3740456.1 | 1333217 | 1333460 | + | 80 | PF13586 PF01609 | DDE\_Tnp\_1\_2 DDE\_Tnp\_1 | Transposase DDE domain Transposase DDE domain | 2.60E-06 6.80E-05 |
| MBX3740457.1 | 1333675 | 1333474 | - | 66 | NO PFAM MATCH | - | - | - |
| MBX3740458.1 | 1334907 | 1333677 | - | 409 | TIGR03942 TIGR04083 TIGR04261 TIGR04163 TIGR03906 | TIGR03942 TIGR04083 TIGR04261 TIGR04163 TIGR03906 | sulfatase\_rSAM: anaerobic sulfatase maturase rSAM\_pep\_methan: putative peptide-modifying radical SAM enzyme, Mhun\_1560 family rSAM\_GlyRichRpt: radical SAM/SPASM domain protein, GRRM system rSAM\_cobopep: peptide-modifying radical SAM enzyme CbpB quino\_hemo\_SAM: quinohemoprotein amine dehydrogenase maturation protein | 1.70E-72 1.10E-49 2.10E-40 2.20E-35 1.00E-32 |

#### Results for WP\_263545016.1 [Tahibacter soli] back to top

#### previous - next

##### Architecture

1000 nucleotides

Link to nucleotide sequence  
  

| Accession | start | end | direction | length (aa) | Pfam/HMM | name | description | E-value |
| --- | --- | --- | --- | --- | --- | --- | --- | --- |
| WP\_263545024.1 | 49503 | 50535 | + | 343 | TIGR02825 PF16884 TIGR02824 PF00107 PF13602 | TIGR02825 ADH\_N\_2 TIGR02824 ADH\_zinc\_N ADH\_zinc\_N\_2 | B4\_12hDH: leukotriene B4 12-hydroxydehydrogenase/15-oxo-prostaglandin 13-reductase N-terminal domain of oxidoreductase quinone\_pig3: putative NAD(P)H quinone oxidoreductase, PIG3 family Zinc-binding dehydrogenase Zinc-binding dehydrogenase | 2.20E-70 1.00E-37 5.50E-19 4.10E-18 3.50E-13 |
| WP\_263545023.1 | 51243 | 50553 | - | 229 | NO PFAM MATCH | - | - | - |
| WP\_263545022.1 | 51567 | 52110 | + | 180 | NO PFAM MATCH | - | - | - |
| WP\_263545021.1 | 54436 | 52153 | - | 760 | TIGR00976 PF02129 PF08530 | TIGR00976 Peptidase\_S15 PepX\_C | /NonD: hydrolase CocE/NonD family protein X-Pro dipeptidyl-peptidase (S15 family) X-Pro dipeptidyl-peptidase C-terminal non-catalytic domain | 7.40E-53 1.90E-40 2.00E-13 |
| WP\_263545020.1 | 54564 | 55485 | + | 306 | TIGR03814 PF04960 | TIGR03814 Glutaminase | Gln\_ase: glutaminase A Glutaminase | 3.70E-119 2.90E-105 |
| WP\_263545019.1 | 57446 | 55478 | - | 655 | PF13229 TIGR04247 TIGR03808 | Beta\_helix TIGR04247 TIGR03808 | Right handed beta helix region NosD\_copper\_fam: nitrous oxide reductase family maturation protein NosD RR\_plus\_rpt\_1: twin-arg-translocated uncharacterized repeat protein | 7.80E-09 5.50E-06 7.90E-05 |
| WP\_263545018.1 | 57566 | 58922 | + | 451 | PF00067 TIGR04538 TIGR04515 TIGR04458 | p450 TIGR04538 TIGR04515 TIGR04458 | Cytochrome P450 P450\_cycloAA\_1: cytochrome P450, cyclodipeptide synthase-associated P450\_rel\_GT\_act: P450-derived glycosyltranferase activator CYP450\_TxtE: 4-nitrotryptophan synthase | 1.00E-24 5.20E-14 5.70E-10 2.20E-08 |
| WP\_263545017.1 | 59176 | 59395 | + | 72 | TIGR03824 | TIGR03824 | FlgM\_jcvi: flagellar biosynthesis anti-sigma factor FlgM | 8.10E-05 |
| WP\_263545016.1 | 59456 | 60302 | + | 281 | PF05114 PF01261 | DUF692 AP\_endonuc\_2 | Protein of unknown function (DUF692) Xylose isomerase-like TIM barrel | 5.90E-43 7.20E-05 |
| WP\_263545015.1 | 60298 | 61075 | + | 258 | NO PFAM MATCH | - | - | - |
| WP\_272841903.1 | 61130 | 63032 | + | 633 | TIGR03471 TIGR02026 TIGR04014 TIGR04013 TIGR04479 | TIGR03471 TIGR02026 TIGR04014 TIGR04013 TIGR04479 | HpnJ: hopanoid biosynthesis associated radical SAM protein HpnJ BchE: magnesium-protoporphyrin IX monomethyl ester anaerobic oxidative cyclase B12\_SAM\_MJ\_0865: B12-binding domain/radical SAM domain protein, MJ\_0865 family B12\_SAM\_MJ\_1487: B12-binding domain/radical SAM domain protein, MJ\_1487 family bcpD\_PhpK\_rSAM: radical SAM P-methyltransferase, PhpK family | 5.80E-40 1.20E-36 6.00E-30 5.80E-27 2.00E-26 |
| WP\_263545013.1 | 63028 | 64009 | + | 326 | TIGR04543 PF13649 TIGR03534 PF13847 PF08241 | TIGR04543 Methyltransf\_25 TIGR03534 Methyltransf\_31 Methyltransf\_11 | ketoArg\_3Met: 2-ketoarginine methyltransferase Methyltransferase domain RF\_mod\_PrmC: protein-(glutamine-N5) methyltransferase, release factor-specific Methyltransferase domain Methyltransferase domain | 2.40E-16 8.50E-10 3.20E-07 5.20E-07 2.90E-06 |
| WP\_263545012.1 | 65947 | 64039 | - | 635 | PF00486 TIGR02154 TIGR04510 PF13414 TIGR02917 | Trans\_reg\_C TIGR02154 TIGR04510 TPR\_11 TIGR02917 | Transcriptional regulatory protein, C terminal PhoB: phosphate regulon transcriptional regulatory protein PhoB mod\_pep\_cyc: putative peptide modification system cyclase TPR repeat PEP\_TPR\_lipo: putative PEP-CTERM system TPR-repeat lipoprotein | 2.40E-19 4.20E-12 8.10E-11 1.20E-08 1.50E-08 |
| WP\_263545011.1 | 66127 | 66910 | + | 260 | PF04116 | FA\_hydroxylase | Fatty acid hydroxylase | 3.70E-27 |
| WP\_263545010.1 | 67651 | 67270 | - | 126 | PF00903 PF18029 | Glyoxalase Glyoxalase\_6 | Glyoxalase/Bleomycin resistance protein/Dioxygenase superfamily Glyoxalase-like domain | 2.20E-06 1.30E-04 |
| WP\_263545009.1 | 68374 | 67807 | - | 188 | PF08818 | DUF1801 | Domain of unknown function (DU1801) | 2.30E-08 |
| WP\_263545008.1 | 68410 | 69967 | + | 518 | PF01494 TIGR01988 TIGR01984 TIGR01989 TIGR02032 | FAD\_binding\_3 TIGR01988 TIGR01984 TIGR01989 TIGR02032 | FAD binding domain Ubi-OHases: ubiquinone biosynthesis hydroxylase, UbiH/UbiF/VisC/COQ6 family UbiH: 2-polyprenyl-6-methoxyphenol 4-hydroxylase COQ6: ubiquinone biosynthesis monooxygenase COQ6 GG-red-SF: geranylgeranyl reductase family | 3.60E-80 3.40E-34 1.50E-21 2.60E-20 7.90E-18 |

#### Results for WP\_167928014.1 [Planosporangium thailandense] back to top

#### previous - next

##### Architecture

1000 nucleotides

Link to nucleotide sequence  
  

| Accession | start | end | direction | length (aa) | Pfam/HMM | name | description | E-value |
| --- | --- | --- | --- | --- | --- | --- | --- | --- |
| WP\_277349805.1 | 74674 | 75805 | + | 376 | PF01041 TIGR04427 TIGR03588 TIGR04181 TIGR02379 | DegT\_DnrJ\_EryC1 TIGR04427 TIGR03588 TIGR04181 TIGR02379 | DegT/DnrJ/EryC1/StrS aminotransferase family PLP\_DesI: dTDP-4-dehydro-6-deoxyglucose aminotransferase PseC: UDP-4-amino-4,6-dideoxy-N-acetyl-beta-L-altrosamine transaminase NHT\_00031: aminotransferase, LLPSF\_NHT\_00031 family ECA\_wecE: TDP-4-keto-6-deoxy-D-glucose transaminase | 3.50E-92 2.10E-88 4.30E-62 4.40E-51 9.40E-24 |
| WP\_277349806.1 | 75882 | 76320 | + | 145 | PF00293 TIGR02705 TIGR00586 | NUDIX TIGR02705 TIGR00586 | NUDIX domain nudix\_YtkD: nucleoside triphosphatase YtkD mutt: mutator mutT protein | 3.50E-18 1.60E-04 1.70E-04 |
| WP\_167928008.1 | 77267 | 76469 | - | 265 | PF13714 TIGR02317 TIGR02320 TIGR02319 | PEP\_mutase TIGR02317 TIGR02320 TIGR02319 | Phosphoenolpyruvate phosphomutase prpB: methylisocitrate lyase PEP\_mutase: phosphoenolpyruvate mutase CPEP\_Pphonmut: carboxyvinyl-carboxyphosphonate phosphorylmutase | 6.30E-67 7.70E-16 5.80E-09 5.50E-07 |
| WP\_240942976.1 | 77459 | 79580 | + | 706 | PF02897 PF00326 | Peptidase\_S9\_N Peptidase\_S9 | Prolyl oligopeptidase, N-terminal beta-propeller domain Prolyl oligopeptidase family | 1.80E-56 1.30E-41 |
| WP\_240942991.1 | 79692 | 80934 | + | 413 | PF00899 TIGR02356 TIGR02355 TIGR02354 RREFam017 | ThiF TIGR02356 TIGR02355 TIGR02354 Pantocin\_Microcin\_RRE | ThiF family adenyl\_thiF: thiazole biosynthesis adenylyltransferase ThiF moeB: molybdopterin synthase sulfurylase MoeB thiF\_fam2: thiamine biosynthesis protein ThiF RRE-containing E1-like protein in a pantocin/microcin cluster | 5.90E-36 3.60E-33 7.50E-23 3.30E-19 4.20E-13 |
| WP\_167928011.1 | 80977 | 82264 | + | 428 | TIGR00900 PF07690 PF05977 TIGR00880 | TIGR00900 MFS\_1 MFS\_3 TIGR00880 | 2A0121: H+ Antiporter protein Major Facilitator Superfamily Transmembrane secretion effector 2\_A\_01\_02: multidrug resistance protein | 1.30E-60 2.30E-25 5.90E-16 8.30E-14 |
| WP\_167928012.1 | 82383 | 83739 | + | 451 | PF00067 TIGR04538 TIGR04515 TIGR04458 | p450 TIGR04538 TIGR04515 TIGR04458 | Cytochrome P450 P450\_cycloAA\_1: cytochrome P450, cyclodipeptide synthase-associated P450\_rel\_GT\_act: P450-derived glycosyltranferase activator CYP450\_TxtE: 4-nitrotryptophan synthase | 1.00E-29 5.10E-12 2.00E-10 9.90E-05 |
| WP\_167928013.1 | 83816 | 83984 | + | 55 | NO PFAM MATCH | - | - | - |
| WP\_167928014.1 | 83999 | 84845 | + | 281 | PF05114 PF01261 | DUF692 AP\_endonuc\_2 | Protein of unknown function (DUF692) Xylose isomerase-like TIM barrel | 9.60E-46 6.10E-04 |
| WP\_167928015.1 | 84846 | 85683 | + | 278 | RREFam007 | Cyanobactin\_D\_RRE | RRE-containing D protein in a cyanobactin cluster | 2.60E-04 |
| WP\_205863251.1 | 85703 | 87626 | + | 640 | TIGR02026 TIGR03471 TIGR04014 TIGR04479 TIGR04013 | TIGR02026 TIGR03471 TIGR04014 TIGR04479 TIGR04013 | BchE: magnesium-protoporphyrin IX monomethyl ester anaerobic oxidative cyclase HpnJ: hopanoid biosynthesis associated radical SAM protein HpnJ B12\_SAM\_MJ\_0865: B12-binding domain/radical SAM domain protein, MJ\_0865 family bcpD\_PhpK\_rSAM: radical SAM P-methyltransferase, PhpK family B12\_SAM\_MJ\_1487: B12-binding domain/radical SAM domain protein, MJ\_1487 family | 6.90E-40 1.40E-36 4.70E-27 3.70E-26 6.40E-25 |
| WP\_167928016.1 | 88618 | 88276 | - | 113 | NO PFAM MATCH | - | - | - |
| WP\_167928017.1 | 89624 | 88751 | - | 290 | NO PFAM MATCH | - | - | - |

#### Results for WP\_114844792.1 [Dyella tabacisoli] back to top

#### previous - next

##### Architecture

1000 nucleotides

Link to nucleotide sequence  
  

| Accession | start | end | direction | length (aa) | Pfam/HMM | name | description | E-value |
| --- | --- | --- | --- | --- | --- | --- | --- | --- |
| WP\_114844784.1 | 10527 | 11676 | + | 382 | PF13378 TIGR02534 PF02746 TIGR01928 TIGR03247 | MR\_MLE\_C TIGR02534 MR\_MLE\_N TIGR01928 TIGR03247 | Enolase C-terminal domain-like mucon\_cyclo: muconate and chloromuconate cycloisomerases Mandelate racemase / muconate lactonizing enzyme, N-terminal domain menC\_lowGC/arch: o-succinylbenzoate synthase glucar-dehydr: glucarate dehydratase | 5.50E-64 2.50E-30 1.10E-26 5.00E-15 1.80E-05 |
| WP\_225427875.1 | 11678 | 12452 | + | 257 | PF13561 PF00106 TIGR01963 TIGR01830 TIGR03206 | adh\_short\_C2 adh\_short TIGR01963 TIGR01830 TIGR03206 | Enoyl-(Acyl carrier protein) reductase short chain dehydrogenase PHB\_DH: 3-hydroxybutyrate dehydrogenase 3oxo\_ACP\_reduc: 3-oxoacyl-[acyl-carrier-protein] reductase benzo\_BadH: 2-hydroxycyclohexanecarboxyl-CoA dehydrogenase | 2.40E-49 8.00E-39 9.80E-39 2.90E-35 4.70E-34 |
| WP\_114844786.1 | 12448 | 13345 | + | 298 | PF08450 | SGL | SMP-30/Gluconolactonase/LRE-like region | 5.70E-68 |
| WP\_114844787.1 | 13398 | 14868 | + | 489 | TIGR00879 PF00083 PF07690 TIGR00895 TIGR00898 | TIGR00879 Sugar\_tr MFS\_1 TIGR00895 TIGR00898 | SP: MFS transporter, sugar porter (SP) family Sugar (and other) transporter Major Facilitator Superfamily 2A0115: MFS transporter, aromatic acid:H+ symporter (AAHS) family 2A0119: cation transport protein | 5.80E-134 7.50E-130 8.70E-38 1.00E-32 4.30E-29 |
| WP\_225427876.1 | 16352 | 14960 | - | 463 | PF03572 TIGR00225 PF14684 TIGR03900 | Peptidase\_S41 TIGR00225 Tricorn\_C1 TIGR03900 | Peptidase family S41 prc: C-terminal processing peptidase Tricorn protease C1 domain prc\_long\_Delta: putative carboxyl-terminal-processing protease, deltaproteobacterial | 1.80E-24 2.80E-10 7.10E-09 1.20E-05 |
| WP\_114844789.1 | 17741 | 16739 | - | 333 | TIGR04543 PF13649 PF08241 PF13847 TIGR03534 | TIGR04543 Methyltransf\_25 Methyltransf\_11 Methyltransf\_31 TIGR03534 | ketoArg\_3Met: 2-ketoarginine methyltransferase Methyltransferase domain Methyltransferase domain Methyltransferase domain RF\_mod\_PrmC: protein-(glutamine-N5) methyltransferase, release factor-specific | 1.10E-12 7.80E-09 6.30E-05 2.70E-04 5.70E-04 |
| WP\_114844790.1 | 19717 | 17737 | - | 659 | TIGR02026 TIGR03471 TIGR04014 TIGR03975 TIGR04479 | TIGR02026 TIGR03471 TIGR04014 TIGR03975 TIGR04479 | BchE: magnesium-protoporphyrin IX monomethyl ester anaerobic oxidative cyclase HpnJ: hopanoid biosynthesis associated radical SAM protein HpnJ B12\_SAM\_MJ\_0865: B12-binding domain/radical SAM domain protein, MJ\_0865 family rSAM\_ocin\_1: ribosomal peptide maturation radical SAM protein 1 bcpD\_PhpK\_rSAM: radical SAM P-methyltransferase, PhpK family | 9.00E-38 1.00E-36 9.70E-28 3.00E-27 2.00E-24 |
| WP\_114844791.1 | 20523 | 19719 | - | 267 | RREFam006 | PqqD\_RRE | RRE-containing protein in a pyrroloquinoline cluster | 1.10E-04 |
| WP\_114844792.1 | 21371 | 20519 | - | 283 | PF05114 | DUF692 | Protein of unknown function (DUF692) | 9.60E-43 |
| WP\_225427877.1 | 21895 | 21631 | - | 87 | TIGR03824 | TIGR03824 | FlgM\_jcvi: flagellar biosynthesis anti-sigma factor FlgM | 1.10E-04 |
| WP\_114844793.1 | 23458 | 22021 | - | 478 | PF00067 TIGR04538 TIGR04515 TIGR04458 | p450 TIGR04538 TIGR04515 TIGR04458 | Cytochrome P450 P450\_cycloAA\_1: cytochrome P450, cyclodipeptide synthase-associated P450\_rel\_GT\_act: P450-derived glycosyltranferase activator CYP450\_TxtE: 4-nitrotryptophan synthase | 6.00E-30 3.20E-11 1.20E-08 3.60E-04 |
| WP\_114844794.1 | 23772 | 24657 | + | 294 | TIGR00950 PF00892 | TIGR00950 EamA | 2A78: carboxylate/amino acid/amine transporter EamA-like transporter family | 9.40E-23 1.00E-22 |
| WP\_114844795.1 | 26230 | 24817 | - | 470 | TIGR01667 PF13515 TIGR01666 | TIGR01667 FUSC\_2 TIGR01666 | YCCS\_YHJK: integral membrane protein, YccS/YhfK family Fusaric acid resistance protein-like YCCS: TIGR01666 family membrane protein | 1.90E-10 2.80E-09 2.10E-08 |
| WP\_162791399.1 | 26562 | 26268 | - | 97 | NO PFAM MATCH | - | - | - |
| WP\_225427878.1 | 26536 | 27712 | + | 391 | PF16499 PF17801 PF02065 PF17450 | Melibiase\_2 Melibiase\_C Melibiase Melibiase\_2\_C | Alpha galactosidase A Alpha galactosidase C-terminal beta sandwich domain Melibiase Alpha galactosidase A C-terminal beta sandwich domain | 2.70E-84 2.00E-20 8.00E-11 4.40E-07 |
| WP\_225427879.1 | 27784 | 28984 | + | 399 | PF16499 PF17801 PF02065 PF17450 | Melibiase\_2 Melibiase\_C Melibiase Melibiase\_2\_C | Alpha galactosidase A Alpha galactosidase C-terminal beta sandwich domain Melibiase Alpha galactosidase A C-terminal beta sandwich domain | 2.20E-81 2.00E-22 1.10E-07 5.50E-05 |
| WP\_114844798.1 | 31044 | 29016 | - | 675 | PF01204 PF03200 PF17389 | Trehalase Glyco\_hydro\_63 Bac\_rhamnosid6H | Trehalase Glycosyl hydrolase family 63 C-terminal domain Bacterial alpha-L-rhamnosidase 6 hairpin glycosidase domain | 9.80E-22 5.90E-10 1.30E-07 |

#### Results for NOT88279.1 [Lysobacter sp.] back to top

#### previous - next

##### Architecture

1000 nucleotides

Link to nucleotide sequence  
  

| Accession | start | end | direction | length (aa) | Pfam/HMM | name | description | E-value |
| --- | --- | --- | --- | --- | --- | --- | --- | --- |
| NOT88271.1 | 1995623 | 1993364 | - | 752 | PF01734 PF07244 TIGR03303 | Patatin POTRA TIGR03303 | Patatin-like phospholipase Surface antigen variable number repeat OM\_YaeT: outer membrane protein assembly complex, YaeT protein | 1.90E-30 5.90E-06 1.20E-05 |
| NOT88272.1 | 1995805 | 1997215 | + | 469 | TIGR00653 PF00120 TIGR03105 PF03951 | TIGR00653 Gln-synt\_C TIGR03105 Gln-synt\_N | GlnA: glutamine synthetase, type I Glutamine synthetase, catalytic domain gln\_synth\_III: glutamine synthetase, type III Glutamine synthetase, beta-Grasp domain | 3.30E-199 1.50E-137 1.50E-69 3.30E-30 |
| NOT88273.1 | 1997362 | 1997818 | + | 151 | PF00583 PF13673 PF13508 TIGR01575 PF13527 | Acetyltransf\_1 Acetyltransf\_10 Acetyltransf\_7 TIGR01575 Acetyltransf\_9 | Acetyltransferase (GNAT) family Acetyltransferase (GNAT) domain Acetyltransferase (GNAT) domain rimI: ribosomal-protein-alanine acetyltransferase Acetyltransferase (GNAT) domain | 6.60E-13 4.90E-09 1.50E-08 3.20E-07 6.30E-04 |
| NOT88274.1 | 1997997 | 1998336 | + | 112 | PF00543 | P-II | Nitrogen regulatory protein P-II | 2.20E-44 |
| NOT88275.1 | 1998357 | 1999905 | + | 515 | TIGR00836 PF00909 TIGR03644 | TIGR00836 Ammonium\_transp TIGR03644 | amt: ammonium transporter Ammonium Transporter Family marine\_trans\_1: probable ammonium transporter, marine subtype | 4.20E-134 4.40E-122 6.30E-54 |
| NOT88276.1 | 2000454 | 1999995 | - | 152 | NO PFAM MATCH | - | - | - |
| NOT88277.1 | 2000712 | 2002113 | + | 466 | PF00067 TIGR04538 TIGR04515 TIGR04458 | p450 TIGR04538 TIGR04515 TIGR04458 | Cytochrome P450 P450\_cycloAA\_1: cytochrome P450, cyclodipeptide synthase-associated P450\_rel\_GT\_act: P450-derived glycosyltranferase activator CYP450\_TxtE: 4-nitrotryptophan synthase | 1.20E-29 6.60E-13 2.30E-08 3.00E-04 |
| NOT88278.1 | 2002364 | 2002550 | + | 61 | NO PFAM MATCH | - | - | - |
| NOT88279.1 | 2002712 | 2003597 | + | 294 | PF05114 PF01261 | DUF692 AP\_endonuc\_2 | Protein of unknown function (DUF692) Xylose isomerase-like TIM barrel | 3.00E-41 4.90E-05 |
| NOT88280.1 | 2003593 | 2004403 | + | 269 | NO PFAM MATCH | - | - | - |
| NOT88281.1 | 2004403 | 2006431 | + | 675 | TIGR03471 TIGR02026 TIGR04479 TIGR04014 TIGR03975 | TIGR03471 TIGR02026 TIGR04479 TIGR04014 TIGR03975 | HpnJ: hopanoid biosynthesis associated radical SAM protein HpnJ BchE: magnesium-protoporphyrin IX monomethyl ester anaerobic oxidative cyclase bcpD\_PhpK\_rSAM: radical SAM P-methyltransferase, PhpK family B12\_SAM\_MJ\_0865: B12-binding domain/radical SAM domain protein, MJ\_0865 family rSAM\_ocin\_1: ribosomal peptide maturation radical SAM protein 1 | 1.20E-37 3.40E-36 1.80E-27 1.60E-26 7.30E-25 |
| NOT88282.1 | 2006493 | 2007477 | + | 327 | TIGR04543 PF13649 PF08241 PF13847 TIGR03534 | TIGR04543 Methyltransf\_25 Methyltransf\_11 Methyltransf\_31 TIGR03534 | ketoArg\_3Met: 2-ketoarginine methyltransferase Methyltransferase domain Methyltransferase domain Methyltransferase domain RF\_mod\_PrmC: protein-(glutamine-N5) methyltransferase, release factor-specific | 2.60E-10 1.50E-09 9.60E-07 1.40E-05 2.80E-05 |
| NOT88283.1 | 2009652 | 2007597 | - | 684 | PF00082 TIGR03921 TIGR03895 | Peptidase\_S8 TIGR03921 TIGR03895 | Subtilase family T7SS\_mycosin: type VII secretion-associated serine protease mycosin protease\_PatA: cyanobactin maturation protease, PatA/PatG family | 1.20E-45 1.10E-42 1.00E-09 |
| NOT88284.1 | 2010846 | 2009898 | - | 315 | NO PFAM MATCH | - | - | - |
| NOT88285.1 | 2013686 | 2011058 | - | 875 | PF00069 TIGR03903 PF07714 TIGR03724 | Pkinase TIGR03903 PK\_Tyr\_Ser-Thr TIGR03724 | Protein kinase domain TOMM\_kin\_cyc: TOMM system kinase/cyclase fusion protein Protein tyrosine and serine/threonine kinase arch\_bud32: Kae1-associated kinase Bud32 | 4.10E-43 1.20E-34 3.50E-29 1.30E-06 |
| NOT88286.1 | 2014266 | 2013693 | - | 190 | TIGR02999 PF07638 TIGR02937 TIGR02983 TIGR02985 | TIGR02999 Sigma70\_ECF TIGR02937 TIGR02983 TIGR02985 | Sig-70\_X6: RNA polymerase sigma factor, TIGR02999 family ECF sigma factor sigma70-ECF: RNA polymerase sigma factor, sigma-70 family SigE-fam\_strep: RNA polymerase sigma-70 factor, sigma-E family Sig70\_bacteroi1: RNA polymerase sigma-70 factor, Bacteroides expansion family 1 | 1.70E-42 1.80E-31 5.70E-22 3.80E-12 2.40E-11 |
| NOT88287.1 | 2016707 | 2014481 | - | 741 | PF10459 PF13365 TIGR02037 PF00089 | Peptidase\_S46 Trypsin\_2 TIGR02037 Trypsin | Peptidase S46 Trypsin-like peptidase domain degP\_htrA\_DO: peptidase Do Trypsin | 3.20E-212 2.10E-09 2.80E-04 5.80E-04 |

#### Results for MBW8851695.1 [Xanthomonadales bacterium] back to top

#### previous - next

##### Architecture

1000 nucleotides

Link to nucleotide sequence  
  

| Accession | start | end | direction | length (aa) | Pfam/HMM | name | description | E-value |
| --- | --- | --- | --- | --- | --- | --- | --- | --- |
| MBW8851687.1 | 10207 | 6868 | - | 1112 | TIGR00540 TIGR02917 | TIGR00540 TIGR02917 | TPR\_hemY\_coli: heme biosynthesis-associated TPR protein PEP\_TPR\_lipo: putative PEP-CTERM system TPR-repeat lipoprotein | 2.90E-05 5.80E-04 |
| MBW8851688.1 | 10524 | 11604 | + | 359 | PF13472 | Lipase\_GDSL\_2 | GDSL-like Lipase/Acylhydrolase family | 6.70E-07 |
| MBW8851689.1 | 11620 | 12304 | + | 227 | NO PFAM MATCH | - | - | - |
| MBW8851690.1 | 12307 | 14833 | + | 841 | TIGR00546 PF00795 PF20154 TIGR03381 | TIGR00546 CN\_hydrolase LNT\_N TIGR03381 | lnt: apolipoprotein N-acyltransferase Carbon-nitrogen hydrolase Apolipoprotein N-acyltransferase N-terminal domain agmatine\_aguB: N-carbamoylputrescine amidase | 1.50E-55 1.00E-15 1.40E-13 7.00E-05 |
| MBW8851691.1 | 15856 | 14839 | - | 338 | PF06674 | DUF1176 | Protein of unknown function (DUF1176) | 1.70E-89 |
| MBW8851692.1 | 17824 | 16030 | - | 597 | TIGR02456 PF00128 TIGR02403 TIGR02402 TIGR02401 | TIGR02456 Alpha-amylase TIGR02403 TIGR02402 TIGR02401 | treS\_nterm: trehalose synthase Alpha amylase, catalytic domain trehalose\_treC: alpha,alpha-phosphotrehalase trehalose\_TreZ: malto-oligosyltrehalose trehalohydrolase trehalose\_TreY: malto-oligosyltrehalose synthase | 6.50E-86 6.30E-78 4.60E-77 4.40E-30 1.50E-16 |
| MBW8851693.1 | 18034 | 19420 | + | 461 | PF00067 TIGR04538 TIGR04515 TIGR04458 | p450 TIGR04538 TIGR04515 TIGR04458 | Cytochrome P450 P450\_cycloAA\_1: cytochrome P450, cyclodipeptide synthase-associated P450\_rel\_GT\_act: P450-derived glycosyltranferase activator CYP450\_TxtE: 4-nitrotryptophan synthase | 5.20E-25 1.10E-10 9.40E-08 2.30E-04 |
| MBW8851694.1 | 19627 | 19897 | + | 89 | TIGR03824 | TIGR03824 | FlgM\_jcvi: flagellar biosynthesis anti-sigma factor FlgM | 1.20E-06 |
| MBW8851695.1 | 20065 | 20917 | + | 283 | PF05114 PF01261 | DUF692 AP\_endonuc\_2 | Protein of unknown function (DUF692) Xylose isomerase-like TIM barrel | 1.90E-40 4.20E-04 |
| MBW8851696.1 | 20913 | 21726 | + | 270 | RREFam006 | PqqD\_RRE | RRE-containing protein in a pyrroloquinoline cluster | 2.90E-04 |
| MBW8851697.1 | 21716 | 23687 | + | 656 | TIGR03471 TIGR02026 TIGR03975 TIGR04014 TIGR04385 | TIGR03471 TIGR02026 TIGR03975 TIGR04014 TIGR04385 | HpnJ: hopanoid biosynthesis associated radical SAM protein HpnJ BchE: magnesium-protoporphyrin IX monomethyl ester anaerobic oxidative cyclase rSAM\_ocin\_1: ribosomal peptide maturation radical SAM protein 1 B12\_SAM\_MJ\_0865: B12-binding domain/radical SAM domain protein, MJ\_0865 family B12\_rSAM\_cofa1: putative variant cofactor biosynthesis B12-binding domain/radical SAM domain protein 1 | 1.90E-36 9.80E-36 4.00E-31 1.40E-26 5.10E-24 |
| MBW8851698.1 | 23683 | 24694 | + | 336 | TIGR04543 PF13649 TIGR03534 PF13847 PF08242 | TIGR04543 Methyltransf\_25 TIGR03534 Methyltransf\_31 Methyltransf\_12 | ketoArg\_3Met: 2-ketoarginine methyltransferase Methyltransferase domain RF\_mod\_PrmC: protein-(glutamine-N5) methyltransferase, release factor-specific Methyltransferase domain Methyltransferase domain | 5.10E-13 2.90E-10 1.30E-08 2.70E-06 2.80E-06 |
| MBW8851699.1 | 24847 | 25276 | + | 142 | PF04143 | Sulf\_transp | Sulphur transport | 3.30E-09 |
| MBW8851700.1 | 25275 | 25722 | + | 148 | PF20398 | DUF6691 | Family of unknown function (DUF6691) | 3.30E-41 |
| MBW8851701.1 | 26733 | 25743 | - | 329 | TIGR01293 PF00248 | TIGR01293 Aldo\_ket\_red | Kv\_beta: voltage-dependent potassium channel beta subunit Aldo/keto reductase family | 5.20E-106 7.50E-66 |
| MBW8851702.1 | 31416 | 26808 | - | 1535 | TIGR01733 TIGR03443 PF00668 PF00501 TIGR01746 | TIGR01733 TIGR03443 Condensation AMP-binding TIGR01746 | AA-adenyl-dom: amino acid adenylation domain alpha\_am\_amid: L-aminoadipate-semialdehyde dehydrogenase Condensation domain AMP-binding enzyme Thioester-redct: thioester reductase domain | 1.00E-124 2.00E-112 4.50E-109 4.30E-94 1.90E-69 |
| MBW8851703.1 | 32039 | 33233 | + | 397 | TIGR02956 TIGR02966 TIGR02938 PF02518 PF00512 | TIGR02956 TIGR02966 TIGR02938 HATPase\_c HisKA | TMAO\_torS: TMAO reductase sytem sensor TorS phoR\_proteo: phosphate regulon sensor kinase PhoR nifL\_nitrog: nitrogen fixation negative regulator NifL Histidine kinase-, DNA gyrase B-, and HSP90-like ATPase His Kinase A (phospho-acceptor) domain | 4.20E-18 2.00E-17 1.70E-16 9.60E-16 4.00E-14 |

#### Results for WP\_162002680.1 [Streptomyces sp. CB01881] back to top

#### previous - next

##### Architecture

1000 nucleotides

Link to nucleotide sequence  
  

| Accession | start | end | direction | length (aa) | Pfam/HMM | name | description | E-value |
| --- | --- | --- | --- | --- | --- | --- | --- | --- |
| WP\_148647889.1 | 4254 | 4692 | + | 145 | NO PFAM MATCH | - | - | - |
| WP\_148647890.1 | 4934 | 5426 | + | 163 | PF16571 | FBP\_C | FBP C-terminal treble-clef zinc-finger | 3.50E-60 |
| WP\_148647891.1 | 5684 | 7934 | + | 749 | TIGR01666 TIGR01667 PF13515 | TIGR01666 TIGR01667 FUSC\_2 | YCCS: TIGR01666 family membrane protein YCCS\_YHJK: integral membrane protein, YccS/YhfK family Fusaric acid resistance protein-like | 1.10E-13 4.80E-12 2.40E-10 |
| WP\_148647892.1 | 8129 | 9338 | + | 402 | PF00291 TIGR00260 TIGR02991 TIGR01136 TIGR01127 | PALP TIGR00260 TIGR02991 TIGR01136 TIGR01127 | Pyridoxal-phosphate dependent enzyme thrC: threonine synthase ectoine\_eutB: ectoine utilization protein EutB cysKM: cysteine synthase ilvA\_1Cterm: threonine ammonia-lyase | 7.90E-50 4.50E-19 5.70E-10 4.10E-09 8.90E-09 |
| WP\_148647893.1 | 12466 | 9619 | - | 948 | PF05147 TIGR03897 PF00069 | LANC\_like TIGR03897 Pkinase | Lanthionine synthetase C-like protein lanti\_2\_LanM: type 2 lantibiotic biosynthesis protein LanM Protein kinase domain | 4.10E-43 6.80E-36 1.50E-16 |
| WP\_148647894.1 | 13136 | 12716 | - | 139 | NO PFAM MATCH | - | - | - |
| WP\_148647895.1 | 13375 | 14776 | + | 466 | PF00067 TIGR04538 | p450 TIGR04538 | Cytochrome P450 P450\_cycloAA\_1: cytochrome P450, cyclodipeptide synthase-associated | 9.30E-25 3.70E-05 |
| WP\_148647896.1 | 14963 | 15206 | + | 80 | NO PFAM MATCH | - | - | - |
| WP\_162002680.1 | 15277 | 16129 | + | 283 | PF05114 PF01261 | DUF692 AP\_endonuc\_2 | Protein of unknown function (DUF692) Xylose isomerase-like TIM barrel | 4.30E-45 2.20E-07 |
| WP\_148647898.1 | 16194 | 17145 | + | 316 | NO PFAM MATCH | - | - | - |
| WP\_148647899.1 | 17135 | 19100 | + | 654 | TIGR02026 TIGR03471 TIGR03975 TIGR04367 TIGR04013 | TIGR02026 TIGR03471 TIGR03975 TIGR04367 TIGR04013 | BchE: magnesium-protoporphyrin IX monomethyl ester anaerobic oxidative cyclase HpnJ: hopanoid biosynthesis associated radical SAM protein HpnJ rSAM\_ocin\_1: ribosomal peptide maturation radical SAM protein 1 HpnR\_B12\_rSAM: hopanoid C-3 methylase HpnR B12\_SAM\_MJ\_1487: B12-binding domain/radical SAM domain protein, MJ\_1487 family | 2.00E-36 4.40E-32 1.40E-30 1.10E-25 4.50E-25 |
| WP\_148647900.1 | 19092 | 20073 | + | 326 | TIGR04543 PF13649 PF13847 TIGR03534 PF08242 | TIGR04543 Methyltransf\_25 Methyltransf\_31 TIGR03534 Methyltransf\_12 | ketoArg\_3Met: 2-ketoarginine methyltransferase Methyltransferase domain Methyltransferase domain RF\_mod\_PrmC: protein-(glutamine-N5) methyltransferase, release factor-specific Methyltransferase domain | 2.40E-19 3.70E-10 1.70E-07 7.30E-06 7.90E-06 |
| WP\_148647901.1 | 20402 | 21146 | + | 247 | NO PFAM MATCH | - | - | - |
| WP\_148647902.1 | 21142 | 22435 | + | 430 | NO PFAM MATCH | - | - | - |
| WP\_148647903.1 | 22431 | 23208 | + | 258 | PF05050 TIGR01444 | Methyltransf\_21 TIGR01444 | Methyltransferase FkbM domain fkbM\_fam: methyltransferase, FkbM family | 4.20E-14 2.50E-07 |
| WP\_241681619.1 | 23669 | 23921 | + | 83 | PF06305 PF03169 | LapA\_dom OPT | Lipopolysaccharide assembly protein A domain OPT oligopeptide transporter protein | 4.40E-07 3.00E-04 |
| WP\_148647904.1 | 24062 | 24611 | + | 182 | PF04525 | LOR | LURP-one-related | 2.40E-16 |

#### Results for WP\_209416722.1 [Kitasatospora sp. RG8] back to top

#### previous - next

##### Architecture

1000 nucleotides

Link to nucleotide sequence  
  

| Accession | start | end | direction | length (aa) | Pfam/HMM | name | description | E-value |
| --- | --- | --- | --- | --- | --- | --- | --- | --- |
| WP\_245206130.1 | 62355 | 61881 | - | 157 | PF04525 | LOR | LURP-one-related | 2.00E-16 |
| WP\_245206132.1 | 62826 | 62574 | - | 83 | PF06305 PF03169 | LapA\_dom OPT | Lipopolysaccharide assembly protein A domain OPT oligopeptide transporter protein | 4.40E-07 5.10E-04 |
| WP\_209416716.1 | 64208 | 63446 | - | 253 | PF05050 TIGR01444 | Methyltransf\_21 TIGR01444 | Methyltransferase FkbM domain fkbM\_fam: methyltransferase, FkbM family | 6.10E-14 2.60E-07 |
| WP\_209416717.1 | 65497 | 64204 | - | 430 | NO PFAM MATCH | - | - | - |
| WP\_209416718.1 | 66261 | 65517 | - | 247 | NO PFAM MATCH | - | - | - |
| WP\_209416719.1 | 67577 | 66596 | - | 326 | TIGR04543 PF13649 PF13847 TIGR03534 PF08242 | TIGR04543 Methyltransf\_25 Methyltransf\_31 TIGR03534 Methyltransf\_12 | ketoArg\_3Met: 2-ketoarginine methyltransferase Methyltransferase domain Methyltransferase domain RF\_mod\_PrmC: protein-(glutamine-N5) methyltransferase, release factor-specific Methyltransferase domain | 2.40E-19 3.80E-10 9.70E-08 1.80E-06 3.30E-06 |
| WP\_209416720.1 | 69534 | 67569 | - | 654 | TIGR02026 TIGR03471 TIGR03975 TIGR04367 TIGR04013 | TIGR02026 TIGR03471 TIGR03975 TIGR04367 TIGR04013 | BchE: magnesium-protoporphyrin IX monomethyl ester anaerobic oxidative cyclase HpnJ: hopanoid biosynthesis associated radical SAM protein HpnJ rSAM\_ocin\_1: ribosomal peptide maturation radical SAM protein 1 HpnR\_B12\_rSAM: hopanoid C-3 methylase HpnR B12\_SAM\_MJ\_1487: B12-binding domain/radical SAM domain protein, MJ\_1487 family | 2.40E-36 1.10E-32 1.70E-30 5.00E-26 5.00E-25 |
| WP\_209416721.1 | 70484 | 69524 | - | 319 | NO PFAM MATCH | - | - | - |
| WP\_209416722.1 | 71434 | 70582 | - | 283 | PF05114 PF01261 | DUF692 AP\_endonuc\_2 | Protein of unknown function (DUF692) Xylose isomerase-like TIM barrel | 2.60E-45 6.50E-07 |
| WP\_209416723.1 | 71748 | 71505 | - | 80 | NO PFAM MATCH | - | - | - |
| WP\_209416724.1 | 73357 | 71956 | - | 466 | PF00067 TIGR04515 TIGR04538 | p450 TIGR04515 TIGR04538 | Cytochrome P450 P450\_rel\_GT\_act: P450-derived glycosyltranferase activator P450\_cycloAA\_1: cytochrome P450, cyclodipeptide synthase-associated | 5.20E-25 3.50E-05 6.70E-05 |
| WP\_209416725.1 | 74231 | 77066 | + | 944 | PF05147 TIGR03897 PF00069 PF14531 | LANC\_like TIGR03897 Pkinase Kinase-like | Lanthionine synthetase C-like protein lanti\_2\_LanM: type 2 lantibiotic biosynthesis protein LanM Protein kinase domain Kinase-like | 8.00E-42 6.10E-35 2.80E-16 2.00E-04 |
| WP\_209416726.1 | 78557 | 77348 | - | 402 | PF00291 TIGR00260 TIGR02991 TIGR01136 TIGR01127 | PALP TIGR00260 TIGR02991 TIGR01136 TIGR01127 | Pyridoxal-phosphate dependent enzyme thrC: threonine synthase ectoine\_eutB: ectoine utilization protein EutB cysKM: cysteine synthase ilvA\_1Cterm: threonine ammonia-lyase | 2.20E-51 2.10E-19 2.40E-10 3.60E-09 4.90E-09 |
| WP\_209416727.1 | 81109 | 78787 | - | 773 | TIGR01666 TIGR01667 PF13515 | TIGR01666 TIGR01667 FUSC\_2 | YCCS: TIGR01666 family membrane protein YCCS\_YHJK: integral membrane protein, YccS/YhfK family Fusaric acid resistance protein-like | 2.00E-14 4.30E-13 7.10E-11 |
| WP\_209416728.1 | 81900 | 81408 | - | 163 | PF16571 | FBP\_C | FBP C-terminal treble-clef zinc-finger | 6.60E-61 |
| WP\_209416729.1 | 82540 | 82102 | - | 145 | NO PFAM MATCH | - | - | - |
| WP\_209416730.1 | 85494 | 83163 | - | 776 | PF13424 PF00931 PF13174 | TPR\_12 NB-ARC TPR\_6 | Tetratricopeptide repeat NB-ARC domain Tetratricopeptide repeat | 6.10E-06 7.30E-05 6.50E-04 |

#### Results for TYC66662.1 [Streptomyces sp. CB01881] back to top

#### previous - next

##### Architecture

1000 nucleotides

Link to nucleotide sequence  
  

| Accession | start | end | direction | length (aa) | Pfam/HMM | name | description | E-value |
| --- | --- | --- | --- | --- | --- | --- | --- | --- |
| TYC66654.1 | 4254 | 4692 | + | 145 | NO PFAM MATCH | - | - | - |
| TYC66655.1 | 4934 | 5426 | + | 163 | PF16571 | FBP\_C | FBP C-terminal treble-clef zinc-finger | 3.50E-60 |
| TYC66656.1 | 5684 | 7934 | + | 749 | TIGR01666 TIGR01667 PF13515 | TIGR01666 TIGR01667 FUSC\_2 | YCCS: TIGR01666 family membrane protein YCCS\_YHJK: integral membrane protein, YccS/YhfK family Fusaric acid resistance protein-like | 1.10E-13 4.80E-12 2.40E-10 |
| TYC66657.1 | 8129 | 9338 | + | 402 | PF00291 TIGR00260 TIGR02991 TIGR01136 TIGR01127 | PALP TIGR00260 TIGR02991 TIGR01136 TIGR01127 | Pyridoxal-phosphate dependent enzyme thrC: threonine synthase ectoine\_eutB: ectoine utilization protein EutB cysKM: cysteine synthase ilvA\_1Cterm: threonine ammonia-lyase | 7.90E-50 4.50E-19 5.70E-10 4.10E-09 8.90E-09 |
| TYC66658.1 | 12466 | 9619 | - | 948 | PF05147 TIGR03897 PF00069 | LANC\_like TIGR03897 Pkinase | Lanthionine synthetase C-like protein lanti\_2\_LanM: type 2 lantibiotic biosynthesis protein LanM Protein kinase domain | 4.10E-43 6.80E-36 1.50E-16 |
| TYC66659.1 | 13136 | 12716 | - | 139 | NO PFAM MATCH | - | - | - |
| TYC66660.1 | 13375 | 14776 | + | 466 | PF00067 TIGR04538 | p450 TIGR04538 | Cytochrome P450 P450\_cycloAA\_1: cytochrome P450, cyclodipeptide synthase-associated | 9.30E-25 3.70E-05 |
| TYC66661.1 | 14963 | 15206 | + | 80 | NO PFAM MATCH | - | - | - |
| TYC66662.1 | 15211 | 16129 | + | 305 | PF05114 PF01261 | DUF692 AP\_endonuc\_2 | Protein of unknown function (DUF692) Xylose isomerase-like TIM barrel | 5.60E-45 2.70E-07 |
| TYC66663.1 | 16194 | 17145 | + | 316 | NO PFAM MATCH | - | - | - |
| TYC66664.1 | 17135 | 19100 | + | 654 | TIGR02026 TIGR03471 TIGR03975 TIGR04367 TIGR04013 | TIGR02026 TIGR03471 TIGR03975 TIGR04367 TIGR04013 | BchE: magnesium-protoporphyrin IX monomethyl ester anaerobic oxidative cyclase HpnJ: hopanoid biosynthesis associated radical SAM protein HpnJ rSAM\_ocin\_1: ribosomal peptide maturation radical SAM protein 1 HpnR\_B12\_rSAM: hopanoid C-3 methylase HpnR B12\_SAM\_MJ\_1487: B12-binding domain/radical SAM domain protein, MJ\_1487 family | 2.00E-36 4.40E-32 1.40E-30 1.10E-25 4.50E-25 |
| TYC66665.1 | 19092 | 20073 | + | 326 | TIGR04543 PF13649 PF13847 TIGR03534 PF08242 | TIGR04543 Methyltransf\_25 Methyltransf\_31 TIGR03534 Methyltransf\_12 | ketoArg\_3Met: 2-ketoarginine methyltransferase Methyltransferase domain Methyltransferase domain RF\_mod\_PrmC: protein-(glutamine-N5) methyltransferase, release factor-specific Methyltransferase domain | 2.40E-19 3.70E-10 1.70E-07 7.30E-06 7.90E-06 |
| TYC66666.1 | 20402 | 21146 | + | 247 | NO PFAM MATCH | - | - | - |
| TYC66667.1 | 21142 | 22435 | + | 430 | NO PFAM MATCH | - | - | - |
| TYC66668.1 | 22431 | 23208 | + | 258 | PF05050 TIGR01444 | Methyltransf\_21 TIGR01444 | Methyltransferase FkbM domain fkbM\_fam: methyltransferase, FkbM family | 4.20E-14 2.50E-07 |
| TYC66742.1 | 23648 | 23921 | + | 90 | PF06305 PF03169 | LapA\_dom OPT | Lipopolysaccharide assembly protein A domain OPT oligopeptide transporter protein | 4.60E-07 1.20E-04 |
| TYC66669.1 | 24062 | 24611 | + | 182 | PF04525 | LOR | LURP-one-related | 2.40E-16 |

#### Results for WP\_129271392.1 [Bradyrhizobium betae] back to top

#### previous - next

##### Architecture

1000 nucleotides

Link to nucleotide sequence  
  

| Accession | start | end | direction | length (aa) | Pfam/HMM | name | description | E-value |
| --- | --- | --- | --- | --- | --- | --- | --- | --- |
| WP\_164988063.1 | 197916 | 200058 | + | 713 | TIGR01398 PF00771 TIGR01399 | TIGR01398 FHIPEP TIGR01399 | FlhA: flagellar biosynthesis protein FlhA FHIPEP family hrcV: type III secretion protein, HrcV family | 1.80E-273 1.00E-256 2.90E-178 |
| WP\_027532768.1 | 200352 | 200553 | + | 66 | NO PFAM MATCH | - | - | - |
| WP\_129271386.1 | 206204 | 201599 | - | 1534 | PF00196 TIGR03020 PF08281 PF00072 TIGR02937 | GerE TIGR03020 Sigma70\_r4\_2 Response\_reg TIGR02937 | Bacterial regulatory proteins, luxR family EpsA: transcriptional regulator EpsA Sigma-70, region 4 Response regulator receiver domain sigma70-ECF: RNA polymerase sigma factor, sigma-70 family | 3.20E-20 4.30E-14 1.30E-11 1.30E-09 9.30E-09 |
| WP\_129271387.1 | 206357 | 206699 | + | 113 | PF12680 PF14534 PF13474 TIGR02246 | SnoaL\_2 DUF4440 SnoaL\_3 TIGR02246 | SnoaL-like domain Domain of unknown function (DUF4440) SnoaL-like domain TIGR02246: conserved hypothetical protein | 8.70E-06 2.30E-04 5.50E-04 5.60E-04 |
| WP\_129271388.1 | 207999 | 206775 | - | 407 | PF07992 TIGR03169 TIGR04369 TIGR02374 TIGR03385 | Pyr\_redox\_2 TIGR03169 TIGR04369 TIGR02374 TIGR03385 | Pyridine nucleotide-disulphide oxidoreductase Nterm\_to\_SelD: pyridine nucleotide-disulfide oxidoreductase family protein fusion\_not\_SelD: oxidoreductase/SelD-related fusion protein nitri\_red\_nirB: nitrite reductase [NAD(P)H], large subunit CoA\_CoA\_reduc: CoA-disulfide reductase | 7.00E-45 2.80E-28 7.10E-14 7.90E-12 2.80E-11 |
| WP\_129271389.1 | 209076 | 208116 | - | 319 | PF12833 PF14525 PF00165 TIGR04094 TIGR02297 | HTH\_18 AraC\_binding\_2 HTH\_AraC TIGR04094 TIGR02297 | Helix-turn-helix domain AraC-binding-like domain Bacterial regulatory helix-turn-helix proteins, AraC family adjacent\_YSIRK: YSIRK-targeted surface antigen transcriptional regulator HpaA: 4-hydroxyphenylacetate catabolism regulatory protein HpaA | 1.70E-21 3.20E-17 8.60E-14 1.20E-06 3.90E-05 |
| WP\_129271390.1 | 209684 | 210770 | + | 361 | PF00067 TIGR04538 TIGR04458 | p450 TIGR04538 TIGR04458 | Cytochrome P450 P450\_cycloAA\_1: cytochrome P450, cyclodipeptide synthase-associated CYP450\_TxtE: 4-nitrotryptophan synthase | 4.50E-32 1.30E-09 9.30E-05 |
| WP\_129271391.1 | 210888 | 211068 | + | 59 | NO PFAM MATCH | - | - | - |
| WP\_129271392.1 | 211139 | 212012 | + | 290 | PF05114 PF01261 | DUF692 AP\_endonuc\_2 | Protein of unknown function (DUF692) Xylose isomerase-like TIM barrel | 2.70E-36 4.40E-04 |
| WP\_129271393.1 | 212004 | 212871 | + | 288 | RREFam006 | PqqD\_RRE | RRE-containing protein in a pyrroloquinoline cluster | 1.50E-04 |
| WP\_129271394.1 | 212848 | 214825 | + | 658 | TIGR02026 TIGR03471 TIGR03975 TIGR04014 TIGR04479 | TIGR02026 TIGR03471 TIGR03975 TIGR04014 TIGR04479 | BchE: magnesium-protoporphyrin IX monomethyl ester anaerobic oxidative cyclase HpnJ: hopanoid biosynthesis associated radical SAM protein HpnJ rSAM\_ocin\_1: ribosomal peptide maturation radical SAM protein 1 B12\_SAM\_MJ\_0865: B12-binding domain/radical SAM domain protein, MJ\_0865 family bcpD\_PhpK\_rSAM: radical SAM P-methyltransferase, PhpK family | 6.10E-39 2.30E-37 8.00E-29 1.20E-28 1.80E-26 |
| WP\_129271395.1 | 214821 | 215817 | + | 331 | TIGR04543 PF13649 TIGR04074 TIGR02469 TIGR03704 | TIGR04543 Methyltransf\_25 TIGR04074 TIGR02469 TIGR03704 | ketoArg\_3Met: 2-ketoarginine methyltransferase Methyltransferase domain bacter\_Hen1: 3' terminal RNA ribose 2'-O-methyltransferase Hen1 CbiT: precorrin-6Y C5,15-methyltransferase (decarboxylating), CbiT subunit PrmC\_rel\_meth: putative protein-(glutamine-N5) methyltransferase, unknown substrate-specific | 4.30E-10 1.60E-07 1.70E-05 1.80E-04 2.60E-04 |
| WP\_129271396.1 | 216911 | 215858 | - | 350 | PF10099 | RskA | Anti-sigma-K factor rskA | 7.80E-19 |
| WP\_129271397.1 | 217494 | 216948 | - | 181 | TIGR02937 TIGR02989 TIGR02954 TIGR02985 TIGR02983 | TIGR02937 TIGR02989 TIGR02954 TIGR02985 TIGR02983 | sigma70-ECF: RNA polymerase sigma factor, sigma-70 family Sig-70\_gvs1: RNA polymerase sigma-70 factor, Rhodopirellula/Verrucomicrobium family Sig70\_famx3: RNA polymerase sigma-70 factor, TIGR02954 family Sig70\_bacteroi1: RNA polymerase sigma-70 factor, Bacteroides expansion family 1 SigE-fam\_strep: RNA polymerase sigma-70 factor, sigma-E family | 4.90E-28 9.50E-24 6.10E-22 7.40E-21 1.50E-17 |
| WP\_129271398.1 | 218090 | 217670 | - | 139 | TIGR02473 PF02050 PF03194 | TIGR02473 FliJ LUC7 | flagell\_FliJ: flagellar export protein FliJ Flagellar FliJ protein LUC7 N\_terminus | 8.20E-26 3.00E-12 8.10E-04 |
| WP\_129271399.1 | 219549 | 218223 | - | 441 | TIGR03498 TIGR03496 TIGR01026 TIGR03497 TIGR02546 | TIGR03498 TIGR03496 TIGR01026 TIGR03497 TIGR02546 | FliI\_clade3: flagellar protein export ATPase FliI FliI\_clade1: flagellar protein export ATPase FliI fliI\_yscN: ATPase, FliI/YscN family FliI\_clade2: flagellar protein export ATPase FliI III\_secr\_ATP: type III secretion apparatus H+-transporting two-sector ATPase | 1.70E-200 5.80E-168 2.60E-160 4.60E-158 4.40E-148 |
| WP\_015688285.1 | 219995 | 220697 | + | 233 | TIGR01387 TIGR02154 PF00072 TIGR03787 PF00486 | TIGR01387 TIGR02154 Response\_reg TIGR03787 Trans\_reg\_C | cztR\_silR\_copR: heavy metal response regulator PhoB: phosphate regulon transcriptional regulatory protein PhoB Response regulator receiver domain marine\_sort\_RR: proteobacterial dedicated sortase system response regulator Transcriptional regulatory protein, C terminal | 4.70E-49 7.40E-46 2.50E-26 3.10E-25 3.50E-20 |

#### Results for WP\_224586504.1 [Mesorhizobium sp. CA14] back to top

#### previous - next

##### Architecture

1000 nucleotides

Link to nucleotide sequence  
  

| Accession | start | end | direction | length (aa) | Pfam/HMM | name | description | E-value |
| --- | --- | --- | --- | --- | --- | --- | --- | --- |
| WP\_224586502.1 | 2509 | 538 | - | 656 | TIGR02026 TIGR03471 TIGR04014 TIGR03975 TIGR04013 | TIGR02026 TIGR03471 TIGR04014 TIGR03975 TIGR04013 | BchE: magnesium-protoporphyrin IX monomethyl ester anaerobic oxidative cyclase HpnJ: hopanoid biosynthesis associated radical SAM protein HpnJ B12\_SAM\_MJ\_0865: B12-binding domain/radical SAM domain protein, MJ\_0865 family rSAM\_ocin\_1: ribosomal peptide maturation radical SAM protein 1 B12\_SAM\_MJ\_1487: B12-binding domain/radical SAM domain protein, MJ\_1487 family | 3.10E-46 1.00E-39 4.80E-32 7.20E-29 1.30E-27 |
| WP\_224586504.1 | 4127 | 3317 | - | 269 | PF05114 PF01261 | DUF692 AP\_endonuc\_2 | Protein of unknown function (DUF692) Xylose isomerase-like TIM barrel | 1.00E-47 9.90E-08 |
| WP\_224586506.1 | 4681 | 5389 | + | 235 | PF04255 | DUF433 | Protein of unknown function (DUF433) | 3.80E-13 |
| WP\_224586508.1 | 5366 | 5783 | + | 138 | PF18480 | DUF5615 | Domain of unknown function (DUF5615) | 5.00E-18 |
| WP\_224586521.1 | 5939 | 5792 | - | 48 | NO PFAM MATCH | - | - | - |
| WP\_076611875.1 | 6540 | 7726 | + | 395 | INFERRED GENE | - | - | - |
| WP\_263488544.1 | 7908 | 7638 | - | 89 | PF08238 | Sel1 | Sel1 repeat | 2.00E-08 |
| WP\_224586692.1 | 8890 | 8641 | - | 82 | PF06412 | TraD | Conjugal transfer protein TraD | 3.40E-24 |
| WP\_224586522.1 | 9218 | 8891 | - | 108 | PF06412 | TraD | Conjugal transfer protein TraD | 5.10E-17 |
| WP\_224586524.1 | 9393 | 12429 | + | 1011 | TIGR02768 PF03389 PF13604 TIGR01448 TIGR01447 | TIGR02768 MobA\_MobL AAA\_30 TIGR01448 TIGR01447 | TraA\_Ti: Ti-type conjugative transfer relaxase TraA MobA/MobL family AAA domain recD\_rel: helicase, RecD/TraA family recD: exodeoxyribonuclease V, alpha subunit | 9.00E-277 1.70E-71 1.90E-55 8.80E-29 7.80E-20 |

#### Results for MQY12875.1 [Streptomyces smaragdinus] back to top

#### previous - next

##### Architecture

1000 nucleotides

Link to nucleotide sequence  
  

| Accession | start | end | direction | length (aa) | Pfam/HMM | name | description | E-value |
| --- | --- | --- | --- | --- | --- | --- | --- | --- |
| MQY12867.1 | 77170 | 77335 | + | 54 | TIGR01206 | TIGR01206 | lysW: lysine biosynthesis protein LysW | 2.10E-14 |
| MQY12868.1 | 77331 | 78219 | + | 295 | TIGR02144 TIGR00768 PF08443 TIGR04356 TIGR04443 | TIGR02144 TIGR00768 RimK TIGR04356 TIGR04443 | LysX\_arch: lysine biosynthesis enzyme LysX rimK\_fam: alpha-L-glutamate ligase, RimK family RimK-like ATP-grasp domain grasp\_GAK: ATP-grasp enzyme, GAK system F420\_CofF: coenzyme gamma-F420-2:alpha-L-glutamate ligase | 6.50E-84 3.10E-77 4.50E-28 1.10E-19 2.70E-19 |
| MQY12869.1 | 78206 | 79241 | + | 344 | TIGR01850 TIGR01851 PF01118 TIGR00978 | TIGR01850 TIGR01851 Semialdhyde\_dh TIGR00978 | argC: N-acetyl-gamma-glutamyl-phosphate reductase argC\_other: N-acetyl-gamma-glutamyl-phosphate reductase Semialdehyde dehydrogenase, NAD binding domain asd\_EA: aspartate-semialdehyde dehydrogenase | 7.80E-96 3.70E-25 9.20E-21 1.40E-04 |
| MQY12870.1 | 79268 | 80111 | + | 280 | TIGR00761 PF00696 TIGR01890 TIGR01027 PF10686 | TIGR00761 AA\_kinase TIGR01890 TIGR01027 YAcAr | argB: acetylglutamate kinase Amino acid kinase family N-Ac-Glu-synth: amino-acid N-acetyltransferase proB: glutamate 5-kinase YspA, cpYpsA-related SLOG family | 5.30E-51 2.60E-32 1.20E-11 4.30E-05 2.10E-04 |
| MQY12871.1 | 80107 | 81214 | + | 368 | TIGR01902 PF01546 TIGR01900 TIGR01910 TIGR01892 | TIGR01902 Peptidase\_M20 TIGR01900 TIGR01910 TIGR01892 | dapE-lys-deAc: N-acetyl-ornithine/N-acetyl-lysine deacetylase Peptidase family M20/M25/M40 dapE-gram\_pos: succinyl-diaminopimelate desuccinylase DapE-ArgE: peptidase, ArgE/DapE family AcOrn-deacetyl: acetylornithine deacetylase (ArgE) | 1.90E-89 2.40E-28 2.20E-26 3.90E-22 1.50E-19 |
| MQY12872.1 | 81210 | 82056 | + | 281 | TIGR00232 PF00456 TIGR00759 TIGR03186 TIGR03182 | TIGR00232 Transketolase\_N TIGR00759 TIGR03186 TIGR03182 | tktlase\_bact: transketolase Transketolase, thiamine diphosphate binding domain aceE: pyruvate dehydrogenase (acetyl-transferring), homodimeric type AKGDH\_not\_PDH: alpha-ketoglutarate dehydrogenase PDH\_E1\_alph\_y: pyruvate dehydrogenase (acetyl-transferring) E1 component, alpha subunit | 6.40E-48 1.80E-45 8.00E-13 5.50E-12 1.10E-11 |
| MQY12873.1 | 82052 | 83021 | + | 322 | TIGR00204 PF02779 PF02780 TIGR00232 | TIGR00204 Transket\_pyr Transketolase\_C TIGR00232 | dxs: 1-deoxy-D-xylulose-5-phosphate synthase Transketolase, pyrimidine binding domain Transketolase, C-terminal domain tktlase\_bact: transketolase | 1.50E-44 6.80E-29 2.50E-28 3.00E-19 |
| MQY12874.1 | 83098 | 83293 | + | 64 | NO PFAM MATCH | - | - | - |
| MQY12875.1 | 83334 | 84204 | + | 289 | PF05114 PF01261 | DUF692 AP\_endonuc\_2 | Protein of unknown function (DUF692) Xylose isomerase-like TIM barrel | 1.00E-40 8.40E-07 |
| MQY12876.1 | 84188 | 85067 | + | 292 | NO PFAM MATCH | - | - | - |
| MQY12877.1 | 85069 | 87043 | + | 657 | TIGR02026 TIGR03471 TIGR04479 TIGR04014 TIGR03975 | TIGR02026 TIGR03471 TIGR04479 TIGR04014 TIGR03975 | BchE: magnesium-protoporphyrin IX monomethyl ester anaerobic oxidative cyclase HpnJ: hopanoid biosynthesis associated radical SAM protein HpnJ bcpD\_PhpK\_rSAM: radical SAM P-methyltransferase, PhpK family B12\_SAM\_MJ\_0865: B12-binding domain/radical SAM domain protein, MJ\_0865 family rSAM\_ocin\_1: ribosomal peptide maturation radical SAM protein 1 | 1.30E-43 7.00E-37 6.20E-31 1.20E-30 2.10E-28 |
| MQY12878.1 | 87062 | 88028 | + | 321 | TIGR04543 PF13649 PF13847 TIGR00091 TIGR03534 | TIGR04543 Methyltransf\_25 Methyltransf\_31 TIGR00091 TIGR03534 | ketoArg\_3Met: 2-ketoarginine methyltransferase Methyltransferase domain Methyltransferase domain TIGR00091: tRNA (guanine-N(7)-)-methyltransferase RF\_mod\_PrmC: protein-(glutamine-N5) methyltransferase, release factor-specific | 6.50E-13 6.10E-07 2.20E-05 2.40E-05 4.90E-05 |
| MQY12879.1 | 88860 | 88032 | - | 275 | PF04672 PF08242 | Methyltransf\_19 Methyltransf\_12 | S-adenosyl methyltransferase Methyltransferase domain | 1.40E-100 2.80E-05 |
| MQY12880.1 | 89495 | 88949 | - | 181 | PF17920 PF00440 TIGR03384 TIGR03613 TIGR03968 | TetR\_C\_16 TetR\_N TIGR03384 TIGR03613 TIGR03968 | Tetracyclin repressor-like, C-terminal domain Bacterial regulatory proteins, tetR family betaine\_BetI: transcriptional repressor BetI RutR: pyrimidine utilization regulatory protein R mycofact\_TetR: mycofactocin system transcriptional regulator | 3.00E-21 2.20E-12 4.30E-10 2.80E-05 5.30E-05 |
| MQY12881.1 | 90976 | 89542 | - | 477 | TIGR00711 PF07690 TIGR00710 TIGR00880 TIGR00893 | TIGR00711 MFS\_1 TIGR00710 TIGR00880 TIGR00893 | efflux\_EmrB: drug resistance MFS transporter, drug:H+ antiporter-2 (14 Spanner) (DHA2) family Major Facilitator Superfamily efflux\_Bcr\_CflA: drug resistance transporter, Bcr/CflA subfamily 2\_A\_01\_02: multidrug resistance protein 2A0114: D-galactonate transporter | 1.30E-45 9.00E-41 3.40E-20 4.50E-17 3.10E-12 |
| MQY12882.1 | 91290 | 91914 | + | 207 | PF14016 | DUF4232 | Protein of unknown function (DUF4232) | 4.10E-09 |
| MQY12883.1 | 94315 | 91990 | - | 774 | TIGR01525 TIGR01512 TIGR01511 TIGR01494 PF00122 | TIGR01525 TIGR01512 TIGR01511 TIGR01494 E1-E2\_ATPase | ATPase-IB\_hvy: heavy metal translocating P-type ATPase ATPase-IB2\_Cd: cadmium-translocating P-type ATPase ATPase-IB1\_Cu: copper-translocating P-type ATPase ATPase\_P-type: HAD ATPase, P-type, family IC E1-E2 ATPase | 2.40E-159 1.60E-153 1.00E-100 1.50E-61 1.00E-36 |

#### Results for MBP1097539.1 [Bradyrhizobium japonicum] back to top

#### previous - next

##### Architecture

1000 nucleotides

Link to nucleotide sequence  
  

| Accession | start | end | direction | length (aa) | Pfam/HMM | name | description | E-value |
| --- | --- | --- | --- | --- | --- | --- | --- | --- |
| MBP1097531.1 | 8397254 | 8397647 | + | 130 | PF01527 PF13518 | HTH\_Tnp\_1 HTH\_28 | Transposase Helix-turn-helix domain | 4.50E-17 2.20E-04 |
| MBP1097532.1 | 8397665 | 8398484 | + | 272 | PF13683 PF13276 PF00665 | rve\_3 HTH\_21 rve | Integrase core domain HTH-like domain Integrase core domain | 4.00E-14 2.20E-13 3.50E-10 |
| MBP1097533.1 | 8399004 | 8398608 | - | 131 | NO PFAM MATCH | - | - | - |
| MBP1097534.1 | 8399552 | 8399288 | - | 87 | NO PFAM MATCH | - | - | - |
| MBP1097535.1 | 8401389 | 8399607 | - | 593 | NO PFAM MATCH | - | - | - |
| MBP1097536.1 | 8403539 | 8402546 | - | 330 | TIGR04543 PF13649 TIGR04074 PF13489 TIGR03534 | TIGR04543 Methyltransf\_25 TIGR04074 Methyltransf\_23 TIGR03534 | ketoArg\_3Met: 2-ketoarginine methyltransferase Methyltransferase domain bacter\_Hen1: 3' terminal RNA ribose 2'-O-methyltransferase Hen1 Methyltransferase domain RF\_mod\_PrmC: protein-(glutamine-N5) methyltransferase, release factor-specific | 1.90E-11 1.90E-08 1.40E-05 2.40E-05 3.10E-05 |
| MBP1097537.1 | 8405515 | 8403535 | - | 659 | TIGR02026 TIGR03471 TIGR04014 TIGR04013 TIGR03975 | TIGR02026 TIGR03471 TIGR04014 TIGR04013 TIGR03975 | BchE: magnesium-protoporphyrin IX monomethyl ester anaerobic oxidative cyclase HpnJ: hopanoid biosynthesis associated radical SAM protein HpnJ B12\_SAM\_MJ\_0865: B12-binding domain/radical SAM domain protein, MJ\_0865 family B12\_SAM\_MJ\_1487: B12-binding domain/radical SAM domain protein, MJ\_1487 family rSAM\_ocin\_1: ribosomal peptide maturation radical SAM protein 1 | 1.10E-35 1.40E-35 1.70E-25 2.50E-25 4.70E-25 |
| MBP1097538.1 | 8406356 | 8405495 | - | 286 | NO PFAM MATCH | - | - | - |
| MBP1097539.1 | 8407221 | 8406348 | - | 290 | PF05114 PF01261 | DUF692 AP\_endonuc\_2 | Protein of unknown function (DUF692) Xylose isomerase-like TIM barrel | 1.80E-38 8.30E-05 |
| MBP1097540.1 | 8407479 | 8407293 | - | 61 | NO PFAM MATCH | - | - | - |
| MBP1097541.1 | 8408691 | 8407599 | - | 363 | PF00067 TIGR04538 TIGR04458 | p450 TIGR04538 TIGR04458 | Cytochrome P450 P450\_cycloAA\_1: cytochrome P450, cyclodipeptide synthase-associated CYP450\_TxtE: 4-nitrotryptophan synthase | 8.50E-33 9.00E-10 1.60E-05 |
| MBP1097542.1 | 8409454 | 8410063 | + | 202 | NO PFAM MATCH | - | - | - |
| MBP1097543.1 | 8411744 | 8410181 | - | 520 | PF20172 TIGR02224 TIGR02225 PF00589 | DUF6538 TIGR02224 TIGR02225 Phage\_integrase | Domain of unknown function (DUF6538) recomb\_XerC: tyrosine recombinase XerC recomb\_XerD: tyrosine recombinase XerD Phage integrase family | 1.50E-12 3.60E-10 4.70E-10 8.20E-07 |
| MBP1097544.1 | 8412033 | 8412987 | + | 317 | TIGR03339 PF03466 TIGR02424 PF00126 TIGR03298 | TIGR03339 LysR\_substrate TIGR02424 HTH\_1 TIGR03298 | phn\_lysR: aminoethylphosphonate catabolism associated LysR family transcriptional regulator LysR substrate binding domain TF\_pcaQ: pca operon transcription factor PcaQ Bacterial regulatory helix-turn-helix protein, lysR family argP: transcriptional regulator, ArgP family | 2.00E-35 1.20E-33 2.90E-21 1.40E-18 1.90E-15 |
| MBP1097545.1 | 8413151 | 8414561 | + | 469 | TIGR00170 PF00330 TIGR01343 TIGR02086 TIGR02083 | TIGR00170 Aconitase TIGR01343 TIGR02086 TIGR02083 | leuC: 3-isopropylmalate dehydratase, large subunit Aconitase family (aconitate hydratase) hacA\_fam: homoaconitate hydratase family protein IPMI\_arch: 3-isopropylmalate dehydratase, large subunit LEU2: 3-isopropylmalate dehydratase, large subunit | 5.90E-194 1.20E-144 6.90E-96 3.70E-86 9.30E-79 |
| MBP1097546.1 | 8414561 | 8415197 | + | 211 | TIGR00171 PF00694 TIGR02087 TIGR02084 TIGR01342 | TIGR00171 Aconitase\_C TIGR02087 TIGR02084 TIGR01342 | leuD: 3-isopropylmalate dehydratase, small subunit Aconitase C-terminal domain LEUD\_arch: 3-isopropylmalate dehydratase, small subunit leud: 3-isopropylmalate dehydratase, small subunit acon\_putative: putative aconitate hydratase | 1.80E-52 6.30E-24 5.10E-18 2.50E-14 1.70E-10 |
| MBP1097547.1 | 8415193 | 8416333 | + | 379 | TIGR00169 PF00180 TIGR02088 TIGR02089 TIGR00175 | TIGR00169 Iso\_dh TIGR02088 TIGR02089 TIGR00175 | leuB: 3-isopropylmalate dehydrogenase Isocitrate/isopropylmalate dehydrogenase LEU3\_arch: isopropylmalate/isohomocitrate dehydrogenases TTC: tartrate dehydrogenase mito\_nad\_idh: isocitrate dehydrogenase, NAD-dependent | 3.60E-122 6.60E-118 2.50E-80 1.10E-70 6.60E-45 |

#### Results for WP\_153452477.1 [Streptomyces smaragdinus] back to top

#### previous - next

##### Architecture

1000 nucleotides

Link to nucleotide sequence  
  

| Accession | start | end | direction | length (aa) | Pfam/HMM | name | description | E-value |
| --- | --- | --- | --- | --- | --- | --- | --- | --- |
| WP\_153452471.1 | 77170 | 77335 | + | 54 | TIGR01206 | TIGR01206 | lysW: lysine biosynthesis protein LysW | 2.10E-14 |
| WP\_228390067.1 | 77376 | 78219 | + | 280 | TIGR02144 TIGR00768 PF08443 TIGR04356 TIGR04443 | TIGR02144 TIGR00768 RimK TIGR04356 TIGR04443 | LysX\_arch: lysine biosynthesis enzyme LysX rimK\_fam: alpha-L-glutamate ligase, RimK family RimK-like ATP-grasp domain grasp\_GAK: ATP-grasp enzyme, GAK system F420\_CofF: coenzyme gamma-F420-2:alpha-L-glutamate ligase | 2.50E-83 1.40E-76 4.20E-28 9.10E-20 2.30E-19 |
| WP\_153452553.1 | 78254 | 79241 | + | 328 | TIGR01850 TIGR01851 PF01118 | TIGR01850 TIGR01851 Semialdhyde\_dh | argC: N-acetyl-gamma-glutamyl-phosphate reductase argC\_other: N-acetyl-gamma-glutamyl-phosphate reductase Semialdehyde dehydrogenase, NAD binding domain | 3.60E-88 3.20E-25 7.70E-14 |
| WP\_153452472.1 | 79268 | 80111 | + | 280 | TIGR00761 PF00696 TIGR01890 TIGR01027 PF10686 | TIGR00761 AA\_kinase TIGR01890 TIGR01027 YAcAr | argB: acetylglutamate kinase Amino acid kinase family N-Ac-Glu-synth: amino-acid N-acetyltransferase proB: glutamate 5-kinase YspA, cpYpsA-related SLOG family | 5.30E-51 2.60E-32 1.20E-11 4.30E-05 2.10E-04 |
| WP\_153452473.1 | 80107 | 81214 | + | 368 | TIGR01902 PF01546 TIGR01900 TIGR01910 TIGR01892 | TIGR01902 Peptidase\_M20 TIGR01900 TIGR01910 TIGR01892 | dapE-lys-deAc: N-acetyl-ornithine/N-acetyl-lysine deacetylase Peptidase family M20/M25/M40 dapE-gram\_pos: succinyl-diaminopimelate desuccinylase DapE-ArgE: peptidase, ArgE/DapE family AcOrn-deacetyl: acetylornithine deacetylase (ArgE) | 1.90E-89 2.40E-28 2.20E-26 3.90E-22 1.50E-19 |
| WP\_153452474.1 | 81210 | 82056 | + | 281 | TIGR00232 PF00456 TIGR00759 TIGR03186 TIGR03182 | TIGR00232 Transketolase\_N TIGR00759 TIGR03186 TIGR03182 | tktlase\_bact: transketolase Transketolase, thiamine diphosphate binding domain aceE: pyruvate dehydrogenase (acetyl-transferring), homodimeric type AKGDH\_not\_PDH: alpha-ketoglutarate dehydrogenase PDH\_E1\_alph\_y: pyruvate dehydrogenase (acetyl-transferring) E1 component, alpha subunit | 6.40E-48 1.80E-45 8.00E-13 5.50E-12 1.10E-11 |
| WP\_153452475.1 | 82052 | 83021 | + | 322 | TIGR00204 PF02779 PF02780 TIGR00232 | TIGR00204 Transket\_pyr Transketolase\_C TIGR00232 | dxs: 1-deoxy-D-xylulose-5-phosphate synthase Transketolase, pyrimidine binding domain Transketolase, C-terminal domain tktlase\_bact: transketolase | 1.50E-44 6.80E-29 2.50E-28 3.00E-19 |
| WP\_153452476.1 | 83098 | 83293 | + | 64 | NO PFAM MATCH | - | - | - |
| WP\_153452477.1 | 83289 | 84204 | + | 304 | PF05114 PF01261 | DUF692 AP\_endonuc\_2 | Protein of unknown function (DUF692) Xylose isomerase-like TIM barrel | 1.20E-40 9.60E-07 |
| WP\_153452478.1 | 84188 | 85067 | + | 292 | NO PFAM MATCH | - | - | - |
| WP\_153452479.1 | 85069 | 87043 | + | 657 | TIGR02026 TIGR03471 TIGR04479 TIGR04014 TIGR03975 | TIGR02026 TIGR03471 TIGR04479 TIGR04014 TIGR03975 | BchE: magnesium-protoporphyrin IX monomethyl ester anaerobic oxidative cyclase HpnJ: hopanoid biosynthesis associated radical SAM protein HpnJ bcpD\_PhpK\_rSAM: radical SAM P-methyltransferase, PhpK family B12\_SAM\_MJ\_0865: B12-binding domain/radical SAM domain protein, MJ\_0865 family rSAM\_ocin\_1: ribosomal peptide maturation radical SAM protein 1 | 1.30E-43 7.00E-37 6.20E-31 1.20E-30 2.10E-28 |
| WP\_153452480.1 | 87044 | 88028 | + | 327 | TIGR04543 PF13649 PF13847 TIGR00091 TIGR03534 | TIGR04543 Methyltransf\_25 Methyltransf\_31 TIGR00091 TIGR03534 | ketoArg\_3Met: 2-ketoarginine methyltransferase Methyltransferase domain Methyltransferase domain TIGR00091: tRNA (guanine-N(7)-)-methyltransferase RF\_mod\_PrmC: protein-(glutamine-N5) methyltransferase, release factor-specific | 5.60E-13 6.30E-07 2.20E-05 2.50E-05 5.00E-05 |
| WP\_153452481.1 | 88860 | 88032 | - | 275 | PF04672 PF08242 | Methyltransf\_19 Methyltransf\_12 | S-adenosyl methyltransferase Methyltransferase domain | 1.40E-100 2.80E-05 |
| WP\_153452482.1 | 89495 | 88949 | - | 181 | PF17920 PF00440 TIGR03384 TIGR03613 TIGR03968 | TetR\_C\_16 TetR\_N TIGR03384 TIGR03613 TIGR03968 | Tetracyclin repressor-like, C-terminal domain Bacterial regulatory proteins, tetR family betaine\_BetI: transcriptional repressor BetI RutR: pyrimidine utilization regulatory protein R mycofact\_TetR: mycofactocin system transcriptional regulator | 3.00E-21 2.20E-12 4.30E-10 2.80E-05 5.30E-05 |
| WP\_153452554.1 | 90913 | 89542 | - | 456 | TIGR00711 PF07690 TIGR00710 TIGR00880 TIGR00893 | TIGR00711 MFS\_1 TIGR00710 TIGR00880 TIGR00893 | efflux\_EmrB: drug resistance MFS transporter, drug:H+ antiporter-2 (14 Spanner) (DHA2) family Major Facilitator Superfamily efflux\_Bcr\_CflA: drug resistance transporter, Bcr/CflA subfamily 2\_A\_01\_02: multidrug resistance protein 2A0114: D-galactonate transporter | 1.10E-45 9.20E-41 2.50E-20 4.70E-17 2.80E-12 |
| WP\_153452483.1 | 91290 | 91914 | + | 207 | PF14016 | DUF4232 | Protein of unknown function (DUF4232) | 4.10E-09 |
| WP\_153452484.1 | 94315 | 91990 | - | 774 | TIGR01525 TIGR01512 TIGR01511 TIGR01494 PF00122 | TIGR01525 TIGR01512 TIGR01511 TIGR01494 E1-E2\_ATPase | ATPase-IB\_hvy: heavy metal translocating P-type ATPase ATPase-IB2\_Cd: cadmium-translocating P-type ATPase ATPase-IB1\_Cu: copper-translocating P-type ATPase ATPase\_P-type: HAD ATPase, P-type, family IC E1-E2 ATPase | 2.40E-159 1.60E-153 1.00E-100 1.50E-61 1.00E-36 |

#### Results for WP\_134006280.1 [Kribbella sp. VKM Ac-2566] back to top

#### previous - next

##### Architecture

1000 nucleotides

Link to nucleotide sequence  
  

| Accession | start | end | direction | length (aa) | Pfam/HMM | name | description | E-value |
| --- | --- | --- | --- | --- | --- | --- | --- | --- |
| WP\_134006265.1 | 103347 | 102786 | - | 186 | NO PFAM MATCH | - | - | - |
| WP\_166680419.1 | 104597 | 103358 | - | 412 | PF13191 TIGR03420 PF00004 | AAA\_16 TIGR03420 AAA | AAA ATPase domain DnaA\_homol\_Hda: DnaA regulatory inactivator Hda ATPase family associated with various cellular activities (AAA) | 5.00E-09 1.90E-06 2.90E-05 |
| WP\_134006269.1 | 106438 | 104956 | - | 493 | NO PFAM MATCH | - | - | - |
| WP\_134006271.1 | 107127 | 106434 | - | 230 | NO PFAM MATCH | - | - | - |
| WP\_134006273.1 | 107365 | 107950 | + | 194 | NO PFAM MATCH | - | - | - |
| WP\_134006274.1 | 109589 | 108377 | - | 403 | PF08007 PF13621 | JmjC\_2 Cupin\_8 | JmjC domain Cupin-like domain | 1.30E-26 8.30E-06 |
| WP\_134006276.1 | 109933 | 109606 | - | 108 | NO PFAM MATCH | - | - | - |
| WP\_134006278.1 | 110631 | 110868 | + | 78 | NO PFAM MATCH | - | - | - |
| WP\_134006280.1 | 110900 | 111758 | + | 285 | PF05114 PF01261 | DUF692 AP\_endonuc\_2 | Protein of unknown function (DUF692) Xylose isomerase-like TIM barrel | 1.70E-40 8.10E-04 |
| WP\_134006282.1 | 111754 | 112633 | + | 292 | NO PFAM MATCH | - | - | - |
| WP\_134006283.1 | 112613 | 114359 | + | 581 | TIGR02026 TIGR03471 TIGR04014 TIGR04479 TIGR03975 | TIGR02026 TIGR03471 TIGR04014 TIGR04479 TIGR03975 | BchE: magnesium-protoporphyrin IX monomethyl ester anaerobic oxidative cyclase HpnJ: hopanoid biosynthesis associated radical SAM protein HpnJ B12\_SAM\_MJ\_0865: B12-binding domain/radical SAM domain protein, MJ\_0865 family bcpD\_PhpK\_rSAM: radical SAM P-methyltransferase, PhpK family rSAM\_ocin\_1: ribosomal peptide maturation radical SAM protein 1 | 2.50E-45 3.20E-37 3.50E-34 6.00E-32 2.70E-30 |
| WP\_134006285.1 | 114340 | 114673 | + | 110 | NO PFAM MATCH | - | - | - |
| WP\_134006287.1 | 114679 | 115684 | + | 334 | TIGR04543 PF13847 PF13649 TIGR04074 PF08241 | TIGR04543 Methyltransf\_31 Methyltransf\_25 TIGR04074 Methyltransf\_11 | ketoArg\_3Met: 2-ketoarginine methyltransferase Methyltransferase domain Methyltransferase domain bacter\_Hen1: 3' terminal RNA ribose 2'-O-methyltransferase Hen1 Methyltransferase domain | 1.40E-14 3.40E-09 5.00E-08 7.10E-06 1.40E-05 |
| WP\_134006289.1 | 117700 | 116242 | - | 485 | PF00589 TIGR02225 TIGR02224 TIGR02249 PF00392 | Phage\_integrase TIGR02225 TIGR02224 TIGR02249 GntR | Phage integrase family recomb\_XerD: tyrosine recombinase XerD recomb\_XerC: tyrosine recombinase XerC integrase\_gron: integron integrase Bacterial regulatory proteins, gntR family | 4.70E-23 8.50E-11 3.00E-10 7.60E-09 1.20E-08 |
| WP\_238177351.1 | 119190 | 117900 | - | 429 | NO PFAM MATCH | - | - | - |
| WP\_134006291.1 | 119889 | 119283 | - | 201 | PF03724 | META | META domain | 1.70E-07 |
| WP\_134006293.1 | 120398 | 119885 | - | 170 | TIGR02983 TIGR02937 TIGR02954 TIGR02985 PF08281 | TIGR02983 TIGR02937 TIGR02954 TIGR02985 Sigma70\_r4\_2 | SigE-fam\_strep: RNA polymerase sigma-70 factor, sigma-E family sigma70-ECF: RNA polymerase sigma factor, sigma-70 family Sig70\_famx3: RNA polymerase sigma-70 factor, TIGR02954 family Sig70\_bacteroi1: RNA polymerase sigma-70 factor, Bacteroides expansion family 1 Sigma-70, region 4 | 9.80E-44 6.80E-21 7.20E-16 8.30E-14 1.60E-12 |

#### Results for MDJ0853098.1 [Myxococcota bacterium] back to top

#### previous - next

##### Architecture

1000 nucleotides

Link to nucleotide sequence  
  

| Accession | start | end | direction | length (aa) | Pfam/HMM | name | description | E-value |
| --- | --- | --- | --- | --- | --- | --- | --- | --- |
| MDJ0853096.1 | 1712 | 44 | - | 555 | PF09423 PF19050 | PhoD PhoD\_2 | PhoD-like phosphatase PhoD related phosphatase | 2.50E-13 2.20E-09 |
| MDJ0853097.1 | 1954 | 2170 | + | 71 | NO PFAM MATCH | - | - | - |
| MDJ0853098.1 | 2180 | 3038 | + | 285 | PF05114 PF01261 | DUF692 AP\_endonuc\_2 | Protein of unknown function (DUF692) Xylose isomerase-like TIM barrel | 4.90E-42 1.50E-05 |
| MDJ0853099.1 | 3037 | 3898 | + | 286 | NO PFAM MATCH | - | - | - |
| MDJ0853100.1 | 3894 | 5862 | + | 655 | TIGR03471 TIGR02026 TIGR04014 TIGR04479 TIGR03975 | TIGR03471 TIGR02026 TIGR04014 TIGR04479 TIGR03975 | HpnJ: hopanoid biosynthesis associated radical SAM protein HpnJ BchE: magnesium-protoporphyrin IX monomethyl ester anaerobic oxidative cyclase B12\_SAM\_MJ\_0865: B12-binding domain/radical SAM domain protein, MJ\_0865 family bcpD\_PhpK\_rSAM: radical SAM P-methyltransferase, PhpK family rSAM\_ocin\_1: ribosomal peptide maturation radical SAM protein 1 | 2.50E-43 6.90E-36 1.10E-29 1.80E-29 6.70E-27 |
| MDJ0853101.1 | 5858 | 6842 | + | 327 | TIGR04543 PF13649 PF13847 TIGR03534 TIGR04074 | TIGR04543 Methyltransf\_25 Methyltransf\_31 TIGR03534 TIGR04074 | ketoArg\_3Met: 2-ketoarginine methyltransferase Methyltransferase domain Methyltransferase domain RF\_mod\_PrmC: protein-(glutamine-N5) methyltransferase, release factor-specific bacter\_Hen1: 3' terminal RNA ribose 2'-O-methyltransferase Hen1 | 1.50E-13 8.90E-10 5.40E-08 1.60E-07 1.10E-06 |
| MDJ0853102.1 | 7937 | 6884 | - | 350 | NO PFAM MATCH | - | - | - |
| MDJ0853103.1 | 9486 | 8037 | - | 482 | TIGR04510 PF00196 TIGR02917 TIGR03020 PF13432 | TIGR04510 GerE TIGR02917 TIGR03020 TPR\_16 | mod\_pep\_cyc: putative peptide modification system cyclase Bacterial regulatory proteins, luxR family PEP\_TPR\_lipo: putative PEP-CTERM system TPR-repeat lipoprotein EpsA: transcriptional regulator EpsA Tetratricopeptide repeat | 2.20E-15 2.40E-14 1.50E-11 3.10E-11 9.30E-11 |
| MDJ0853104.1 | 10497 | 9573 | - | 307 | PF03466 TIGR03339 TIGR03418 PF00126 TIGR02424 | LysR\_substrate TIGR03339 TIGR03418 HTH\_1 TIGR02424 | LysR substrate binding domain phn\_lysR: aminoethylphosphonate catabolism associated LysR family transcriptional regulator chol\_sulf\_TF: putative choline sulfate-utilization transcription factor Bacterial regulatory helix-turn-helix protein, lysR family TF\_pcaQ: pca operon transcription factor PcaQ | 1.60E-29 1.00E-24 3.60E-24 2.70E-19 5.30E-14 |
| MDJ0853105.1 | 10623 | 11271 | + | 215 | PF04075 TIGR00026 | F420H2\_quin\_red TIGR00026 | F420H(2)-dependent quinone reductase hi\_GC\_TIGR00026: deazaflavin-dependent oxidoreductase, nitroreductase family | 7.40E-05 2.20E-04 |
| MDJ0853106.1 | 12508 | 11449 | - | 352 | PF01609 PF13751 | DDE\_Tnp\_1 DDE\_Tnp\_1\_6 | Transposase DDE domain Transposase DDE domain | 1.50E-19 1.10E-12 |

#### Results for WP\_141343163.1 [Bradyrhizobium sp. USDA 3458] back to top

#### previous - next

##### Architecture

1000 nucleotides

Link to nucleotide sequence  
  

| Accession | start | end | direction | length (aa) | Pfam/HMM | name | description | E-value |
| --- | --- | --- | --- | --- | --- | --- | --- | --- |
| WP\_012561414.1 | 503 | 871 | + | 122 | INFERRED GENE | - | - | - |
| WP\_159109487.1 | 1544 | 953 | - | 196 | TIGR04543 PF13649 PF13847 PF08241 TIGR02021 | TIGR04543 Methyltransf\_25 Methyltransf\_31 Methyltransf\_11 TIGR02021 | ketoArg\_3Met: 2-ketoarginine methyltransferase Methyltransferase domain Methyltransferase domain Methyltransferase domain BchM-ChlM: magnesium protoporphyrin O-methyltransferase | 1.50E-11 4.00E-09 6.90E-08 8.50E-06 1.20E-05 |
| WP\_141343160.1 | 1960 | 1540 | - | 139 | NO PFAM MATCH | - | - | - |
| WP\_141343161.1 | 3899 | 1928 | - | 656 | TIGR02026 TIGR03471 TIGR04014 TIGR04479 TIGR03975 | TIGR02026 TIGR03471 TIGR04014 TIGR04479 TIGR03975 | BchE: magnesium-protoporphyrin IX monomethyl ester anaerobic oxidative cyclase HpnJ: hopanoid biosynthesis associated radical SAM protein HpnJ B12\_SAM\_MJ\_0865: B12-binding domain/radical SAM domain protein, MJ\_0865 family bcpD\_PhpK\_rSAM: radical SAM P-methyltransferase, PhpK family rSAM\_ocin\_1: ribosomal peptide maturation radical SAM protein 1 | 7.20E-40 8.10E-39 2.80E-28 1.20E-23 3.00E-23 |
| WP\_141343162.1 | 4756 | 3889 | - | 288 | PF14407 | Frankia\_peptide | Ribosomally synthesized peptide prototyped by Frankia Franean1\_4349. | 5.50E-04 |
| WP\_141343163.1 | 5622 | 4752 | - | 289 | PF05114 PF01261 | DUF692 AP\_endonuc\_2 | Protein of unknown function (DUF692) Xylose isomerase-like TIM barrel | 1.90E-41 8.20E-07 |
| WP\_141343164.1 | 5864 | 5636 | - | 75 | NO PFAM MATCH | - | - | - |
| WP\_141343165.1 | 6326 | 5987 | - | 112 | NO PFAM MATCH | - | - | - |
| WP\_141343166.1 | 6410 | 9131 | + | 906 | NO PFAM MATCH | - | - | - |
| WP\_141343167.1 | 9112 | 9445 | + | 110 | NO PFAM MATCH | - | - | - |
| WP\_141343168.1 | 9525 | 10395 | + | 289 | TIGR04187 TIGR04185 TIGR04184 TIGR04192 TIGR00768 | TIGR04187 TIGR04185 TIGR04184 TIGR04192 TIGR00768 | GRASP\_SAV\_5884: ATP-grasp ribosomal peptide maturase, SAV\_5884 family ATPgraspMvdC: ATP-grasp ribosomal peptide maturase, MvdC family ATPgraspMvdD: ATP-grasp ribosomal peptide maturase, MvdD family GRASP\_w\_spasm: ATP-GRASP peptide maturase, grasp-with-spasm system rimK\_fam: alpha-L-glutamate ligase, RimK family | 2.90E-38 1.00E-28 1.80E-18 2.00E-10 3.50E-05 |
| WP\_141343169.1 | 10585 | 10402 | - | 60 | NO PFAM MATCH | - | - | - |

#### Results for WP\_160330237.1 [Streptomyces roseifaciens] back to top

#### previous - next

##### Architecture

1000 nucleotides

Link to nucleotide sequence  
  

| Accession | start | end | direction | length (aa) | Pfam/HMM | name | description | E-value |
| --- | --- | --- | --- | --- | --- | --- | --- | --- |
| WP\_063804260.1 | 2248737 | 2248290 | - | 148 | NO PFAM MATCH | - | - | - |
| WP\_058043659.1 | 2249359 | 2249923 | + | 187 | PF15567 | Imm35 | Immunity protein 35 | 5.00E-20 |
| WP\_079110361.1 | 2250449 | 2252000 | + | 516 | PF00561 PF08386 TIGR01249 TIGR03056 TIGR02427 | Abhydrolase\_1 Abhydrolase\_4 TIGR01249 TIGR03056 TIGR02427 | alpha/beta hydrolase fold TAP-like protein pro\_imino\_pep\_1: prolyl aminopeptidase bchO\_mg\_che\_rel: putative magnesium chelatase accessory protein protocat\_pcaD: 3-oxoadipate enol-lactonase | 1.30E-16 2.10E-14 5.40E-12 2.30E-06 1.20E-05 |
| WP\_058043662.1 | 2252012 | 2253635 | + | 540 | PF00561 TIGR01249 PF08386 TIGR01250 | Abhydrolase\_1 TIGR01249 Abhydrolase\_4 TIGR01250 | alpha/beta hydrolase fold pro\_imino\_pep\_1: prolyl aminopeptidase TAP-like protein pro\_imino\_pep\_2: proline-specific peptidase | 9.40E-15 1.50E-11 2.50E-11 7.40E-05 |
| WP\_144440827.1 | 2255570 | 2253683 | - | 628 | NO PFAM MATCH | - | - | - |
| WP\_079110362.1 | 2256900 | 2255724 | - | 391 | TIGR02032 PF04820 PF01494 TIGR02023 PF13450 | TIGR02032 Trp\_halogenase FAD\_binding\_3 TIGR02023 NAD\_binding\_8 | GG-red-SF: geranylgeranyl reductase family Tryptophan halogenase FAD binding domain BchP-ChlP: geranylgeranyl reductase NAD(P)-binding Rossmann-like domain | 6.20E-24 1.40E-13 1.10E-10 3.90E-07 3.40E-04 |
| WP\_058043664.1 | 2259935 | 2256896 | - | 1012 | PF17990 PF18417 | LodA\_N LodA\_C | L-Lysine epsilon oxidase N-terminal L-lysine epsilon oxidase C-terminal domain | 2.50E-79 1.50E-37 |
| WP\_144440828.1 | 2260064 | 2260373 | + | 102 | NO PFAM MATCH | - | - | - |
| WP\_160330237.1 | 2260392 | 2261262 | + | 289 | PF05114 PF01261 | DUF692 AP\_endonuc\_2 | Protein of unknown function (DUF692) Xylose isomerase-like TIM barrel | 7.30E-40 1.20E-07 |
| WP\_144440829.1 | 2261270 | 2262137 | + | 288 | RREFam006 | PqqD\_RRE | RRE-containing protein in a pyrroloquinoline cluster | 2.00E-04 |
| WP\_079110364.1 | 2262139 | 2264161 | + | 673 | TIGR02026 TIGR03471 TIGR03975 TIGR04014 TIGR04479 | TIGR02026 TIGR03471 TIGR03975 TIGR04014 TIGR04479 | BchE: magnesium-protoporphyrin IX monomethyl ester anaerobic oxidative cyclase HpnJ: hopanoid biosynthesis associated radical SAM protein HpnJ rSAM\_ocin\_1: ribosomal peptide maturation radical SAM protein 1 B12\_SAM\_MJ\_0865: B12-binding domain/radical SAM domain protein, MJ\_0865 family bcpD\_PhpK\_rSAM: radical SAM P-methyltransferase, PhpK family | 2.40E-40 4.60E-36 5.00E-31 4.80E-28 2.10E-27 |
| WP\_144440830.1 | 2264162 | 2265290 | + | 375 | TIGR04543 PF13649 TIGR00091 PF13847 | TIGR04543 Methyltransf\_25 TIGR00091 Methyltransf\_31 | ketoArg\_3Met: 2-ketoarginine methyltransferase Methyltransferase domain TIGR00091: tRNA (guanine-N(7)-)-methyltransferase Methyltransferase domain | 1.10E-13 3.00E-06 3.80E-05 7.20E-04 |
| WP\_245700162.1 | 2266595 | 2265197 | - | 465 | PF04820 TIGR02032 PF12831 PF01266 PF01494 | Trp\_halogenase TIGR02032 FAD\_oxidored DAO FAD\_binding\_3 | Tryptophan halogenase GG-red-SF: geranylgeranyl reductase family FAD dependent oxidoreductase FAD dependent oxidoreductase FAD binding domain | 4.30E-20 9.60E-20 6.60E-08 1.70E-07 2.70E-06 |
| WP\_107105587.1 | 2268037 | 2266648 | - | 462 | TIGR03906 TIGR04064 TIGR03974 TIGR04068 TIGR04163 | TIGR03906 TIGR04064 TIGR03974 TIGR04068 TIGR04163 | quino\_hemo\_SAM: quinohemoprotein amine dehydrogenase maturation protein rSAM\_nif11: nif11-like peptide radical SAM maturase rSAM\_six\_Cys: SCIFF radical SAM maturase rSAM\_ocin\_clost: Cys-rich peptide radical SAM maturase CcpM rSAM\_cobopep: peptide-modifying radical SAM enzyme CbpB | 1.10E-93 1.60E-70 1.10E-63 3.30E-43 6.40E-37 |
| WP\_058043672.1 | 2268912 | 2268033 | - | 292 | PF18417 | LodA\_C | L-lysine epsilon oxidase C-terminal domain | 3.20E-05 |
| WP\_058043673.1 | 2270327 | 2268908 | - | 472 | PF17990 | LodA\_N | L-Lysine epsilon oxidase N-terminal | 2.20E-04 |
| WP\_160330239.1 | 2271696 | 2270367 | - | 442 | PF12902 TIGR04492 | Ferritin-like TIGR04492 | Ferritin-like VioB: iminophenyl-pyruvate dimer synthase VioB | 1.70E-35 1.50E-09 |

#### Results for WP\_306383405.1 [Pseudomonas protegens] back to top

#### previous - next

##### Architecture

1000 nucleotides

Link to nucleotide sequence  
  

| Accession | start | end | direction | length (aa) | Pfam/HMM | name | description | E-value |
| --- | --- | --- | --- | --- | --- | --- | --- | --- |
| WP\_076612215.1 | 144643 | 144837 | + | 64 | INFERRED GENE | - | - | - |
| WP\_306383399.1 | 146800 | 145174 | - | 541 | PF00383 TIGR02571 PF14437 TIGR00326 PF14439 | dCMP\_cyt\_deam\_1 TIGR02571 MafB19-deam TIGR00326 Bd3614-deam | Cytidine and deoxycytidylate deaminase zinc-binding region ComEB: ComE operon protein 2 MafB19-like deaminase eubact\_ribD: riboflavin biosynthesis protein RibD Bd3614-like deaminase | 5.00E-22 3.40E-15 5.40E-13 2.50E-04 5.50E-04 |
| WP\_306383400.1 | 147182 | 148175 | + | 330 | PF01381 | HTH\_3 | Helix-turn-helix | 7.60E-05 |
| WP\_010955146.1 | 148398 | 149849 | + | 483 | INFERRED GENE | - | - | - |
| WP\_306383401.1 | 150236 | 149900 | - | 111 | PF05717 | TnpB\_IS66 | IS66 Orf2 like protein | 1.60E-25 |
| WP\_306383402.1 | 150550 | 150232 | - | 105 | PF01527 | HTH\_Tnp\_1 | Transposase | 4.30E-12 |
| WP\_306383403.1 | 150629 | 150785 | + | 51 | NO PFAM MATCH | - | - | - |
| WP\_306383404.1 | 150884 | 151082 | + | 65 | NO PFAM MATCH | - | - | - |
| WP\_306383405.1 | 151109 | 151952 | + | 280 | PF05114 | DUF692 | Protein of unknown function (DUF692) | 6.80E-41 |
| WP\_306383406.1 | 151948 | 152770 | + | 273 | NO PFAM MATCH | - | - | - |
| WP\_306383407.1 | 153092 | 154118 | + | 341 | TIGR04543 PF13847 PF05175 TIGR03534 TIGR00536 | TIGR04543 Methyltransf\_31 MTS TIGR03534 TIGR00536 | ketoArg\_3Met: 2-ketoarginine methyltransferase Methyltransferase domain Methyltransferase small domain RF\_mod\_PrmC: protein-(glutamine-N5) methyltransferase, release factor-specific hemK\_fam: methyltransferase, HemK family | 1.50E-14 2.40E-11 1.50E-09 5.30E-09 3.20E-08 |
| WP\_306383408.1 | 155287 | 154114 | - | 390 | TIGR04269 TIGR03942 TIGR04496 TIGR04261 TIGR03906 | TIGR04269 TIGR03942 TIGR04496 TIGR04261 TIGR03906 | SAM\_SPASM\_FxsB: radical SAM/SPASM domain protein, FxsB family sulfatase\_rSAM: anaerobic sulfatase maturase rSAM\_XyeB: radical SAM/SPASM domain peptide maturase, XyeB family rSAM\_GlyRichRpt: radical SAM/SPASM domain protein, GRRM system quino\_hemo\_SAM: quinohemoprotein amine dehydrogenase maturation protein | 2.80E-105 3.40E-54 2.30E-52 6.10E-39 7.50E-24 |
| WP\_076612005.1 | 155854 | 157005 | + | 383 | INFERRED GENE | - | - | - |
| WP\_306383409.1 | 158370 | 157083 | - | 428 | PF00589 TIGR02224 TIGR02225 | Phage\_integrase TIGR02224 TIGR02225 | Phage integrase family recomb\_XerC: tyrosine recombinase XerC recomb\_XerD: tyrosine recombinase XerD | 1.20E-15 9.60E-13 2.70E-10 |
| WP\_011063767.1 | 159011 | 158501 | - | 169 | TIGR00621 PF00436 TIGR04418 | TIGR00621 SSB TIGR04418 | ssb: single-stranded DNA-binding protein Single-strand binding protein family PriB\_gamma: primosomal replication protein PriB | 1.80E-58 1.00E-37 3.80E-05 |
| WP\_110596282.1 | 160415 | 159020 | - | 464 | PF07690 TIGR00880 TIGR00710 TIGR00711 TIGR00895 | MFS\_1 TIGR00880 TIGR00710 TIGR00711 TIGR00895 | Major Facilitator Superfamily 2\_A\_01\_02: multidrug resistance protein efflux\_Bcr\_CflA: drug resistance transporter, Bcr/CflA subfamily efflux\_EmrB: drug resistance MFS transporter, drug:H+ antiporter-2 (14 Spanner) (DHA2) family 2A0115: MFS transporter, aromatic acid:H+ symporter (AAHS) family | 2.30E-45 3.40E-22 8.70E-20 1.80E-15 3.40E-15 |
| WP\_102882401.1 | 160592 | 163427 | + | 944 | TIGR00630 PF17760 TIGR04521 TIGR03608 TIGR02857 | TIGR00630 UvrA\_inter TIGR04521 TIGR03608 TIGR02857 | uvra: excinuclease ABC subunit A UvrA interaction domain ECF\_ATPase\_2: energy-coupling factor transporter ATPase L\_ocin\_972\_ABC: putative bacteriocin export ABC transporter, lactococcin 972 group CydD: thiol reductant ABC exporter, CydD subunit | 0.00E+00 1.00E-38 2.60E-36 2.50E-35 3.50E-35 |

#### Results for WP\_257551926.1 [Streptomyces sp. NBC\_00162] back to top

#### previous - next

##### Architecture

1000 nucleotides

Link to nucleotide sequence  
  

| Accession | start | end | direction | length (aa) | Pfam/HMM | name | description | E-value |
| --- | --- | --- | --- | --- | --- | --- | --- | --- |
| WP\_257551918.1 | 16586 | 16127 | - | 152 | NO PFAM MATCH | - | - | - |
| WP\_257551919.1 | 16895 | 16658 | - | 78 | NO PFAM MATCH | - | - | - |
| WP\_257551920.1 | 17349 | 17007 | - | 113 | PF18977 | DUF5713 | Family of unknown function (DUF5713) | 2.60E-45 |
| WP\_257551921.1 | 17544 | 18816 | + | 423 | PF00872 | Transposase\_mut | Transposase, Mutator family | 9.10E-115 |
| WP\_257551922.1 | 20035 | 19099 | - | 311 | TIGR04543 PF13649 PF13847 TIGR03534 TIGR03533 | TIGR04543 Methyltransf\_25 Methyltransf\_31 TIGR03534 TIGR03533 | ketoArg\_3Met: 2-ketoarginine methyltransferase Methyltransferase domain Methyltransferase domain RF\_mod\_PrmC: protein-(glutamine-N5) methyltransferase, release factor-specific L3\_gln\_methyl: protein-(glutamine-N5) methyltransferase, ribosomal protein L3-specific | 2.10E-14 1.20E-07 3.00E-07 1.80E-06 4.90E-06 |
| WP\_257551923.1 | 20437 | 20092 | - | 114 | RREFam003 | Lasso\_Fused\_RRE | RRE-containing fused peptidase in a lasso peptide cluster | 2.10E-04 |
| WP\_257551924.1 | 22086 | 20391 | - | 564 | TIGR02026 TIGR03471 TIGR03975 TIGR04014 TIGR04479 | TIGR02026 TIGR03471 TIGR03975 TIGR04014 TIGR04479 | BchE: magnesium-protoporphyrin IX monomethyl ester anaerobic oxidative cyclase HpnJ: hopanoid biosynthesis associated radical SAM protein HpnJ rSAM\_ocin\_1: ribosomal peptide maturation radical SAM protein 1 B12\_SAM\_MJ\_0865: B12-binding domain/radical SAM domain protein, MJ\_0865 family bcpD\_PhpK\_rSAM: radical SAM P-methyltransferase, PhpK family | 1.20E-42 4.90E-39 1.10E-30 2.30E-30 7.00E-30 |
| WP\_257551925.1 | 22985 | 22088 | - | 298 | NO PFAM MATCH | - | - | - |
| WP\_257551926.1 | 23839 | 22969 | - | 289 | PF05114 PF01261 | DUF692 AP\_endonuc\_2 | Protein of unknown function (DUF692) Xylose isomerase-like TIM barrel | 1.00E-38 4.40E-05 |
| WP\_257551927.1 | 24091 | 23896 | - | 64 | NO PFAM MATCH | - | - | - |
| WP\_020460478.1 | 24281 | 24641 | + | 120 | INFERRED GENE | - | - | - |
| WP\_257551928.1 | 25574 | 24929 | - | 214 | NO PFAM MATCH | - | - | - |
| WP\_012377440.1 | 26541 | 26848 | + | 102 | INFERRED GENE | - | - | - |
| WP\_257551929.1 | 27761 | 28268 | + | 168 | NO PFAM MATCH | - | - | - |
| WP\_257551906.1 | 28541 | 28823 | + | 93 | NO PFAM MATCH | - | - | - |
| WP\_257551930.1 | 28819 | 29149 | + | 109 | PF19746 | DUF6233 | Family of unknown function (DUF6233) | 1.70E-18 |
| WP\_257551931.1 | 29938 | 29257 | - | 226 | NO PFAM MATCH | - | - | - |

#### Results for MBV8367743.1 [Candidatus Eremiobacteraeota bacterium] back to top

#### previous - next

##### Architecture

1000 nucleotides

Link to nucleotide sequence  
  

| Accession | start | end | direction | length (aa) | Pfam/HMM | name | description | E-value |
| --- | --- | --- | --- | --- | --- | --- | --- | --- |
| MBV8367735.1 | 45655 | 45217 | - | 145 | NO PFAM MATCH | - | - | - |
| MBV8367736.1 | 45738 | 46458 | + | 239 | NO PFAM MATCH | - | - | - |
| MBV8367737.1 | 46462 | 47398 | + | 311 | TIGR01473 PF01040 TIGR01474 TIGR01475 | TIGR01473 UbiA TIGR01474 TIGR01475 | cyoE\_ctaB: protoheme IX farnesyltransferase UbiA prenyltransferase family ubiA\_proteo: 4-hydroxybenzoate polyprenyl transferase ubiA\_other: putative 4-hydroxybenzoate polyprenyltransferase | 7.40E-102 1.30E-64 4.80E-24 2.80E-18 |
| MBV8367738.1 | 47412 | 48822 | + | 469 | TIGR00711 PF07690 TIGR00880 TIGR00710 TIGR00895 | TIGR00711 MFS\_1 TIGR00880 TIGR00710 TIGR00895 | efflux\_EmrB: drug resistance MFS transporter, drug:H+ antiporter-2 (14 Spanner) (DHA2) family Major Facilitator Superfamily 2\_A\_01\_02: multidrug resistance protein efflux\_Bcr\_CflA: drug resistance transporter, Bcr/CflA subfamily 2A0115: MFS transporter, aromatic acid:H+ symporter (AAHS) family | 1.40E-58 3.00E-54 8.40E-28 1.10E-21 3.10E-21 |
| MBV8367739.1 | 48863 | 50039 | + | 391 | TIGR04269 TIGR04496 TIGR03942 TIGR03906 TIGR04261 | TIGR04269 TIGR04496 TIGR03942 TIGR03906 TIGR04261 | SAM\_SPASM\_FxsB: radical SAM/SPASM domain protein, FxsB family rSAM\_XyeB: radical SAM/SPASM domain peptide maturase, XyeB family sulfatase\_rSAM: anaerobic sulfatase maturase quino\_hemo\_SAM: quinohemoprotein amine dehydrogenase maturation protein rSAM\_GlyRichRpt: radical SAM/SPASM domain protein, GRRM system | 2.80E-109 1.70E-53 3.70E-49 3.00E-26 6.80E-26 |
| MBV8367740.1 | 51074 | 50042 | - | 343 | TIGR04543 PF13649 PF13847 TIGR03534 TIGR00091 | TIGR04543 Methyltransf\_25 Methyltransf\_31 TIGR03534 TIGR00091 | ketoArg\_3Met: 2-ketoarginine methyltransferase Methyltransferase domain Methyltransferase domain RF\_mod\_PrmC: protein-(glutamine-N5) methyltransferase, release factor-specific TIGR00091: tRNA (guanine-N(7)-)-methyltransferase | 1.70E-09 1.90E-09 5.70E-09 2.70E-07 9.10E-07 |
| MBV8367741.1 | 53065 | 51076 | - | 662 | TIGR02026 TIGR03471 TIGR04014 TIGR03975 TIGR04013 | TIGR02026 TIGR03471 TIGR04014 TIGR03975 TIGR04013 | BchE: magnesium-protoporphyrin IX monomethyl ester anaerobic oxidative cyclase HpnJ: hopanoid biosynthesis associated radical SAM protein HpnJ B12\_SAM\_MJ\_0865: B12-binding domain/radical SAM domain protein, MJ\_0865 family rSAM\_ocin\_1: ribosomal peptide maturation radical SAM protein 1 B12\_SAM\_MJ\_1487: B12-binding domain/radical SAM domain protein, MJ\_1487 family | 1.60E-42 3.60E-38 4.80E-31 1.10E-30 1.40E-30 |
| MBV8367742.1 | 53838 | 53055 | - | 260 | NO PFAM MATCH | - | - | - |
| MBV8367743.1 | 54734 | 53807 | - | 308 | PF05114 PF01261 | DUF692 AP\_endonuc\_2 | Protein of unknown function (DUF692) Xylose isomerase-like TIM barrel | 3.90E-41 1.20E-09 |
| MBV8367744.1 | 54894 | 54708 | - | 61 | NO PFAM MATCH | - | - | - |
| MBV8367745.1 | 55978 | 54970 | - | 335 | PF02630 PF11604 PF00578 PF08534 | SCO1-SenC CusF\_Ec AhpC-TSA Redoxin | SCO1/SenC Copper binding periplasmic protein CusF AhpC/TSA family Redoxin | 1.00E-17 5.00E-10 1.00E-04 1.10E-04 |
| MBV8367746.1 | 56116 | 56416 | + | 99 | PF01355 | HIPIP | High potential iron-sulfur protein | 1.30E-06 |
| MBV8367747.1 | 56629 | 56443 | - | 61 | NO PFAM MATCH | - | - | - |
| MBV8367748.1 | 56799 | 57114 | + | 104 | TIGR02945 TIGR03406 PF01883 TIGR02159 TIGR00411 | TIGR02945 TIGR03406 FeS\_assembly\_P TIGR02159 TIGR00411 | SUF\_assoc: FeS assembly SUF system protein FeS\_long\_SufT: probable FeS assembly SUF system protein SufT Iron-sulfur cluster assembly protein PA\_CoA\_Oxy4: phenylacetate-CoA oxygenase, PaaJ subunit redox\_disulf\_1: redox-active disulfide protein 1 | 1.30E-28 1.90E-25 2.40E-20 3.00E-10 3.50E-04 |
| MBV8367749.1 | 57596 | 57110 | - | 161 | TIGR04110 TIGR04109 PF01243 TIGR03618 | TIGR04110 TIGR04109 Putative\_PNPOx TIGR03618 | heme\_HutZ: heme utilization protein HutZ heme\_ox\_HugZ: heme oxygenase, HugZ family Pyridoxamine 5'-phosphate oxidase Rv1155\_F420: PPOX class probable F420-dependent enzyme | 5.60E-27 3.60E-18 1.40E-08 1.00E-04 |
| MBV8367750.1 | 58014 | 57651 | - | 120 | NO PFAM MATCH | - | - | - |
| MBV8367751.1 | 58400 | 58010 | - | 129 | NO PFAM MATCH | - | - | - |

#### Results for WP\_299281768.1 [uncultured Tateyamaria sp.] back to top

#### previous - next

##### Architecture

1000 nucleotides

Link to nucleotide sequence  
  

| Accession | start | end | direction | length (aa) | Pfam/HMM | name | description | E-value |
| --- | --- | --- | --- | --- | --- | --- | --- | --- |
| WP\_299281753.1 | 272474 | 271919 | - | 184 | PF13521 TIGR01526 PF02223 | AAA\_28 TIGR01526 Thymidylate\_kin | AAA domain nadR\_NMN\_Atrans: nicotinamide-nucleotide adenylyltransferase Thymidylate kinase | 2.10E-18 7.50E-05 2.20E-04 |
| WP\_299281755.1 | 273357 | 272661 | - | 231 | NO PFAM MATCH | - | - | - |
| WP\_299281757.1 | 273527 | 273353 | - | 57 | TIGR02607 | TIGR02607 | antidote\_HigA: addiction module antidote protein, HigA family | 2.30E-10 |
| WP\_299281844.1 | 274803 | 273639 | - | 387 | PF06114 PF01381 | Peptidase\_M78 HTH\_3 | IrrE N-terminal-like domain Helix-turn-helix | 7.70E-06 1.30E-04 |
| WP\_299281760.1 | 275171 | 274802 | - | 122 | NO PFAM MATCH | - | - | - |
| WP\_299281762.1 | 275311 | 275971 | + | 219 | PF14452 | Multi\_ubiq | Multiubiquitin | 4.80E-33 |
| WP\_299281764.1 | 275945 | 277127 | + | 393 | PF20590 PF00899 TIGR02356 TIGR02354 | DUF6791 ThiF TIGR02356 TIGR02354 | Domain of unknown function (DUF6791) ThiF family adenyl\_thiF: thiazole biosynthesis adenylyltransferase ThiF thiF\_fam2: thiamine biosynthesis protein ThiF | 2.20E-51 8.30E-10 8.70E-07 9.00E-06 |
| WP\_299281766.1 | 278020 | 278206 | + | 61 | NO PFAM MATCH | - | - | - |
| WP\_299281768.1 | 278222 | 279152 | + | 309 | PF05114 PF01261 | DUF692 AP\_endonuc\_2 | Protein of unknown function (DUF692) Xylose isomerase-like TIM barrel | 7.70E-39 9.00E-07 |
| WP\_299281770.1 | 279148 | 279967 | + | 272 | NO PFAM MATCH | - | - | - |
| WP\_299281772.1 | 279963 | 281889 | + | 641 | TIGR02026 TIGR03471 TIGR04014 TIGR04479 TIGR03975 | TIGR02026 TIGR03471 TIGR04014 TIGR04479 TIGR03975 | BchE: magnesium-protoporphyrin IX monomethyl ester anaerobic oxidative cyclase HpnJ: hopanoid biosynthesis associated radical SAM protein HpnJ B12\_SAM\_MJ\_0865: B12-binding domain/radical SAM domain protein, MJ\_0865 family bcpD\_PhpK\_rSAM: radical SAM P-methyltransferase, PhpK family rSAM\_ocin\_1: ribosomal peptide maturation radical SAM protein 1 | 4.70E-39 2.50E-36 6.40E-29 6.00E-26 1.30E-25 |
| WP\_299281773.1 | 281927 | 282443 | + | 171 | TIGR02469 | TIGR02469 | CbiT: precorrin-6Y C5,15-methyltransferase (decarboxylating), CbiT subunit | 2.20E-04 |
| WP\_299281774.1 | 282522 | 282909 | + | 128 | PF01527 PF13518 | HTH\_Tnp\_1 HTH\_28 | Transposase Helix-turn-helix domain | 9.50E-16 3.30E-05 |
| WP\_299281775.1 | 282938 | 283259 | + | 106 | PF05717 | TnpB\_IS66 | IS66 Orf2 like protein | 1.70E-36 |
| WP\_076612405.1 | 283313 | 284783 | + | 490 | INFERRED GENE | - | - | - |
| WP\_299281777.1 | 285259 | 286489 | + | 409 | TIGR04269 TIGR04496 TIGR03942 TIGR04261 TIGR03906 | TIGR04269 TIGR04496 TIGR03942 TIGR04261 TIGR03906 | SAM\_SPASM\_FxsB: radical SAM/SPASM domain protein, FxsB family rSAM\_XyeB: radical SAM/SPASM domain peptide maturase, XyeB family sulfatase\_rSAM: anaerobic sulfatase maturase rSAM\_GlyRichRpt: radical SAM/SPASM domain protein, GRRM system quino\_hemo\_SAM: quinohemoprotein amine dehydrogenase maturation protein | 4.20E-105 2.40E-53 1.20E-49 3.60E-36 3.00E-27 |
| WP\_299281779.1 | 287973 | 286938 | - | 344 | PF13391 | HNH\_2 | HNH endonuclease | 9.40E-08 |

#### Results for MBP6189622.1 [Azonexus sp.] back to top

#### previous - next

##### Architecture

1000 nucleotides

Link to nucleotide sequence  
  

| Accession | start | end | direction | length (aa) | Pfam/HMM | name | description | E-value |
| --- | --- | --- | --- | --- | --- | --- | --- | --- |
| MBP6189614.1 | 1233 | 3327 | + | 697 | PF02624 TIGR03604 TIGR03266 TIGR00702 TIGR03882 | YcaO TIGR03604 TIGR03266 TIGR00702 TIGR03882 | YcaO cyclodehydratase, ATP-ad Mg2+-binding TOMM\_cyclo\_SagD: thiazole/oxazole-forming peptide maturase, SagD family component methan\_mark\_1: putative methanogenesis marker protein 1 TIGR00702: YcaO-type kinase domain cyclo\_dehyd\_2: bacteriocin biosynthesis cyclodehydratase domain | 2.20E-86 1.70E-67 1.60E-41 7.00E-31 8.50E-24 |
| MBP6189615.1 | 3357 | 3993 | + | 211 | PF20043 | DUF6445 | Family of unknown function (DUF6445) | 1.60E-09 |
| MBP6189616.1 | 3993 | 5370 | + | 458 | TIGR03605 TIGR04511 PF00881 | TIGR03605 TIGR04511 Nitroreductase | antibiot\_sagB: SagB-type dehydrogenase domain SagB\_rel\_DH\_2: putative peptide maturation dehydrogenase Nitroreductase family | 2.00E-35 1.10E-11 8.30E-08 |
| MBP6189617.1 | 5440 | 6085 | + | 214 | PF13621 | Cupin\_8 | Cupin-like domain | 3.90E-04 |
| MBP6189618.1 | 6242 | 7448 | + | 401 | TIGR04269 TIGR03942 TIGR04496 TIGR04261 TIGR03906 | TIGR04269 TIGR03942 TIGR04496 TIGR04261 TIGR03906 | SAM\_SPASM\_FxsB: radical SAM/SPASM domain protein, FxsB family sulfatase\_rSAM: anaerobic sulfatase maturase rSAM\_XyeB: radical SAM/SPASM domain peptide maturase, XyeB family rSAM\_GlyRichRpt: radical SAM/SPASM domain protein, GRRM system quino\_hemo\_SAM: quinohemoprotein amine dehydrogenase maturation protein | 6.00E-103 6.30E-51 1.50E-47 8.00E-36 1.10E-25 |
| MBP6189619.1 | 8480 | 7457 | - | 340 | TIGR04543 PF13649 PF13847 TIGR03534 TIGR02469 | TIGR04543 Methyltransf\_25 Methyltransf\_31 TIGR03534 TIGR02469 | ketoArg\_3Met: 2-ketoarginine methyltransferase Methyltransferase domain Methyltransferase domain RF\_mod\_PrmC: protein-(glutamine-N5) methyltransferase, release factor-specific CbiT: precorrin-6Y C5,15-methyltransferase (decarboxylating), CbiT subunit | 6.20E-16 3.60E-09 3.90E-09 1.70E-08 1.00E-06 |
| MBP6189620.1 | 10471 | 8476 | - | 664 | TIGR02026 TIGR03471 TIGR04014 TIGR03975 TIGR04013 | TIGR02026 TIGR03471 TIGR04014 TIGR03975 TIGR04013 | BchE: magnesium-protoporphyrin IX monomethyl ester anaerobic oxidative cyclase HpnJ: hopanoid biosynthesis associated radical SAM protein HpnJ B12\_SAM\_MJ\_0865: B12-binding domain/radical SAM domain protein, MJ\_0865 family rSAM\_ocin\_1: ribosomal peptide maturation radical SAM protein 1 B12\_SAM\_MJ\_1487: B12-binding domain/radical SAM domain protein, MJ\_1487 family | 1.40E-35 4.60E-34 3.10E-29 5.80E-28 2.80E-27 |
| MBP6189621.1 | 11274 | 10467 | - | 268 | NO PFAM MATCH | - | - | - |
| MBP6189622.1 | 12128 | 11273 | - | 284 | PF05114 PF01261 | DUF692 AP\_endonuc\_2 | Protein of unknown function (DUF692) Xylose isomerase-like TIM barrel | 8.60E-43 1.80E-05 |
| MBP6189623.1 | 12350 | 12152 | - | 65 | NO PFAM MATCH | - | - | - |
| MBP6189624.1 | 14564 | 13133 | - | 476 | PF05598 PF13751 PF01609 | DUF772 DDE\_Tnp\_1\_6 DDE\_Tnp\_1 | Transposase domain (DUF772) Transposase DDE domain Transposase DDE domain | 2.80E-17 2.70E-16 2.30E-13 |

#### Results for WP\_074971115.1 [Paracoccus aminovorans] back to top

#### previous - next

##### Architecture

1000 nucleotides

Link to nucleotide sequence  
  

| Accession | start | end | direction | length (aa) | Pfam/HMM | name | description | E-value |
| --- | --- | --- | --- | --- | --- | --- | --- | --- |
| WP\_007803351.1 | 0 | 205 | + | 68 | INFERRED GENE | - | - | - |
| WP\_231964699.1 | 333 | 657 | + | 107 | NO PFAM MATCH | - | - | - |
| WP\_083412974.1 | 758 | 1979 | + | 406 | TIGR04269 TIGR04496 TIGR03942 TIGR04261 TIGR03906 | TIGR04269 TIGR04496 TIGR03942 TIGR04261 TIGR03906 | SAM\_SPASM\_FxsB: radical SAM/SPASM domain protein, FxsB family rSAM\_XyeB: radical SAM/SPASM domain peptide maturase, XyeB family sulfatase\_rSAM: anaerobic sulfatase maturase rSAM\_GlyRichRpt: radical SAM/SPASM domain protein, GRRM system quino\_hemo\_SAM: quinohemoprotein amine dehydrogenase maturation protein | 1.20E-101 3.00E-49 4.80E-47 7.00E-31 6.40E-29 |
| WP\_170848994.1 | 2928 | 1956 | - | 323 | TIGR04543 PF13649 PF13847 TIGR00091 TIGR00740 | TIGR04543 Methyltransf\_25 Methyltransf\_31 TIGR00091 TIGR00740 | ketoArg\_3Met: 2-ketoarginine methyltransferase Methyltransferase domain Methyltransferase domain TIGR00091: tRNA (guanine-N(7)-)-methyltransferase TIGR00740: tRNA (cmo5U34)-methyltransferase | 7.80E-12 4.30E-09 1.80E-08 2.40E-07 1.20E-05 |
| WP\_139218153.1 | 4925 | 2960 | - | 654 | TIGR03471 TIGR02026 TIGR04014 TIGR04479 TIGR04367 | TIGR03471 TIGR02026 TIGR04014 TIGR04479 TIGR04367 | HpnJ: hopanoid biosynthesis associated radical SAM protein HpnJ BchE: magnesium-protoporphyrin IX monomethyl ester anaerobic oxidative cyclase B12\_SAM\_MJ\_0865: B12-binding domain/radical SAM domain protein, MJ\_0865 family bcpD\_PhpK\_rSAM: radical SAM P-methyltransferase, PhpK family HpnR\_B12\_rSAM: hopanoid C-3 methylase HpnR | 1.90E-37 1.70E-35 1.40E-26 1.40E-26 1.80E-26 |
| WP\_139218154.1 | 5740 | 4921 | - | 272 | PF11590 | DNAPolymera\_Pol | DNA polymerase catalytic subunit Pol | 2.70E-04 |
| WP\_074971115.1 | 6621 | 5736 | - | 294 | PF05114 | DUF692 | Protein of unknown function (DUF692) | 4.00E-39 |
| WP\_074971105.1 | 6868 | 6682 | - | 61 | NO PFAM MATCH | - | - | - |
| WP\_074971107.1 | 7777 | 6973 | - | 267 | PF05721 | PhyH | Phytanoyl-CoA dioxygenase (PhyH) | 7.70E-25 |
| WP\_074971109.1 | 9648 | 7752 | - | 631 | TIGR03975 TIGR04367 TIGR04013 TIGR03471 TIGR04014 | TIGR03975 TIGR04367 TIGR04013 TIGR03471 TIGR04014 | rSAM\_ocin\_1: ribosomal peptide maturation radical SAM protein 1 HpnR\_B12\_rSAM: hopanoid C-3 methylase HpnR B12\_SAM\_MJ\_1487: B12-binding domain/radical SAM domain protein, MJ\_1487 family HpnJ: hopanoid biosynthesis associated radical SAM protein HpnJ B12\_SAM\_MJ\_0865: B12-binding domain/radical SAM domain protein, MJ\_0865 family | 9.40E-168 1.60E-11 2.40E-09 3.50E-09 3.20E-08 |
| WP\_074971111.1 | 9895 | 9658 | - | 78 | NO PFAM MATCH | - | - | - |
| WP\_170848995.1 | 10874 | 10671 | - | 66 | PF01695 | IstB\_IS21 | IstB-like ATP binding protein | 2.00E-26 |

#### Results for MTH36435.1 [Paracoccus limosus] back to top

#### previous - next

##### Architecture

1000 nucleotides

Link to nucleotide sequence  
  

| Accession | start | end | direction | length (aa) | Pfam/HMM | name | description | E-value |
| --- | --- | --- | --- | --- | --- | --- | --- | --- |
| WP\_017998486.1 | 126 | 603 | + | 159 | INFERRED GENE | - | - | - |
| MTH36428.1 | 599 | 1034 | + | 144 | TIGR02219 PF00877 | TIGR02219 NLPC\_P60 | phage\_NlpC\_fam: putative phage cell wall peptidase, NlpC/P60 family NlpC/P60 family | 5.50E-47 4.40E-06 |
| MTH36429.1 | 1043 | 5009 | + | 1321 | PF13547 PF13550 | GTA\_TIM Phage-tail\_3 | GTA TIM-barrel-like domain Putative phage tail protein | 1.80E-151 3.70E-39 |
| MTH36430.1 | 5019 | 6945 | + | 641 | PF10983 | DUF2793 | Protein of unknown function (DUF2793) | 8.80E-34 |
| MTH36431.1 | 7231 | 8452 | + | 406 | TIGR04269 TIGR04496 TIGR03942 TIGR04261 TIGR03906 | TIGR04269 TIGR04496 TIGR03942 TIGR04261 TIGR03906 | SAM\_SPASM\_FxsB: radical SAM/SPASM domain protein, FxsB family rSAM\_XyeB: radical SAM/SPASM domain peptide maturase, XyeB family sulfatase\_rSAM: anaerobic sulfatase maturase rSAM\_GlyRichRpt: radical SAM/SPASM domain protein, GRRM system quino\_hemo\_SAM: quinohemoprotein amine dehydrogenase maturation protein | 3.60E-107 4.00E-52 4.30E-46 3.20E-32 1.00E-30 |
| MTH36432.1 | 9440 | 8429 | - | 336 | TIGR04543 PF13649 TIGR00091 PF13847 PF05175 | TIGR04543 Methyltransf\_25 TIGR00091 Methyltransf\_31 MTS | ketoArg\_3Met: 2-ketoarginine methyltransferase Methyltransferase domain TIGR00091: tRNA (guanine-N(7)-)-methyltransferase Methyltransferase domain Methyltransferase small domain | 3.10E-12 1.20E-07 4.50E-06 6.10E-05 6.50E-05 |
| MTH36433.1 | 11401 | 9436 | - | 654 | TIGR02026 TIGR03471 TIGR04014 TIGR03975 TIGR04367 | TIGR02026 TIGR03471 TIGR04014 TIGR03975 TIGR04367 | BchE: magnesium-protoporphyrin IX monomethyl ester anaerobic oxidative cyclase HpnJ: hopanoid biosynthesis associated radical SAM protein HpnJ B12\_SAM\_MJ\_0865: B12-binding domain/radical SAM domain protein, MJ\_0865 family rSAM\_ocin\_1: ribosomal peptide maturation radical SAM protein 1 HpnR\_B12\_rSAM: hopanoid C-3 methylase HpnR | 5.70E-37 3.20E-36 7.00E-31 2.40E-25 1.90E-24 |
| MTH36434.1 | 12216 | 11397 | - | 272 | NO PFAM MATCH | - | - | - |
| MTH36435.1 | 13142 | 12212 | - | 309 | PF05114 | DUF692 | Protein of unknown function (DUF692) | 8.40E-38 |
| MTH36436.1 | 13345 | 13159 | - | 61 | NO PFAM MATCH | - | - | - |
| WP\_008335259.1 | 13626 | 13959 | + | 111 | INFERRED GENE | - | - | - |
| MTH36437.1 | 14036 | 15119 | + | 360 | PF05598 PF01609 PF13586 | DUF772 DDE\_Tnp\_1 DDE\_Tnp\_1\_2 | Transposase domain (DUF772) Transposase DDE domain Transposase DDE domain | 2.30E-21 2.10E-18 7.50E-04 |
| MTH36438.1 | 15784 | 17236 | + | 483 | TIGR03962 TIGR04250 TIGR04545 TIGR04051 TIGR04403 | TIGR03962 TIGR04250 TIGR04545 TIGR04051 TIGR04403 | mycofact\_rSAM: mycofactocin radical SAM maturase SCM\_rSAM\_ScmE: SynChlorMet cassette radical SAM/SPASM protein ScmE rSAM\_ahbD\_hemeb: heme b synthase rSAM\_NirJ: heme d1 biosynthesis radical SAM protein NirJ rSAM\_skfB: sporulation killing factor system radical SAM maturase | 2.00E-33 3.00E-31 1.70E-29 3.20E-25 1.80E-24 |
| MTH36439.1 | 17492 | 18326 | + | 277 | NO PFAM MATCH | - | - | - |
| MTH36440.1 | 19256 | 19499 | + | 80 | TIGR03070 PF13560 PF01381 TIGR02612 | TIGR03070 HTH\_31 HTH\_3 TIGR02612 | couple\_hipB: transcriptional regulator, y4mF family Helix-turn-helix domain Helix-turn-helix mob\_myst\_A: mobile mystery protein A | 7.00E-06 8.70E-06 1.40E-04 3.00E-04 |
| MTH36441.1 | 19701 | 19980 | + | 92 | TIGR01764 PF12728 | TIGR01764 HTH\_17 | excise: DNA binding domain, excisionase family Helix-turn-helix domain | 2.90E-04 5.50E-04 |
| MTH36442.1 | 20267 | 20093 | - | 57 | NO PFAM MATCH | - | - | - |

#### Results for WP\_170294018.1 [Paracoccus limosus] back to top

#### previous - next

##### Architecture

1000 nucleotides

Link to nucleotide sequence  
  

| Accession | start | end | direction | length (aa) | Pfam/HMM | name | description | E-value |
| --- | --- | --- | --- | --- | --- | --- | --- | --- |
| WP\_017998486.1 | 126 | 603 | + | 159 | INFERRED GENE | - | - | - |
| WP\_013654520.1 | 599 | 1034 | + | 145 | INFERRED GENE | - | - | - |
| WP\_155065942.1 | 1043 | 5009 | + | 1321 | PF13547 PF13550 | GTA\_TIM Phage-tail\_3 | GTA TIM-barrel-like domain Putative phage tail protein | 1.80E-151 3.70E-39 |
| WP\_155065943.1 | 5019 | 6945 | + | 641 | PF10983 | DUF2793 | Protein of unknown function (DUF2793) | 8.80E-34 |
| WP\_170294017.1 | 7255 | 8452 | + | 398 | TIGR04269 TIGR04496 TIGR03942 TIGR04261 TIGR03906 | TIGR04269 TIGR04496 TIGR03942 TIGR04261 TIGR03906 | SAM\_SPASM\_FxsB: radical SAM/SPASM domain protein, FxsB family rSAM\_XyeB: radical SAM/SPASM domain peptide maturase, XyeB family sulfatase\_rSAM: anaerobic sulfatase maturase rSAM\_GlyRichRpt: radical SAM/SPASM domain protein, GRRM system quino\_hemo\_SAM: quinohemoprotein amine dehydrogenase maturation protein | 4.20E-107 4.20E-52 4.10E-46 3.00E-32 1.40E-30 |
| WP\_155065945.1 | 9440 | 8429 | - | 336 | TIGR04543 PF13649 TIGR00091 PF13847 PF05175 | TIGR04543 Methyltransf\_25 TIGR00091 Methyltransf\_31 MTS | ketoArg\_3Met: 2-ketoarginine methyltransferase Methyltransferase domain TIGR00091: tRNA (guanine-N(7)-)-methyltransferase Methyltransferase domain Methyltransferase small domain | 3.10E-12 1.20E-07 4.50E-06 6.10E-05 6.50E-05 |
| WP\_246175382.1 | 11392 | 9436 | - | 651 | TIGR02026 TIGR03471 TIGR04014 TIGR03975 TIGR04367 | TIGR02026 TIGR03471 TIGR04014 TIGR03975 TIGR04367 | BchE: magnesium-protoporphyrin IX monomethyl ester anaerobic oxidative cyclase HpnJ: hopanoid biosynthesis associated radical SAM protein HpnJ B12\_SAM\_MJ\_0865: B12-binding domain/radical SAM domain protein, MJ\_0865 family rSAM\_ocin\_1: ribosomal peptide maturation radical SAM protein 1 HpnR\_B12\_rSAM: hopanoid C-3 methylase HpnR | 6.00E-37 3.10E-36 6.90E-31 2.40E-25 2.10E-24 |
| WP\_155065947.1 | 12216 | 11397 | - | 272 | NO PFAM MATCH | - | - | - |
| WP\_170294018.1 | 13097 | 12212 | - | 294 | PF05114 | DUF692 | Protein of unknown function (DUF692) | 1.80E-37 |
| WP\_155065949.1 | 13345 | 13159 | - | 61 | NO PFAM MATCH | - | - | - |
| WP\_076611756.1 | 13453 | 13570 | + | 39 | INFERRED GENE | - | - | - |
| GL279\_RS19630 | 13641 | 13959 | + | 106 | INFERRED GENE | - | - | - |
| WP\_155065950.1 | 14036 | 15119 | + | 360 | PF05598 PF01609 PF13586 | DUF772 DDE\_Tnp\_1 DDE\_Tnp\_1\_2 | Transposase domain (DUF772) Transposase DDE domain Transposase DDE domain | 2.30E-21 2.10E-18 7.50E-04 |
| WP\_170294019.1 | 15453 | 15855 | + | 133 | NO PFAM MATCH | - | - | - |
| WP\_170294020.1 | 15841 | 17236 | + | 464 | TIGR03962 TIGR04250 TIGR04545 TIGR04051 TIGR04403 | TIGR03962 TIGR04250 TIGR04545 TIGR04051 TIGR04403 | mycofact\_rSAM: mycofactocin radical SAM maturase SCM\_rSAM\_ScmE: SynChlorMet cassette radical SAM/SPASM protein ScmE rSAM\_ahbD\_hemeb: heme b synthase rSAM\_NirJ: heme d1 biosynthesis radical SAM protein NirJ rSAM\_skfB: sporulation killing factor system radical SAM maturase | 1.80E-33 2.50E-31 1.50E-29 3.00E-25 1.60E-24 |
| WP\_155065952.1 | 17492 | 18326 | + | 277 | NO PFAM MATCH | - | - | - |
| WP\_281348998.1 | 19083 | 18951 | - | 43 | NO PFAM MATCH | - | - | - |

#### Results for MDH5676845.1 [Myxococcales bacterium] back to top

#### previous - next

##### Architecture

1000 nucleotides

Link to nucleotide sequence  
  

| Accession | start | end | direction | length (aa) | Pfam/HMM | name | description | E-value |
| --- | --- | --- | --- | --- | --- | --- | --- | --- |
| MDH5676843.1 | 312 | 2616 | + | 767 | PF03712 | Cu2\_monoox\_C | Copper type II ascorbate-dependent monooxygenase, C-terminal domain | 2.00E-06 |
| MDH5676844.1 | 2962 | 3100 | + | 45 | NO PFAM MATCH | - | - | - |
| MDH5676845.1 | 3111 | 3657 | + | 181 | PF05114 | DUF692 | Protein of unknown function (DUF692) | 2.00E-18 |
| MDH5676846.1 | 4117 | 4528 | + | 136 | NO PFAM MATCH | - | - | - |
| MDH5676847.1 | 4530 | 5250 | + | 239 | NO PFAM MATCH | - | - | - |
| MDH5676848.1 | 5233 | 7138 | + | 634 | TIGR03942 TIGR03906 TIGR04463 TIGR03978 TIGR04163 | TIGR03942 TIGR03906 TIGR04463 TIGR03978 TIGR04163 | sulfatase\_rSAM: anaerobic sulfatase maturase quino\_hemo\_SAM: quinohemoprotein amine dehydrogenase maturation protein rSAM\_vs\_C\_rich: radical SAM/SPASM domain protein maturase rSAM\_paired\_1: His-Xaa-Ser system radical SAM maturase HxsB rSAM\_cobopep: peptide-modifying radical SAM enzyme CbpB | 4.40E-26 1.90E-19 6.20E-15 8.10E-15 1.20E-14 |
| MDH5676849.1 | 7708 | 7288 | - | 139 | NO PFAM MATCH | - | - | - |
| MDH5676850.1 | 8690 | 7700 | - | 329 | NO PFAM MATCH | - | - | - |
| MDH5676851.1 | 9232 | 8751 | - | 159 | NO PFAM MATCH | - | - | - |

#### Results for WP\_083412975.1 [Paracoccus aminovorans] back to top

#### previous - next

##### Architecture

1000 nucleotides

Link to nucleotide sequence  
  

| Accession | start | end | direction | length (aa) | Pfam/HMM | name | description | E-value |
| --- | --- | --- | --- | --- | --- | --- | --- | --- |
| WP\_231964698.1 | 1379593 | 1379143 | - | 149 | PF05521 | Phage\_H\_T\_join | Phage head-tail joining protein | 8.80E-06 |
| WP\_074970890.1 | 1380034 | 1381252 | + | 405 | TIGR00937 PF02417 | TIGR00937 Chromate\_transp | 2A51: chromate efflux transporter Chromate transporter | 2.10E-99 1.00E-68 |
| WP\_090271472.1 | 1381891 | 1383391 | + | 499 | PF00665 | rve | Integrase core domain | 8.60E-10 |
| WP\_028030902.1 | 1383390 | 1384119 | + | 242 | PF01695 TIGR00362 TIGR03420 PF00308 TIGR03499 | IstB\_IS21 TIGR00362 TIGR03420 Bac\_DnaA TIGR03499 | IstB-like ATP binding protein DnaA: chromosomal replication initiator protein DnaA DnaA\_homol\_Hda: DnaA regulatory inactivator Hda Bacterial dnaA protein FlhF: flagellar biosynthesis protein FlhF | 5.40E-75 3.30E-09 1.10E-07 1.60E-06 7.40E-06 |
| WP\_074971111.1 | 1384895 | 1385132 | + | 78 | NO PFAM MATCH | - | - | - |
| WP\_074971109.1 | 1385142 | 1387038 | + | 631 | TIGR03975 TIGR04367 TIGR04013 TIGR03471 TIGR04014 | TIGR03975 TIGR04367 TIGR04013 TIGR03471 TIGR04014 | rSAM\_ocin\_1: ribosomal peptide maturation radical SAM protein 1 HpnR\_B12\_rSAM: hopanoid C-3 methylase HpnR B12\_SAM\_MJ\_1487: B12-binding domain/radical SAM domain protein, MJ\_1487 family HpnJ: hopanoid biosynthesis associated radical SAM protein HpnJ B12\_SAM\_MJ\_0865: B12-binding domain/radical SAM domain protein, MJ\_0865 family | 9.40E-168 1.60E-11 2.40E-09 3.50E-09 3.20E-08 |
| WP\_074971107.1 | 1387013 | 1387817 | + | 267 | PF05721 | PhyH | Phytanoyl-CoA dioxygenase (PhyH) | 7.70E-25 |
| WP\_074971105.1 | 1387922 | 1388108 | + | 61 | NO PFAM MATCH | - | - | - |
| WP\_083412975.1 | 1388124 | 1389054 | + | 309 | PF05114 | DUF692 | Protein of unknown function (DUF692) | 1.90E-39 |
| WP\_145981073.1 | 1389050 | 1389869 | + | 272 | PF11590 | DNAPolymera\_Pol | DNA polymerase catalytic subunit Pol | 2.70E-04 |
| WP\_139218153.1 | 1389865 | 1391830 | + | 654 | TIGR03471 TIGR02026 TIGR04014 TIGR04479 TIGR04367 | TIGR03471 TIGR02026 TIGR04014 TIGR04479 TIGR04367 | HpnJ: hopanoid biosynthesis associated radical SAM protein HpnJ BchE: magnesium-protoporphyrin IX monomethyl ester anaerobic oxidative cyclase B12\_SAM\_MJ\_0865: B12-binding domain/radical SAM domain protein, MJ\_0865 family bcpD\_PhpK\_rSAM: radical SAM P-methyltransferase, PhpK family HpnR\_B12\_rSAM: hopanoid C-3 methylase HpnR | 1.90E-37 1.70E-35 1.40E-26 1.40E-26 1.80E-26 |
| WP\_170848994.1 | 1391862 | 1392834 | + | 323 | TIGR04543 PF13649 PF13847 TIGR00091 TIGR00740 | TIGR04543 Methyltransf\_25 Methyltransf\_31 TIGR00091 TIGR00740 | ketoArg\_3Met: 2-ketoarginine methyltransferase Methyltransferase domain Methyltransferase domain TIGR00091: tRNA (guanine-N(7)-)-methyltransferase TIGR00740: tRNA (cmo5U34)-methyltransferase | 7.80E-12 4.30E-09 1.80E-08 2.40E-07 1.20E-05 |
| WP\_083412974.1 | 1394032 | 1392811 | - | 406 | TIGR04269 TIGR04496 TIGR03942 TIGR04261 TIGR03906 | TIGR04269 TIGR04496 TIGR03942 TIGR04261 TIGR03906 | SAM\_SPASM\_FxsB: radical SAM/SPASM domain protein, FxsB family rSAM\_XyeB: radical SAM/SPASM domain peptide maturase, XyeB family sulfatase\_rSAM: anaerobic sulfatase maturase rSAM\_GlyRichRpt: radical SAM/SPASM domain protein, GRRM system quino\_hemo\_SAM: quinohemoprotein amine dehydrogenase maturation protein | 1.20E-101 3.00E-49 4.80E-47 7.00E-31 6.40E-29 |
| WP\_231964699.1 | 1394457 | 1394133 | - | 107 | NO PFAM MATCH | - | - | - |
| WP\_090271472.1 | 1394585 | 1396085 | + | 499 | PF00665 | rve | Integrase core domain | 8.60E-10 |
| WP\_028030902.1 | 1396084 | 1396813 | + | 242 | PF01695 TIGR00362 TIGR03420 PF00308 TIGR03499 | IstB\_IS21 TIGR00362 TIGR03420 Bac\_DnaA TIGR03499 | IstB-like ATP binding protein DnaA: chromosomal replication initiator protein DnaA DnaA\_homol\_Hda: DnaA regulatory inactivator Hda Bacterial dnaA protein FlhF: flagellar biosynthesis protein FlhF | 5.40E-75 3.30E-09 1.10E-07 1.60E-06 7.40E-06 |
| WP\_074971043.1 | 1396940 | 1397342 | + | 133 | NO PFAM MATCH | - | - | - |

#### Results for MDI9342369.1 [Sediminibacterium sp.] back to top

#### previous - next

##### Architecture

1000 nucleotides

Link to nucleotide sequence  
  

| Accession | start | end | direction | length (aa) | Pfam/HMM | name | description | E-value |
| --- | --- | --- | --- | --- | --- | --- | --- | --- |
| MDI9342361.1 | 4123 | 5947 | + | 607 | PF04024 TIGR02978 | PspC TIGR02978 | PspC domain phageshock\_pspC: phage shock protein C | 8.30E-25 8.00E-18 |
| MDI9342362.1 | 6224 | 6044 | - | 59 | NO PFAM MATCH | - | - | - |
| MDI9342363.1 | 7609 | 6274 | - | 444 | TIGR00797 PF01554 PF01943 PF14667 | TIGR00797 MatE Polysacc\_synt Polysacc\_synt\_C | matE: MATE efflux family protein MatE Polysaccharide biosynthesis protein Polysaccharide biosynthesis C-terminal domain | 4.10E-73 7.90E-50 6.60E-08 1.40E-07 |
| MDI9342364.1 | 7651 | 8257 | + | 201 | PF13640 PF13661 | 2OG-FeII\_Oxy\_3 2OG-FeII\_Oxy\_4 | 2OG-Fe(II) oxygenase superfamily 2OG-Fe(II) oxygenase superfamily | 3.30E-20 1.50E-14 |
| MDI9342365.1 | 8318 | 10367 | + | 682 | TIGR02278 PF00171 TIGR03216 TIGR01804 TIGR02299 | TIGR02278 Aldedh TIGR03216 TIGR01804 TIGR02299 | PaaN-DH: phenylacetic acid degradation protein paaN Aldehyde dehydrogenase family OH\_muco\_semi\_DH: 2-hydroxymuconic semialdehyde dehydrogenase BADH: betaine-aldehyde dehydrogenase HpaE: 5-carboxymethyl-2-hydroxymuconate semialdehyde dehydrogenase | 1.50E-275 3.70E-55 9.50E-31 3.50E-27 3.00E-26 |
| MDI9342366.1 | 10403 | 11018 | + | 204 | PF05154 | TM2 | TM2 domain | 5.40E-18 |
| MDI9342367.1 | 11134 | 11344 | + | 69 | PF10825 | DUF2752 | Protein of unknown function (DUF2752) | 8.30E-15 |
| MDI9342368.1 | 11580 | 11880 | + | 99 | NO PFAM MATCH | - | - | - |
| MDI9342369.1 | 11904 | 12774 | + | 289 | PF05114 | DUF692 | Protein of unknown function (DUF692) | 3.90E-42 |
| MDI9342370.1 | 12763 | 13543 | + | 259 | NO PFAM MATCH | - | - | - |
| MDI9342371.1 | 13547 | 15428 | + | 626 | TIGR03471 TIGR02026 TIGR04014 TIGR04479 TIGR03975 | TIGR03471 TIGR02026 TIGR04014 TIGR04479 TIGR03975 | HpnJ: hopanoid biosynthesis associated radical SAM protein HpnJ BchE: magnesium-protoporphyrin IX monomethyl ester anaerobic oxidative cyclase B12\_SAM\_MJ\_0865: B12-binding domain/radical SAM domain protein, MJ\_0865 family bcpD\_PhpK\_rSAM: radical SAM P-methyltransferase, PhpK family rSAM\_ocin\_1: ribosomal peptide maturation radical SAM protein 1 | 6.30E-45 3.40E-43 6.40E-33 4.00E-32 4.30E-29 |
| MDI9342372.1 | 15446 | 16415 | + | 322 | PF13847 TIGR04543 PF13649 TIGR01934 TIGR04074 | Methyltransf\_31 TIGR04543 Methyltransf\_25 TIGR01934 TIGR04074 | Methyltransferase domain ketoArg\_3Met: 2-ketoarginine methyltransferase Methyltransferase domain MenG\_MenH\_UbiE: ubiquinone/menaquinone biosynthesis methyltransferase bacter\_Hen1: 3' terminal RNA ribose 2'-O-methyltransferase Hen1 | 4.60E-15 4.10E-14 1.40E-10 6.30E-09 1.10E-07 |
| MDI9342373.1 | 16445 | 17873 | + | 475 | PF00515 PF07719 TIGR02521 PF13432 TIGR02917 | TPR\_1 TPR\_2 TIGR02521 TPR\_16 TIGR02917 | Tetratricopeptide repeat Tetratricopeptide repeat type\_IV\_pilW: type IV pilus biogenesis/stability protein PilW Tetratricopeptide repeat PEP\_TPR\_lipo: putative PEP-CTERM system TPR-repeat lipoprotein | 3.20E-14 2.00E-13 1.70E-12 8.90E-12 7.10E-11 |
| MDI9342374.1 | 19685 | 18074 | - | 536 | PF00082 TIGR03921 TIGR03895 | Peptidase\_S8 TIGR03921 TIGR03895 | Subtilase family T7SS\_mycosin: type VII secretion-associated serine protease mycosin protease\_PatA: cyanobactin maturation protease, PatA/PatG family | 8.50E-56 2.70E-43 1.60E-20 |
| MDI9342375.1 | 21171 | 19800 | - | 456 | TIGR01137 TIGR01136 TIGR01139 TIGR01138 TIGR03945 | TIGR01137 TIGR01136 TIGR01139 TIGR01138 TIGR03945 | cysta\_beta: cystathionine beta-synthase cysKM: cysteine synthase cysK: cysteine synthase A cysM: cysteine synthase B PLP\_SbnA\_fam: 2,3-diaminopropionate biosynthesis protein SbnA | 3.90E-147 9.70E-90 4.20E-84 3.50E-77 2.70E-74 |
| MDI9342376.1 | 22145 | 21290 | - | 284 | PF00682 TIGR02146 TIGR01108 TIGR03217 TIGR00977 | HMGL-like TIGR02146 TIGR01108 TIGR03217 TIGR00977 | HMGL-like LysS\_fung\_arch: homocitrate synthase oadA: oxaloacetate decarboxylase alpha subunit 4OH\_2\_O\_val\_ald: 4-hydroxy-2-oxovalerate aldolase citramal\_synth: citramalate synthase | 6.10E-23 2.60E-10 5.90E-08 1.40E-06 2.10E-06 |
| MDI9342377.1 | 22760 | 22214 | - | 181 | NO PFAM MATCH | - | - | - |


### RODEO2

##### Parameters

|  |  |
| --- | --- |
| Run Time | 06:58PM on October 26, 2023 |
| Version | 2.3.3 |
| Gene Window | +/-8 CDS |
| Peptide Range | 30-300 aa |
| Fetch Distance | 150bp |
| Peptide Type | general |

Annotation Legend

| Appearance | Accession/Name |
| --- | --- |

##### Input Queries (click to navigate)

- TAY82299.1

#### Results for TAY82299.1 [Rhizobium leguminosarum] back to top

#### previous - next

##### Architecture

1000 nucleotides

Link to nucleotide sequence  
  

| Accession | start | end | direction | length (aa) | Pfam/HMM | name | description | E-value |
| --- | --- | --- | --- | --- | --- | --- | --- | --- |
| TAY81972.1 | 4999636 | 5000659 | + | 340 | PF02653 | BPD\_transp\_2 | Branched-chain amino acid transport system / permease component | 6.80E-46 |
| TAY81973.1 | 5000805 | 5001753 | + | 315 | PF02955 PF02951 PF08443 PF14398 | GSH-S\_ATP GSH-S\_N RimK ATPgrasp\_YheCD | Prokaryotic glutathione synthetase, ATP-grasp domain Prokaryotic glutathione synthetase, N-terminal domain RimK-like ATP-grasp domain YheC/D like ATP-grasp | 1.50E-71 1.50E-40 8.30E-17 1.50E-05 |
| TAY81974.1 | 5002075 | 5003731 | + | 551 | PF00239 PF07508 PF13408 | Resolvase Recombinase Zn\_ribbon\_recom | Resolvase, N terminal domain Recombinase Recombinase zinc beta ribbon domain | 1.10E-19 5.80E-11 5.30E-08 |
| TAY81975.1 | 5004187 | 5005096 | + | 302 | NO PFAM MATCH | - | - | - |
| TAY81976.1 | 5007012 | 5005092 | - | 639 | PF02384 PF13847 PF01170 | N6\_Mtase Methyltransf\_31 UPF0020 | N-6 DNA Methylase Methyltransferase domain Putative RNA methylase family UPF0020 | 5.90E-29 1.50E-08 4.90E-07 |
| TAY81977.1 | 5007931 | 5007298 | - | 210 | PF00239 PF07508 | Resolvase Recombinase | Resolvase, N terminal domain Recombinase | 8.40E-24 1.70E-04 |
| TAY81978.1 | 5010204 | 5008224 | - | 659 | PF04055 PF02310 | Radical\_SAM B12-binding | Radical SAM superfamily B12 binding domain | 4.10E-19 4.50E-13 |
| TAY81979.1 | 5011017 | 5010222 | - | 264 | RREFam006 | PqqD\_RRE | RRE-containing protein in a pyrroloquinoline cluster | 8.60E-05 |
| TAY82299.1 | 5011876 | 5011006 | - | 289 | PF05114 PF01261 | DUF692 AP\_endonuc\_2 | Protein of unknown function (DUF692) Xylose isomerase-like TIM barrel | 2.10E-40 2.60E-04 |
| TAY81980.1 | 5013633 | 5012157 | - | 491 | PF00067 | p450 | Cytochrome P450 | 2.90E-35 |
| TAY81981.1 | 5013777 | 5014791 | + | 337 | PF01370 | Epimerase | NAD dependent epimerase/dehydratase family | 7.70E-09 |
| TAY81982.1 | 5015308 | 5014912 | - | 131 | NO PFAM MATCH | - | - | - |
| TAY82300.1 | 5015809 | 5015326 | - | 160 | PF03733 | YccF | Inner membrane component domain | 5.10E-32 |
| TAY81983.1 | 5015938 | 5016313 | + | 124 | PF01381 PF13560 PF12844 | HTH\_3 HTH\_31 HTH\_19 | Helix-turn-helix Helix-turn-helix domain Helix-turn-helix domain | 1.70E-14 6.00E-10 7.90E-09 |
| TAY81984.1 | 5016359 | 5019182 | + | 940 | PF02399 PF01807 | Herpes\_ori\_bp zf-CHC2 | Origin of replication binding protein CHC2 zinc finger | 1.90E-10 3.70E-04 |
| TAY81985.1 | 5019759 | 5019390 | - | 122 | PF20346 | DUF6641 | Family of unknown function (DUF6641) | 3.60E-06 |
| TAY81986.1 | 5019978 | 5020437 | + | 152 | NO PFAM MATCH | - | - | - |
