## Supporting Information for "Biosynthesis of macrocyclic peptides with C-terminal β-amino-α-keto acid groups by three different metalloenzymes"

[1]Department of Chemistry, [2]Carl R. Woese Institute for Genomic Biology, University of Illinois at Urbana-Champaign, Urbana, Illinois, 61801, USA, [3]School of Chemical Sciences NMR Laboratory, University of Illinois at Urbana-Champaign, Urbana, 61801, IL, USA, [4]School of Chemical Sciences George L. Clark X-Ray Facility and 3M Materials Laboratory, University of Illinois at Urbana-Champaign, Urbana, 61801, IL, USA, [5]Materials Research Laboratory, University of Illinois at Urbana-Champaign, Urbana, 61801, IL, USA

* To whom correspondence should be addressed.

### Supplementary Methods

**Molecular cloning**. All PCR primers were purchased from Integrated DNA Technologies Inc. A primer table can be found in Supplementary Dataset 2. Genes optimized for *Escherichia coli* expression were purchased from Twist Bioscience. Genes used for *Burkholderia.* sp FERM BP-3421 expression were amplified from the genomic DNA of *Burkholderia thailandensis* E264, after extraction using Qiagen gDNA Isolation Kits. The sequences of all genes are provided in Tables S1-S3.

Except for the linearized pRSF backbone generated using BamHI and XhoI (NEB), all DNA fragments were generated by PCR using Q5 polymerase (NEB) following the recommended protocol of the manufacturer. Specifically, 5 cycles of 98 °C denaturation (10 s), annealing (30 s), and 72 °C extension (30 s) were performed with annealing temperature calculated by the NEB $T_m$ calculator. Then, 30 cycles of 98 °C denaturation (10 s), 72 °C annealing (30 s), and 72 °C extension (30 s) were performed. All DNA fragments prior to assembly were purified using agarose gel electrophoresis [0.7% (w/v)] followed by gel extraction (GeneJET). A table summarizing ligation pieces for each construct can be found in Supplementary Dataset 2.

The vectors and inserts were then ligated using Gibson Assembly Master Mix (NEB) at 50 °C for 1 h. The reaction mixtures were then used directly to transform chemically competent *E. coli* DH5α cells, which were then placed on Luria-Bertani-Miller (LB-Miller, 10 g/L tryptone, 5 g/L yeast extract, 10 g/L NaCl) agar containing 50 µg/mL kanamycin (for pRSF-based constructs) or 50 µg/mL trimethoprim (for pSCrhaB2-based constructs). All antibiotics were purchased from Gold Biotechnology Inc. (GoldBio, St. Louis, MO). The cells were grown at 37 °C, and random colonies were picked for overnight cultures (12–16 h, 37 °C, 220 rpm) in LB medium. Plasmids were isolated using the GeneJET Plasmid Miniprep Kit and subjected to Sanger sequencing (UIUC Roy J. Carver Biotechnology Center) or whole plasmid sequencing (Plasmidsaurus).

**Expression in *E. coli***. The pRSF constructs co-expressing ApyA with modifying enzymes were used to transform chemically competent *E. coli* BL21(DE3) and placed on an LB-Miller agar plate containing 50 µg/mL kanamycin. Constructs containing *apyD* were used for co-transformation with a pCDF-based plasmid overexpressing the B12-uptake operon *btu* (*btuB*, *btuC*, *btuD*, *btuE*, *btuF*),[1] and placed on LB-Miller agar plate containing 50 µg/mL kanamycin and 50 µg/mL streptomycin. Post-transformation, the plates were incubated at 37 °C for 12–15 h. For each expression, a few colonies were picked for an overnight culture (12 h, 37 °C, 220 rpm) in LB-Miller broth containing the same amount of antibiotics used in the LB-Miller plate. The cultures were then diluted at a 1:100 (v/v) ratio in terrific broth (24 g/L yeast extract, 12 g/L tryptone, 0.4% glycerol (v/v), 17 mM $KH_2PO_4$ and 72 mM $K_2HPO_4$) containing the same antibiotic concentrations and grown at 37 °C with 200 rpm shaking.

For constructs containing *apyD*, the cultures were grown in a baffled 2.8 L Fernbach flask (1 L in each flask) to an optical density at 600 nm ($OD_{600}$) of 0.6–1.0. At this point, 0.2 % (w/v) L-arabinose (GoldBio) and 2 µM of vitamin B12a were added (hydroxocobalamin, Sigma) to induce *btu* expression. The cultures then continued growing at 30 °C and 200 rpm shaking until $OD_{600}$ 1.8–3.0. Isopropyl ß-D-1-thiogalactopyranoside (IPTG, 0.5 mM final concentration) was added to induce the expression of genes from the pRSF construct. Iron(II) citrate [final concentration 255 μM; stock: 100 mg/mL iron(II) ammonium sulfate heptahydrate (Sigma) mixed with 1 M sodium citrate (Fisher Scientific)] and 200 μM L-cysteine (Sigma) were also added during IPTG induction to support the function of the B12-rSAM ApyD. The cultures were grown at 18 °C and 200 rpm for 18–20 h before collection.

For constructs containing ApyO, the cultures were grown to OD of 1.8 - 3. IPTG (0.5 mM) was then added to induce gene expression. 5-Aminolevulinic acid (1 mM; GoldBio) was added to support the biosynthesis of heme[2] along with 255 μM iron (II) citrate. The cultures then continued to grow at 18 °C and 200 rpm for 18–20 h before collection. For constructs encoding both ApyD and ApyO, the expression followed the same protocol as constructs containing ApyD, except that 1 mM 5-aminolevulinic acid was also added during IPTG induction. The cells were collected by centrifugation at 4500 × g for 15 min. The supernatant was discarded, and the cell pellet was frozen and stored at -80 °C until needed.

**Expression in *B.* sp FERM BP-3421**. A glycerol stock [20% (v/v)] of *B.* sp FERM BP-3421 fr9DEF– was revived on a Luria-Bertani-Lennox (LB-Lennox 10 g/L tryptone, 5 g/L yeast extract, 5 g/L NaCl) agar plate at 30 °C for 24 h. A single colony was picked and grown overnight at 30 °C in LB-Lennox liquid media for 24 h. The cells were rendered electrocompetent following a previously reported protocol.[3] Specifically, the cell solution was split into 1.5 mL Eppendorf tubes and cells were harvested by centrifugation at 16000 × g for 2 min. The supernatant was discarded, and each tube was washed with 1 mL of 300 mM sucrose (Sigma) twice. After washing, the cells were resuspended in 100 μL of 300 mM sucrose, and 100–500 ng of pSCrhaB2 plasmid[4] containing the desired insert (Tables S2 and S3) was added.

The resulting solution was then placed into a 0.1 cm-gap Gene Pulser Electroporation Cuvette and electroporated using a Gene Pulser Xcell electroporator (Biorad) using the following condition: C = 25 μF; PC = 200 ohm; V = 1.8 kV. Then 1 mL of LB-Lennox was added, and the cells were rescued at 30 °C for 1 h. Afterwards, the cells were collected at 16000 × g for 2 min, and the supernatant was discarded until there was 100 μL of solution remaining. The cells were then resuspended, plated on an LB-Lennox Agar plate containing 120 μg/mL trimethoprim, and grown at 30 °C and 200 rpm shaking for 22–26 h.

A single colony was then grown in LB-Lennox liquid media containing 120 μg/mL trimethoprim at 30 °C and 200 rpm shaking for 22–26 h. The overnight culture was diluted at a 1:100 (v/v) ratio to 1 L of 2S4G media containing: 20 g/L phytone peptone (Gibco), 40 g/L glycerol (Fisher Scientific), 2 g/L ammonium sulfate (Sigma), 2 g/L $CaCO_3$ (Sigma), and 405 μM $MgSO_4$. The culture was grown at 30 °C and 200 rpm shaking in a Thomson Ultra Yield 2.5 L flask enclosed with a vent cap containing a 0.2 μm filter. When the $OD_{600}$ reached 0.6–1.0, 0.2% (w/v) of L-rhamnose (ChemImpex) was added to induce gene expression. The cells continued to grow at 30 °C and 200 rpm shaking for 24–26 h. This condition was used to express constructs listed in Table S2B (pSCrhaB2_*apy*OAHIDS, Figure S14), S2C (ApyO omitted), S2D (ApyH omitted), S2E (ApyI omitted), S2F (ApyS omitted), and all single mutants of precursor peptides (Supplementary Dataset 2). ApyD activity was low or undetectable except for the construct with ApyO excluded. This condition was utilized to scale up production of ApyO/H/I/S-modified ApyA.

To produce ApyD/O/H/I/S-modified ApyA, another condition was employed for constructs in Table S2B and Table S3 (pSCrhaB2_*apy*D_RBS_*apy*OAHIS, Figure S14), with the construct in Table S3 giving higher efficiency of ApyD-catalyzed methylation. First, expressions were performed with 1.5 L of 2S4G media instead of 1 L in Thomson Ultra Yield 2.5 L flasks. To facilitate ApyD activity, vitamin B12a (2 µM) was added after induction. Iron(II) citrate (255 µM) and L-cysteine (200 µM) were also supplemented. The cells continued to grow at 30 °C and 200 rpm shaking for 24–26 h.

The cell collection involved two steps, starting with centrifugation at 1000 × g for 2 min. The supernatant, which still contained the cells, was then carefully decanted not to carry over $CaCO_3$. After the removal of the majority of $CaCO_3$, the cells were collected by centrifuging at 7000 × g for 20 min. The supernatant was then discarded, and the cell pellet was frozen and stored at -80 °C before use.

**Cell lysis and Ni-NTA affinity purification.** The cells were resuspended in 50 mL of lysis buffer (6 M guanidinium hydrochloride, 50 mM phosphate, 500 mM NaCl, pH 7.5) per 10 g of estimated weight of wet cells. After resuspension, the cell solution was lysed using a high-pressure homogenizer (Avestin, Inc) at an operating pressure between 10–15 kpsi (for *E. coli)* or 15-20k psi (for *Burkholderia* sp FERM BP-3421). The homogenization was repeated 3-4 times. The supernatant was then clarified by centrifugation at 36000 × g for 1 h. The lysate was then stirred with 1 mL of Ni-NTA resin (HisPur, Thermo) per 75 mL of lysate using a magnetic stirrer for 2 h. For the product containing the α-keto acid moiety, 5 mM of sodium pyruvate (Fisher Scientific) was added to every purification buffer. The supernatant was then removed by centrifugation at 1000 × g for 2 h followed by decantation. The Ni-NTA resin was then loaded onto a column, washed with 18 column volumes (CV) of lysis buffer [with 1 mM tris(2-carboxyethyl)phosphine (TCEP)], 8 CV of wash buffer 1 (4 M guanidinium hydrochloride, 50 mM sodium phosphate, 300 mM NaCl, 30 mM imidazole, pH 7.5, 1 mM TCEP), and 8 CV of wash buffer 2 (50 mM sodium phosphate, 1M NaCl, 30 mM imidazole, pH 7.5, 1 mM TCEP). The peptide was then eluted using 6 CV of elution buffer (50 mM sodium phosphate, 300 mM NaCl, 1 M imidazole, pH 7.5, 1 mM TCEP).

**High-performance liquid chromatography (HPLC) of full-length peptides.** The eluted peptide solutions were acidified using formic acid (FA, Optima) until the pH reached ~3. The solutions were then injected into a Macherey Nagel-C18 HTec, 250 × 10 mm, 5 µm column, or a Phenomenex Luna-C5, 250 × 10 mm, 10 µm column. The flow rate of the run was 4 mL/min. Solvent A was $H_2O$ with 0.1% trifluoroacetic acid (TFA), and solvent B was $CH_3CN$ with 0.1% TFA. Gradient: 0 – 5 min: 2% B, 5 – 35 min: 2% B to 60% B, 35 – 40 min: 60% B to 90% B, 40 min to 42 min: 90% B to 2% B. After elution, the solution was then lyophilized to dryness. Lyophilization was performed using a Labconco (Kansas City, MO) freeze dryer.

**Proteolysis with endoproteinase.** The HPLC-purified peptide was dissolved in $H_2O$ after lyophilization. The concentration of the peptide was roughly estimated using dry weight or absorbance at 280 nm using a Nanodrop. Endoproteinase GluC (Worthington) or LysC (NEB) was added to the peptide in an estimated ratio of 1:20 (w/v) for GluC and 1:100 (w/v) for LysC, and the reaction was run in digestion buffer [50 mM Tris(hydroxymethyl)aminomethane (Tris), 1 mM TCEP, pH 8] at 37 °C for 12 h. For products containing the α-keto acid moiety, 20 µg/mL of catalase (Sigma) and 4 mM sodium pyruvate were added to the digestion reaction to prevent oxidative breakdown as they readily oxidatively decarboxylate.[5,6]

**HPLC of GluC-digested peptides.**

a) ApyO-modified ApyA, ApyD/ApyO-modified ApyA and ApyD/ApyO/ApyHI/ApyS-modified ApyA. After digestion, the solutions were injected into a Macherey Nagel-C18 HTec, 250 × 10 mm, 5 µm column. The flow rate of the run was 4 mL/min. Solvent A was 20 mM $NH_4OAc$, and solvent B was $CH_3CN$. Gradient: 0 – 5 min: 2% B, 5 – 35 min: 2% B to 60% B, 35 – 40 min: 60% B to 90% B, 40 min to 42 min: 90% B to 2% B. After elution, the solution was lyophilized to dryness prior to downstream application (e.g., NMR analysis).

For GluC-cleaved ApyD/ApyO-modified ApyA, prior to NMR analysis, the lyophilized product was first dissolved in a small amount of water (1 mL for products purified from 16 L of expression). The supernatant was discarded after centrifugation at 4500 × g for 15 min. The remaining pellet was dissolved in 40% $CD_3CN$/60% $H_2O$ for NMR analysis and was found to have significantly improved purity.

b) ApyD-modified ApyA. After digestion, the solutions were injected into a Macherey Nagel-C18 HTec, 250 × 10 mm, 5 µm column. The flow rate was 4 mL/min, and solvent A was 20 mM $NH_4OAc$, and solvent B was $CH_3CN$. Gradient: 0 – 5 min: 2% B, 5 – 35 min: 2% B to 60% B, 35 – 40 min: 60% B to 90% B, 40 min to 42 min: 90% B to 2% B. After elution, the solution was lyophilized to dryness. The dried powder was dissolved in $H_2O$ and subjected to another round of HPLC with the same column but with solvent A as $H_2O$ with 0.1% TFA and solvent B as $CH_3CN$ with 0.1% TFA. After elution, the solution was lyophilized to dryness prior to downstream application (e.g., NMR analysis).

c) ApyA modified with ApyO, ApyHI, and ApyS. After digestion, formic acid was used to adjust the pH to 3. The solutions were injected into a Macherey Nagel-C18 HTec, 250 × 10 mm, 5 µm column. The flow rate of the run was 4 mL/min, and solvent A was $H_2O$ with 0.1% TFA, and solvent B was $CH_3CN$ with 0.1% TFA. Gradient: 0 – 5 min: 2% B, 5 – 35 min: 2% B to 60% B, 35 – 40 min: 60% B to 90% B, 40 min to 42 min: 90% B to 2% B. After elution, the solution was lyophilized to dryness. The sample remained impure with other fragments from GluC digestion. Thus, the mixture was treated with 1 mg of trypsin (Thermo) at 37 °C for 2 h. The resulting mixture was then subjected to another round of HPLC with the same C18 column. After elution, the solution was then lyophilized to dryness prior to downstream application (e.g., NMR analysis).

**Liquid chromatography and high-resolution electrospray ionization tandem mass spectrometry (LC-HR-ESI-MS/MS) analysis of peptides.** Samples (2–20 µL) were injected into an Agilent AdvanceBio Peptide Plus column (2.1 × 150 mm, 2.7 µm), a Kinetex C18 LC column (2.1 × 150 mm, 2.6 µm, 100 Å), or an Agilent C18 Poroshell 120 column (3.0 × 100 mm, 2.7 µm) on an Agilent 6545B Q-TOF interfaced with an Agilent 1290 Infinity II LC system. Mobile phase solvent A was $H_2O$ with 0.1% formic acid (FA) and solvent B was $CH_3CN$ with 0.1% FA. The flow rate was 0.4 mL/min. The gradient was as follows: first 2-5 min 5% B, next 8 min 5% B - 70% B, last 2 min 70–90% B. The column was equilibrated with 3 min of 5% B before the next samples were injected. The samples were run to waste in the first 2-5 min (depending on the time equilibrated with 5% B) before applying to the mass spectrometer. MS parameters were as follows:

Mass range 100 to 1700 *m/z* ; 325 °C at 10 L/min; nebulizer, 35 psig; nozzle voltage, 0 V; sheath gas, 375 °C at 11 L/min; capillary, 3500 V; fragmentor, 120 V; skimmer, 65 V; MS scan rate (10 spectra/s); MS-MS scan rate (5 spectra/s); and isolation width (MS/MS), 1.3 *m/z*. Positive ionization mode was utilized for all MS experiments except Marfey's stereochemical analysis. Fragmentation was performed using collision-induced dissociation (CID) at 25 eV. The data analysis was performed using Agilent MassHunter Qualitative Analysis 10.0. The exact mass lists are exported and analyzed using IPSA and mMass.[7,8]

**Marfey's analysis of β-methyltyrosine in ApyD-modified ApyA product.** GluC-cleaved and HPLC-purified methylated ApyA (0.4 mg) was equilibrated under nitrogen gas and subjected to one freeze-pump-thaw cycle in a Schlenk flask. The sample was then frozen, put under vacuum, reagents were added (1 mL of 6 M DCl in $D_2O$ (Sigma), 1% (w/v) phenol and 3% (w/v) of 2-mercaptoethanol), the tube was sealed, and equilibrated back to room temperature. The peptide was then hydrolyzed at 100 °C for 2 h. The solvents and volatile components were removed using a SpeedVac concentrator (ThermoFisher Scientific) at 35 °C overnight. The hydrolyzed peptides were then dissolved in 150 µL of 1 M $NaHCO_3$ followed by addition of 150 µL acetone containing 4 mg/mL of L-FDAA and D-FDAA (FDAA = 1-fluoro-2-4-dinitrophenyl-5-L-alanine amide). The reactions proceeded for 2.5 h at 60 °C and were then quenched using 1 mL of 150 mM ammonium formate. The solutions were concentrated to 100 µL for LC analysis. The standards (*2S,3R*)-2-amino-3-(4-hydroxy-phenyl)-butyric acid and (*2S,3S*)-2-amino-3-(4-hydroxy-phenyl)-butyric acid (JW Pharma) were subjected to analogous hydrolysis and derivatization conditions.

Aliquots (1 µL) of the samples were injected onto an Accucore PFP column (ThermoFisher Scientific) 2.6 µm, 80 Å, 150 × 4.6 column attached to an Agilent 6545B Q-TOF interfaced with an Agilent 1290 Infinity II LC system. Mobile phase solvent A was $H_2O$ with 0.1% FA and solvent B was $CH_3CN$ with 0.1% FA. The flow rate was 1 mL/min. The gradient was as follows: 0 - 3 min 2% B, 3 – 7 min 2% B – 43% B, 7 – 16 min 30% B – 70% B, 16 – 17 min 70% B – 90% B, 17 – 18 min 90% B – 2% B. MS parameters were as follows: Mass range 100 to 1700 *m/z* ; 200 °C at 13 L/min; nebulizer, 35 psig; nozzle voltage, 1000 V; sheath gas, 350 °C at 11 L/min; capillary, 3500 V; fragmentor, 125 V; skimmer, 65 V; MS scan rate (5 spectra/s). Negative ionization mode was utilized.

**Marfey's analysis of ApyO/ApyHI/ApyS-modified ApyA product.** Full-length ApyO/ApyHI/ApyS-modified ApyA product was subjected to 1 mM $H_2O_2$ in 50 mM Tris buffer pH 7.0 for 2 h at room temperature. Then, endoproteinase GluC (Worthington) was added to the mixture at an estimated ratio of 1:20 (w/v) enzyme: substrate, and the reaction proceeded for 2 h at 37 °C. Afterwards, 10 mM of sodium pyruvate was added, and the mixture was injected into a Macherey Nagel-C18 HTec, 250 × 10 mm, 5 µm column for HPLC (with 0.1% TFA as solvent A and $CH_3CN$ 0.1% TFA as solvent B). The flow rate of the run was 4 mL/min. After elution, the solution was lyophilized to dryness. The dried powder was dissolved in $H_2O$ and subjected to a round of ultra-pressure liquid chromatography (UPLC) on a Vanquish Core HPLC System (ThermoFisher) with an Accucore aQ-C18 column (ThermoFisher Scientific) 2.6 µm, 80 Å, 150 × 4.6 column. The flow rate of the run was 1.5 mL/min, and solvent A was $H_2O$ with 0.1% FA, and solvent B was $CH_3CN$ with 0.1% FA. Gradient: 0 – 5 min: 5% B, 5 – 15 min: 5% B to 35% B, 15 – 15.1 min: 35% B to 95% B, 15.1 – 20 min: 95% B, 20 – 20.1 min: 95% B to 5% B, 20.1 –

25 min: 5% B. After elution, the solution was then lyophilized to dryness prior to Marfey's analysis.

Marfey's analysis was performed following a protocol used for lanthipeptide stereochemical analysis.[9] Specifically, after lyophilization, approximately 0.05-0.1 mg was dissolved in 0.8 mL of 6 M $DCl/D_2O$ having 1% (w/v) phenol and transferred to a Pyrex 15 mL screw cap glass tube. The sample was sparged with $N_2$ for 2 min, the tube was sealed, and the mixture was heated at 110 °C for 2 h. The resulting mixture was then immersed in a water bath of 80 °C and dried using a rotary evaporator, 1 mL of water was then added, and the rotary evaporation was repeated. After lyophilization overnight, the dried solid was dissolved in 0.6 mL of 0.8 M $NaHCO_3$ and derivatized with 0.6 mL of 3 mg/mL L-FDLA dissolved in $CH_3CN$ (FDLA = 1-fluoro-2-4-dinitrophenyl-5-L-leucine amide). The reaction was then stirred in the dark at 65 °C for 1.5 h. Then, 0.1 mL of 6 M HCl was slowly added to quench the reaction, and the mixture was frozen and lyophilized. The dried sample was then redissolved in 1 mL of $CH_3CN$ and subjected to LC-MS analysis. The standards L- and D-alanine were subjected to analogous conditions.

Aliquots of samples (1 µL) were injected onto a Kinetex 1.7 µm F5 100 Å, 100 × 2.1 mm LC column attached to an Agilent 6545B Q-TOF interfaced with an Agilent 1290 Infinity II LC system. Mobile phase solvent A was $H_2O$ with 0.1% FA, and solvent B was $CH_3CN$. The flow rate was 0.3 mL/min. The gradient was as follows: 0 - 1 min 2% B, 1 – 4 min 2% B – 40% B, 4 – 13 min 40% B – 70% B, 13– 16 min 70% B – 95% B. MS parameters were as follows: Mass range 100 to 1700 *m/z* ; 325 °C at 13 L/min; nebulizer, 35 psig; nozzle voltage, 1000 V; sheath gas, 350 °C at 11 L/min; capillary, 3500 V; fragmentor, 125 V; skimmer, 65 V; MS scan rate (5 spectra/s). Negative ionization mode was utilized.

**Purification of iodoacetamide-alkylated Cys-Ile-Glu tripeptide for NMR and Marfey's analysis.** ApyA/O/H/I/S-modified ApyA was treated with 2 mM TCEP at 37 °C for 30 min followed by alkylation at 37 °C with 30 mM iodoacetamide (IAA) for 1 h in 50 mM Tris at pH 8.0. The reaction was quenched with 20 mM dithiothreitol (DTT) prior to proteolytic digestion with GluC and HPLC purification as above. The fraction containing the alkylated Cys was lyophilized to dryness and dissolved in 50 mM Tris pH 8.0 buffer for digestion at 37 °C for 12 h using trypsin with an estimated ratio of 1:20 (w/v) enzyme: substrate. The digestion mixture was then subjected to a round of UPLC on a Vanquish Core HPLC System (ThermoFisher) with an Accucore C18 (ThermoFisher Scientific) 2.6 µm, 80 Å, 150 × 4.6 mm column. The flow rate was 1.5 mL/min, and solvent A was $H_2O$ with 0.1% FA, and solvent B was $CH_3CN$ with 0.1% FA. Gradient: 0 – 5 min: 5% B, 5 – 10 min: 5% B to 95 % B, 10 – 15 min: 95% B, 15 – 17 min: 95% B to 5% B, 17 – 27 min: 5% B. The fraction containing the IAA-alkylated Cys-Ile-Glu tripeptide was then lyophilized to dryness prior to NMR and Marfey's analysis.

The purified tripeptide was subjected to a similar Marfey's protocol for ApyO/ApyHI/ApyS-modified ApyA product as mentioned above but with L-FDAA as the derivatization reagent for 3.5 h instead of L-FDLA for 1.5 h. Aliquots of samples (1 µL) were injected onto a Kinetex 1.7 µm F5 100 Å, 100 × 2.1 mm LC column attached to an Agilent 6545B Q-TOF interfaced with an Agilent 1290 Infinity II LC system. Mobile phase solvent A was $H_2O$ with 0.1% FA, and solvent B was $CH_3CN$. The flow rate was 0.3 mL/min. The gradient was as follows: 0 - 2 min 5% B, 2 – 5 min 2% B – 40% B, 5 – 14 min 40% B – 60% B, 14 – 17 min 60% B – 80% B, 17 – 19 min 80%

B to 95% B. MS parameters were as follows: Mass range 100 to 1700 *m/z* ; 325 °C at 13 L/min; nebulizer, 35 psig; nozzle voltage, 1000 V; sheath gas, 350 °C at 11 L/min; capillary, 3500 V; fragmentor, 125 V; skimmer, 65 V; MS scan rate (5 spectra/s). Negative ionization mode was utilized. Fragmentation was performed with collision-induced dissociation at 18 eV for masses corresponding to the FDAA-derivatized alkylated Cys residue.

**Exciton-coupled circular dichroism.** Spectra were measured on a J-1500 CD system (Jasco, Inc). (*R*)-(+)- and (*S*)-(-)-5,5′,6,6′,7,7′,8,8′-octahydro-1,1′-bi-2-naphthol were from TCI America. (*R*)-(+)-5,5',6,6'-Tetramethyl-3,3'-di-t-butyl-1,1'-biphenyl-2,2'-diol was from VWR Strem Chemical Inc. (*S*)-(−)-5,5′,6,6′-Tetramethyl-3,3′-di-tert-butyl-1,1′-biphenyl-2,2′-diol was from Sigma. The standards were dissolved in spectroscopic grade MeOH (Fisher). The ApyO-modified ApyA pentapeptide was dried with a SpeedVac concentrator twice from a MeOH solution before being redissolved in MeOH as a sample for circular dichroism. The estimated concentration of standards (from dry weight) and ApyO-modified ApyA pentapeptide (by qNMR) was 250 µM. Then 200 µL of each sample was added to a 0553 Micro Rectangular Quartz Micro Cell (10 mm pathlength) for data collection. The parameters of data collection were as follows: temperature, 23 °C; measured range 400–240 nm; CD scale 200 mdeg/0.1 dOD; bandwidth 1.00 nm; scanning speed 50 nm/min; data pitch 0.2 nm; digital integration time: 1 s.

**Crystallography screening, microcrystal electron diffraction (MicroED) data collection and analysis.** Approximately 400 µL of 0.3 mM of HPLC-purified ApyD- and ApyO-modified ApyA pentapeptide was subjected to Microbatch-Under-Oil crystallization with 1,536 crystallization conditions in microtitre plates using a protocol and service offered by the National High-Throughput Crystallization Center.[10] All plates were visualized by brightfield imaging, ultraviolet two-photon excited fluorescence microscopy, and second harmonic generation microscopy to determine appropriate crystals. Conditions determined to form peptide crystals were then incrementally modified to a total of 24 conditions with variations in pH and additive concentrations. These conditions were then subjected to sitting-drop crystallization with the pentapeptide. Specifically, 2 µL of 0.3 mM ApyD- and ApyO-modified ApyA pentapeptide was added to 2 µL of the buffers (Hampton Research) for each condition in 24-well CrysChem M plates, with the reservoir filled with 500 µL of buffers. The plates were then sealed with ClearSeal Film (Hampton Research) and left for a week in areas with minimal vibration. The crystals were then visualized using a brightfield microscope, and 10 µL of 0.3 mM ApyD- and ApyO-modified ApyA pentapeptide was added to wells with crystals. The plates were then left for a week, and the crystals formed in the well with 30 mM zinc acetate and 17.86% PEG3350 were subjected to MicroED data collection.

The solution was frozen on glow discharge 200 mesh Cu TEM grids (Ted Pella Cat. No. 01801). For each grid, 3 µL of sample was applied at 22 °C and 90–95% ambient temperature and humidity, respectively. The FEI Mark IV Vitrobot was used for freezing and blotting. After applying the sample, grids were blotted once with blot force 3 and immediately plunged into liquid ethane. Once frozen, TEM grids were clipped into autogrids and loaded into a ThermoFisher Glacios CryoTEM for analysis. Data was collected at 200 kV on a ThermoFisher Ceta-D CMOS camera using the ThermoFisher Scientific Software package EPU-D (version 1.11.0.4164RE). Data was collected using the full available angle range of the ThermoFisher tilt stage (-60 to 60 degrees). Prior to data collection, the gun lenses were changed from 4 to 8 to reduce the electron current and avoid crystal damage during data collection at 77 K. Cell refinement, integration of intensity data, scaling, absorption correction, and merging of multiple data sets of two nanocrystals (electron dose rate 0.00427 $e^{-}/Å^2/s$ and 0.00712 $e^{-}/Å^2/s$) and was carried out using DIALS software.[11] The structures were phased with dual-space methods using SHELXD[12] and refined with the full-matrix least-squares program SHELXL.[13]

A structural model consisting of two complete peptide molecules, one Zn atom, one PEG fragment, and numerous water molecules in the asymmetric unit was developed; however, positions for the PEG fragment and water molecules not involved in hydrogen bonding were poorly determined. This model converged with wR2 = 0.3867 and R1 = 0.1487 for 1083 parameters with 2030 restraints against 9167 data. Since positions for the PEG fragment and non-hydrogen bonded water molecules were poorly determined a second structural model was refined with contributions from these molecules removed from the diffraction data using the solvent mask procedure in Olex2 v1.5.[14] No positions for the host network differed by more than two su's (standard uncertainty) between these two refined models. The electron count from the "squeeze" model converged in good agreement with the number of PEG/water molecules predicted by the complete refinement. 316 electrons were removed from one continuous void space in the unit cell, consistent with 79 electrons per $Zn(peptide)_2$ assembly.

Rigid-bond restraints (esd 0.004) were imposed on displacement parameters for all anisotropic atoms in the structure. Similar displacement amplitudes (esd 0.01) were imposed on all atoms in the peptide chains overlapping by less than the sum of van der Waals radii. All phenyl rings in the peptide chains were constrained to be perfect hexagons. Similarity restraints were imposed on all 1,2 and 1,3 distances of the two peptide chains in the asymmetric unit (esd 0.01, 0.02 Å). When the refinement was essentially complete, a bond distance survey was performed with Mogul[15] (CCDC, Sept 2023 database update). Several bond distances showed significant deviations from a typical bond length of that type. For these deviant bonds, the bond distance was restrained to the median value reported by the Mogul geometry check (esd 0.02 Å).

The Zn atom was modeled as disordered over two orientations with the displacement parameters of the two sites constrained to be the same. The site occupancy ratio was allowed to freely refine. The carboxylate substituents on the C-terminus of both unique peptide chains were modeled as disordered over two orientations. The two site occupancy ratios were allowed to freely refine. The water oxygen atoms that were included in the final refinement were isotropically refined.

H atom treatment - Methyl H atom positions, $R-CH_3$, were optimized by rotation about R-C bonds with idealized C-H, R--H and H--H distances. Residual density peaks consistent with the hydroxyl H atom positions were visible in the difference map. To maintain a higher data to parameter ratio, hydroxyl H atom positions, R-OH, were optimized by rotation about R-O bonds with idealized O-H and R--H distances for the final refinement. Residual density peaks consistent with H atom positions for amine atoms N2, N3, and N4 were visible in the difference map. The amine H atoms were refined as riding idealized contributors for the final refinement to maintain a higher data to parameter ratio. For N5, H atoms were initially assigned using a riding model. This resulted in two overlapping H atom positions due to the disorder of the carboxylate groups bound to N5. For the final refinement, only one of the idealized H atom positions was included and its position was allowed to freely refine with the N---H distance restrained to 0.86(2) Å. At convergence, the majority of the hydroxyl and amine H atoms were in good hydrogen bonding geometries.[16]~~5~~ Remaining H atoms were included as riding idealized contributors. Methyl, hydroxyl, and amine H atom U's were assigned as 1.5 times $U_{eq}$ of the carrier atom; remaining H atom U's were assigned as 1.2 times carrier $U_{eq}$. No attempt was made to assign H atoms to the water oxygen atoms.

*Note on charge balance*: Due to the presence of Zn, which is assumed to be in the 2+ state, each peptide chain must bear a 1- charge. No residual density peaks were visible near the N-terminus nitrogen atom or the C-terminus carboxylate oxygen atoms. In solution, the most reasonable charge distribution based on pKa considerations is a neutral N-terminus and a negatively charged C-terminus. There are two carboxylate groups at the C-terminus and it is unclear whether the negative charge is localized to one of the carboxylate groups or disordered over both. Due to this ambiguity, no H atoms were assigned to the N-terminus amine or the C-terminus carboxylate oxygen atoms. The absolute structure was assigned to be consistent with the known stereochemistry of C3, C18, C20, and C24.

One distinct cell was identified using DIALS.[11] Data from two separate crystallites were integrated and filtered for statistical outliers using DIALS v3-7-2. The co-symmetrizing routing in DIALS was used to sort, merge, and scale the combined data. Decay corrections are included as needed in the DIALS scaling engine.

The structure was phased by dual space methods.[12] Systematic conditions suggested the unambiguous space group P2(1)2(1)2(1). The space group choice was confirmed by successful convergence of the full-matrix least-squares refinement on $F^2$. The final difference Fourier map had no significant features. A final analysis of variance between observed and calculated structure factors showed little dependence on amplitude or resolution. The crystallographic data and structure refinements can be found in Table S7.

**NMR data acquisition and analysis.** Sample information: ApyD-modified ApyA pentapeptide (~ 5.5 mM) in 60% $H_2O$ and 40% $CD_3CN$, ApyO-modified ApyA pentapeptide (~4.4 mM) in 90% $H_2O$ and 10% $D_2O$, ApyD/O-modified ApyA pentapeptide (3.0 mM) in 60% $H_2O$ and 40% $CD_3CN$, and IAA-alkylated Cys-Ile-Glu tripeptide in $CD_3OH$ were dissolved in 550 μL volume and transferred into a 5-mm Wilmad 535-pp NMR tube. ApyO/H/I/S-modified ApyA pentapeptide (total ~6.0 mM, containing ~ 3.6 mM keto form, ~ 2.0 mM gem-diol form, and trace rotamers and impurities) were dissolved in 320 μL 90% $H_2O$ and 10% $D_2O$ in a Shigemi Advanced NMR microtube assembly matched with $D_2O$ (Sigma, bottom L 8 mM). NMR data were collected at 25 °C on an Agilent VNMRS 750 MHz NMR spectrometer equipped with an RT indirect detection 5-mm HCN probe or a Bruker Avance NEO 600 MHz spectrometer equipped with a 5-mm BBO prodigy probe. One-dimensional (1D) $^1$H NMR, two-dimensional (2D) homonuclear $^1$H-$^1$H COSY (correlation spectroscopy which reveals the correlation between two neighboring protons), $^1$H-$^1$H TOCSY (total correlation spectroscopy which reveals the correlation of protons in the same spin system) at 80 ms mixing time, $^1$H-$^1$H NOESY (Nuclear Overhauser Effect spectroscopy which reveals close proximity in space between two protons) at 400 ms mixing time, or $^1$H-$^1$H ROESY at 300 ms mixing time (NOESY in rotating frame, revealing the close proximity in space between two protons), $^1$H-$^{13}$C HSQC (heteronuclear single quantum coherence spectroscopy, revealing one-bond correlation between $^1$H and $^{13}$C), and $^1$H-$^{13}$C HMBC (heteronuclear multiple bond correlation, revealing long range $^1$H-$^{13}$C connectivity such as 2-bond, 3-bond, or 4 or more bonds) spectra were acquired for each sample using either the Biopack pulse sequences in the VNMRJ 4.2A software or pulse programs in Bruker Topspin 4.1.3. The spectra were processed and analyzed in Mnova (version 14.3.0. Mestrelab Research). The sample concentrations were determined by $^{1}H$ spectra collected under the qNMR condition on an Agilent VNMRS 750 MHz spectrometer with a HCN probe that was calibrated with a known standard. The calibrated parameters were saved to the probe file and used to calculate the sample concentration based on the integration values of the proton peaks.

**Table S1:** Sequence of the pRSF-based vectors used in this study for expression in *E. coli*. Each gene is highlighted as follows. *apyA*: bolded black; *apyO*: pink; *apyD*: green; *apyH*: orange; *apyI*: black underlined, *apyS*: blue. The N-terminus of ApyA contains a hexa-His tag.

A) Backbone of pRSF vector

```
CTCGAGTCTGGTAAAGAAACCGCTGCTGCGAAATTTGAACGCCAGCACATGGACTCGTCTACTAGCGCAGCTTAATT
AACCTAGGCTGCTGCCACCGCTGAGCAATAACTAGCATAACCCCTTGGGGCCTCTAAACGGGTCTTGAGGGGTTTTT
TGCTGAAACCTCAGGCATTTGAGAAGCACACGGTCACACTGCTTCCGGTAGTCAATAAACCGGTAAACCAGCAATAG
ACATAAGCGGCTATTTAACGACCCTGCCCTGAACCGACGACAAGCTGACGACCGGGTCTCCGCAAGTGGCACTTTTC
GGGGAAATGTGCGCGGAACCCCTATTTGTTTATTTTTCTAAATACATTCAAATATGTATCCGCTCATGAATTAATTC
TTAGAAAAACTCATCGAGCATCAAATGAAACTGCAATTTATTCATATCAGGATTATCAATACCATATTTTTGAAAAA
GCCGTTTCTGTAATGAAGGAGAAAACTCACCGAGGCAGTTCCATAGGATGGCAAGATCCTGGTATCGGTCTGCGATT
CCGACTCGTCCAACATCAATACAACCTATTAATTTCCCCTCGTCAAAAATAAGGTTATCAAGTGAGAAATCACCATG
AGTGACGACTGAATCCGGTGAGAATGGCAAAAGTTTATGCATTTCTTTCCAGACTTGTTCAACAGGCCAGCCATTAC
GCTCGTCATCAAAATCACTCGCATCAACCAAACCGTTATTCATTCGTGATTGCGCCTGAGCGAGACGAAATACGCGG
TCGCTGTTAAAAGGACAATTACAAACAGGAATCGAATGCAACCGGCGCAGGAACACTGCCAGCGCATCAACAATATT
TTCACCTGAATCAGGATATTCTTCTAATACCTGGAATGCTGTTTTCCCGGGGATCGCAGTGGTGAGTAACCATGCAT
CATCAGGAGTACGGATAAAATGCTTGATGGTCGGAAGAGGCATAAATTCCGTCAGCCAGTTTAGTCTGACCATCTCA
TCTGTAACATCATTGGCAACGCTACCTTTGCCATGTTTCAGAAACAACTCTGGCGCATCGGGCTTCCCATACAATCG
ATAGATTGTCGCACCTGATTGCCCGACATTATCGCGAGCCCATTTATACCCATATAAATCAGCATCCATGTTGGAAT
TTAATCGCGGCCTAGAGCAAGACGTTTCCCGTTGAATATGGCTCATACTCTTCCTTTTTCAATATTATTGAAGCATT
TATCAGGGTTATTGTCTCATGAGCGGATACATATTTGAATGTATTTAGAAAAATAAACAAATAGGCATGCAGCGCTC
TTCCGCTTCCTCGCTCACTGACTCGCTACGCTCGGTCGTTCGACTGCGGCGAGCGGTGTCAGCTCACTCAAAAGCGG
TAATACGGTTATCCACAGAATCAGGGGATAAAGCCGGAAAGAACATGTGAGCAAAAAGCAAAGCACCGGAAGAAGCC
AACGCCGCAGGCGTTTTTCCATAGGCTCCGCCCCCCTGACGAGCATCACAAAAATCGACGCTCAAGCCAGAGGTGGC
GAAACCCGACAGGACTATAAAGATACCAGGCGTTTCCCCCTGGAAGCTCCCTCGTGCGCTCTCCTGTTCCGACCCTG
CCGCTTACCGGATACCTGTCCGCCTTTCTCCCTTCGGGAAGCGTGGCGCTTTCTCATAGCTCACGCTGTTGGTATCT
CAGTTCGGTGTAGGTCGTTCGCTCCAAGCTGGGCTGTGTGCACGAACCCCCCGTTCAGCCCGACCGCTGCGCCTTAT
CCGGTAACTATCGTCTTGAGTCCAACCCGGTAAGACACGACTTATCGCCACTGGCAGCAGCCATTGGTAACTGATTT
AGAGGACTTTGTCTTGAAGTTATGCACCTGTTAAGGCTAAACTGAAAGAACAGATTTTGGTGAGTGCGGTCCTCCAA
CCCACTTACCTTGGTTCAAAGAGTTGGTAGCTCAGCGAACCTTGAGAAAACCACCGTTGGTAGCGGTGGTTTTTCTT
TATTTATGAGATGATGAATCAATCGGTCTATCAAGTCAACGAACAGCTATTCCGTTACTCTAGATTTCAGTGCAATT
TATCTCTTCAAATGTAGCACCTGAAGTCAGCCCCATACGATATAAGTTGTAATTCTCATGTTAGTCATGCCCCGCGC
CCACCGGAAGGAGCTGACTGGGTTGAAGGCTCTCAAGGGCATCGGTCGAGATCCCGGTGCCTAATGAGTGAGCTAAC
TTACATTAATTGCGTTGCGCTCACTGCCCGCTTTCCAGTCGGGAAACCTGTCGTGCCAGCTGCATTAATGAATCGGC
CAACGCGCGGGGAGAGGCGGTTTGCGTATTGGGCGCCAGGGTGGTTTTTCTTTTCACCAGTGAGACGGGCAACAGCT
GATTGCCCTTCACCGCCTGGCCCTGAGAGAGTTGCAGCAAGCGGTCCACGCTGGTTTGCCCCAGCAGGCGAAAATCC
TGTTTGATGGTGGTTAACGGCGGGATATAACATGAGCTGTCTTCGGTATCGTCGTATCCCACTACCGAGATGTCCGC
ACCAACGCGCAGCCCGGACTCGGTAATGGCGCGCATTGCGCCCAGCGCCATCTGATCGTTGGCAACCAGCATCGCAG
TGGGAACGATGCCCTCATTCAGCATTTGCATGGTTTGTTGAAAACCGGACATGGCACTCCAGTCGCCTTCCCGTTCC
GCTATCGGCTGAATTTGATTGCGAGTGAGATATTTATGCCAGCCAGCCAGACGCAGACGCGCCGAGACAGAACTTAA
TGGGCCCGCTAACAGCGCGATTTGCTGGTGACCCAATGCGACCAGATGCTCCACGCCCAGTCGCGTACCGTCTTCAT
GGGAGAAAATAATACTGTTGATGGGTGTCTGGTCAGAGACATCAAGAAATAACGCCGGAACATTAGTGCAGGCAGCT
TCCACAGCAATGGCATCCTGGTCATCCAGCGGATAGTTAATGATCAGCCCACTGACGCGTTGCGCGAGAAGATTGTG
CACCGCCGCTTTACAGGCTTCGACGCCGCTTCGTTCTACCATCGACACCACCACGCTGGCACCCAGTTGATCGGCGC
GAGATTTAATCGCCGCGACAATTTGCGACGGCGCGTGCAGGGCCAGACTGGAGGTGGCAACGCCAATCAGCAACGAC
TGTTTGCCCGCCAGTTGTTGTGCCACGCGGTTGGGAATGTAATTCAGCTCCGCCATCGCCGCTTCCACTTTTTCCCG
CGTTTTCGCAGAAACGTGGCTGGCCTGGTTCACCACGCGGGAAACGGTCTGATAAGAGACACCGGCATACTCTGCGA
CATCGTATAACGTTACTGGTTTCACATTCACCACCCTGAATTGACTCTCTTCCGGGCGCTATCATGCCATACCGCGA
AAGGTTTTGCGCCATTCGATGGTGTCCGGGATCTCGACGCTCTCCCTTATGCGACTCCTGCATTAGGAAATTAATAC
GACTCACTATAGGGGAATTGTGAGCGGATAACAATTCCCCTGTAGAAATAATTTTGTTTAACTTTAATAAGGAGATA
TACC-Insert
```

B) Insert of ApyA co-expressed with ApyD, ApyS, ApyI, ApyH, and ApyO (see panel a for color coding).

**ATGGGCAGCAGCCATCACCATCATCACCACAGCCAGATGGCGACCAAACCAAAAAAGACCAAGGGCGTGAGCGTATC**
**TATTGAAGGGAAATTACCGAAAATGACGTTAGATATGCCTGTCGATGCAAAAAAAATCAAAGCTATTCAGAAATGTC**
**TGGAGAACGGTAAACTCACCATTACCATGTCTAAAGTAGATCTGGCAGGCGGTCGTATGGGGAAGGTTATCTGTAC**
**GATTGA**GATTCAAGGAGATATACCATGAAACTGTTGCTGGTTGATAACTTGGTAATGCCGGAGGAGGGCTCGCTGGC
ATATTTGGACGTGCATCCGCATCTGGGACTGCTGGCGCTGGCGGCCGTCGCAGAAGCAGACGGTCACCGCGTGCAAA
TTTACGATCCGAAACGCCTGATTCGTGATGGCACACTGGCTTATGATGCCACCCTTTATGAACGCGCATCTCAAGCG
ATTCTGGCAGCGCGTCCTGATGGCGTCGGTTTCACTGCACTGGGCTGTTCCTTCCTGTTCGCGTTGAATGTGGCGGC
GTTACTGAAACGTCGCGAACCCGATCTGCCGGTGTTGCTGGGCGGTCCGCACGCGACCATGCTGCATCGCCAGATTC
TCGAACGTTTTCCGCAGTTTGATATCATCGTTCGCTATGAAGCCGATGAAATTCTGCCGGCCGTCCTCGATTGCCTG
CCACACCGCACCTTCGACGTGATCCCTGGCCTGAGTTGGCGTGCAACCGGTCGCGGCAGCCCGCTTCGGTTCACGGA
TGGCAAGCCGAAAGTTGAGGATCTGGACCTGCTGCCCATCGCTAGTTACGACCATTATCCGGTTGAAGAACTGGGAC
TGTCGATGCTGCGTATCGAAGCCGGCCGCGGCTGCCCATTTGCGTGCACCTTTTGTTCCACGGCCGGTTTCTTTCAG
CGTTCCTTCCGCCTGAAGTCCGCTGAACGGCTGGTGCGTGAACTTGATATCTTACACCAACGTTACCGTGTCTCCGA
TTTCAAACTGGATCATGATATGTTTACTGTGAATCGTCGTAAAGTCATGGAATTCTGTGAAGCGGTTGCAGGTCGTG
ATTATCGCTGGCGTGCCTCGGCCCGCATCGACTGTGTGGATGAAGCCCTGCTGAAGAAAATGGCGGATGCGGGTTGC
GTTAACCTGTACTTTGGTGTTGAAACAGGTTCAGAACGCATGCAGAAACTGTGTAAAAAGCGTCTGGATCTTCAACG
TGTGGAGCCAATCCTGGCGGCCGCGGACAGCTTCGGCATTGAAACCACCGCCTCGTTCATCACCGGCTACCCGGAAG
AAACGGGTGAAGATCAAGACGATACGCTCGATATGATTGGCCGCTGCGCCCGTCGTGCTAGTTGTCTGACCCAGCTG
CATATGCTGGCGCCGGAACCGGGGACGCCGTTGTTCGACGAACGCGGCGCGGAAATCGCGTATGACGGTTGTGGCGG
TCCCTATAACACGCGTTTATTATCTAGCTCAGATGAACGTGCGGTACTCGGTCACCCTGATATTTTCCAGACTTACT
ACCACTATCCGGCAGCTTTACCGCGTGCACGTTATATCTTTGCTGTTGGGGCTGCCGATCTCCTGCGTCGTGTGGGT
CCAGTAATTTTTGCGTATGCGCTGCGTGGATTCGGCGGCCGTCTGTCCACTCTGATTAGTGCGTGGCGCCAGTTCGC
CGACGAGACCAGTCCGGGAGCCCCGGCGGACGCAGCGGGTCTGGAAGCGTTTATCGCTGCTCGTTTTGGCCGGGGCC
ACCACCTGCATAGTCTGTTCCGTTATGCGTTGCGCCTGTACGCCGCCCGTGCCCCTGCAGATGTGATCGTCGAAGGT
CCATATGAGCCAGATCGTCCATGCATGCTTGCTGAAAACGTCGGGTTGTTGGAAGACATTCACAATTGTGATGCCTT
GGTTGATCGTATTCGCCGTTCAACCGAAGGCGCCCCGATGCTGGGCGATGCCGATGTCGGGGGCTTGGGCACGTATA
TCGTGCAGGGTCACGGCGAAGCCGCCACAGGTTACTGGGTCGAATCCGGTATTGCGACTCTGCTGGATCTGTTTCGT
TCTCCTCGTAGTTGCCGGGCTGTGGCGCAATGGCTGGCAGACGCAACGCGTAACGATGGCATTGATGCCAGCATCTT
TGAACCTCTGCTCCGCCGCGGTATTCTCGTGAGCGCAGGGTAAGATTCAAGGAGATATACCATGCGTCGCACGCTCG
AGCGCTGGTCAGCATTAGACTTGGCTGAGGGTCTGCAATTAGCGCACGCGGTGGCTGCGCTTCAGGCGTTGGGTGTC
CTGGATGCGATGCGTGAGCCAGTTACGGCAGAGACATTGTCTGCCTCTCACGATCTCGACCCGGAATTGCTGCGTGG
AATTCTTGAATACGCCGCATCCCGCACTAATTTGGTGCGTAAGACGGACCGTGGATTCGCGATTTCTGAGCATTATA
CCGCCCAAGCCCGTTTTCTCCTGAATTTATATGCCGGTGCTTACGCAGACAACGCCAGCCGTTTGGCCACATTATTA
CGTCGTCCCGCACTGGCTGGGAGCATGATCGATCTTGTGGCGCACGCTCGTGCATTCGATGCGAGTGAGGCTGGTGG
TGGTGCTCTTGCGTCTATTGTCACCCAATTGCGTCTCAACCATGTGTTGGATCTTGGTTGCGGCGCGGGCGCACTGT
TACGTGAGTTGGCAGCGGATGATCGTGGCTTCGTCGGCTGGGGTCTCGATCGCAATCCTGCGATGTGTAAGGCCGCA
CGTGTCCGTGCACGCCAGGCTGGATTGGCCGCGCGCGTCAAGGTTTTCCACGGAGACGGTTGGAATCCTGGTGCCAG
CGTGCCACCGCGCGTTTTAGCGCGCGTACGTAACGTCGTTGCTAGTCAATTCGTGAACGAAATGTTCCGCGGCGGTA
CGTCCCGCGTTGAGACCTGGTTGCGCCGTATGCGTCGCCTCTTGCCCGGCCGTATGTTGGTTATTAGTGACTATTAC
GGTCGTCTCGGGCACGGCACCCACACCATGCGTCGCGAGACGCTTTTGCATGATTACGTGCAGTTGATCTCAGGTCA
AGGTGTGCCGCCACCTGACTCCGATGCGTGGTTGGCTATGTACCGTGCCGCAGGTTGCCGTCCGCTTCATGTGATCG
AGGACCGCGCGGCGACTACTTGCTTCGTTCATTTAGTTGGTTTATGAGTAGCGAAGCGGTCTAGCATAACCCCTTGG
GGCCTCTAAACGGGTCTTGAGGGGTTTTTTGCCCTTAATACGACTCACTATAGGGGAATTGTGAGCGGATAACAATT
CCAAGGAGATATACAATGAACGCAGGCAAAATTCACGCGGTTATTGCGGCTGGTGCTAAAGACCCACGGTTGATTGA
ACATTGGCGTCGCCGTCCAGATCTGCTGCGGCGCGCGGGTATCGAGCCCGATCATCTGGATCTTCGCGTCCTGGAGA
ATTTTGCGGGGCTGTCCCTTAAAGTTCGCCACAATGGATTGCGCGGTGATTTGCCGTTAAGCTTTCGGTTGCTCTCT
GTGGCGGGATTGGAAATTGAAGTTTTTGCAGCGTACGCATCGGATTGCGCCGAATCAGGGCACGTCTTTGCCGCCAC
ACCGCGTGAACGTGCTCGCGACCTCATGGCCTTCCTTGACCGCTGGCTGGATCCGGCCCAGCGCAACCACGTGCTGT
TATGGGACATCCTGCGCCATGAATACGCTCTCTCGTGCCTGCGTCAGGAATCTGCGGCCCCGCTGCCGGCAGCGCGC
CGTATTATTCGCCACCGTGATCGTGCAACTGGCGGTTGCATTCCAATGCTGAATGGAAGTGTCGCCCAGCACGTTAT
GCATTGTAATCCAGAAGATCTGGCGTCGGCCTTACGCCAAAAAGAACCGCCCCTGGATGCTGTTCCGGCCGCGGACA
CCTACTGTTGCTACTGGCGCAGTGCTGAGACGGGCGAGATCCGTTTTCTCCAGGCAGATACCCTGGGTTATTTTGCA
ATTGAGGCAATCGATGGCCGCCGCTCTGCGGCGGACATCTGTGATCGCCTGCTGGGCGGTCGTCGCCTCCTGCCCCA
GTTTCTGCGCAGCCTGGACCAGCTCGCGGGCATTGGTATTGTACGGTTCGAGCGTCCACTTGGAAGCGCGGCAGCAT

AAGATTCAAGGAGATATACCATGAAAGCTTTTGATCGCGATGCATCAGAACCACGTATCGGCCTGGCCTATGGTCCG
GGCATGGCGGAGTTTGCAGCCCGGAACGCGCATCTCGTGGACTACATTGAAGTTCCGTTTGAACAACTGCGTTTCAG
CCCAGCAGTAGCCGAGCTGCAGCAGACGATTCCGTTCGTGCTGCATTGCGCGAGCTTGAGCGTGGCTGGCTTTGTGC
CGCCGGATGACTCGACTGTAGATGCAATTGAACGCACCGCAGTGCAAACTGGTACCCCGTGGATCGGCGAGCACCTG
GCGTATATCTCGGCGGACCCCGTTGGTGAAGCATTAGGTGGTACCGGGGAGCCGACTTCGCTGTCATACACCCTCTG
TCCGCAGTTATCCGACGAGACCGTCCGTCGCGTGGTGGATAATTTAGCCGCACTGCGTCCTCACTTTCCGGTGCCCT
TGATCGTAGAAAACTCGCCGCAATATTTTCCAATCCCCGGCAGCACTATGGGCATGACCGATTTTATCCGTGCCATT
ACGGATCGCTGTGATGTTGGCTTATTACTTGATTTATCTCACTTTCTGATCACTGCCCACAATACGGGCGCAGAAGT
TCACCGCGAACTGGCGCGCCTGCCACTGGAACGCGTGGTGGAGGTACACTTGTCCGGCATGTCTGTGCAAAGCGGTA
CCGCCTGGGATGATCATTCATTACCGGCCTCACCTATTTTATTTGAACTGTTGGAACGGTTGCTGGACGTGGCTCGC
CCGCGCGCCCTGACATTTGAATACAATTGGAGTCCCTACTTTCCACTGAGCGTGTTGACTACCCACATCGAACGCGC
CCGTCAGCTGATGGGGCTGGCTTAAGATTCAAGGAGATATACCATGAACGGTCTCCCGCCTTCTCTGCCTCGTGTGG
ACTCAACAGAAAGCCTGTTCGCGGAGCCTTTAGCATTCTTAGCGCAAGCCCGTTCTCGTCACGGAGACGTGTTCGTT
ATGCGCGAGCACGGCCCCATCTTTTCCCGCGCATCGGATTGTTCGGGTGTCATTGCTGTCTTTGGAGAGCACCGTTT
ACGTCAAATCCTTACGGACATCGACAACTTCGCGTTACCGATGAGTGCCGCCGCCAAGATGGCATTACCTAAGAACC
TCGTCAATTTAAACCGTGGCCTGCATAGTATGCGCGAGCCGGAGCATGGTCGTCACAAGCGTTTGCTTACCGGTACA
ATTAACCGCGAGCTCTTTGACGCCCACCGTTTCGAAATTCGCGCAGCGCTCAACCGCTTTTGTGAAATGCTCAAGGT
AGATCGCCGTATCAGTGTGGTGAGCCGTATGCGCGAGCTGACGGTCGAGATGGCATCTCATATCTTTCTGGGTGCAC
AATGTCAGGAAGACGATGAGTTGGCGTTTCTTCTTTCAGCCTATTTCACCCTGCGCCGCGAGGCTAGCTCATTAAAC
GCTCGTGACCCGTTGCTTTATCGTGACGAGCTGATCGGTGTGGGACAACAGCTGGACCGCACATTACGCGAACGTAT
CCGCCGCTATCGTAAACGCCCCGTCGACGCCCGCGCCGGTCTGTTACAGCGTCTTGCTACGGCAGGTCCGCCGGGCT
CGCCTGCGTTGTCGGAAGATGAAATTGTGGGTCACGCCAACGTAATGTTCGTGTCTTCCACGGAGCCGGTTGCCATG
TCTCTTACGTGGTTGTTGCTGGTCTTAAGTCAACTGCCTGACTTACGTCGTGCGCTGCGTGCGGAGATCGCGGACCG
TGCGTCTATGCCGGCCAGTACTAATGGCGCGTCGTGGTTAGAGAACGTTGTCAACGAGACGTTGCGCTTATTGACCC
CAAATGCCCTGATGGTCCGCGCCACAACACGCGCAGTGTCACTGCAAGGTGTTGCACTCCCGGCCCGCTGTGAGATT
GTGGTATGTCCATTTCTGGCTCACCGCGAGGCCAAACCATTTCCCGACCCCCACGCATTCTCACCATCGCGCTGGGA
GACCGCGCGTCCCAGTCCTTATGAGTACTTTCCTTTCGGGGCGGGCGGACACTTCTGTGCGGGGCGTAATTTGGCCC
TCTCACTTATTCGCGAAGTGTTGTCGACGCTGTTAAGCCGTTTCGATTTCGTTCTTGACGGTGAGCAAAGCATCGAC
TGGCGCATTCATATTATGCTGATGCCCAAGGGAGACCCGGCTTTGATTGCTCACCCTGTGGATGAACGTGGCGATAC
CCCGAGTCCAAAGTGGCGTGGGCCCATTACCGATTTATTTCACTTCGCCCCTGGGCTTTCATGA

C) Insert of ApyA co-expressed with ApyD

**ATGGGCAGCAGCCATCACCATCATCACCACAGCCAGATGGCGACCAAACCAAAAAAGACCAAGGGCGTGAGCGTATC**
**TATTGAAGGGAAATTACCGAAAATGACGTTAGATATGCCTGTCGATGCAAAAAAAATCAAAGCTATTCAGAAATGTC**
**TGGAGAACGGTAAACTCACCATTACCATGTCTAAAGTAGATCTGGCAGGCGGTCGTATGGGGGAAGGTTATCTGTAC**
**GATTGA**GATTCAAGGAGATATACCATGAAACTGTTGCTGGTTGATAACTTGGTAATGCCGGAGGAGGGCTCGCTGGC
ATATTTGGACGTGCATCCGCATCTGGGACTGCTGGCGCTGGCGGCCGTCGCAGAAGCAGACGGTCACCGCGTGCAAA
TTTACGATCCGAAACGCCTGATTCGTGATGGCACACTGGCTTATGATGCCACCCTTTATGAACGCGCATCTCAAGCG
ATTCTGGCAGCGCGTCCTGATGGCGTCGGTTTCACTGCACTGGGCTGTTCCTTCCTGTTCGCGTTGAATGTGGCGGC
GTTACTGAAACGTCGCGAACCCGATCTGCCGGTGTTGCTGGGCGGTCCGCACGCGACCATGCTGCATCGCCAGATTC
TCGAACGTTTTCCGCAGTTTGATATCATCGTTCGCTATGAAGCCGATGAAATTCTGCCGGCCGTCCTCGATTGCCTG
CCACACCGCACCTTCGACGTGATCCCTGGCCTGAGTTGGCGTGCAACCGGTCGCGGCAGCCCGCTTCGGTTCACGGA
TGGCAAGCCGAAAGTTGAGGATCTGGACCTGCTGCCCATCGCTAGTTACGACCATTATCCGGTTGAAGAACTGGGAC
TGTCGATGCTGCGTATCGAAGCCGGCCGCGGCTGCCCATTTGCGTGCACCTTTTGTTCCACGGCCGGTTTCTTTCAG
CGTTCCTTCCGCCTGAAGTCCGCTGAACGGCTGGTGCGTGAACTTGATATCTTACACCAACGTTACCGTGTCTCCGA
TTTCAAACTGGATCATGATATGTTTACTGTGAATCGTCGTAAAGTCATGGAATTCTGTGAAGCGGTTGCAGGTCGTG
ATTATCGCTGGCGTGCCTCGGCCCGCATCGACTGTGTGGATGAAGCCCTGCTGAAGAAAATGGCGGATGCGGGTTGC
GTTAACCTGTACTTTGGTGTTGAAACAGGTTCAGAACGCATGCAGAAACTGTGTAAAAAGCGTCTGGATCTTCAACG
TGTGGAGCCAATCCTGGCGGCCGCGGACAGCTTCGGCATTGAAACCACCGCCTCGTTCATCACCGGCTACCCGGAAG
AAACGGGTGAAGATCAAGACGATACGCTCGATATGATTGGCCGCTGCGCCCGTCGTGCTAGTTGTCTGACCCAGCTG
CATATGCTGGCGCCGGAACCGGGGACGCCGTTGTTCGACGAACGCGGCGCGGAAATCGCGTATGACGGTTGTGGCGG
TCCCTATAACACGCGTTTATTATCTAGCTCAGATGAACGTGCGGTACTCGGTCACCCTGATATTTTCCAGACTTACT
ACCACTATCCGGCAGCTTTACCGCGTGCACGTTATATCTTTGCTGTTGGGGCTGCCGATCTCCTGCGTCGTGTGGGT
CCAGTAATTTTTGCGTATGCGCTGCGTGGATTCGGCGGCCGTCTGTCCACTCTGATTAGTGCGTGGCGCCAGTTCGC
CGACGAGACCAGTCCGGGAGCCCCGGCGGACGCAGCGGGTCTGGAAGCGTTTATCGCTGCTCGTTTTGGCCGGGGCC
ACCACCTGCATAGTCTGTTCCGTTATGCGTTGCGCCTGTACGCCGCCCGTGCCCCTGCAGATGTGATCGTCGAAGGT

CCATATGAGCCAGATCGTCCATGCATGCTTGCTGAAAACGTCGGGTTGTTGGAAGACATTCACAATTGTGATGCCTT
GGTTGATCGTATTCGCCGTTCAACCGAAGGCGCCCCGATGCTGGGCGATGCCGATGTCGGGGGCTTGGGCACGTATA
TCGTGCAGGGTCACGGCGAAGCCGCCACAGGTTACTGGGTCGAATCCGGTATTGCGACTCTGCTGGATCTGTTTCGT
TCTCCTCGTAGTTGCCGGGCTGTGGCGCAATGGCTGGCAGACGCAACGCGTAACGATGGCATTGATGCCAGCATCTT
TGAACCTCTGCTCCGCCGCGGTATTCTCGTGAGCGCAGGGTAA

D) ApyA co-expressed with ApyO
**ATGGGCAGCAGCCATCACCATCATCACCACAGCCAGATGGCGACCAAACCAAAAAAGACCAAGGGCGTGAGCGTATC
TATTGAAGGGAAATTACCGAAAATGACGTTAGATATGCCTGTCGATGCAAAAAAAATCAAAGCTATTCAGAAATGTC
TGGAGAACGGTAAACTCACCATTACCATGTCTAAAGTAGATCTGGCAGGCGGTCGTATGGGGGAAGGTTATCTGTAC
GATTGA**GATTCAAGGAGATATACCATGAACGGTCTCCCGCCTTCTCTGCCTCGTGTGGACTCAACAGAAAGCCTGTT
CGCGGAGCCTTTAGCATTCTTAGCGCAAGCCCGTTCTCGTCACGGAGACGTGTTCGTTATGCGCGAGCACGGCCCCA
TCTTTTCCCGCGCATCGGATTGTTCGGGTGTCATTGCTGTCTTTGGAGAGCACCGTTTACGTCAAATCCTTACGGAC
ATCGACAACTTCGCGTTACCGATGAGTGCCGCCGCCAAGATGGCATTACCTAAGAACCTCGTCAATTTAAACCGTGG
CCTGCATAGTATGCGCGAGCCGGAGCATGGTCGTCACAAGCGTTTGCTTACCGGTACAATTAACCGCGAGCTCTTTG
ACGCCCACCGTTTCGAAATTCGCGCAGCGCTCAACCGCTTTTGTGAAATGCTCAAGGTAGATCGCCGTATCAGTGTG
GTGAGCCGTATGCGCGAGCTGACGGTCGAGATGGCATCTCATATCTTTCTGGGTGCACAATGTCAGGAAGACGATGA
GTTGGCGTTTCTTCTTTCAGCCTATTTCACCCTGCGCCGCGAGGCTAGCTCATTAAACGCTCGTGACCCGTTGCTTT
ATCGTGACGAGCTGATCGGTGTGGGACAACAGCTGGACCGCACATTACGCGAACGTATCCGCCGCTATCGTAAACGC
CCCGTCGACGCCCGCGCCGGTCTGTTACAGCGTCTTGCTACGGCAGGTCCGCCGGGCTCGCCTGCGTTGTCGGAAGA
TGAAATTGTGGGTCACGCCAACGTAATGTTCGTGTCTTCCACGGAGCCGGTTGCCATGTCTCTTACGTGGTTGTTGC
TGGTCTTAAGTCAACTGCCTGACTTACGTCGTGCGCTGCGTGCGGAGATCGCGGACCGTGCGTCTATGCCGGCCAGT
ACTAATGGCGCGTCGTGGTTAGAGAACGTTGTCAACGAGACGTTGCGCTTATTGACCCCAAATGCCCTGATGGTCCG
CGCCACAACACGCGCAGTGTCACTGCAAGGTGTTGCACTCCCGGCCCGCTGTGAGATTGTGGTATGTCCATTTCTGG
CTCACCGCGAGGCCAAACCATTTCCCGACCCCCACGCATTCTCACCATCGCGCTGGGAGACCGCGCGTCCCAGTCCT
TATGAGTACTTTCCTTTCGGGGCGGGCGGACACTTCTGTGCGGGGCGTAATTTGGCCCTCTCACTTATTCGCGAAGT
GTTGTCGACGCTGTTAAGCCGTTTCGATTTCGTTCTTGACGGTGAGCAAAGCATCGACTGGCGCATTCATATTATGC
TGATGCCCAAGGGAGACCCGGCTTTGATTGCTCACCCTGTGGATGAACGTGGCGATACCCCGAGTCCAAAGTGGCGT
GGGCCCATTACCGATTTATTTCACTTCGCCCCTGGGCTTTCATGA

E) ApyA co-expressed with ApyD and ApyO
**ATGGGCAGCAGCCATCACCATCATCACCACAGCCAGATGGCGACCAAACCAAAAAAGACCAAGGGCGTGAGCGTATC
TATTGAAGGGAAATTACCGAAAATGACGTTAGATATGCCTGTCGATGCAAAAAAAATCAAAGCTATTCAGAAATGTC
TGGAGAACGGTAAACTCACCATTACCATGTCTAAAGTAGATCTGGCAGGCGGTCGTATGGGGGAAGGTTATCTGTAC
GATTGA**GATTCAAGGAGATATACCATGAAACTGTTGCTGGTTGATAACTTGGTAATGCCGGAGGAGGGCTCGCTGGC
ATATTTGGACGTGCATCCGCATCTGGGACTGCTGGCGCTGGCGGCCGTCGCAGAAGCAGACGGTCACCGCGTGCAAA
TTTACGATCCGAAACGCCTGATTCGTGATGGCACACTGGCTTATGATGCCACCCTTTATGAACGCGCATCTCAAGCG
ATTCTGGCAGCGCGTCCTGATGGCGTCGGTTTCACTGCACTGGGCTGTTCCTTCCTGTTCGCGTTGAATGTGGCGGC
GTTACTGAAACGTCGCGAACCCGATCTGCCGGTGTTGCTGGGCGGTCCGCACGCGACCATGCTGCATCGCCAGATTC
TCGAACGTTTTCCGCAGTTTGATATCATCGTTCGCTATGAAGCCGATGAAATTCTGCCGGCCGTCCTCGATTGCCTG
CCACACCGCACCTTCGACGTGATCCCTGGCCTGAGTTGGCGTGCAACCGGTCGCGGCAGCCCGCTTCGGTTCACGGA
TGGCAAGCCGAAAGTTGAGGATCTGGACCTGCTGCCCATCGCTAGTTACGACCATTATCCGGTTGAAGAACTGGGAC
TGTCGATGCTGCGTATCGAAGCCGGCCGCGGCTGCCCATTTGCGTGCACCTTTTGTTCCACGGCCGGTTTCTTTCAG
CGTTCCTTCCGCCTGAAGTCCGCTGAACGGCTGGTGCGTGAACTTGATATCTTACACCAACGTTACCGTGTCTCCGA
TTTCAAACTGGATCATGATATGTTTACTGTGAATCGTCGTAAAGTCATGGAATTCTGTGAAGCGGTTGCAGGTCGTG
ATTATCGCTGGCGTGCCTCGGCCCGCATCGACTGTGTGGATGAAGCCCTGCTGAAGAAAATGGCGGATGCGGGTTGC
GTTAACCTGTACTTTGGTGTTGAAACAGGTTCAGAACGCATGCAGAAACTGTGTAAAAAGCGTCTGGATCTTCAACG
TGTGGAGCCAATCCTGGCGGCCGCGGACAGCTTCGGCATTGAAACCACCGCCTCGTTCATCACCGGCTACCCGGAAG
AAACGGGTGAAGATCAAGACGATACGCTCGATATGATTGGCCGCTGCGCCCGTCGTGCTAGTTGTCTGACCCAGCTG
CATATGCTGGCGCCGGAACCGGGGACGCCGTTGTTCGACGAACGCGGCGCGGAAATCGCGTATGACGGTTGTGGCGG
TCCCTATAACACGCGTTTATTATCTAGCTCAGATGAACGTGCGGTACTCGGTCACCCTGATATTTTCCAGACTTACT
ACCACTATCCGGCAGCTTTACCGCGTGCACGTTATATCTTTGCTGTTGGGGCTGCCGATCTCCTGCGTCGTGTGGGT
CCAGTAATTTTTGCGTATGCGCTGCGTGGATTCGGCGGCCGTCTGTCCACTCTGATTAGTGCGTGGCGCCAGTTCGC
CGACGAGACCAGTCCGGGAGCCCCGGCGGACGCAGCGGGTCTGGAAGCGTTTATCGCTGCTCGTTTTGGCCGGGGCC
ACCACCTGCATAGTCTGTTCCGTTATGCGTTGCGCCTGTACGCCGCCCGTGCCCCTGCAGATGTGATCGTCGAAGGT
CCATATGAGCCAGATCGTCCATGCATGCTTGCTGAAAACGTCGGGTTGTTGGAAGACATTCACAATTGTGATGCCTT

GGTTGATCGTATTCGCCGTTCAACCGAAGGCGCCCCGATGCTGGGCGATGCCGATGTCGGGGGCTTGGGCACGTATA
TCGTGCAGGGTCACGGCGAAGCCGCCACAGGTTACTGGGTCGAATCCGGTATTGCGACTCTGCTGGATCTGTTTCGT
TCTCCTCGTAGTTGCCGGGCTGTGGCGCAATGGCTGGCAGACGCAACGCGTAACGATGGCATTGATGCCAGCATCTT
TGAACCTCTGCTCCGCCGCGGTATTCTCGTGAGCGCAGGGTAAGATTCAAGGAGATATACCATGAACGGTCTCCCGC
CTTCTCTGCCTCGTGTGGACTCAACAGAAAGCCTGTTCGCGGAGCCTTTAGCATTCTTAGCGCAAGCCCGTTCTCGT
CACGGAGACGTGTTCGTTATGCGCGAGCACGGCCCCATCTTTTCCCGCGCATCGGATTGTTCGGGTGTCATTGCTGT
CTTTGGAGAGCACCGTTTACGTCAAATCCTTACGGACATCGACAACTTCGCGTTACCGATGAGTGCCGCCGCCAAGA
TGGCATTACCTAAGAACCTCGTCAATTTAAACCGTGGCCTGCATAGTATGCGCGAGCCGGAGCATGGTCGTCACAAG
CGTTTGCTTACCGGTACAATTAACCGCGAGCTCTTTGACGCCCACCGTTTCGAAATTCGCGCAGCGCTCAACCGCTT
TTGTGAAATGCTCAAGGTAGATCGCCGTATCAGTGTGGTGAGCCGTATGCGCGAGCTGACGGTCGAGATGGCATCTC
ATATCTTTCTGGGTGCACAATGTCAGGAAGACGATGAGTTGGCGTTTCTTCTTTCAGCCTATTTCACCCTGCGCCGC
GAGGCTAGCTCATTAAACGCTCGTGACCCGTTGCTTTATCGTGACGAGCTGATCGGTGTGGGACAACAGCTGGACCG
CACATTACGCGAACGTATCCGCCGCTATCGTAAACGCCCCGTCGACGCCCGCGCCGGTCTGTTACAGCGTCTTGCTA
CGGCAGGTCCGCCGGGCTCGCCTGCGTTGTCGGAAGATGAAATTGTGGGTCACGCCAACGTAATGTTCGTGTCTTCC
ACGGAGCCGGTTGCCATGTCTCTTACGTGGTTGTTGCTGGTCTTAAGTCAACTGCCTGACTTACGTCGTGCGCTGCG
TGCGGAGATCGCGGACCGTGCGTCTATGCCGGCCAGTACTAATGGCGCGTCGTGGTTAGAGAACGTTGTCAACGAGA
CGTTGCGCTTATTGACCCCAAATGCCCTGATGGTCCGCGCCACAACACGCGCAGTGTCACTGCAAGGTGTTGCACTC
CCGGCCCGCTGTGAGATTGTGGTATGTCCATTTCTGGCTCACCGCGAGGCCAAACCATTTCCCGACCCCCACGCATT
CTCACCATCGCGCTGGGAGACCGCGCGTCCCAGTCCTTATGAGTACTTTCCTTTCGGGGCGGGCGGACACTTCTGTG
CGGGGCGTAATTTGGCCCTCTCACTTATTCGCGAAGTGTTGTCGACGCTGTTAAGCCGTTTCGATTTCGTTCTTGAC
GGTGAGCAAAGCATCGACTGGCGCATTCATATTATGCTGATGCCCAAGGGAGACCCGGCTTTGATTGCTCACCCTGT
GGATGAACGTGGCGATACCCCGAGTCCAAAGTGGCGTGGGCCCATTACCGATTTATTTCACTTCGCCCCTGGGCTTT
CATGA

F) ApyA co-expressed with ApyI and ApyH

**ATGGGCAGCAGCCATCACCATCATCACCACAGCCAGATGGCGACCAAACCAAAAAAGACCAAGGGCGTGAGCGTATC
TATTGAAGGGAAATTACCGAAAATGACGTTAGATATGCCTGTCGATGCAAAAAAAATCAAAGCTATTCAGAAATGTC
TGGAGAACGGTAAACTCACCATTACCATGTCTAAAGTAGATCTGGCAGGCGGTCGTATGGGGGAAGGTTATCTGTAC
GATTGA**GTAGCGAAGCGGTCTAGCATAACCCCTTGGGGCCTCTAAACGGGTCTTGAGGGGTTTTTTGCCCTTAATAC
GACTCACTATAGGGGAATTGTGAGCGGATAACAATTCCAAGGAGATATACAATGAACGCAGGCAAAATTCACGCGGT
TATTGCGGCTGGTGCTAAAGACCCACGGTTGATTGAACATTGGCGTCGCCGTCCAGATCTGCTGCGGCGCGCGGGTA
TCGAGCCCGATCATCTGGATCTTCGCGTCCTGGAGAATTTTGCGGGGCTGTCCCTTAAAGTTCGCCACAATGGATTG
CGCGGTGATTTGCCGTTAAGCTTTCGGTTGCTCTCTGTGGCGGGATTGGAAATTGAAGTTTTTGCAGCGTACGCATC
GGATTGCGCCGAATCAGGGCACGTCTTTGCCGCCACACCGCGTGAACGTGCTCGCGACCTCATGGCCTTCCTTGACC
GCTGGCTGGATCCGGCCCAGCGCAACCACGTGCTGTTATGGGACATCCTGCGCCATGAATACGCTCTCTCGTGCCTG
CGTCAGGAATCTGCGGCCCCGCTGCCGGCAGCGCGCCGTATTATTCGCCACCGTGATCGTGCAACTGGCGGTTGCAT
TCCAATGCTGAATGGAAGTGTCGCCCAGCACGTTATGCATTGTAATCCAGAAGATCTGGCGTCGGCCTTACGCCAAA
AAGAACCGCCCCTGGATGCTGTTCCGGCCGCGGACACCTACTGTTGCTACTGGCGCAGTGCTGAGACGGGCGAGATC
CGTTTTCTCCAGGCAGATACCCTGGGTTATTTTGCAATTGAGGCAATCGATGGCCGCCGCTCTGCGGCGGACATCTG
TGATCGCCTGCTGGGCGGTCGTCGCCTCCTGCCCCAGTTTCTGCGCAGCCTGGACCAGCTCGCGGGCATTGGTATTG
TACGGTTCGAGCGTCCACTTGGAAGCGCGGCAGCATAAGATTCAAGGAGATATACCATGAAAGCTTTTGATCGCGAT
GCATCAGAACCACGTATCGGCCTGGCCTATGGTCCGGGCATGGCGGAGTTTGCAGCCCGGAACGCGCATCTCGTGGA
CTACATTGAAGTTCCGTTTGAACAACTGCGTTTCAGCCCAGCAGTAGCCGAGCTGCAGCAGACGATTCCGTTCGTGC
TGCATTGCGCGAGCTTGAGCGTGGCTGGCTTTGTGCCGCCGGATGACTCGACTGTAGATGCAATTGAACGCACCGCA
GTGCAAACTGGTACCCCGTGGATCGGCGAGCACCTGGCGTATATCTCGGCGGACCCCGTTGGTGAAGCATTAGGTGG
TACCGGGGAGCCGACTTCGCTGTCATACACCCTCTGTCCGCAGTTATCCGACGAGACCGTCCGTCGCGTGGTGGATA
ATTTAGCCGCACTGCGTCCTCACTTTCCGGTGCCCTTGATCGTAGAAAACTCGCCGCAATATTTTCCAATCCCCGGC
AGCACTATGGGCATGACCGATTTTATCCGTGCCATTACGGATCGCTGTGATGTTGGCTTATTACTTGATTTATCTCA
CTTTCTGATCACTGCCCACAATACGGGCGCAGAAGTTCACCGCGAACTGGCGCGCCTGCCACTGGAACGCGTGGTGG
AGGTACACTTGTCCGGCATGTCTGTGCAAAGCGGTACCGCCTGGGATGATCATTCATTACCGGCCTCACCTATTTTA
TTTGAACTGTTGGAACGGTTGCTGGACGTGGCTCGCCCGCGCGCCCTGACATTTGAATACAATTGGAGTCCCTACTT
TCCACTGAGCGTGTTGACTACCCACATCGAACGCGCCCGTCAGCTGATGGGGCTGGCTTAA

**Table S2:** Sequence of the initial pSCrhaB2-based vectors used in this study for expression in *Burkholderia* sp. FERM BP-3421. Each gene is highlighted as follows. *apyA*: bolded black; *apyO*: pink; *apyD*: green; *apyH*: orange; *apyI*: black underlined; *apyS*: blue. The N-terminus of ApyA contains a hexa-His tag.

A) Backbone of the pSCrhaB2 vector.

TCTAGAGTCGACCTGCAGGCATGCAAGCTTGGCTGTTTTGGCGGATGAGAGAAGATTTTCAGCCTGATACAGATTAA
ATCAGAACGCAGAAGCGGTCTGATAAAACAGAATTTGCCTGGCGGCAGTAGCGCGGTGGTCCCACCTGACCCCATGC
CGAACTCAGAAGTGAAACGCCGTAGCGCCGATGGTAGTGTGGGGTCTCCCCATGCGAGAGTAGGGAACTGCCAGGCA
TCAAATAAAACGAAAGGCTCAGTCGAAAGACTGGGCCTTTCGTTTTATCTGTTGTTTGTCGGTGAACGCTCTCCTGA
GTAGGACAAATCCGCCGGGAGCGGATTTGAACGTTGCGAAGCAACGGCCCGGAGGGTGGCGGGCAGGACGCCCGCCA
TAAACTGCCAGGCATCAAATTAAGCAGAAGGCCATCCTGACGGATGGCCTTTTTGCGTTTCTACAAACTCTTGTTAT
TGGTCGACTTAAACGCCTGGTGCTACGCCTGAATAAGTGATAATAAGCGGATGAATGGCAGAAATTCGAAAGCAAAT
TCGACCCGGTCGTCGGTTCAGGGCAGGGTCGTTAAATAGCCGCTTATGTCTATTGCTGGTTTACCGGTTTATTGACT
ACCGGAAGCAGTGTGACCGTGTGCTTCTCAAATGCCTGAGGCCAGTTTGCTCAGGCTCTCCCCGTGGAGGTAATAAT
TGACGATATGATCATTTATTCTGCCTCCCAGAGCCTGATAAAAACGGTGAATCCGTTAGCGAGGTGCCGCCGGCTTC
CATTCAGGTCGAGGTGGCCCGGCTCCATGCACCGCGACGCAACGCGGGGAGGCAGACAAGGTATAGGGCGGCGAGGC
GGCTACAGCCGATAGTCTGGAACAGCGCACTTACGGGTTGCTGCGCAACCCAAGTGCTACCGGCGCGGCAGCGTGAC
CCGTGTCGGCGGCTCCAACGGCTCGCCATCGTCCAGAAAACACGGCTCATCGGGCATCGGCAGGCGCTGCTGCCCGC
GCCGTTCCCATTCCTCCGTTTCGGTCAAGGCTGGCAGGTCTGGTTCCATGCCCGGAATGCCGGGCTGGCTGGGCGGC
TCCTCGCCGGGGCCGGTCGGTAGTTGCTGCTCGCCCGGATACAGGGTCGGGATGCGGCGCAGGTCGCCATGCCCCAA
CAGCGATTCGTCCTGGTCGTCGTGATCAACCACCACGGCGGCACTGAACACCGACAGGCGCAACTGGTCGCGGGGCT
GGCCCCACGCCACGCGGTCATTGACCACGTAGGCCGACACGGTGCCGGGGCCGTTGAGCTTCACGACGGAGATCCAG
CGCTCGGCCACCAAGTCCTTGACTGCGTATTGGACCGTCCGCAAAGAACGTCCGATGAGCTTGGAAAGTGTCTTCTG
GCTGACCACCACGGCGTTCTGGTGGCCCATCTGCGCCACGAGGTGATGCAGCAGCATTGCCGCCGTGGGTTTCCTCG
CAATAAGCCCGGCCCACGCCTCATGCGCTTTGCGTTCCGTTTGCACCCAGTGACCGGGCTTGTTCTTGGCTTGAATG
CCGATTTCTCTGGACTGCGTGGCCATGCTTATCTCCATGCGGTAGGGGTGCCGCACGGTTGCGGCACCATGCGCAAT
CAGCTGCAACTTTTCGGCAGCGCGACAACAATTATGCGTTGCGTAAAAGTGGCAGTCAATTACAGATTTTCTTTAAC
CTACGCAATGAGCTATTGCGGGGGGTGCCGCAATGAGCTGTTGCGTACCCCCCTTTTTTAAGTTGTTGATTTTTAAG
TCTTTCGCATTTCGCCCTATATCTAGTTCTTTGGTGCCCAAAGAAGGGCACCCCTGCGGGGTTCCCCCACGCCTTCG
GCGCGGCTCCCCCTCCGGCAAAAAGTGGCCCCTCCGGGGCTTGTTGATCGACTGCGCGGCCTTCGGCCTTGCCCAAG
GTGGCGCTGCCCCCTTGGAACCCCCGCACTCGCCGCCGTGAGGCTCGGGGGGCAGGCGGGCGGGCTTCGCCCTTCGA
CTGCCCCCACTCGCATAGGCTTGGGTCGTTCCAGGCGCGTCAAGGCCAAGCCGCTGCGCGGTCGCTGCGCGAGCCTT
GACCCGCCTTCCACTTGGTGTCCAACCGGCAAGCGAAGCGCGCAGGCCGCAGGCCGGAGGCTTTTCCCCAGAGAAAA
TTAAAAAAATTGATGGGGCAAGGCCGCAGGCCGCGCAGTTGGAGCCGGTGGGTATGTGGTCGAAGGCTGGGTAGCCG
GTGGGCAATCCCTGTGGTCAAGCTCGTGGGCAGGCGCAGCCTGTCCATCAGCTTGTCCAGCAGGGTTGTCCACGGGC
CGAGCGAAGCGAGCCAGCCGGTGGCCGCTCGCGGCCATCGTCCACATATCCACGGGCTGGCAAGGGAGCGCAGCGAC
CGCGCAGGGCGAAGCCCGGAGAGCAAGCCCGTAGGGCGCCGCAGCCGCCGTAGGCGGTCACGACTTTGCGAAGCAAA
GTCTAGTGAGTATACTCAAGCATTGAGTGGCCCGCCGGAGGCACCGCCTTGCGCTGCCCCCGTCGAGCCGGTTGGAC
ACCAAAAGGGAGGGGCAGGCATGGCGGCATACGCGATCATGCGATGCAAGAAGCTGGCGAAAATGGGCAACGTGGCG
GCCAGTCTCAAGCACGCCTACCGCGAGCGCGAGACGCCCAACGCTGACGCCAGCAGGACGCCAGAGAACGAGCACTG
GGCGGCCAGCAGCACCGATGAAGCGATGGGCCGACTGCGCGAGTTGCTGCCAGAGAAGCGGCGCAAGGACGCTGTGT
TGGCGGTCGAGTACGTCATGACGGCCAGCCCGGAATGGTGGAAGTCGGCCAGCCAAGAACAGCAGGCGGCGTTCTTC
GAGAAGGCGCACAAGTGGCTGGCGGACAAGTACGGGGCGGATCGCATCGTGACGGCCAGCATCCACCGTGACGAAAC
CAGCCCGCACATGACCGCGTTCGTGGTGCCGCTGACGCAGGACGGCAGGCTGTCGGCCAAGGAGTTCATCGGCAACA
AAGCGCAGATGACCCGCGACCAGACCACGTTTGCGGCCGCTGTGGCCGATCTAGGGCTGCAACGGGGCATCGAGGGC
AGCAAGGCACGTCACACGCGCATTCAGGCGTTCTACGAGGCCCTGGAGCGGCCACCAGTGGGCCACGTCACCATCAG
CCCGCAAGCGGTCGAGCCACGCGCCTATGCACCGCAGGGATTGGCCGAAAAGCTGGGAATCTCAAAGCGCGTTGAGA
CGCCGGAAGCCGTGGCCGACCGGCTGACAAAAGCGGTTCGGCAGGGGTATGAGCCTGCCCTACAGGCCGCCGCAGGA
GCGCGTGAGATGCGCAAGAAGGCCGATCAAGCCCAAGAGACGGCCCGAGACCTTCGGGAGCGCCTGAAGCCCGTTCT
GGACGCCCTGGGGCCGTTGAATCGGGATATGCAGGCCAAGGCCGCCGCGATCATCAAGGCCGTGGGCGAAAAGCTGC
TGACGGAACAGCGGGAAGTCCAGCGCCAGAAACAGGCCCAGCGCCAGCAGGAACGCGGGCGCGCACATTTCCCCGAA
AAGTGCCACCTGACGTCTAAGAAACCATTATTATCATGACATTAACCTATAAAAATAGGCGTATCACGAGGCCCTTT
CGTCTTCGAATAAATACCTGTGACGGAAGATCACTTCGCAGAATAAATAAATCCTGGTGTCCCTGTTGATACCGGGA
AGCCCTGGGCCAACTTTTGGCGAAAATGAGACGTTGATCGGCACGTAAGAGGTTCCAACTTTCACCATAATGAAATA
AGATCACTACCGGGCGTATTTTTTGAGTTATCGAGATTTTCAGGAGCTAAGGAAGCTAAAATGGAGAAAAAAATCAC

TGGATATACCACCGTTGATATATCCCAATGGCATCGTAAAGAACATTTTGAGGCATTTCAGTCAGTTGCTCAATGTA
CCTATAACCAGACCGTTCAGCTGGATATTACGGCCTTTTTAGCTTTTGCCATTCTCACCGGATTCAGTCGTCACTCA
TGGTGATTTCTCACTTGATAACCTTATTTTTGACGAGGGGAAATTAATAGGTTGTATTGATGTTGGACGAGTCGGAA
TCGCAGACCGATACCAGGATCTTGCCATCCTATGGAACTGCCTCGGTGAGTTTTCTCCTTCATTACAGAAACGGCTT
TTTCAAAAATATGGTATTGATAATCCTGATATGAATAAATTGCAGTTTCATTTGATGCTCGATGAGTTTTTCTAATC
AGAATTGGTTAATTGGTTGTAACACTGGCAGAGCATTACGCTGACTTGACGGGACGGCGGCTTTGTTGAATAAATCG
AACTTTTGCTGAGTTGAAGGATCAGATCACGCATCTTCCCGACAACGCAGACCGTTCCGTGGCAAAGCAAAAGTTCA
AAATCACCAACTGGTCCACCTACAACAAAGCTCTCATCAACCGTGGCTCCCTCACTTTCTGGCTGGATGATGGGGCG
ATTCAGGCCTGGTATGAGTCAGCAACACCTTCTTCACGAGGCAGACCTCAGCGCCAGAAGGCCGCCAGAGAGGCCGA
GCGCGGCCGTGAGGCTTGGACGCTAGGGCAGGGCATGAAAAAGCCCGTAGCGGGCTGCTACGGGCGTCTGACGCGGT
GGAAAGGGGGAGGGGATGTTGTTTAAATGGCTCTGCTGTAGTGAGTGGTCTACGGGGTCTGACGCTCAGTGGAACGA
AATCGATGATAAGCTGTCAAACATGAGCAGATCCTCTACGCCGGACGCATCGTGGCCGGCATCACCGGCGCCACAGG
TGCGGTTGCTGGCGCCTATATCGCCGACATCACCGATGGGGAGATCCTAAGATATCGCTTAGGCCACACGTTCAAGTG
CAGCCACAGGATAAATTTGCACTGAGCCTGGGTGGGATTCGGACTCGACCGCATAGCCTTCAGGAGTGAGTTTTGTG
CAATACCAACCGACGACTTGACCCTGCCAAGCGGCACCAGATTTCTTGCGTACGCGATCCCCTAAGCCAAAGGTGGC
ACTCAGGGGAAGCGCAAACTGCCCTGCAACGGGAGCGTTGGCTTCATCGCTACTTTGACCCATGTCGAATCCTTCTT
GTGAATCTATTATGGCGACAAGCAAATCGAGCTCTGACTGCCTACCCCACAACAACTATCAGAAAGCACCAGCACAA
CGGCTGCCTAACTTTGTTTTAGGGCGACTGCCCTGCTGCGTAACATCGTTGCTGCTCCATAACATCAAACATCGACC
CACGGCGTAACGCGCTTGCTGCTTGGATGCCCGAGGCATAGACTGTACAAAAAAACAGTCATAACAAGCCATGAAAA
CCGCCACTGCGCCGTTACCACCGCTGCGTTCGGTCAAGGTTCTGGACCAGTTGCGTGAGCGCATACGCTACTTGCAT
TACAGTTTACGAACCGAACAGGCTTATGTCAACTGGGTTCGTGCGAGCTCATCGATGCATTTAATCTTTCTGCGAAT
TGAGATGACGCCACTGGCTGGGCGTCATCCCGGTTTCCCGGGTAAACACCACCGAAAAATAGTTACTATCTTCAAAG
CCACATTCGGTCGAAATATCACTGATTAACAGGCGGCTATGCTGGAGAAGATATTGCGCATGACACACTCTGACCTG
TCGCAGATATTGATTGATGGTCATTCCAGTCTGCTGGCGAAATTGCTGACGCAAAACGCGCTCACTGCACGATGCCT
CATCACAAAATTTATCCAGCGCAAAGGGACTTTTCAGGCTAGCCGCCAGCCGGGTAATCAGCTTATCCAGCAACGTT
TCGCTGGATGTTGGCGGCAACGAATCACTGGTGTAACGATGGCGATTCAGCAACATCACCAACTGCCCGAACAGCAA
CTCAGCCATTTCGTTAGCAAACGGCACATGCTGACTACTTTCATGCTCAAGCTGACCGATAACCTGCCGCGCCTGCG
CCATCCCCATGCTACCTAAGCGCCAGTGTGGTTGCCCTGCGCTGGCGTTAAATCCCGGAATCGCCCCCTGCCAGTCA
AGATTCAGCTTCAGACGCTCCGGGCAATAAATAATATTCTGCAAAACCAGATCGTTAACGGAAGCGTAGGAGTGTTT
ATCGTCAGCATGAATGTAAAAGAGATCGCCACGGGTAATGCGATAAGGGCGATCGTTGAGTACATGCAGGCCATTAC
CGCGCCAGACAATCACCAGCTCACAAAAATCATGTGTATGTTCAGCAAAGACATCTTGCGGATAACGGTCAGCCACA
GCGACTGCCTGCTGGTCGCTGGCAAAAAAATCATCTTTGAGAAGTTTTAACTGATGCGCCACCGTGGCTACCTCGGC
CAGAGAACGAAGTTGATTATTCGCAATATGGCGTACAAATACGTTGAGAAGATTCGCGTTATTGCAGAAAGCCATCC
CGTCCCTGGCGAATATCACGCGGTGACCAGTTAAACTCTCGGCGAAAAAGCGTCGAAAAGTGGTTACTGTCGCTGAA
TCCACAGCGATAGGCGATGTCAGTAACGCTGGCCTCGCTGTGGCGTAGCAGATGTCGGGCTTTCATCAGTCGCAGGC
GGTTCAGGTATCGCTGAGGCGTCAGTCCCGTTTGCTGCTTAAGCTGCCGATGTAGCGTACGCAGTGAAAGAGAAAAT
TGATCCGCCACGGCATCCCAATTCACCTCATCGGCAAAATGGTCCTCCAGCCAGGCCAGAAGCAAGTTGAGACGTGA
TGCGCTGTTTTCCAGGTTCTCCTGCAAACTGCTTTTACGCAGCAAGAGCAGTAATTGCATAAACAAGATCTCGCGAC
TGGCGGTCGAGGGTAAATCATTTTCCCCTTCCTGCTGTTCCATCTGTGCAACCAGCTGTCGCACCTGCTGCAATACG
CTGTGGTTAACGCGCCAGTGAGACGGATACTGCCCATCCAGCTCTTGTGGCAGCAACTGATTCAGCCCGGCGAGAAA
CTGAAATCGATCCGGCGAGCGATACAGCACATTGGTCAGACACAGATTATCGGTATGTTCATACAGATGCCGATCAT
GATCGCGTACGAAACAGACCGTGCCACCGGTGATGGTATAGGGCTGCCCATTAAACACATGAATACCCGTGCCATGT
TCGACAATCACAATTTCATGAAAATCATGATGATGTTCAGGAAAATCCGCCTGCGGGAGCCGGGGTTCTATCGCCAC
GGACGCGTTACCAGACGGAAAAAAATCCACACTATGTAATACGGTCATACTGGCCTCCTGATGTCGTCAACACGGCG
AAATAGTAATCACGAGGTCAGGTTCTTACCTTAAATTTTCGACGGAAAACCACGTAAAAAACGTCGATTTTTCAAGA
TACAGCGTGAATTTTCAGGAAATGCGGTGAGCATCACATCACCACAATTCAGCAAATTGTGAACATCATCACGTTCA
TCTTTCCCTGGTTGCCAATGGCCCATTTTCCTGTCAGTAACGAGAAGGTCGCGAATTCAGGCGCTTTTTAGACTGGT
CGTAATGAAATTCAGCAGGATCACATA

B) Insert from genomic DNA of *B. thailandensis* E264 encoding all proteins. Removal of the first 833 bp from this insert (yellow) did not change the mass of modified ApyA after co-expression. Full construct name: pSCrhaB2_ApyOAHIDS

ATGCAAACAAGTGGTTTCCTCGACCTCGGACGGTGGCTGGGTCCACCGGTGGATGGGATTTCTGCCGAATCGAATGT
CGCCGACAGCGACAGGTTTTCGGATGTGGCGGTCTGGCTCGAGTCGTCTGCGAAGACATTTCTTTCCGCCGCAGCGC
GCCGTGCCGTGCGTGTCCGCCACGAGCCAGGGCAATTGCTGGTGCGGCCGACAGGCCGCGCCCGGGCGCGGCGAGGG

TTCGGGAATGCGGCGTGTCCGCTCACGCATGAGATCCGCGCGCATCGGGTATGGCTCGGGCGCTCGCGCCGGGGCCT
TCGTCCTCGACAAGCAGGAGGACGCTCGCGCGCCGGACGTCGCGGAACCCCCGGGGCGGAGCATGAAGGCCTGCTGT
CGTCTCCGATCGAGGGCGCGCTCATGAAGGCCGCACATGCCCGGGCATTTTGCGCGGCGCCGGACGTTTGGATGATT
TCAGGTGCGGCCGCTGATTATCGACTTCGCATTCGTTTGGCGTTTCCATCGGATGCCGACACCTCCGAGCCGGGAAC
GCTGTCGAACGAATTCGACGGGCCGCGCATATTTTCCGCGCCGTCTTGCCGGCGTATTGGCGCGAGCGGCGCTGCGC
GCGAATTTCCGATCGAGAAGCCGTATCGCGATGTCCACCCGGCGCCGATCGAGGACGATGCGACGAAGCTCGAACGT
CATGCGCGAGCGCGCGGGCGGCGTGGCGCGCGCGGTAACGTAGGCGCCGTCAGACGTGATACTGTGACGACGCCATG
CGCCATGGCGATCAGCCGGCGTAGCCAGGATCGGCACGAGTCGTTCACGGATACCGCTTGCAAATGAACGGTCTACC
TCCGTCGCTTCCGCGGGTCGACAGCACCGAGTCGCTTTTTGCGGAGCCGCTCGCGTTTCTCGCACAAGCGCGCTCGC
GGCATGGTGATGTCTTCGTCATGCGGGAGCATGGTCCGATTTTTTCCCGCGCGAGCGATTGCAGCGGCGTGATAGCG
GTATTCGGCGAGCATCGGCTTCGACAGATATTGACCGATATAGACAATTTCGCGTTGCCGATGTCCGCGGCGGCGAA
AATGGCATTGCCGAAAAACCTCGTCAACCTCAATCGCGGCTTGCACAGCATGCGCGAGCCGGAGCATGGCCGGCACA
AGCGCCTCTTGACGGGAACGATCAATCGCGAGTTGTTCGACGCGCACCGCTTCGAAATACGAGCGGCTTTGAACCGC
TTTTGCGAAATGCTGAAAGTGGACCGCCGGATTTCAGTGGTCAGCCGGATGCGCGAGCTGACCGTGGAGATGGCGTC
TCATATCTTTCTTGGGGCGCAGTGCCAAGAGGATGACGAACTGGCGTTTCTTTTAAGCGCCTATTTTACTTTGCGAC
GCGAAGCGTCTTCGCTGAACGCGCGCGATCCCCTGCTGTACCGGGACGAACTGATCGGCGTCGGGCAGCAACTCGAC
CGCACATTGCGGGAGCGGATCAGGCGGTATCGAAAGCGTCCCGTCGACGCTCGTGCGGGGCTGCTGCAACGTCTGGC
AACGGCCGGACCGCCGGGCTCGCCCGCGCTTTCCGAGGACGAGATAGTCGGCCATGCCAACGTGATGTTCGTGTCGA
GCACCGAGCCCGTCGCGATGTCGCTGACGTGGCTGCTGCTCGTTCTGTCGCAGCTGCCCGATCTGAGGCGCGCGCTT
CGCGCGGAAATTGCCGATCGCGCGTCGATGCCGGCATCGACGAACGGCGCGTCGTGGCTCGAGAACGTCGTCAACGA
GACGCTCCGGCTCCTGACGCCCAACGCGCTGATGGTTCGCGCGACGACGCGCGCGGTCTCGCTGCAAGGCGTCGCGC
TTCCGGCGCGTTGCGAGATCGTCGTGTGTCCGTTCCTCGCGCATCGCGAAGCCAAGCCGTTTCCGGACCCGCATGCA
TTCTCGCCGTCCCGGTGGGAGACGGCCAGGCCGTCTCCGTACGAGTATTTTCCGTTCGGCGCCGGCGGTCATTTTTG
CGCGGGGCGAAATCTCGCGCTCTCGCTGATTCGCGAAGTACTGTCAACGCTGCTATCGCGCTTCGATTTCGTTTTGG
ATGGCGAACAGTCTATCGACTGGCGCATTCATATCATGTTGATGCCGAAAGGCGACCCGGCACTCATCGCGCATCCC
GTCGATGAGCGCGGTGACACGCCGTCGCCGAAATGGCGCGGGCCGATAACCGATCTATTCCATTTCGCGCCGGGGCT
TTCTTGACATTTACCAGGGCGCATCCTTTAATCAGAAGCGAATCGGCATGCATGCCCATCGGGATTTTTTCCAGTCA
TTGATCCGCATGGTTCTTTTCACGGGTTATTCCGCGCTCACACGCTAAGCGGCGCGTCCTGCCTACAACAAAGGAAC
AATC**ATGCATCACCATCACCATCACATGGCTACGAAACCCAAAAAAACAAAGGGCGTTTCTGTTTCCATCGAAGGGA**
**AGCTGCCCAAGATGACCCTGGATATGCCGGTGGACGCGAAAAAGATCAAGGCCATCCAGAAATGCCTGGAAAACGGC**
**AAGCTGACCATCACGATGAGCAAGGTGGATCTTGCCGGCGGCCGCATGGGCGAGGGCTATCTCTACGACTGA**TGTCC
GGCGGCGCTCTGGTTTTCGCCGGCGGCGTCGCGATCTCGCCGCCTCGATGCTCGGGCCTCGGAGGGAACGTGCCGCC
GAACGCGTGCGTTTGGCGCGCGGTCGCGCGATAACGGGCGATGAATTCCGGGCTTCGGCAGTTCGACCTTAAGGGCG
CGGGAAGCCTATGAAAGCCTTCGATCGCGATGCATCGGAGCCTCGGATCGGTCTCGCGTATGGGCCCGGGATGGCGG
AATTCGCCGCACGGAACGCGCACCTCGTCGATTACATCGAAGTGCCGTTCGAGCAATTGCGATTCTCGCCGGCGGTC
GCCGAGCTTCAGCAAACCATTCCGTTCGTTCTGCACTGCGCCAGCTTGTCCGTCGCGGGATTCGTGCCGCCCGACGA
TTCGACGGTCGACGCAATCGAGCGCACGGCCGTGCAGACCGGCACGCCATGGATAGGCGAACATCTCGCGTACATTT
CGGCCGATCCCGTTGGCGAAGCGCTGGGCGGGACCGGTGAGCCCACCTCGTTGTCGTACACGTTGTGCCCTCAGTTG
AGCGACGAGACGGTACGCCGCGTCGTCGACAATCTCGCCGCGCTTCGTCCGCATTTTCCGGTTCCACTGATCGTCGA
AAACTCCCCGCAGTACTTTCCGATACCGGGCAGCACGATGGGAATGACGGACTTCATTCGCGCGATCACGGACCGAT
GCGACGTCGGTTTGCTCCTGGACCTGAGTCACTTCCTGATCACGGCGCACAACACGGGCGCAGAGGTGCATCGGGAA
CTCGCCCGGCTTCCGCTGGAGCGCGTGGTCGAAGTTCATCTGTCCGGAATGAGCGTGCAGTCCGGCACGGCCTGGGA
CGACCATTCGCTGCCGGCATCGCCGATCCTGTTCGAACTGCTCGAGCGCCTGCTCGACGTGGCGCGGCCTCGCGCGC
TGACCTTCGAATACAACTGGTCGCCATACTTCCCGCTGTCGGTCCTGACGACGCATATCGAGCGCGCACGCCAGTTG
ATGGGGCTCGCATGAACGCAGGGAAAATACACGCCGTGATCGCCGCCGGCGCCAAGGACCCGCGGCTCATCGAGCAT
TGGCGGCGGCGTCCGGACCTGCTGCGCCGCGCAGGAATCGAGCCGGATCATCTCGATTTGCGCGTATTGGAGAATTT
CGCCGGGTTGAGCCTGAAGGTGCGTCACAACGGCCTGCGCGGGGATCTGCCGCTGAGTTTCAGGTTGTTGAGCGTGG
CCGGCCTCGAGATCGAGGTCTTCGCCGCGTATGCGTCGGACTGCGCGGAAAGCGGCCATGTATTCGCCGCCACGCCG
CGGGAACGCGCGCGCGATCTGATGGCGTTCCTCGACAGATGGCTGGACCCCGCGCAACGAAATCACGTTCTGCTGTG
GGACATCCTTCGTCATGAATATGCGCTTTCGTGCCTGCGGCAAGAAAGCGCCGCGCCGCTACCCGCCGCTCGCCGCA
TCATCCGCCATCGGGATCGCGCGACAGGCGGCTGCATTCCGATGCTCAACGGCTCCGTCGCTCAGCACGTGATGCAT
TGCAATCCGGAAGACCTGGCGTCGGCGCTGCGTCAGAAGGAGCCGCCGCTCGATGCGGTTCCCGCCGCCGACACTTA
CTGCTGCTATTGGCGCTCCGCCGAAACGGGCGAAATCCGGTTTCTGCAGGCCGATACGCTCGGCTATTTCGCGATCG
AAGCGATCGACGGCCGGCGATCCGCCGCCGACATCTGCGATCGGCTGCTCGGCGGCCGCCGTCTGCTGCCGCAGTTC
CTCCGGTCGCTCGACCAACTGGCCGGAATCGGGATCGTGCGGTTCGAGAGGCCGCTCGGAAGCGCTGCGGCATGAAA
CTGCTGCTCGTCGACAACCTCGTCATGCCGGAAGAGGGCTCGCTCGCGTACCTCGACGTGCATCCGCATCTCGGGCT
GCTCGCGCTGGCCGCGGTCGCCGAAGCCGACGGCCATCGCGTGCAGATCTACGATCCGAAACGACTGATCAGGGACG

GCACGCTGGCTTACGATGCGACGCTTTACGAGCGGGCGAGCCAAGCGATTCTCGCGGCGCGTCCGGACGGGGTCGGC
TTCACGGCGCTCGGCTGCAGCTTTCTGTTTGCGCTCAATGTCGCGGCGCTGTTGAAGCGGCGCGAGCCCGACCTGCC
CGTCTTGCTCGGCGGGCCGCACGCGACGATGCTGCACCGGCAGATCCTCGAGCGGTTCCCGCAGTTCGACATCATCG
TCCGGTACGAGGCGGATGAAATCCTGCCCGCCGTGCTCGACTGCCTCCCGCACCGGACATTCGACGTGATTCCCGGC
TTGAGCTGGCGCGCGACGGGCCGCGGCTCGCCGTTGCGCTTCACCGACGGCAAGCCCAAGGTCGAGGATCTGGACCT
GCTGCCGATCGCCTCGTACGACCATTATCCGGTTGAGGAACTGGGGCTCTCGATGCTGCGCATCGAGGCCGGCCGAG
GGTGTCCGTTCGCATGCACATTCTGCTCGACGGCCGGTTTCTTTCAACGCAGCTTCCGCCTCAAATCGGCCGAGCGC
CTGGTTCGCGAACTGGATATCCTTCACCAGCGTTATCGCGTGTCGGACTTCAAGCTCGACCACGACATGTTCACCGT
GAATCGCCGCAAGGTAATGGAGTTTTGCGAGGCGGTCGCCGGGCGCGACTATCGGTGGCGCGCGTCGGCGCGGATCG
ACTGCGTCGACGAAGCATTGTTGAAGAAGATGGCCGACGCGGGCTGCGTGAATCTTTATTTCGGCGTGGAGACGGGC
TCCGAGCGCATGCAAAAGCTCTGCAAGAAGCGGCTGGATCTGCAACGGGTCGAGCCCATTCTGGCGGCGGCGGATTC
ATTCGGAATCGAGACGACCGCGTCGTTCATCACCGGCTATCCCGAGGAAACCGGGGAGGATCAGGACGACACGCTGG
ACATGATCGGGCGCTGCGCTCGGCGAGCGTCTTGCCTCACGCAACTGCACATGCTCGCGCCGGAGCCCGGCACACCG
CTGTTCGACGAACGCGGCGCGGAAATCGCCTACGACGGCTGTGGCGGGCCTTACAACACGCGCCTGCTGAGTTCGTC
GGACGAGCGTGCGGTGCTCGGCCATCCCGACATATTCCAAACCTACTATCACTACCCCGCCGCGCTGCCGCGTGCGC
GATACATCTTTGCCGTCGGGGCGGCGGATCTGCTGCGCCGGGTCGGGCCGGTCATTTTCGCGTATGCGTTGCGGGGG
TTCGGAGGGCGGCTGAGCACGCTGATATCGGCGTGGCGACAATTCGCCGACGAAACGTCGCCCGGCGCGCCCGCGGA
CGCGGCGGGGCTCGAGGCCTTCATTGCCGCGCGGTTCGGCCGCGGCCATCACCTGCATTCTCTTTTTCGGTACGCGT
TGCGGCTCTATGCGGCCCGCGCTCCGGCGGACGTAATCGTCGAGGGGCCCTATGAGCCGGACCGGCCTTGCATGCTG
GCCGAAAACGTCGGTCTGCTGGAGGACATTCACAATTGCGACGCATTGGTCGACCGGATCAGGCGATCGACCGAAGG
CGCGCCGATGCTCGGCGACGCGGATGTCGGCGGGCTCGGAACATATATCGTTCAGGGGCACGGCGAGGCCGCCACGG
GCTATTGGGTGGAGTCCGGCATCGCTACGCTTCTCGATCTGTTCAGGTCGCCCCGTTCGTGCCGCGCCGTTGCGCAA
TGGCTGGCGGACGCCACCCGCAATGACGGCATAGACGCTTCGATCTTCGAGCCGCTGCTGCGCAGAGGCATCCTGGT
GTCGGCCGGTTAAATCCATTTCGTCCATGGTGCGTCGCACACTCGAACGTTGGTCCGCGCTGGATCTCGCGGAAGGG
CTTCAACTGGCGCATGCCGTCGCCGCGCTGCAGGCGCTCGGCGTTCTGGACGCGATGAGGGAGCCGGTCACGGCGGA
AACGTTGAGCGCCAGCCATGATCTCGATCCCGAGTTGCTGCGCGGCATCCTCGAATATGCGGCGTCGAGAACGAATC
TGGTTCGCAAGACGGACCGCGGATTCGCGATTTCCGAGCATTACACCGCGCAAGCGCGCTTTCTGTTGAACCTGTAC
GCCGGCGCTTATGCCGATAACGCTTCACGGCTGGCGACGCTGCTCCGGCGTCCGGCATTGGCGGGCTCCATGATCGA
CCTCGTCGCGCACGCGCGCGCCTTCGACGCGAGCGAGGCCGGCGGTGGAGCGCTGGCGTCGATCGTCACGCAATTGC
GTCTCAATCATGTGCTGGATCTGGGATGCGGCGCCGGCGCGCTGCTGCGCGAGTTGGCGGCGGACGATCGCGGATTC
GTCGGCTGGGGACTGGATCGCAATCCCGCGATGTGCAAGGCGGCGCGGGTTCGAGCGCGGCAAGCGGGGCTCGCCGC
GCGCGTGAAGGTGTTCCACGGCGACGGCTGGAATCCCGGGGCGTCGGTCCCGCCGCGCGTGCTGGCGCGCGTGCGCA
ACGTCGTCGCGAGCCAGTTCGTCAACGAGATGTTTCGCGGCGGCACATCGCGAGTGGAAACGTGGCTTCGGCGCATG
CGGCGCTTGCTGCCCGGCCGCATGCTCGTGATATCGGACTATTACGGCCGCCTCGGCCACGGCACTCACACGATGCG
CCGCGAGACGCTGCTGCATGACTATGTGCAGCTCATATCGGGGCAGGGCGTGCCGCCGCCCGATTCCGACGCGTGGT
TGGCCATGTATCGAGCAGCCGGTTGCCGGCCTCTGCACGTGATCGAAGACCGGGCGGCCACGACATGCTTCGTCCAT
CTGGTGGGGCTATAG

C) Insert from genomic DNA of *B. thailandensis* E264 encoding all proteins, with the gene encoding ApyO removed. Full construct name: pSCrhaB2_ApyAHIDS

ATGAACGGTCTACCTCCGTCGCTTCCGCGGGTCGACAGCACCGAGTCGCTTTTTGCGGAGCCGCTCGCGTTTCTCGC
ACAAGCGCGCTCGCGGCATGGTGATCCGGCACTCATCGCGCATCCCGTCGATGAGCGCGGTGACACGCCGTCGCCGA
AATGGCGCGGGCCGATAACCGATCTATTCCATTTCGCGCCGGGGCTTTCTTGACATTTACCAGGGCGCATCCTTTAA
TCAGAAGCGAATCGGCATGCATGCCCATCGGGATTTTTTCCAGTCATTGATCCGCATGGTTCTTTTCACGGGTTATT
CCGCGCTCACACGCTAAGCGGCGCGTCCTGCCTACAACAAAGGAACAATC**ATGCATCACCATCACCATCACATGGCT**
**ACGAAACCCAAAAAAACAAAGGGCGTTTCTGTTTCCATCGAAGGGAAGCTGCCCAAGATGACCCTGGATATGCCGGT**
**GGACGCGAAAAAGATCAAGGCCATCCAGAAATGCCTGGAAAACGGCAAGCTGACCATCACGATGAGCAAGGTGGATC**
**TTGCCGGCGGCCGCATGGGCGAGGGCTATCTCTACGACTGA**TGTCCGGCGGCGCTCTGGTTTTCGCCGGCGGCGTCG
CGATCTCGCCGCCTCGATGCTCGGGCCTCGGAGGGAACGTGCCGCCGAACGCGTGCGTTTGGCGCGCGGTCGCGCGA
TAACGGGCGATGAATTCCGGGCTTCGGCAGTTCGACCTTAAGGGCGCGGGAAGCCTATGAAAGCCTTCGATCGCGAT
GCATCGGAGCCTCGGATCGGTCTCGCGTATGGGCCCGGGATGGCGGAATTCGCCGCACGGAACGCGCACCTCGTCGA
TTACATCGAAGTGCCGTTCGAGCAATTGCGATTCTCGCCGGCGGTCGCCGAGCTTCAGCAAACCATTCCGTTCGTTC
TGCACTGCGCCAGCTTGTCCGTCGCGGGATTCGTGCCGCCCGACGATTCGACGGTCGACGCAATCGAGCGCACGGCC
GTGCAGACCGGCACGCCATGGATAGGCGAACATCTCGCGTACATTTCGGCCGATCCCGTTGGCGAAGCGCTGGGCGG
GACCGGTGAGCCCACCTCGTTGTCGTACACGTTGTGCCCTCAGTTGAGCGACGAGACGGTACGCCGCGTCGTCGACA

ATCTCGCCGCGCTTCGTCCGCATTTTCCGGTTCCACTGATCGTCGAAAACTCCCCGCAGTACTTTCCGATACCGGGC
AGCACGATGGGAATGACGGACTTCATTCGCGCGATCACGGACCGATGCGACGTCGGTTTGCTCCTGGACCTGAGTCA
CTTCCTGATCACGGCGCACAACACGGGCGCAGAGGTGCATCGGGAACTCGCCCGGCTTCCGCTGGAGCGCGTGGTCG
AAGTTCATCTGTCCGGAATGAGCGTGCAGTCCGGCACGGCCTGGGACGACCATTCGCTGCCGGCATCGCCGATCCTG
TTCGAACTGCTCGAGCGCCTGCTCGACGTGGCGCGGCCTCGCGCGCTGACCTTCGAATACAACTGGTCGCCATACTT
CCCGCTGTCGGTCCTGACGACGCATATCGAGCGCGCACGCCAGTTGATGGGGCTCGCATGAACGCAGGGAAAATACA
CGCCGTGATCGCCGCCGGCGCCAAGGACCCGCGGCTCATCGAGCATTGGCGGCGGCGTCCGGACCTGCTGCGCCGCG
CAGGAATCGAGCCGGATCATCTCGATTTGCGCGTATTGGAGAATTTCGCCGGGTTGAGCCTGAAGGTGCGTCACAAC
GGCCTGCGCGGGGATCTGCCGCTGAGTTTCAGGTTGTTGAGCGTGGCCGGCCTCGAGATCGAGGTCTTCGCCGCGTA
TGCGTCGGACTGCGCGGAAAGCGGCCATGTATTCGCCGCCACGCCGCGGGAACGCGCGCGCGATCTGATGGCGTTCC
TCGACAGATGGCTGGACCCCGCGCAACGAAATCACGTTCTGCTGTGGGACATCCTTCGTCATGAATATGCGCTTTCG
TGCCTGCGGCAAGAAAGCGCCGCGCCGCTACCCGCCGCTCGCCGCATCATCCGCCATCGGGATCGCGCGACAGGCGG
CTGCATTCCGATGCTCAACGGCTCCGTCGCTCAGCACGTGATGCATTGCAATCCGGAAGACCTGGCGTCGGCGCTGC
GTCAGAAGGAGCCGCCGCTCGATGCGGTTCCCGCCGCCGACACTTACTGCTGCTATTGGCGCTCCGCCGAAACGGGC
GAAATCCGGTTTCTGCAGGCCGATACGCTCGGCTATTTCGCGATCGAAGCGATCGACGGCCGGCGATCCGCCGCCGA
CATCTGCGATCGGCTGCTCGGCGGCCGCCGTCTGCTGCCGCAGTTCCTCCGGTCGCTCGACCAACTGGCCGGAATCG
GGATCGTGCGGTTCGAGAGGCCGCTCGGAAGCGCTGCGGCATGAAACTGCTGCTCGTCGACAACCTCGTCATGCCGG
AAGAGGGCTCGCTCGCGTACCTCGACGTGCATCCGCATCTCGGGCTGCTCGCGCTGGCCGCGGTCGCCGAAGCCGAC
GGCCATCGCGTGCAGATCTACGATCCGAAACGACTGATCAGGGACGGCACGCTGGCTTACGATGCGACGCTTTACGA
GCGGGCGAGCCAAGCGATTCTCGCGGCGCGTCCGGACGGGGTCGGCTTCACGGCGCTCGGCTGCAGCTTTCTGTTTG
CGCTCAATGTCGCGGCGCTGTTGAAGCGGCGCGAGCCCGACCTGCCCGTCTTGCTCGGCGGGCCGCACGCGACGATG
CTGCACCGGCAGATCCTCGAGCGGTTCCCGCAGTTCGACATCATCGTCCGGTACGAGGCGGATGAAATCCTGCCCGC
CGTGCTCGACTGCCTCCCGCACCGGACATTCGACGTGATTCCCGGCTTGAGCTGGCGCGCGACGGGCCGCGGCTCGC
CGTTGCGCTTCACCGACGGCAAGCCCAAGGTCGAGGATCTGGACCTGCTGCCGATCGCCTCGTACGACCATTATCCG
GTTGAGGAACTGGGGCTCTCGATGCTGCGCATCGAGGCCGGCCGAGGGTGTCCGTTCGCATGCACATTCTGCTCGAC
GGCCGGTTTCTTTCAACGCAGCTTCCGCCTCAAATCGGCCGAGCGCCTGGTTCGCGAACTGGATATCCTTCACCAGC
GTTATCGCGTGTCGGACTTCAAGCTCGACCACGACATGTTCACCGTGAATCGCCGCAAGGTAATGGAGTTTTGCGAG
GCGGTCGCCGGGCGCGACTATCGGTGGCGCGCGTCGGCGCGGATCGACTGCGTCGACGAAGCATTGTTGAAGAAGAT
GGCCGACGCGGGCTGCGTGAATCTTTATTTCGGCGTGGAGACGGGCTCCGAGCGCATGCAAAAGCTCTGCAAGAAGC
GGCTGGATCTGCAACGGGTCGAGCCCATTCTGGCGGCGGCGGATTCATTCGGAATCGAGACGACCGCGTCGTTCATC
ACCGGCTATCCCGAGGAAACCGGGGAGGATCAGGACGACACGCTGGACATGATCGGGCGCTGCGCTCGGCGAGCGTC
TTGCCTCACGCAACTGCACATGCTCGCGCCGGAGCCCGGCACACCGCTGTTCGACGAACGCGGCGCGGAAATCGCCT
ACGACGGCTGTGGCGGGCCTTACAACACGCGCCTGCTGAGTTCGTCGGACGAGCGTGCGGTGCTCGGCCATCCCGAC
ATATTCCAAACCTACTATCACTACCCCGCCGCGCTGCCGCGTGCGCGATACATCTTTGCCGTCGGGGCGGCGGATCT
GCTGCGCCGGGTCGGGCCGGTCATTTTCGCGTATGCGTTGCGGGGGTTCGGAGGGCGGCTGAGCACGCTGATATCGG
CGTGGCGACAATTCGCCGACGAAACGTCGCCCGGCGCGCCCGCGGACGCGGCGGGGCTCGAGGCCTTCATTGCCGCG
CGGTTCGGCCGCGGCCATCACCTGCATTCTCTTTTTCGGTACGCGTTGCGGCTCTATGCGGCCCGCGCTCCGGCGGA
CGTAATCGTCGAGGGGCCCTATGAGCCGGACCGGCCTTGCATGCTGGCCGAAAACGTCGGTCTGCTGGAGGACATTC
ACAATTGCGACGCATTGGTCGACCGGATCAGGCGATCGACCGAAGGCGCGCCGATGCTCGGCGACGCGGATGTCGGC
GGGCTCGGAACATATATCGTTCAGGGGCACGGCGAGGCCGCCACGGGCTATTGGGTGGAGTCCGGCATCGCTACGCT
TCTCGATCTGTTCAGGTCGCCCCGTTCGTGCCGCGCCGTTGCGCAATGGCTGGCGGACGCCACCCGCAATGACGGCA
TAGACGCTTCGATCTTCGAGCCGCTGCTGCGCAGAGGCATCCTGGTGTCGGCCGGTTAAATCCATTTCGTCCATGGT
GCGTCGCACACTCGAACGTTGGTCCGCGCTGGATCTCGCGGAAGGGCTTCAACTGGCGCATGCCGTCGCCGCGCTGC
AGGCGCTCGGCGTTCTGGACGCGATGAGGGAGCCGGTCACGGCGGAAACGTTGAGCGCCAGCCATGATCTCGATCCC
GAGTTGCTGCGCGGCATCCTCGAATATGCGGCGTCGAGAACGAATCTGGTTCGCAAGACGGACCGCGGATTCGCGAT
TTCCGAGCATTACACCGCGCAAGCGCGCTTTCTGTTGAACCTGTACGCCGGCGCTTATGCCGATAACGCTTCACGGC
TGGCGACGCTGCTCCGGCGTCCGGCATTGGCGGGCTCCATGATCGACCTCGTCGCGCACGCGCGCGCCTTCGACGCG
AGCGAGGCCGGCGGTGGAGCGCTGGCGTCGATCGTCACGCAATTGCGTCTCAATCATGTGCTGGATCTGGGATGCGG
CGCCGGCGCGCTGCTGCGCGAGTTGGCGGCGGACGATCGCGGATTCGTCGGCTGGGGACTGGATCGCAATCCCGCGA
TGTGCAAGGCGGCGCGGGTTCGAGCGCGGCAAGCGGGGCTCGCCGCGCGCGTGAAGGTGTTCCACGGCGACGGCTGG
AATCCCGGGGCGTCGGTCCCGCCGCGCGTGCTGGCGCGCGTGCGCAACGTCGTCGCGAGCCAGTTCGTCAACGAGAT
GTTTCGCGGCGGCACATCGCGAGTGGAAACGTGGCTTCGGCGCATGCGGCGCTTGCTGCCCGGCCGCATGCTCGTGA
TATCGGACTATTACGGCCGCCTCGGCCACGGCACTCACACGATGCGCCGCGAGACGCTGCTGCATGACTATGTGCAG
CTCATATCGGGGCAGGGCGTGCCGCCGCCCGATTCCGACGCGTGGTTGGCCATGTATCGAGCAGCCGGTTGCCGGCC
TCTGCACGTGATCGAAGACCGGGCGGCCACGACATGCTTCGTCCATCTGGTGGGGCTATAG

D) Insert from genomic DNA of *B. thailandensis* E264 encoding all proteins, with the gene encoding ApyH removed. Full construct name: pSCrhaB2_ApyOAIDS

ATGAACGGTCTACCTCCGTCGCTTCCGCGGGTCGACAGCACCGAGTCGCTTTTTGCGGAGCCGCTCGCGTTTCTCGC
ACAAGCGCGCTCGCGGCATGGTGATGTCTTCGTCATGCGGGAGCATGGTCCGATTTTTTCCCGCGCGAGCGATTGCA
GCGGCGTGATAGCGGTATTCGGCGAGCATCGGCTTCGACAGATATTGACCGATATAGACAATTTCGCGTTGCCGATG
TCCGCGGCGGCGAAAATGGCATTGCCGAAAAACCTCGTCAACCTCAATCGCGGCTTGCACAGCATGCGCGAGCCGGA
GCATGGCCGGCACAAGCGCCTCTTGACGGGAACGATCAATCGCGAGTTGTTCGACGCGCACCGCTTCGAAATACGAG
CGGCTTTGAACCGCTTTTGCGAAATGCTGAAAGTGGACCGCCGGATTTCAGTGGTCAGCCGGATGCGCGAGCTGACC
GTGGAGATGGCGTCTCATATCTTTCTTGGGGCGCAGTGCCAAGAGGATGACGAACTGGCGTTTCTTTTAAGCGCCTA
TTTTACTTTGCGACGCGAAGCGTCTTCGCTGAACGCGCGCGATCCCCTGCTGTACCGGGACGAACTGATCGGCGTCG
GGCAGCAACTCGACCGCACATTGCGGGAGCGGATCAGGCGGTATCGAAAGCGTCCCGTCGACGCTCGTGCGGGGCTG
CTGCAACGTCTGGCAACGGCCGGACCGCCGGGCTCGCCCGCGCTTTCCGAGGACGAGATAGTCGGCCATGCCAACGT
GATGTTCGTGTCGAGCACCGAGCCCGTCGCGATGTCGCTGACGTGGCTGCTGCTCGTTCTGTCGCAGCTGCCCGATC
TGAGGCGCGCGCTTCGCGCGGAAATTGCCGATCGCGCGTCGATGCCGGCATCGACGAACGGCGCGTCGTGGCTCGAG
AACGTCGTCAACGAGACGCTCCGGCTCCTGACGCCCAACGCGCTGATGGTTCGCGCGACGACGCGCGCGGTCTCGCT
GCAAGGCGTCGCGCTTCCGGCGCGTTGCGAGATCGTCGTGTGTCCGTTCCTCGCGCATCGCGAAGCCAAGCCGTTTC
CGGACCCGCATGCATTCTCGCCGTCCCGGTGGGAGACGGCCAGGCCGTCTCCGTACGAGTATTTTCCGTTCGGCGCC
GGCGGTCATTTTTGCGCGGGGCGAAATCTCGCGCTCTCGCTGATTCGCGAAGTACTGTCAACGCTGCTATCGCGCTT
CGATTTCGTTTTGGATGGCGAACAGTCTATCGACTGGCGCATTCATATCATGTTGATGCCGAAAGGCGACCCGGCAC
TCATCGCGCATCCCGTCGATGAGCGCGGTGACACGCCGTCGCCGAAATGGCGCGGGCCGATAACCGATCTATTCCAT
TTCGCGCCGGGGCTTTCTTGACATTTACCAGGGCGCATCCTTTAATCAGAAGCGAATCGGCATGCATGCCCATCGGG
ATTTTTTCCAGTCATTGATCCGCATGGTTCTTTTCACGGGTTATTCCGCGCTCACACGCTAAGCGGCGCGTCCTGCC
TACAACAAAGGAACAATC**ATGCATCACCATCACCATCACATGGCTACGAAACCCAAAAAAACAAAGGGCGTTTCTGT**
**TTCCATCGAAGGGAAGCTGCCCAAGATGACCCTGGATATGCCGGTGGACGCGAAAAAGATCAAGGCCATCCAGAAAT**
**GCCTGGAAAACGGCAAGCTGACCATCACGATGAGCAAGGTGGATCTTGCCGGCGGCCGCATGGGCGAGGGCTATCTC**
**TACGACTGA**TGTCCGGCGGCGCTCTGGTTTTCGCCGGCGGCGTCGCGATCTCGCCGCCTCGATGCTCGGGCCTCGGA
GGGAACGTGCCGCCGAACGCGTGCGTTTGGCGCGCGGTCGCGCGATAACGGGCGATGAATTCCGGGCTTCGGCAGTT
CGACCTTAAGGGCGCGGGAAGCCTATGAAAGCCTTCGATCGCGATGCATCGGAGCCTCGGATCGGTCTCGCGTATGG
GCCCGGGATGGCGGAATTCGCCGCACGGAACGCGCACCTCGTCGATTACCGCGCGCTGACCTTCGAATACAACTGGT
CGCCATACTTCCCGCTGTCGGTCCTGACGACGCATATCGAGCGCGCACGCCAGTTGATGGGGCTCGCATGAACGCAG
GGAAAATACACGCCGTGATCGCCGCCGGCGCCAAGGACCCGCGGCTCATCGAGCATTGGCGGCGGCGTCCGGACCTG
CTGCGCCGCGCAGGAATCGAGCCGGATCATCTCGATTTGCGCGTATTGGAGAATTTCGCCGGGTTGAGCCTGAAGGT
GCGTCACAACGGCCTGCGCGGGGATCTGCCGCTGAGTTTCAGGTTGTTGAGCGTGGCCGGCCTCGAGATCGAGGTCT
TCGCCGCGTATGCGTCGGACTGCGCGGAAAGCGGCCATGTATTCGCCGCCACGCCGCGGGAACGCGCGCGCGATCTG
ATGGCGTTCCTCGACAGATGGCTGGACCCCGCGCAACGAAATCACGTTCTGCTGTGGGACATCCTTCGTCATGAATA
TGCGCTTTCGTGCCTGCGGCAAGAAAGCGCCGCGCCGCTACCCGCCGCTCGCCGCATCATCCGCCATCGGGATCGCG
CGACAGGCGGCTGCATTCCGATGCTCAACGGCTCCGTCGCTCAGCACGTGATGCATTGCAATCCGGAAGACCTGGCG
TCGGCGCTGCGTCAGAAGGAGCCGCCGCTCGATGCGGTTCCCGCCGCCGACACTTACTGCTGCTATTGGCGCTCCGC
CGAAACGGGCGAAATCCGGTTTCTGCAGGCCGATACGCTCGGCTATTTCGCGATCGAAGCGATCGACGGCCGGCGAT
CCGCCGCCGACATCTGCGATCGGCTGCTCGGCGGCCGCCGTCTGCTGCCGCAGTTCCTCCGGTCGCTCGACCAACTG
GCCGGAATCGGGATCGTGCGGTTCGAGAGGCCGCTCGGAAGCGCTGCGGCATGAAACTGCTGCTCGTCGACAACCTC
GTCATGCCGGAAGAGGGCTCGCTCGCGTACCTCGACGTGCATCCGCATCTCGGGCTGCTCGCGCTGGCCGCGGTCGC
CGAAGCCGACGGCCATCGCGTGCAGATCTACGATCCGAAACGACTGATCAGGGACGGCACGCTGGCTTACGATGCGA
CGCTTTACGAGCGGGCGAGCCAAGCGATTCTCGCGGCGCGTCCGGACGGGGTCGGCTTCACGGCGCTCGGCTGCAGC
TTTCTGTTTGCGCTCAATGTCGCGGCGCTGTTGAAGCGGCGCGAGCCCGACCTGCCCGTCTTGCTCGGCGGGCCGCA
CGCGACGATGCTGCACCGGCAGATCCTCGAGCGGTTCCCGCAGTTCGACATCATCGTCCGGTACGAGGCGGATGAAA
TCCTGCCCGCCGTGCTCGACTGCCTCCCGCACCGGACATTCGACGTGATTCCCGGCTTGAGCTGGCGCGCGACGGGC
CGCGGCTCGCCGTTGCGCTTCACCGACGGCAAGCCCAAGGTCGAGGATCTGGACCTGCTGCCGATCGCCTCGTACGA
CCATTATCCGGTTGAGGAACTGGGGCTCTCGATGCTGCGCATCGAGGCCGGCCGAGGGTGTCCGTTCGCATGCACAT
TCTGCTCGACGGCCGGTTTCTTTCAACGCAGCTTCCGCCTCAAATCGGCCGAGCGCCTGGTTCGCGAACTGGATATC
CTTCACCAGCGTTATCGCGTGTCGGACTTCAAGCTCGACCACGACATGTTCACCGTGAATCGCCGCAAGGTAATGGA
GTTTTGCGAGGCGGTCGCCGGGCGCGACTATCGGTGGCGCGCGTCGGCGCGGATCGACTGCGTCGACGAAGCATTGT
TGAAGAAGATGGCCGACGCGGGCTGCGTGAATCTTTATTTCGGCGTGGAGACGGGCTCCGAGCGCATGCAAAAGCTC
TGCAAGAAGCGGCTGGATCTGCAACGGGTCGAGCCCATTCTGGCGGCGGCGGATTCATTCGGAATCGAGACGACCGC
GTCGTTCATCACCGGCTATCCCGAGGAAACCGGGGAGGATCAGGACGACACGCTGGACATGATCGGGCGCTGCGCTC

GGCGAGCGTCTTGCCTCACGCAACTGCACATGCTCGCGCCGGAGCCCGGCACACCGCTGTTCGACGAACGCGGCGCG
GAAATCGCCTACGACGGCTGTGGCGGGCCTTACAACACGCGCCTGCTGAGTTCGTCGGACGAGCGTGCGGTGCTCGG
CCATCCCGACATATTCCAAACCTACTATCACTACCCCGCCGCGCTGCCGCGTGCGCGATACATCTTTGCCGTCGGGG
CGGCGGATCTGCTGCGCCGGGTCGGCCGGTCATTTTCGCGTATGCGTTGCGGGGGTTCGGAGGGCGGCTGAGCACG
CTGATATCGGCGTGGCGACAATTCGCCGACGAAACGTCGCCCGGCGCGCCCGCGGACGCGGCGGGGCTCGAGGCCTT
CATTGCCGCGCGGTTCGGCCGCGGCCATCACCTGCATTCTCTTTTTCGGTACGCGTTGCGGCTCTATGCGGCCCGCG
CTCCGGCGGACGTAATCGTCGAGGGGCCCTATGAGCCGGACCGGCCTTGCATGCTGGCCGAAAACGTCGGTCTGCTG
GAGGACATTCACAATTGCGACGCATTGGTCGACCGGATCAGGCGATCGACCGAAGGCGCGCCGATGCTCGGCGACGC
GGATGTCGGCGGGCTCGGAACATATATCGTTCAGGGGCACGGCGAGGCCGCCACGGGCTATTGGGTGGAGTCCGGCA
TCGCTACGCTTCTCGATCTGTTCAGGTCGCCCCGTTCGTGCCGCGCCGTTGCGCAATGGCTGGCGGACGCCACCCGC
AATGACGGCATAGACGCTTCGATCTTCGAGCCGCTGCTGCGCAGAGGCATCCTGGTGTCGGCCGGTTAAATCCATTT
CGTCCATGGTGCGTCGCACACTCGAACGTTGGTCCGCGCTGGATCTCGCGGAAGGGCTTCAACTGGCGCATGCCGTC
GCCGCGCTGCAGGCGCTCGGCGTTCTGGACGCGATGAGGGAGCCGGTCACGGCGGAAACGTTGAGCGCCAGCCATGA
TCTCGATCCCGAGTTGCTGCGCGGCATCCTCGAATATGCGGCGTCGAGAACGAATCTGGTTCGCAAGACGGACCGCG
GATTCGCGATTTCCGAGCATTACACCGCGCAAGCGCGCTTTCTGTTGAACCTGTACGCCGGCGCTTATGCCGATAAC
GCTTCACGGCTGGCGACGCTGCTCCGGCGTCCGGCATTGGCGGGCTCCATGATCGACCTCGTCGCGCACGCGCGCGC
CTTCGACGCGAGCGAGGCCGGCGGTGGAGCGCTGGCGTCGATCGTCACGCAATTGCGTCTCAATCATGTGCTGGATC
TGGGATGCGGCGCCGGCGCGCTGCTGCGCGAGTTGGCGGCGGACGATCGCGGATTCGTCGGCTGGGGACTGGATCGC
AATCCCGCGATGTGCAAGGCGGCGCGGGTTCGAGCGCGGCAAGCGGGGCTCGCCGCGCGCGTGAAGGTGTTCCACGG
CGACGGCTGGAATCCCGGGGCGTCGGTCCCGCCGCGCGTGCTGGCGCGCGTGCGCAACGTCGTCGCGAGCCAGTTCG
TCAACGAGATGTTTCGCGGCGGCACATCGCGAGTGGAAACGTGGCTTCGGCGCATGCGGCGCTTGCTGCCCGGCCGC
ATGCTCGTGATATCGGACTATTACGGCCGCCTCGGCCACGGCACTCACACGATGCGCCGCGAGACGCTGCTGCATGA
CTATGTGCAGCTCATATCGGGGCAGGGCGTGCCGCCGCCCGATTCCGACGCGTGGTTGGCCATGTATCGAGCAGCCG
GTTGCCGGCCTCTGCACGTGATCGAAGACCGGGCGGCCACGACATGCTTCGTCCATCTGGTGGGGCTATAG

E) Insert from genomic DNA of *B. thailandensis* E264 encoding all proteins, with the gene encoding ApyI removed. Full construct name: pSCrhaB2_ApyOAHDS

ATGAACGGTCTACCTCCGTCGCTTCCGCGGGTCGACAGCACCGAGTCGCTTTTTGCGGAGCCGCTCGCGTTTCTCGC
ACAAGCGCGCTCGCGGCATGGTGATGTCTTCGTCATGCGGGAGCATGGTCCGATTTTTTCCCGCGCGAGCGATTGCA
GCGGCGTGATAGCGGTATTCGGCGAGCATCGGCTTCGACAGATATTGACCGATATAGACAATTTCGCGTTGCCGATG
TCCGCGGCGGCGAAAATGGCATTGCCGAAAAACCTCGTCAACCTCAATCGCGGCTTGCACAGCATGCGCGAGCCGGA
GCATGGCCGGCACAAGCGCCTCTTGACGGGAACGATCAATCGCGAGTTGTTCGACGCGCACCGCTTCGAAATACGAG
CGGCTTTGAACCGCTTTTGCGAAATGCTGAAAGTGGACCGCCGGATTTCAGTGGTCAGCCGGATGCGCGAGCTGACC
GTGGAGATGGCGTCTCATATCTTTCTTGGGGCGCAGTGCCAAGAGGATGACGAACTGGCGTTTCTTTTAAGCGCCTA
TTTTACTTTGCGACGCGAAGCGTCTTCGCTGAACGCGCGCGATCCCCTGCTGTACCGGGACGAACTGATCGGCGTCG
GGCAGCAACTCGACCGCACATTGCGGGAGCGGATCAGGCGGTATCGAAAGCGTCCCGTCGACGCTCGTGCGGGGCTG
CTGCAACGTCTGGCAACGGCCGGACCGCCGGGCTCGCCCGCGCTTTCCGAGGACGAGATAGTCGGCCATGCCAACGT
GATGTTCGTGTCGAGCACCGAGCCCGTCGCGATGTCGCTGACGTGGCTGCTGCTCGTTCTGTCGCAGCTGCCCGATC
TGAGGCGCGCGCTTCGCGCGGAAATTGCCGATCGCGCGTCGATGCCGGCATCGACGAACGGCGCGTCGTGGCTCGAG
AACGTCGTCAACGAGACGCTCCGGCTCCTGACGCCCAACGCGCTGATGGTTCGCGCGACGACGCGCGCGGTCTCGCT
GCAAGGCGTCGCGCTTCCGGCGCGTTGCGAGATCGTCGTGTGTCCGTTCCTCGCGCATCGCGAAGCCAAGCCGTTTC
CGGACCCGCATGCATTCTCGCCGTCCCGGTGGGAGACGGCCAGGCCGTCTCCGTACGAGTATTTTCCGTTCGGCGCC
GGCGGTCATTTTTGCGCGGGGCGAAATCTCGCGCTCTCGCTGATTCGCGAAGTACTGTCAACGCTGCTATCGCGCTT
CGATTTCGTTTTGGATGGCGAACAGTCTATCGACTGGCGCATTCATATCATGTTGATGCCGAAAGGCGACCCGGCAC
TCATCGCGCATCCCGTCGATGAGCGCGGTGACACGCCGTCGCCGAAATGGCGCGGGCCGATAACCGATCTATTCCAT
TTCGCGCCGGGGCTTTCTTGACATTTACCAGGGCGCATCCTTTAATCAGAAGCGAATCGGCATGCATGCCCATCGGG
ATTTTTTCCAGTCATTGATCCGCATGGTTCTTTTCACGGGTTATTCCGCGCTCACACGCTAAGCGGCGCGTCCTGCC
TACAACAAAGGAACAATC**ATGCATCACCATCACCATCACATGGCTACGAAACCCAAAAAAACAAAGGGCGTTTCTGT**
**TTCCATCGAAGGGAAGCTGCCCAAGATGACCCTGGATATGCCGGTGGACGCGAAAAAGATCAAGGCCATCCAGAAAT**
**GCCTGGAAAACGGCAAGCTGACCATCACGATGAGCAAGGTGGATCTTGCCGGCGGCCGCATGGGCGAGGGCTATCTC**
**TACGACTGA**TGTCCGGCGGCGCTCTGGTTTTCGCCGGCGGCGTCGCGATCTCGCCGCCTCGATGCTCGGGCCTCGGA
GGGAACGTGCCGCCGAACGCGTGCGTTTGGCGCGCGGTCGCGCGATAACGGGCGATGAATTCCGGGCTTCGGCAGTT
CGACCTTAAGGGCGCGGGAAGCCTATGAAAGCCTTCGATCGCGATGCATCGGAGCCTCGGATCGGTCTCGCGTATGG
GCCCGGGATGGCGGAATTCGCCGCACGGAACGCGCACCTCGTCGATTACATCGAAGTGCCGTTCGAGCAATTGCGAT
TCTCGCCGGCGGTCGCCGAGCTTCAGCAAACCATTCCGTTCGTTCTGCACTGCGCCAGCTTGTCCGTCGCGGGATTC
GTGCCGCCCGACGATTCGACGGTCGACGCAATCGAGCGCACGGCCGTGCAGACCGGCACGCCATGGATAGGCGAACA

TCTCGCGTACATTTCGGCCGATCCCGTTGGCGAAGCGCTGGGCGGGACCGGTGAGCCCACCTCGTTGTCGTACACGT
TGTGCCCTCAGTTGAGCGACGAGACGGTACGCCGCGTCGTCGACAATCTCGCCGCGCTTCGTCCGCATTTTCCGGTT
CCACTGATCGTCGAAAACTCCCCGCAGTACTTTCCGATACCGGGCAGCACGATGGGAATGACGGACTTCATTCGCGC
GATCACGGACCGATGCGACGTCGGTTTGCTCCTGGACCTGAGTCACTTCCTGATCACGGCGCACAACACGGGCGCAG
AGGTGCATCGGGAACTCGCCCGGCTTCCGCTGGAGCGCGTGGTCGAAGTTCATCTGTCCGGAATGAGCGTGCAGTCC
GGCACGGCCTGGGACGACCATTCGCTGCCGGCATCGCCGATCCTGTTCGAACTGCTCGAGCGCCTGCTCGACGTGGC
GCGGCCTCGCGCGCTGACCTTCGAATACAACTGGTCGCCATACTTCCCGCTGTCGGTCCTGACGACGCATATCGAGC
GCGCACGCCAGTTGATGGGGCTCGCATGAAACGCAGGGAAAATACACGCCGTGATCGCCGCCGGCGCCAAGGACCCG
CGGCTCATCGAGCATTGGCGGCGGCGTCCGGACCTGCTGCGCCGCGCAGGAATCGAGCCGGATGATCGGCTGCTCGG
CGGCCGCCGTCTGCTGCCGCAGTTCCTCCGGTCGCTCGACCAACTGGCCGGAATCGGGATCGTGCGGTTCGAGAGGC
CGCTCGGAAGCGCTGCGGCATGAAACTGCTGCTCGTCGACAACCTCGTCATGCCGGAAGAGGGCTCGCTCGCGTACC
TCGACGTGCATCCGCATCTCGGGCTGCTCGCGCTGGCCGCGGTCGCCGAAGCCGACGGCCATCGCGTGCAGATCTAC
GATCCGAAACGACTGATCAGGGACGGCACGCTGGCTTACGATGCGACGCTTTACGAGCGGGCGAGCCAAGCGATTCT
CGCGGCGCGTCCGGACGGGGTCGGCTTCACGGCGCTCGGCTGCAGCTTTCTGTTTGCGCTCAATGTCGCGGCGCTGT
TGAAGCGGCGCGAGCCCGACCTGCCCGTCTTGCTCGGCGGGCCGCACGCGACGATGCTGCACCGGCAGATCCTCGAG
CGGTTCCCGCAGTTCGACATCATCGTCCGGTACGAGGCGGATGAAATCCTGCCCGCCGTGCTCGACTGCCTCCCGCA
CCGGACATTCGACGTGATTCCCGGCTTGAGCTGGCGCGCGACGGGCCGCGGCTCGCCGTTGCGCTTCACCGACGGCA
AGCCCAAGGTCGAGGATCTGGACCTGCTGCCGATCGCCTCGTACGACCATTATCCGGTTGAGGAACTGGGGCTCTCG
ATGCTGCGCATCGAGGCCGGCCGAGGGTGTCCGTTCGCATGCACATTCTGCTCGACGGCCGGTTTCTTTCAACGCAG
CTTCCGCCTCAAATCGGCCGAGCGCCTGGTTCGCGAACTGGATATCCTTCACCAGCGTTATCGCGTGTCGGACTTCA
AGCTCGACCACGACATGTTCACCGTGAATCGCCGCAAGGTAATGGAGTTTTGCGAGGCGGTCGCCGGGCGCGACTAT
CGGTGGCGCGCGTCGGCGCGGATCGACTGCGTCGACGAAGCATTGTTGAAGAAGATGGCCGACGCGGGCTGCGTGAA
TCTTTATTTCGGCGTGGAGACGGGCTCCGAGCGCATGCAAAAGCTCTGCAAGAAGCGGCTGGATCTGCAACGGGTCG
AGCCCATTCTGGCGGCGGCGGATTCATTCGGAATCGAGACGACCGCGTCGTTCATCACCGGCTATCCCGAGGAAACC
GGGGAGGATCAGGACGACACGCTGGACATGATCGGGCGCTGCGCTCGGCGAGCGTCTTGCCTCACGCAACTGCACAT
GCTCGCGCCGGAGCCCGGCACACCGCTGTTCGACGAACGCGGCGCGGAAATCGCCTACGACGGCTGTGGCGGGCCTT
ACAACACGCGCCTGCTGAGTTCGTCGGACGAGCGTGCGGTGCTCGGCCATCCCGACATATTCCAAACCTACTATCAC
TACCCCGCCGCGCTGCCGCGTGCGCGATACATCTTTGCCGTCGGGGCGGCGGATCTGCTGCGCCGGGTCGGGCCGGT
CATTTTCGCGTATGCGTTGCGGGGGTTCGGAGGGCGGCTGAGCACGCTGATATCGGCGTGGCGACAATTCGCCGACG
AAACGTCGCCCGGCGCGCCCGCGGACGCGGCGGGGCTCGAGGCCTTCATTGCCGCGCGGTTCGGCCGCGGCCATCAC
CTGCATTCTCTTTTTCGGTACGCGTTGCGGCTCTATGCGGCCCGCGCTCCGGCGGACGTAATCGTCGAGGGGCCCTA
TGAGCCGGACCGGCCTTGCATGCTGGCCGAAAACGTCGGTCTGCTGGAGGACATTCACAATTGCGACGCATTGGTCG
ACCGGATCAGGCGATCGACCGAAGGCGCGCCGATGCTCGGCGACGCGGATGTCGGCGGGCTCGGAACATATATCGTT
CAGGGGCACGGCGAGGCCGCCACGGGCTATTGGGTGGAGTCCGGCATCGCTACGCTTCTCGATCTGTTCAGGTCGCC
CCGTTCGTGCCGCGCCGTTGCGCAATGGCTGGCGGACGCCACCCGCAATGACGGCATAGACGCTTCGATCTTCGAGC
CGCTGCTGCGCAGAGGCATCCTGGTGTCGGCCGGTTAAATCCATTTCGTCCATGGTGCGTCGCACACTCGAACGTTG
GTCCGCGCTGGATCTCGCGGAAGGGCTTCAACTGGCGCATGCCGTCGCCGCGCTGCAGGCGCTCGGCGTTCTGGACG
CGATGAGGGAGCCGGTCACGGCGGAAACGTTGAGCGCCAGCCATGATCTCGATCCCGAGTTGCTGCGCGGCATCCTC
GAATATGCGGCGTCGAGAACGAATCTGGTTCGCAAGACGGACCGCGGATTCGCGATTTCCGAGCATTACACCGCGCA
AGCGCGCTTTCTGTTGAACCTGTACGCCGGCGCTTATGCCGATAACGCTTCACGGCTGGCGACGCTGCTCCGGCGTC
CGGCATTGGCGGGCTCCATGATCGACCTCGTCGCGCACGCGCGCGCCTTCGACGCGAGCGAGGCCGGCGGTGGAGCG
CTGGCGTCGATCGTCACGCAATTGCGTCTCAATCATGTGCTGGATCTGGGATGCGGCGCCGGCGCGCTGCTGCGCGA
GTTGGCGGCGGACGATCGCGGATTCGTCGGCTGGGGACTGGATCGCAATCCCGCGATGTGCAAGGCGGCGCGGGTTC
GAGCGCGGCAAGCGGGGCTCGCCGCGCGCGTGAAGGTGTTCCACGGCGACGGCTGGAATCCCGGGGCGTCGGTCCCG
CCGCGCGTGCTGGCGCGCGTGCGCAACGTCGTCGCGAGCCAGTTCGTCAACGAGATGTTTCGCGGCGGCACATCGCG
AGTGGAAACGTGGCTTCGGCGCATGCGGCGCTTGCTGCCCGGCCGCATGCTCGTGATATCGGACTATTACGGCCGCC
TCGGCCACGGCACTCACACGATGCGCCGCGAGACGCTGCTGCATGACTATGTGCAGCTCATATCGGGGCAGGGCGTG
CCGCCGCCCGATTCCGACGCGTGGTTGGCCATGTATCGAGCAGCCGGTTGCCGGCCTCTGCACGTGATCGAAGACCG
GGCGGCCACGACATGCTTCGTCCATCTGGTGGGGCTATAG

F) Insert from genomic DNA of *B. thailandensis* E264 encoding all proteins, with the gene encoding ApyS removed. Full construct name: pSCrhaB2_ApyOAIHD

ATGAACGGTCTACCTCCGTCGCTTCCGCGGGTCGACAGCACCGAGTCGCTTTTTGCGGAGCCGCTCGCGTTTCTCGC
ACAAGCGCGCTCGCGGCATGGTGATGTCTTCGTCATGCGGGAGCATGGTCCGATTTTTTCCCGCGCGAGCGATTGCA
GCGGCGTGATAGCGGTATTCGGCGAGCATCGGCTTCGACAGATATTGACCGATATAGACAATTTCGCGTTGCCGATG

TCCGCGGCGGCGAAAATGGCATTGCCGAAAAACCTCGTCAACCTCAATCGCGGCTTGCACAGCATGCGCGAGCCGGA
GCATGGCCGGCACAAGCGCCTCTTGACGGGAACGATCAATCGCGAGTTGTTCGACGCGCACCGCTTCGAAATACGAG
CGGCTTTGAACCGCTTTTGCGAAATGCTGAAAGTGGACCGCCGGATTTCAGTGGTCAGCCGGATGCGCGAGCTGACC
GTGGAGATGGCGTCTCATATCTTTCTTGGGGCGCAGTGCCAAGAGGATGACGAACTGGCGTTTCTTTTAAGCGCCTA
TTTTACTTTGCGACGCGAAGCGTCTTCGCTGAACGCGCGCGATCCCCTGCTGTACCGGGACGAACTGATCGGCGTCG
GGCAGCAACTCGACCGCACATTGCGGGAGCGGATCAGGCGGTATCGAAAGCGTCCCGTCGACGCTCGTGCGGGGCTG
CTGCAACGTCTGGCAACGGCCGGACCGCCGGGCTCGCCCGCGCTTTCCGAGGACGAGATAGTCGGCCATGCCAACGT
GATGTTCGTGTCGAGCACCGAGCCCGTCGCGATGTCGCTGACGTGGCTGCTGCTCGTTCTGTCGCAGCTGCCCGATC
TGAGGCGCGCGCTTCGCGCGGAAATTGCCGATCGCGCGTCGATGCCGGCATCGACGAACGGCGCGTCGTGGCTCGAG
AACGTCGTCAACGAGACGCTCCGGCTCCTGACGCCCAACGCGCTGATGGTTCGCGCGACGACGCGCGCGGTCTCGCT
GCAAGGCGTCGCGCTTCCGGCGCGTTGCGAGATCGTCGTGTGTCCGTTCCTCGCGCATCGCGAAGCCAAGCCGTTTC
CGGACCCGCATGCATTCTCGCCGTCCCGGTGGGAGACGGCCAGGCCGTCTCCGTACGAGTATTTTCCGTTCGGCGCC
GGCGGTCATTTTTGCGCGGGGCGAAATCTCGCGCTCTCGCTGATTCGCGAAGTACTGTCAACGCTGCTATCGCGCTT
CGATTTCGTTTTGGATGGCGAACAGTCTATCGACTGGCGCATTCATATCATGTTGATGCCGAAAGGCGACCCGGCAC
TCATCGCGCATCCCGTCGATGAGCGCGGTGACACGCCGTCGCCGAAATGGCGCGGGCCGATAACCGATCTATTCCAT
TTCGCGCCGGGGCTTTCTTGACATTTACCAGGGCGCATCCTTTAATCAGAAGCGAATCGGCATGCATGCCCATCGGG
ATTTTTTCCAGTCATTGATCCGCATGGTTCTTTTCACGGGTTATTCCGCGCTCACACGCTAAGCGGCGCGTCCTGCC
TACAACAAAGGAACAATC**ATGCATCACCATCACCATCACATGGCTACGAAACCCAAAAAAACAAAGGGCGTTTCTGT**
**TTCCATCGAAGGGAAGCTGCCCAAGATGACCCTGGATATGCCGGTGGACGCGAAAAAGATCAAGGCCATCCAGAAAT**
**GCCTGGAAAACGGCAAGCTGACCATCACGATGAGCAAGGTGGATCTTGCCGGCGGCCGCATGGGCGAGGGCTATCTC**
**TACGACTGA**TGTCCGGCGGCGCTCTGGTTTTCGCCGGCGGCGTCGCGATCTCGCCGCCTCGATGCTCGGGCCTCGGA
GGGAACGTGCCGCCGAACGCGTGCGTTTGGCGCGCGGTCGCGCGATAACGGGCGATGAATTCCGGGCTTCGGCAGTT
CGACCTTAAGGGCGCGGGAAGCCTATGAAAGCCTTCGATCGCGATGCATCGGAGCCTCGGATCGGTCTCGCGTATGG
GCCCGGGATGGCGGAATTCGCCGCACGGAACGCGCACCTCGTCGATTACATCGAAGTGCCGTTCGAGCAATTGCGAT
TCTCGCCGGCGGTCGCCGAGCTTCAGCAAACCATTCCGTTCGTTCTGCACTGCGCCAGCTTGTCCGTCGCGGGATTC
GTGCCGCCCGACGATTCGACGGTCGACGCAATCGAGCGCACGGCCGTGCAGACCGGCACGCCATGGATAGGCGAACA
TCTCGCGTACATTTCGGCCGATCCCGTTGGCGAAGCGCTGGGCGGGACCGGTGAGCCCACCTCGTTGTCGTACACGT
TGTGCCCTCAGTTGAGCGACGAGACGGTACGCCGCGTCGTCGACAATCTCGCCGCGCTTCGTCCGCATTTTCCGGTT
CCACTGATCGTCGAAAACTCCCCGCAGTACTTTCCGATACCGGGCAGCACGATGGGAATGACGGACTTCATTCGCGC
GATCACGGACCGATGCGACGTCGGTTTGCTCCTGGACCTGAGTCACTTCCTGATCACGGCGCACAACACGGGCGCAG
AGGTGCATCGGGAACTCGCCCGGCTTCCGCTGGAGCGCGTGGTCGAAGTTCATCTGTCCGGAATGAGCGTGCAGTCC
GGCACGGCCTGGGACGACCATTCGCTGCCGGCATCGCCGATCCTGTTCGAACTGCTCGAGCGCCTGCTCGACGTGGC
GCGGCCTCGCGCGCTGACCTTCGAATACAACTGGTCGCCATACTTCCCGCTGTCGGTCCTGACGACGCATATCGAGC
GCGCACGCCAGTTGATGGGGCTCGCATGAACGCAGGGAAAATACACGCCGTGATCGCCGCCGGCGCCAAGGACCCGC
GGCTCATCGAGCATTGGCGGCGGCGTCCGGACCTGCTGCGCCGCGCAGGAATCGAGCCGGATCATCTCGATTTGCGC
GTATTGGAGAATTTCGCCGGGTTGAGCCTGAAGGTGCGTCACAACGGCCTGCGCGGGGATCTGCCGCTGAGTTTCAG
GTTGTTGAGCGTGGCCGGCCTCGAGATCGAGGTCTTCGCCGCGTATGCGTCGGACTGCGCGGAAAGCGGCCATGTAT
TCGCCGCCACGCCGCGGGAACGCGCGCGCGATCTGATGGCGTTCCTCGACAGATGGCTGGACCCCGCGCAACGAAAT
CACGTTCTGCTGTGGGACATCCTTCGTCATGAATATGCGCTTTCGTGCCTGCGGCAAGAAAGCGCCGCGCCGCTACC
CGCCGCTCGCCGCATCATCCGCCATCGGGATCGCGCGACAGGCGGCTGCATTCCGATGCTCAACGGCTCCGTCGCTC
AGCACGTGATGCATTGCAATCCGGAAGACCTGGCGTCGGCGCTGCGTCAGAAGGAGCCGCCGCTCGATGCGGTTCCC
GCCGCCGACACTTACTGCTGCTATTGGCGCTCCGCCGAAACGGGCGAAATCCGGTTTCTGCAGGCCGATACGCTCGG
CTATTTCGCGATCGAAGCGATCGACGGCCGGCGATCCGCCGCCGACATCTGCGATCGGCTGCTCGGCGGCCGCCGTC
TGCTGCCGCAGTTCCTCCGGTCGCTCGACCAACTGGCCGGAATCGGGATCGTGCGGTTCGAGAGGCCGCTCGGAAGC
GCTGCGGCATGAAACTGCTGCTCGTCGACAACCTCGTCATGCCGGAAGAGGGCTCGCTCGCGTACCTCGACGTGCAT
CCGCATCTCGGGCTGCTCGCGCTGGCCGCGGTCGCCGAAGCCGACGGCCATCGCGTGCAGATCTACGATCCGAAACG
ACTGATCAGGGACGGCACGCTGGCTTACGATGCGACGCTTTACGAGCGGGCGAGCCAAGCGATTCTCGCGGCGCGTC
CGGACGGGGTCGGCTTCACGGCGCTCGGCTGCAGCTTTCTGTTTGCGCTCAATGTCGCGGCGCTGTTGAAGCGGCGC
GAGCCCGACCTGCCCGTCTTGCTCGGCGGGCCGCACGCGACGATGCTGCACCGGCAGATCCTCGAGCGGTTCCCGCA
GTTCGACATCATCGTCCGGTACGAGGCGGATGAAATCCTGCCCGCCGTGCTCGACTGCCTCCCGCACCGGACATTCG
ACGTGATTCCCGGCTTGAGCTGGCGCGCGACGGGCCGCGGCTCGCCGTTGCGCTTCACCGACGGCAAGCCCAAGGTC
GAGGATCTGGACCTGCTGCCGATCGCCTCGTACGACCATTATCCGGTTGAGGAACTGGGGCTCTCGATGCTGCGCAT
CGAGGCCGGCCGAGGGTGTCCGTTCGCATGCACATTCTGCTCGACGGCCGGTTTCTTTCAACGCAGCTTCCGCCTCA
AATCGGCCGAGCGCCTGGTTCGCGAACTGGATATCCTTCACCAGCGTTATCGCGTGTCGGACTTCAAGCTCGACCAC
GACATGTTCACCGTGAATCGCCGCAAGGTAATGGAGTTTTGCGAGGCGGTCGCCGGGCGCGACTATCGGTGGCGCGC
GTCGGCGCGGATCGACTGCGTCGACGAAGCATTGTTGAAGAAGATGGCCGACGCGGGCTGCGTGAATCTTTATTTCG
GCGTGGAGACGGGCTCCGAGCGCATGCAAAAGCTCTGCAAGAAGCGGCTGGATCTGCAACGGGTCGAGCCCATTCTG

GCGGCGGCGGATTCATTCGGAATCGAGACGACCGCGTCGTTCATCACCGGCTATCCCGAGGAAACCGGGGAGGATCA
GGACGACACGCTGGACATGATCGGGCGCTGCGCTCGGCGAGCGTCTTGCCTCACGCAACTGCACATGCTCGCGCCGG
AGCCCGGCACACCGCTGTTCGACGAACGCGGCGCGGAAATCGCCTACGACGGCTGTGGCGGGCCTTACAACACGCGC
CTGCTGAGTTCGTCGGACGAGCGTGCGGTGCTCGGCCATCCCGACATATTCCAAACCTACTATCACTACCCCGCCGC
GCTGCCGCGTGCGCGATACATCTTTGCCGTCGGGGCGGCGGATCTGCTGCGCCGGGTCGGGCCGGTCATTTTCGCGT
ATGCGTTGCGGGGGTTCGGAGGGCGGCTGAGCACGCTGATATCGGCGTGGCGACAATTCGCCGACGAAACGTCGCCC
GGCGCGCCCGCGGACGCGGCGGGGCTCGAGGCCTTCATTGCCGCGCGGTTCGGCCGCGGCCATCACCTGCATTCTCT
TTTTCGGTACGCGTTGCGGCTCTATGCGGCCCGCGCTCCGGCGGACGTAATCGTCGAGGGGCCCTATGAGCCGGACC
GGCCTTGCATGCTGGCCGAAAACGTCGGTCTGCTGGAGGACATTCACAATTGCGACGCATTGGTCGACCGGATCAGG
CGATCGACCGAAGGCGCGCCGATGCTCGGCGACGCGGATGTCGGCGGGCTCGGAACATATATCGTTCAGGGGCACGG
CGAGGCCGCCACGGGCTATTGGGTGGAGTCCGGCATCGCTACGCTTCTCGATCTGTTCAGGTCGCCCCGTTCGTGCC
GCGCCGTTGCGCAATGGCTGGCGGACGCCACCCGCAATGACGGCATAGACGCTTCGATCTTCGAGCCGCTGCTGCGC
AGAGGCATCCTGGTGTCGGCCGGTTAAATCCATTTCGTCCATGGTGCGTCGCACACTCGAACGTTGGTCCGCGCTGG
ATCTCGCGGAAGGGCTTCAACTGGCGCATGCCGTCGCCGCGCTGCAGGCGCTCGGCGTTCTGGACTAG

**Table S3:** Sequence of the reconstructed pSCrhaB2-based vectors used for more efficient ApyD-catalyzed methylation. Construct name: pSCrhaB2_ApyD_RBS_ApyOAHIDS. Each gene is highlighted as follows. *apyA*: bolded black; *apyO*: pink; *apyD*: green; *apyH*: orange; *apyI*: underlined; *apyS*: blue. The N-terminus of ApyA contains a hexa-His tag. In this construct, ApyD is placed right after the ribosome binding site downstream of the rhamnose-inducible promoter in the pSCrhaB2 backbone.

ATGAAACTGCTGCTCGTCGACAACCTCGTCATGCCGGAAGAGGGCTCGCTCGCGTACCTCGACGTGCATCCGCATCT
CGGGCTGCTCGCGCTGGCCGCGGTCGCCGAAGCCGACGGCCATCGCGTGCAGATCTACGATCCGAAACGACTGATCA
GGGACGGCACGCTGGCTTACGATGCGACGCTTTACGAGCGGGCGAGCCAAGCGATTCTCGCGGCGCGTCCGGACGGG
GTCGGCTTCACGGCGCTCGGCTGCAGCTTTCTGTTTGCGCTCAATGTCGCGGCGCTGTTGAAGCGGCGCGAGCCCGA
CCTGCCCGTCTTGCTCGGCGGGCCGCACGCGACGATGCTGCACCGGCAGATCCTCGAGCGGTTCCCGCAGTTCGACA
TCATCGTCCGGTACGAGGCGGATGAAATCCTGCCCGCCGTGCTCGACTGCCTCCCGCACCGGACATTCGACGTGATT
CCCGGCTTGAGCTGGCGCGCGACGGGCCGCGGCTCGCCGTTGCGCTTCACCGACGGCAAGCCCAAGGTCGAGGATCT
GGACCTGCTGCCGATCGCCTCGTACGACCATTATCCGGTTGAGGAACTGGGGCTCTCGATGCTGCGCATCGAGGCCG
GCCGAGGGTGTCCGTTCGCATGCACATTCTGCTCGACGGCCGGTTTCTTTCAACGCAGCTTCCGCCTCAAATCGGCC
GAGCGCCTGGTTCGCGAACTGGATATCCTTCACCAGCGTTATCGCGTGTCGGACTTCAAGCTCGACCACGACATGTT
CACCGTGAATCGCCGCAAGGTAATGGAGTTTTGCGAGGCGGTCGCCGGGCGCGACTATCGGTGGCGCGCGTCGGCGC
GGATCGACTGCGTCGACGAAGCATTGTTGAAGAAGATGGCCGACGCGGGCTGCGTGAATCTTTATTTCGGCGTGGAG
ACGGGCTCCGAGCGCATGCAAAAGCTCTGCAAGAAGCGGCTGGATCTGCAACGGGTCGAGCCCATTCTGGCGGCGGC
GGATTCATTCGGAATCGAGACGACCGCGTCGTTCATCACCGGCTATCCCGAGGAAACCGGGGAGGATCAGGACGACA
CGCTGGACATGATCGGGCGCTGCGCTCGGCGAGCGTCTTGCCTCACGCAACTGCACATGCTCGCGCCGGAGCCCGGC
ACACCGCTGTTCGACGAACGCGGCGCGGAAATCGCCTACGACGGCTGTGGCGGGCCTTACAACACGCGCCTGCTGAG
TTCGTCGGACGAGCGTGCGGTGCTCGGCCATCCCGACATATTCCAAACCTACTATCACTACCCCGCCGCGCTGCCGC
GTGCGCGATACATCTTTGCCGTCGGGGCGGCGGATCTGCTGCGCCGGGTCGGGCCGGTCATTTTCGCGTATGCGTTG
CGGGGGTTCGGAGGGCGGCTGAGCACGCTGATATCGGCGTGGCGACAATTCGCCGACGAAACGTCGCCCGGCGCGCC
CGCGGACGCGGCGGGGCTCGAGGCCTTCATTGCCGCGCGGTTCGGCCGCGGCCATCACCTGCATTCTCTTTTTCGGT
ACGCGTTGCGGCTCTATGCGGCCCGCGCTCCGGCGGACGTAATCGTCGAGGGGCCCTATGAGCCGGACCGGCCTTGC
ATGCTGGCCGAAAACGTCGGTCTGCTGGAGGACATTCACAATTGCGACGCATTGGTCGACCGGATCAGGCGATCGAC
CGAAGGCGCGCCGATGCTCGGCGACGCGGATGTCGGCGGGCTCGGAACATATATCGTTCAGGGGCACGGCGAGGCCG
CCACGGGCTATTGGGTGGAGTCCGGCATCGCTACGCTTCTCGATCTGTTCAGGTCGCCCCGTTCGTGCCGCGCCGTT
GCGCAATGGCTGGCGGACGCCACCCGCAATGACGGCATAGACGCTTCGATCTTCGAGCCGCTGCTGCGCAGAGGCAT
CCTGGTGTCGGCCGGTTAAATAGCAGGAGGAATTCACCATGAACGGTCTACCTCCGTCGCTTCCGCGGGTCGACAGC
ACCGAGTCGCTTTTTGCGGAGCCGCTCGCGTTTCTCGCACAAGCGCGCTCGCGGCATGGTGATGTCTTCGTCATGCG
GGAGCATGGTCCGATTTTTTCCCGCGCGAGCGATTGCAGCGGCGTGATAGCGGTATTCGGCGAGCATCGGCTTCGAC
AGATATTGACCGATATAGACAATTTCGCGTTGCCGATGTCCGCGGCGGCGAAAATGGCATTGCCGAAAAACCTCGTC
AACCTCAATCGCGGCTTGCACAGCATGCGCGAGCCGGAGCATGGCCGGCACAAGCGCCTCTTGACGGGAACGATCAA
TCGCGAGTTGTTCGACGCGCACCGCTTCGAAATACGAGCGGCTTTGAACCGCTTTTGCGAAATGCTGAAAGTGGACC
GCCGGATTTCAGTGGTCAGCCGGATGCGCGAGCTGACCGTGGAGATGGCGTCTCATATCTTTCTTGGGGCGCAGTGC
CAAGAGGATGACGAACTGGCGTTTCTTTTAAGCGCCTATTTTACTTTGCGACGCGAAGCGTCTTCGCTGAACGCGCG
CGATCCCCTGCTGTACCGGGACGAACTGATCGGCGTCGGGCAGCAACTCGACCGCACATTGCGGGAGCGGATCAGGC
GGTATCGAAAGCGTCCCGTCGACGCTCGTGCGGGGCTGCTGCAACGTCTGGCAACGGCCGGACCGCCGGGCTCGCCC
GCGCTTTCCGAGGACGAGATAGTCGGCCATGCCAACGTGATGTTCGTGTCGAGCACCGAGCCCGTCGCGATGTCGCT
GACGTGGCTGCTGCTCGTTCTGTCGCAGCTGCCCGATCTGAGGCGCGCGCTTCGCGCGGAAATTGCCGATCGCGCGT
CGATGCCGGCATCGACGAACGGCGCGTCGTGGCTCGAGAACGTCGTCAACGAGACGCTCCGGCTCCTGACGCCCAAC
GCGCTGATGGTTCGCGCGACGACGCGCGCGGTCTCGCTGCAAGGCGTCGCGCTTCCGGCGCGTTGCGAGATCGTCGT
GTGTCCGTTCCTCGCGCATCGCGAAGCCAAGCCGTTTCCGGACCCGCATGCATTCTCGCCGTCCCGGTGGGAGACGG
CCAGGCCGTCTCCGTACGAGTATTTTCCGTTCGGCGCCGGCGGTCATTTTTGCGCGGGGCGAAATCTCGCGCTCTCG
CTGATTCGCGAAGTACTGTCAACGCTGCTATCGCGCTTCGATTTCGTTTTGGATGGCGAACAGTCTATCGACTGGCG
CATTCATATCATGTTGATGCCGAAAGGCGACCCGGCACTCATCGCGCATCCCGTCGATGAGCGCGGTGACACGCCGT
CGCCGAAATGGCGCGGGCCGATAACCGATCTATTCCATTTCGCGCCGGGGCTTTCTTGACATTTACCAGGGCGCATC
CTTTAATCAGAAGCGAATCGGCATGCATGCCCATCGGGATTTTTTCCAGTCATTGATCCGCATGGTTCTTTTCACGG
GTTATTCCGCGCTCACACGCTAAGCGGCGCGTCCTGCCTACAACAAAGGAACAATC**ATGCATCACCATCACCATCAC**
**ATGGCTACGAAACCCAAAAAAACAAAGGGCGTTTCTGTTTCCATCGAAGGGAAGCTGCCCAAGATGACCCTGGATAT**
**GCCGGTGGACGCGAAAAAGATCAAGGCCATCCAGAAATGCCTGGAAAACGGCAAGCTGACCATCACGATGAGCAAGG**
**TGGATCTTGCCGGCGGCCGCATGGGCGAGGGCTATCTCTACGACTGA**TGTCCGGCGGCGCTCTGGTTTTCGCCGGCG

GCGTCGCGATCTCGCCGCCTCGATGCTCGGGCCTCGGAGGGAACGTGCCGCCGAACGCGTGCGTTTGGCGCGCGGTC
GCGCGATAACGGGCGATGAATTCCGGGCTTCGGCAGTTCGACCTTAAGGGCGCGGGAAGCCTATGAAAGCCTTCGAT
CGCGATGCATCGGAGCCTCGGATCGGTCTCGCGTATGGGCCCGGGATGGCGGAATTCGCCGCACGGAACGCGCACCT
CGTCGATTACATCGAAGTGCCGTTCGAGCAATTGCGATTCTCGCCGGCGGTCGCCGAGCTTCAGCAAACCATTCCGT
TCGTTCTGCACTGCGCCAGCTTGTCCGTCGCGGGATTCGTGCCGCCCGACGATTCGACGGTCGACGCAATCGAGCGC
ACGGCCGTGCAGACCGGCACGCCATGGATAGGCGAACATCTCGCGTACATTTCGGCCGATCCCGTTGGCGAAGCGCT
GGGCGGGACCGGTGAGCCCACCTCGTTGTCGTACACGTTGTGCCCTCAGTTGAGCGACGAGACGGTACGCCGCGTCG
TCGACAATCTCGCCGCGCTTCGTCCGCATTTTCCGGTTCCACTGATCGTCGAAAACTCCCCGCAGTACTTTCCGATA
CCGGGCAGCACGATGGGAATGACGGACTTCATTCGCGCGATCACGGACCGATGCGACGTCGGTTTGCTCCTGGACCT
GAGTCACTTCCTGATCACGGCGCACAACACGGGCGCAGAGGTGCATCGGGAACTCGCCCGGCTTCCGCTGGAGCGCG
TGGTCGAAGTTCATCTGTCCGGAATGAGCGTGCAGTCCGGCACGGCCTGGGACGACCATTCGCTGCCGGCATCGCCG
ATCCTGTTCGAACTGCTCGAGCGCCTGCTCGACGTGGCGCGGCCTCGCGCGCTGACCTTCGAATACAACTGGTCGCC
ATACTTCCCGCTGTCGGTCCTGACGACGCATATCGAGCGCGCACGCCAGTTGATGGGGCTCGCATGAACGCAGGGAA
AATACACGCCGTGATCGCCGCCGGCGCCAAGGACCCGCGGCTCATCGAGCATTGGCGGCGGCGTCCGGACCTGCTGC
GCCGCGCAGGAATCGAGCCGGATCATCTCGATTTGCGCGTATTGGAGAATTTCGCCGGGTTGAGCCTGAAGGTGCGT
CACAACGGCCTGCGCGGGGATCTGCCGCTGAGTTTCAGGTTGTTGAGCGTGGCCGGCCTCGAGATCGAGGTCTTCGC
CGCGTATGCGTCGGACTGCGCGGAAAGCGGCCATGTATTCGCCGCCACGCCGCGGGAACGCGCGCGCGATCTGATGG
CGTTCCTCGACAGATGGCTGGACCCCGCGCAACGAAATCACGTTCTGCTGTGGGACATCCTTCGTCATGAATATGCG
CTTTCGTGCCTGCGGCAAGAAAGCGCCGCGCCGCTACCCGCCGCTCGCCGCATCATCCGCCATCGGGATCGCGCGAC
AGGCGGCTGCATTCCGATGCTCAACGGCTCCGTCGCTCAGCACGTGATGCATTGCAATCCGGAAGACCTGGCGTCGG
CGCTGCGTCAGAAGGAGCCGCCGCTCGATGCGGTTCCCGCCGCCGACACTTACTGCTGCTATTGGCGCTCCGCCGAA
ACGGGCGAAATCCGGTTTCTGCAGGCCGATACGCTCGGCTATTTCGCGATCGAAGCGATCGACGGCCGGCGATCCGC
CGCCGACATCTGCGATCGGCTGCTCGGCGGCCGCCGTCTGCTGCCGCAGTTCCTCCGGTCGCTCGACCAACTGGCCG
GAATCGGGATCGTGCGGTTCGAGAGGCCGCTCGGAAGCGCTGCGGCATGAAACTGCTGCTCGTCGACAACCTCGTCA
TGCCGGAAGAGGGCTCGCTCGCGTACCTCGACGTGCATGCCGTTGCGCAATGGCTGGCGGACGCCACCCGCAATGAC
GGCATAGACGCTTCGATCTTCGAGCCGCTGCTGCGCAGAGGCATCCTGGTGTCGGCCGGTTAAATCCATTTCGTCCA
TGGTGCGTCGCACACTCGAACGTTGGTCCGCGCTGGATCTCGCGGAAGGGCTTCAACTGGCGCATGCCGTCGCCGCG
CTGCAGGCGCTCGGCGTTCTGGACGCGATGAGGGAGCCGGTCACGGCGGAAACGTTGAGCGCCAGCCATGATCTCGA
TCCCGAGTTGCTGCGCGGCATCCTCGAATATGCGGCGTCGAGAACGAATCTGGTTCGCAAGACGGACCGCGGATTCG
CGATTTCCGAGCATTACACCGCGCAAGCGCGCTTTCTGTTGAACCTGTACGCCGGCGCTTATGCCGATAACGCTTCA
CGGCTGGCGACGCTGCTCCGGCGTCCGGCATTGGCGGGCTCCATGATCGACCTCGTCGCGCACGCGCGCGCCTTCGA
CGCGAGCGAGGCCGGCGGTGGAGCGCTGGCGTCGATCGTCACGCAATTGCGTCTCAATCATGTGCTGGATCTGGGAT
GCGGCGCCGGCGCGCTGCTGCGCGAGTTGGCGGCGGACGATCGCGGATTCGTCGGCTGGGGACTGGATCGCAATCCC
GCGATGTGCAAGGCGGCGCGGGTTCGAGCGCGGCAAGCGGGGCTCGCCGCGCGCGTGAAGGTGTTCCACGGCGACGG
CTGGAATCCCGGGGCGTCGGTCCCGCCGCGCGTGCTGGCGCGCGTGCGCAACGTCGTCGCGAGCCAGTTCGTCAACG
AGATGTTTCGCGGCGGCACATCGCGAGTGGAAACGTGGCTTCGGCGCATGCGGCGCTTGCTGCCCGGCCGCATGCTC
GTGATATCGGACTATTACGGCCGCCTCGGCCACGGCACTCACACGATGCGCCGCGAGACGCTGCTGCATGACTATGT
GCAGCTCATATCGGGGCAGGGCGTGCCGCCGCCCGATTCCGACGCGTGGTTGGCCATGTATCGAGCAGCCGGTTGCC
GGCCTCTGCACGTGATCGAAGACCGGGCGGCCACGACATGCTTCGTCCATCTGGTGGGGCTATAG

**Figure S1:** Deconvoluted HR-ESI-MS of ApyA co-expressed with modifying enzymes in *E. coli*, cleaved with GluC or LysC. The sequence of the constructs can be found in Table S1. The sequence of the peptide is shown below with the cut site bolded. *: sodium adduct, **: potassium adduct, ***: serine phosphorylation of the GSS sequence on the N-terminus.[17] ****: gluconylation on the N-terminus.[18] A table with observed masses and the error compared to calculated values can be found in the Supplementary Dataset 3 (Excel).

`GSSHHHHHHMATKPKKTKGVSVSIEGKLPKMTLDMPVDAKKIKAIQKCLENGKLTITMS`**`K`**`VDLA`
`GGRMG`**`E`**`GYLYD`

A) ApyA (no modifying enzymes). Calculated exact mass for ApyA: 8421.3619 Da. Calculated exact mass for GluC-cleaved ApyA: 629.2696 Da

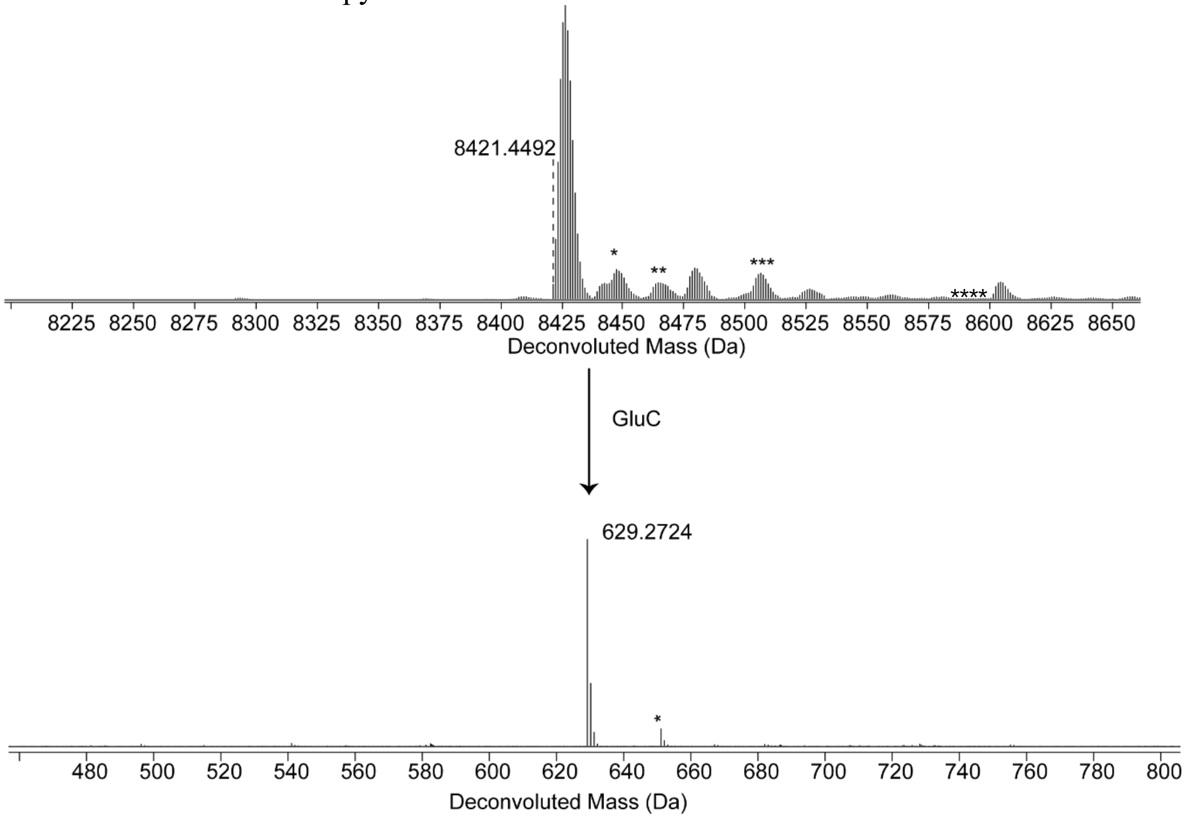

B) ApyA co-expressed with ApyD. Calculated exact mass for ApyA: 8421.3619 Da. Calculated exact mass for GluC-cleaved ApyA: 629.2696 Da. Calculated exact mass for ApyD-modified ApyA: 8435.3775 Da. Calculated exact mass for GluC-cleaved ApyD-modified ApyA: 643.2852 Da.

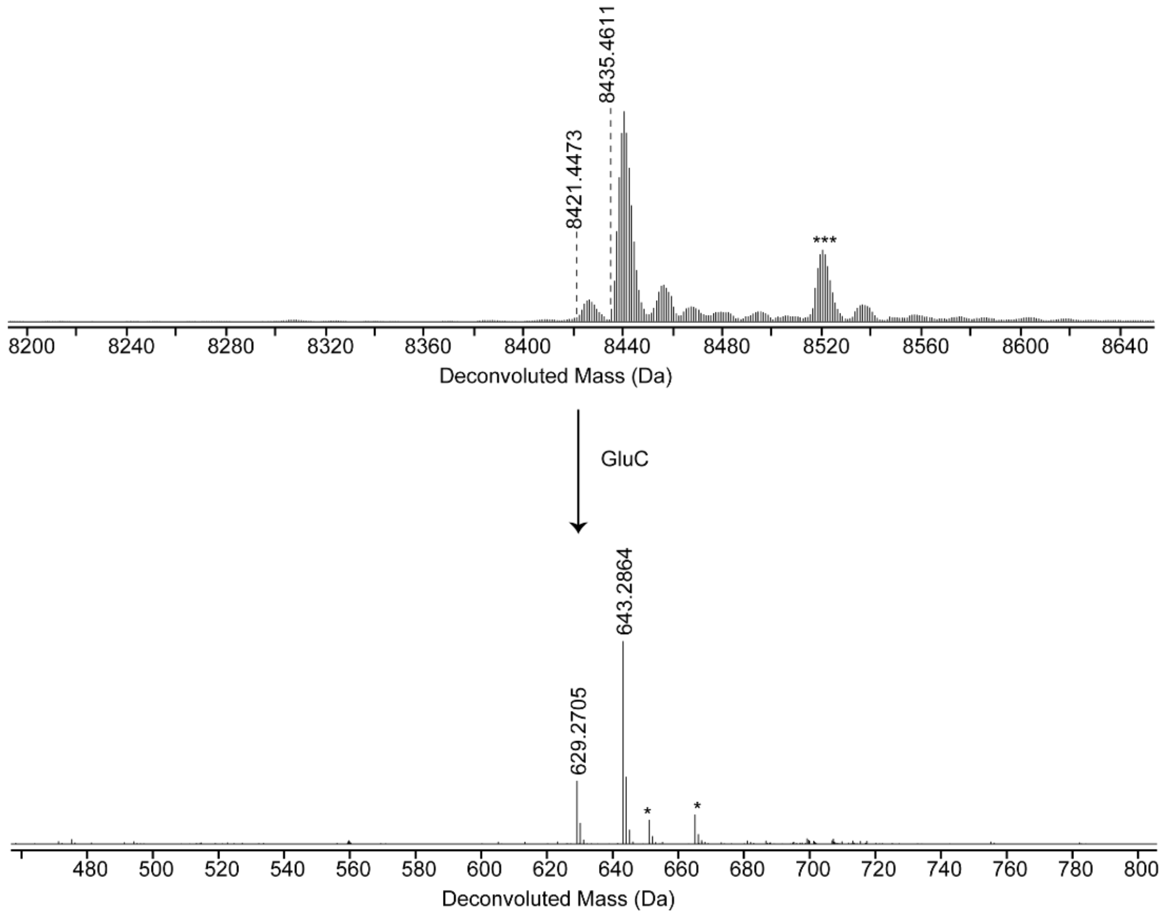

C) ApyA co-expressed with ApyO. Calculated exact mass for ApyA: 8421.3619 Da. Calculated exact mass for GluC-cleaved ApyA: 629.2696 Da. Calculated exact mass for ApyO-modified ApyA: 8419.3463 Da. Calculated exact mass for GluC-cleaved ApyO-modified ApyA: 627.2540 Da.

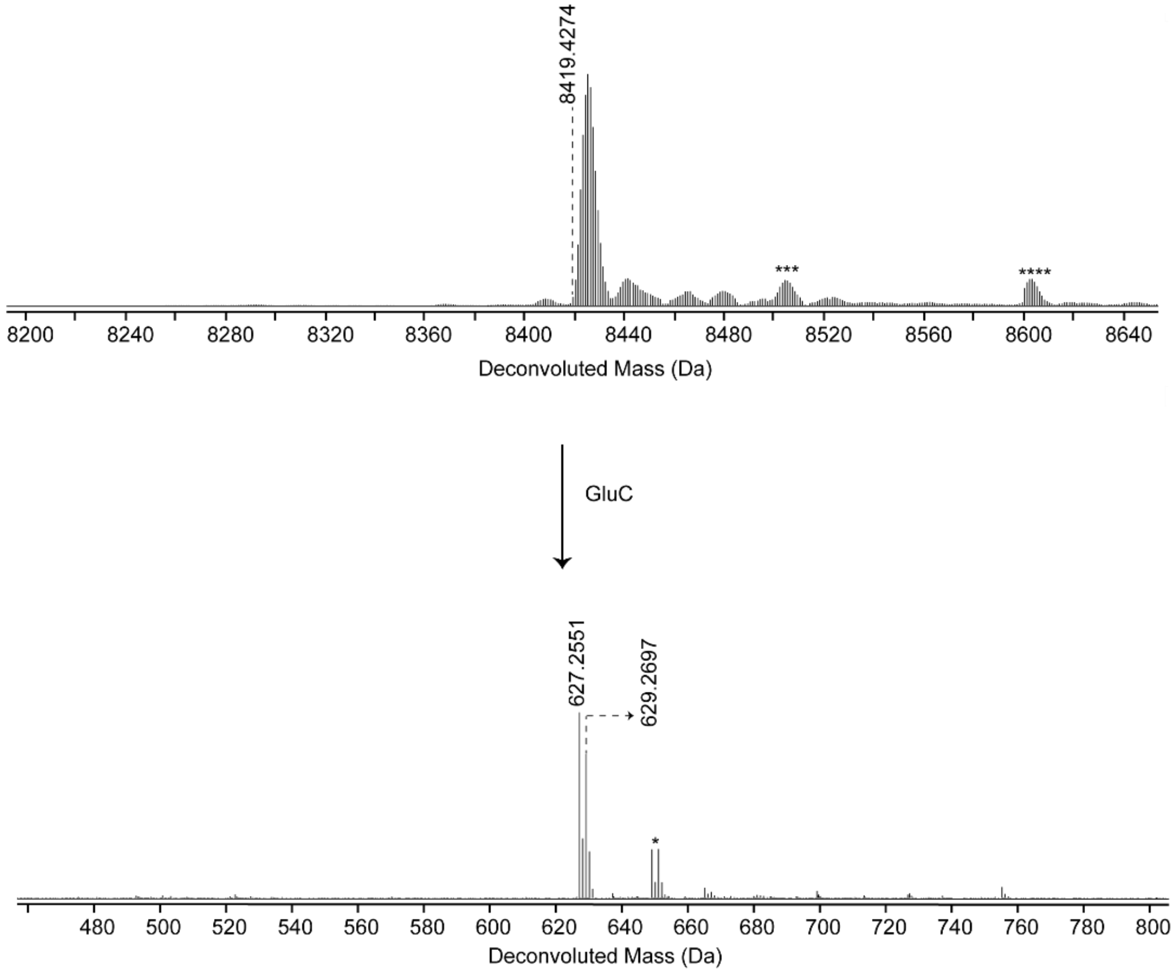

D) ApyA co-expressed with ApyD and ApyO. Calculated exact mass for GluC-cleaved ApyA: 629.2696 Da. Calculated exact mass for GluC-cleaved ApyO-modified ApyA: 627.2540 Da. Calculated exact mass for GluC-cleaved ApyD-modified ApyA: 643.2852 Da. Calculated exact mass for GluC-cleaved ApyD/O-modified ApyA: 641.2696 Da.

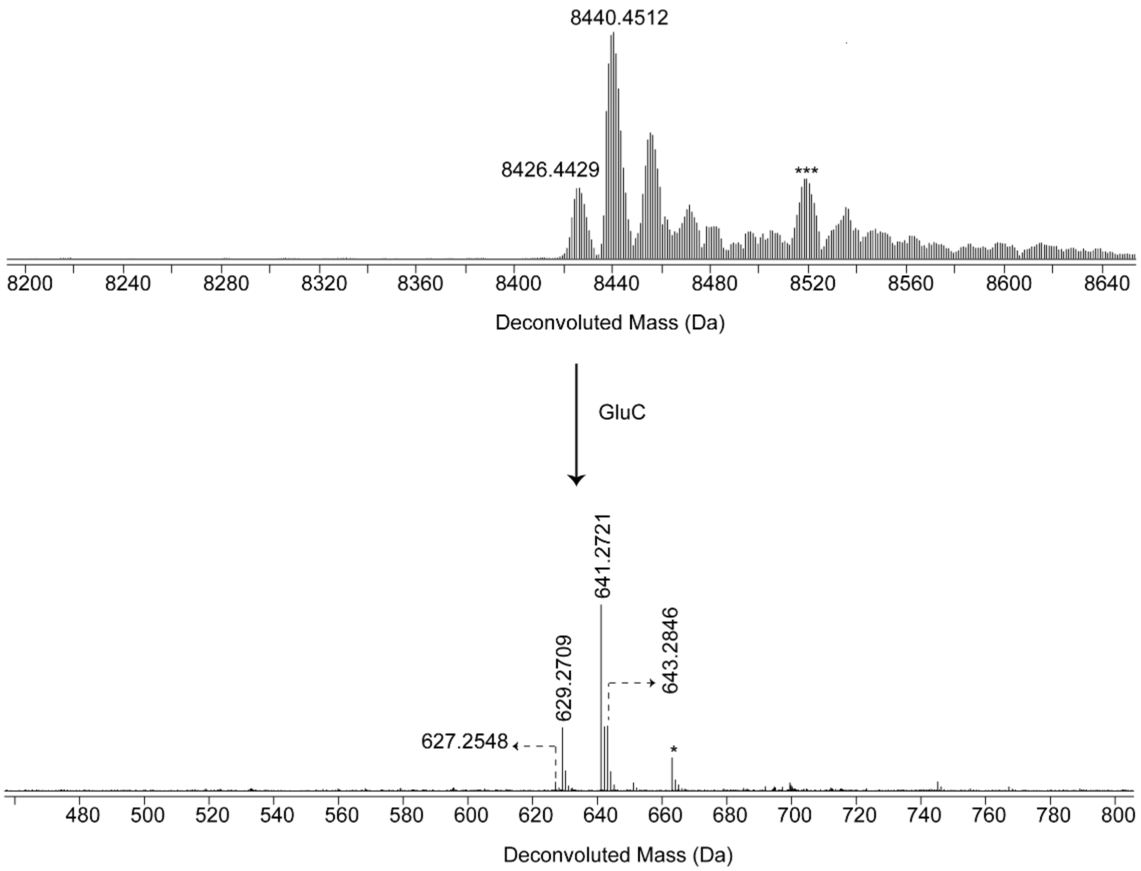

E) ApyA co-expressed with ApyD, ApyO, ApyH, ApyI, and ApyS. ApyHI and ApyS were not functional. Calculated exact mass for LysC-cleaved ApyA: 1614.7347 Da. Calculated exact mass for LysC-cleaved ApyO-modified ApyA: 1612.7191 Da. Calculated exact mass for LysC-cleaved ApyD-modified ApyA: 1628.7503 Da. Calculated exact mass for LysC-cleaved ApyD/O/H/I/S-modified ApyA: 1626.7347 Da

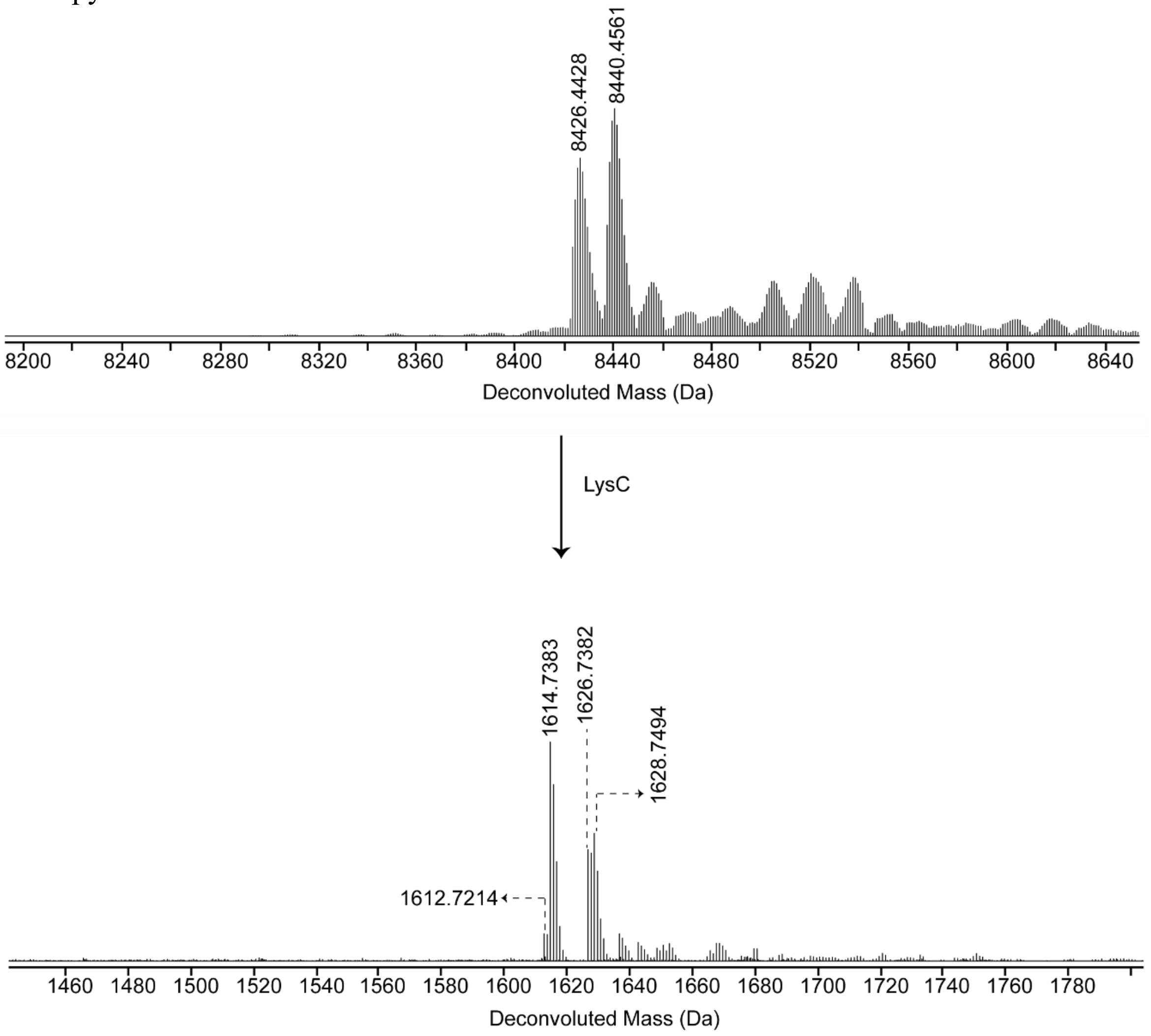

F) ApyA co-expressed with ApyH and ApyI. ApyHI were non-functional. Calculated exact mass for LysC-cleaved ApyA: 1614.7347 Da.

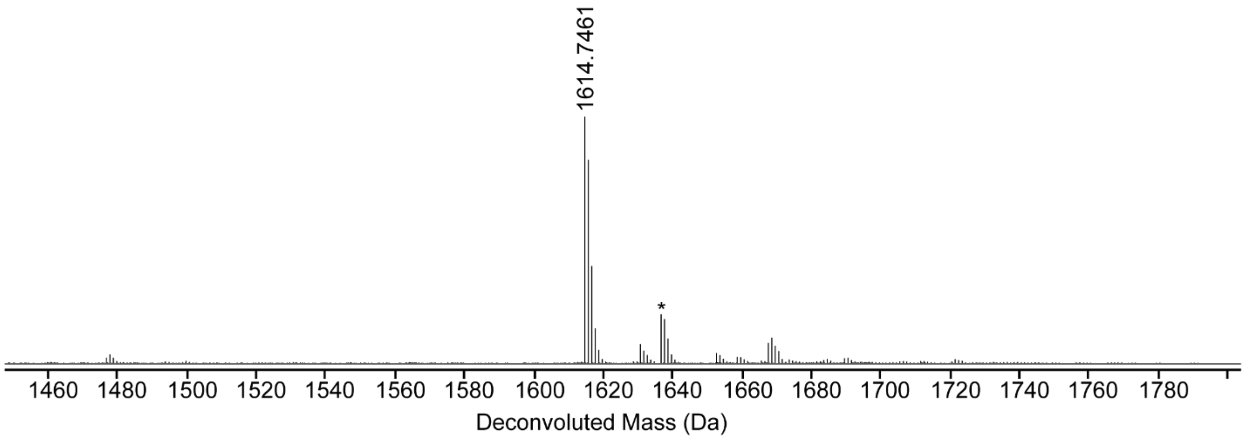

**Figure S2:** HR-ESI-MS/MS spectra of ApyA co-expressed with modifying enzymes in *E. coli* followed by GluC cleavage. The sequence of the constructs can be found in Table S1. Assigned single fragmentations are colored. Structures and the corresponding observed mass resulting from multiple fragmentations following the rules of a, b, c, x, y, and z ions typically observed with CID are shown.[19-21] Note that there could be multiple isomers associated with certain mass values. In such cases, only one isomer is shown. A table with observed masses and the error compared to calculated values can be found in the Supplementary Dataset 3 (Excel).

A) ApyA co-expressed with ApyO.

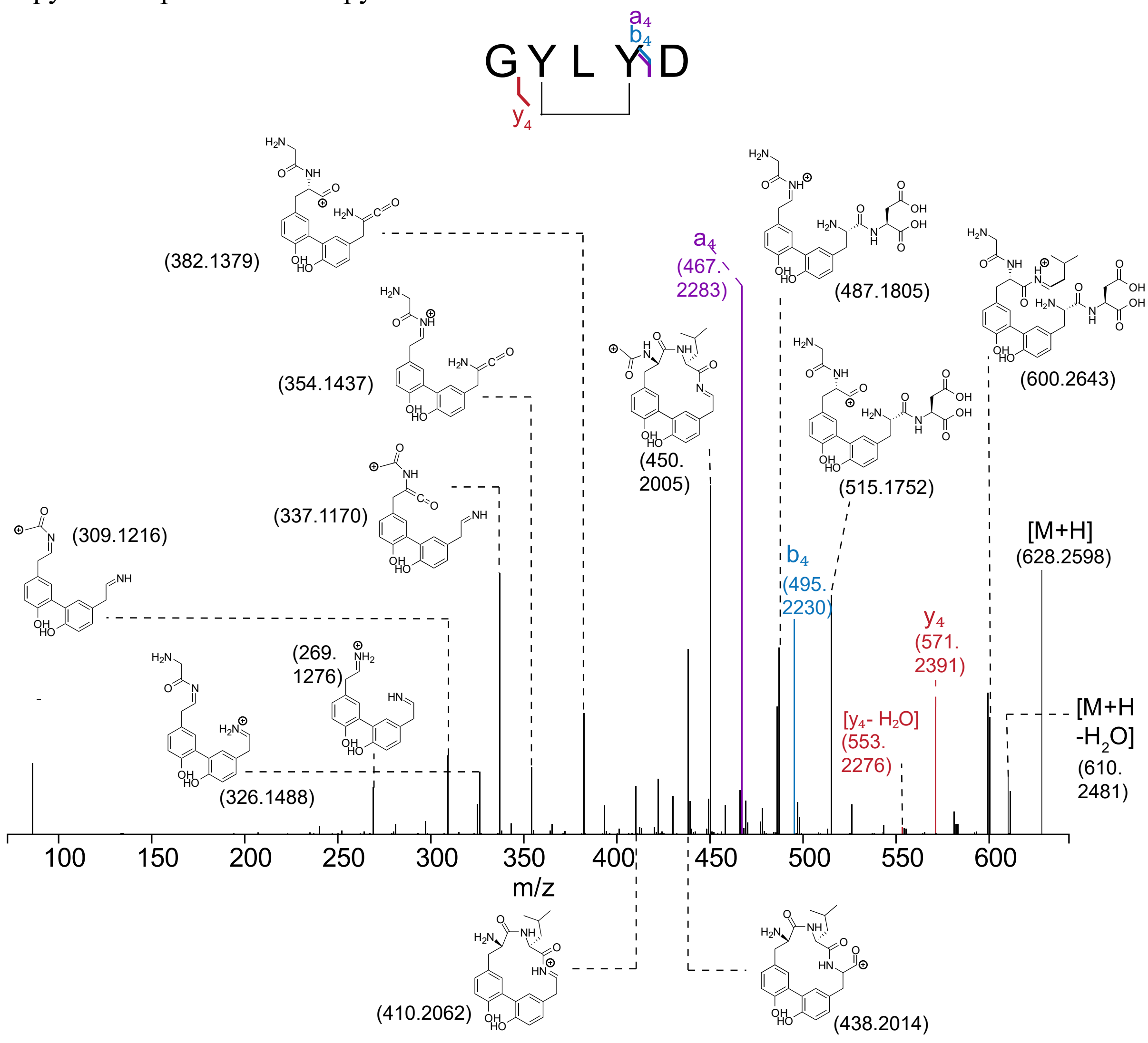

B) ApyA co-expressed with ApyD and ApyO

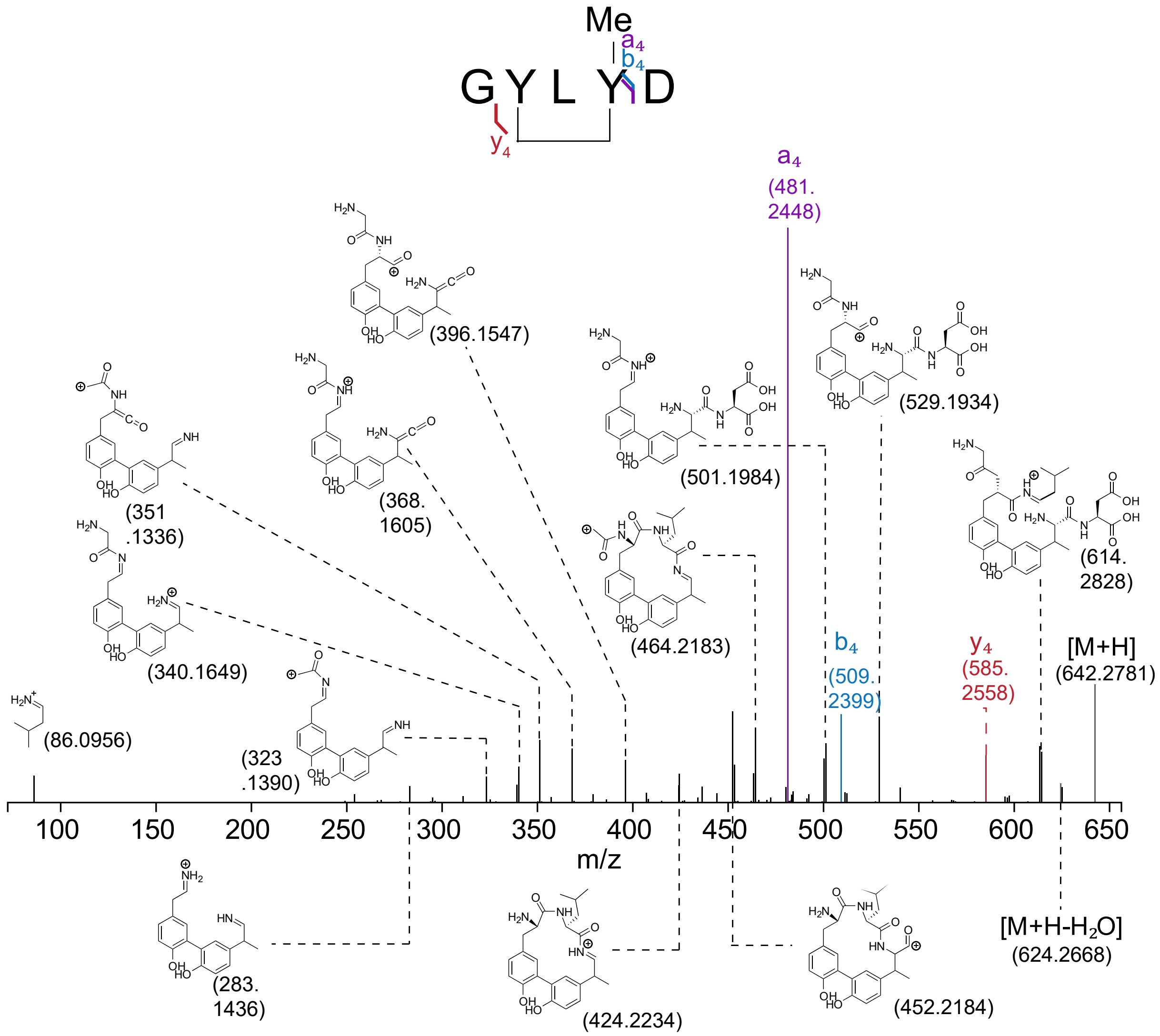

**Figure S3:** Deconvoluted HR-ESI-MS of ApyA co-expressed with all modifying enzymes (pSCrhaB2_ApyOAHIDS) in *Burkholderia* sp. FERM BP-3421. The construct sequence can be found in Table S2B. The two constructs (with or without the first 833 bp) yielded similar results (panels A and B). ApyO, ApyHI, and ApyS were active. In this expression condition, ApyD has low activity (see Figure S11 and supplementary methods). The sequence of the peptide is shown below with the cut site bolded (endoproteinase GluC was used). *: sodium adduct, #: potential methionine oxidation; absent in fragment after proteolysis. Calculated exact mass for ApyO/H/I/S-modified ApyA: 8088.2156 Da. Calculated exact mass for GluC-cleaved ApyO/H/I/S-modified ApyA: 611.2591 Da. Calculated exact mass for ApyD/O/H/I/S-modified ApyA: 625.2747 Da. A table with observed masses and the error compared to calculated values can be found in the Supplementary Dataset 3 (Excel).

`MHHHHHHMATKPKKTKGVSVSIEGKLPKMTLDMPVDAKKIKAIQKCLENGKLTITMSKVDLAGG`
`RMG`**`E`**`GYLYD`

A) Product resulting from the construct in Table S2B

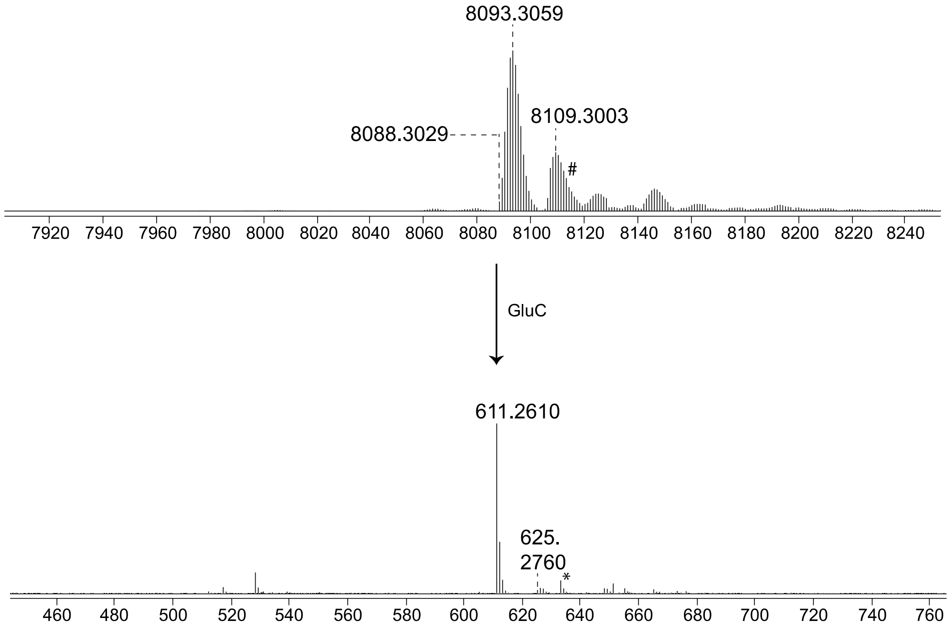

B) Construct in Table S2B but with the first 833 bp of the insert removed.

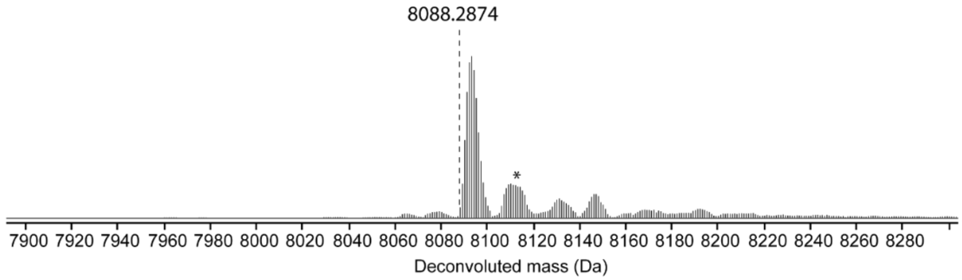

**Figure S4:** HR-ESI-MS/MS of ApyA co-expressed with all modifying enzymes (pSCrhaB2_ApyOAHIDS) in *Burkholderia* sp. FERM BP-3421. The sequence of the construct can be found in Table S2B. Endoproteinase GluC was used for proteolysis of modified ApyA. ApyS, ApyHI and ApyO were active. In this expression condition, ApyD has low activity (see Figure S11 and supplementary methods). D* here shows the modified aspartate (3-amino-2-oxobutanoic acid). Observable single fragmentations are colored. Structures and the corresponding observed mass resulting from multiple fragmentations following the rules of a, b, c, x, y, and z ions typically observed with CID are shown.[19-21] Note that there could be multiple isomers associated with certain mass values. In such cases, only one isomer is shown. A table with observed masses and the error compared to calculated values can be found in the Supplementary Dataset 3 (Excel).

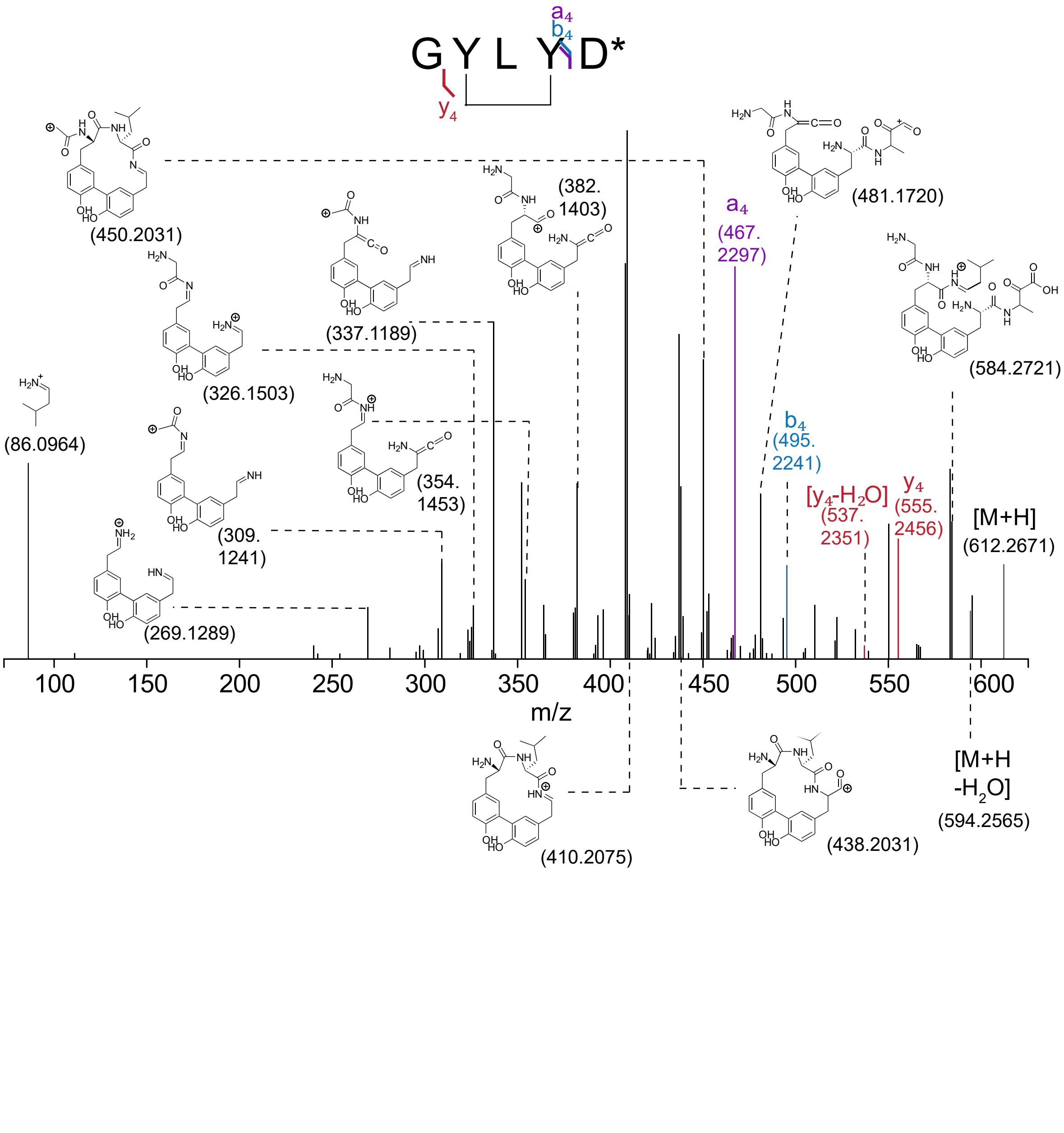

**Figure S5:** Deconvoluted HR-ESI-MS of ApyA co-expressed with all modifying enzymes except ApyH (pSCrhaB2_ApyOAIDS) in *Burkholderia* sp. FERM BP-3421. The sequence of the construct can be found in Table S2D. In this expression condition, ApyD has low activity (see Figure S11 and supplementary methods). The sequence of the peptide is shown below with the cut site bolded (endoproteinase GluC was used). *: sodium adduct. **: potassium adduct. Calculated exact mass for GluC-cleaved ApyO-modified ApyA: 627.2540 Da. A table with observed masses and the error compared to calculated values can be found in the Supplementary Dataset 3 (Excel).

MHHHHHHMATKPKKTKGVSVSIEGKLPKMTLDMPVDAKKIKAIQKCLENGKLTITMSKVDLAGG
RMG**E**GYLYD

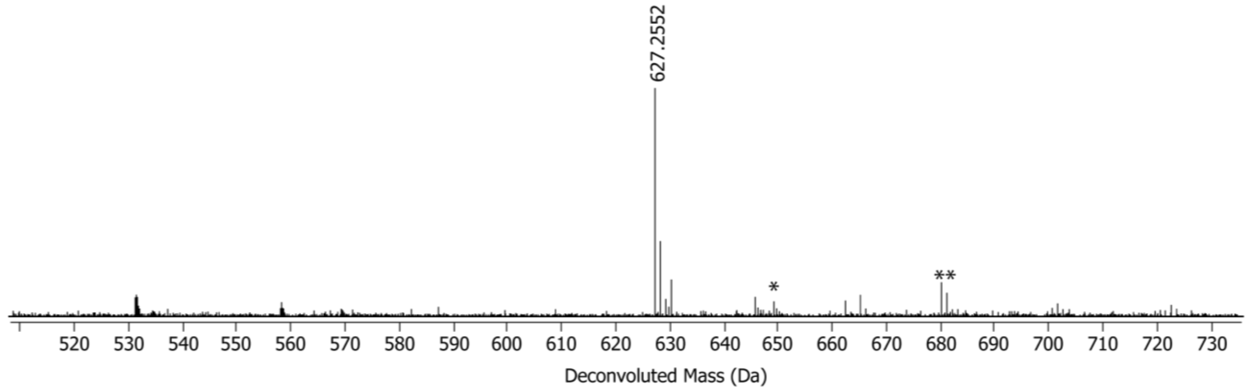

**Figure S6:** Deconvoluted HR-ESI-MS of ApyA co-expressed with all modifying enzymes except ApyI (pSCrhaB2_ApyOAHDS) in *Burkholderia* sp. FERM BP-3421. The sequence of the construct can be found in Table S2E. In this expression condition, ApyD has low activity (see Figure S11 and supplementary methods). The sequence of the peptide is shown below with the cut site bolded (endoproteinase GluC was used). *: sodium adduct. **: potassium adduct. Calculated exact mass for GluC-cleaved ApyO-modified ApyA: 627.2540 Da. A table with observed masses and the error compared to calculated values can be found in the Supplementary Dataset 3 (Excel).

MHHHHHHMATKPKKTKGVSVSIEGKLPKMTLDMPVDAKKIKAIQKCLENGKLTITMSKVDLAGG
RMG**E**GYLYD

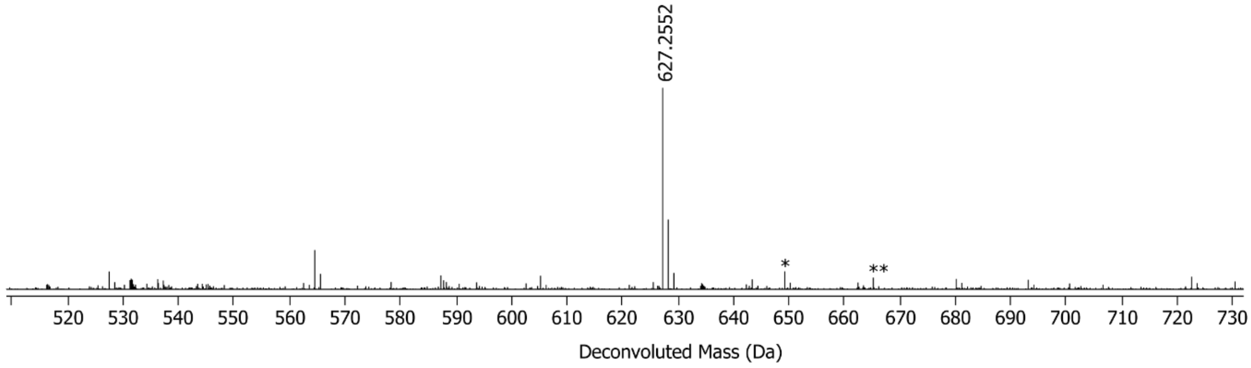

**Figure S7:** The HR-ESI-MS/MS of ApyA co-expressed with all modifying enzymes except ApyH (pSCrhaB2_ApyOAIDS) in *Burkholderia sp. FERM* BP-3421. The sequence can be found in Table S2D. Endoproteinase GluC was used for the proteolysis of modified ApyA. In this expression condition, ApyD has low activity (see Figure S11 and supplementary methods). Structures and the corresponding observed mass resulting from multiple fragmentations following the rules of a, b, c, x, y, and z ions typically observed with CID are shown. [19-21] Note that there could be multiple isomers associated with certain mass values. In such cases, only one isomer is shown. A table with observed masses and the error compared to calculated values can be found in the Supplementary Dataset 3 (Excel).

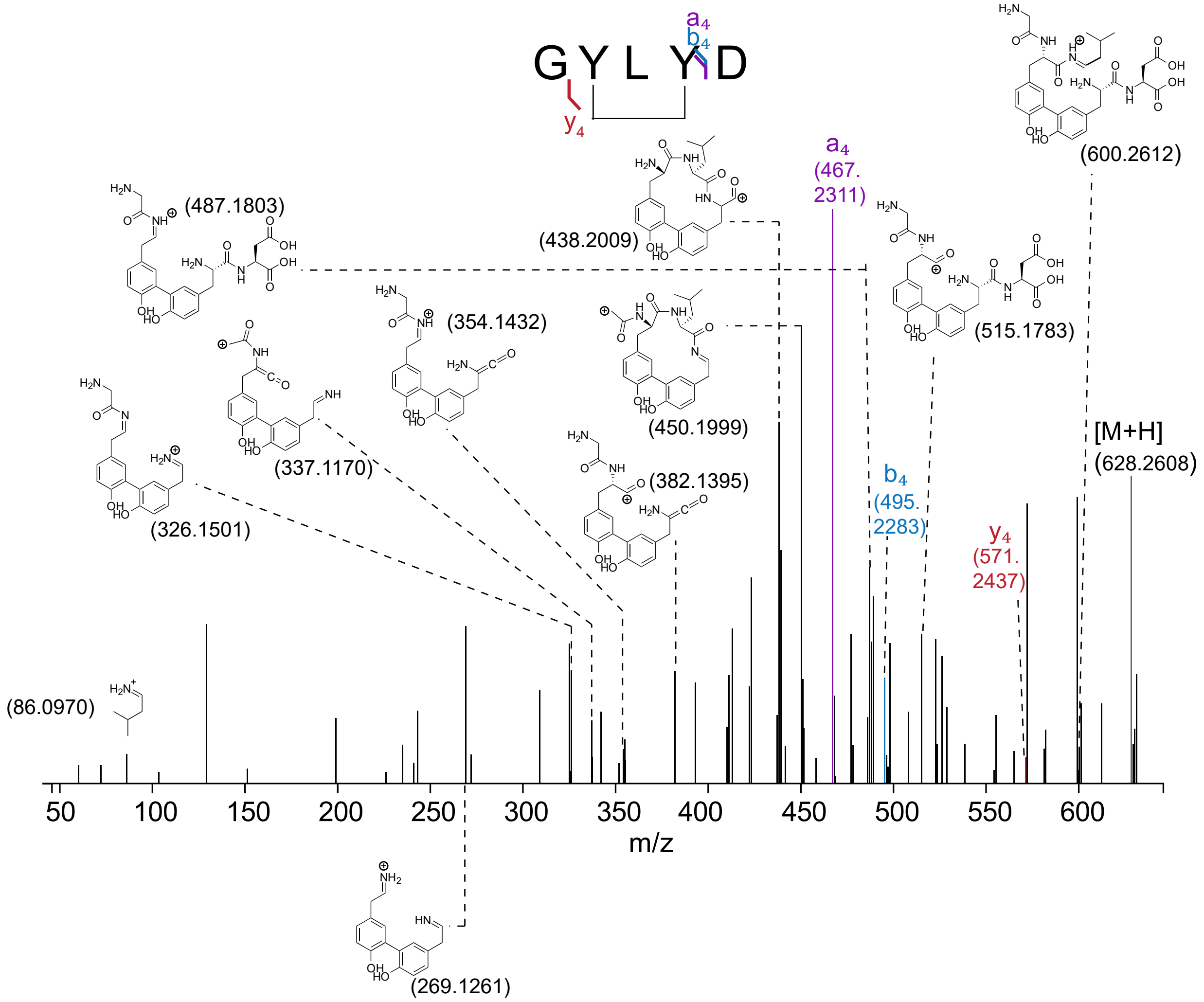

**Figure S8:** The HR-ESI-MS/MS of ApyA co-expressed with all modifying enzymes except ApyI (pSCrhaB2_ApyOAHDS) in *Burkholderia* sp. FERM BP-3421. The sequence of the construct can be found in Table S2E. In this expression condition, ApyD has low activity (see Figure S11 and supplementary methods). Structures and the corresponding observed mass resulting from multiple fragmentations following the rules of a, b, c, x, y, and z ions typically observed with CID are shown. [19-21] Note that there could be multiple isomers associated with certain mass values. In such cases, only one isomer is shown. A table with observed masses and the error compared to calculated values can be found in the Supplementary Dataset 3 (Excel).

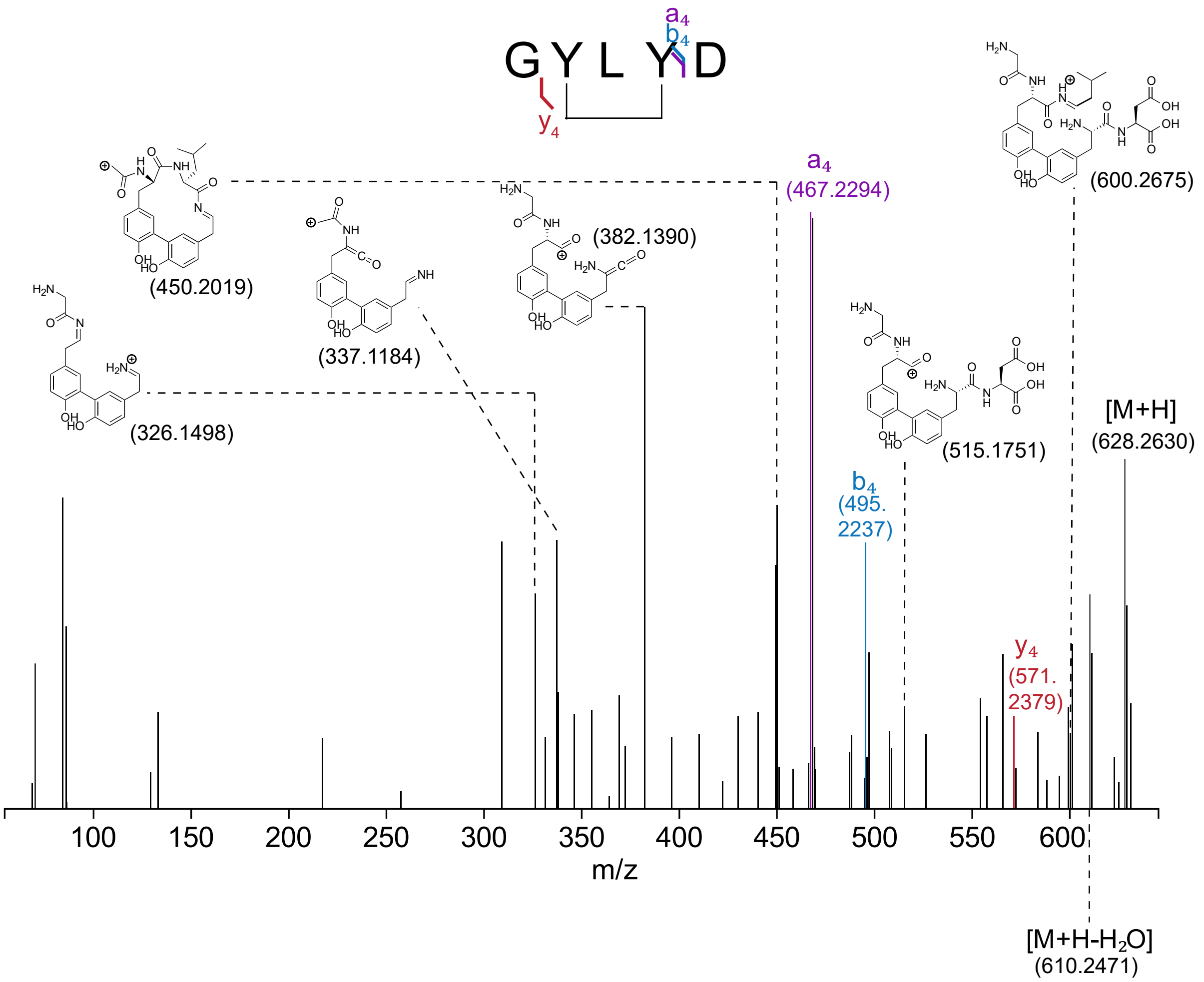

**Figure S9:** Deconvoluted HR-ESI-MS of ApyA co-expressed with all modifying enzymes except ApyS (pSCrhaB2_ApyOAHID) in *Burkholderia* sp. FERM BP-3421. The sequence of the construct can be found in Table S2F. In this expression condition, ApyD has low activity (see Figure S11 and supplementary methods). The sequence of the peptide is shown below with the cut site bolded (endoproteinase GluC was used). *: sodium adduct, **: potassium adduct, #: potential methionine oxidation (consistent with the peak disappearing in the GluC-digested peptide. Calculated exact mass for ApyO/H/I-modified ApyA: 8074.2000 Da. Calculated exact mass for GluC-cleaved ApyO/H/I-modified ApyA: 597.2435 Da. A table with observed masses and the error compared to calculated values can be found in the Supplementary Dataset 3 (Excel).

MHHHHHHMATKPKKTKGVSVSIEGKLPKMTLDMPVDAKKIKAIQKCLENGKLTITMSKVDLAGG RMG**E**GYLYD

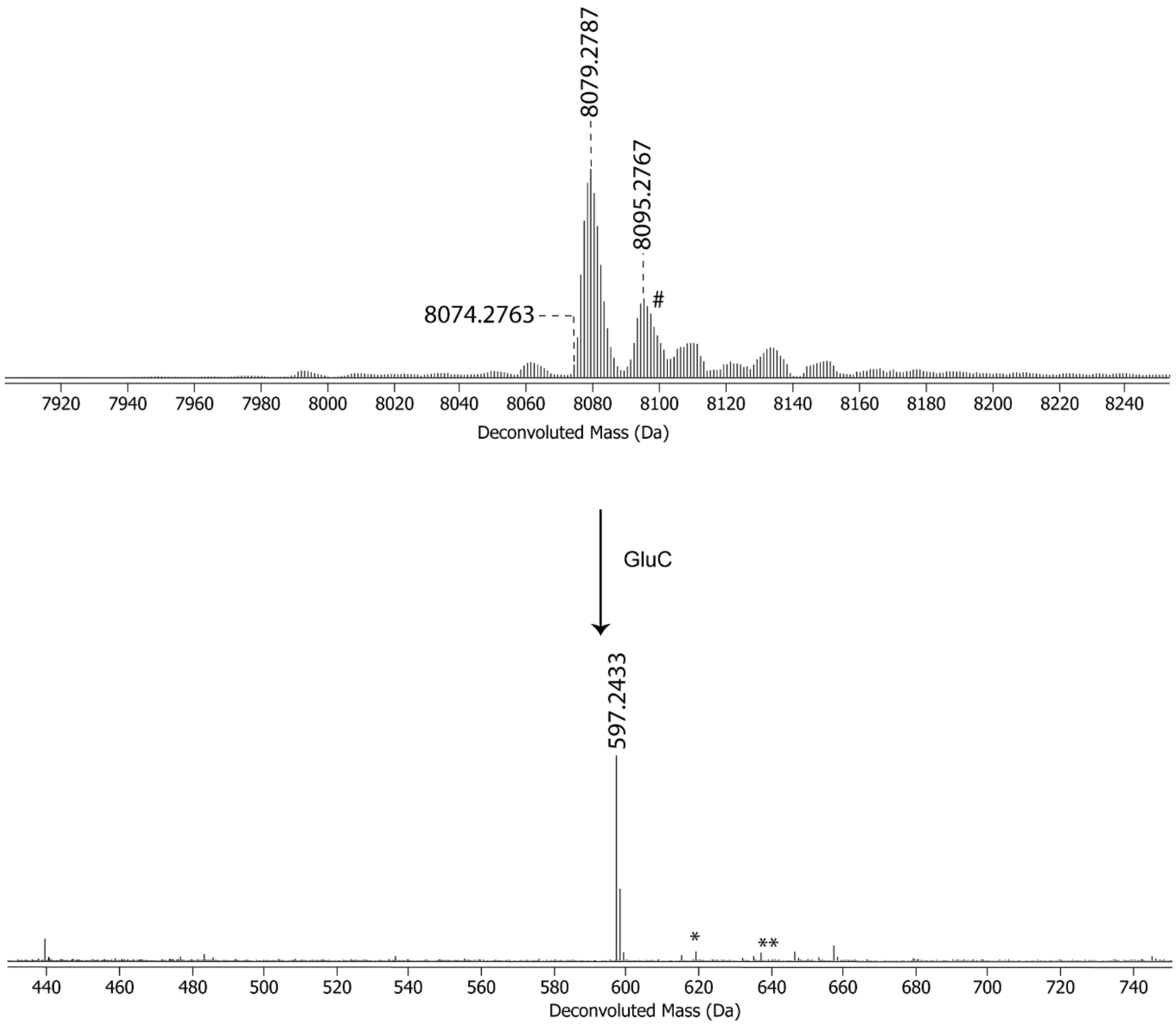

**Figure S10:** HR-ESI-MS/MS of ApyA co-expressed with all modifying enzymes except ApyS (pSCrhaB2_ApyOAHID) in *Burkholderia* sp. FERM BP-3421. D** represents the modified aspartate (aminopyruvic acid) in *Burkholderia* sp. FERM BP-3421. The sequence of the construct can be found in Table S2F. In this expression condition, ApyD has low activity (see Figure S11 and supplementary methods). Structures and the corresponding observed mass resulting from multiple fragmentations following the rules of a, b, c, x, y, and z ions typically observed with CID are shown.[19-21] Note that there could be multiple isomers associated with certain mass values. In such cases, only one isomer is shown. A table with observed masses and the error compared to calculated values can be found in the Supplementary Dataset 3 (Excel).

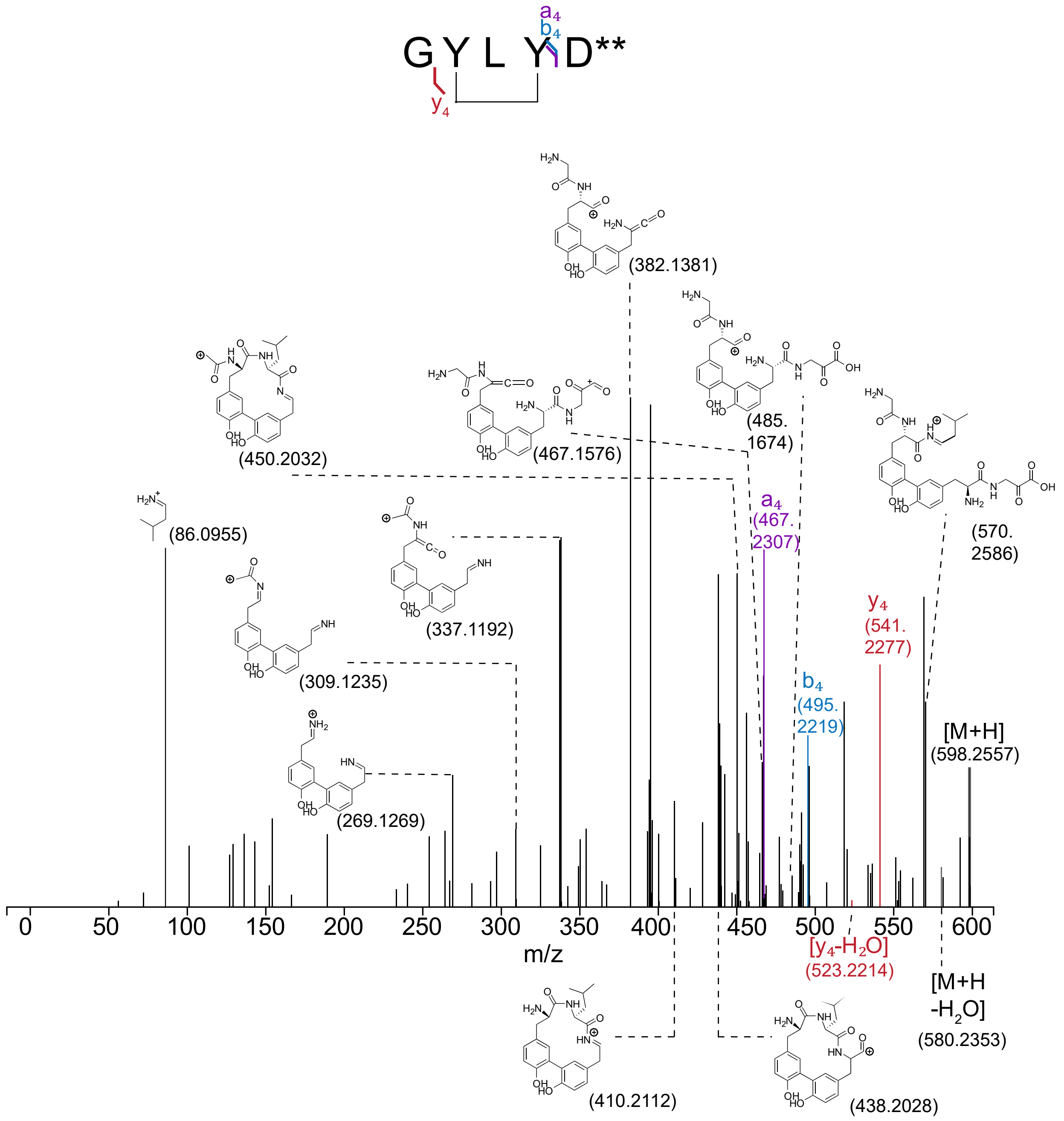

**Figure S11:** Deconvoluted HR-ESI-MS of ApyA co-expressed with all modifying enzymes in *Burkholderia.* sp FERM BP-3421 using different expression conditions. The conditions can be found in the supplementary methods. The sequence and vector map of the constructs are in Table S2B, Table S3, and Figure S14. The sequence of the peptide is shown below with the cut site bolded (endoproteinase LysC was used). Calculated exact mass for ApyO/H/I//S-modified ApyA cleaved with LysC without ApyD catalyzed methylation: 1596.7242 Da (1614.7347 Da for the hydrated diol form). Calculated exact mass for LysC-cleaved ApyO/H/I/D/S-modified ApyA with ApyD-catalyzed methylation: 1610.7398 Da (1628.7503 Da for the hydrated diol form). *: sodium adduct of ApyO/H/I/S-modified ApyA, **: sodium adduct of ApyD/O/H/I/S-modified ApyA. Masses with +18 Da of the modified species are observed. These masses correspond to the water adduct of the modified species (Figure S12-S13) due to the presence of a ketone. A table with observed masses and the error compared to calculated values can be found in the Supplementary Dataset 3 (Excel).

MHHHHHHMATKPKKTKGVSVSIEGKLPKMTLDMPVDAKKIKAIQKCLENGKLTITMS**K**VDLAGG
RMGEGYLYD

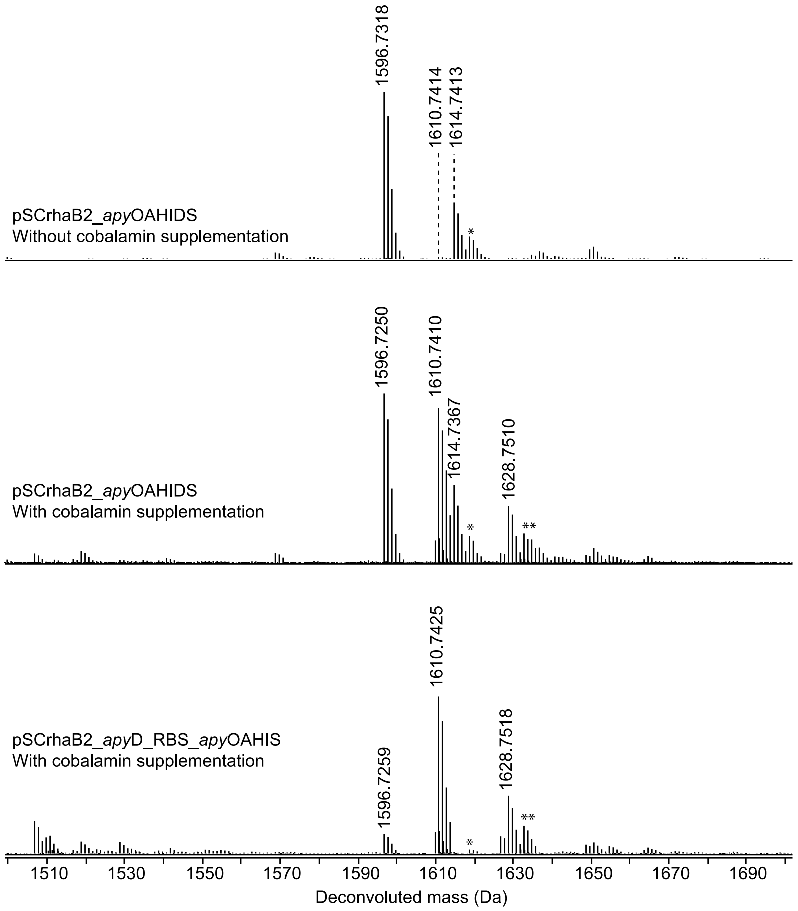

**Figure S12**: HR-ESI-MS/MS of the water adduct of LysC-cleaved ApyO/H/I/S-modified ApyA. A table with observed masses and the error compared to calculated values can be found in the Supplementary Dataset 3 (Excel). D* here shows the modified aspartate (3-amino-2-oxobutanoic acid).

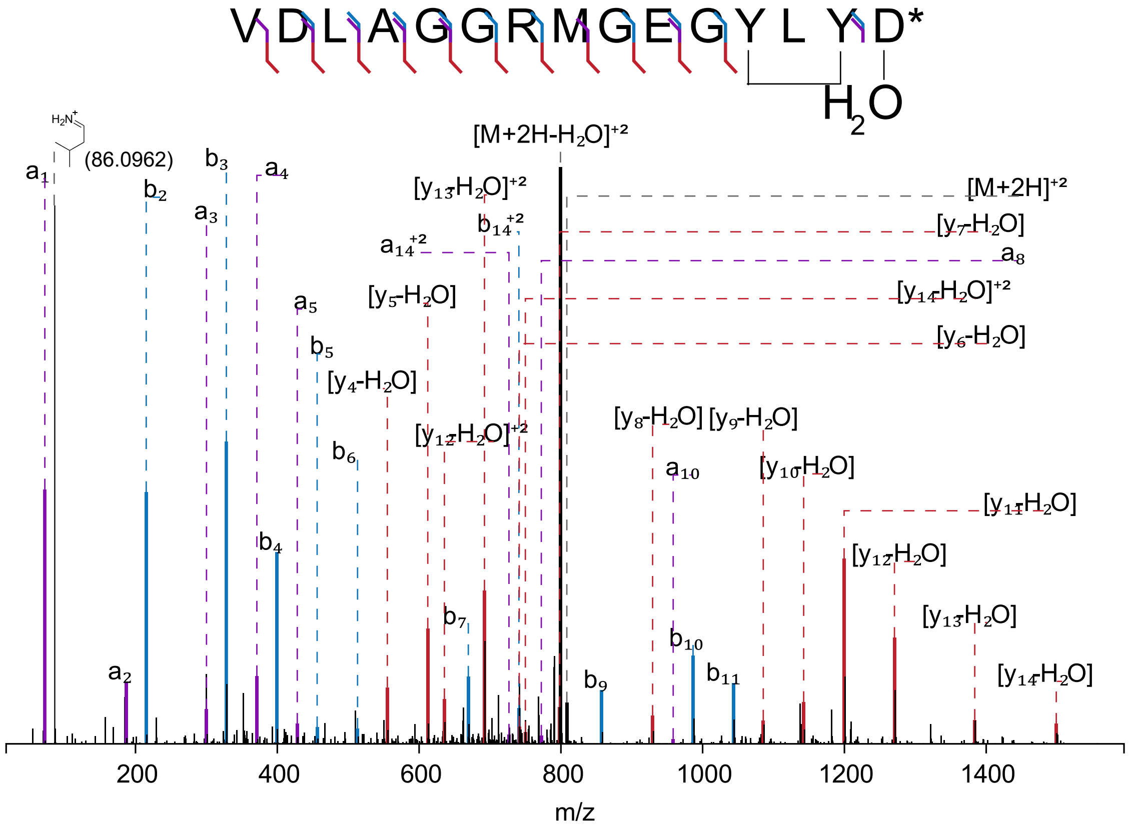

**Figure S13:** HR-ESI-MS/MS of the water adduct of LysC-cleaved ApyD/O/H/I/S-modified ApyA. A table with observed masses and the error compared to calculated values can be found in the Supplementary Dataset 3 (Excel). D* here shows the modified aspartate (3-amino-2-oxobutanoic acid).

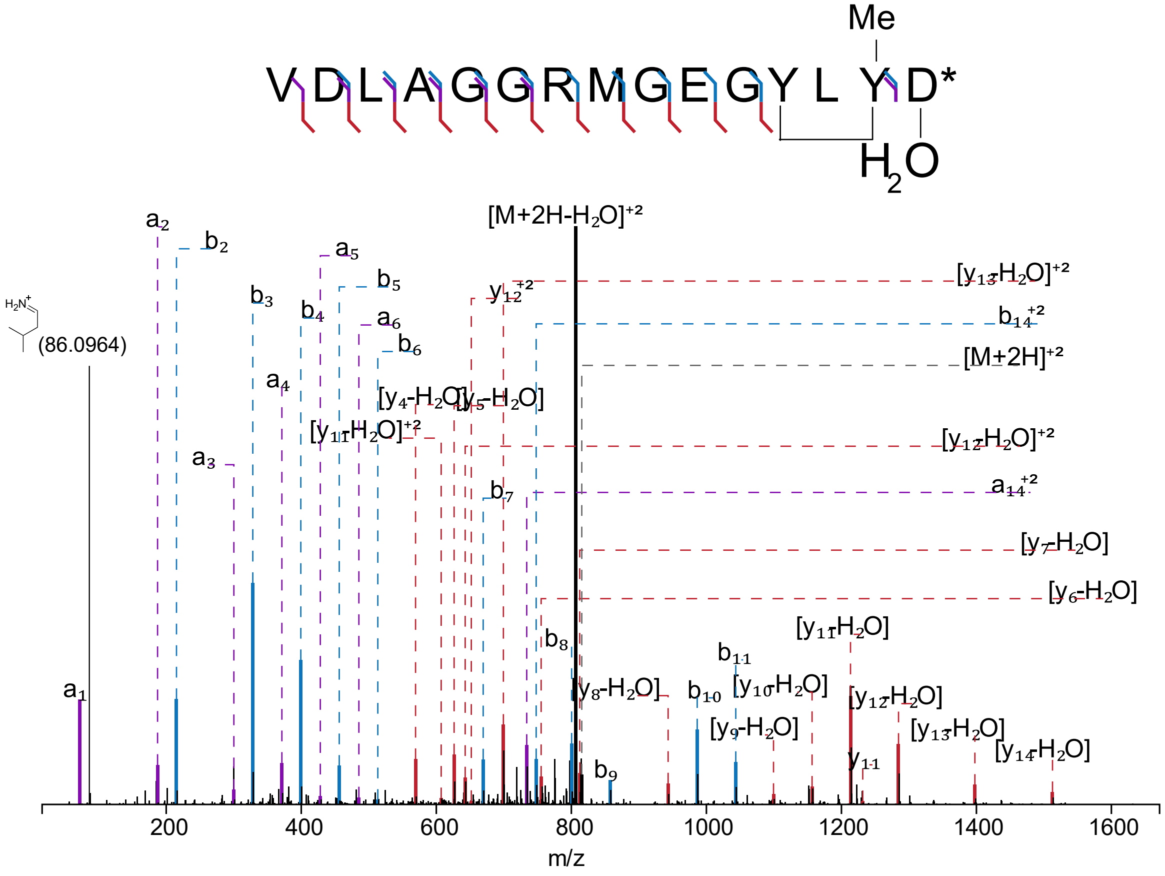

**Figure S14:** Plasmid map of constructs in Table S2B (A) and Table S3 (B). Artificial RBS (ribosome binding sites) here represent RBS that are not present in the genomic DNA from *B. thailandensis* E264 encoding ApyO/A/H/I/D/S. These RBS are either from the plasmid vector pSCrhaB2 or introduced for expression optimization.

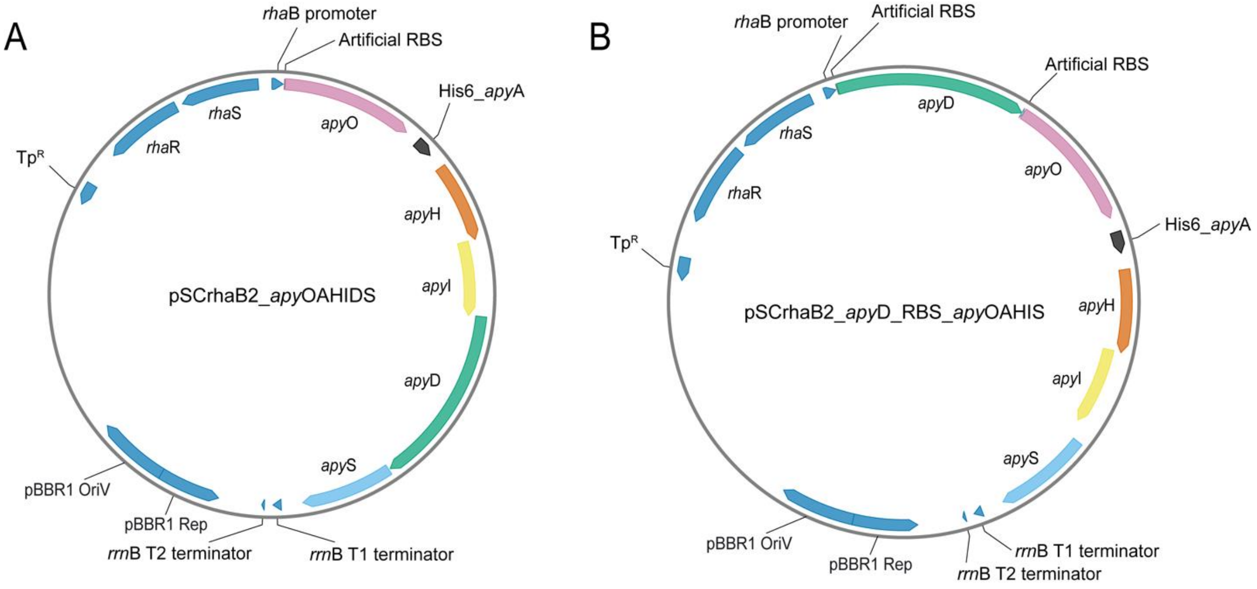

**Figure S15:** Deconvoluted HR-ESI-MS of ApyA co-expressed with modifying enzymes in *Burkholderia* sp. FERM BP-3421 (pSCrhaB2_ApyD_RBS_ApyOAHIS). The sequence of the construct can be found in Table S3. The sequence of the peptide is shown below with the cut site bolded (endoproteinase GluC was used). #: potential methionine oxidation (consistent with the oxidation peak not being present in the GluC-digested peptide). Calculated exact mass for ApyO/H/I/D/S-modified ApyA: 8102.2312 Da. Calculated exact mass for GluC-cleaved ApyO/H/I/D/S-modified ApyA: 625.2747 Da. A table with observed masses and the error compared to calculated values can be found in the Supplementary Dataset 3 (Excel).

MHHHHHHMATKPKKTKGVSVSIEGKLPKMTLDMPVDAKKIKAIQKCLENGKLTITMSKVDLAGG
RMG**E**GYLYD

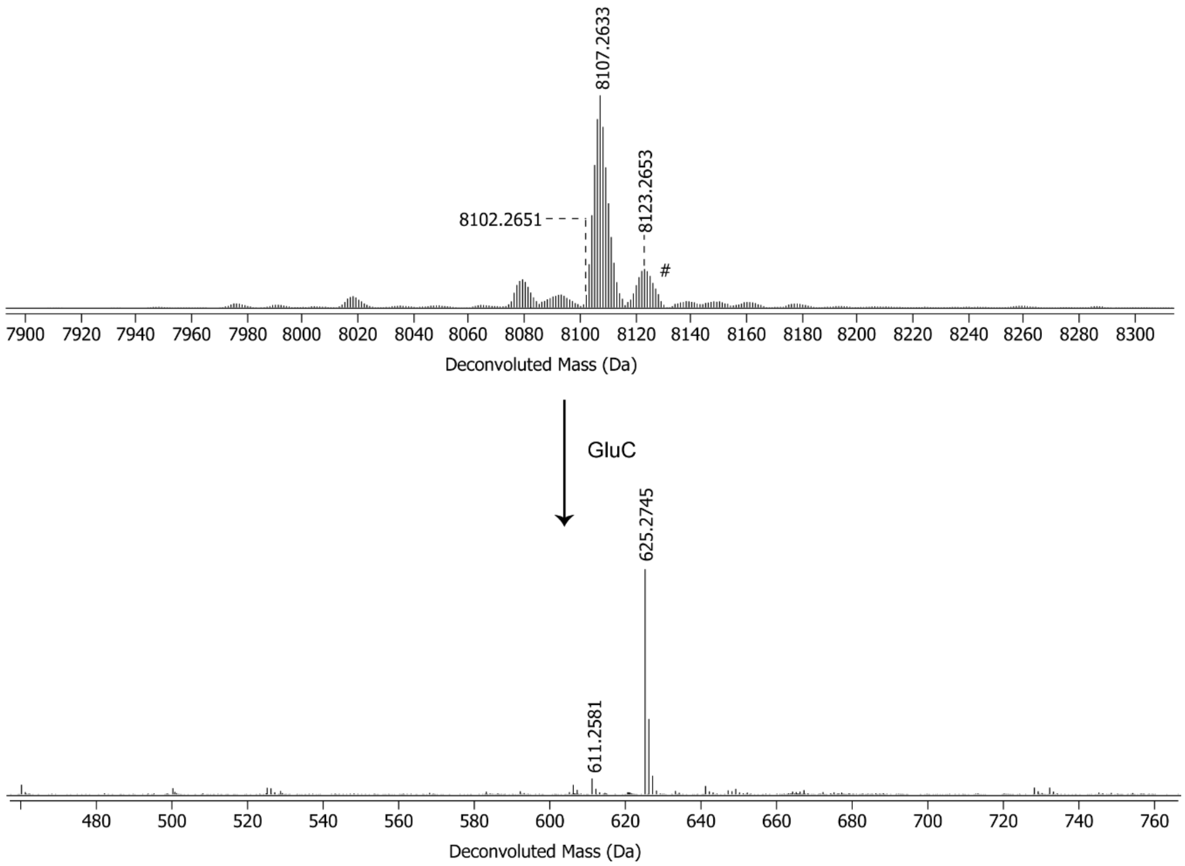

**Figure S16:** The HR-ESI-MS/MS of ApyA co-expressed with the modifying enzymes in *Burkholderia* sp. FERM BP-3421 (pSCrhaB2_ApyD_RBS_ApyAOHIS). The sequence of the construct can be found in Table S3. D* represents the modified aspartate (3-amino-2-oxobutanoic acid in this experiment). Structures and the corresponding observed mass resulting from multiple fragmentations following the rules of a, b, c, x, y, and z ions typically observed with CID are shown.[19-21] Note that there could be multiple isomers associated with certain mass values. In such cases, only one isomer is shown. A table with observed masses and the error compared to calculated values can be found in the Supplementary Dataset 3 (Excel).

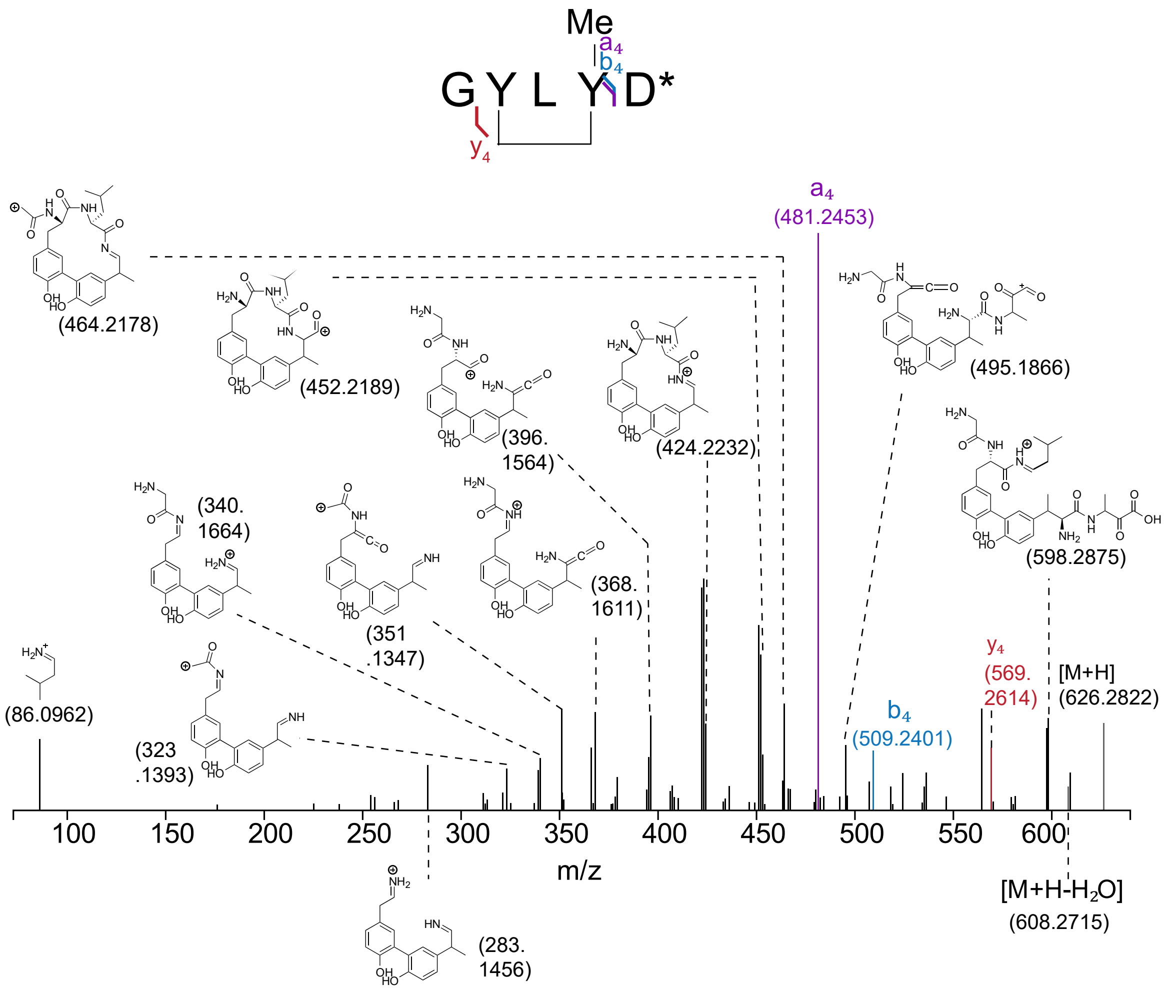

**Figure S17:** NMR spectra of ApyA/D-modified product after GluC cleavage. The solvent used was 40% ACN-d3/60% $H_2O$. The structure of the product is shown in Figure 3 (main text).

A) $^1$H-NMR

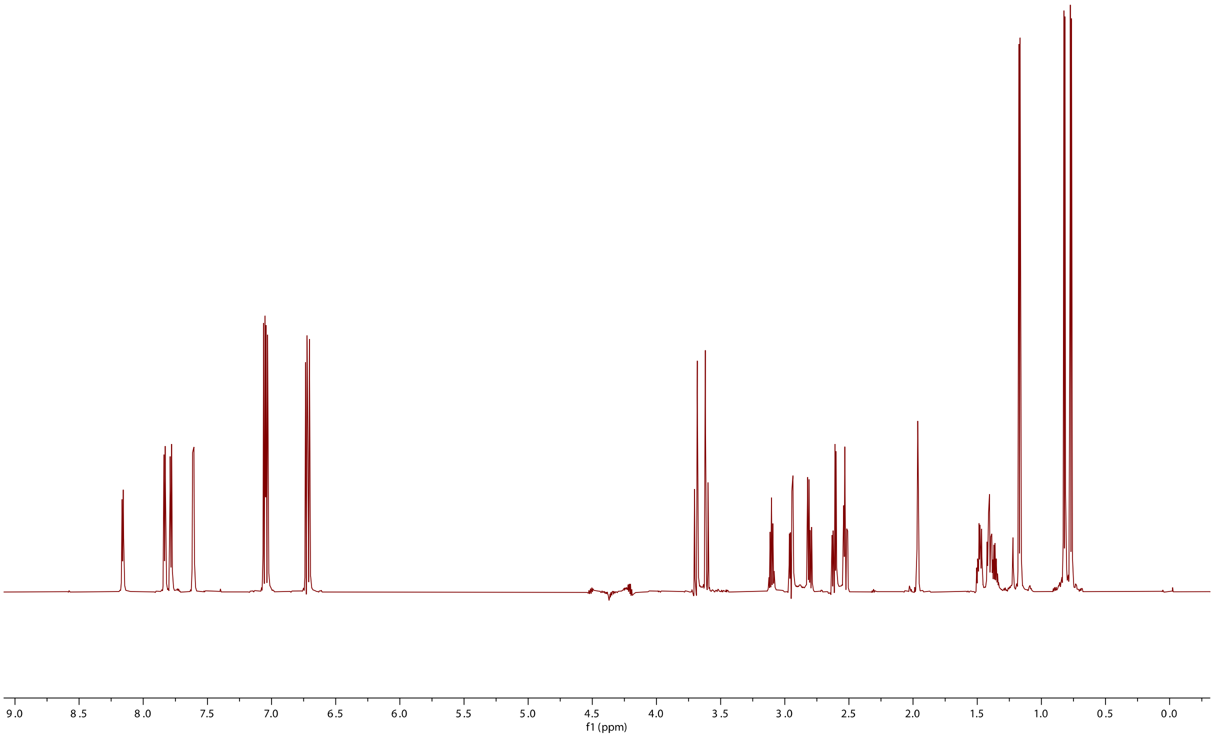

B) $^{1}$H-$^{1}$H TOCSY

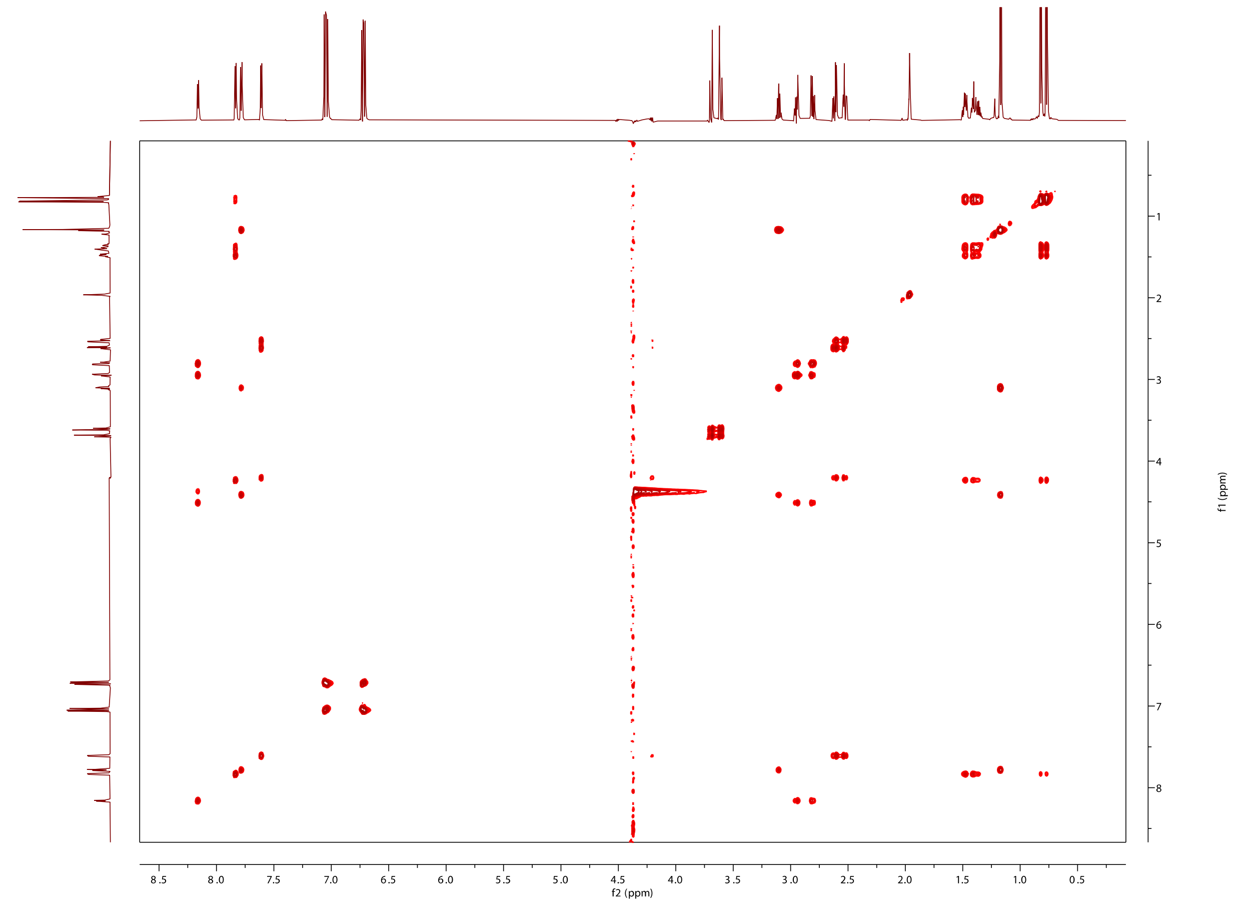

8.5 8.0 7.5 7.0 6.5 6.0 5.5 5.0 4.5 4.0 3.5 3.0 2.5 2.0 1.5 1.0 0.5
f2 (ppm)
1 2 3 4 5 6 7 8
f1 (ppm)

C) $^{1}$H-$^{1}$H ROESY

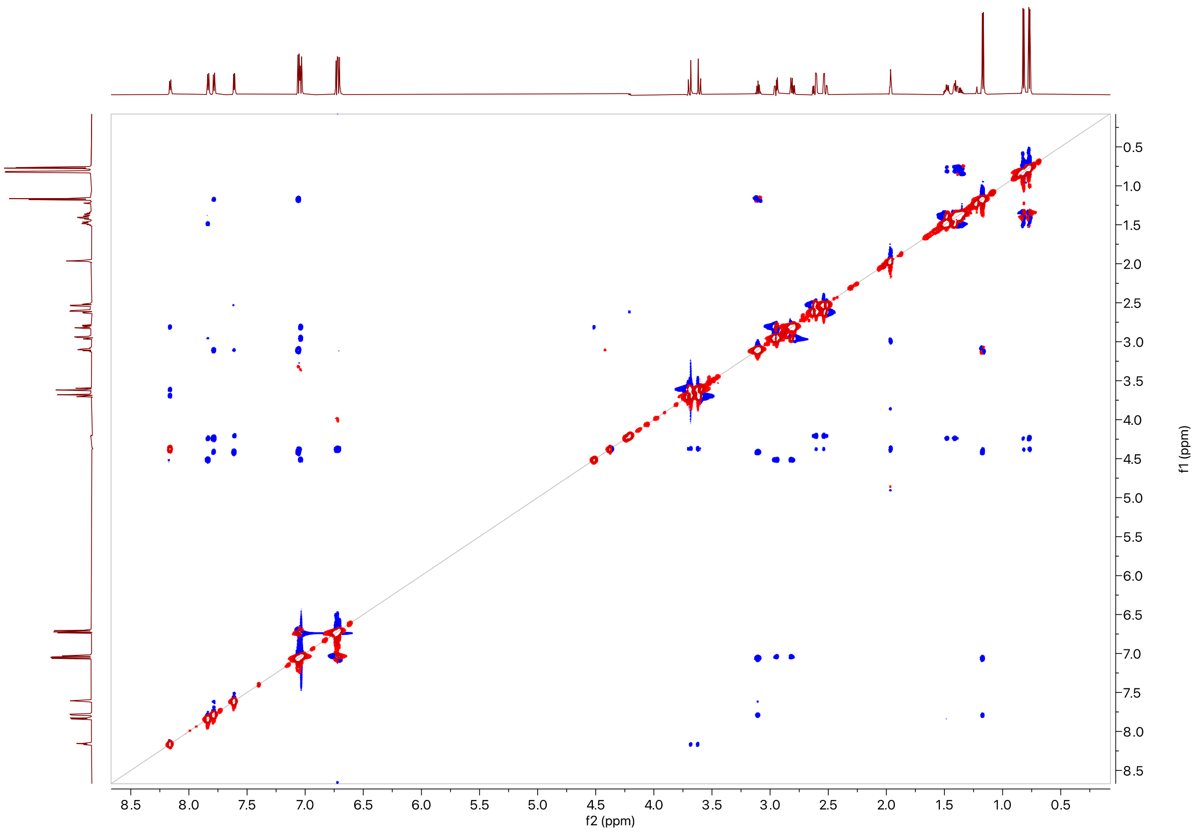

D) $^{1}H$-$^{13}C$ HSQC

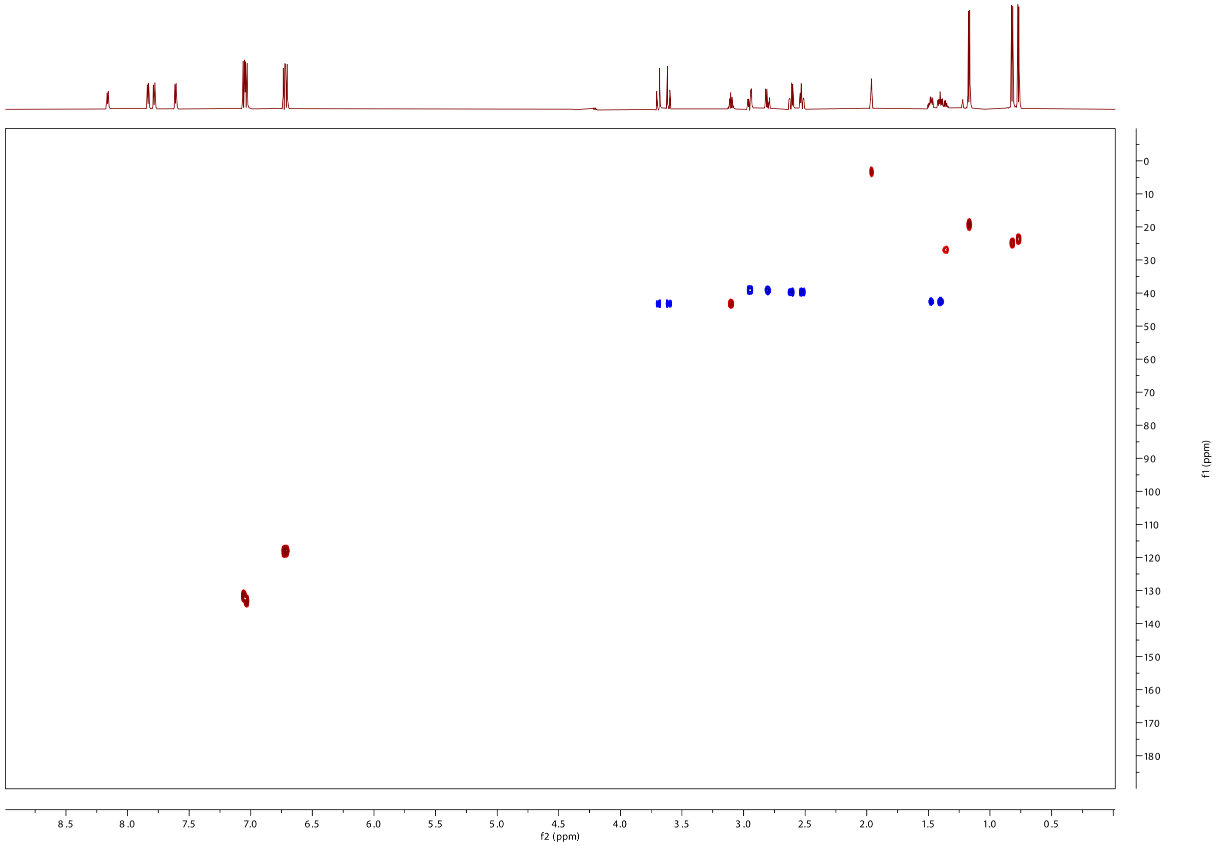
8.5 8.0 7.5 7.0 6.5 6.0 5.5 5.0 4.5 4.0 3.5 3.0 2.5 2.0 1.5 1.0 0.5
f2 (ppm)
0 10 20 30 40 50 60 70 80 90 100 110 120 130 140 150 160 170 180
f1 (ppm)

E) $^{1}H$-$^{13}C$ HMBC

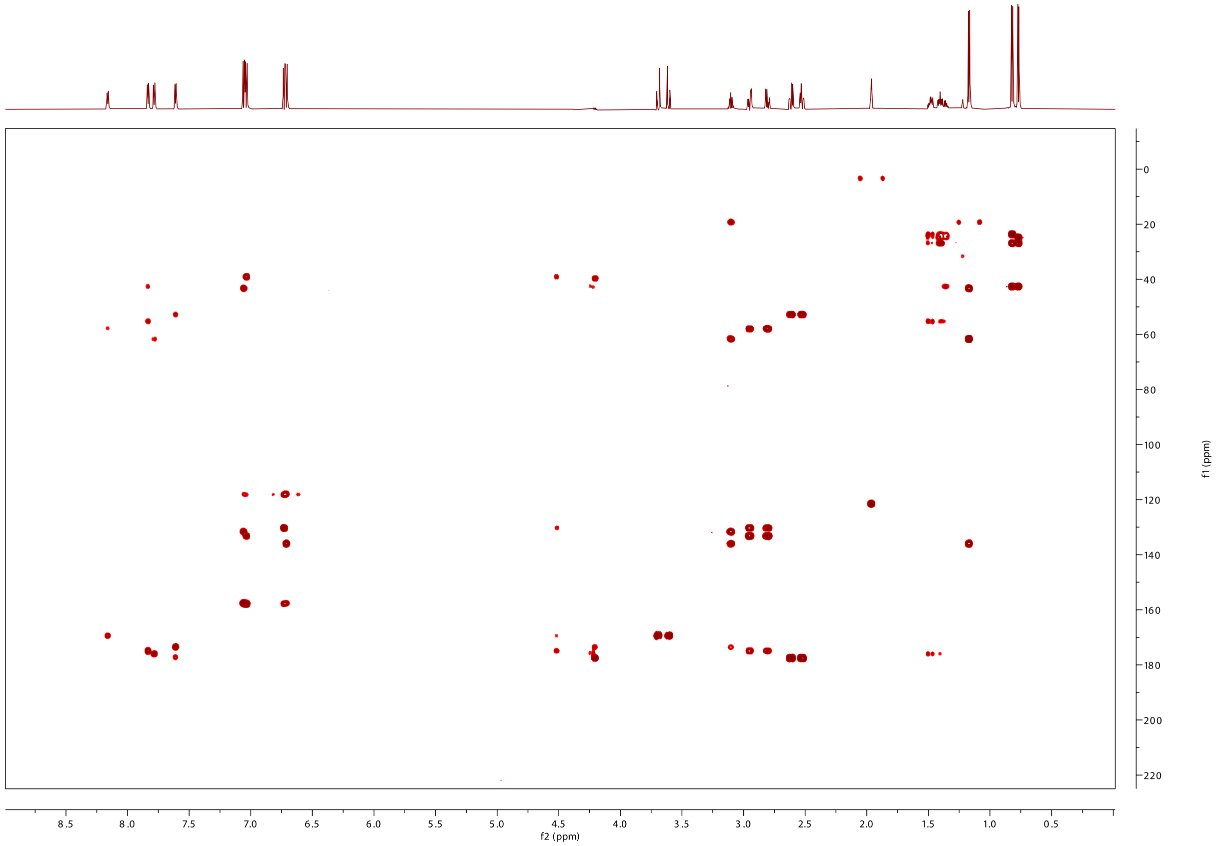

F) $^{1}H$-$^{13}C$ HMBC key cross peaks to assign the structure to β-methyltyrosine. See also Figure 3 (main text).

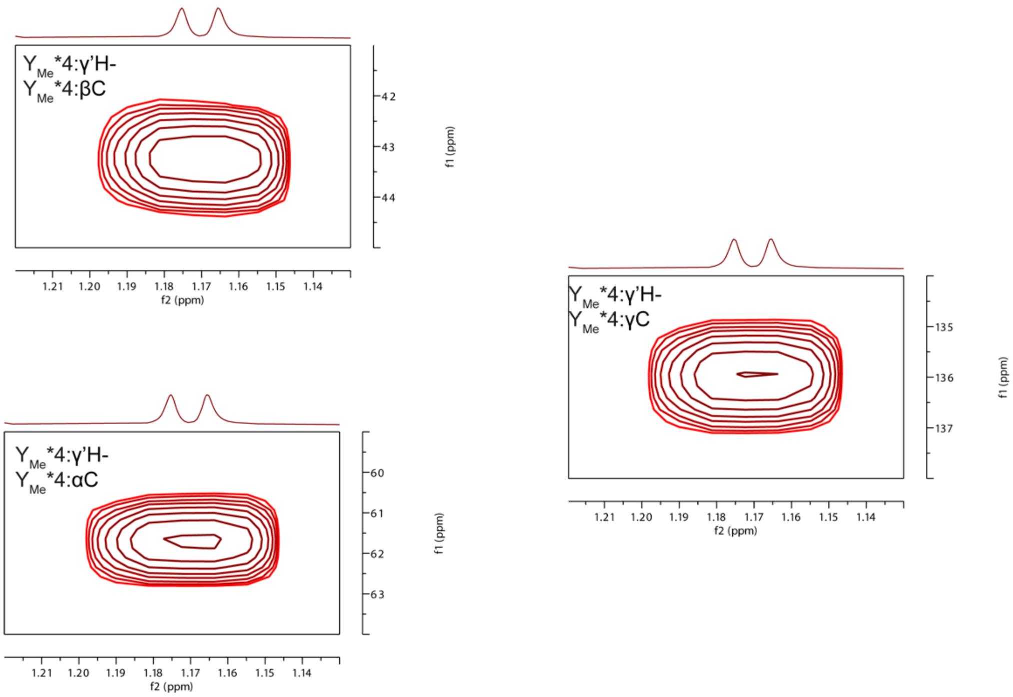

**Table S4:** Summary of $^{1}$H and $^{13}$C chemical shifts for the ApyA/D-modified product after GluC cleavage.

| residue number | AA | NH/C=O | αH/C | βH/C | γH/C | δH/C and others |
|---|---|---|---|---|---|---|
| 1 | Gly | 169.4 | 3.69, 3.62<br>43.3 | | | |
| 2 | Tyr | 8.16<br>175.0 | 4.50<br>58.0 | 2.94, 2.80<br>39.2 | 130.3 | 7.04, 133.5 (ortho)<br>6.73, 118.4 (meta)<br>C-OH: 157.6 |
| 3 | Leu | 7.83<br>176.0 | 4.23<br>55.3 | 1.48, 1.40<br>42.7 | 1.36<br>27.1 | 0.82, 25.1<br>0.77, 23.7 |
| 4 | (β–Me)Tyr | 7.78<br>173.6 | 4.42<br>61.6 | 3.10<br>43.1 | 135.9 | 1.17, 19.3 (β–Methyl)<br>7.06, 131.6 (ortho)<br>6.71, 118.0 (meta)<br>C-OH: 158.0 |
| 5 | Asp | 7.61<br>177.2 | 4.20<br>52.8 | 2.61, 2.52<br>39.8 | | 177.6 |

**Figure S18:** Marfey's analysis of the ApyAD-modified product. A) Marfey's derivatization scheme to determine the stereochemistry of β-methyltyrosine. B) Extracted ion chromatogram (EIC *m/z* = 700.1937) of hydrolyzed ApyA and subsequently derivatized β-methyltyrosine using L-FDAA as the derivatization reagent. Commercial isomers of β-methyltyrosine were derivatized in the same manner and co-injected with the ApyA-derived material. Results using D-FDAA are shown in Figure 3D (main text). In this experiment, treatment of modified ApyA and standards in 6 M DCL resulted in exchange of the ortho protons of the Tyr residues with solvent.[22,23] DCl is used such that any potential epimerization can be detected by incorporation of additional deuterium.

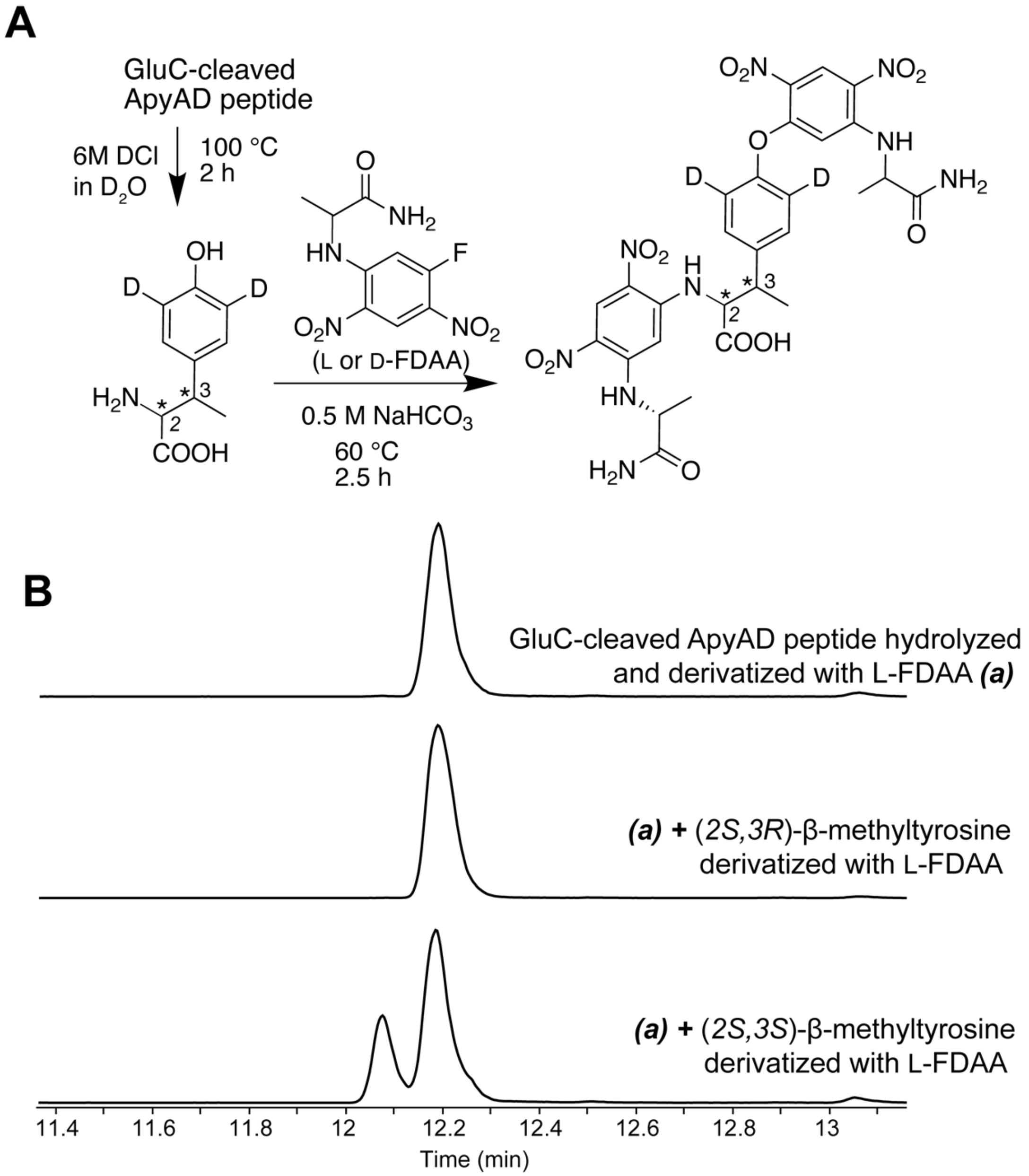

**Figure S18:** NMR spectra of ApyA after ApyO modification and GluC cleavage. The solvent was 10% $D_2O$/90% $H_2O$.

A) Depiction of elucidated structure based on key correlations shown

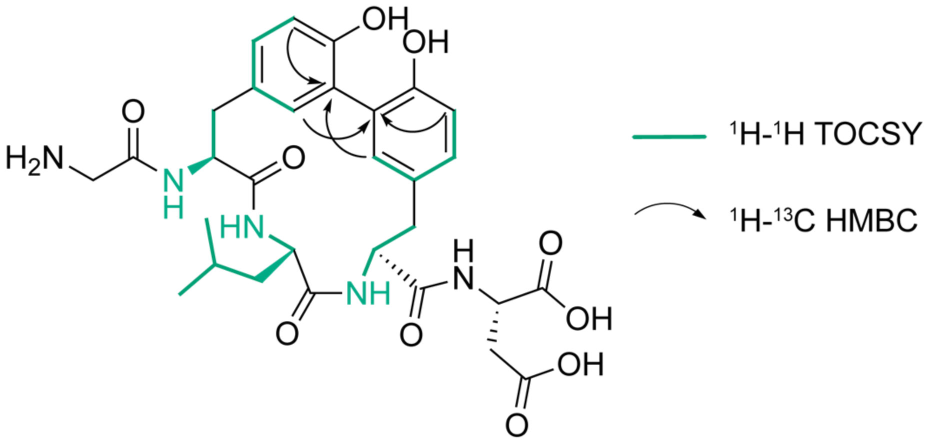

B) [1]H-NMR spectrum. The major singlet at 1.76 ppm is residual ammonium acetate from HPLC purification.

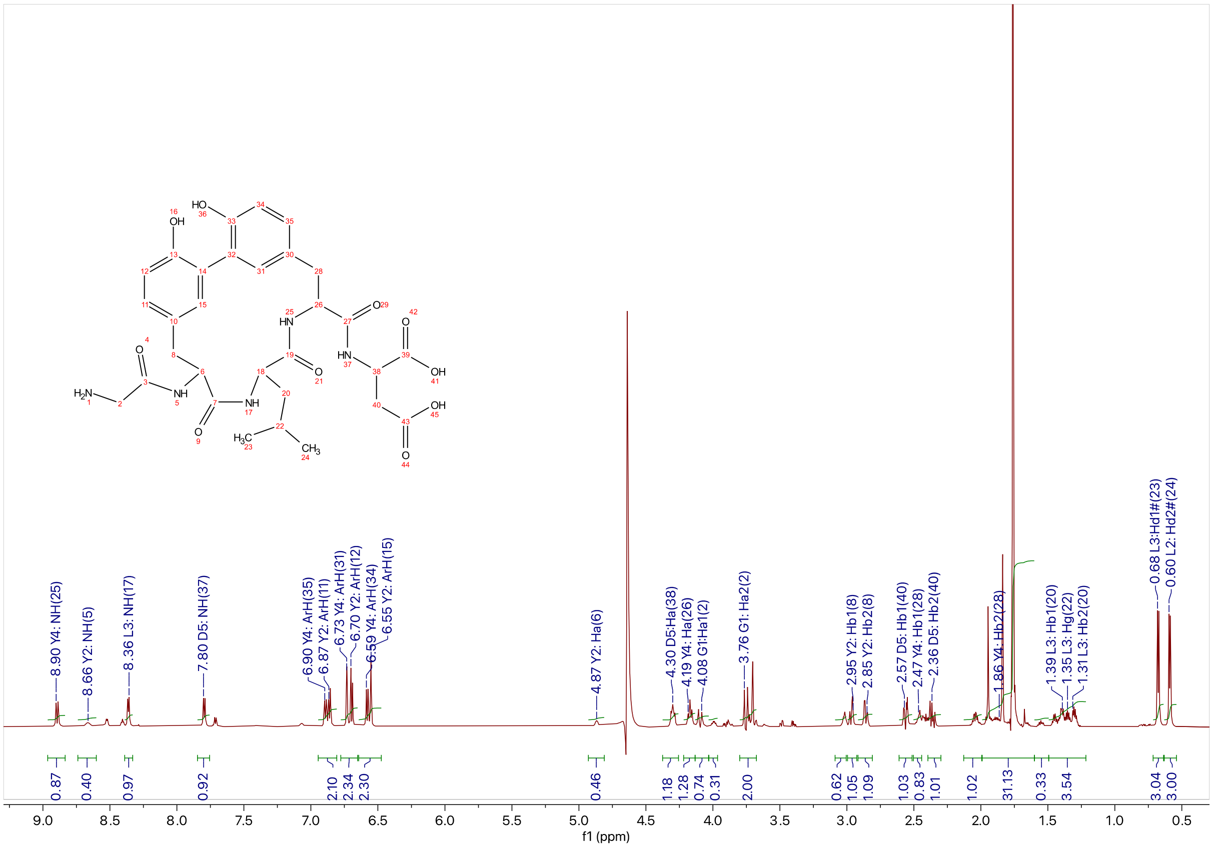

C) $^{1}H$-$^{1}H$ TOCSY

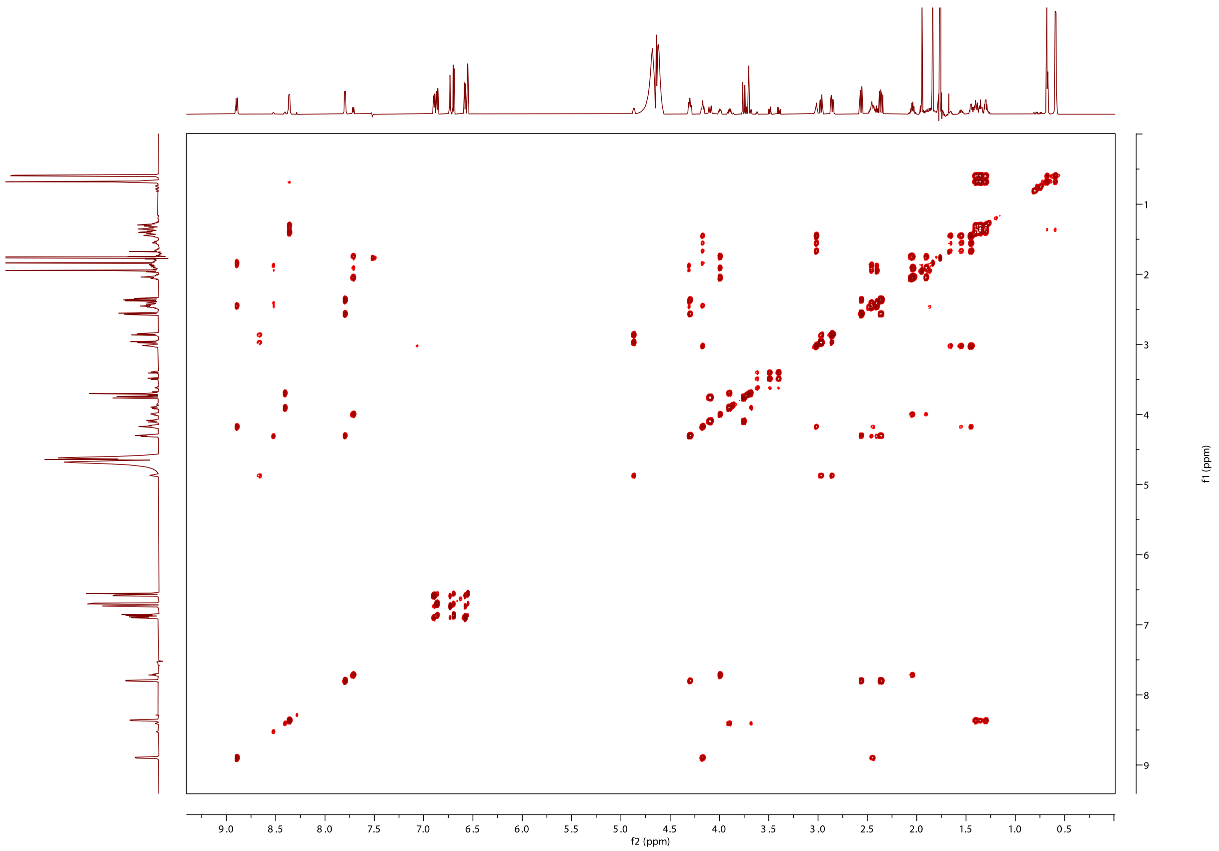

D) $^{1}H$-$^{1}H$ NOESY

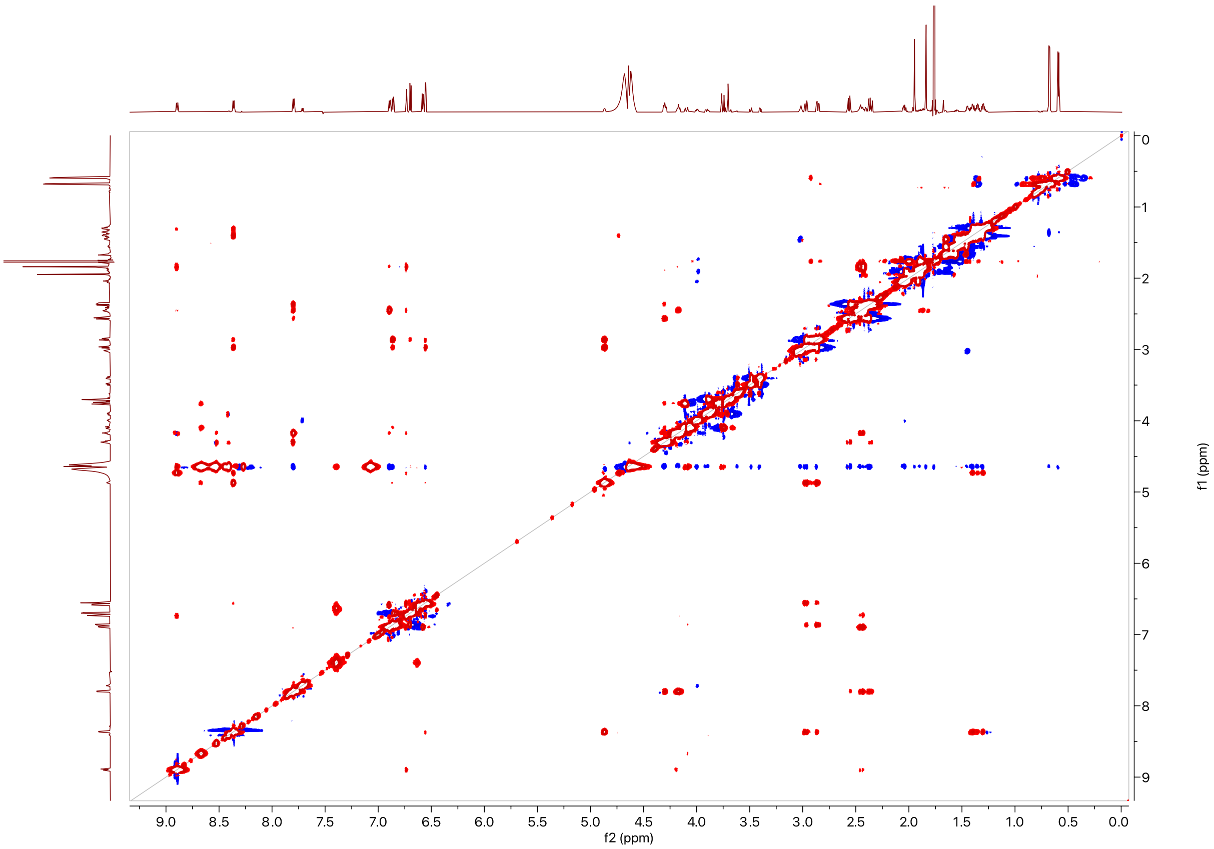

f2 (ppm)
f1 (ppm)
9.0
8.5
8.0
7.5
7.0
6.5
6.0
5.5
5.0
4.5
4.0
3.5
3.0
2.5
2.0
1.5
1.0
0.5
0.0
0
1
2
3
4
5
6
7
8
9

E) $^{1}H$-$^{13}C$ HSQC

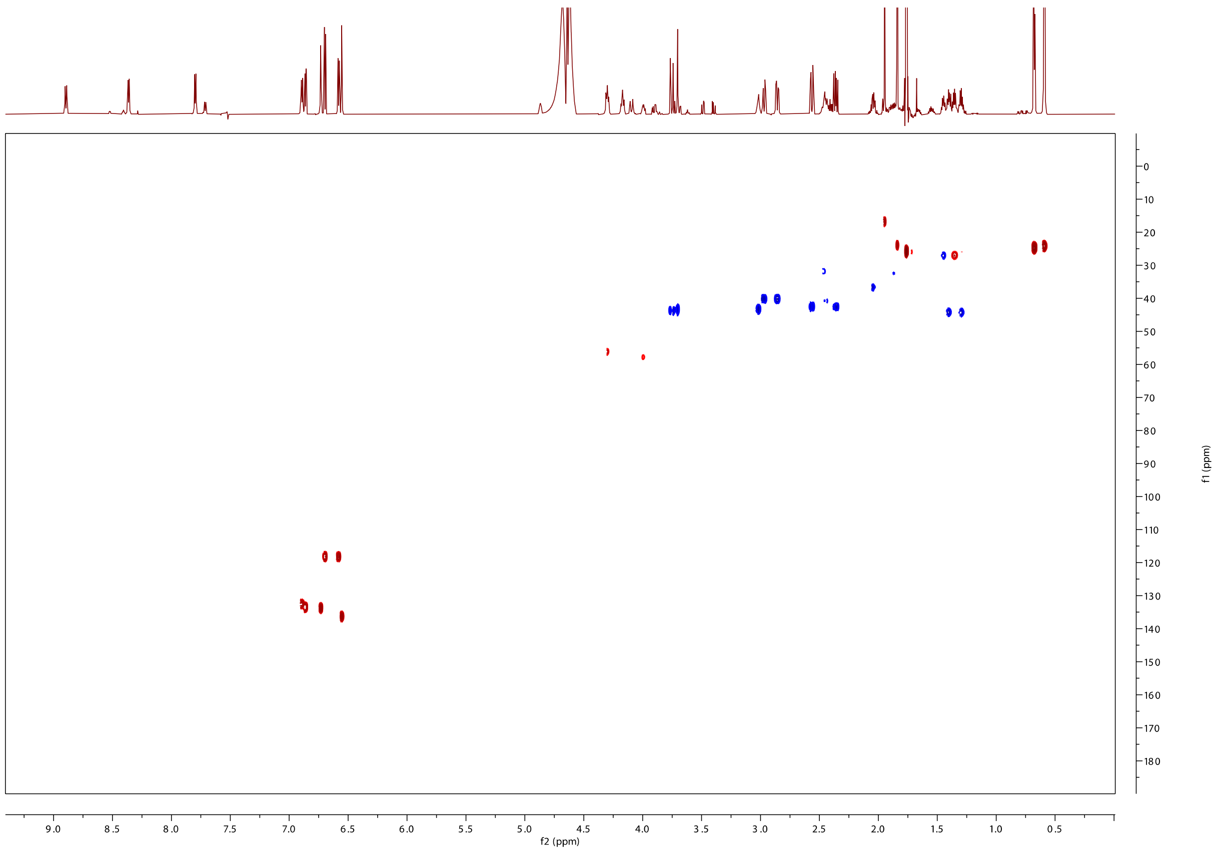

F) $^{1}H$-$^{13}C$ HMBC

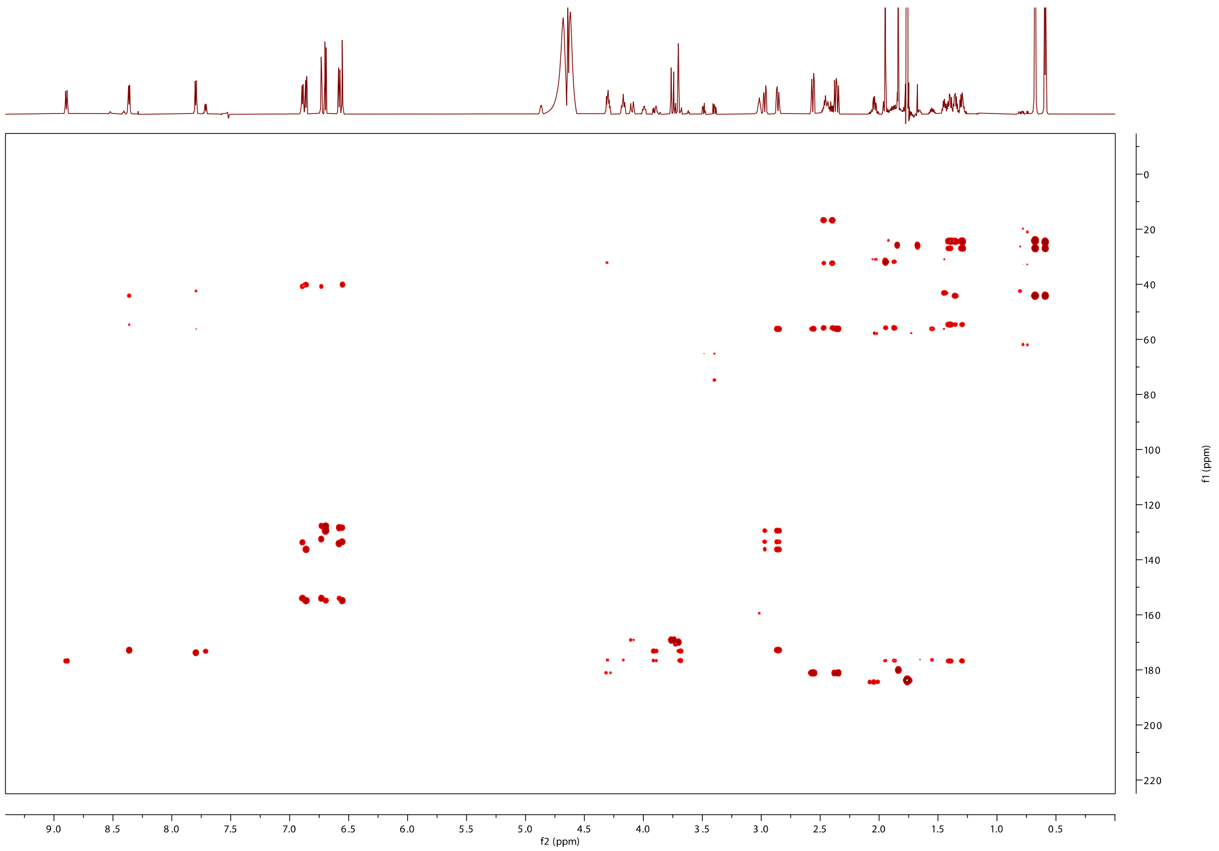

### G) Assignment of amino acids using $^1H$-$^1H$ TOCSY data

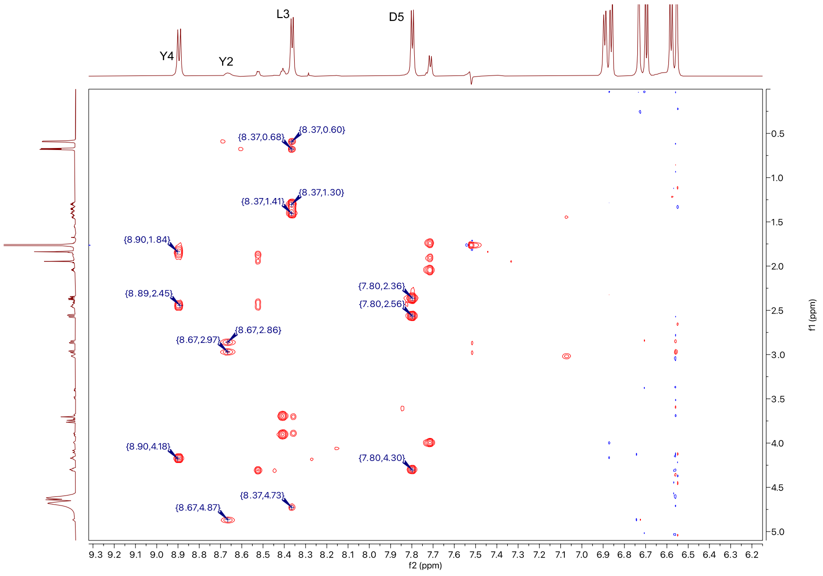

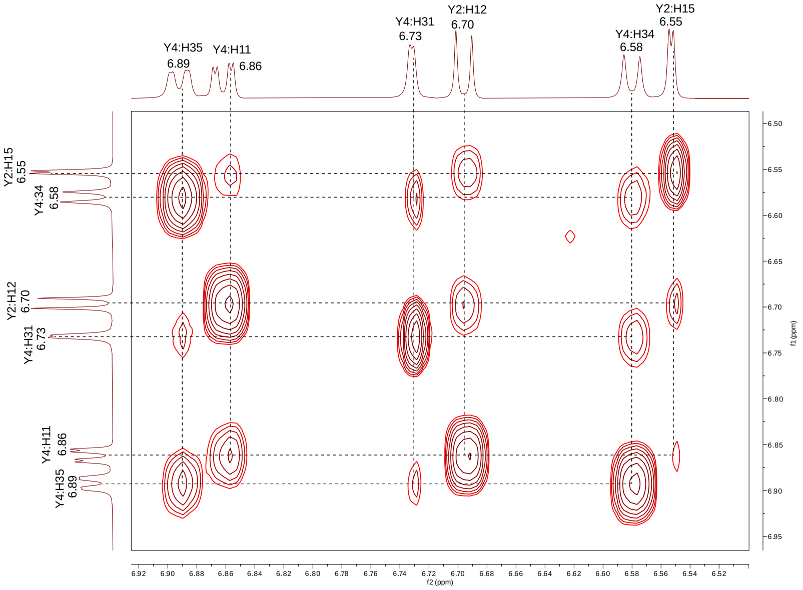

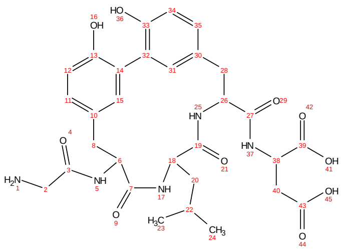

H) Annotated $^{1}$H-$^{1}$H NOESY correlations

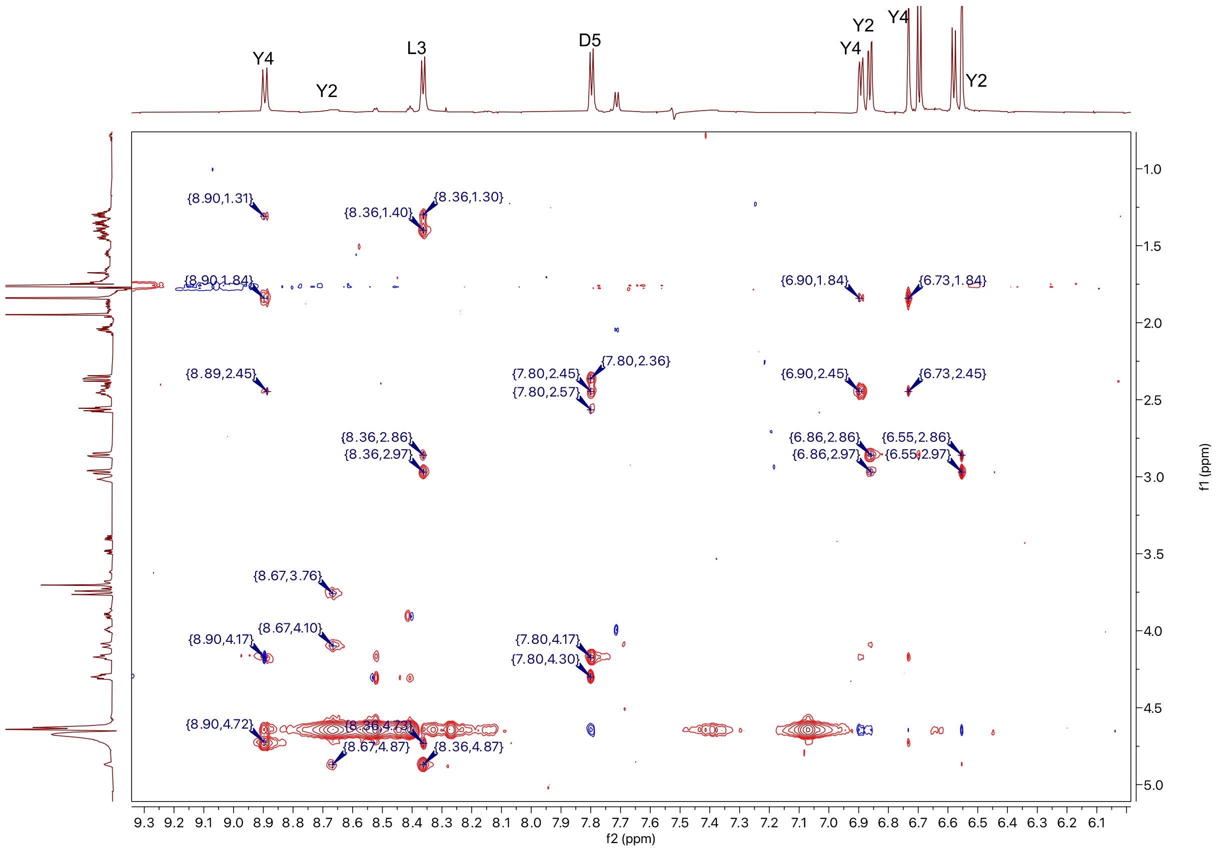

### I) Annotated $^{1}H$-$^{13}C$ HSQC correlations

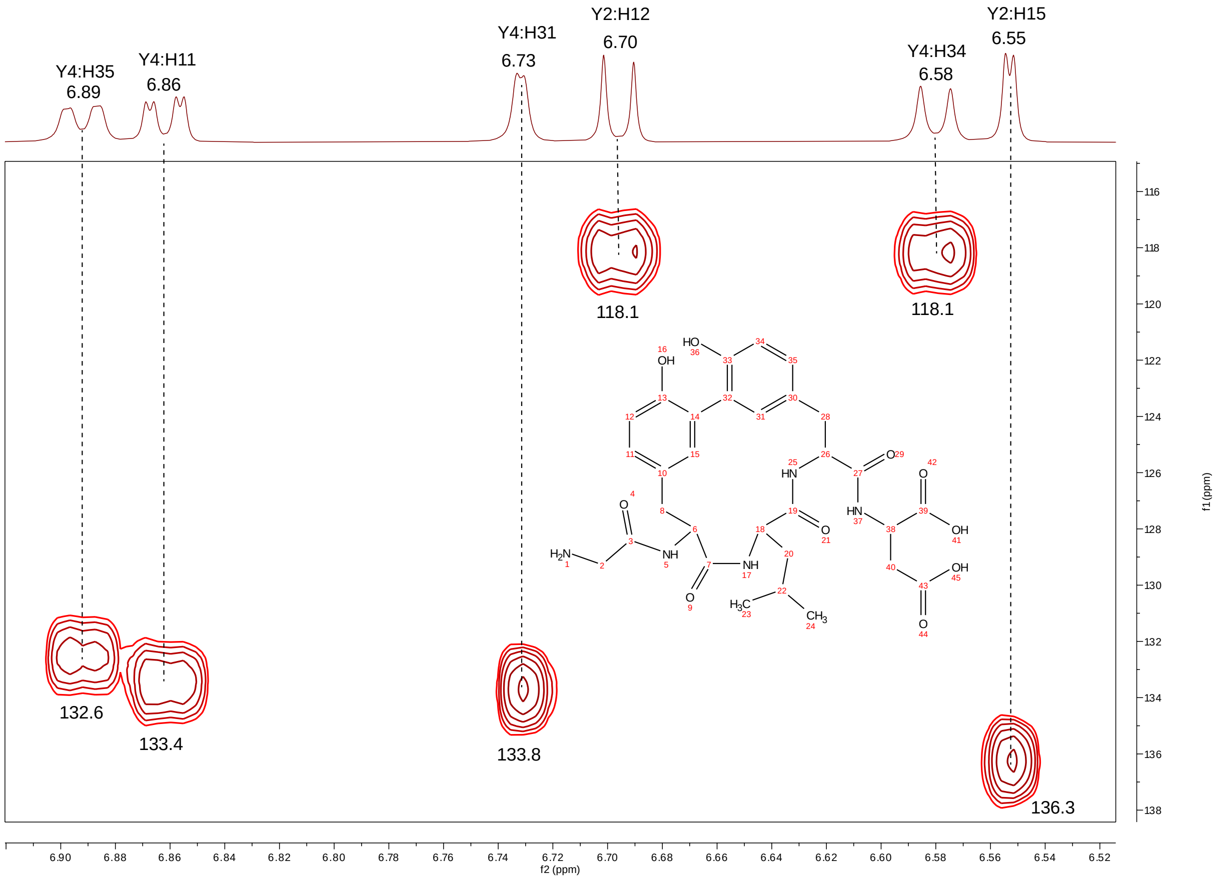

J) Key $^{1}H$-$^{13}C$ HMBC correlations for structural elucidation.

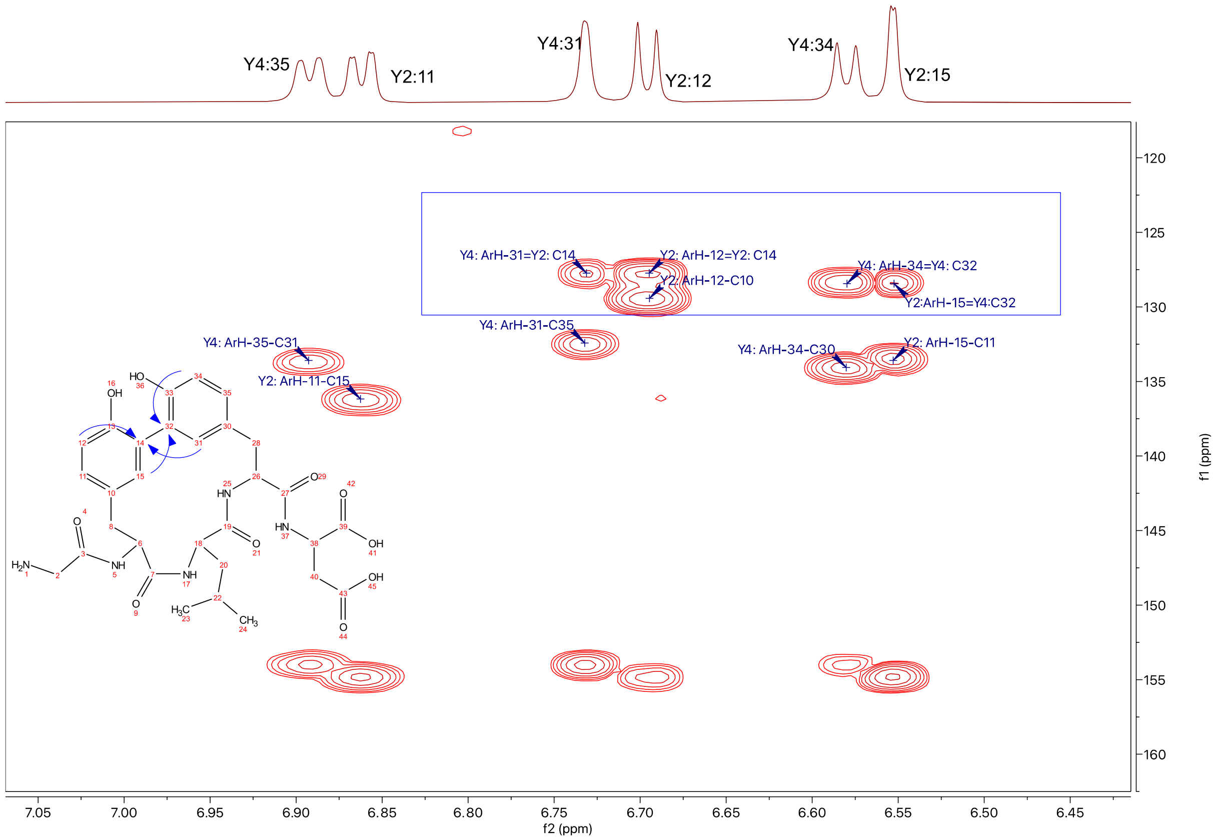

**Table S5:** Summary of $^{1}H$ and $^{13}C$ chemical shifts for the ApyA/O-modified product after GluC cleavage.

| residue number | AA | NH/C=O | αH/C | βH/C | γH/C | δH/C and others |
|---|---|---|---|---|---|---|
| 1 | Gly | 169.2 | 3.75, 3.70<br>43.3 | | | |
| 2 | Tyr | 8.67<br>172.8 | 4.87<br>56.2 | 2.98, 2.87<br>40.2 | 129.6<br>(Aryl C4) | 6.86 (dd), 133.4 (Aryl C5)<br>6.70 (d), 118.1 (Aryl C6)<br>6.55 (d, 2Hz), 136.3 (Aryl C3).<br>154.9 (C-OH, Aryl C1)<br>127.8 (Aryl C2, crosslinked) |
| 3 | Leu | 8.36<br>176.8 | 4.72<br>54.9 | 1.40, 1.30<br>44.1 | 1.36<br>26.9 | 0.68, 24,6<br>0.59, 24.2 |
| 4 | Tyr | 8.90<br>173.8 | 4.17<br>57.2 | 2.45, 1.85<br>40.8 | 134.1<br>(Aryl C4) | 6.89 (dd), 132.6 (Aryl C5)<br>6.58 (d), 118.1 (Aryl C6)<br>6.73(d, 2Hz), 133.8 (Aryl C3)<br>153.9 (C-OH, Aryl C1)<br>128.4 (Aryl C2, crosslinked) |
| 5 | Asp | 7.80<br>181.0 | 4.30<br>56.0 | 2.57, 2.36<br>42.4 | | 181.1 |

**Figure S19:** NMR spectra of ApyA after ApyD/O modification and GluC cleavage. The solvent used was 40% $CD_3CN$/60% $H_2O$. The structure of the product is shown below:

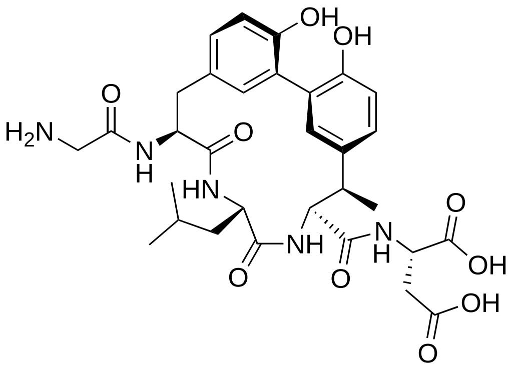

A) $^{1}$H-NMR spectrum

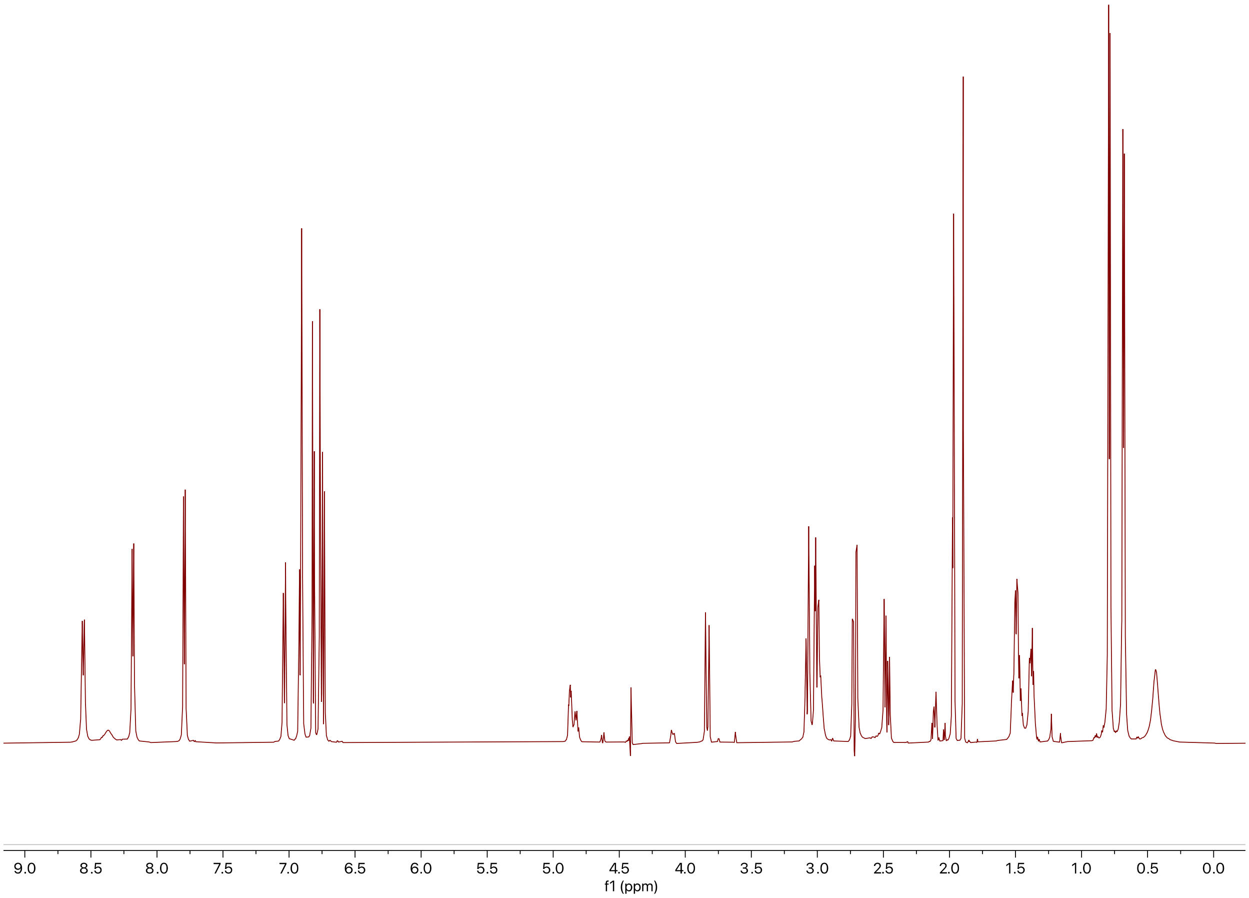

B) $^{1}H$-$^{1}H$ TOCSY

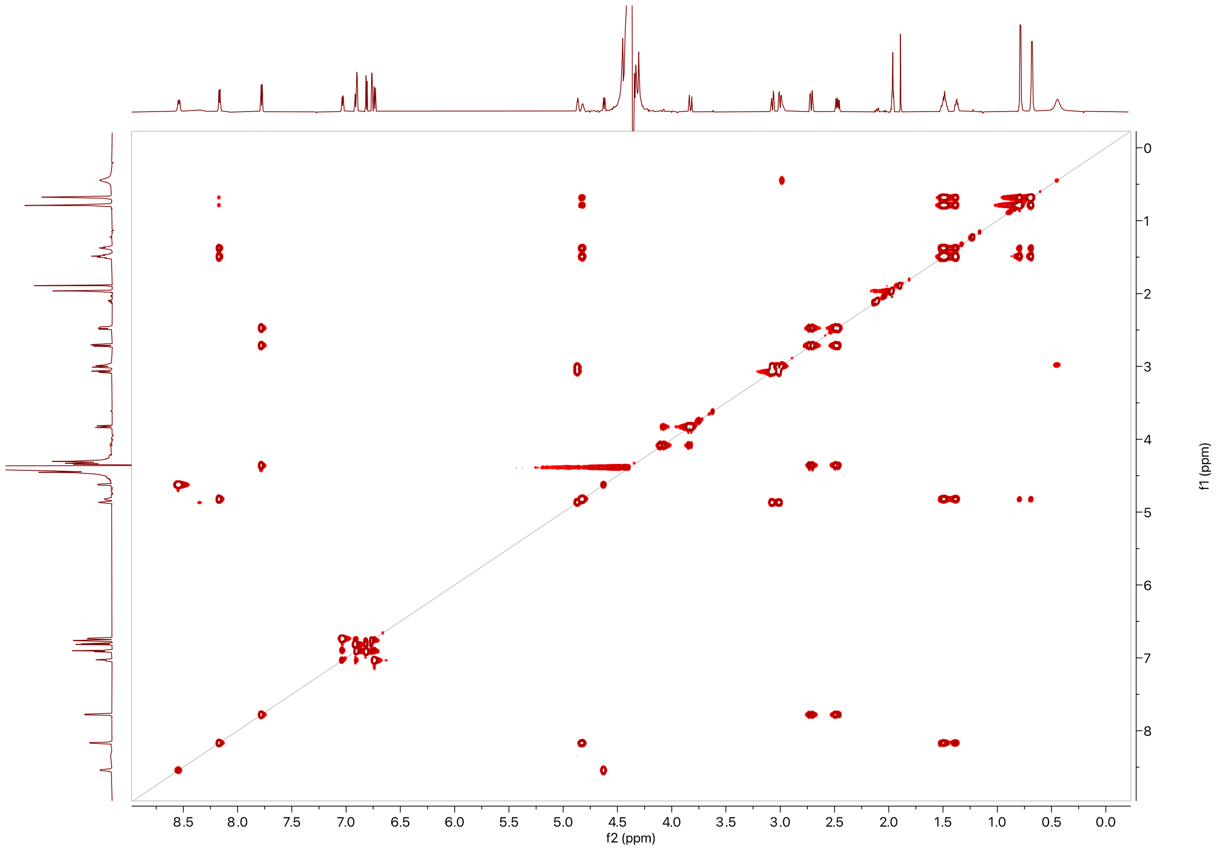

C) $^{1}H$-$^{1}H$ NOESY

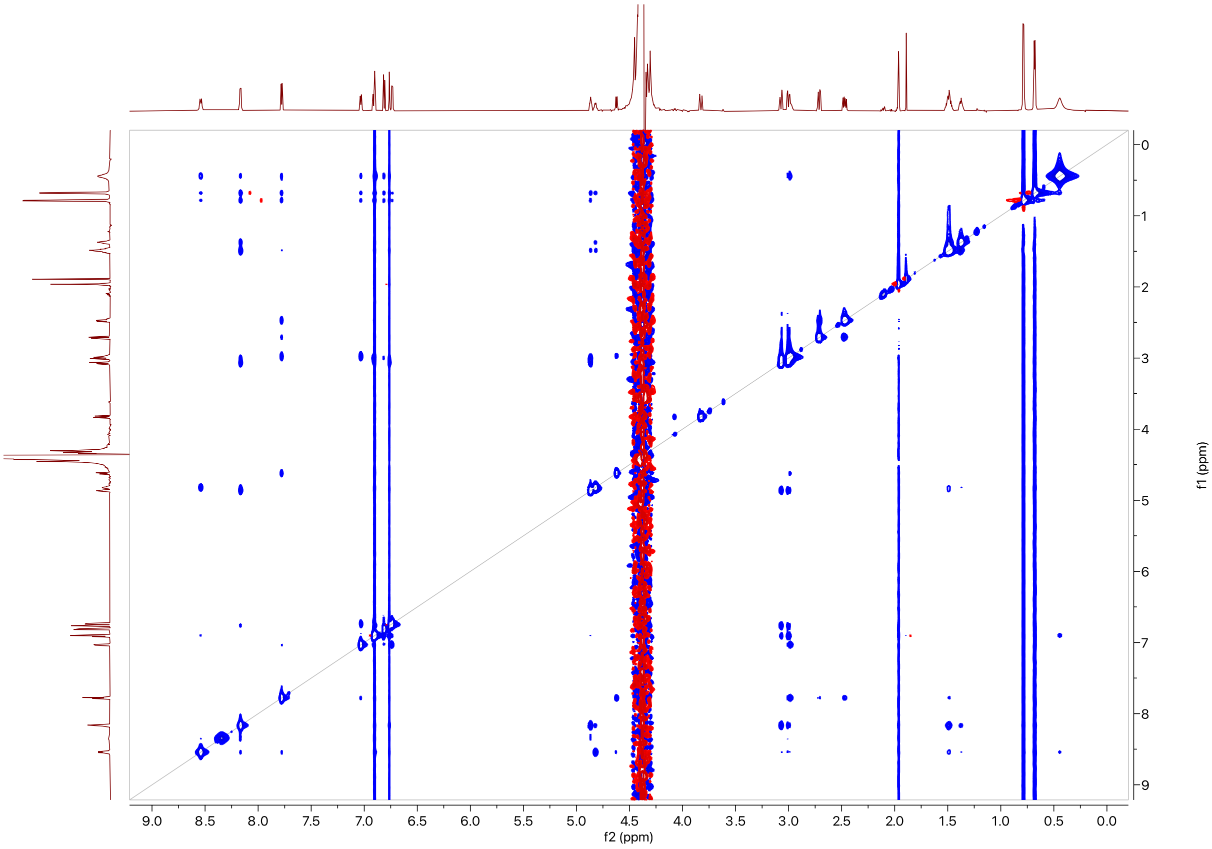

D) $^{1}H$-$^{13}C$ HSQC

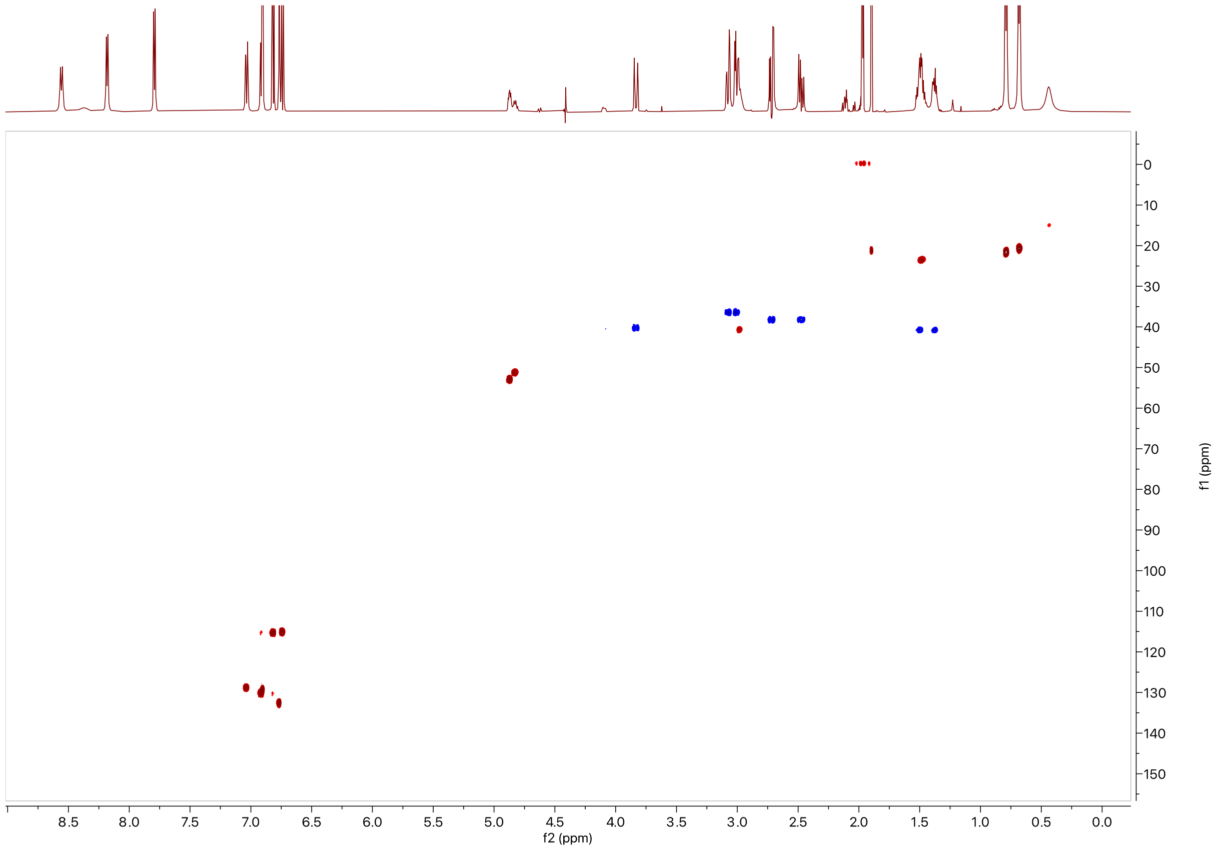

8.5
8.0
7.5
7.0
6.5
6.0
5.5
5.0
4.5
4.0
3.5
3.0
2.5
2.0
1.5
1.0
0.5
0.0
f2 (ppm)
0
10
20
30
40
50
60
70
80
90
100
110
120
130
140
150
f1 (ppm)

E) $^{1}H$-$^{13}C$ HMBC

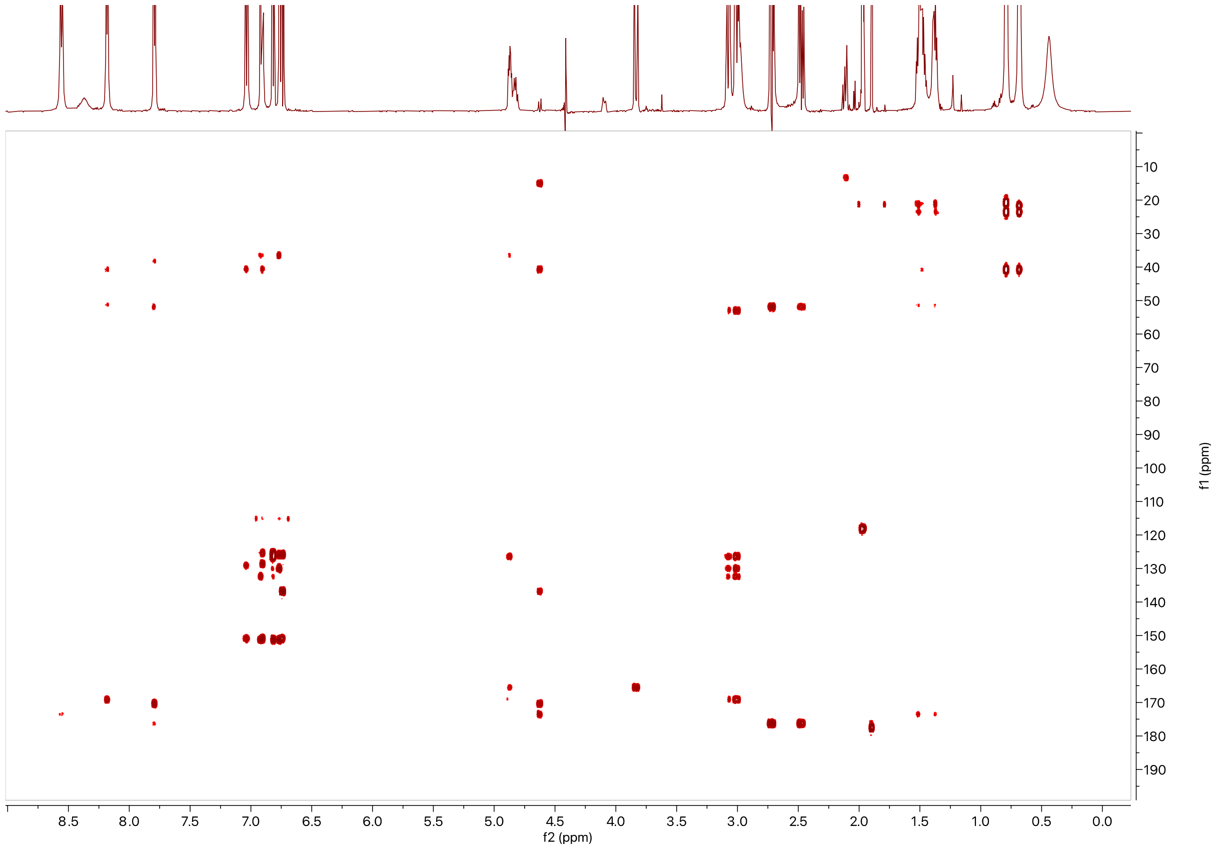

**Table S6:** Summary of $^1$H and $^{13}$C chemical shifts for ApyA after ApyD/O modification and GluC cleavage.

| number | AA | NH/C=O | αH/C | βH/C | γH/C | δH/C and others |
|---|---|---|---|---|---|---|
| 1 | Gly | 167.2 | 4.08, 3.83<br>42.1 | | | |
| 2 | Tyr | 8.35 (br)<br>170.8 | 4.87<br>54.8 | 3.08, 3.01<br>38.2 | 127.1<br>(Aryl C4) | 6.92(dd), 131.8 (Aryl C5)<br>6.82(d), 117.0 (Aryl C6)<br>6.77(d, 2Hz), 134.3 (Aryl C3)<br>153.0 (C-OH, Aryl C1)<br>128.3 (Aryl C2, crosslinked) |
| 3 | Leu | 8.17<br>175.1 | 4.83<br>53.0 | 1.51, 1.38<br>42.6 | 1.51<br>25.3 | 0.79, 23.4<br>0.69, 22.4 |
| 4 | (β–Me)Tyr | 8.54<br>172.1 | 4.63<br>57.6 | 3.00 (br)<br>42.4 | 138.5<br>(Aryl C4) | 0.45 (br), 16.8 (β–methyl)<br>7.04(dd), 130.3 (Aryl C5)<br>6.74(d), 116.8 (Aryl C6)<br>6.91(d, 2Hz), 131.1 (Aryl C3)<br>152.7 (C-OH, Aryl C1)<br>127.6 (Aryl C2, crosslinked) |
| 5 | Asp | 7.78<br>N/A | 4.36<br>53.6 | 2.72, 2.47<br>40.0 | | 178.0 (γ) |

**Figure S20:** Exciton-coupled circular dichroism (ECD) spectra of synthetic standards.

**A** (*R*)-(+)-5,5′,6,6′,7,7′,8,8′-Octahydro-1,1′-bi-2-naphthol

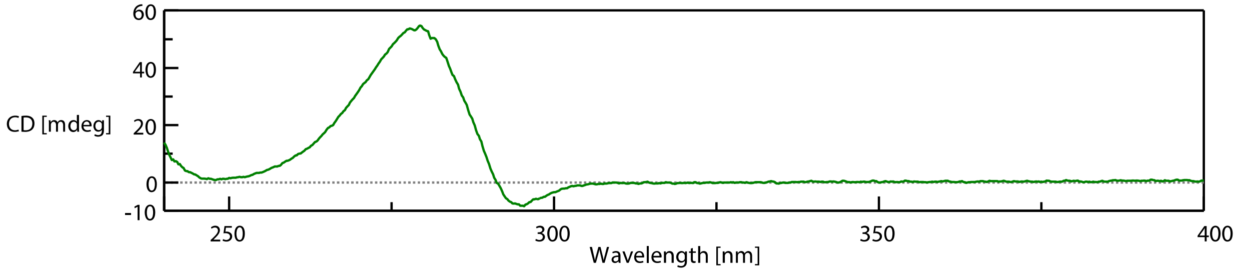

**B** (*S*)-(-)-5,5′,6,6′,7,7′,8,8′-Octahydro-1,1′-bi-2-naphthol

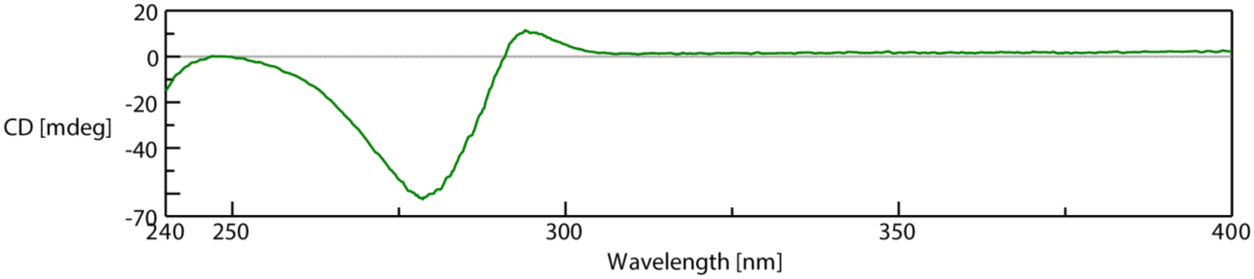

**C** (*R*)-(+)-5,5',6,6'-Tetramethyl-3,3'-di-t-butyl-1,1'-biphenyl-2,2'-diol

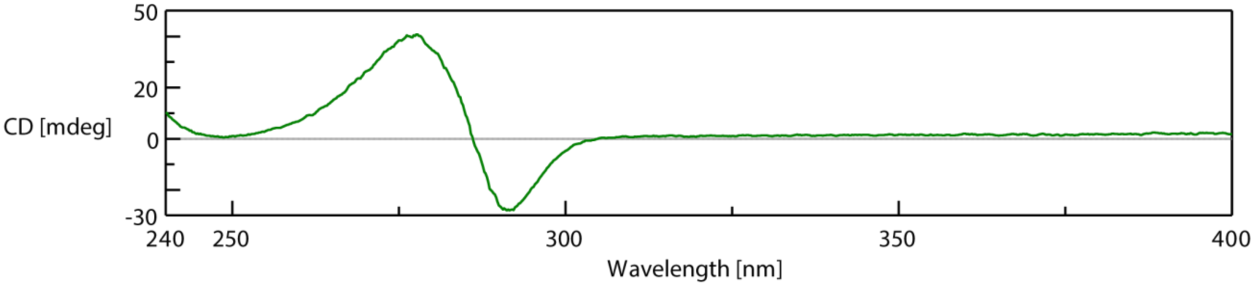

**D** (*S*)-(-)-5,5',6,6'-Tetramethyl-3,3'-di-t-butyl-1,1'-biphenyl-2,2'-diol

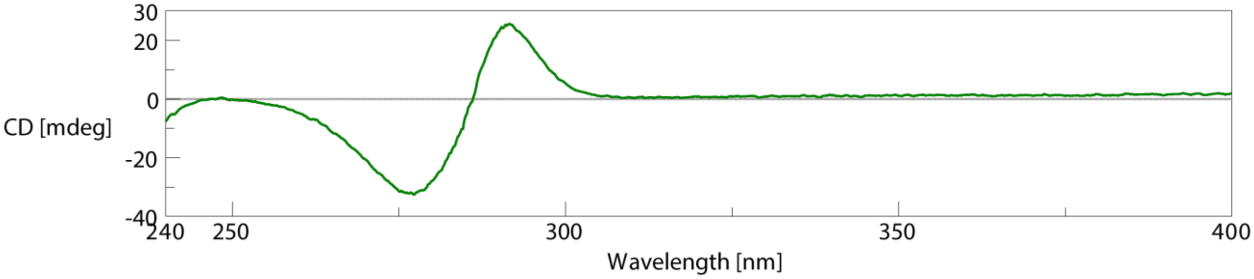

**Figure S21:** ECD spectrum (top) and UV spectrum (bottom) of the GluC-cleaved, ApyA modified by ApyO. Abs: absorbance.

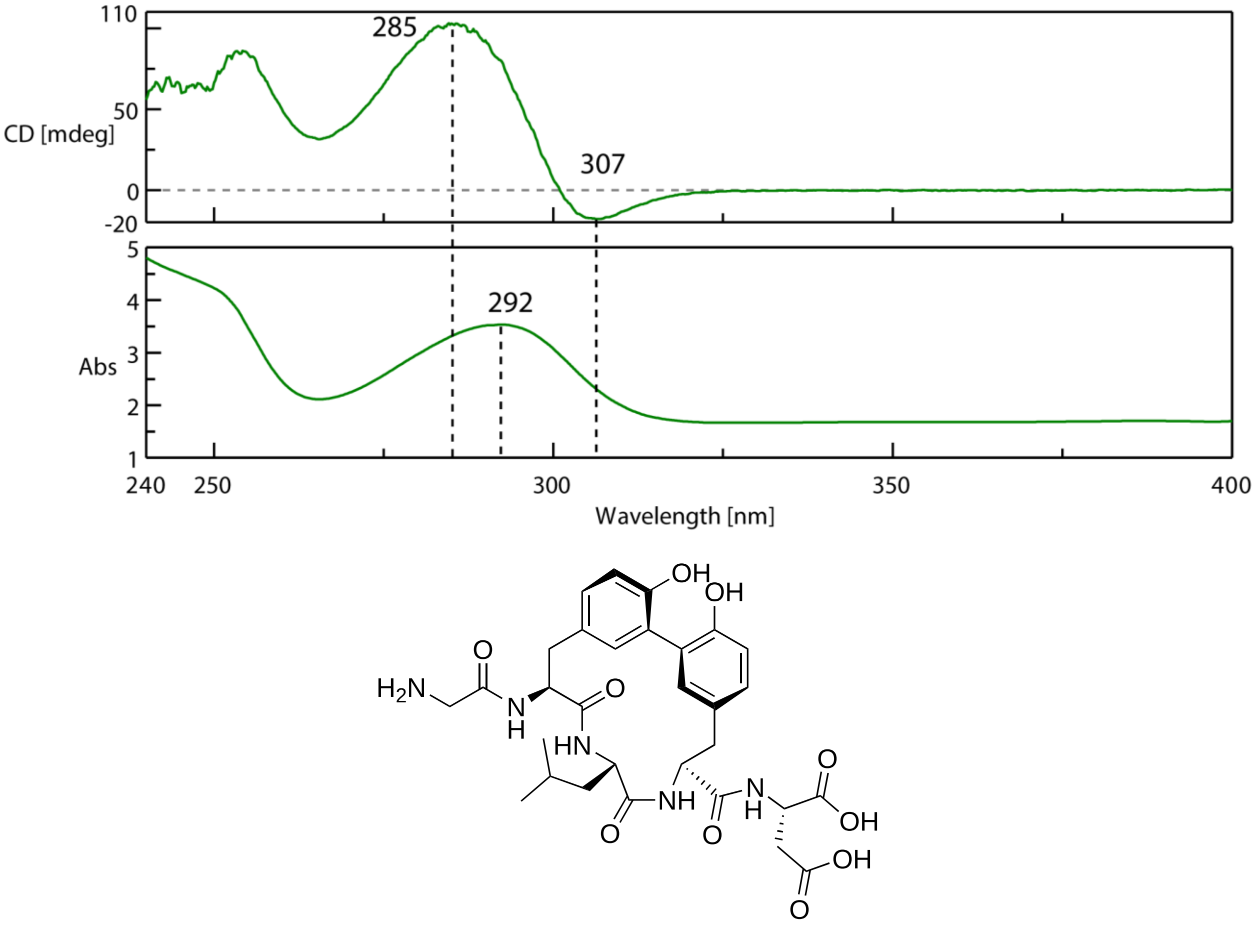

**Figure S22:** ECD spectrum (top) and UV spectrum (bottom) of the GluC-cleaved ApyA modified by ApyD and ApyA modified by ApyD and ApyO. (A) Spectra of the the GluC-cleaved ApyA modified by ApyD, which does not contain the biaryl crosslink. (B) Spectra of the GluC-cleaved ApyA modified by ApyD and ApyO. The structure of each product can be found in Figure 2 (Main Text). Abs: absorbance

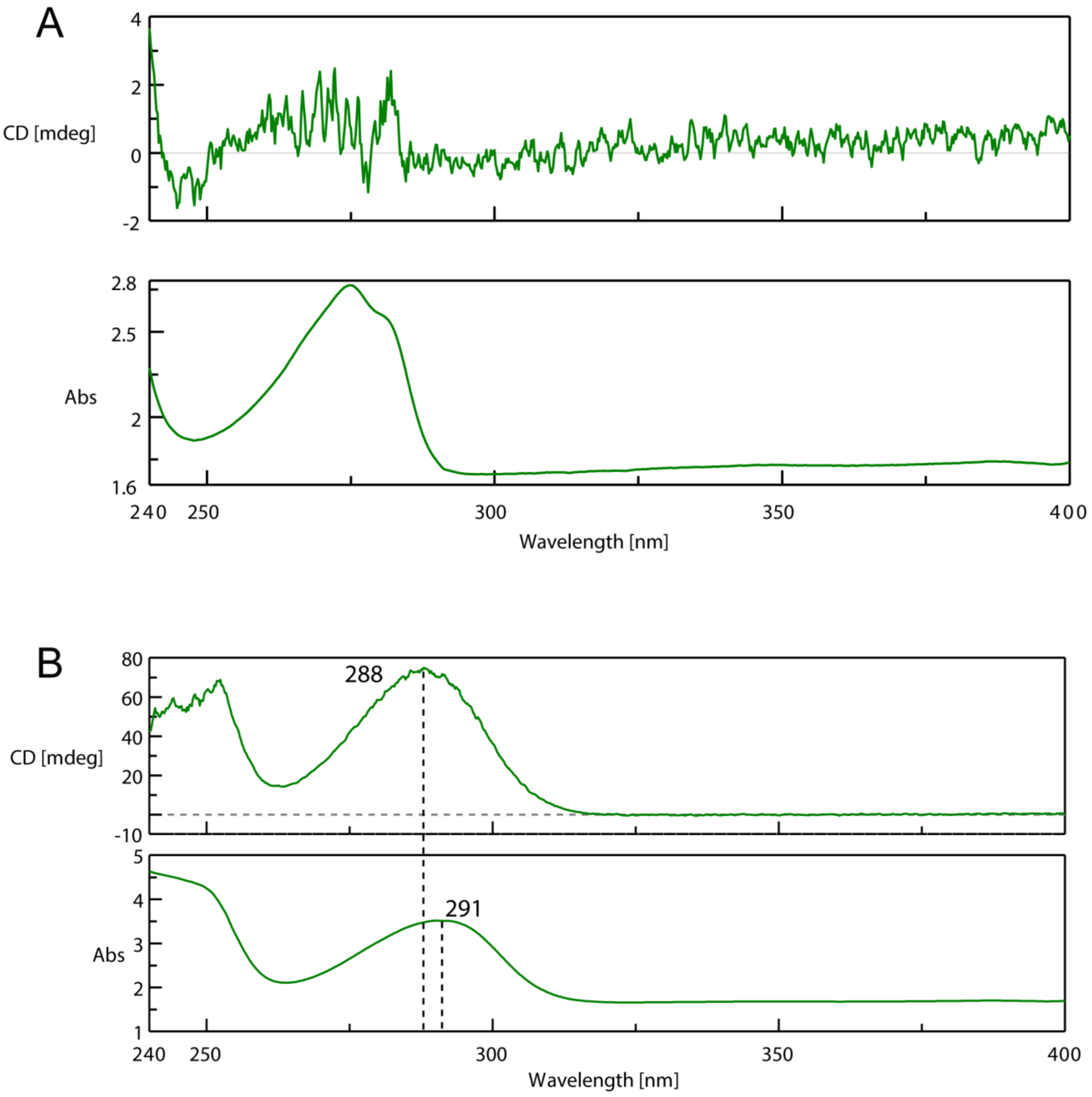

**Figure S23:** Fully refined MicroED structure of ApyD/O-modified ApyA pentapeptide with an overlaid density map ($F_{observed}$). Oxygen = Red, Nitrogen = Blue, Carbon = Grey, Hydrogen = White, and Zinc = Pink.

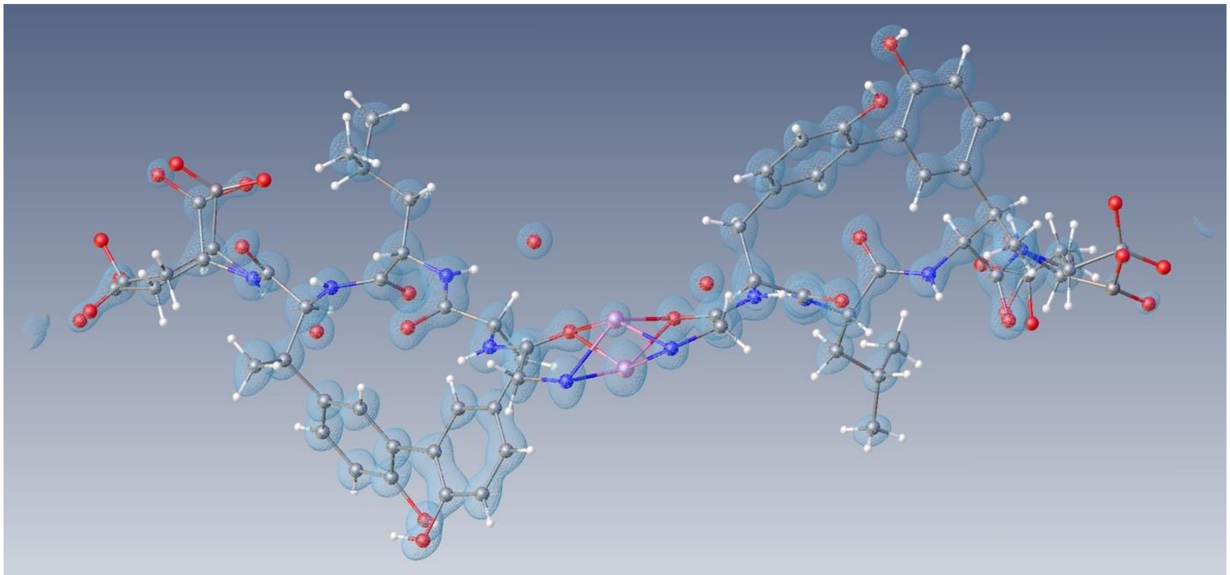

**Table S7:** Crystallographic data and structure refinements for the ApyA pentapeptide modified by ApyD and ApyO. The data analysis and refinement process can be found in the supplementary methods. The structure was deposited under CCDC ID 2324739.

| | | |
|---|---|---|
| Empirical formula | C62 H70 N10 O23 Zn | |
| Formula weight | 1388.68 | |
| Temperature | 77 K | |
| Wavelength | 0.02805 ≈ | |
| Crystal system | Orthorhombic | |
| Space group | $P2_12_12_1$ | |
| Unit cell dimensions | a = 11.130(2) ≈ | $\alpha$= 90∞. |
| | b = 27.400(6) ≈ | $\beta$= 90∞. |
| | c = 33.980(13) ≈ | $\gamma$ = 90∞. |
| Volume | 10363(5) $≈^3$ | |
| Z | 4 | |
| Density (calculated) | 0.890 $Mg/m^3$ | |
| Absorption coefficient | 0.000 $mm^{-1}$ | |
| F(000) | 1065 | |
| Crystal size | ? x ? x ? $mm^3$ ; not determined | |
| Theta range for data collection | 0.038 to 0.803∞. | |
| Index ranges | -11<=h<=11, -27<=k<=27, -29<=l<=28 | |
| Reflections collected | 51679 | |
| Independent reflections | 9167 [R(int) = 0.1730] | |
| Completeness to theta = 0.803∞ | 83.5 % | |
| Absorption correction | Semi-empirical from equivalents | |
| Refinement method | Full-matrix least-squares on $F^2$ | |
| Data / restraints / parameters | 9167 / 2166 / 969 | |
| Goodness-of-fit on $F^2$ | 0.913 | |
| Final R indices [I>2sigma(I)] | R1 = 0.0819, wR2 = 0.1988 | |
| R indices (all data) | R1 = 0.1171, wR2 = 0.2159 | |
| Absolute structure parameter | ? | |
| Extinction coefficient | 7(2) | |
| Largest diff. peak and hole | 0.126 and -0.078 e.$≈^{-3}$ | |

**Figure S24:** Deconvoluted HR-ESI-MS of ApyA co-expressed with all modifying enzymes except ApyO (pSCrhaB2_ApyAHIDS) in *Burkholderia* sp. FERM BP-3421. The sequence of the construct can be found in Table S2C. The sequence of the peptide is shown below with the cut site bolded (endoproteinase GluC was used). This expression was performed using similar conditions where ApyD had low activity (see Figure S11 and supplementary methods), showing that the removal of ApyO abolished ApyHI activity but enhanced ApyD activity. Calculated exact mass for GluC-cleaved ApyA: 629.2696 Da. Calculated exact mass for GluC-cleaved ApyD-modified ApyA: 643.2852 Da. A table with observed masses and the error compared to calculated values can be found in the Supplementary Dataset 3 (Excel).

MHHHHHHMATKPKKTKGVSVSIEGKLPKMTLDMPVDAKKIKAIQKCLENGKLTITMSKVDLAGG
RMG**E**GYLYD

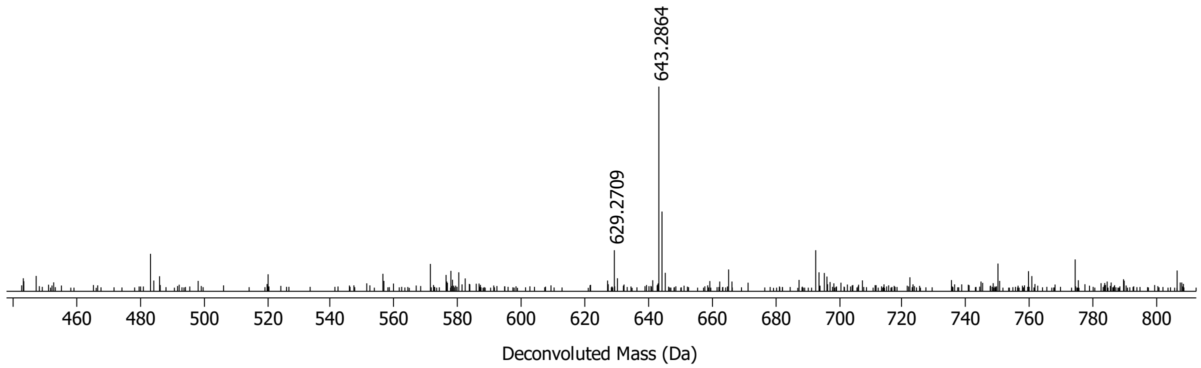

**Figure S25:** HR-ESI-MS/MS of ApyA co-expressed with all modifying enzymes except ApyO (pSCrhaB2_ApyAHIDS) in *Burkholderia* sp. FERM BP-3421. The sequence of the construct can be found in Table S2C. This expression is done using similar conditions where ApyD has low activity (see Figure S11 and supplementary methods), showing that the removal of ApyO abolished ApyHI activity but enhanced ApyD activity. A table with observed masses and the error compared to calculated values can be found in the Supplementary Dataset 3 (Excel).

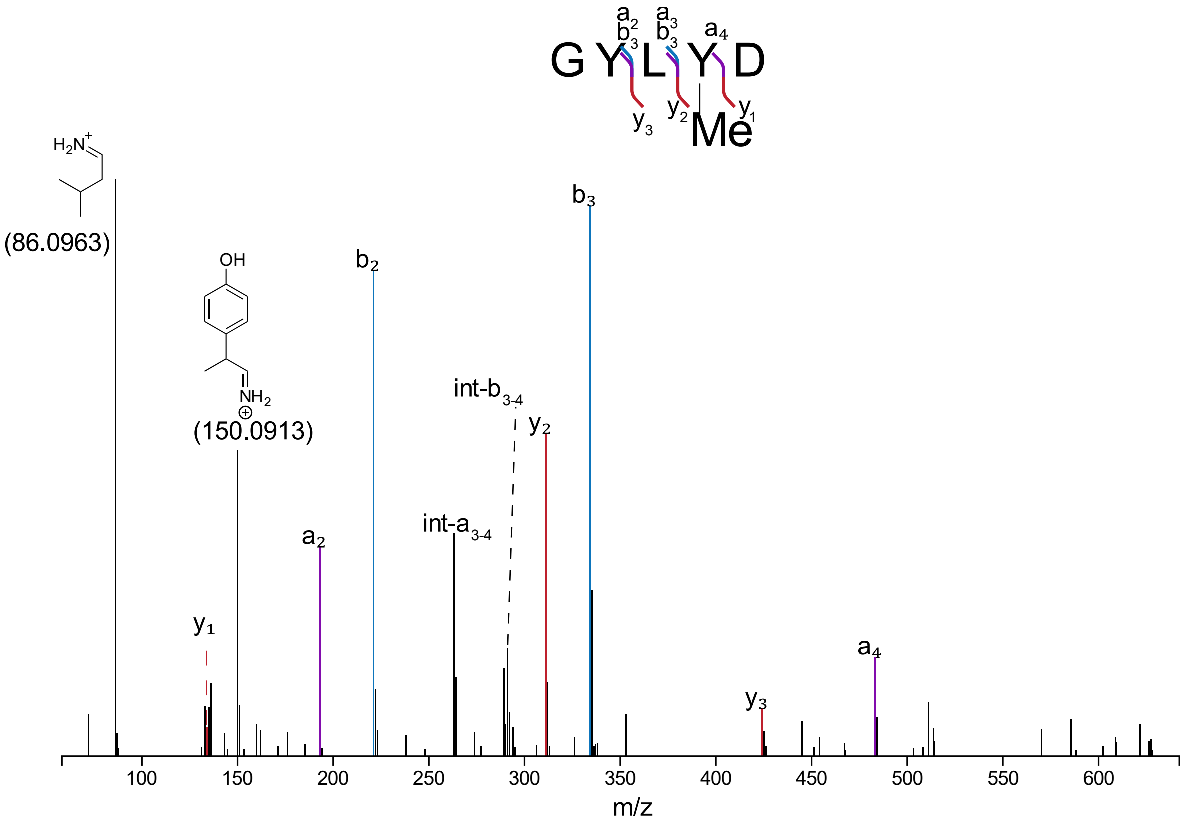

**Figure S26:** NMR spectra of ApyA modified by ApyO/H/I/S after GluC cleavage. The solvent was 10% $D_2O$/ 90% $H_2O$. The structure of the products and key NMR depiction are depicted below and in Figure 4 (main text). The NMR data depict the ketone and the gem-diol form of the peptide.

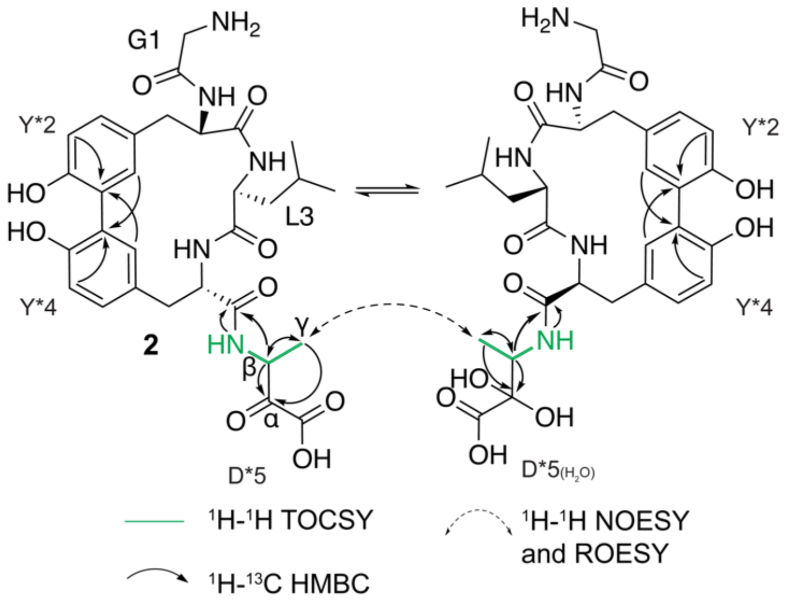

A) $^{1}$H NMR

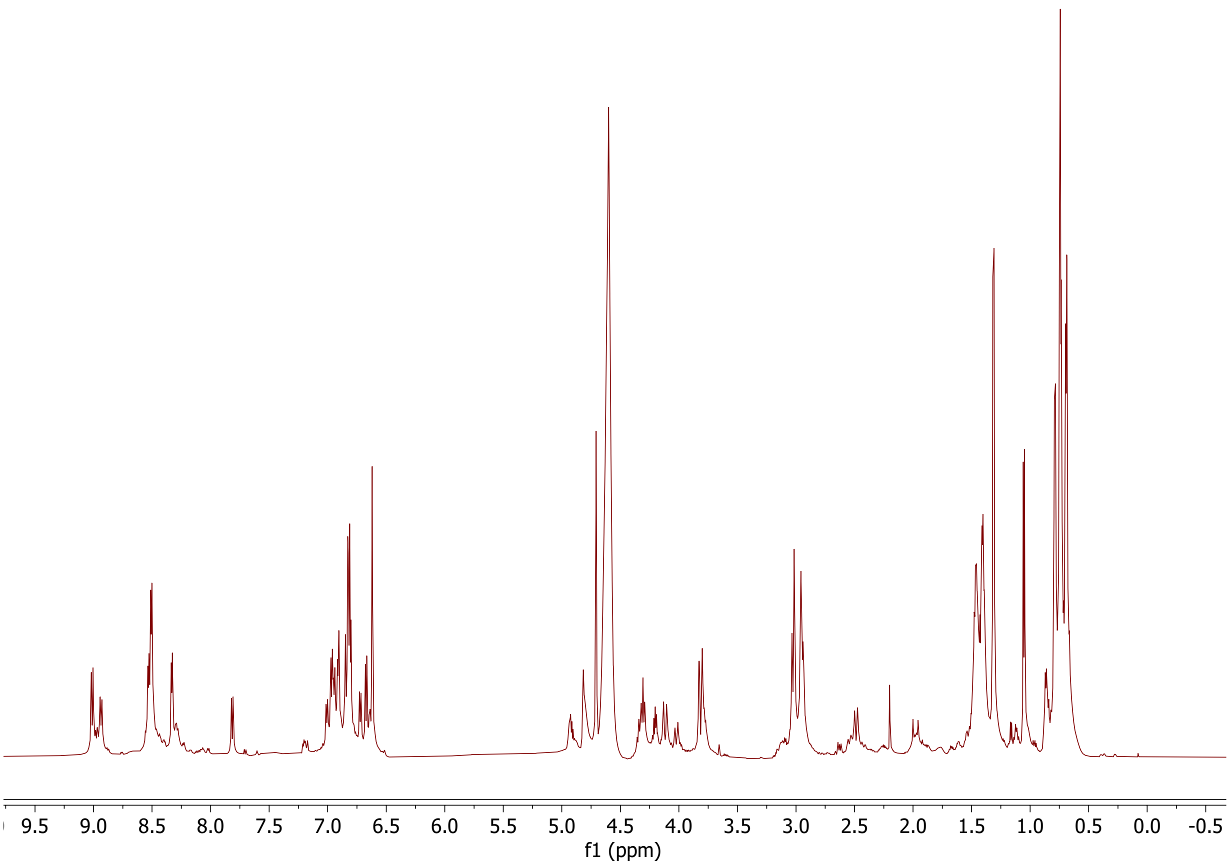

B) $^{1}H$-$^{1}H$ TOCSY

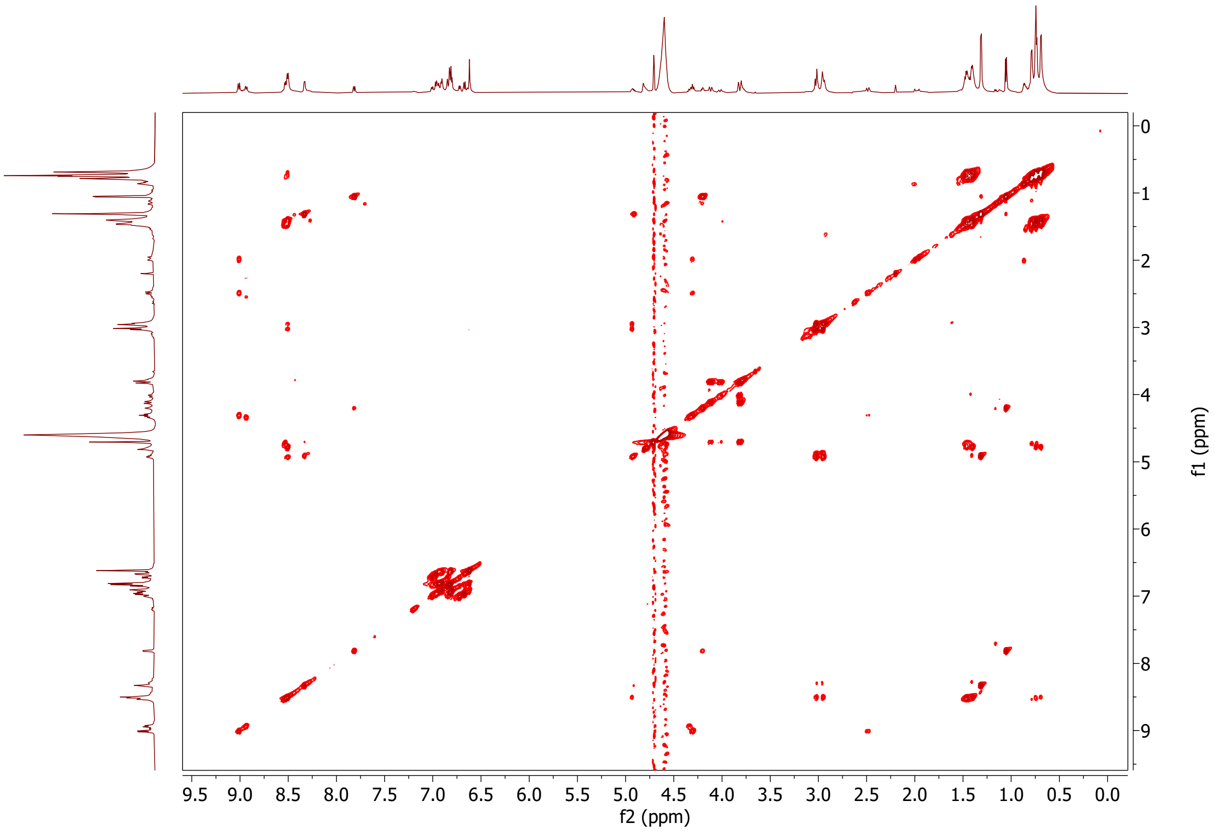

C) $^{1}H$-$^{1}H$ NOESY

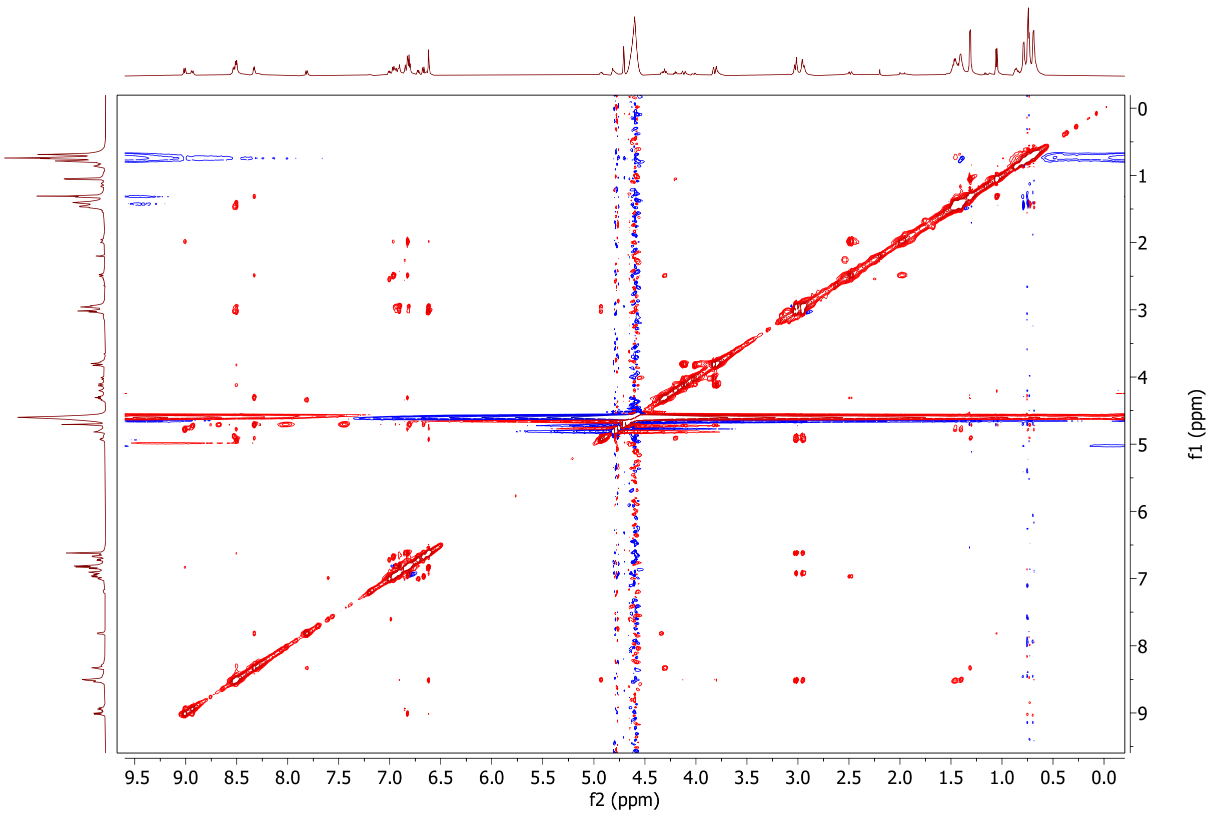

f1 (ppm)
f2 (ppm)
9.5 9.0 8.5 8.0 7.5 7.0 6.5 6.0 5.5 5.0 4.5 4.0 3.5 3.0 2.5 2.0 1.5 1.0 0.5 0.0
0 1 2 3 4 5 6 7 8 9

D) $^1H$-$^1H$ ROESY

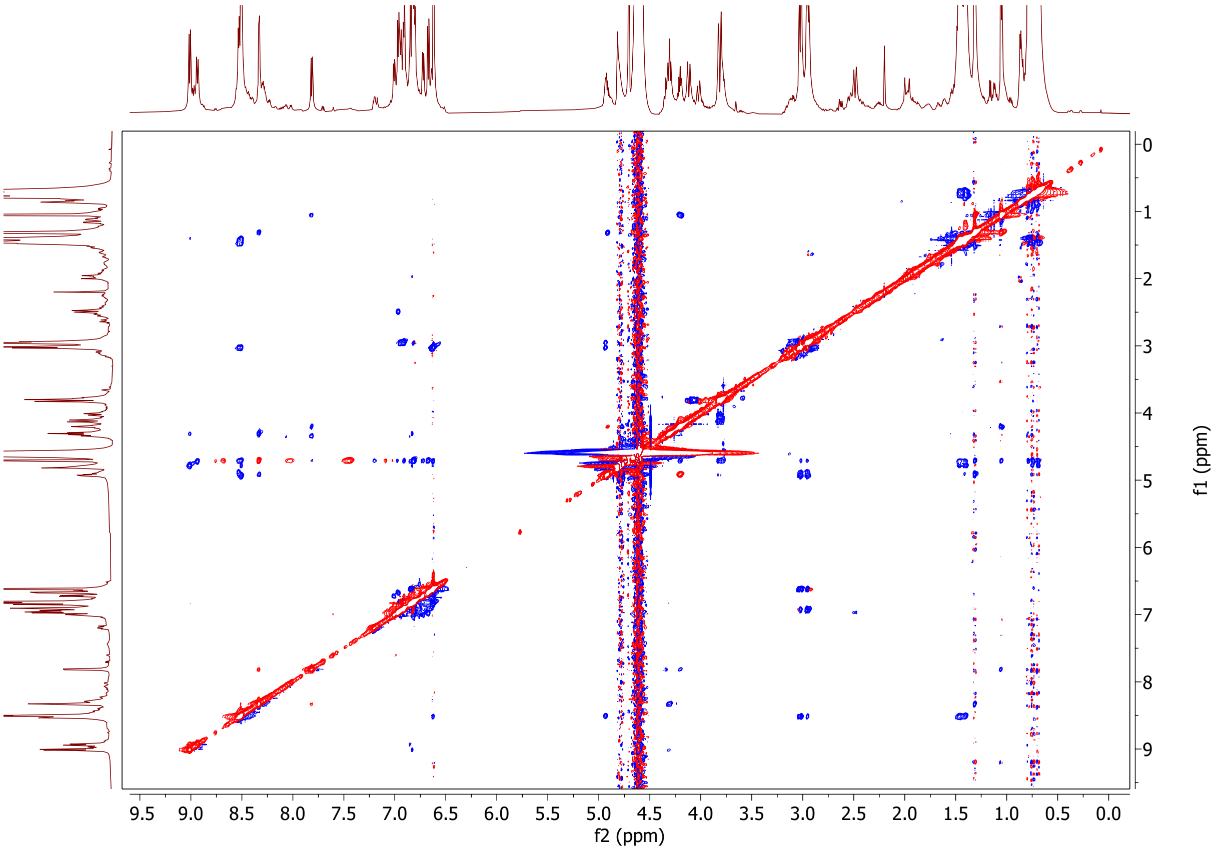

9.5 9.0 8.5 8.0 7.5 7.0 6.5 6.0 5.5 5.0 4.5 4.0 3.5 3.0 2.5 2.0 1.5 1.0 0.5 0.0
f2 (ppm)
0 1 2 3 4 5 6 7 8 9
f1 (ppm)

E) $^{1}H$-$^{13}C$ HSQC

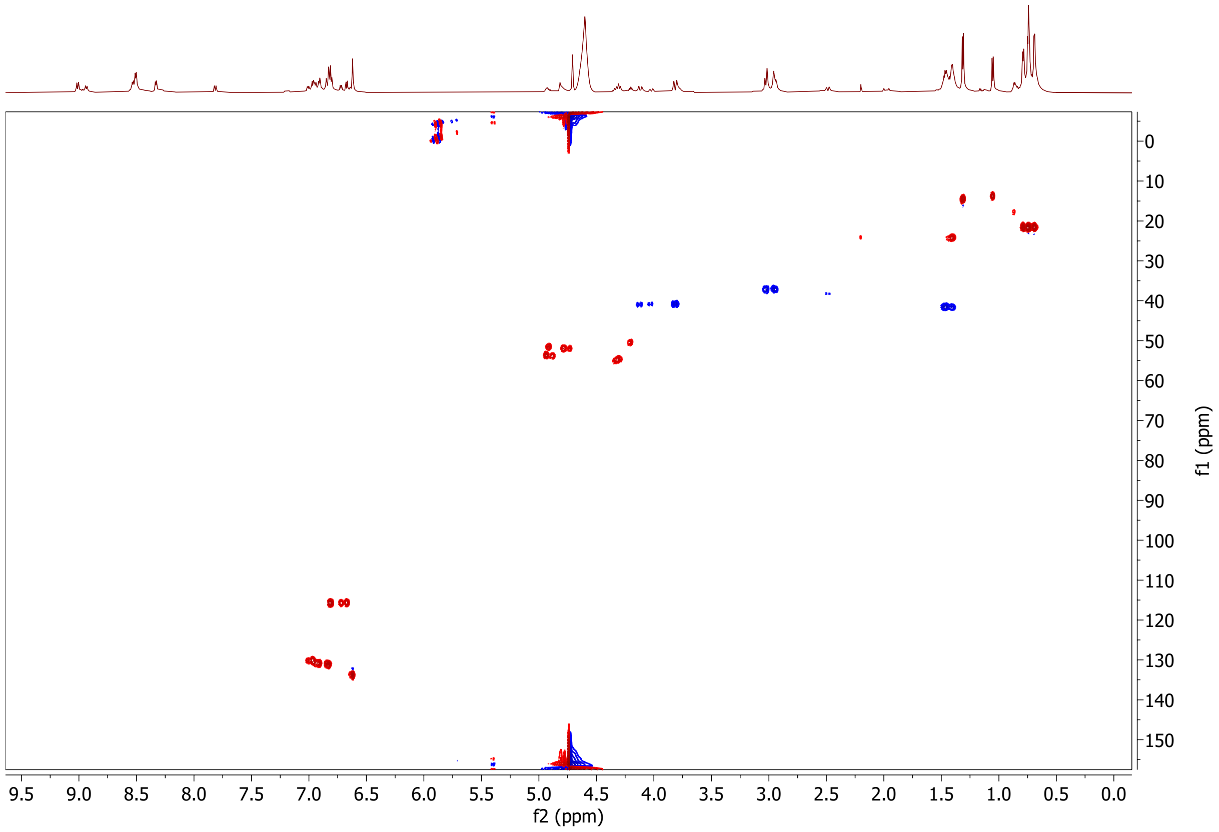

0
10
20
30
40
50
60
70
80
90
100
110
120
130
140
150
f1 (ppm)
9.5 9.0 8.5 8.0 7.5 7.0 6.5 6.0 5.5 5.0 4.5 4.0 3.5 3.0 2.5 2.0 1.5 1.0 0.5 0.0
f2 (ppm)

F) $^{1}H$-$^{13}C$ HMBC

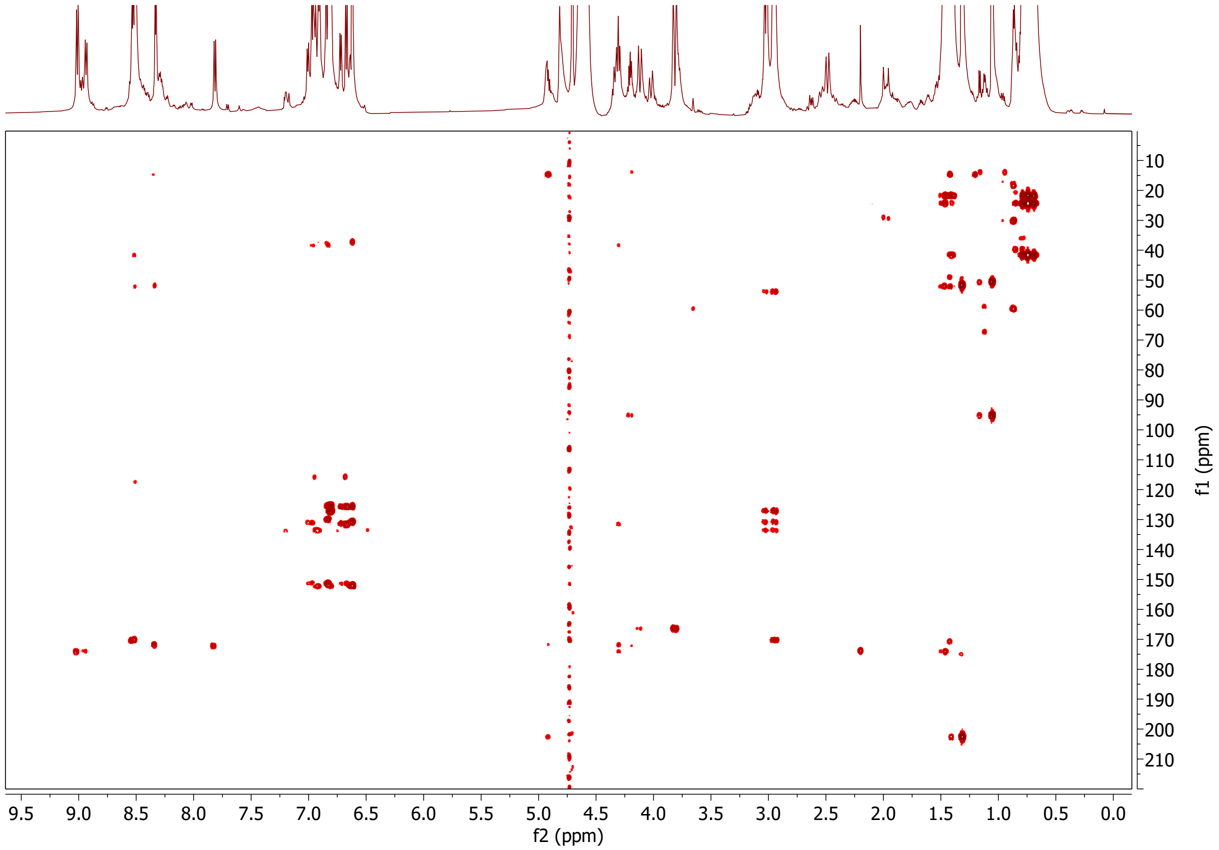

9.5 9.0 8.5 8.0 7.5 7.0 6.5 6.0 5.5 5.0 4.5 4.0 3.5 3.0 2.5 2.0 1.5 1.0 0.5 0.0
f2 (ppm)
10 20 30 40 50 60 70 80 90 100 110 120 130 140 150 160 170 180 190 200 210
f1 (ppm)

G) $^{1}H$-$^{13}C$ HSQC assignment of the aromatic region.

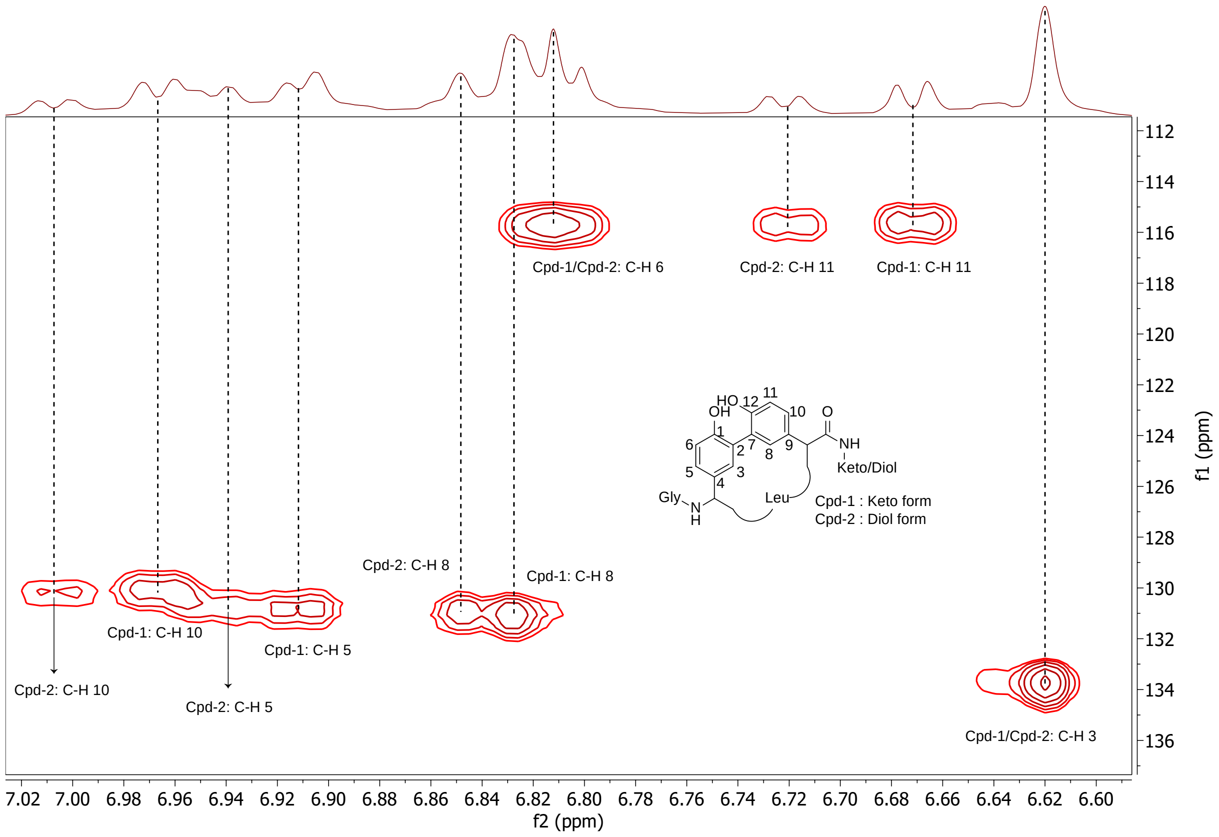

H) $^{1}$H-$^{13}$C HMBC assignment of the aromatic region.

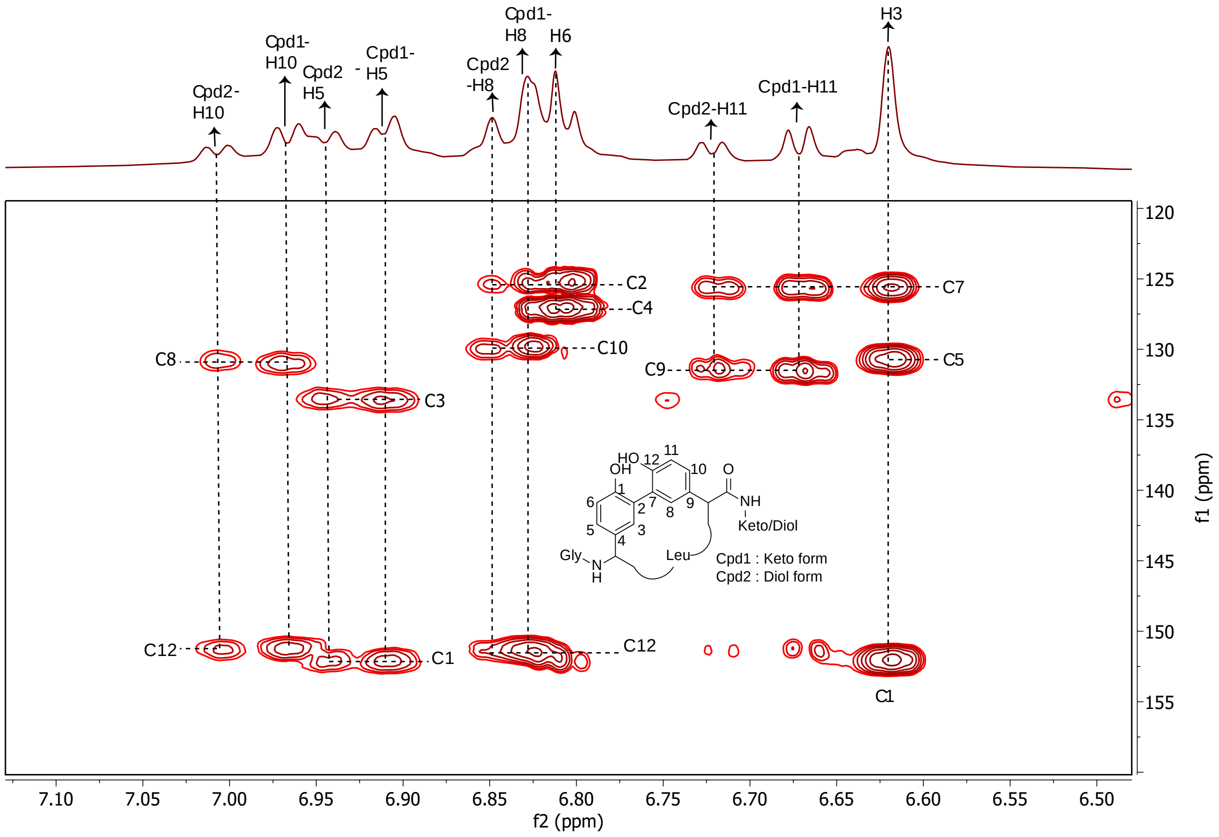

Observed cross peaks:

| Hydrogen | Carbon | | |
|---|---|---|---|
| Cpd1/Cpd2-H3 | C7 | C5 | C1 |
| Cpd1-H11 | C7 | C9 | |
| Cdp2-H11 | C7 | C9 | |
| Cpd1/Cpd2-H6 | C2 | C4 | |
| Cpd1-H8 | C2 | C10 | C12 |
| Cpd2-H8 | C2 | C10 | C12 |
| Cpd1-H5 | C3 | C10 | |
| Cpd2-H5 | C3 | C10 | |
| Cpd1-H10 | C8 | C12 | |
| Cpd2-H10 | C8 | C12 | |

I) $^{1}H$-$^{1}H$ TOCSY and $^{1}H$-$^{13}C$ HMBC of the keto form of the product. Other key cross peaks are in Figure 5 (main text).

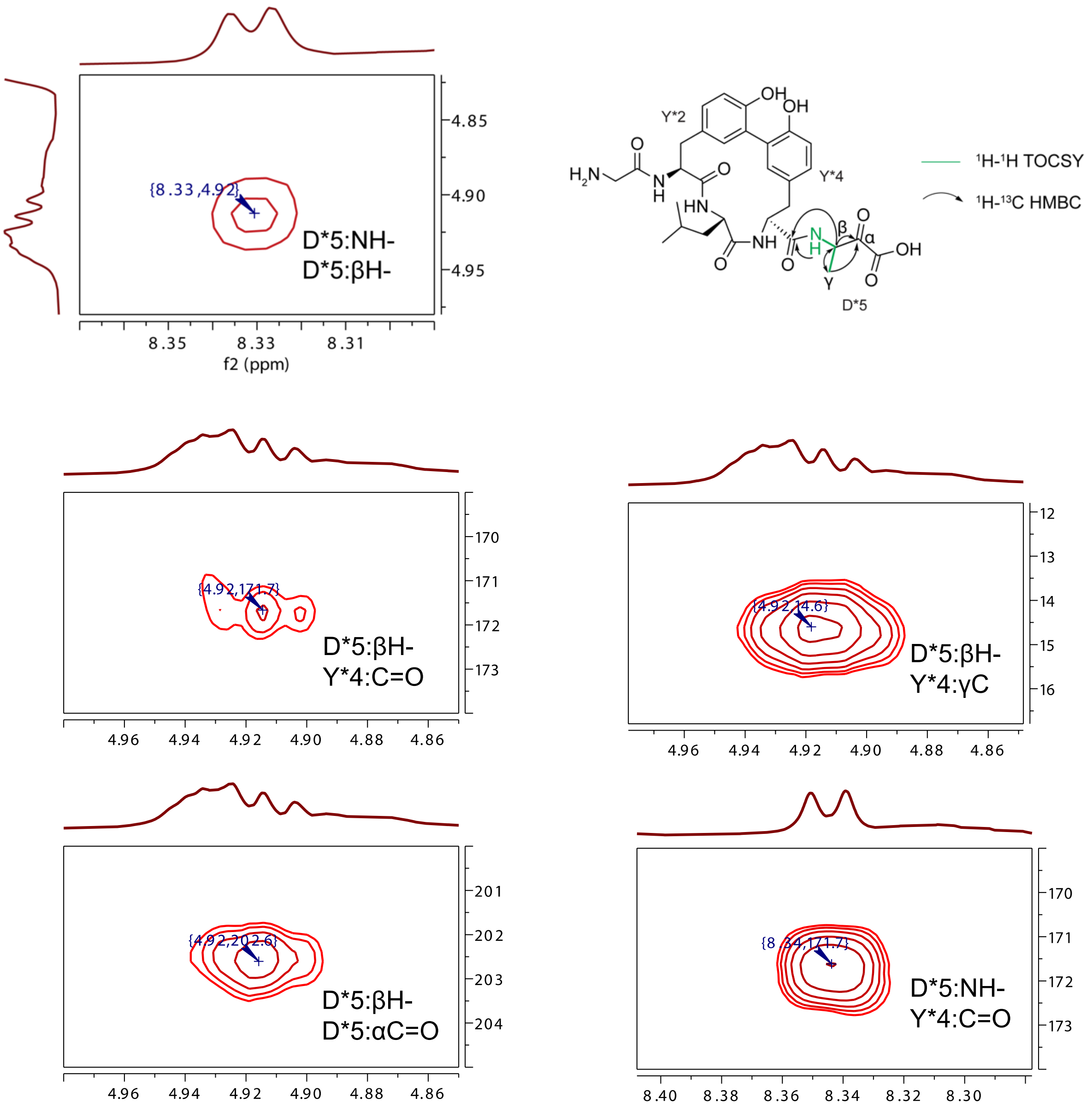

J) $^1H$-$^1H$ TOCSY and $^1H$-$^{13}C$ HMBC correlations of D*5:γH of the gem-diol form of the product. Figure 5 (Main text) depicts similar correlations of the keto form.

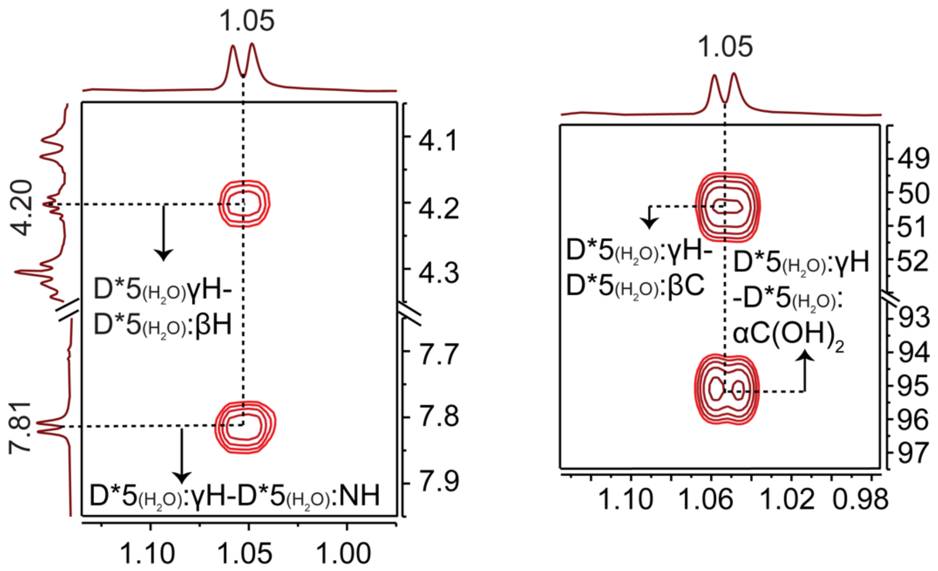

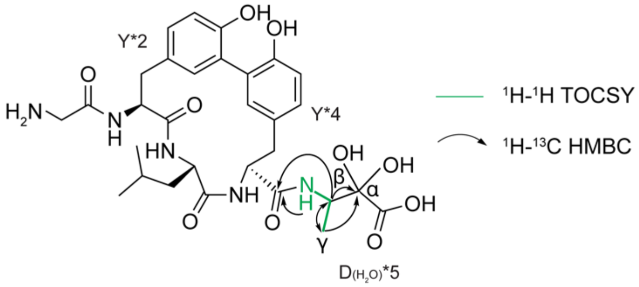

K) $^{1}H$-$^{1}H$ TOCSY and $^{1}H$-$^{13}C$ HMBC correlations of the D*5:βH and D*5:NH of the gem-diol form of the product

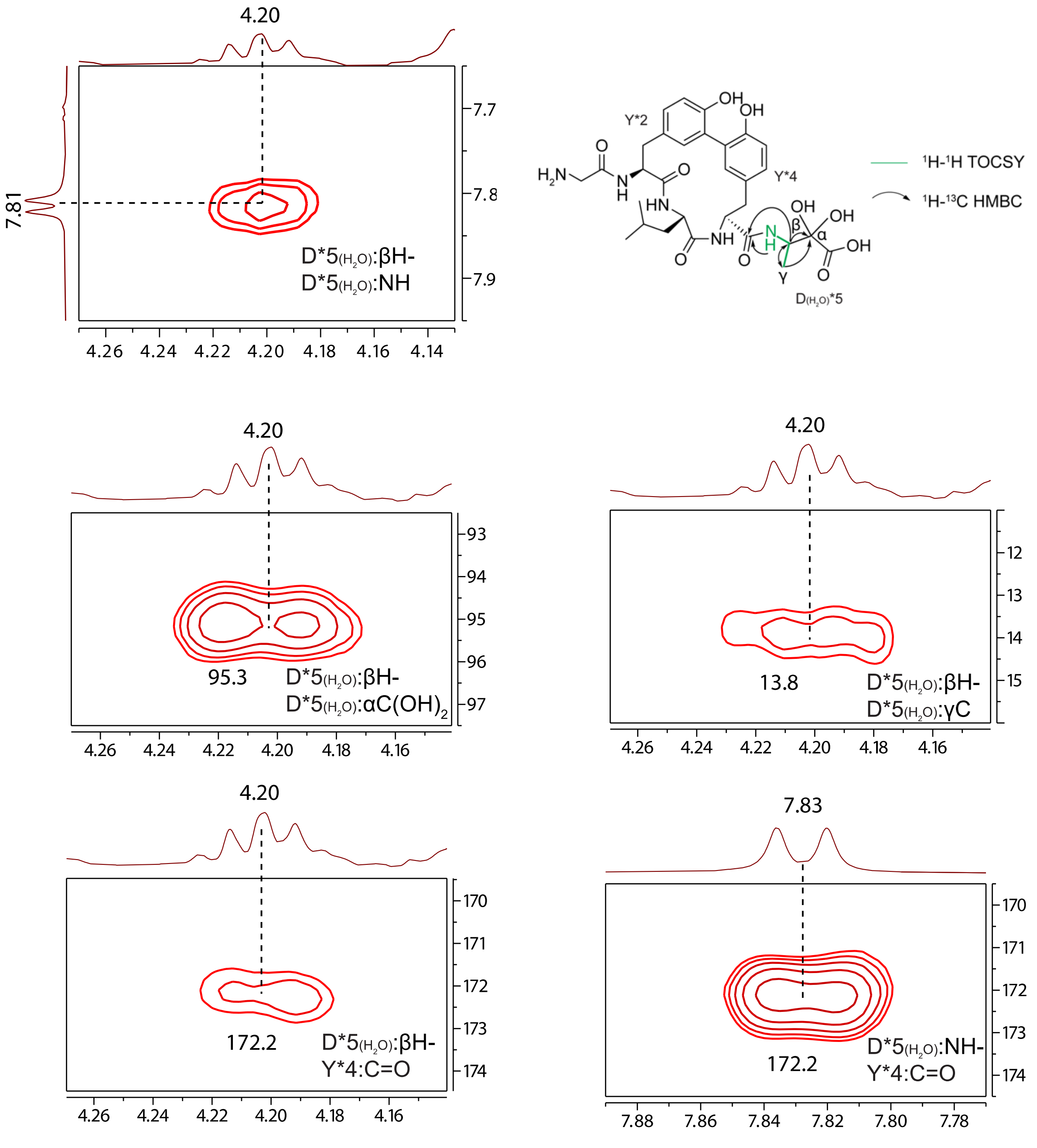

L) $^{1}H$-$^{1}H$ NOESY and $^{1}H$-$^{1}H$ ROESY cross peak between two methyl groups in the keto and the gem-diol form. These cross peaks suggested that the two species undergo chemical exchange.

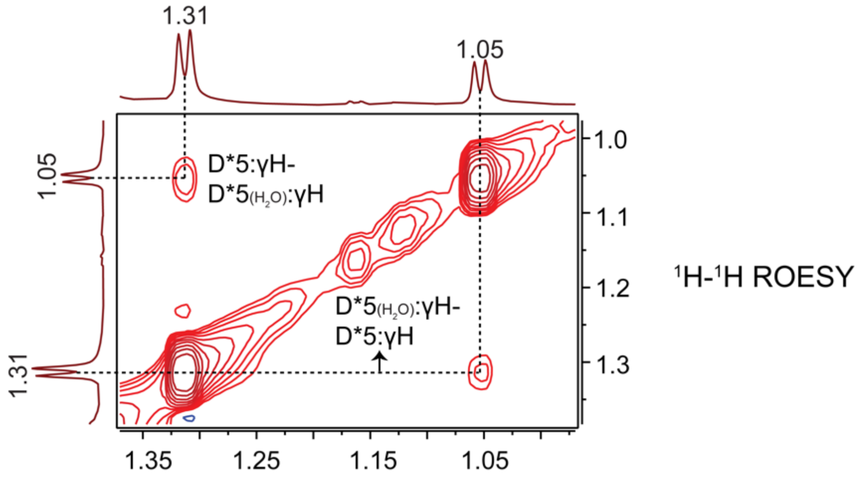

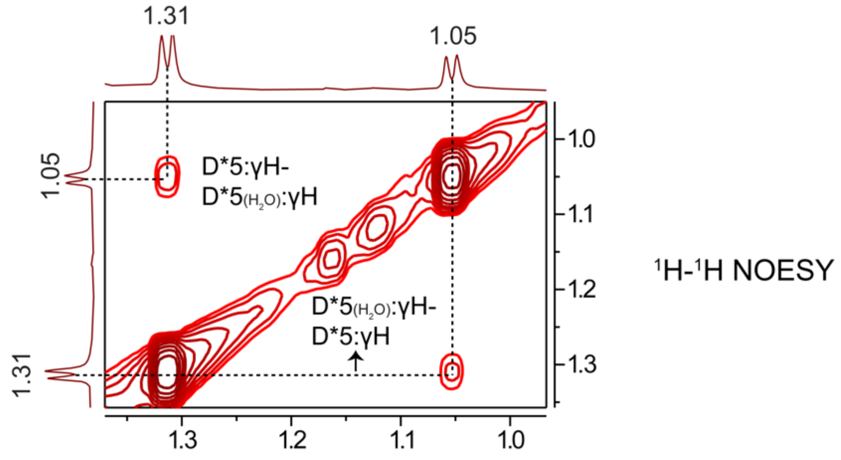

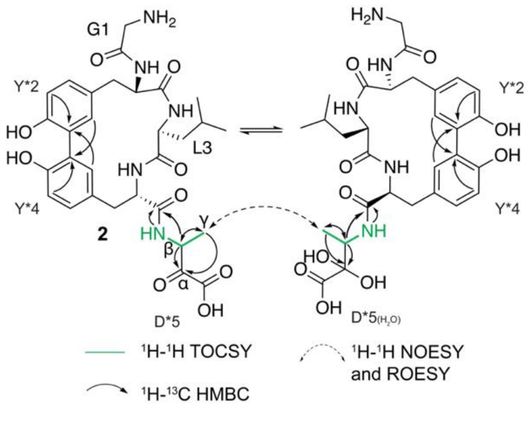

**Table S8:** Summary of $^{1}H$ and $^{13}C$ chemical shifts of ApyA modified by ApyO/H/I/S after GluC cleavage. Note: The chemical shifts of backbone NHs deviate up to 0.02 ppm depending on the data acquisition process.

A) Keto form

| number | AA | NH/C=O | αH/C | βH/C | γH/C | δH/C and others |
|---|---|---|---|---|---|---|
| 1 | G | <br>166.4 | 4.06, 3.79<br>40.9 | | | |
| 2 | Y | 8.51<br>170.2 | 4.93<br>53.7 | 3.03, 2.95<br>37.1 | 127.2<br>(Aryl C4) | 6.91(dd), 130.9 (Aryl C5)<br>6.81 (d), 115.7 (Aryl C6)<br>6.62 (d, 2Hz), 133.7 (Aryl C3)<br>C-OH: 152.1 (Aryl C1)<br>Ar-C (crosslinked): 125.3 (Aryl C2) |
| 3 | L | 8.51<br>174.2 | 4.78<br>52.0 | 1.47<br>41.6 | 1.41<br>24.2 | 0.74, 21.6<br>0.69, 21.6 |
| 4 | Y | 9.01<br>171.7 | 4.31<br>54.7 | 2.49, 1.98<br>38.2 | 131.6<br>(Aryl C4) | 6.97 (dd), 130.2 (Aryl C5)<br>6.67 (d), 115.7 (Aryl C6)<br>6.83 (d, 2Hz), 131.1 (Aryl C3)<br>151.2 (C-OH, Aryl C1)<br>125.6 (Crosslinked, Aryl C2) |
| 5 | D | 8.33 | 202.5 | 4.92<br>51.6 | 1.31<br>14.6 | |

B) Gem-diol form

| number | AA | NH/C=O | αH/C | βH/C | γH/C | δH/C and others |
|---|---|---|---|---|---|---|
| 1 | G | <br>166.4 | 4.06, 3.79<br>40.9 | | | |
| 2 | Y | 8.30<br>(br)<br>170.3 | 4.87<br>54.0 | 3.02, 2.95<br>37.1 | 127.2<br>(Aryl C4) | 6.94 (dd), 130.9 (Aryl C5)<br>6.81 (d), 115.7(Aryl C6)<br>6.62 (d, 2Hz), 133.7(Aryl C3)<br>OH-C: 152.1 (Aryl C1)<br>Ar-C (linker): 125.3(Aryl C2) |
| 3 | L | 8.53<br>174.0 | 4.73<br>52.0 | 1.41<br>41.6 | 1.41<br>24.2 | 0.79, 21.6, 0.74, 21.6 |
| 4 | Y | 8.93<br>172.2 | 4.34<br>55.1 | 2.54, 2.26<br>37.4 | 131.6<br>(Aryl C4) | 7.00 (dd), 130.2(Aryl C5)<br>6.72 (d), 115.7(Aryl C6)<br>6.85 (d, 2Hz), 131.1(Aryl C3)<br>151.2(C-OH, Aryl C1)<br>125.6(Crosslinked, Aryl C2) |
| 5 | D | 7.81 | 95.3 | 4.20<br>50.5 | 1.05<br>13.8 | |

**Figure S27:** HR-ESI-MS of oxidation of the α-keto acid in ApyO/H/I/S-modified ApyA pentapeptide by $H_2O_2$. The full structure of **2** and **3** can be found in Figure 5 (Main Text). A table with observed masses and the error compared to calculated values can be found in the Supplementary Dataset 3 (Excel). *: Sodium adduct. **: Potassium adduct.

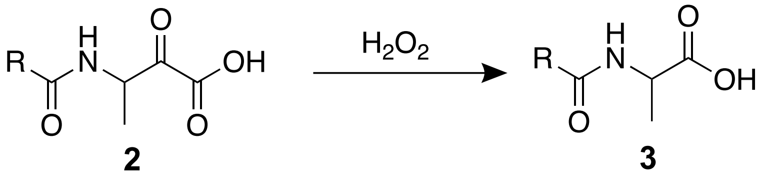

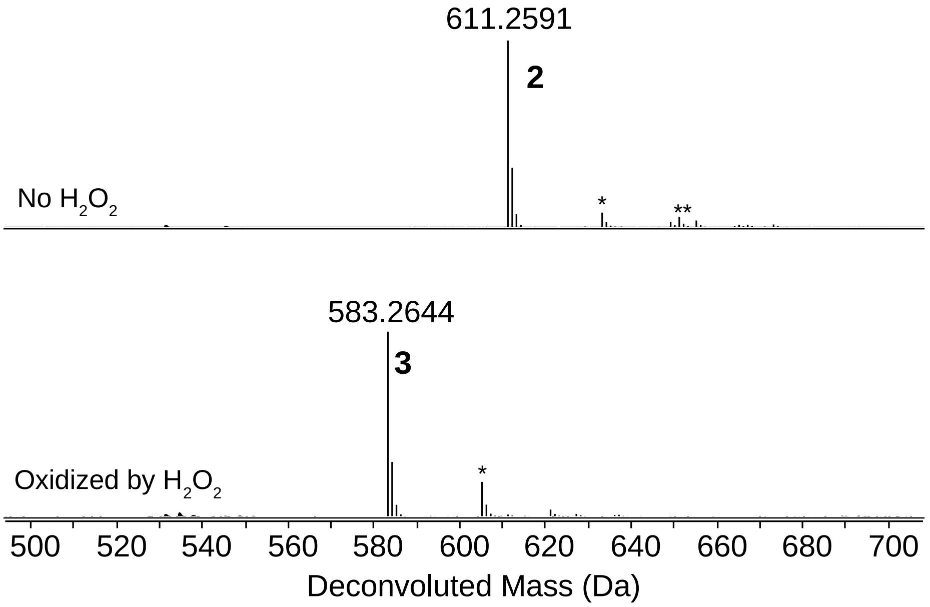

**Figure S28:** AlphaFold model of ApyS (pink) superimposed with the crystal structure of MppJ (green, PDB: 4KIB).[24] The active site of MppJ contains *S*-adenosyl-L-methionine, phenylpyruvic acid, and ferric ion. Parameters generated from DALI: Z-score = 25.7, RMSD = 3.6, number of aligned C-alpha atoms = 309, number of residues in target structure = 340. UCSF Chimera was used to generate this image.[25]

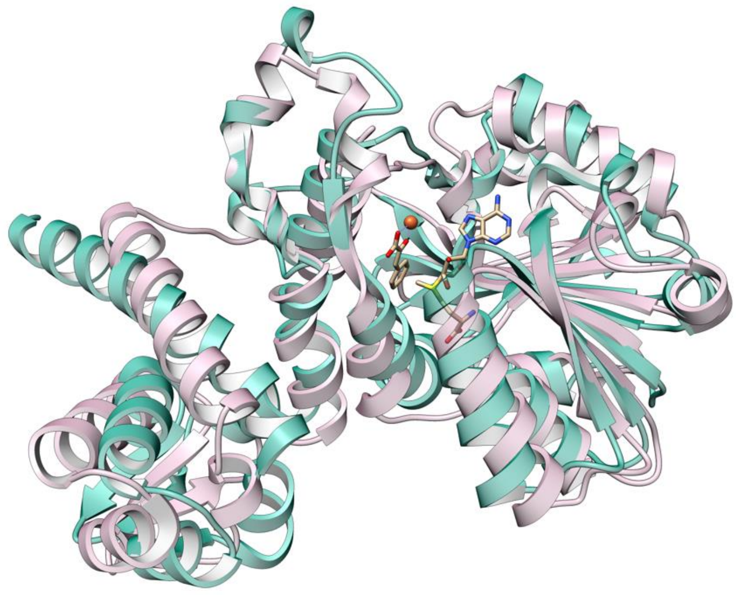

**Figure S29:** HR-ESI-MS of ApyO/H/I/S-modified ApyA after GluC cleavage. The Cys residue was alkylated by iodoacetamide. The MS spectrum of Fragment 4 is in Figure S3. Only the last four residues of the precursor peptides were modified by the enzymes. *: sodium adduct. CAM = Carbamidomethyl. A table with observed masses and the error compared to calculated values can be found in the Supplementary Dataset 3 (Excel).

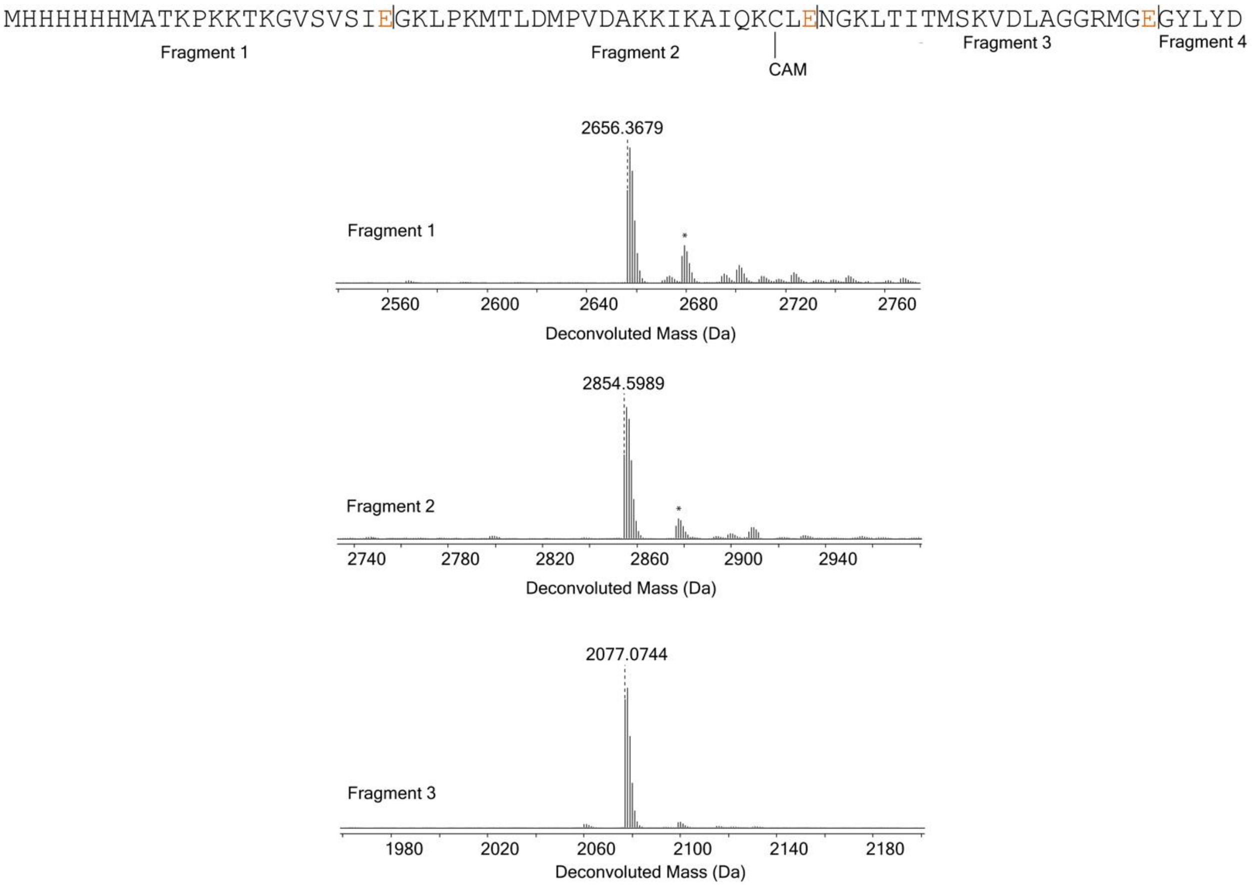

**Figure S30:** HR-ESI-MS/MS of fragments 1, 2, and 3 (Figure S30) of ApyO/H/I/S-modified ApyA after GluC cleavage. The Cys residue was alkylated by IAA.A table with observed masses and the error compared to calculated values can be found in the Supplementary Dataset 3 (Excel).

A) Fragment 1.

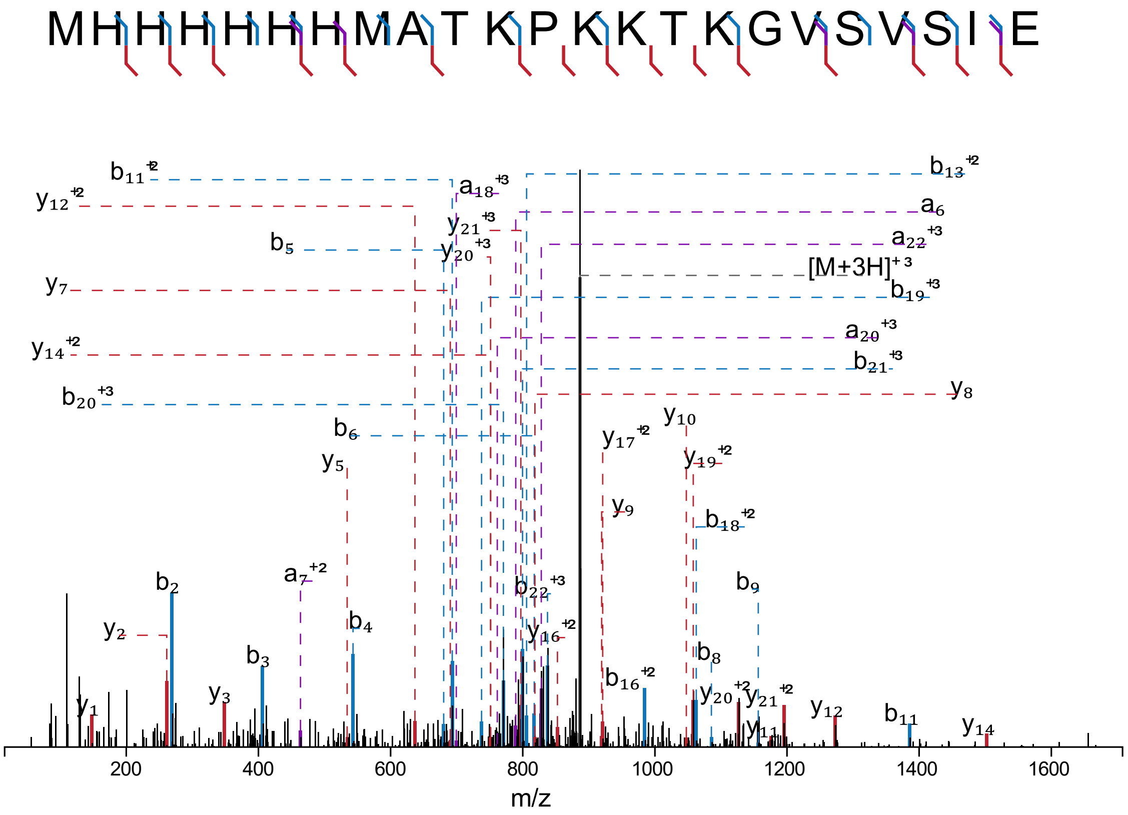

B) Fragment 2. c = carbamidomethylated Cys.

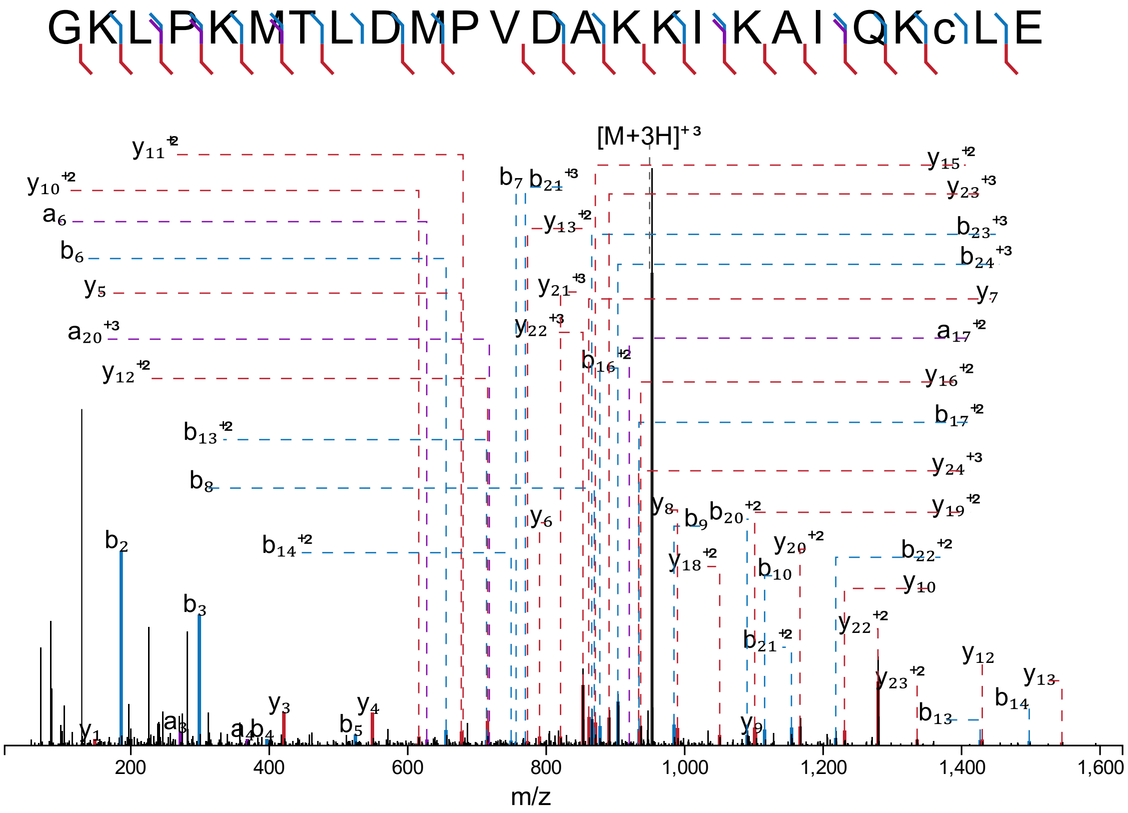

C) Fragment 3.

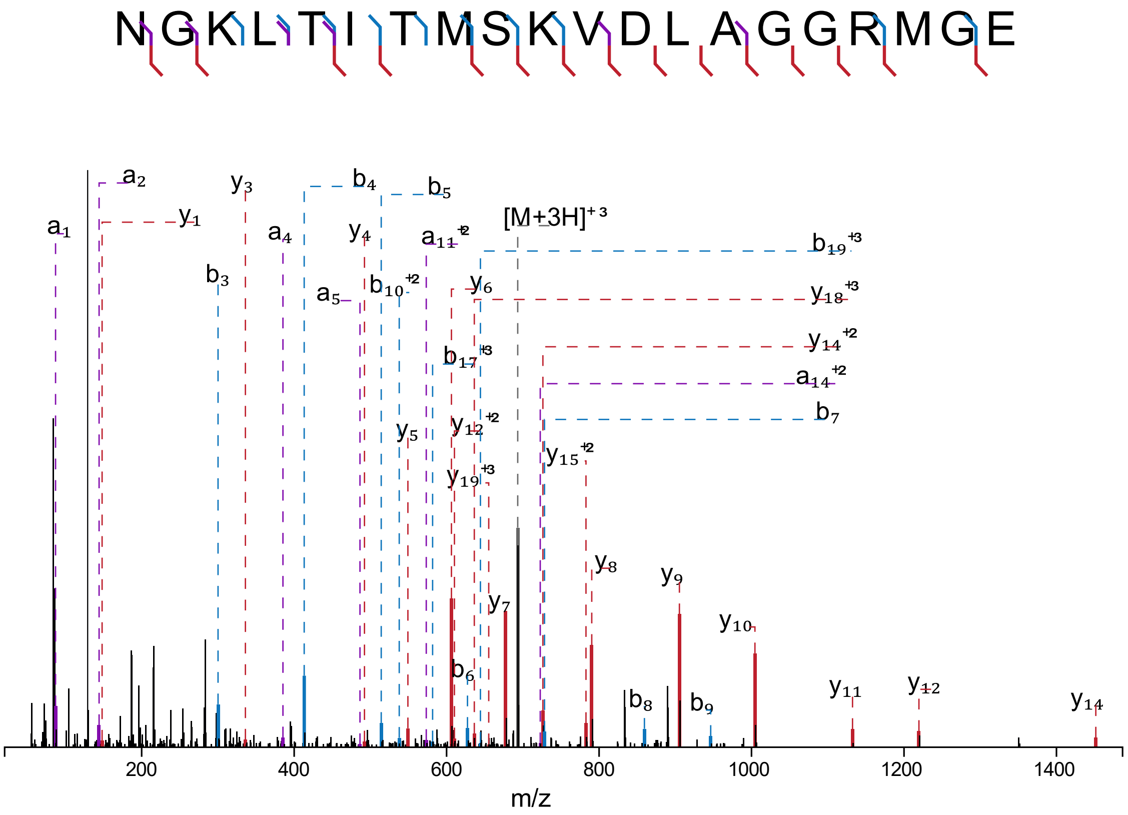

N G K L T I T M S K V D L A G G R M G E
a1
a2
y1
y3
b3
a4
b4
y4
a5
b5
b10 +2
a11 +2
[M+3H]+3
b19 +3
y6
y18 +3
b17 +3
y14 +2
a14 +2
b7
y5
y12 +2
y19 +3
y15 +2
y8
y9
y7
y10
b6
b8
b9
y11
y12
y14
200
400
600
800
1000
1200
1400
m/z

**Figure S31:** NMR spectra of the alkylated Cys-Leu-Glu tripeptide after GluC and trypsin cleavage of ApyO/H/I/S-modified ApyA. The solvent was $CD_3OH$. The structure of the product and the key NMR depiction are depicted below.

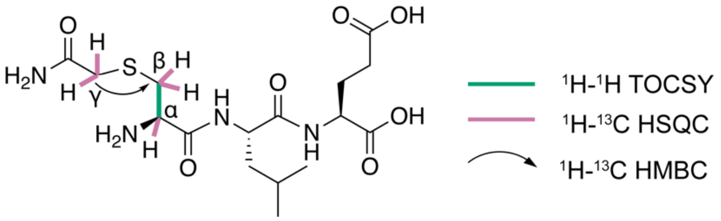

A) $^1$H NMR

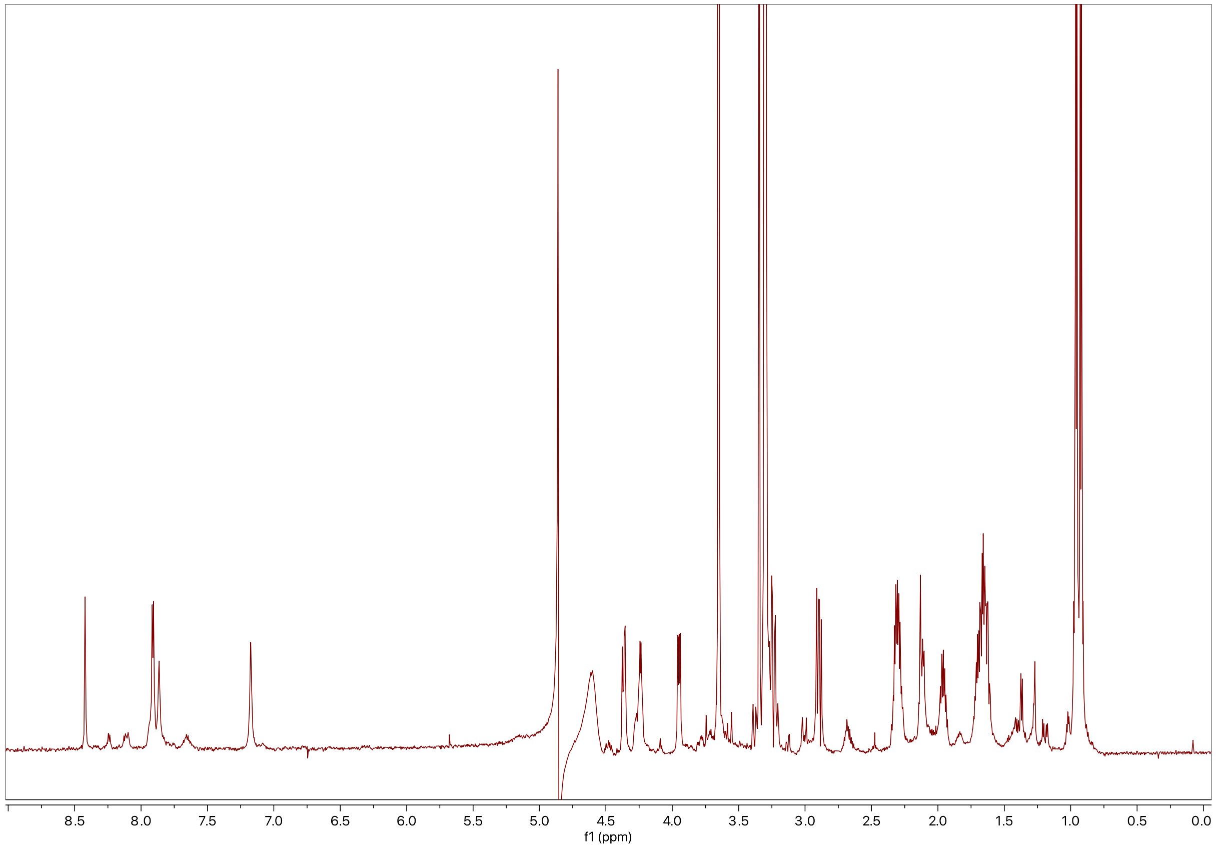

B) $^{1}H$-$^{1}H$ TOCSY

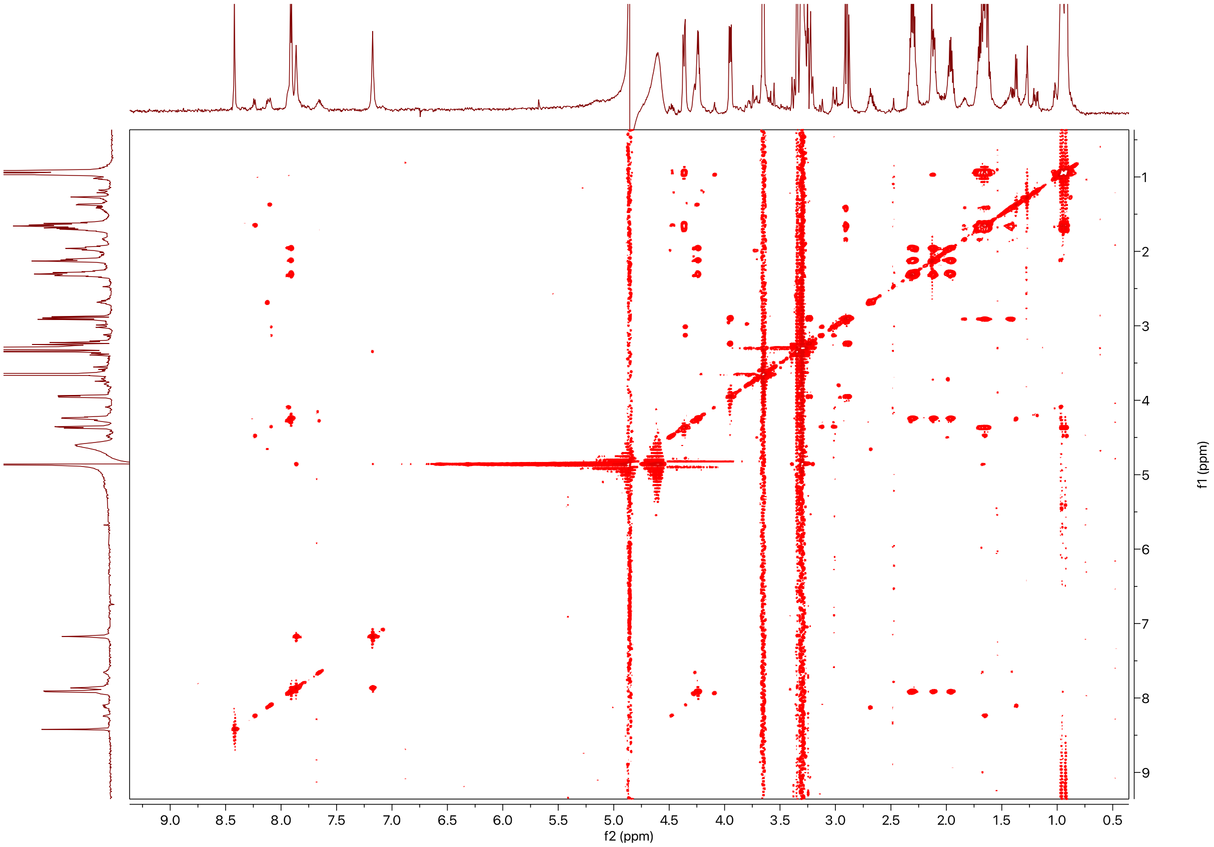

9.0
8.5
8.0
7.5
7.0
6.5
6.0
5.5
5.0
4.5
4.0
3.5
3.0
2.5
2.0
1.5
1.0
0.5
f2 (ppm)
1
2
3
4
5
6
7
8
9
f1 (ppm)

C) $^{1}H$-$^{13}C$ HSQC

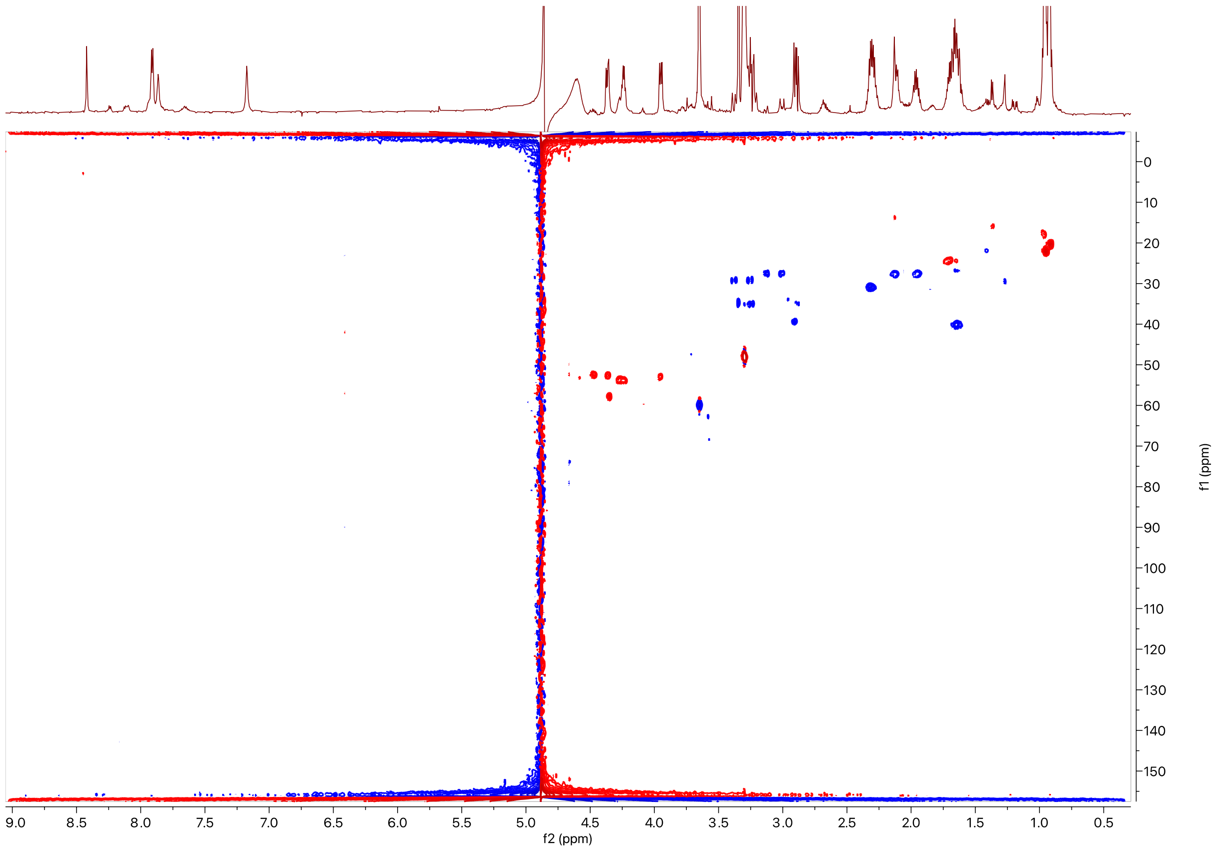

0
10
20
30
40
50
60
70
80
90
100
110
120
130
140
150
f1 (ppm)
9.0
8.5
8.0
7.5
7.0
6.5
6.0
5.5
5.0
4.5
4.0
3.5
3.0
2.5
2.0
1.5
1.0
0.5
f2 (ppm)

D) $^{1}H$-$^{13}C$ HMBC

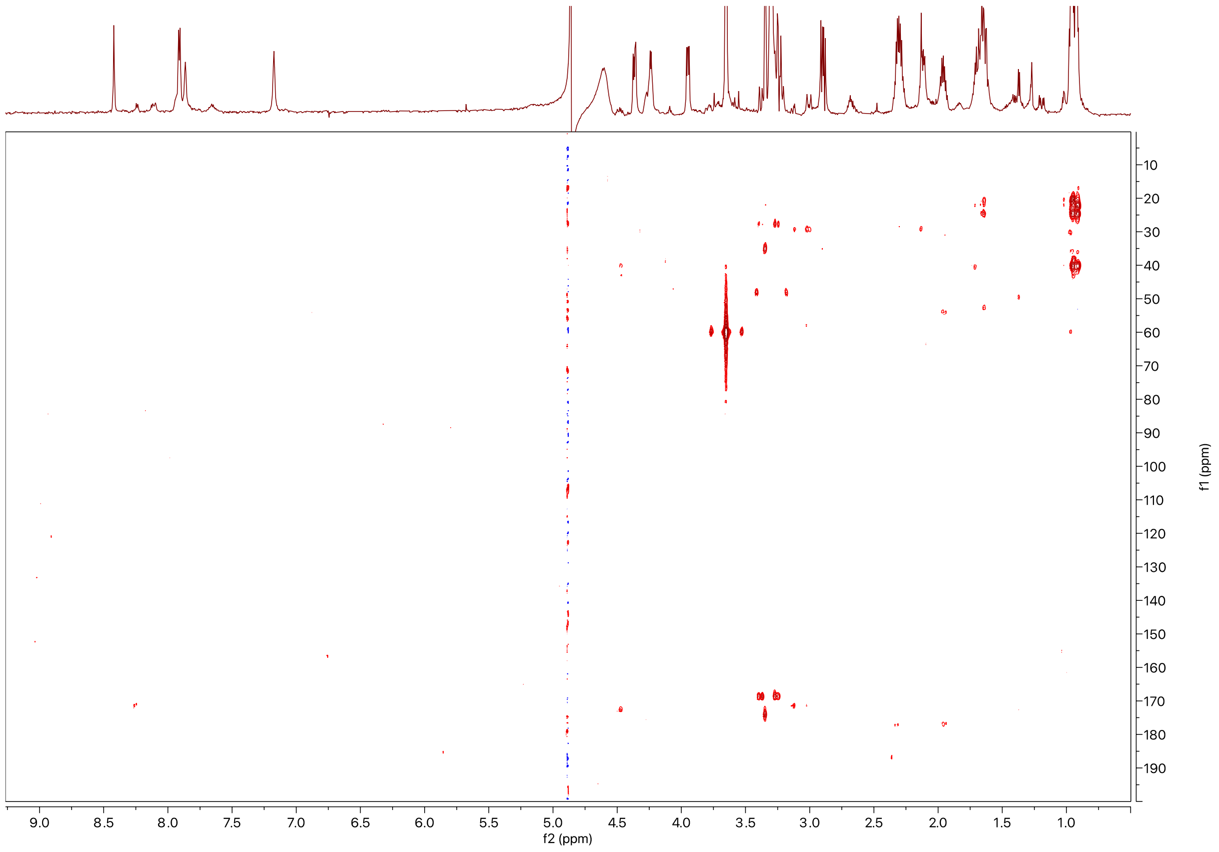

E) Key cross peaks in $^{1}H$-$^{1}H$ TOCSY, $^{1}H$-$^{13}C$ HSQC and $^{1}H$-$^{13}C$ HMBC suggesting that the Cys residue is unmodified by the enzymes.

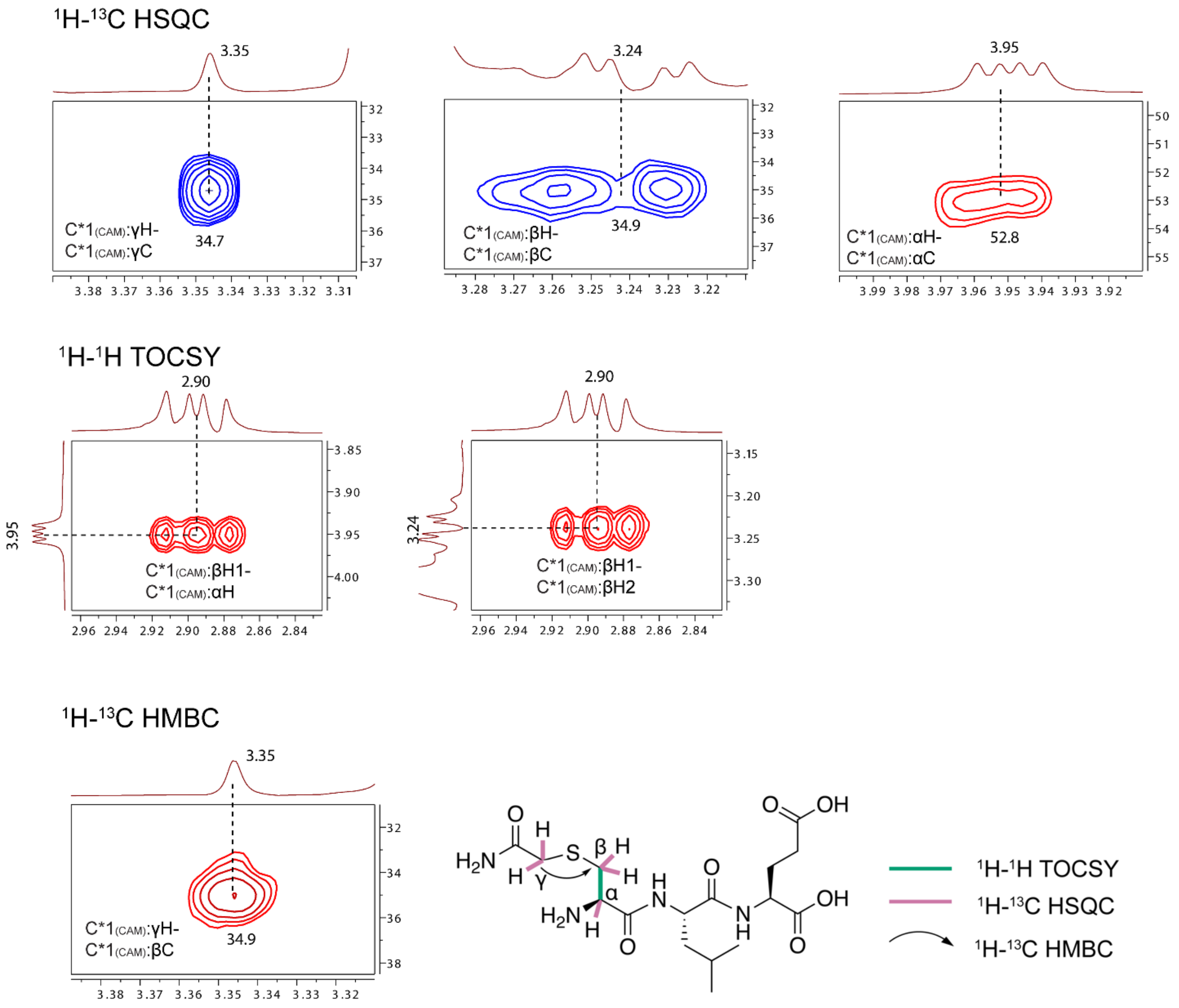

**Table S9:** Summary of $^{1}H$ and $^{13}C$ chemical shifts of iodoacetamide-alkylated Cys-Leu-Glu tripeptide after GluC and trypsin cleavage of ApyO/H/I//S-modified ApyA.

| Number | AA | NH, C=O | αH | βH | γH | δH |
|---|---|---|---|---|---|---|
| 1 | C | | 3.95<br>52.8 | 3.24, 2.90<br>34.9 | 3.35<br>34.7 | Sidechain $NH_2$:<br>7.86, 7.17<br>C=O 174.1 |
| 2 | L | 8.24<br>172.5 | 4.47<br>52.5 | 1.65<br>40.2 | 1.71<br>24.5 | 0.96, 22.3<br>0.92, 20.4 |
| 3 | E | 7.91<br>176.8 | 4.24<br>53.9 | 2.12, 1.96<br>27.6 | 2.30<br>31.1 | (d)C=O:<br>177.1 |

**Figure S32:** Marfey's stereochemical analysis of the IAA-alkylated Cys-Leu-Glu tripeptide. (A). Marfey's hydrolysis and derivatization scheme with L-FDAA. B) Extracted ion chromatogram (EIC *m/z* = 432.0780) of hydrolyzed alkylated Cys-Leu-Glu tripeptide and subsequently derivatized alkylated Cys using L-FDAA as the derivatization reagent. Commercial isomers of Cys were derivatized in the same manner and co-injected with the tripeptide-derived material. In this experiment, treatment of alkylated Cys-Leu-Glu tripeptide and standards in 6 M DCL resulted in the exchange of the protons of the carbamidomethyl group with solvent (see Figure S34). DCl is used such that any potential epimerization can be detected by the incorporation of additional deuterium.

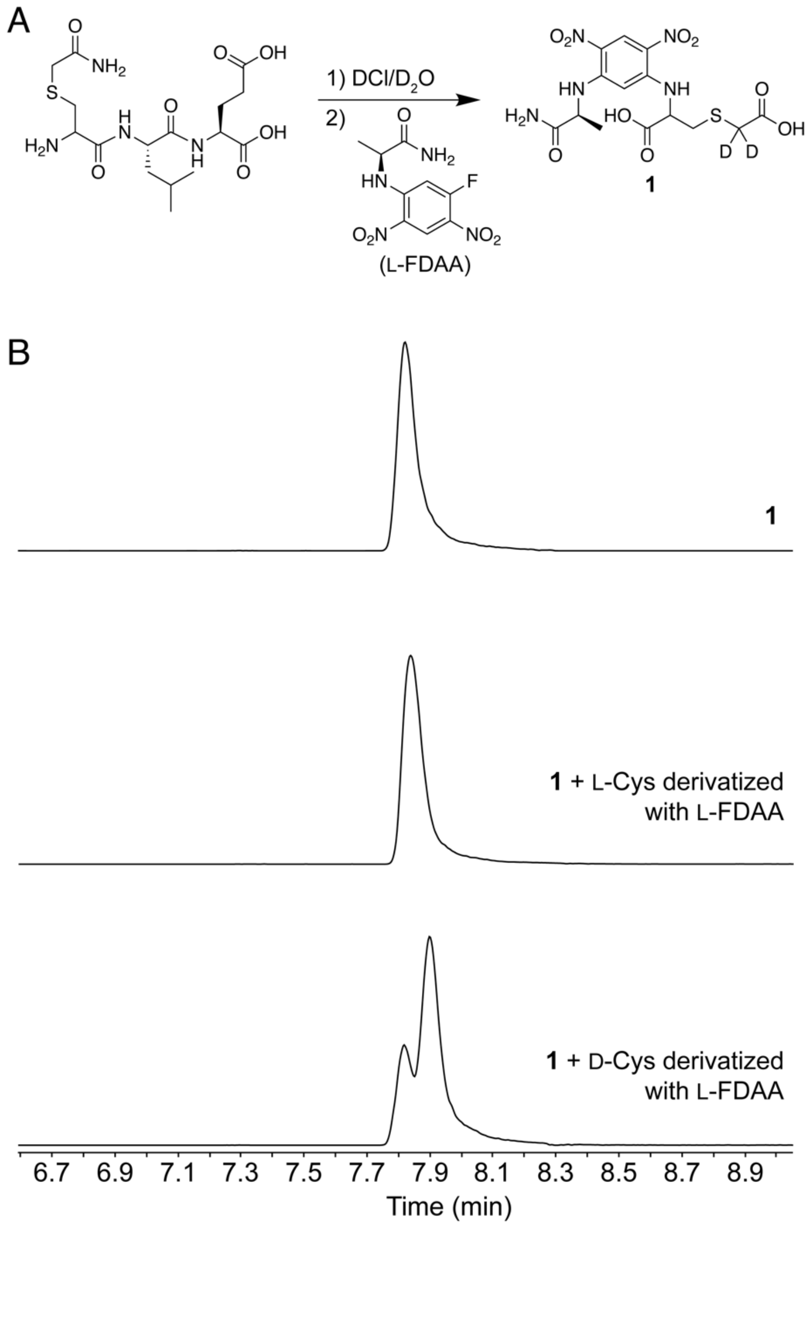

**Figure S33:** HR-ESI-MS/MS of the L-FDAA derivatized alkylated Cys. A) ESI-MS spectra showing deuterium incorporation. B) Targeted CID of each isotope showing that the deuterium(s) was incorporated in the proton of the carbamidomethyl moiety. A table with observed masses and the error compared to calculated values can be found in the Supplementary Dataset 3 (Excel).

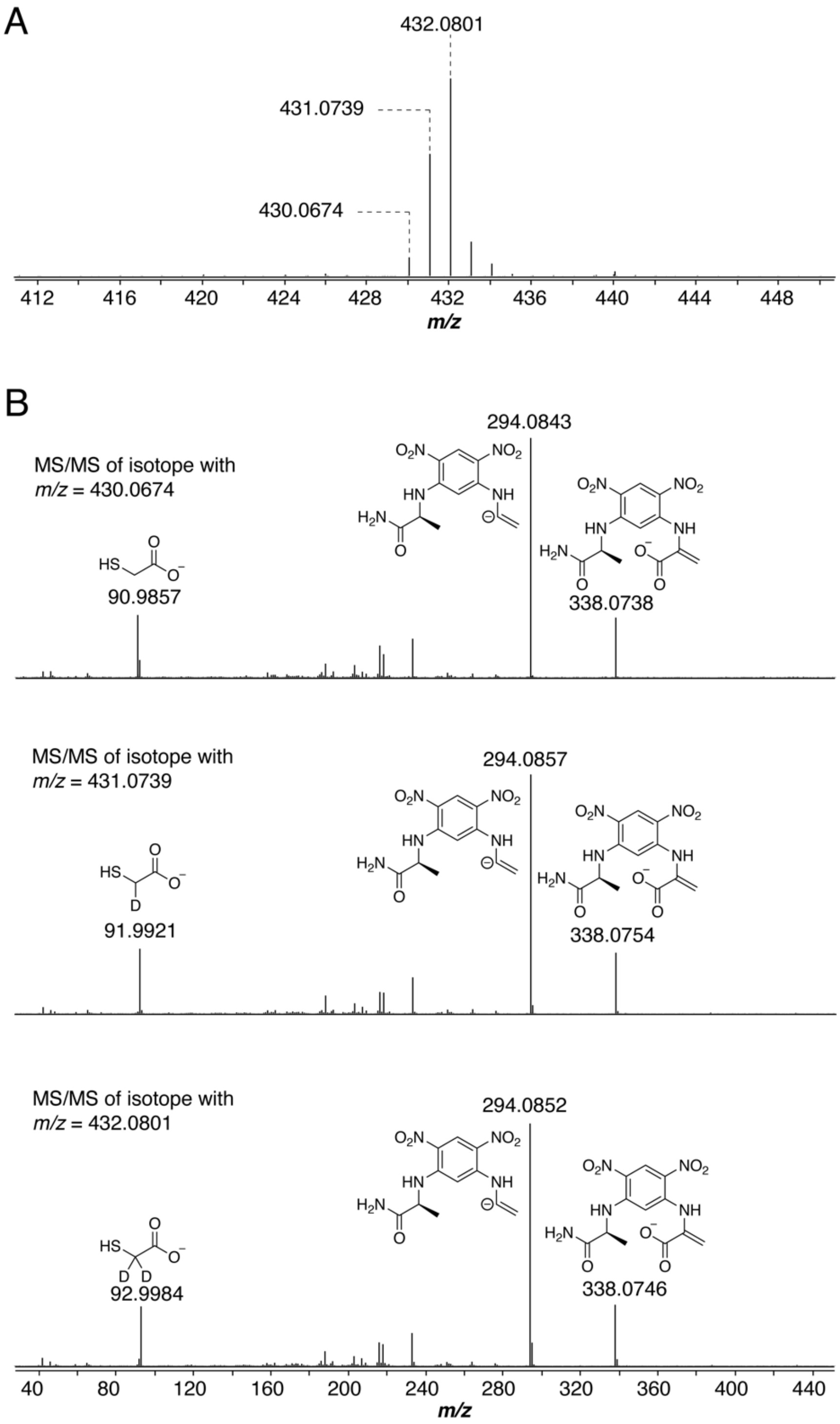

**Figure S34:** EICs of the C46A, C46S, G50A, G50V variants and wild type ApyA subjected to modification by ApyO, ApyHI, and ApyS. Endoproteinase LysC was used in this experiment. EIC *m/z* = 792.3621 represents ApyO/H/I-modified ApyA (z = 2). EIC *m/z* = 799.3699 represents ApyO/H/I/S-modified ApyA (z = 2). EIC *m/z* = 807.3674 represents ApyO-modified ApyA (z = 2).

46 50
MHHHHHHMATKPKKTKGVSVSIEGKLPKMTLDMPVDAKKIKAIQK**C**LEN**G**KLTITMSKVDLAGGRMGEGYLYD

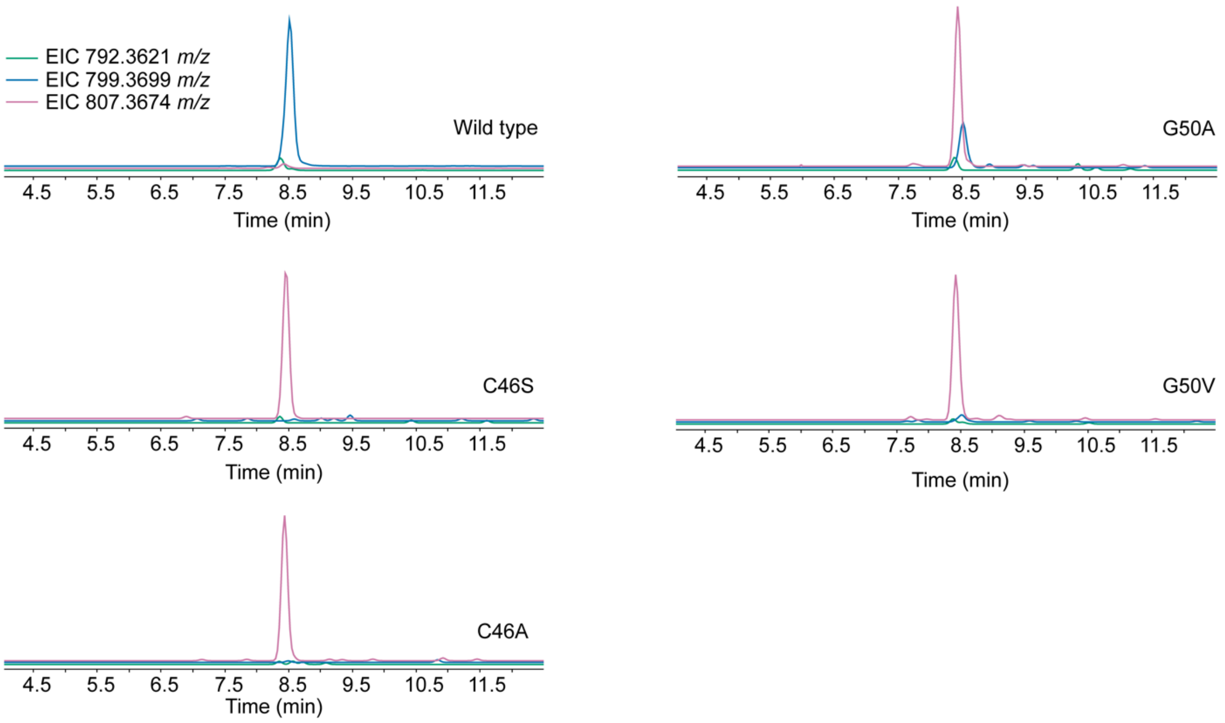

**Figure S35:** HR-ESI-MS/MS of LysC-cleaved ApyO/H/I/S-modified ApyA (G50A). A table with observed masses and the error compared to calculated values can be found in the Supplementary Dataset 3 (Excel).

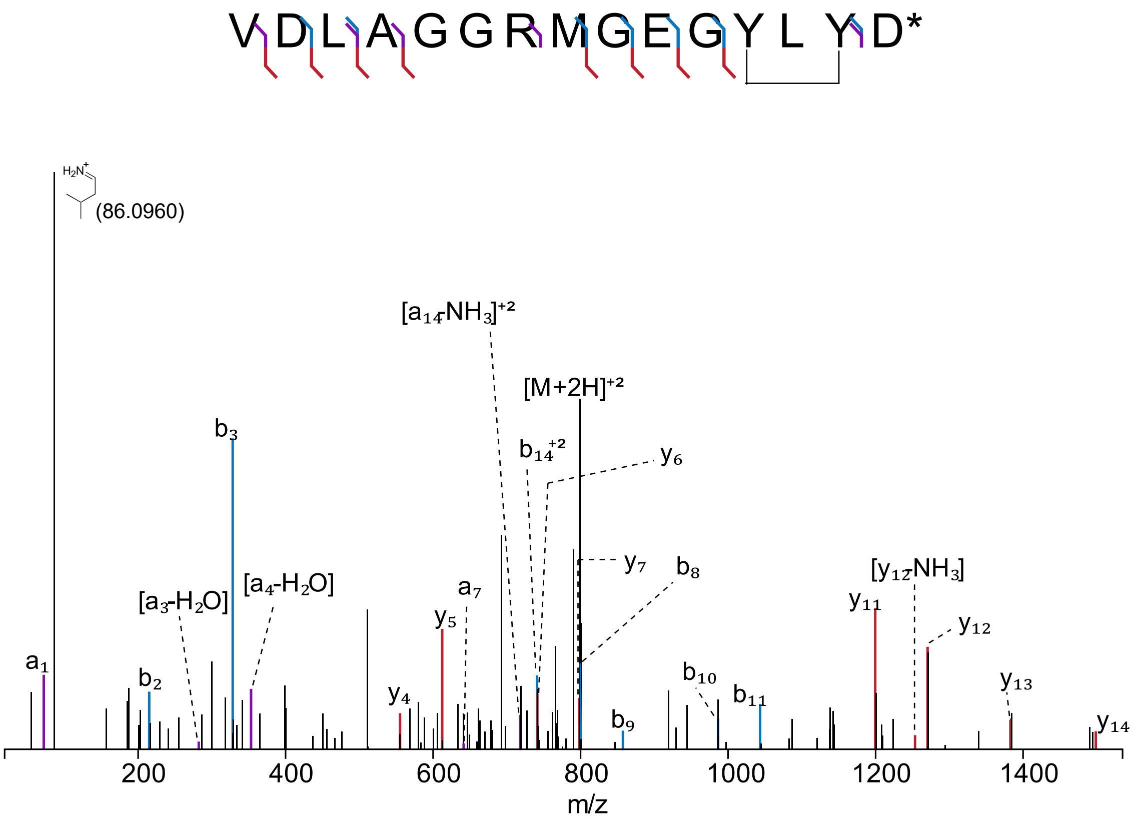

**Figure S36:** EICs of C46H and C46D variants of ApyA subjected to modification by ApyO and ApyHI. Endoproteinase LysC was used in this experiment. EIC *m/z* = 792.3621 represents ApyO/H/I-modified ApyA (z = 2). EIC *m/z* = 807.3674 represents ApyO-modified ApyA (z = 2). Note that the samples were subjected to similar liquid chromatography gradients (see supplementary methods) except that the run with the wild type ApyA underwent 5 min of desalting at 5% $CH_3CN$ (0.1% FA) instead of 4 min like in ApyA(C46H) and ApyA(C46D).

```
                                                      46  50
MHHHHHHMATKPKKTKGVSVSIEGKLPKMTLDMPVDAKKIKAIQKCLENGKLTITMSKVDLAGGRMGEGYLYD
```

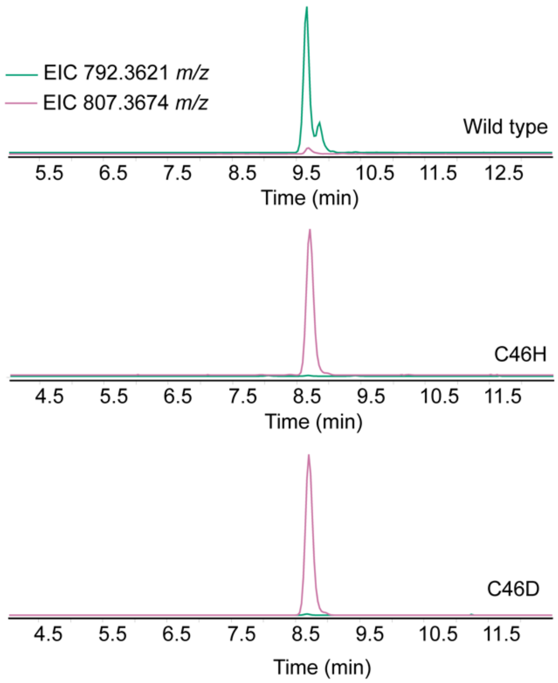

**Figure S37:** Proposed mechanisms for the transformation catalyzed by ApyHI. The ApyH protein likely has two or three iron ions in its active site,[26-28] but only one is drawn. The others may aid in binding the substrate or could be involved in catalysis. The decarboxylation may or may not take place in the enzyme active site. Decarboxylation from a radical intermediate like **II** to provide an alkene has precedence in other iron dependent enzymes.[29-32]

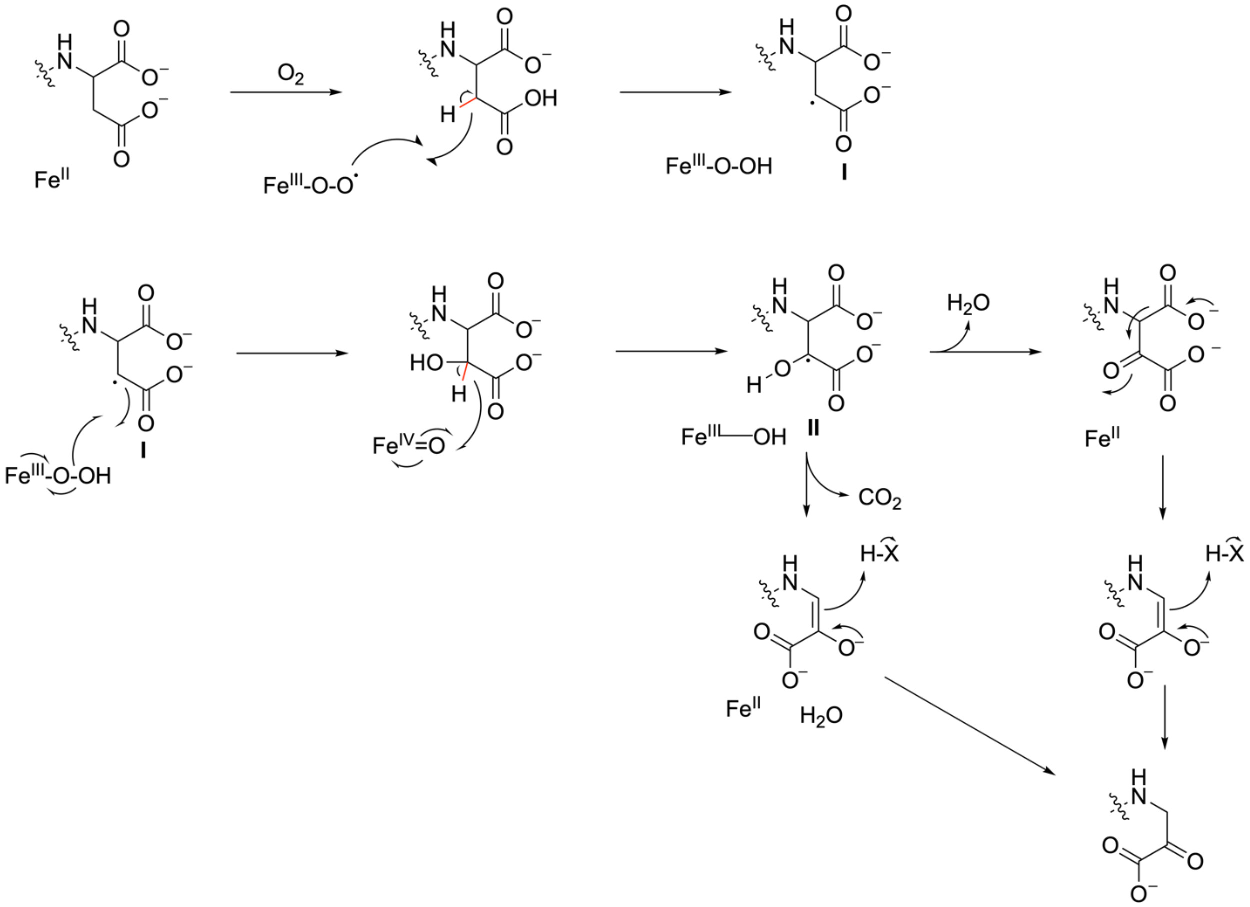
